# CGGBP1-regulated heterogeneous C–T transition rates relate with G-quadruplex potential of terrestrial vertebrate genomes

**DOI:** 10.64898/2025.12.10.693595

**Authors:** Praveen Kumar, Umashankar Singh

## Abstract

The formation of G-quadruplexes (G4) is fundamentally linked to GC content of the DNA. There is little evidence for evolutionary selection of G4 stabilizers; G4 formation depends strictly on inherent sequence properties. Vertebrate promoters are notable for their consistently high GC-content as well as pronounced G/C strand asymmetry. Genomic GC content has changed in the course of evolution of terrestrial vertebrates, especially the amniotes but its relationship with the potential to form G4 remains less well understood. By analyzing genomes of 101 amniotes we report that lineage-specific differences in association between GC-content and G4 formation potential (pG4) in amniotes are concentrated at 1kb promoter regions. To understand possible mechanisms underlying high GC-concentrations in pG4 of mammalian and avian promoters, we test the possibility of selective cytosine methylation restriction leading to G/C-retention by CGGBP1, a protein involved in mitigation of cytosine methylation as well as G4-formation. By analyzing promoterome-wide C-T transition rates at TFBSs in pG4s we show that mammalian and avian CGGBP1s preserve C through methylation restriction. Our approach involves a combined meta-analysis of (i) genomes and promoters of 101 amniote genomes classified into reptilian, avian and mammalian classes through PQS finder and FIMO for 1019 JASPAR vertebrate motifs, (ii) recently reported global cytosine methylation patterns affected by overexpression of amniotic or non-amniotic forms of CGGBPs, (iii) and PWM reconstruction of motifs differentially enriched in pG4s showing homeothermic specific GC-retention. Our findings suggest that cytosine methylation restriction by CGGBP1 shapes G4 forming profiles of vertebrate promoters by preserving C on the G4-complementary strand resulting in minute differences in TFBSs across amniotes.

**Highlights:**

- Evolution of vertebrate genome G4-forming potential mirrors that of CGGBP1.
- CGGBP1 prevents meC-to-T mutations at pG4 sites, preserving G4-forming potential specifically in amniote promoters.
- pG4-TFBS mutations reveal homeothermic GC-retention bias, mirroring CGGBP1-mediated pG4 protection patterns.
- The G4-forming regions most responsive to CGGBP1 exemplify this methylation-linked defense against C-to-T substitutions, distinguishing homeotherms from poikilotherms.

## Introduction

DNA is classically recognized as a right-handed double helix (B-DNA) formed by Watson-Crick base pairing between purines and pyrimidines (Bochman et al. 2012). However, DNA can adopt diverse non-B DNA secondary structures driven by the complexity of the underlying sequence (Wang and Vasquez 2023). Beyond primary sequence alone, their formation depends on various cellular factors and processes, illustrating the dynamic transition from regular B-DNA to alternative conformations (Wang and Vasquez 2023). A well-studied example is the G-quadruplex (G4), which forms in guanine-rich sequences where runs of guanines are repetitively interrupted by other nucleotides (Wang and Vasquez 2023; Rhodes and Lipps 2015). G4s are formed through Hoogsteen hydrogen bonding between guanines in stacked tetrads (G-quartets) derived from the G-rich strand of DNA or RNA (Bochman et al. 2012; Varshney et al. 2020). These structures typically comprise at least two stacked G-quartets for stability and conform to the consensus motif G_3–5_N_1–7_G_3–5_N_1–7_G_3–5_N_1–7_G_3–5_, where G_3–5_ indicates runs of 3–5 guanines separated by loops (N) of 1–7 nucleotides (Bochman et al. 2012). Loop lengths vary, enabling diverse topologies such as propeller or hybrid forms (Bochman et al. 2012).

It is important to note that the genomic DNA is predominantly maintained as a double helix because this conformation is energetically stable, even when sequence patterns could form alternative structures like G4s (Wang and Vasquez 2023). Thus, beyond the DNA sequence itself, G4 formation depends on multiple factors and is a dynamic and transient event (Varshney et al. 2020). In the native cellular environment, G4 formation is triggered by strand separation during processes like replication and transcription, and is further influenced by the open chromatin states, negative supercoiling, and specific binding proteins(Shen et al. 2021; De and Michor 2011; Wang and Vasquez 2023) . Upon folding, G4-forming sequences act as molecular switches regulating all major steps of gene expression, from DNA to protein (Wang and Vasquez 2023). In the nucleus, these structures work hand-in-hand with the transcription machinery. They’re not just roadblocks that slow down transcription (Robinson et al. 2021), instead they act as special promoter elements facilitating nucleosomes exclusion and make RNA polymerase II pause briefly, setting genes up for quick activation when needed (Eddy et al. 2011; Esnault et al. 2025). Beyond promoters, their presence in introns and splice sites expands the proteome diversity by modulating alternative splicing (Georgakopoulos-Soares et al. 2022). These structures continue their regulatory influence even after transcription; G4s in the 5’ untranslated region (UTR) can modulate translation in size dependent manner (Chow et al. 2025; Lee et al. 2024), while their presence in 3’ UTR regulates mRNA stability by modulating microRNA binding (Rouleau et al. 2017; Qi et al. 2021). Through these mechanisms, G-forming sequences serve as hubs of transcription factor binding and shape cell type specific transcriptomes (Spiegel et al. 2021; Lago et al. 2021). Furthermore, regardless of their indisputable role in telomere maintenance and recombination (Bochman et al. 2012; Varshney et al. 2020), these structures can be considered as double-edge swords. If left unresolved, G4s can impede replication by stalling the fork progression (Lemmens et al. 2015), lead to mutagenic R-loop formation (Sollier and Cimprich 2015), and could promote genomic instability associated with disease like cancer (De and Michor 2011; Wang and Vasquez 2014; Kim and Jinks-Robertson 2012).

Although G4 formation remains highly dynamic and transient, their potential genomic locations can be reliably predicted computationally from the underlying DNA sequence (Li et al. 2023; Vannutelli et al. 2023). The capacity of specific DNA sequences to form G4s represents their G4-forming potential (pG4). The genome-wide G4-forming potential can thus reveal evolutionary patterns in G4 distribution. Various studies have experimentally confirmed G4s across the tree of life (Wu et al. 2021; Marsico et al. 2019; Vannutelli et al. 2023). Researchers are now studying how these structures evolved across species and came to regulate such diverse biological processes. Studies have shown that these sequences face positive selection pressure (Brázda et al. 2021; Mohanty et al. 2025). They appear far more often than random DNA would predict, and they’re linked to greater developmental complexity, despite their risk of causing genomic instability (Smith 2010; Makova and Weissensteiner 2023). While B-DNA offers simple sites for protein recognition, secondary structures like G4s provide many more binding conformations, increasing the genome’s epigenetic potential (De and Michor 2011; Spiegel et al. 2021). The transition from poikilothermy to homeothermy brought higher body temperatures and a modest 35-45% GC content increase, suggesting evolutionary selection favored G-rich motifs over temperature-driven genome stabilization (Smith 2010).

Whether G4 structures actually form i*n vivo* remains highly variable and context-dependent, much like other epigenetic marks. G4 formation is fundamentally driven by DNA sequence properties but regulated through two distinct molecular mechanisms. First, specialized helicases like BLM, WRN, and DHX36 actively resolve G4s while preserving the underlying sequence’s G4-forming potential (Mendoza et al. 2016; Varshney et al. 2020; Wu et al. 2018). Second, the intrinsic sequence features enabling G4 formation are themselves conserved under evolutionary selection, maintaining the DNA’s capacity for G4 structure formation (Du et al. 2008). This sequence conservation in gene regulatory regions underscores their functional importance (Du et al. 2008). However, mutagenic processes like methylation-driven C-to-T transitions can disrupt this conservation, abolishing G4-forming potential entirely (Mao et al. 2018; Neupane et al. 2023). Such mutations can occur both in G-rich runs critical for tetrads (GGG regions) and in loop regions between runs, with transitions accounting for approximately 22% of G4 structure-destabilizing mutations (Neupane et al. 2023). Furthermore, the relationship between DNA methylation and G4 formation is bidirectional. While methylation can destabilize certain G4 structures or prevent their formation, G4 structures themselves protect CpG islands from methylation by sequestering and inhibiting DNA methyltransferase 1 (Mao et al. 2018). This interplay suggests that G4 formation and the preservation of G4-forming sequence potential represent a critical regulatory interface between genomic sequence architecture and epigenomic regulation (Mao et al. 2018).

Cytosine methylation is one of the most well-conserved epigenetic mechanisms known to occur in almost all organisms with double-stranded DNA as their genetic material. Traditionally, it has been studied as a chemical modification that affects DNA-protein interactions, chromatin compaction, gene expression, and a molecular memory for genome function over and above the DNA sequence (Baubec and Schübeler 2014; Jones 2012; Moore et al. 2013). More recently, cytosine methylation has been studied in the context of non-canonical DNA structures, most importantly the G4s (Mao et al. 2018; Niu et al. 2025; Wang et al. 2020). The formation of G4s primarily depends on short and spaced repeats of G on the same DNA strand. Although cross-strand annealings of G-rich segments do yield inter-strand quadruplex-like structures, it only affects the length of G-rich segments on a particular strand of DNA without negating the necessity of skewed G-richness on the strands participating in G4 formation. The G4-forming region inherently has a local GC-skew with the C-rich strand offering substrates for non-CpG cytosine methylation. In the event of G4 folding of the G-rich strand, the C-rich strand, which is left single-stranded, runs the risk of a higher mutation load, which is exacerbated by cytosine methylation. The meC-to-T transitions occur spontaneously at a rate much lower than possible otherwise due to the double-stranded nature of the DNA. While most of the U (generated by spontaneous deamination of C) is rectified back to C guided by G:::C pairing on dsDNA, the T (generated by spontaneous deamination of 5meC) poses a G-A mutational load on the complementary strand. GC-richness combined with GC-skew is rich in transcription factor binding sites (TFBSs), often concentrated in the cis promoters and enhancers genome-wide (Deaton and Bird 2011; Hartono et al. 2015).

TFBSs in vertebrate genomes are abundantly GC-rich, with cytosine methylation functioning as a mitigator of TFBS-DNA binding (Yin et al. 2017). The primarily unmethylated status of CpG dinucleotides in CpG islands is a classic demonstration of the cooperation between TFBS activity, gene expression, and the unmethylated status of cytosines (Yin et al. 2017). The palindromic nature of the CpG context ensures symmetrical cytosine content on complementary strands, as well as the maintenance of methylation across replication and repair. The CpG dinucleotides thus abrogate any inter-strand GC-skew and associated secondary structures such as G4, which are potentially formed at regions with non-CpG cytosines (Agarwal et al. 2015; Patel et al. 2019; Datta et al. 2024). There is evidence that vertebrate TFBSs have an abundance of non-CpG GC-contents (Khuu et al. 2007), which are prone to strand-asymmetric cytosine methylation, loss of C to T, and eventual loss of TFBS GC-content. The concentration of non-CpG TFBSs in the *cis* promoters, as well as the “dark matter” of the genome, increases the chances of GC-skew and resultant G4 formation potential. Since the sparse non-CpG cytosines tend to be predominantly methylated, due to an ancient mechanism of guarding against non-self genetic material, it combines two distinct processes: a chemical mark that is methylation of cytosines and a physical structure that is the G quadruplex. Biochemical studies have shown that G4 stability is dually affected by cytosine methylation, with evidence that cytosine methylation generally enhances G4 stability (Lin et al. 2013). Since cytosines are not involved in the G4 Hoogsteen bonding directly, their influence on G4 stability is likely to be indirect and as yet poorly understood.

Given the above premise, there is a strong case for the loss of GC-content in the course of evolution through the meC-to-T mutation. However, against all odds, the course of terrestrial vertebrate evolution has not only preserved the GC-content, but also enriched it (Morbia et al. 2024; Kumar et al. 2025). As compared to the coelacanths, the sarcopterygian lineage ancestral to the terrestrial vertebrates, and non-amniotes, the avian and mammalian genomes generally have higher GC-content (Kumar et al. 2025). The various lineages of the divergent reptilian groups show highly divergent GC-contents. Such a retention of GC-content, especially in the non-CpG contexts, stands at odds with the default nature of non-CpG island cytosines. Such a retention of GC-content, with implicit conserving effects on TFBS-DNA interactions, could not have occurred without an active mechanism. The transition of vertebrates from aquatic to terrestrial habitats has been accompanied by the evolution of CGGBP1, a gene that seems to fill this functional void. CGGBP1 binds to GC-rich DNA and, amongst a host of other effects, manifests functions pertinent to our discussion: mitigation of cytosine methylation in non-CpG contexts, prevention of G4 formation at regions of GC-skew, and counteraction of R-loop accumulation and transcription-replication conflicts at short CGG-repeat tracts prone to R-loop formation (Agarwal et al. 2015; Datta et al. 2024; Ummethum et al. 2025).

Recent reports have shown that the TFBS landscapes have retained/gained GC in higher amniotes (which are also endothermic homeotherms), whereas the poikilothermic lower amniotes have progressively lost TFBS GC-contents under the influence of CGGBP1 (Kumar et al. 2025). Bernardi and colleagues demonstrated that DNA methylation levels are inversely correlated with body temperature across vertebrate species, with poikilothermic (cold-blooded) organisms exhibiting significantly higher DNA methylation levels compared to homeothermic (warm-blooded) organisms (Varriale and Bernardi 2006). This temperature-dependent methylation pattern, reflecting evolutionary adaptations to organismal physiology, appears to be fundamentally encoded within the CGGBP1 protein itself. Recent experimental evidence demonstrates that simply expressing CGGBP1 orthologs derived from poikilothermic species (*Latimeria chalumnae* and *Anolis carolinensis*) versus homeothermic species (*Homo sapiens* and *Gallus gallus*) recapitulates the ancestral methylation signatures associated with their organismal origin (Kumar et al. 2025) in human somatic cell cultures. Remarkably, when poikilothermic CGGBP1 orthologs are overexpressed, the resulting genome-wide methylation profile exhibits elevated methylation levels characteristic of cold-adapted species, whereas homeothermic CGGBP1 orthologs impose methylation restriction patterns typical of warm-blooded organisms (Kumar et al. 2025). We assume and argue that a similar cytosine methylation regulatory function of CGGBP1 would have operated in the germline, giving rise to methylation associated with mutational biases and genomic base compositional changes. To the extent this holds true, comparisons of genome-wide G4-forming potentials and CGGBP1-driven point mutation rates between major vertebrate phyla can inform us about how cytosine methylation has affected G4-forming potentials in the course of vertebrate evolution.

The GC-richness of higher amniotic genomes, especially the promoters, inherently increases their G4-forming capacity. The evolutionary expansion and conservation of G-quadruplex motifs suggest they play crucial roles in gene regulation and genome stability (Wu et al. 2021). Particularly, the progressive enrichment of G4 motifs in regulatory regions and their increasing density and diversity during species evolution indicate that higher organisms have evolved G-quadruplexes into an elaborate transcriptional regulatory mechanism to meet complex physiological and behavioral demands (Wu et al. 2021). Apparently, the evolution of terrestrial vertebrates has the following underlying coincidental genomic feature enrichment, which is likely to be more concentrated at promoters: an increase in G4-forming potential reflected as GC-skew, GC-richness, and an enhanced property of cytosine methylation mitigation. Evidently, the evolution of G4-forming potential of genomes has an epigenetic basis in the form of cytosine methylation, at the TFBS-rich proximal promoters, as well as the genomic dark matter and gene expression. A systematic assessment of the evolution of the G4-forming potential of genomes and its relationship with cytosine methylation is required to understand this mechanism. Through a simple yet rigorous meta-analysis of 105 genomes, we demonstrate an evolutionary path of G4-forming potential in taxon groups ranging from coelacanths to eutherians. We provide evidence that the evolution of the cytosine methylation mitigator protein CGGBP1 correlates with this enhanced G4-forming potential of mammalian genomes. We show that CGGBP1 expression specifically restricts meC-to-T transition mutations (not other types of transitions or transversions) strongly at genome-wide regions with GC-skew and high G4-forming potential, which are concentrated at *cis-*proximal promoters. Our findings are based on the predictable G4-forming potential, actual GC-skew, and a base-level analysis of the point mutation landscape. These results explain the pattern of evolution of the G4-forming potential of terrestrial vertebrate genomes and its evolution alongside the progressive emergence of amniotes, homeotherms, and mammals. The results presented here provide an epigenetic basis for the lineage-specific differences in G4-forming potentials, present C deamination as a key molecular mechanism that indirectly drives the evolution of G4-forming potential, and posit CGGBP1 as a key arbiter of G4-forming potential by determining the mutability of G4-complementary C-rich strand.

## Results

### Genomic GC content is the primary driver of G-quadruplex potential and evolution in vertebrates

While DNA is predominantly recognized as a B-DNA double helix, its underlying sequence complexity enables the formation of alternative non-B DNA structures. G-quadruplexes (G4s), one of the most widely studied, form in guanine-rich sequences where runs of guanines are interrupted by loops of variable length formed by other bases. Because G4 formation is sequence-dependent, these motifs serve as markers for studying G4 evolution across genomes. Computational tools like QGRS Mapper, G4Hunter, and pqsfinder reliably predict these regions. In this study, we used pqsfinder to identify potential G4-forming regions in vertebrate genomes (Hon et al. 2017). This tool detects these regions based on sequence signatures characterized by four consecutive (though sometimes imperfect) guanine runs, separated by variable-length loops (Labudová et al. 2020). The classical G4-forming motif typically fits the pattern G≥3N1-7G≥3N1-7G≥3N1-7G≥3, where ’G’ denotes guanine tracts of at least three residues, and ’N’ denotes intervening loop sequences of 1-7 nucleotides (Hon et al. 2017). Unlike earlier methods, pqsfinder can also detect non-canonical G4s that include imperfections, such as bulges or longer loops, broadening the range of identifiable G4 structures (Labudová et al. 2020).

To map the G4 landscape across vertebrates, we quantified the G4-formation potential across 105 species spanning four major clades (including 57 mammals, 27 aves, 17 reptiles, and 4 non-amniotes). We compared G4 potential across species using the whole-genome as background (Tables S1). Analysis of G4-forming potential across whole-genome backgrounds reveals its dependence on GC content fluctuations in vertebrate genomes. However, since these species share evolutionary history and aren’t independent data points, we also calculated Phylogeny Independent Contrasts (PICs) alongside raw correlations (Species phylogenetic tree in Fig. S1 and tree file as additional file 1). We further quantified shared ancestry effects using Pagel’s Lambda (λ) phylogenetic signal.

To investigate how genomic background influences G4 frequency, we plotted pG4 density (per Mb) against GC content. First, we examined pG4 density (per Mb) variation across all vertebrates, followed by each clade’s specific contribution to the overall trend (Fig. 1a- e; Panel 1: pG4 density vs Genomic GC content). We found a strong positive correlation between pG4 density and genomic GC content (Fig. 1a; ρ = 0.56, P < 0.001; full statistics for Fig. 1 in Table S2). A strong phylogenetic signal (λ ≈ 0.90) confirms that global pG4 patterns are heavily shaped by ancestry. PIC analysis showing evolutionary link: for every 1% rise in GC content, pG4 density increases by ∼118.9 units (Fig. 1a; PIC R_adj_² = 0.35). To identify key clade contributions, we examined deviations from this trend: Mammals have a strong ancestral constraint (λ ≈ 1.0) on pG4 density in their genomes with moderate correlation (Fig. 1b; ρ = 0.44, P < 0.001), but a lower sensitivity to the GC-drifts in the genomes (Fig. 1b; PIC R_adj_² = 0.17, β = 47.89). Avian genomes uniquely exhibit weak phylogeny (λ ≈ 0.16) with a strong correlation (Fig. 1c; ρ = 0.66, P < 0.001) and the highest evolutionary sensitivity (Fig. 1c; PIC R_adj_² = 0.68, β = 283.12) towards the pG4 density with GC-flux of the genome. Reptilian genomes display the tightest raw association (Fig. 1d; ρ = 0.87, P < 0.001) along with high phylogenetic signal (λ ≈ 1.09), and strong evolutionary coupling (Fig. 1d; PIC R_adj_² = 0.57, β = 103.23). While in the early vertebrates (Non-amniotes), the relationship is non-significant across all metrics, likely due to high genomic heterogeneity (Fig. 1e; ρ = 0.40, λ = 1.25, PIC P > 0.05).

**Figure 1.**
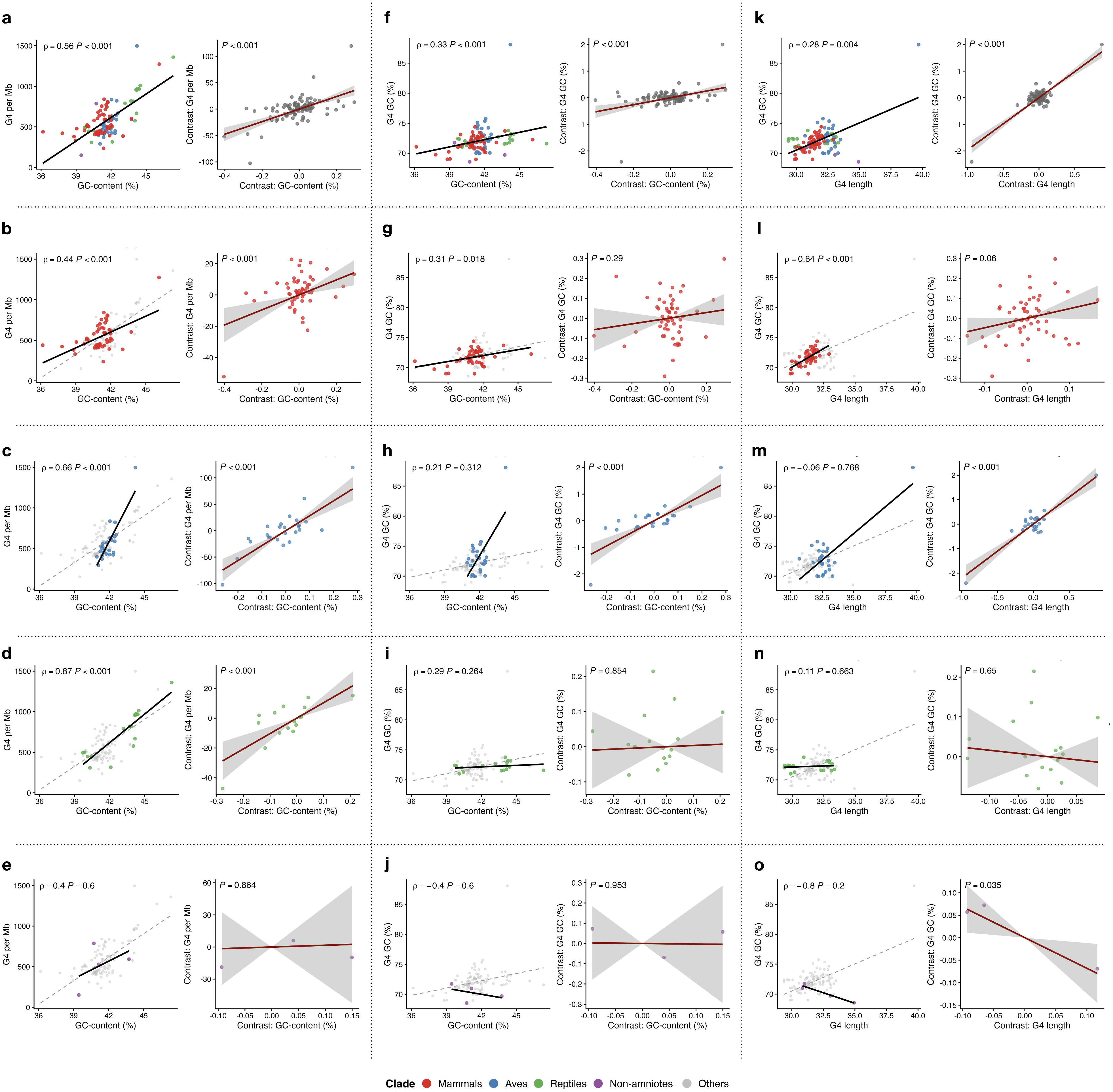
Potential G4-forming regions density coevolve with genomic GC content across vertebrates. Three panel scatter plots comparing pG4s features across 105 vertebrate genomes (57 mammals, 27 aves, 17 reptiles, 4 non-amniotes). Each panel left column represents raw correlation plots (a-o) and right column (a-o) represents Phylogeny Independent Contrasts (PICs). Panel 1: pG4 density per Mb vs. Genomic GC content. (a): Across the vertebrates pG4 density is strongly correlated with genomic GC content (ρ=0.56 (P < 0.001), PIC R_adj_² = 0.35, P < 0.001, β=118.9 ). (b): Mammals shows moderate correlation (ρ = 0.44, P < 0.001), with lower sensitivity to the GC-drifts in the genomes (PIC R_adj_² = 0.17, β = 47.89, P < 0.001). (c): Avian genomes show hypersensitivity with a strong raw correlation (ρ = 0.66, P < 0.001) and higher evolutionary coupling (PIC R_adj_² = 0.68, β = 283.12, P < 0.001). (d): Reptiles show very strong correlation (ρ = 0.87, P < 0.001) along with a strong evolutionary coupling (PIC R_adj_² = 0.57, β = 103.23, P < 0.001). (e): Non-amniotes show no significant relationship for this feature (ρ = 0.40, PIC P > 0.05). Panel 2: pG4 GC content vs. Genomic GC content correlation represents if pG4s GC content tracks genomic GC changes. (f): Globally there is moderate correlation with a weak evolutionary coupling (ρ = 0.33, PIC R_adj_² = 0.16, P < 0.001). Clade specific results shows the contribution to raw correlation from homeotherms (mammals and aves) while evolutionary coupling only with aves. (g): Mammals genomes GC show no evolutionary coupling with pG4s GC (ρ = 0.31, P < 0.018; PIC R_adj_² ≈ 0, P > 0.05). (h): Aves show stronger evolutionary coupling (ρ = 0.21, P > 0.05; PIC R_adj_² ≈ 0.62, P < 0.001). In poikilotherms (reptiles and non-amniotes) pG4 GC content is evolutionary decoupled from genomic GC-flux (i): Reptiles (ρ = 0.29, P > 0.05; PIC R_adj_² ≈ -0.06, P > 0.05), (j): Non-amniotes (ρ = -0.4, P > 0.05; PIC R_adj_² ≈ -0.5, P > 0.05). Panel 3: pG4 GC content vs. pG4 length, represents how pG4s GC content varies along with their length. (k): Global trends shows that pG4 GC content increases with increase in its length (ρ = 0.28, P < 0.001; PIC R_adj_² = 0.69, P < 0.001). (l): Mammals shows strong raw correlation but fails to show evolutionary coupling (ρ = 0.64, P < 0.001; PIC R_adj_² = 0.047, P = 0.06). (m): Aves show no significant correlation (ρ = -0.06, P > 0.05), but PIC shows strong evolutionary coupling (R_adj_² = 0.81, P < 0.001). (n): Reptiles show no correlation (ρ = 0.1, P > 0.05, PIC R_adj_² = - 0.05, P > 0.05) between pG4s GC content variation with its length. (o): Non-amniotes on the other hand shows that longer pG4s tend to have lower GC content (ρ = -0.8, P > 0.05, PIC R_adj_² = 0.9, P = 0.03). Clades colors are represented as follows: Mammals (Red), Aves (blue), Reptiles (green), and Non-amniotes (purple) and other are shown as grey points highlighting color of the specific clade. Dashed line in raw correlation represents global trends, shaded area around the trend line in PIC represents 95% confidence interval for evolutionary regression. Full statistics in Table S2.

Second, to test whether G4 motif GC content tracks background genomic GC changes, we plotted pG4 GC against genomic GC (Fig. 1f-j, Panel 2). Globally, this shows moderate correlation (Fig. 1f; ρ = 0.33, P < 0.001; full statistics for Fig. 1 in Table S2), with a weak but positive evolutionary coupling (Fig. 1f; PIC R_adj_² = 0.16, P < 0.001), suggesting pG4s become GC-rich as genomes do. Clade-specific analysis reveals striking differences. Mammalian pG4 composition is decoupled from the background GC-drift (Fig. 1g; ρ = 0.31, P < 0.018; PIC R_adj_² ≈ 0, P > 0.05), with strong phylogenetic conservation (λ ≈ 0.999) resisting genomic GC changes. Avian pG4s show hypersensitivity to background GC (Fig. 1h; ρ = 0.21, P > 0.05; PIC R_adj_² ≈ 0.62, P < 0.001), with near-zero λ indicating composition responds purely to current genomic environment. Like mammals, pG4 composition is uncoupled from the background GC-flux (Fig. 1i; ρ = 0.29, P > 0.05; PIC R_adj_² ≈ - 0.06, P > 0.05, λ ≈ 0). Compositional evolution of pG4 in non-amniotes was not significant and showed no correlation with the background genome composition (Fig. 1j).

Third, we examined whether pG4 GC content varies with length (Fig. 1k-o, Panel 3: pG4 GC content vs length). A universal feature emerged across vertebrates, depicting that longer pG4s are more GC-rich (Fig. 1k; ρ = 0.28, P < 0.001; PIC R_adj_² = 0.69, P < 0.001, full statistics for Fig. 1 in Table S2). Mammals show a strong raw correlation (Fig. 1l; ρ = 0.64, P < 0.001), but the relationship disappears after phylogeny correction (Fig. 1d; PIC R_adj_² = 0.047, P = 0.06), suggesting ancestral history (λ ≈ 0.999) rather than active evolutionary optimization. While raw correlation shows no significant correlation in aves (Fig. 1m; ρ = -0.06, P > 0.05), PIC analysis depicts strong evolutionary coupling (Fig. 1m; R_adj_² = 0.81, P < 0.001). It appears that in aves the composition of pG4s adjusts optimally regardless of their phylogenetic position (λ ≈ 0). Poikilotherms (reptiles and non-amniotes) on the other hand show no evolutionary coupling (reptiles) or show negative correlation (non-amniotes). Reptiles show no significant evolutionary coupling of pG4 length and its GC composition (Fig. 1n; ρ = 0.1, P > 0.05, PIC R_adj_² = -0.05, P > 0.05). Interestingly, non-amniotes show longer pG4s becoming GC-poorer (Fig. 1o; ρ = -0.8, P > 0.05, PIC R_adj_² = 0.9, P = 0.03).

These results suggest pG4s is not randomly distributed but is strongly influenced by the underlying GC-content of the genome. Although the global pattern shows GC-content as a primary driver of G4 evolution, our clade-level breakdown reveals that different lineages respond differently to the genomic influence, from relative conservation in mammals to higher evolutionary flexibility observed in birds. While the genome-wide analysis establishes GC content as the main engine propelling pG4 evolution, it does not account for the arrangement of these motifs within the functional architecture of the genomes. If pG4s were simply mutational byproducts, we would expect a uniform distribution across all genomic compartments. Their established roles in biological regulation predict non-random enrichment in key regulatory regions instead. Therefore, we mapped pG4 density across functional elements of the genome, including UTRs, exons, introns, transcription start sites (TSSs), and entire genes, to determine whether G4 potential is prioritized in response to selective pressures in specific regulatory regions.

### Lineage-specific enrichment and positional bias of pG4 density within functional genomic elements

While mapping the pG4 density across localized genomic elements, we normalized all feature-specific pG4 densities to the species-specific genomic GC content to distinguish functional pG4 selection from background GC-bias-driven artifacts.

We began by examining G4 formation potential flanking transcription start sites (TSSs), as prior studies have shown selective constraints in promoter regions associated with gene activation. We therefore tested whether this feature evolved at specific timepoints in vertebrates. We analyzed pG4 density in 50-bp windows across TSS ±2 kb regions, normalized to species-specific GC content (Fig. 2a; details in Table S3). Each bin represents the mean normalized pG4 density across all TSSs per species. Interestingly, pG4 density flanking TSSs shows a bimodal distribution preserved across amniotes but absent in non-amniotes (Fig. 2a). Consistent with prior studies, we observed significant enrichment in proximal promoter regions with a characteristic trough at the TSS, but demonstrate for the first time its conservation across vertebrates, likely maintaining transcriptional accessibility. A clear evolutionary gradient emerges in normalized G4 potential at ±2 kb flanking TSSs: highest in aves, then mammals, reptiles, and lowest in non-amniotes (Fig. S2, P < 0.05 vs mammals, Table S4). This pattern suggests regulatory pG4 roles in homeothermic promoters (mammals and aves), while poikilothermic promoters, particularly non-amniotes, show reduced pG4 dependence. The corresponding heatmap illustrates this progressive dissipation from homeotherms to poikilotherms (Fig. S3).

**Figure 2.**
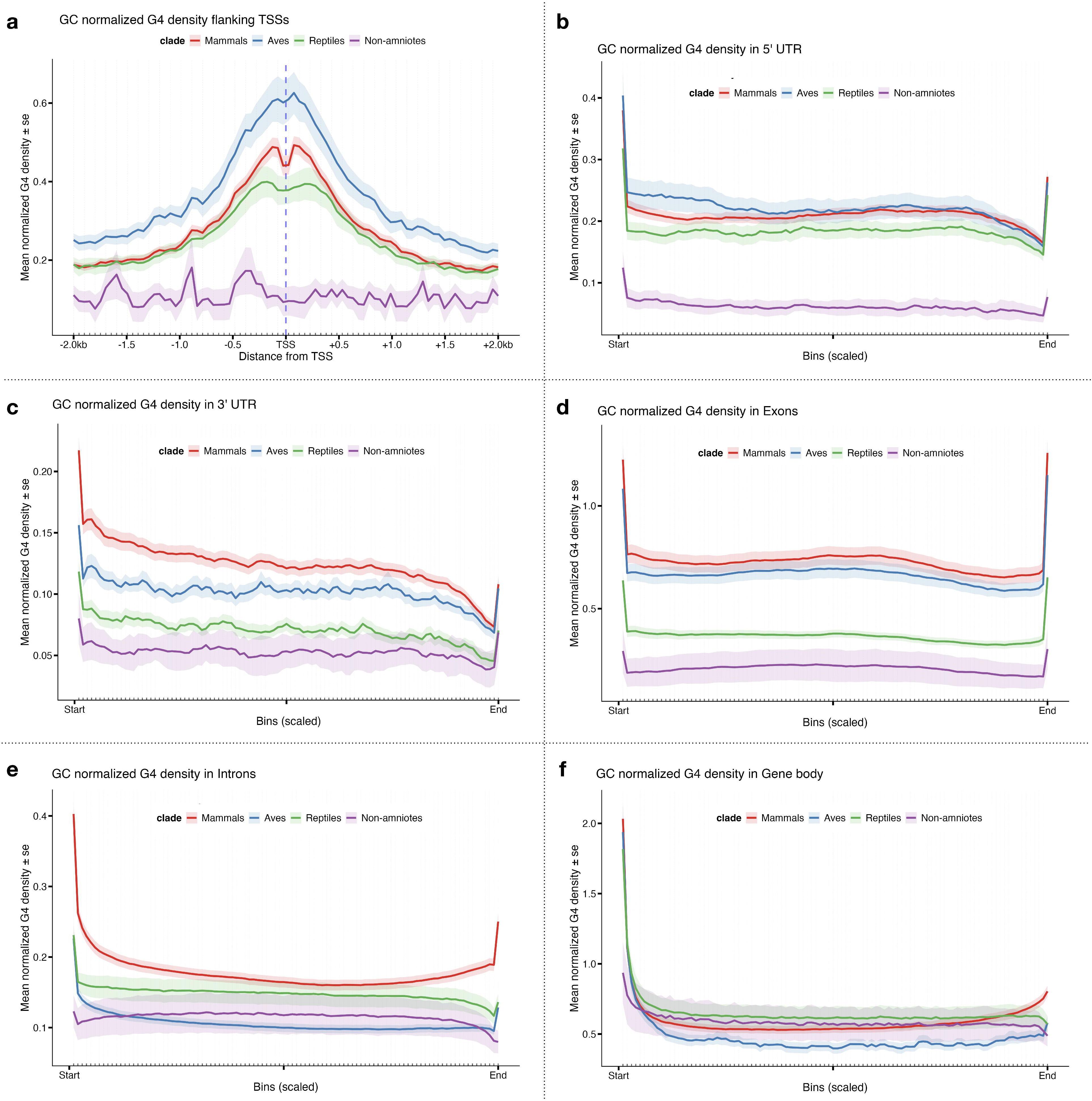
pG4 density across functional genomic elements reveals clade specific evolutionary patterns in vertebrates. Plot profiles showing GC-normalized mean pG4 density across 105 vertebrate genomes (57 mammals, 27 aves, 17 reptiles, 4 non-amniotes). (a): GC-normalized mean pG4 density flanking TSS ±2kb shows bimodal distribution conserved in amniotes and showing enrichment at proximal promoters and a peculiar trough at TSS. Aves highest, then mammals > reptiles > non-amniotes (Fig. S2, P < 0.05 vs mammals, Table S4). GC normalized mean pG4 density ± se is represented on y-axis in bins of 50 bp and vertical blue dashed lines represents TSS, with x-axis representing distance from TSS upstream and downstream. (b): GC-normalized mean pG4 density in 5’ UTRs represents scaled signal of pG4 density in 100 bins with ±se. It shows remarkable amniote conservation, mammals ∼ aves >> reptiles > non-amniotes (Fig. S4, P < 0.05 vs mammals, Table S4). The lack of significant difference between aves and mammals suggests it as unified homeothermic trait. (c): GC-normalized mean pG4 density in 3’ UTRs represents scaled signal of pG4 density in 100 bins with ±se. It shows remarkable evolutionary gradient with mammals > aves > reptiles > non-amniotes (Fig. S4, P < 0.05 vs mammals, Table S4). (d): GC-normalized mean pG4 density in exons represents scaled signal of pG4 density in 100 bins with ±se, show homeotherms enhanced pG4 density across exons, mammals ∼ aves >>> reptiles > non-amniotes (Fig. S4, P < 0.05 vs mammals, Table S4). (e): GC-normalized mean pG4 density in introns represents scaled signal of pG4 density in 100 bins with ±se. Introns shows mammalian specific peaks for pG4 density, mammals > reptiles > aves ∼ non-amniotes (Fig. S7, Table S4), suggesting complex role of pG4 in splicing events in mammals. (f): GC-normalized mean pG4 density in entire gene bodies represents scaled signal of pG4 density in 100 bins with ±se, show no distinct gradient, mammals modestly > aves ≈ reptiles ≈ non-amniotes (Fig. S8, Table S4). Clade specific colors: mammals (red), aves (blue), reptiles(green) and non-amniotes (purple). To compare pG4 density across clades, we performed Wilcoxon rank-sum tests with Benjamini-Hochberg (BH) p-value adjustment for multiple comparisons, using Mammals as the reference group. Clade-stratified heatmaps illustrating species-specific pG4 density distributions across all analyzed functional elements are provided in the supplementary material (Figs. S3, S5–6, and S9–11).

Building on this promoter analysis, we examined pG4 density across transcript regions (UTRs, exons, introns, full gene body), scaling regions into 100 bins with GC normalization (details in methods, Table S3), to test whether their reported roles in transcription, splicing, and translation show clade-specific evolutionary patterns. Remarkable evolutionary conservation of pG4 density appears in 5’ UTRs of terrestrial vertebrates (Fig. 2b), suggesting an important regulatory role in gene expression. The lack of difference between mammals and aves indicates a unified homeothermic trait, while the massive drop in non-amniotes shows that high pG4 density at translation initiation sites became standardized during amniote evolution (Fig. S4, P < 0.05 vs mammals, Table S4). In homeotherms, elevated pG4 density likely controls translation efficiency. Conversely, 3’ UTRs reveal a statistically significant evolutionary gradient in pG4 density (Fig. 2c), mammals > aves > reptiles > non-amniotes (P < 0.05 vs mammals, Table S4). Clade-stratified heatmaps confirm 5’ UTR enrichment in homeotherms and mammalian 3’ UTR expansion (Fig. S5-6, respectively). Since RNA pG4s in 3’ UTRs regulate post-transcriptional processes, this gradient suggests increasing regulatory complexity in vertebrate post-transcriptional networks.

Exonic regions show a striking homeothermic bias: pG4 density in mammals and aves is indistinguishable, while poikilotherms exhibit a significant drop to nearly negligible levels (Fig. 2d; P < 0.05 vs mammals, Table S4). This pattern positions exonic pG4s as a warm-blooded hallmark, likely linked to thermal stability requirements of transcripts in homeotherms (Fig. S4). The most striking lineage divergence appears in introns. Mammals show significantly higher G4 potential than all other clades (Fig. 2e), with reptiles as an evolutionary midpoint between the lower, similar densities of aves/non-amniotes (Fig. S7, Table S4). Metagene profiles reveal sharp pG4 peaks at splice sites, suggesting mammals have uniquely recruited these structures to support their complex alternative splicing patterns (Fig. 2e). Analysis of entire gene bodies reveals no distinct enrichment gradient (Fig. 2f). Mammals exhibit modestly elevated total pG4 density relative to aves, yet comparable levels to reptiles and non-amniotes (Fig. S8, Table S4). This uniformity reflects compartment-specific sparsity in regulatory regions (5’ UTRs, exons) that offsets higher densities elsewhere across the full gene length. Lineage-specific heatmaps confirm homeothermic enrichment in exons, mammalian enrichment in introns, and amniote-specific conservation in gene bodies (Fig. S9-11). These patterns demonstrate that positional enrichment in regulatory regions drives pG4 evolutionary conservation more than genome-wide density.

While total gene body profiles show no clear evolutionary gradient, most functional elements reveal distinct clade-specific patterns, from amniote-specific segregation (5’ UTRs), homeotherm-specific enrichment (exons, promoters), to mammal-specific high signal (introns). The M-shaped curve near TSSs (Fig. 2a) underscores G4s’ regulatory roles. We therefore examined 1 kb core promoters to characterize how pG4 density and sequence composition evolve relative to genomic background.

### Intensified selection for G4 potential defines the evolutionary landscape of amniote core promoters

The striking bimodal distribution of pG4 density flanking TSSs shows that these regions in the amniotes are subjected to different evolutionary pressures. To determine if this localized significant enrichment represents a specialised regulatory mechanism, we isolated 1 kb core proximal promoters for 105 genomes and subjected them to the same phylogenetic and correlative framework used for the whole genome (Table S5). This allowed us to assess how GC drive is intensified at regulatory hubs of transcription compared to the rest of the genome.

We next examined how pG4 density varies within 1 kb core promoters across vertebrates (Fig. 3 a-e; Panel 1: pG4 density vs Promoter GC content). Across 105 species, promoter pG4 density was strongly influenced by local GC content (Fig. 3a; ρ = 0.90, P < 0.001; full statistics for Fig. 3 in Table S6). Promoters GC content explains 83% of evolutionary pG4 variation (Fig. 3a; PIC R_adj_² = 0.834, P < 0.001), twice as sensitive as the genomic average (β = 235.01). Moderate phylogenetic signal (λ ≈ 0.44) suggests pG4 promoter evolution balances ancestral inheritance with active tuning. We next examined clade-specific promoter evolution. It revelas that pG4 density dependence on promoter GC content as an amniote-specific evolutionary trait. Mammalian promoter pG4 density strongly correlates with local GC content (Fig. 3b; ρ = 0.82, P < 0.001), explaining 80% of evolutionary variance (Fig. 3b; PIC R_adj_² = 0.80, P < 0.001, β = 179.20). Unlike genome-wide patterns, where mammals resist GC drifts, promoters show heightened evolutionary sensitivity to GC changes. Moderate phylogenetic signal (λ ≈ 0.53) suggests that mammals actively recruit pG4s into GC-rich promoters beyond simple inheritance. Aves show the strongest pG4-GC co-evolution in promoters (Fig. 3c; ρ = 0.94, P < 0.001), with virtually no phylogenetic signal (λ ≈ 0). Their promoters exhibit extreme evolutionary sensitivity (Fig. 3c; PIC R_adj_² = 0.78, P < 0.001, β = 333.35), 2.5 times higher than mammals, confirming pG4 density as a highly plastic feature that rapidly responds to GC content changes. Reptiles show strong pG4-GC correlation (Fig. 3d; ρ = 0.88, P < 0.001) but strong phylogenetic constraint (λ ≈ 0.95), yielding moderate evolutionary coupling (Fig. 3d; PIC R_adj_² = 0.65, P < 0.001, β = 185.02), similar to that of mammals. Non-amniotes promoters show no evolutionary coupling of pG4 density with increase in promoter GC content (Fig. 3e; ρ = 0.80, P > 0.05, PIC R_adj_² = -0.02, P > 0.05). Thus, pG4-GC coupling in core promoters evolved and stabilized specifically in amniotes.

**Figure 3.**
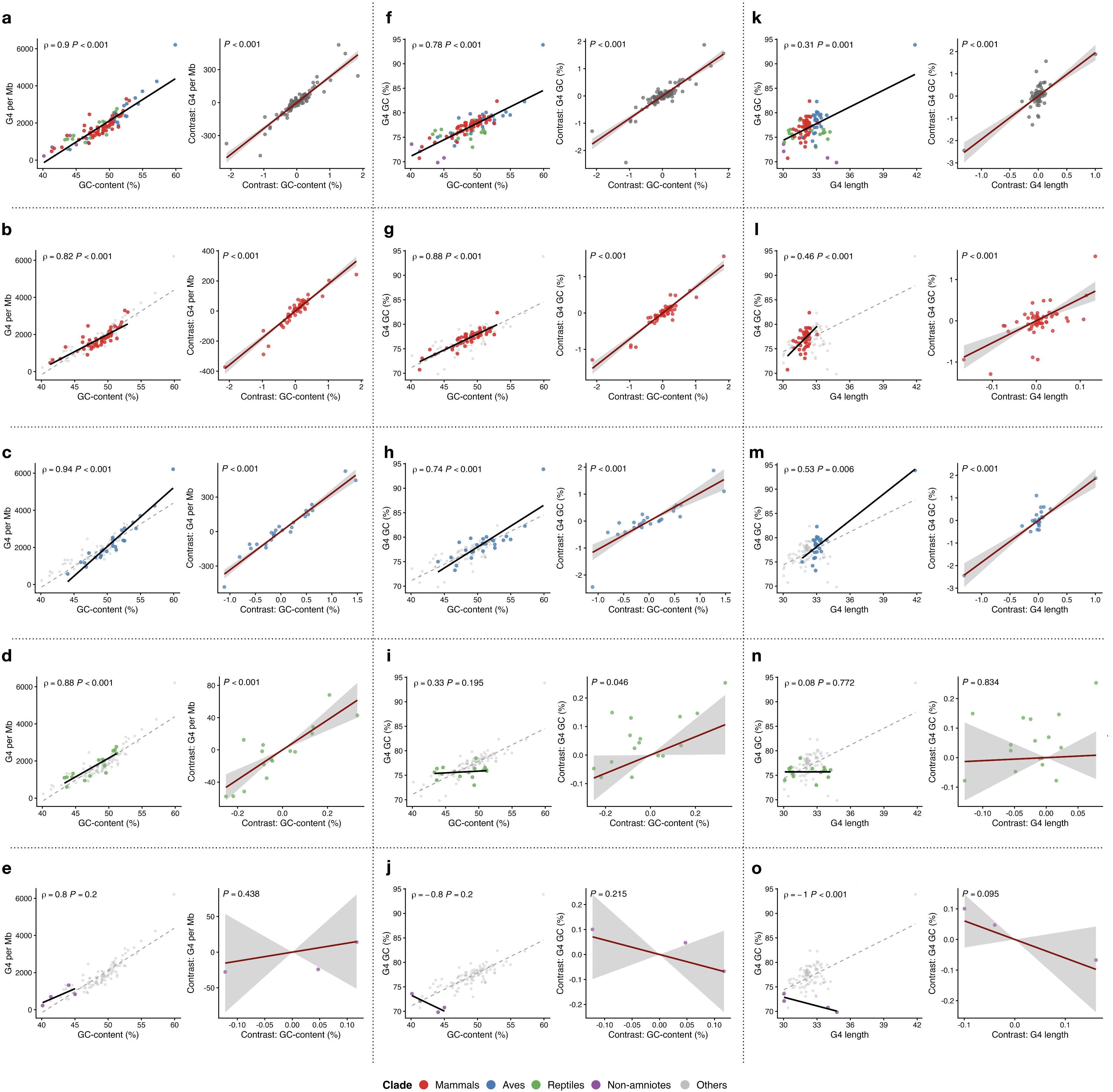
Amniote promoters show enhanced evolutionary constraints linking pG4 features to local GC content. Three panel scatter plots comparing pG4s features across 105 vertebrate genomes (57 mammals, 27 aves, 17 reptiles, 4 non-amniotes). Each panel left column represents raw correlation plots (a-o) and right column (a-o) represents Phylogeny Independent Contrasts (PICs).Panel 1: pG4 density per Mb vs. Promoter GC content. (a): Across vertebrates promoter pG4 density strongly correlates with promoter GC content (ρ = 0.90 P < 0.001, PIC R_adj_² = 0.834, P < 0.001, β = 235.01). Clade specific breakdown shows that it holds true only for amniotes. (b): Mammals show strong correlation (ρ=0.82, P < 0.001), explaining 80% evolutionary variance (PIC R_adj_² = 0.80, β = 179.20, P < 0.001). (c): Aves show strongest correlation (ρ=0.94, P < 0.001) with extreme evolutionary sensitivity (PIC R_adj_² = 0.78, β=333.35, P < 0.001). (d): Reptiles show strong correlation (ρ = 0.88, P < 0.001) with moderate evolutionary coupling (PIC R_adj_² = 0.65, β = 185.02, P < 0.001). (e): Non-amniotes show no correlation (ρ = 0.80, P > 0.05; PIC R_adj_² = -0.02, P > 0.05). Panel 2: pG4 GC content vs. Promoter GC content correlation represents if pG4s GC content tracks promoter GC changes. (f): Globally pG4 GC tightly couples with promoter GC (ρ = 0.78, P < 0.001; PIC R_adj_² = 0.751, P < 0.001, β = 0.83). (g): Mammals show reversal from genome-wide trend with strong coupling (ρ = 0.98, P < 0.001; PIC R_adj_² = 0.88, P < 0.001, β = 0.72). (h): Aves show highest plasticity (ρ = 0.74, P < 0.001; PIC R_adj_² = 0.72, P < 0.001, β=1.05). (i): Reptiles exhibit weaker coupling (ρ = 0.33, P > 0.05; PICR_adj_² = 0.18, P < 0.045, β = 0.32). (j): Non-amniotes show no significant relationship (P > 0.05). Panel 3: pG4 GC content vs. pG4 length, represents how pG4s GC content varies along with their length. (k): Global trend shows longer pG4s have higher GC content (ρ=0.31, P < 0.001; PIC R_adj_² = 0.542, P < 0.001). While clade specific analysis shows that it is evolved in homeotherms. (l): Mammals show moderate correlation with strong evolutionary slope (ρ=0.46, P < 0.001; PIC R_adj_² = 0.41, P < 0.001, β = 5.35). (m): Aves exhibit strong coupling (ρ=0.46, P < 0.001; PIC R_adj_² = 0.781, P < 0.001, β = 1.85). (n): Reptiles lack significant relationships (P > 0.05). (o): Non-amniotes lack significant relationships (P > 0.05). Clades colors are represented as follows: Mammals (Red), Aves (blue), Reptiles (green), and Non-amniotes (purple) and others are shown as grey points highlighting color of the specific clade. Dashed line in raw correlation represents global trends, shaded area around the trend line in PIC represents 95% confidence interval for evolutionary regression. Full statistics in Table S6.

Further, we tested whether pG4 GC content varies with promoter GC across vertebrates (Fig. 3f-j, Panel 2). Globally, pG4 GC tightly couples with promoter GC (Fig. 3f; ρ = 0.78, P < 0.001; Table S6). Phylogenetically, background GC explains 75% of pG4 GC variation (Fig. 3f; PIC R_adj_² = 0.751, P < 0.001, β = 0.83). Clade breakdown reveals striking patterns: Mammals show a striking reversal in their genome-wide trend, where pG4 composition was ancestry-locked, promoters show near-zero phylogenetic signal (λ ≈ 0) with strong coupling (Fig. 3g; ρ = 0.98, P < 0.001) and 88% evolutionary variance explained (Fig. 3g; PIC R_adj_² = 0.88, P < 0.001, β = 0.72). This suggests mammalian promoter pG4s freely adapt to local GC environments. Aves show even higher plasticity (Fig. 3h; ρ = 0.74, P < 0.001; PIC R_adj_² = 0.72, P < 0.001, β = 1.05), every 1% GC increase triggers pG4 GC response with no phylogenetic constraint (λ ≈ 0), marking avian promoters as most evolutionarily flexible. Reptiles exhibit weaker coupling (Fig. 3i; ρ = 0.33, P > 0.05; PIC R_adj_² = 0.18, P < 0.045, β = 0.32), while non-amniotes show no significant relationship (Fig. 3j, P > 0.05). These patterns demonstrate that tight co-evolution between pG4 composition and promoter GC content represents a derived feature of homeothermic amniotes.

Finally, we tested whether pG4 GC content correlates with pG4 length in promoters (Fig. 3k-o, Panel 3). Across vertebrates, longer pG4s have higher GC content (Fig. 3k; ρ = 0.31, P < 0.001; Table S6), with length predicting 54% of evolutionary GC variation (Fig. 3k; PIC R_adj_² = 0.542, P < 0.001). Clade patterns reveal homeothermic specialization: Mammals show moderate correlation (Fig. 3l; ρ = 0.46, P < 0.001) but striking evolutionary slope (Fig. 3l; PIC R_adj_² = 0.41, P < 0.001, β = 5.35). Every unit length increase drives GC enrichment, free from ancestral constraints (λ ≈ 0). Aves exhibit even stronger coupling (Fig. 3m; ρ = 0.46, P < 0.001), with length explaining 78% of GC variance (Fig. 3m; PIC R_adj_² = 0.781, P < 0.001, β = 1.85), and cannot be explained by phylogeny (λ ≈ 0). In stark contrast, poikilotherms lack this coupling. Neither reptiles nor non-amniotes show significant pG4 length-GC relationships (Fig. 3n-o, P > 0.05). pG4 GC-length coupling thus represents a homeotherm-specific innovation, tightly regulated in amniote promoters.

These results reveal two distinct pG4 evolutionary modes in vertebrates. Genome-wide, pG4 density couples tightly to GC content but remains phylogenetically constrained, particularly in mammals. In core promoters, G4 potential instead evolves in lockstep with local GC changes, free from ancestral constraints (λ ≈ 0). Both mammals and aves show this dynamic promoter regulation, contrasting their more conserved genome-wide patterns and highlighting powerful selective pressure on promoter pG4 architecture. The intense selective pressure maintaining promoter GC richness suggests dedicated mechanisms actively preserve these GC-rich, G4-prone regions. CGGBP1, a GC-rich binding protein, is known to prevent GC-loss genome-wide and specifically within transcription factor binding sites in 1 kb cis promoters of certain target genes. The GC retention function of CGGBP1 is associated with its role in preventing the meC-to-T transition. Additionally, CGGBP1 preferentially binds regions with GC asymmetry that have the potential to form G4 structures. Notably, depletion of CGGBP1 leads to increased G4 formation, indicating that CGGBP1 acts to suppress G4 formation. Therefore, potential G4-forming regions in the human genome likely exhibit these features and show dependence on CGGBP1 activity.

### Genome-wide GC preservation in pG4 regions via meC-to-T suppression is shaped by CGGBP1 evolutionary divergence

We next focused on potential G4-forming regions identified by pqsfinder in the human genome to investigate whether evolutionary forms of CGGBP1 influence methylation-associated GC-loss in these regions. We leveraged recently published MeDIP-seq datasets (GSE281704) to examine genome-wide cytosine methylation alterations mediated by vertebrate CGGBP1 orthologs expressed in human HEK293T cells. Following endogenous human CGGBP1 knockdown, orthologs from key evolutionary lineages, non-amniote (*Latimeria chalumnae*), reptile (*Anolis carolinensis*), avian (*Gallus gallus*), and mammalian (*Homo sapiens*), were ectopically expressed in HEK293T cells. We quantified point mutation rates in G4-forming regions using the Methylation Associated Point-Mutation Assessment Pipeline (MAP-MAP) pipeline to map methylation-driven point mutations affecting genome-wide cytosine methylation, a computational framework designed to profile mutation dynamics in methylation patterns across the experimental samples, we saw an overall increase in mutation rates ( Table S7), with the highest rates observed in *Homo sapiens* (Hs) CGGBP1 (Fig. S12, Table S8).

CGGBP1 is known to restrict cytosine methylation and meC-to-T transition rates genome-wide. Analysis of meC-to-T transitions within potential G4-forming regions revealed that endogenous CGGBP1 (Ev) limits these transitions, thereby reducing GC-loss in these sequences (Fig. 4a). Evolutionary forms of CGGBP1 differ in this capacity: evolutionary forms of CGGBP1 exhibit differing abilities in this regard: *Latimeria chalumnae* (Lc) and *Anolis carolinensis* (Ac) forms were unable to prevent meC-to-T transitions in human cells, *Gallus gallus* (Gg) showed partial inhibition, and *Homo sapiens* (Hs) displayed the strongest restriction, minimizing these transitions to the greatest extent (Fig. 4a, Table S8). Notably, over-expression of Hs markedly minimized all mutations leading to GC-loss (Fig. S12). These findings suggest that human CGGBP1 prevents genome-wide GC-loss, including in potential G4-forming regions, by restricting meC-to-T transitions, a function likely acquired through evolution. This mechanism may contribute to the preservation of both G4-forming potential and the GC-content of G4 sequences in vertebrate genomes. Our earlier findings showed that CGGBP1 preserves GC content at transcription factor binding sites in repressed promoters by limiting meC-to-T transitions, a property developed over evolutionary time. This underscores the importance of examining promoter regions, their G4-forming potential, and the impact of CGGBP1 on GC retention both in G4 structures and the broader promoter context.

**Figure 4.**
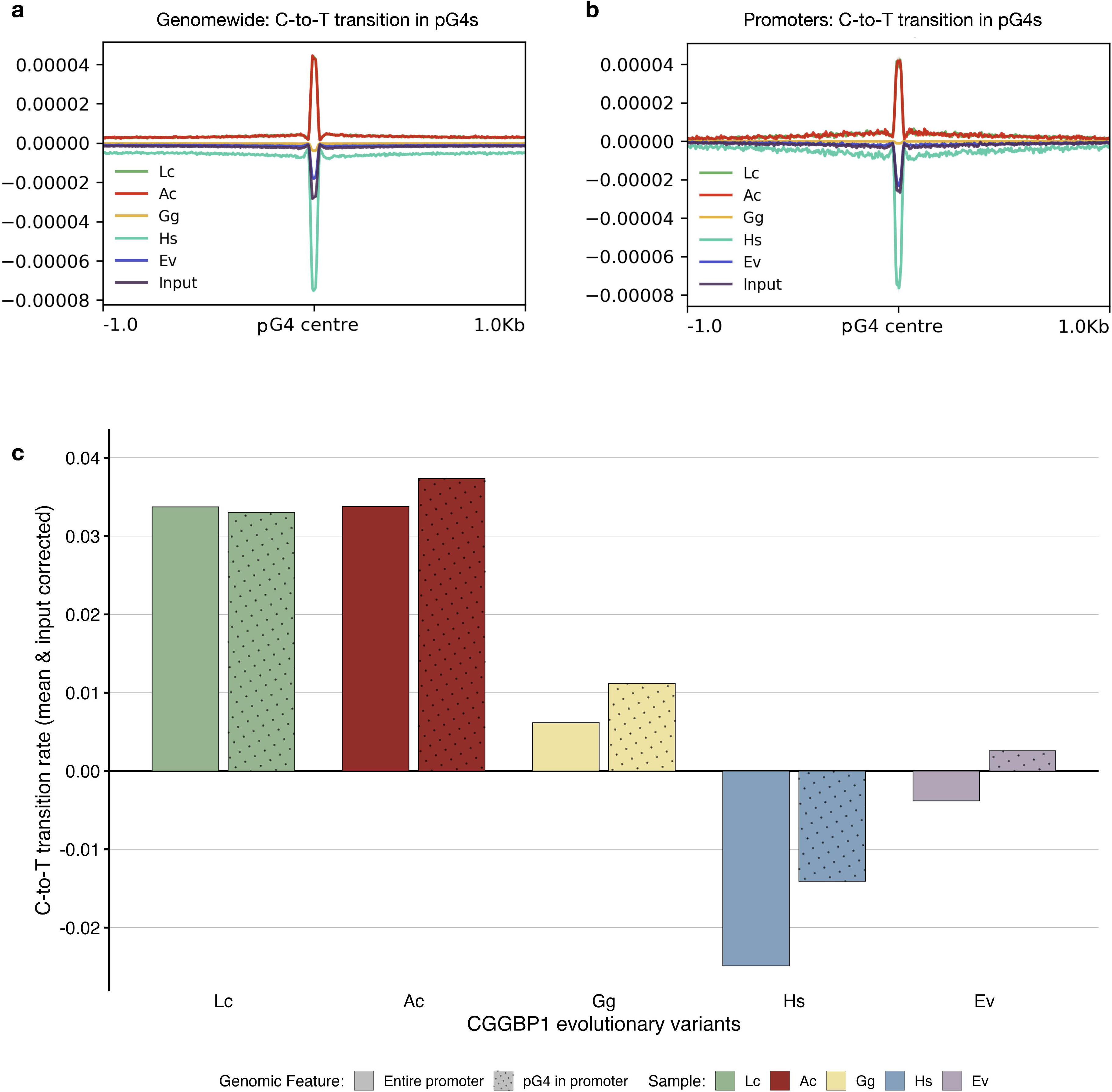
CGGBP1 evolutionary forms regulate the restriction of meC-to-T transition in methylation-enriched, pG4-forming genomic regions, distinguishing between poikilothermic and homeothermic species. (a): Genome-wide analysis of potential G4-forming regions, defined by 1 kb flanks around pG4 coordinates, reveals that meC-to-T transition rates are restricted in a CGGBP1 form-dependent manner. Samples expressing human (Hs), avian (Gg) forms of CGGBP1, and empty vector (Ev) representing endogenous CGGBP1, as well as the input control, exhibit lower meC-to-T transition rates, with Hs showing the strongest restriction (lowest transition rate). In contrast, poikilothermic forms (Lc and Ac) display positive meC-to-T transition rates, indicating less restriction. The plotted data show genome-wide meC-to-T transition signals for all samples (Lc, Ac, Gg, Hs, Ev, and Input) across potential G4-forming regions. (b): Potential G4-forming motifs within 1 kb promoters of the human genome also exhibit CGGBP1 form-dependent restriction of meC-to-T transitions. Human CGGBP1 is the most effective at limiting methyl-cytosine to thymine mutation rates, mirroring the genome-wide pattern, whereas Lc and Ac CGGBP1 forms do not achieve similar restriction. Gg and Ev reduce transition rates to a degree, but both still maintain a higher mutation rate than the input sample, which reflects the baseline background mutation rate in non-methylated DNA. (c): meC-to-T transition rates as a fraction of all point mutations in 1 kb gene promoters and in predicted G4-forming regions (P < 0.01; Table S9-12) show significant differences depending upon evolutionary forms of CGGBP1, as indicated by chi-square testing (P < 0.01; Table S13). pG4-forming regions within promoters display differential meC-to-T transition rates as compared to background promoters, and both endogenous and overexpressed human CGGBP1 demonstrate strong restriction of meC-to-T transitions relative to other forms. Gg and Ev moderately restrict mutation rates, while poikilothermic forms (Lc and Ac) fail to do so. Hs CGGBP1 maintains the lowest meC-to-T rates in both pG4-forming regions and background promoters, though meC-to-T rates are still marginally but significantly higher in pG4-forming regions (Table S13), whereas Lc shows higher meC-to-T in promoters versus corresponding pG4-forming regions (Table S13). Ac, Gg, and Ev do not show statistically significant differences in promoters, though for Gg and Ev, meC-to-T appears higher in pG4-forming regions, opposite to Ac (Table S13). To highlight methylation-specific changes, input as background was subtracted from both promoters and pG4-forming regions, and these normalized meC-to-T transition rates are shown as bar graphs: Lc (green), Ac (red), Gg (yellow), Hs (blue), and Ev (purple), with solid fill for promoters and dots fill for potential G4-forming regions found in promoters.

We further focused on all annotated 1 kb promoters in the human genome and their potential G4-forming regions. The predicted G4 sequences within these promoters exhibited a CGGBP1 evolutionary form-dependent restriction of meC-to-T transition rates (Fig. 4b). This pattern mirrored the genome-wide trend, with Hs CGGBP1 acting as the strongest suppressor of cytosine methylation-induced transitions in predicted G4-forming sequences. The endogenous CGGBP1 represented by empty vector (Ev) showed moderate restriction compared to input controls, Gg displayed a weaker effect, while Lc and Ac failed to restrict these transitions effectively (Fig. 4b). These differences correspond to previously reported distinctions between homeothermic and poikilothermic vertebrates in GC retention mediated by CGGBP1’s evolutionary forms, particularly through the restriction of meC-to-T transitions in transcription factor binding sites. GC asymmetry is a characteristic sequence feature associated with CGGBP1-regulated regions. GC asymmetry measured as interstrand GC-skew is known to overcome the ds of DNA and form G4. To explore this further, we examined GC-skew in potential G4-forming regions genome-wide, within 1 kb cis promoter G4-forming regions, and in regions lacking potential G4 formation. Our analysis revealed that potential G4-forming regions exhibit higher GC-skew overall. Specifically, genome-wide predicted G4-forming regions display greater GC-skew than those within 1 kb promoter G4-forming regions (Fig. S13). In contrast, regions without potential G4 formation showed no significant enrichment of GC-skew (Fig. S13). Mutation-free regions across MeDIP samples exhibit varying G4 formation potential depending on the over-expressed CGGBP1 variant (Fig. S14). Hs maintains the highest G4-forming potential among these, accompanied by detectable GC-skew in these regions, although the skew is not centrally enriched (Fig. S14). In contrast, methylation-enriched, mutation-free regions lack GC-skew, with only Hs showing GC-skew comparable to the genomic background. For Lc, Ac, Gg, and Ev, G4-forming potential decreases significantly alongside a marked reduction in GC-skew (Fig. S14). This means that mutation-free regions enriched in MeDIP DNA in different samples in Hs show the highest G4 formation capability with retention of GC-skew as that of the genomic background.

Next, we compared meC-to-T transition rates in potential G4-forming regions to those in background promoter sequences by examining overall point mutation rates across entire 1 kb promoter regions, as well as within G4-forming segments located in promoters (Table S9). In both analyses, we observed that homeothermic samples (Hs and Gg) displayed increased point mutation rates that favored GC retention, distinguishing them from the poikilotherms (Lc and Ac) (Fig. S15a-b). This trend, preferential GC retention, was evident both at the level of whole promoters and within their potential G4-forming regions. All observed point mutations, expressed as fractions of all mutations within each sample, were statistically significant for both predicted G4s in promoters and the background (Table S10-11). In all samples, meC-to-T transition rates differed significantly in both promoter regions and their potential G4s (Table S12).

To further discern whether meC-to-T transitions are more prevalent in promoters as a background or specifically within G4s, we compared meC-to-T to all point mutations in both contexts. Results showed that in homeotherms (Hs and Gg), G4-forming regions were more prone to meC-to-T transitions than the broader promoter background, whereas in poikilotherms (Lc and Ac), the reverse was true. This difference reached significance only for Hs among homeotherms and for Lc among poikilotherms (Table S13). After adjusting for input controls, potential G4-forming sequences maintained higher meC-to-T transition rates compared to background regions, in line with the evolutionary dependence on CGGBP1, except for Lc (Fig. 4c). Overexpression of Hs CGGBP1 resulted in the strongest restriction of meC-to-T transitions in both promoters and predicted G4 regions, followed by the endogenous CGGBP1 level and then Gg, while Lc and Ac showed the highest mutation rates in both regions (Fig. 4c). This elevated mutation rate in G4 regions is expected, as G4 formation requires the local unwinding and separation of the two DNA strands, exposing single-stranded DNA (ssDNA). ssDNA is more vulnerable to various mutagenic processes, including spontaneous deamination of methylated cytosines, which leads to increased meC-to-T transitions. Thus, the structural dynamics essential for G4 formation inherently increase the susceptibility of these regions to methylation-associated mutations. After confirming the impact of CGGBP1 on GC preservation via limiting meC-to-T transitions in potential G4-forming promoters and genome-wide regions, it is essential to realize that these sites harbor G4-forming potential but may not always form G4 structures. Consequently, to study methylation effects on G4s, experiments must account for both chromatin-dependent environments influenced by CGGBP1 and settings devoid of nuclear constraints.

We analyzed a publicly available dataset (GSE202456) of experimentally validated G4 regions formed under native nuclear conditions (G4-IP-nat), which capture true functional G4 sites despite chromatin favoring dsDNA. These differ from *in vitro* G4 potential (G4-IP-Denat) and dsDNA controls. Homeothermic CGGBP1 restricts meC-to-T mutations at native G4 sites. Human (Hs) and chicken (Gg) CGGBP1 strongly suppressed C-to-T transitions here, while coelacanth (Lc) and lizard (Ac) variants did not (Fig. S16a). In contrast, denatured G4 sites showed no such restriction by any ortholog, confirming CGGBP1 targets chromatin-constrained G4s (Fig. S16b). Native G4 regions also showed high central GC-skew amid low-skew flanks, unlike denatured sites with background skew (Fig. S17). This local skew likely overcomes dsDNA barriers to enable functional G4s. We then examined Differentially Captured Regions (DCRs) from CGGBP1 modulation studies, GC-rich sites with a strong G4 signal upon CGGBP1 depletion. Homeothermic CGGBP1 (Hs, Gg) restricted C-to-T mutations in DCRs, while poikilothermic forms did not (Fig. S16c; Table S14). Even endogenous Hs CGGBP1 suppressed these transitions versus input.

These results establish homeothermic CGGBP1 orthologs as potent suppressors of meC-to-T mutations at functional G4-forming regions. Human and avian CGGBP1 (Hs, Gg) consistently restrict C-to-T transitions across native G4 sites, while poikilothermic variants (Lc, Ac) lack this capacity.

This evolutionary innovation directly explains the enhanced GC retention we documented in homeotherm promoters and G4 landscapes. Since CGGBP1 also protects transcription factor binding sites from GC loss in targeted promoters, we next asked whether this homeothermic bias shows up in regulatory elements too.

### Potential G4-forming regions in vertebrates show homeotherm-specific GC retention in TFBS

CGGBP1 protects TFBS from GC erosion by restricting C-to-T transitions in target promoters. We analyzed MeDIP datasets (GSE281704) from cells expressing evolutionary CGGBP1 orthologs. We performed FIMO motif enrichment for GC-rich motifs (GC > 50%) genome-wide, extracted motif hit coordinates, and applied the MAP-MAP pipeline to quantify meC-to-T transition rates within these sites (Table S15). Interestingly, homeothermic variants (Hs, Gg) showed GC retention bias across most TFBS (Fig. S18), unlike poikilothermic forms (Lc, Ac). G4-forming sequences are known hubs for transcription factor binding. We therefore asked whether the homeothermic GC retention bias extends to TFBS motifs within genome-wide pG4 regions across vertebrates, including their 1 kb *cis* promoter contexts. If CGGBP1 drives the promoter GC retention we observed, TFBS in pG4 regions should exhibit pronounced homeotherm-specific GC preservation, the functional signature of mutation suppression at regulatory elements.

We next performed motif enrichment analysis across 105 vertebrate genomes, focusing on potential G4-forming regions (pG4) genome-wide and within 1 kb promoter regions (identified by pqsfinder). For each, we extracted FASTA sequences from pG4 regions and ran FIMO to identify TFBS motifs (Table S16). Controls matched pG4 regions by length and number using G4-free sequences, genome-wide non-pG4 regions, and 1 kb promoter regions lacking pG4 as backgrounds (Table S16). We subtracted background motif densities from pG4 motif densities (Δ = pG4 motif density - pG4-free background density) to identify true pG4-associated motifs (Table S17). Homeotherms vs poikilotherms were analyzed separately to test our hypothesis of homeotherm-specific GC retention. All homeotherm species were pooled per motif, as were all poikilotherms. We calculated mean motif densities in pG4 regions vs their respective backgrounds, then computed fold-changes and statistical significance after background subtraction (Fig. 5a; Table S18). We observed differentially enriched or depleted motifs in pG4 regions of homeotherms versus poikilotherms after background correction (Fig. 5a). We next asked whether these differentially enriched motifs (either homeotherm- or poikilotherm-specific) exhibit GC content bias that could explain their evolutionary patterns. To test this, we quantified GC retention bias separately for homeotherms and poikilotherms across both genome-wide and 1 kb promoter contexts. From pG4 regions identified by pqsfinder, we extracted all FIMO-reported motif match sequences for each thermal group and calculated the mean GC content per motif across species (Table S19).

**Figure 5.**
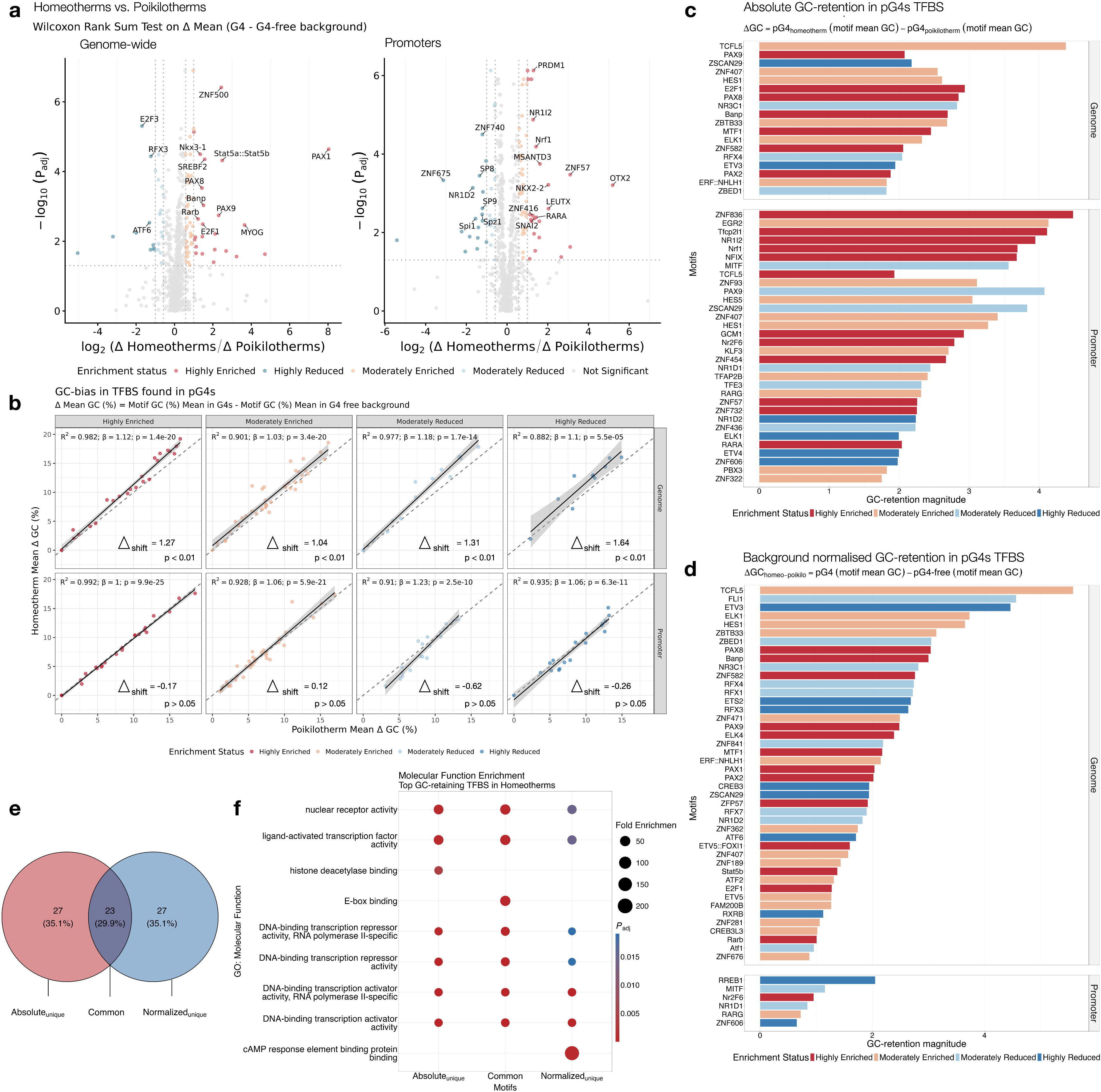
Figure 5. Enrichment of TFBS within GC-retaining pG4 regions reveals homeotherm-specific evolutionary coupling. (a): Volcano plots representing TFBS enrichment in pG4 across genomewide (left) and promoters (right) as compared to the background (pG4 free) in homeotherms vs. poikilotherms. Enrichment was calculated using the Wilcoxon Rank Sum Test comparing the mean Δ density (pG4-associated density minus pG4-free background density) between homeotherms and poikilotherms. Significance was adjusted for multiple testing using the Benjamini-Hochberg (BH) method (P_adj_). The X-axis shows homeotherm vs poikilotherm motif preference (log_2_(Homeotherm Δ / Poikilotherm Δ)). Positive values indicate motifs preferred in homeotherms, while negative values indicate preference in poikilotherms as compared to the background. The y-axis represents significance (-log_10_ P_adj_). Horizontal dashed line represents the significance cutoff (P_adj_ = 0.05). Vertical dotted line represents the biological effective thresholds for moderate ± 0.58 log_2_ FC and ± 1.0 log_2_ FC for high enrichment level. Motifs are color coded into five tiers based on their enrichment status defined by cutoff values: Highly enriched (red; P_adj_ < 0.05 and log_2_ FC > 1.0), Moderately enriched (light red; P_adj_ < 0.05 and log_2_ FC > 0.58), Moderately reduced (light blue; P_adj_ < 0.05 and log_2_ FC > -0.58), and Highly reduced (blue; P_adj_ < 0.05 and log_2_ FC > -1.0) in homeotherms. Motifs failing to meet the criteria are classified as non-significant (Grey). Top motifs with the highest divergence are labelled (details in Table S18). (b): Background-normalized GC-retention (ΔGC) in pG4-associated TFBS. Scatter plots depicting the ΔGC-bias, defined as the mean GC-content of a motif within pG4s minus its mean GC-content in pG4-free background regions. Genome-wide motifs show highly significant systematic GC-retention in Homeotherms compared to Poikilotherms across all enrichment tiers (P < 0.01, Paired Wilcoxon test), with positive mean shifts ranging from +1.04% to +1.64%. Promoter motifs exhibit higher variance in ΔGC-retention, though linear regressions confirm a strong correlation (R^2^ > 0.91) and thermal group preservation trends (β ≥ 1.0). Genome-wide (upper) and promoters (lower) are shown in different panels, with motif enrichment status in homeotherms shown below with different colors: highly enriched (Red), moderately enriched (light red), highly reduced (blue), and moderately reduced (light blue). The consistent positioning of genome-wide points above the dashed parity line demonstrates that pG4 contexts specifically amplify GC-preservation in homeothermic lineages beyond local background drift. Statistical metrics, including R-squared (R^2^), beta-slope (β), and P-values, are indicated within each panel (Table S20). (c & d): Top 50 TFBS ranked by absolute and background normalized GC retention in homeotherms vs. pikilotherms, respectively. Horizontal bar plot ranks the top 50 motifs with the largest GC increase in homeotherm pG4 regions compared to poikilotherms for absolute: (ΔGC = homeotherm mean GC% - poikilotherm mean GC%) and for background normalized: ΔGC = (homeotherm pG4 GC - homeotherm pG4-free GC) - (poikilotherm pG4 GC - poikilotherm pG4-free GC). Only Chi-square validated motifs (BH-adjusted p < 0.05) made the cut. The upper panel represents the enrichment status of these motifs in the genome, and the lower panel represents the same for promoters, with colors showing enrichment status in homeotherms (highly enriched (red), highly reduced (blue), moderately enriched (light red), moderately reduced (light blue)). Each motif in the list exhibits positive homeotherm retention, as indicated by the GC retention magnitude shown on the x-axis. Motif names are labelled on the y-axis (Table S21). (e): Common and unique absolute and background normalized overall top 50 GC retentive motifs in homeotherms as compared to poikilotherms. (f): GO enrichment of top GC-retaining pG4 TFBS. The dot plot shows Gene Ontology (GO) molecular function enrichment for top-ranked GC retentive TFBS, grouped into three categories: Common Core (top 50 in both absolute + background normalized), absolute-unique, and normalized-unique. Dot size represents fold enrichment (observed vs expected genes), dot color depicts significance (red = lower BH-adjusted p-values, all < 0.05). Top hits include “DNA-binding transcription factor activity” across all groups, plus nuclear receptors and E-box binding, showing significant enrichment.

Further, we performed three separate analyses proving homeotherms actively retain GC content in pG4 TFBS : (1) absolute GC bias in pG4-free background TFBS, (2) absolute GC bias in pG4 TFBS, and (3) background-normalized ΔGC (pG4 minus pG4-free TFBS) between homeotherms and poikilotherms (Table S20). Interestingly, raw absolute GC content shows homeotherm GC-retention bias specifically within TFBS of pG4 regions. Paired Wilcoxon tests identified significant homeotherm GC excess in 14/16 categories (p < 0.001 for 10 genome + 4 promoter; shifts +0.41-1.97%; Table S20), with pG4 regions systematically higher than pG4-free backgrounds (pG4 genome: +0.60-1.05% vs pG4-free genome: -0.22 to -1.04%; pG4 promoters: +0.41-1.80% vs pG4-free promoters: +0.67-1.97%). Furthermore, linear regressions confirmed this pattern (Fig. S19, R² > 0.94), with pG4 slopes consistently elevated (genome β = 1.01-1.07 vs pG4-free β = 0.97-1.02; promoters β = 0.93-1.07 vs pG4-free β ≈ 1.0, Table S20), demonstrating pG4 contexts amplify intrinsic homeotherm GC-retention beyond background expectations (P < 0.001, Table S20). Finally, our background-normalized ΔGC analysis (pG4 minus pG4-free TFBS) provided strong evidence of systematic homeotherm GC retention (Fig. 5b). Paired Wilcoxon tests revealed highly significant homeotherm advantage across all 4 genome-wide categories (P < 0.01, Δ-shifts +1.04-1.64%, Table S20), while promoters showed mixed patterns (3/4 suggestive, p < 0.10). Linear regressions confirmed genome-wide amplification (R² > 0.88, β = 1.03-1.18, p < 0.001, Table S20) versus promoter variability (β = 1.0-1.23), demonstrating pG4 contexts drive targeted GC preservation beyond local sequence composition (Table S20). More importantly, this positive ΔGC bias demonstrates that homeotherms actively preserve GC content in functional pG4 contexts beyond local sequence composition expectations. This mutation suppression signature was consistent regardless of whether motifs were enriched or depleted in homeotherm pG4 regions, confirming the generality of homeotherm GC retention at regulatory TFBS.

We then identified the top 50 motifs with the most significant GC-retention using absolute ΔGC and background-corrected ΔGC individually, with Chi-square validation (BH-adjusted p < 0.05), regardless of enrichment status (genome-wide/promoters; Table S21). Strikingly, all 50 top motifs exhibited higher GC-retention in homeotherms vs poikilotherms (Fig. 5c-d). Notably, 23 out of 50 motifs were common to both absolute ΔGC and background-corrected ΔGC categories (Fig. 5e, Table S21). Functional enrichment analysis of the 23 common core motifs revealed exceptional enrichment of DNA-binding transcription activators, repressors, and nuclear receptors (Fig. 5f, ), suggesting that these sequences are evolutionarily conserved in pG4s for complex epigenetic control, regardless of the background GC content (P <0.001, Table S22). Among the top 50 motifs, particularly those with absolute ΔGC bias, 32 out of 50 exhibited promoter specificity. This disproportionate promoter enrichment provides direct evidence of intensified purifying selection targeting regulatory TFBS within pG4 regions to control gene regulation. (Table S22).

Finally, since GC bias in TFBS likely arises from suppressed meC-to-T deamination (and the corresponding G-to-A transitions on the opposite strand), the primary mechanism driving GC loss in vertebrates, we specifically examined position weight matrices (PWMs) of differentially enriched/depleted motifs to detect these direct mutation signatures (Additional file 2). We generated sequence logos for all significantly enriched or depleted motifs (homeotherm vs poikilotherm), comparing those in pG4 regions vs pG4-free backgrounds (both genome-wide and promoters), alongside JASPAR reference logos (Fig. S20-21, Table S23). We observed clear suppression of mutation rates for the differentially enriched and significantly GC-retentive motifs (Fig. S20-21). The visible position-specific C-to-T transition signatures in PWM logos confirm that the reduced cytosine deamination drives this homeotherm-specific evolutionary pattern (Fig. S20-21. This provides the final molecular evidence that homeotherms are actively preventing these mutations at different rates across pG4 TFBS positions.

CGGBP1 emerges as a key contributor to this process, restricting meC-to-T mutations at GC-rich TFBS in pG4 contexts as shown by our ortholog expression data (Fig. 4, Figs. S15-16 and S18). However, CGGBP1 is not acting alone; these findings reveal a broader molecular machinery that maintains GC-rich regions genome-wide, including TFBS in pG4s, ultimately sustaining vertebrate G4 potential through coordinated mutation suppression. We propose a co-evolutionary mechanism linking CGGBP1 function, heterogeneous C-to-T transition rates, and G4-forming potential across vertebrate genomes (Fig. 6). This represents an evolutionary innovation of CGGBP1: since G4 formation emerges as a chemical byproduct of GC-rich DNA sequences, CGGBP1 has adapted to selectively restrict meC-to-T mutations, thereby preserving genome-wide GC content and sustaining G4 landscapes where these structures serve regulatory roles. This framework provides a novel evolutionary perspective on CGGBP1’s cellular function, revealing its role as a mutation suppressor tuned to vertebrate genomic architecture.

## Discussion

CGGBP1 is reported or predicted in all terrestrial vertebrate classes, with indications of gene-loss in most amphibian groups and retention only in amniotes (Singh and Westermark 2015; Morbia et al. 2024; Kumar et al. 2025). Within the amniotes, the evolutionary conservation pattern of CGGBP1 follows the endo-homeothermic versus poikilothermic divide. The G4-forming potential, its GC-richness, and GC-skew of G4-forming regions also follow the same pattern. The best bifurcation of the G4-forming potential genome-wide is achieved only when we cluster the 105 species through a simple homeothermic-poikilothermic divide, not otherwise. These findings establish that with the evolution of the terrestrial vertebrates, the epigenome (in the form of cytosine methylation landscape) has affected the genome, its GC-skew, and G4-forming potential under the influence of CGGBP1. This enrichment of G4-forming potential in homeotherms coincides with a G4-preventing function of CGGBP1. The increase in G4-forming potential has been under an evolutionary check by the same agent that facilitates it, the homeothermic CGGBP1, best represented by its highly conserved mammalian form.

Our analysis establishes that the G4 GC-contents have increased and correlated with genomic GC-contents during vertebrate evolution. The evolution of CGGBP1 and its C-retentive effects appears to be causal to this phenomenon. Interestingly, of all the GC-relevant point mutations analyzed, all C-retentive (secondarily G-retentive) point mutations are antagonized by CGGBP1. This function of CGGBP1 ascends from non-amniotes to mammals, mimicking the GC-content increase in G4s as shown in Figure 1. A direct test of the potentially causal role of CGGBP1 in G4 GC-retention is to test C-retention rates in actual G4-formed regions *in cellulo* for which a regulatory role of CGGBP1 is established. We discover that the acute expression of different CGGBP1 forms inside HEK293T cells affects C-retention differently very strongly at CGGBP1-mitigated G4s. When compared against the G4-forming potential of the genomes, not under chromatin constraints, we observed a distinction between G4 structures in actuality, rather than the potential of DNA sequences to form G4 reflected as DNA sequence properties. Specific meC-to-T transition restriction by CGGBP1 is observed only at regions that can overcome the constraints on G4 formation by double-strandedness. These regions have strong G4 folding properties such that, given a loss of chromatin constraints, *in vitro,* they would form G4. In contrast, the regions with potential for G4 formation, which can be realized only when the double-strandedness is physically overcome, are not targets for meC-to-T restriction by CGGBP1. *In vivo*, the G4-forming potential realization into actual G4s is impeded not only by double double-strandedness of the DNA but also by chromatin constraints. Nucleosomes, DNA-binding proteins, and epigenetic features render the G4-forming potential into actual G4 folding depending on a combination of the following: innate G4-forming potential, default state of double-strandedness, chromatin constraints, transitory single-stranded state due to processes like replication, transcription, repair, and recombination (Eddy et al. 2011; Maizels 2012; Johnson 2020). Hs CGGBP1 has been shown to be required for exerting the chromatin constraints on G4 formation (Datta et al. 2024). *In vitro*, recombinant CGGBP1 can impede G4 folding of ssDNA by physically associating with it through mechanisms that are not completely understood. CGGBP1 could be collaborating with additional factors in mitigating G4 folding; however, it remains clear that just by itself, CGGBP1 can interfere with G4 formation (Ummethum et al. 2025; Datta et al. 2024). The current work sheds light on the biochemical nature of this function of CGGBP1 and the effect of its evolution on the G4 landscape of higher vertebrates.

It is interesting to note that through the evolution of higher vertebrates, there has been no selection against G4-forming potential. To the contrary, as our results demonstrate, genomic features that favour G4 formation seem to have become enriched. One such primary feature, apart from GC-richness, is that of inter-strand GC compositional asymmetry. Without invoking the role of CGGBP1 in G4 biology, it remains unclear what the true evolutionary force underlies this G4-forming potential enrichment in higher vertebrates. It could either be the actual G4 formation, or perhaps the GC-skew offers other functional roles, such as TFBSs and DNA biophysical properties compatible with replication and transcription bubble formation. The latter would imply that G4 folding of dsDNA is an unavoidable outcome of the DNA composition actually selected for functions requiring DNA structural turnovers. The absence of any known G4-stabilizing proteins or domains and an abundance of helicases and G4-resolving proteins lend support to this possibility. The role of CGGBP1 provides an additional layer of regulation on the G4-forming potential: it interferes with the G4 folding and, at the same time, protects and helps retain G4-forming potential. Interestingly, the mechanism underlying this dual effect of CGGBP1 on G4-forming regions also favours GC-retention at TFBSs in target gene promoters (Kumar et al. 2025). Biochemically, CGGBP1 interferes with cytosine methylation, restricts meC-to-T transitions, and thereby affects GC-contents locally (Agarwal et al. 2015; Singh and Westermark 2015; Morbia et al. 2024; Kumar et al. 2025). In terms of G4-forming potential retention and regions with high GC-skew, this attributes a unique function to CGGBP1: It is not the G-rich G4-forming strand, rather the C-rich complementary strand, which is under the influence of cytosine methylation and a higher load of meC-to-T transitions. A primary loss of C on the C-rich strand would secondarily lead to G-loss on the G4-forming strand. Such a mechanism of G4 regulation is not reported and is unique to CGGBP1. Through cytosine methylation restriction, CGGBP1 ensures that C and eventually G are retained, which is reflected as G4-forming potential. Since this effect of CGGBP1 is directly proportional to the GC-content of the genomes, one would expect that the increasing correlations between GC-contents of pG4s and respective genomes from non-amniotes and lower amniotes to the higher amniotes are due to a parallel evolution of CGGBP1. Our findings here show that representative forms of CGGBP1 from these groups mimic the biochemical effects of CGGBP1 on meC-to-T transitions in acute overexpression experiments. Remarkably, these acute biochemical effects of CGGBP1 on the cytosine methylome reflect the evolutionary changes in genomic GC-content and G4-forming potential associated sequence features.

Together, these findings show that CGGBP1 mediates the evolution of G4-forming potential of the genomes through two mechanisms. One is by exerting a chromatin constraint and disallowing G4 folding, and the second, by minimizing the meC-to-T loss on the C-rich non-G4 strand through a cytosine methylation restriction. Apparently, this function of CGGBP1 is enriched in vertebrate evolution and observed as concentrated only at regions with a strong tendency to form G4s, not at all locations genome-wide, with a varying range of G4-forming potentials. The emergence of higher vertebrates, which are homeothermic, coincides with genomic GC-richness, favouring TFBS richness against a G4-forming potential which can affect TF-binding and genome function. The evolutionary pressures on establishing a balance between these two opposing features apparently operate not through regulation of G-richness, but rather through maintenance of the complementary C-richness. The evolution of CGGBP1 has played an important role in reconciling non-CpG cytosine methylation, GC-skew, GC-richness, and G4-forming potential with functional outcomes of the genome compatible with the emergence of higher amniotes and homeothermy.

## Conclusions

- Our findings position CGGBP1 as a key evolutionary safeguard of GCJrich, methylationJvulnerable regions, where it limits meC-to-T transitions and stabilizes GC content at promoters, TFBSs, and G4Jforming elements, particularly in homeotherms.
- By integrating vertebrate comparative genomics with baseJlevel mutation profiling, we show that CGGBP1 activity relates with G4-forming potential and GC stability, with human CGGBP1 displaying a uniquely strong capacity to suppress methylationJdriven GC loss at structurally complex regulatory loci.
- Overall, this work supports a model in which CGGBP1 protects G4Jprone, GCJdense regulatory DNA from methylationJassociated mutagenesis, thereby contributing to the maintenance of genomic integrity and regulatory complexity in higher vertebrates.

## Limitations of the study

1. A direct cause-effect relationship between G4 formation and evolutionary forms of CGGBP1 needs targeted experiments.
2. The time-series experiments, through which cytosine methylation and meC-to-T transition restrictions effects of various CGGBP1 forms at high G4-forming potential regions have been deduced, shall be extended to *in-cellulo* G4 landscape characterization experimentally.

## Methods

### 1. Mapping the G4 landscape in vertebrate genomes and their 1 kb cis promoters via pqsfinder

To assess G4-forming potential across vertebrate genomes, we analyzed 105 reference genomes, including 57 mammals, 27 birds, 17 reptiles, and 4 non-amniotes, sourced from Ensembl as more details in Supplementary Table S17 (Kumar et al. 2025). Genome FASTA files were systematically processed using pqsfinder, a G4 prediction tool uniquely capable of identifying not only canonical G-quadruplex-forming sequences but also those with imperfections, such as bulges, mismatches, and atypically long loops (Hon et al. 2017; Labudová et al. 2020). In contrast to previous algorithms that enforce the strict [G3N1-7]4 motif, pqsfinder accommodates the greater sequence diversity observed in experimentally validated G4 structures, including longer loops and non-standard spacers. Its high computational efficiency and parameterization using high-throughput G4-seq experimental datasets make it particularly suited for large-scale genomic analysis. We applied a recommended score threshold of 52 to prioritize reliability and specificity, effectively reducing false positives and aligning predictions with experimentally validated G4 motifs (Hon et al. 2017; Labudová et al. 2020). We applied the same analysis to all annotated promoters across vertebrate genomes. We supplied 1kb *cis* promoter sequence FASTA files for each species, which were retrieved using coordinates previously extracted from genome transfer format (GTF) files, as described elsewhere (Kumar et al. 2025).

### 2. Genome-wide sequence feature characterization of predicted G-quadruplex potential across vertebrate genomes

In addition to obtaining the raw output generated by pqsfinder, which includes coordinates, sequences, and scores of potential G-quadruplex (G4)-forming regions, we developed a customized pipeline to calculate comprehensive summary statistics and associated features for both the predicted G4-forming sequences and their respective input sequences. These metrics encompass genome size or promoter region size (defined as the total nucleotide length of all annotated promoters combined), GC-content of the entire genome and promoter regions, the total number of predicted G4 motifs within genomes and promoters, and the proportion of the genome or promoter sequences covered by these G4-forming regions. Furthermore, we computed the density of G4 motifs per megabase, average G4 motif length, average G4 prediction scores, average GC-content of the predicted G4 sequences, and the number of chromosomes or scaffolds per species. This statistical characterization was carried out both genome-wide and across all annotated promoters for each species analyzed. These summary statistics formed the basis for downstream analyses and enabled us to draw biologically meaningful conclusions supported by statistical significance testing (details in Supplementary Table S1 and Table S5). All these features for a particular species represent the average value of all potential G4-forming sequences reported by the tool pqsfinder within the genomes or promoters analysed.

### 3. Phylogenetic comparative analysis of pG4 distribution in vertebrate genomes

To examine how putative G-quadruplex (pG4) landscapes varies across the vertebrate phylogeny, we performed a comparative analysis using two distinct genomic contexts: Whole-Genome (Genomes) and Promoters (1 kb *cis* upstream of TSS).

#### 3.1 Phylogeny and data integration

We utilized a time-calibrated vertebrate species tree in Newick format (Species.nwk) generated from the TimeTree web tool from 105 species. To ensure data-tree consistency, we employed the *treedata* function from the *geiger* R package to prune the phylogeny and align the species metadata. We standardized names with underscores (like Homo_sapiens) so every species lined up correctly for downstream analysis.

#### 3.2 Characterization of phylogenetic signal

We calculated Pagel’s Lambda (λ) on all pG4 metrics (density, GC content, length), to determine the extent to which these pG4 features are influenced by shared ancestry, using the *phylosig* function in the *phytools* package. λ = 1 means traits evolve exactly following the phylogeny (Brownian motion), λ = 0 means no phylogenetic signal, species evolve independently. This told us how much shared ancestry shaped current pG4 patterns.

#### 3.3 Raw correlation (No Phylogeny Correction)

To see current patterns across living species, we calculated Spearman’s rank correlation (ρ) between background GC content and pG4 features (density, composition, length). We fit ordinary least squares regression lines with 95% confidence intervals, color-coding species by clade (mammals red, birds orange, reptiles green, etc.). This showed the baseline relationship without correcting for evolutionary history.

#### 3.4 Phylogenetic Independent Contrasts (PIC)

Since closely related species share evolutionary history and tend to cluster together, we used Phylogenetic Independent Contrasts (PIC) to account for this non-independence. We used Felsenstein’s PIC method (ape package) to transform all variables into (n-1) independent contrasts representing evolutionary divergences along each branch. We then ran origin-forced regression (through 0,0 coordinate) between PIC-transformed background GC vs pG4 features. The adjusted R² showed what fraction of evolutionary change in pG4 patterns was actually driven by background GC shifts, not just phylogenetic inheritance. This was performed independently for each dataset (Genomes vs. Promoters) to determine the baseline correlation between background nucleotide composition and pG4 density, composition, and length.

### 4. Mapping pG4 density across vertebrate genomic landscapes

#### 4.1 Extraction and categorization of genomic features

To map putative G-quadruplex (pG4) distribution variation across vertebrate lineages, we extracted coordinates for six specific genomic features from GTF annotation files for 105 vertebrate species. We specifically extracted coordinates for Transcription Start Sites (TSS), 5’ Untranslated Regions (5’UTR), Coding Sequences (CDS), 3’ Untranslated Regions (3’UTR), Introns, and entire Gene Bodies. TSS coordinates were defined as the start position of a transcript, with a surrounding window of ±2 kb. Intronic coordinates were derived by subtracting exon intervals from the total transcript.

#### 4.2 Metagene profiling and metrical scaling

To account for the varying lengths of genomic features across species, we implemented a metagene scaling approach using bedtools makewindows. Around each transcription start site (TSS), we analyzed the ±2 kb promoter region by dividing it into 50 bp bins, this gave us high-resolution signal patterns right at the core promoter and just upstream. For gene features like coding sequences (CDS), UTRs, and introns, we scaled each one into 100 proportional bins (0-100% of feature length). This ensures that pG4 density signals are comparable between a short 5’UTR and a long intronic region by representing the signal as a relative position within the feature (0% to 100%).

#### 4.3 GC-content normalization and signal quantification

pG4 density mapping was performed using bedtools intersect to count the number of predicted pG4 motifs within each specific genomic bin. For all gene-associated regions (TSS, UTRs, CDS, and Introns), G4 density was normalized relative to the background GC-content of the respective species’ genome. We utilized a normalization ratio derived from the global GC-content (extracted from species metadata) to calculate the Normalized pG4 Density:

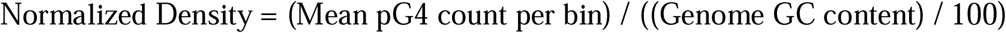

This correction ensures that the observed enrichment in promoters or UTRs is a biological signal rather than a mathematical artifact of high local GC-content. We also adjusted bin numbering based on strand orientation for each gene feature (CDS, UTRs, introns). For features on the negative strand, we automatically flipped the bin order so the biological 5’→3’ direction stayed consistent. This means Bin 1 always marks the true start of each feature, the start codon for CDS, 5’ end for UTRs, regardless of whether it’s on the + or - strand.

### 5. Characterization of point mutations within potential G4-forming sites in human genomic and promoter regions

Leveraging our recently published methylation-enriched DNA point mutation analysis pipeline (MAP-MAP) (see additional file 11 (Kumar et al. 2025)), we delved deep into the mutation landscape of potential G4-forming regions within the human genome. Our study focused on methylation-associated point mutation analysis in MeDIP DNA and the role of orthologous representative CGGBP1 forms in modulating the mutability of potential G4-forming sequences.We primarily aimed to test whether G4 motifs display elevated mutation rates by changing evolutionary variant of CGGBP1. We retrieved mutation profiles for predicted G4 regions across the genome, including entire promoters and potential G4 sequences therein. We also examined two experimental conditions, G4-IP-native (GSM6122749) and G4-IP-denatured (GSM6122750), that reflect G4 formation in the presence and absence of nuclear constraints, respectively, with the denatured condition representing DNA disrupted by heat treatment (Datta et al. 2024). Mutation patterns in these regions were thoroughly plotted and analyzed. Further, we concentrated on differentially captured regions (DCRs) in GSE202456 (Datta et al. 2024), which are genomic loci strongly influenced by CGGBP1 levels and possess potential G4-forming capability. For these DCRs, we calculated mutation profiles with a specific focus on methylation-dependent C-to-T transitions and assessed how different CGGBP1 forms impact these mutational processes.

### 6. Point mutation rate calculation in GC-rich TFBS genomewide

We first selected all GC-rich motifs (GC > 50%) from the JASPAR vertebrate database and scanned entire genomes for these motifs using FIMO. We then aligned this motif mapping with MeDIP sequencing data from cells overexpressing different evolutionary variants of CGGBP proteins. For each motif hit, we ran MAP-MAP analysis on those exact coordinates to count every mutation type, both transitions (like meC-to-T) and transversions, across all samples (Table S15). Our main focus was meC-to-T mutation rates specifically within these TFBS. We calculated mutation rates as the fraction of each mutation type per sample, then computed the mean rate across all samples for each motif, and subtracted this mean from individual sample rates to reveal variant-specific effects. Finally, we generated heatmaps with unsupervised clustering to visualize how CGGBP variants alter mutation patterns across GC-rich motifs.

### 7. Mapping TFBS density in pG4 regions of vertebrate genomes

#### 7.1 Scanning motifs across genomes and promoters

We started by scanning 105 vertebrate genomes and their 1 kb promoter regions for TFBS using FIMO (default parameters) against JASPAR vertebrate motifs. As a control, we randomly sampled exactly matching non-pG4 regions from each genome, same number and length as the pG4 regions, and ran FIMO on those too. For promoters, we extracted pG4-free promoter sequences by removing pG4 coordinates from the full 1 kb regions, then counted TFBS in both pG4 and pG4-free promoters. To compare across species with different genome sizes, we normalized all counts to TFBS per Mb using the total base pairs searched in each region (pG4 vs background) from our Promoter Search Space Summary. which represents the total physical base pairs of the pG4-containing and equivalent pG4-free regions used during the FIMO search in promoter as well as genomewide. This gave us clean TFBS densities for pG4 regions (both genome-wide and promoters) versus their matched backgrounds.

#### 7.2 Thermal clade enrichment analysis

Next, we grouped species into homeotherms (57 mammals + 27 birds) vs poikilotherms (17 reptiles + 4 non-amniotes) and calculated Δ Density (pG4 minus background TFBS density) for each motif. We computed mean retention for each motif within thermal groups, then used Wilcoxon Rank Sum tests (pseudocount 0.1 for stable log2FC) to evaluate the significance of differences between, homeotherm vs poikilotherm. To account for multiple testing across the JASPAR vertebrate motif library, raw P-values were adjusted using the Benjamini-Hochberg (BH) False Discovery Rate (FDR) correction. Motifs were subsequently classified into discrete enrichment tiers based on an FDR-adjusted P-value threshold of 0.05 and effect size: “Highly Enriched” or “Highly Reduced” (log2FC > 1.0), and “Moderately Enriched” or “Moderately Reduced” (0.58 < log2FC ≤ 1.0). Further, these results were visualized through dual-panel volcano plots, where statistical significance (-log_10_ P_adj_) was mapped against the magnitude of cladal divergence, using this strategy we were able to isolate the “Global Core” motifs that exhibit superior preservation within homeothermic and poikilothermic pG4 structures.

#### 7.3 Extracting motif sequences for GC content analysis

Next, to investigate if the differentially enriched motif is associated with the compositional difference in the GC content of the motifs, we implemented a targeted sequence extraction pipeline. Using the significantly differentially enriched above (highly/moderately enriched or reduced), we extracted raw nucleotide sequences (the actual string matched string reported by FIMO) from the FIMO output files. Which were then organized into a hierarchical FASTA database structured by thermal group (Homeotherm vs. Poikilotherm), genomic context (pG4-containing vs. pG4-free), and enrichment status. For each motif, we calculated the mean GC content separately for homeotherm vs poikilotherm species, breaking it down by genomic context (pG4 regions in both genome-wide and promoters) versus their matched pG4-free backgrounds. We then summarized the mean GC% and total hit counts across all these categories. This let us test whether homeotherm pG4 motifs showed specific GC retention beyond general genomic trends.

#### 7.4 Comparison of GC content of motifs across thermal groups to assess GC retention bias

To quantify the relationship between significantly differentially enriched motifs in promoteers and genome has any association with GC retention, we conducted a comparative statistical analysis of GC-content across homeothermic and poikilothermic lineages. We utilized two different approach to analyse it: first assessing Absolute GC-content within pG4 and pG4-free regions, and second by calculating Δ GC-content, defined as the mean GC-fraction of a motif in pG4 regions minus its mean in pG4-free background sequences. We ran paired Wilcoxon tests on absolute GC% (pG4 and pG4-free) and ΔGC (pG4 minus pG4-free) between homeotherms and poikilotherms. Furthermore, we applied linear regression analysis (R², β slopes) to observe the conservation of GC-content between the thermal groups and visualized them via faced scatter plots, where the observed GC-content of homeotherms was mapped against poikilotherms relative to a 1:1 parity line. This statistical analysis revealed whether differentially enriched motifs in homeotherms had elevated GC signatures specifically in pG4 contexts and deviate from the general genomic GC-drift. We also created detailed profiles at two levels: species-by-species and thermal group averages (homeotherm vs poikilotherm). Each profile compared promoter vs genome-wide pG4 motif hits alongside the average GC content for each motif, either across all species within that group or averaged across thermal clades. We annotated everything with JASPAR IDs and enrichment status (highly/moderately enriched or reduced), enabling direct side-by-side comparisons between homeotherms and poikilotherms. This captured both individual species differences and the broader thermal evolutionary trends in a single framework.

#### 7.5 Statistical validation of GC shifts in differentially enriched motifs

To validate the GC bias in motifs of pG4s between homeotherms and poikilotherms, we performed a chi-square test of proportions on the aggregated motif hit counts and their respective GC-fractions. We derived the thermal group statistics by integrating homeotherm and poikilotherm mean GC-content and total hit counts from both pG4 and pG4-free regions. We calculated Absolute GC Shift (homeotherm minus poikilotherm) and background-corrected shift (ΔGC difference). BH-adjusted P < 0.05 identified the top 50 motifs which shows highest GC retention from all differentially enriched motifs, by both metrics (absolutes and ΔGC difference), characterized them by genomic region (promoter or genome) along with thermal dominance (homeotherm or poikilotherm specific GC retention). Followed by we plotted these motifs using bar plots and labelled them enrichment category wise (promoter or genomewide), both for absolute and background-corrected.

#### 7.6 Functional enrichment analysis of top 50 GC retentive motifs in pG4s

To determine the biological significance of top GC retentive motifs found in pG4s of homeotherms, we performed a Gene Ontology (GO) functional enrichment analysis. We characterized top 50 motifs from absolute as well as background corrected motifs into three categories (Absolute_unique_, Normalized_unique_,and Common motifs), based on their commonality and uniqueness. Next, motif symbols were mapped to ENTREZ IDs using the org.Hs.eg.db database, and over-representation analysis for Molecular Functions (MF) was conducted using the clusterProfiler package. Significance was determined using a Benjamini-Hochberg adjusted P-value cutoff of 0.05. Results were visualized using dot plot. Dot plots showed fold enrichment vs significance of specific molecular function across these three categories of motifs.

#### 7.7 Generation of position weight matrices for differentially enriched motifs

To enable the thermal group specific comparison of motif composition and visualization of GC retention, we built thermal group-specific Position Weight Matrices (PWMs) in MEME format for each TFBS. We aggregated all extracted sequences for a given motif, classified by thermal group, genomic region, and enrichment status, and calculated the nucleotide distribution at each position. Also, to prevent zero-probability values, we applied a pseudocount before normalizing the raw counts into a Position Probability Matrix (PPM). This strategy was utilised for each motif found in pG4s of promoter as well as genome and their respective background in homeotherm and poikilotherms. This allowed us to quantify subtle sequence-level shifts between homeotherms and poikilotherms.

#### 7.7 Comparison of motif sequence logos for GC retention

Finally, we performed a structural comparative analysis of Position Weight Matrices (PWMs) to visualize how motif architecture varies between homeotherms and poikilotherms. For each motif, five distinct PWMs were compared: the JASPAR 2026 Reference, Homeotherm (pG4/Free), and Poikilotherm (pG4/Free). Using hierarchical clustering (Pearson correlation) to assess the similarity between these matrices, visualizing the relationships through integrated dendrogram-logo plots. We also calculated the location wise GC retention at each nucleotide position, identifying specific “hotspots” where the homeothermic pG4-motif significantly deviated (>10%) from the JASPAR reference. We also made sure that observed GC retention is not a product of random variation by performing proportion tests (P_adj_ < 0.05). This analysis provided the final evidence that in pG4s TFBS specific GC retention is actively preserved from GC loss via meC-to-T mutations in homeotherms as compared to poikilotherms.

### 8. Data visualization

To characterize the spatial distribution and statistical relationships of putative G-quadruplexes (pG4s), we visualized results using heatmaps, profile plots, and comparative distributions. Heatmaps were generated to visualize pG4 density across specific genomic intervals, using color intensity to represent signal enrichment across 100 scaled bins per feature. To complement these, profile plots were constructed by averaging the normalized pG4 density across all species for each bin, with shaded areas representing the standard error to illustrate taxonomic consistency. Inter-clade variations in absolute pG4 density across discrete genomic regions were assessed using boxplots, where the center line represents the median and whiskers extend to 1.5 times the interquartile range, allowing for the identification of significant outliers and taxonomic shifts. Finally, we generated correlation plots (scatterplots) to evaluate the coupling between pG4 features and background GC-content, incorporating both Ordinary Least Squares (OLS) and Phylogenetic Independent Contrast (PIC) regression lines to distinguish between contemporary patterns and evolutionary trajectories.

To visualize C-to-T transitions across the genome, and specifically within promoter regions and potential G4-forming regions of the human genome, we first produced normalized bigWig files for each MeDIP sample (Lc, Ac, Gg, Hs, Ev, and Input). These files represent the distribution of C-to-T transitions genome-wide, with each bigWig encoding the signal as the fraction of all transitions identified in each sample, normalized by subtracting the sample mean to account for global differences. These scaled bigWig files were subsequently used to plot C-to-T transition signals for each sample in 1kb windows upstream and downstream of predicted G4-forming regions in a bin size of 10 bp. Similarly, we plotted these scaled signals in relation to G4-IP-native (GSM6122749) and G4-IP-denatured (GSM6122750) samples, examining their distributions within 0.1Mb flanking intervals in a bin size of 1000 bp from each peak center.

To further investigate the distribution of potential G4 signals in regions devoid of mutations, we generated additional scaled bigWig tracks in 10bp bins for mutation-free sites genome-wide, using the predicted G4 bed coordinates for each sample. For inter-regional comparisons, the number of mutation-free regions per million bases was normalized for each flank by dividing each value by the highest observed value in the dataset (after subtracting the Input signal), resulting in relative values ranging from 0 (minimum) to 1 (maximum). In parallel, we created normalized bigWig tracks for GC-skew (calculated as the modulus of GC skew) in 25bp bins genome-wide. We plotted these signals in relation to mutation-free regions for each sample. The GC-skew signal was also assessed for enrichment by plotting the 25bp bin GC-skew modulus bigWig track in 0.1Mb flanking intervals in a bin size of 1000 bp around G4-IP-native and G4-IP-denatured peak centers. Finally, to study sequence features linked to potential pG4-forming regions identified by pqsfinder, we plotted GC-skew signals in 1kb intervals both upstream and downstream of these coordinates. As a control, GC-skew signals were also plotted for pG4-free regions, defined as genome-wide regions where potential G4-forming motifs were absent.

#### 9. Statistical analysis

We ran all analyses in R, building a pipeline that handled both species differences and evolutionary relationships. To compare pG4 density across vertebrate groups without assuming normal data distribution, we used Kruskal-Wallis tests (non-parametric ANOVA alternative) followed by Dunn’s post-hoc tests for specific pairwise comparisons. Since we made many taxonomic comparisons, we applied Benjamini-Hochberg correction to control false discoveries and ensure our P-values were reliable. For metagene profile plots, we applied loess smoothing to capture non-linear signal patterns, adding standard error ribbons to show how consistent pG4 enrichment peaks were across species. To measure pG4-genomic coupling strength, we calculated Adjusted R² and Spearman’s rank correlation (ρ). Finally, Phylogenetic Independent Contrasts (PIC) regressions confirmed these correlations reflected real evolutionary signals, not just shared ancestry artifacts

The proportions of transition and transversion mutations, calculated as fractions of total point mutation entries in each MeDIP sample within potential G4-forming regions, were found to be highly significant (P-value < 0.001) based on chi-square tests. In parallel, for each sample, methylated C-to-T (meC-to-T) transitions, expressed as a fraction of total transitions, were also highly significant (P-value < 0.001) upon application of the same statistical test. These statistical comparisons were performed across three genomic contexts within the human genome: potential G4-forming regions genome-wide, predicted G4 motifs located within 1kb promoter regions, and the full 1kb promoters themselves. This was possible because MeDIP data corresponding to various CGGBP1 forms were specifically available for the human genome.

To assess C-to-T transition rates genome-wide within potential G4-forming regions, we calculated the proportion of C-to-T transitions among all transition events for each sample. We subtracted the average value, representing deviations from the mean. To illustrate differences in point mutation rates across samples, each point mutation type was expressed as a percentage of total point mutations within the respective sample; this percentage was then normalized by subtracting the corresponding value obtained from the Input sample. This approach allowed a clear demonstration of methylation-associated point mutation rates in potential G4-forming regions located within 1kb annotated promoter regions of the human genome. Finally, analysis revealed that the percentage of point mutations occurring in potential G4-forming regions within 1kb promoter intervals, relative to the total point mutation count per sample, varied significantly among samples (P-value < 0.001, chi-square test), further supporting the specificity and reproducibility of these findings.

### 10. Software

All analyses were implemented using bash, Python, and R scripts (available in Code Availability). The pipeline was implemented in R (version 4.3.2). Putative G-quadruplexes (PQS) were predicted across 105 genomes using pqsfinder (version 2.0) with a stringent min_score = 52. We used Biostrings v2.70 and stringr v1.5 for sequence/GC calculations, BEDTools v2.31 and deepTools v3.5 for coordinates, ape v5.7/geiger v2.0 with TimeTree v5.0 for phylogenetics, and tidyverse/patchwork for visualization. TFBS discovery used FIMO (MEME Suite) against JASPAR 2026 motifs. clusterProfiler (org.Hs.eg.db) handled GO enrichment, Biopython extracted sequences, and universalmotif/ggseqlogo generated clade-specific PWMs with Pearson clustering for logo comparisons.

## Code availability

Custom scripts and code utilized in this study are publicly accessible at the following GitHub repository: https://github.com/umash-singh/MAP-MAP-analysis.

## Author contributions

PK performed all the analyses, generated the data, and participated in manuscript writing. US provided supervision and participated in writing the manuscript.

## Statements and Declarations

The authors declare no conflict of interest.

## Acknowledgments

This work was carried out with funds from ANRF (erstwhile SERB) [CRG/2021/000375] to US with additional support from IIT Gandhinagar. PK was supported by the MHRD fellowship.

## Additional information

This manuscript includes 6 main figures (Fig. 1-6 as PNG files), 21 supplementary figures (Fig. S1-S19 as PNG files; Fig. S20-S21 as PDF files), 23 supplementary tables (Table S1-S23 in Excel format), and two additional files: Additional file 1 contains the phylogenetic tree of all 105 vertebrate species analyzed in NEWICK format, while Additional file 2 is a zipped archive of motif PWMs in MEME format.

## Supplementary Figure legends

Figure S1. Phylogenetic relationship among 105 vertebrate species. Unrooted phylogenetic tree showing relationships among 57 mammals, 27 aves, 17 reptiles, and 4 non-amniotes, generated using <u>TimeTree.org</u> from our species list. The lower panel displays the geological timescale with key attributes evolving across phylogenetic branches.

Figure S2. GC-normalized mean pG4 density flanking TSSs across vertebrate clades. Boxplot showing mean normalized pG4 density (±2 kb flanking TSSs) for mammals (red), aves (blue), reptiles (green), and non-amniotes (purple). Aves exhibit the highest density, followed by mammals, with reptiles intermediate and non-amniotes lowest (P < 0.05 vs mammals, Wilcoxon rank-sum test). Asterisks indicate statistical significance. Data derived from 50-bp bins normalized to species-specific GC content (see methods, Table S4).

Figure S3. Clade-specific pG4 density variation surrounding TSSs (±2 kb). Heatmap of mean normalized pG4 density in 50-bp bins across individual vertebrate species, with clades clustered separately (mammals, aves, reptiles, non-amniotes). Clade colors: mammals (red), aves (blue), reptiles (green), non-amniotes (purple). The upper plot shows the mean profile across all vertebrates ± SE. The dashed blue line represents TSS. Continuous bin numbers are marked below the plot (1-80). Color scale transitions from blue (low) → yellow (moderate) → red (high) GC-normalized pG4 density (see methods, Tables S3-S4).

Figure S4. GC normalized pG4 density across transcript regions (UTRs and exons). Boxplots show mean normalized pG4 density in 5’ UTRs, 3’ UTRs, and exons across mammals (red), aves (blue), reptiles (green), and non-amniotes (purple). 5’ UTRs exhibit amniote-specific conservation with no significant differences among mammals, aves, and reptiles (P > 0.05), but reduced density in non-amniotes (P < 0.05 vs mammals, Wilcoxon rank-sum test). Exons show homeotherm-specific enrichment (mammals = aves) with progressive decline through reptiles to non-amniotes (P < 0.05 vs mammals). 3’ UTRs display a clear evolutionary gradient: mammals > aves > reptiles > non-amniotes (P < 0.05 vs mammals). Asterisks indicate statistical significance. Data from scaled binning with species-specific GC normalization (see methods, Table S4).

Figure S5. Clade-specific pG4 density variation across 5’ UTRs. Heatmap of mean normalized pG4 density in scaled bins across individual vertebrate species, with clades clustered separately (mammals, aves, reptiles, non-amniotes). Clade colors: mammals (red), aves (blue), reptiles (green), non-amniotes (purple). The upper plot shows the mean profile across all vertebrates ± SE. 5’ UTR start/end are marked below the plot; bin numbers (1-100) are shown beneath. Color scale: blue (low) → yellow (moderate) → red (high) GC-normalized pG4 density (see methods, Table S3).

Figure S6. Clade-specific pG4 density variation across 3’ UTRs. Heatmap of mean normalized pG4 density in scaled bins across individual vertebrate species, with clades clustered separately (mammals, aves, reptiles, non-amniotes). Clade colors: mammals (red), aves (blue), reptiles (green), non-amniotes (purple). The upper plot shows the mean profile across all vertebrates ± SE. 3’ UTR start/end are marked below the plot; bin numbers (1-100) are shown beneath. Color scale: blue (low) → yellow (moderate) → red (high) GC-normalized pG4 density (see methods, Table S3).

Figure S7. GC normalized pG4 density across introns. Boxplot showing mean normalized pG4 density in introns for mammals (red), aves (blue), reptiles (green), and non-amniotes (purple). Mammals exhibit the highest density, followed by reptiles, then non-amniotes, with aves lowest (P < 0.05 vs mammals, Wilcoxon rank-sum test). Asterisks indicate statistical significance. Data from scaled binning with species-specific GC normalization (see methods, Table S4).

Figure S8. GC normalized pG4 density across the gene body. Boxplot showing mean normalized pG4 density in introns for mammals (red), aves (blue), reptiles (green), and non-amniotes (purple). Mammals, reptiles, and non-amniotes show no significant differences (P > 0.05), while aves exhibit significantly lower density (P < 0.05 vs mammals, Wilcoxon rank-sum test). Asterisks indicate statistical significance. Data from scaled binning with species-specific GC normalization (see methods, Table S4).

Figure S9. Clade-specific pG4 density variation across exons. Heatmap of mean normalized pG4 density in scaled bins across individual vertebrate species, with clades clustered separately (mammals, aves, reptiles, non-amniotes). Clade colors: mammals (red), aves (blue), reptiles (green), non-amniotes (purple). The upper plot shows the mean profile across all vertebrates ± SE. Exon start/end are marked below the plot; bin numbers (1-100) are shown beneath. Color scale: blue (low) → yellow (moderate) → red (high) GC-normalized pG4 density (see methods, Table S3).

Figure S10. Clade-specific pG4 density variation across introns. Heatmap of mean normalized pG4 density in scaled bins across individual vertebrate species, with clades clustered separately (mammals, aves, reptiles, non-amniotes). Clade colors: mammals (red), aves (blue), reptiles (green), non-amniotes (purple). The upper plot shows the mean profile across all vertebrates ± SE. Intron start/end are marked below the plot; bin numbers (1-100) are shown beneath. Color scale: blue (low) → yellow (moderate) → red (high) GC-normalized pG4 density (see methods, Table S3).

Figure S11. Clade-specific pG4 density variation across the gene body. Heatmap of mean normalized pG4 density in scaled bins across individual vertebrate species, with clades clustered separately (mammals, aves, reptiles, non-amniotes). Clade colors: mammals (red), aves (blue), reptiles (green), non-amniotes (purple). The upper plot shows the mean profile across all vertebrates ± SE. The gene body start/end are marked below the plot; bin numbers (1-100) are shown beneath. Color scale: blue (low) → yellow (moderate) → red (high) GC-normalized pG4 density (see methods, Table S3).

Figure S12. Genome-wide point mutation rates in potential pG4-forming regions show evolution-dependent enhancement favoring GC retention by CGGBP1. Point mutation rates were determined as the percentage of each mutation type out of the total mutations for each sample, adjusted by subtracting the input control background. Different mutation types are illustrated as bar graphs. The analysis reveals an increased mutation rate in Hs, reflecting a genome-wide bias toward GC-content preservation in potential pG4-forming regions. Bars are color-coded by sample: Lc (blue), Ac (green), Gg (red), Hs (yellow), and Ev (grey). All mutation type proportions relative to total mutations are statistically significant (Chi-square test, P < 0.01; Table S12).

Figure S13. Potential pG4-forming regions are distinguished by their notable GC asymmetry. (a): pG4-free regions lack central enrichment of GC-skew within their ±1 kb flanks. (b): Predicted G4s in promoters exhibit pronounced central GC-skew enrichment. (c): Potential G4-forming regions show strong G/C asymmetry enrichment. GC-skew signals were plotted across ±1 kb flanks around the center of pG4-free regions, promoter-associated predicted G4s, and genome-wide potential G4-forming regions. Genome-wide potential G4-forming regions display higher GC-skew than those found specifically in promoters.

Figure S14. Human CGGBP1 maintains elevated G/C asymmetry and G4-forming potential in mutation-free regions. Scaled and normalized counts of potential G4-forming regions were plotted ±1 kb around one million randomly selected mutation-free regions across samples. Hs exhibits higher genome-wide G4-forming potential compared to Lc, Ac, Gg, and Ev. Similarly, GC-skew signals, also scaled and normalized by the number of mutation-free regions per sample, were plotted around the same set of regions. Regions with higher associated G4-forming potential display greater G/C asymmetry, with Hs maintaining significantly higher genome-wide G/C asymmetry than the other samples.

Figure S15. CGGBP1 drives evolutionary-dependent GC bias in point mutations within promoters and associated potential G4-forming regions. (a) & (b): Point mutation rates were calculated as the percentage of each mutation type relative to the total mutations within each sample, with background correction applied by subtracting the input control. Different mutation types are shown as bar graphs. Overall, the analysis reveals an elevated mutation rate in Hs, reflecting a bias toward preserving GC content in both potential G4-forming (b) regions and background promoters (a). Bars are color-coded by sample: Lc (blue), Ac (green), Gg (red), Hs (yellow), and Ev (grey). All mutation type proportions relative to total point mutations are statistically significant (Chi-square test, P < 0.01; Table S10 for promoters and Table S11 for potential G4s in promoters).

Figure S16. CGGBP1 maintains GC-content in DNA regions known to form G-quadruplexes by restricting methylation-associated C-to-T transitions. (a): Regions capable of forming G-quadruplexes within the chromatin-constrained nuclear environment (G4-IP-Nat; GSM6122749) show sensitivity to CGGBP1-dependent methylation-associated C-to-T transitions. Human CGGBP1 (Hs) strongly restricts these meC-to-T transitions, while empty vector control (Ev) and avian (Gg) forms exhibit slightly higher transition rates compared to input (background). Poikilothermic forms (Lc and Ac) display elevated meC-to-T transition rates in these verified G4-forming regions. (b): Potential G-quadruplex regions identified from heat-denatured genomic DNA, which lacks chromatin constraints and double-stranded structure (G4-IP-Denat; GSM6122750), do not associate with methylation patterns. No central enrichment or CGGBP1-dependent differences in meC-to-T transitions are observed in these regions. C-to-T transition signals, calculated as the fraction of all transition mutations with average background subtraction, are plotted ±0.1 Mb around peak centers for both G4-IP-Nat and G4-IP-Denat samples. (c): Differentially captured regions (DCRs), experimentally validated G4-forming loci highly sensitive to CGGBP1 levels (GSE202456), display species-specific meC-to-T transition patterns distinguishing poikilotherms (Lc, Ac) from homeotherms (Hs, Gg, Ev). Poikilotherms show poor restriction of meC-to-T transitions, whereas homeotherms, particularly Hs CGGBP1, strongly restrict these transitions. The empty vector control (Ev) represents endogenous CGGBP1 and displays intermediate restriction, with Gg showing moderate restriction relative to background input. Transition rates were calculated as the fraction of C-to-T mutations among all transitions, with averages subtracted per sample. Differences between groups are statistically significant (Chi-square test, P < 0.01; Table S14).

Figure S17. Chromatin-constrained G4-forming regions are defined by pronounced G/C asymmetry. G4-forming regions within the chromatin-constrained nuclear environment (G4-IP-Nat, GSM6122749) exhibit pronounced GC-skew enrichment, flanked by broader areas of low GC-skew. In contrast, G4-IP-Denat regions (GSM6122750) display only background genomic GC-skew levels without any central enrichment. These patterns are illustrated by heat maps generated from ±0.1 Mb flanks around the peak centers of each region.

Figure S18. Heatmap showing meC-to-T mutation rates across genome-wide GC-rich TFBS. Unsupervised hierarchical clustering clearly separates homeotherm (Gg, Hs) from poikilotherm (Lc, Ac) CGGBP orthologs, plus controls (Ev, Input). Human cells overexpressing these orthologs were MeDIP-sequenced and analyzed via MAP-MAP pipeline. Each row represents one TFBS (names on left), each column one sample. C-to-T rates were calculated as fraction of total transitions per motif per sample, then normalized by subtracting the mean rate for that motif across all samples. Blue-to-red scale shows low-to-high normalized mutation rates. Boxes below highlight homeotherms (red), poikilotherms (green), and controls (grey).

Figure S19. Absolute GC retention bias in TFBS in pG4s and their background. Scatter plots showing the correlation of mean absolute GC-content (%) for transcription factor binding sites (TFBS) between Homeotherms (y-axis) and Poikilotherms (x-axis). Genomewide (upper) and promoters (lower) are shown in different panels with motif enrichment status in homeotherms shown below with different colors highly enriched (Red), moderately enriched (light red), highly reduced (blue), and moderately reduced (light blue). (a): pG4-free background regions: Correlation of GC-content in TFBS located outside of predicted G4 regions. Slopes (β ∼ 1.0) and mean shifts (ranging from -0.22% to +1.97%) indicate baseline thermal group differences in genomic composition. (b): pG4 regions: Correlation of GC-content for TFBS overlapping with potential G4-forming sequences (pG4s). Homeotherms exhibit a systematic increase in absolute GC-content across almost all categories (e.g., +1.05% shift for highly enriched genomic motifs; P < 0.001, Paired Wilcoxon test). The dashed line represents the 1:1 parity line (y = x); the solid black line represents the linear regression fit. Statistical metrics including R-squared (R^2^), beta-slope (β), and P-values are indicated within each panel (Table S20).

Figure S20. Mutation protection hotspots in homeotherm pG4 TFBS across vertebrate genomes. Dendrogram sequence logo plots show sequence logos for 91 significantly enriched/reduced motifs (24 Highly Enriched, 39 Moderately Enriched, 10 Highly Reduced, 18 Moderately Reduced) from vertebrate genomes classified into thermal groups, comparing pG4 regions vs pG4-free backgrounds in homeotherms vs poikilotherms, clustered alongside JASPAR 2026 reference logos. Hierarchical clustering (Pearson correlation) groups similar motif architectures. Light red highlighting marks GC-loss hotspots where homeotherm motifs show C-enrichment (suppressed C-to-T) and G-enrichment (suppressed G-to-A), direct evidence of mutation protection at functionally critical positions.

Figure S21. Mutation protection hotspots in homeotherm pG4 TFBS across vertebrate promoters. Dendrogram sequence logo plots show sequence logos for 97 significantly enriched/reduced motifs (24 Highly Enriched, 36 Moderately Enriched, 18 Highly Reduced, 19 Moderately Reduced) in vertebrate promoters classified into thermal groups, comparing pG4 regions vs pG4-free backgrounds in homeotherms vs poikilotherms, clustered alongside JASPAR 2026 reference logos. Hierarchical clustering (Pearson correlation) groups similar motif architectures. Light red highlighting marks GC-loss hotspots where homeotherm motifs show C-enrichment (suppressed C-to-T) and G-enrichment (suppressed G-to-A), direct evidence of mutation protection at functionally critical positions.

## Supplementary Table legends

Table S1. Summary of G4-forming potential and associated features across 105 vertebrate genomes

Table S2. Genome-wide PIC and correlation statistics for pG4-associated features across vertebrate genomes

Table S3. GC-normalized pG4 density across functional regions in vertebrate genomes Table S4. Statistical metrics for pG4 density across vertebrate functional genomic regions

Table S5. Summary of G4-forming potential and associated features across 105 vertebrate 1 kb *cis* promoter

Table S6. Promoter PIC and correlation statistics for pG4-associated features across vertebrate genomes

Table S7. Summary of point mutation in potential G4-forming regions genome-wide in publicly available MeDIP dataset (GSE281704) after over-expression of evolutionary forms of CGGBP1

Table S8. Statistical summary of individual point mutations in potential G4-forming regions genomewide in publicly available MeDIP dataset (GSE281704) after over-expression of evolutionary forms of CGGBP1

Table S9. Summary of point mutation in pG4s in 1kb promoters and entire promoters in publicly available MeDIP dataset (GSE281704) after over-expression of evolutionary forms of CGGBP1

Table S10. Summary of point mutation in pG4s of 1 kb *cis* promoters in publicly available MeDIP dataset (GSE281704) after over-expression of evolutionary forms of CGGBP1

Table S11. Summary of point mutation in 1 kb *cis* promoters in publicly available MeDIP dataset (GSE281704) after over-expression of evolutionary forms of CGGBP1

Table S12. Summary and stats of C-to-T transition mutation in pG4s of 1kb promoters and entire promoters in publicly available MeDIP dataset (GSE281704) after over-expression of evolutionary forms of CGGBP1

Table S13. Statistical summary of C-to-T transition in publicly available MeDIP samples after over-expression of evolutionary forms of CGGBP1

Table S14. Summary and stats of C-to-T transition in Differentially Captured Regions (DCRs) obtained from publicly available dataset (GSE202456)

Table S15. Genomewide normalised point mutation rates in GC-rich TFBS in publicly available MeDIP dataset (GSE281704) after over-expression of evolutionary forms of CGGBP1

Table S16. TFBS raw counts in pG4 of promoters and genomes along with their respective pG4 free background across 105 vertebrates and summary stats of promoter pG4 and free regions

Table S17. TFBS density in pG4s of promoters and genomes along with their respective pG4-free backgrounds

Table S18. TFBS density in pG4s of promoters and genome after subtraction of pG4-free background in homeothermic vs poikilothermic comparison

Table S19. Detailed summary of mean GC content of each TFBS, species-wise and thermal group wise in pG4s of promoters as well as genomes along with their respective background

Table S20. Statistical summary differentially enriched TFBS showing absolute and background normalised GC-retention in homeotherm vs. poikilotherm pG4s

Table S21. Comprehensive detailed summary of differentially enriched TFBS GC-retention and overall top 50 GC-retentive motifs showing absolute and background normalised GC-retention in pG4s

Table S22. Functional enrichment analysis summary of top 50 GC-retentive motifs in pG4s overall in absolute and background normalised category

Table S23. Detailed summary of GC-retention in TFBS in and out of pG4s of promoters and genomes along with the enrichment status

## Abbreviations

Ac: Anolis carolinensis
CGGBP1: CGG triplet repeat-binding protein 1
DCRs: Differentially Captured Regions
Ev: Empty vector
G4s: G-quadruplexes
Gg: Gallus gallus
Hs: Homo sapiens
Lc: Latimeria chalumane
meC-to-T: methylated cytosine to thymine transition
MAP-MAP: Methylation-associated point mutation assessment pipeline
MeDIP: Methylated DNA Immuno-precipitation
pG4: Putative/Potential G4-forming regions
TFBSs: Transcription Factor Binding Sites
TSS: Transcription Start Sites
UTR: Untranslated Region
dsDNA: double-stranded DNA
ssDNA: single-stranded DNA

---

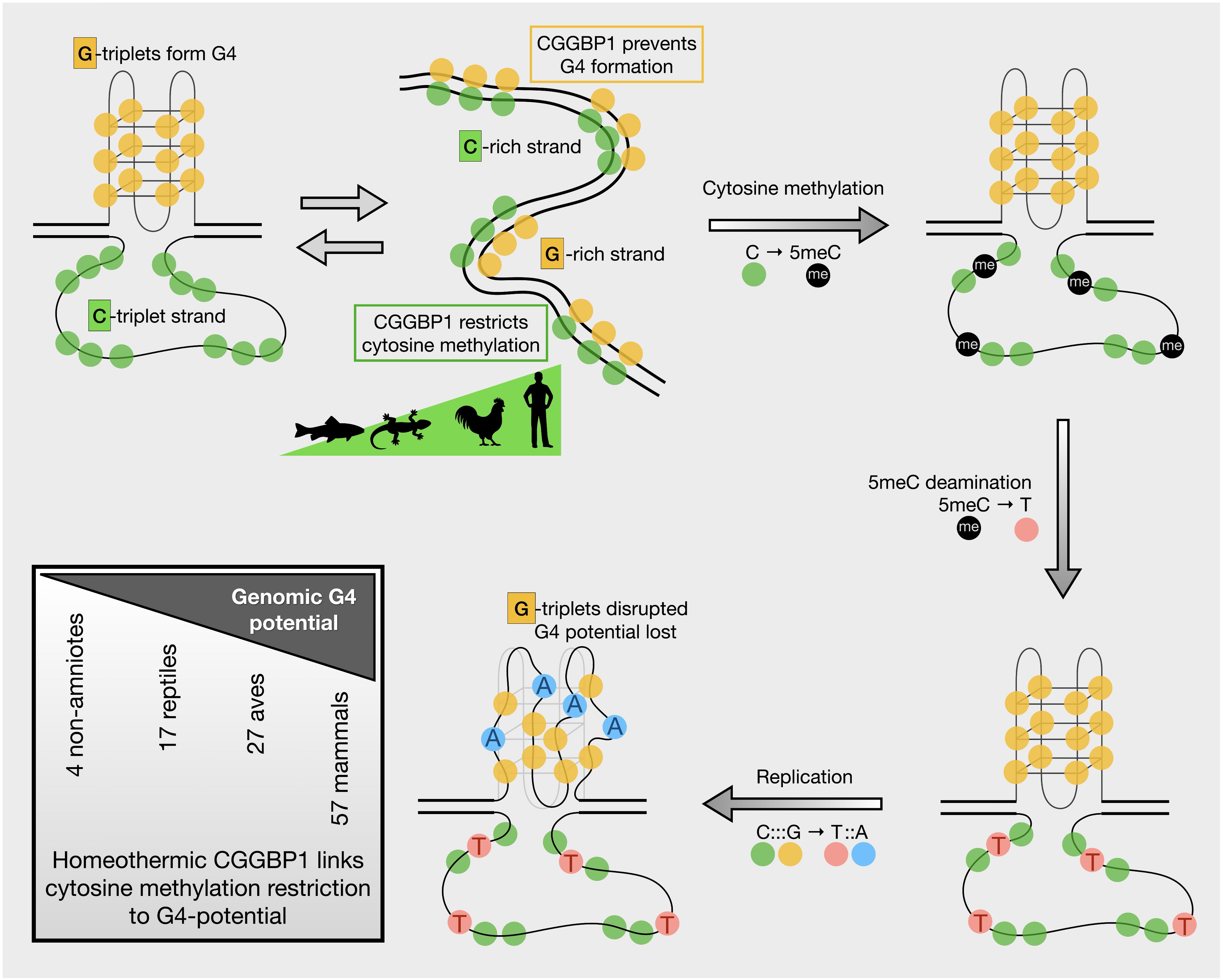

