## Supplementary figures and images for "CGGBP1-regulated heterogeneous C–T transition rates relate with G-quadruplex potential of terrestrial vertebrate genomes"

### Fig. S1

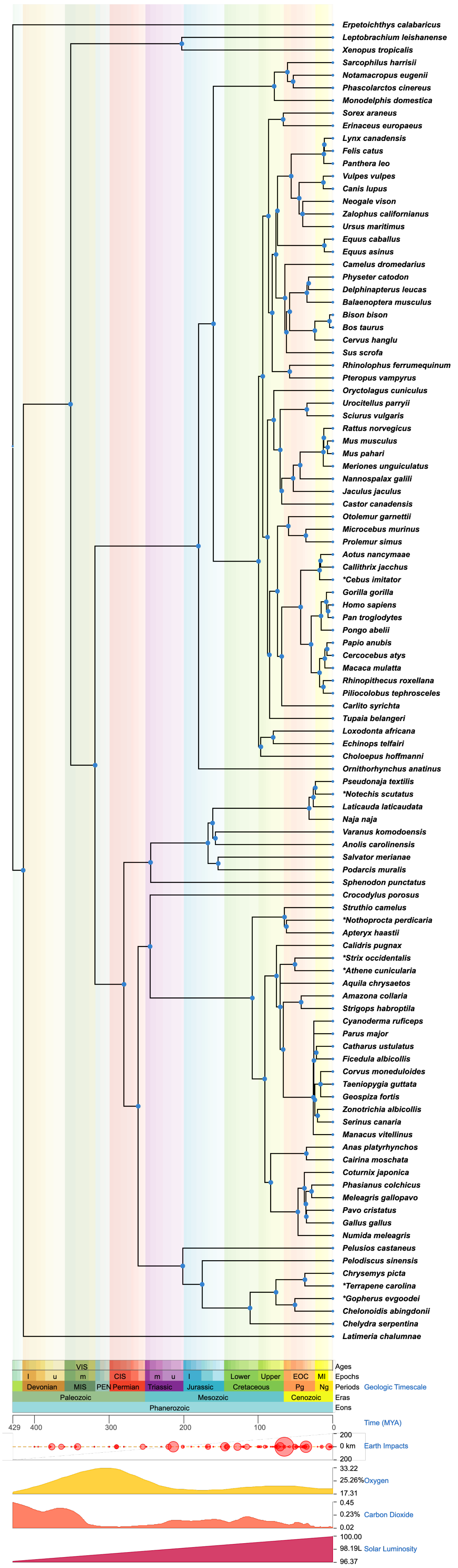

### Fig. S2

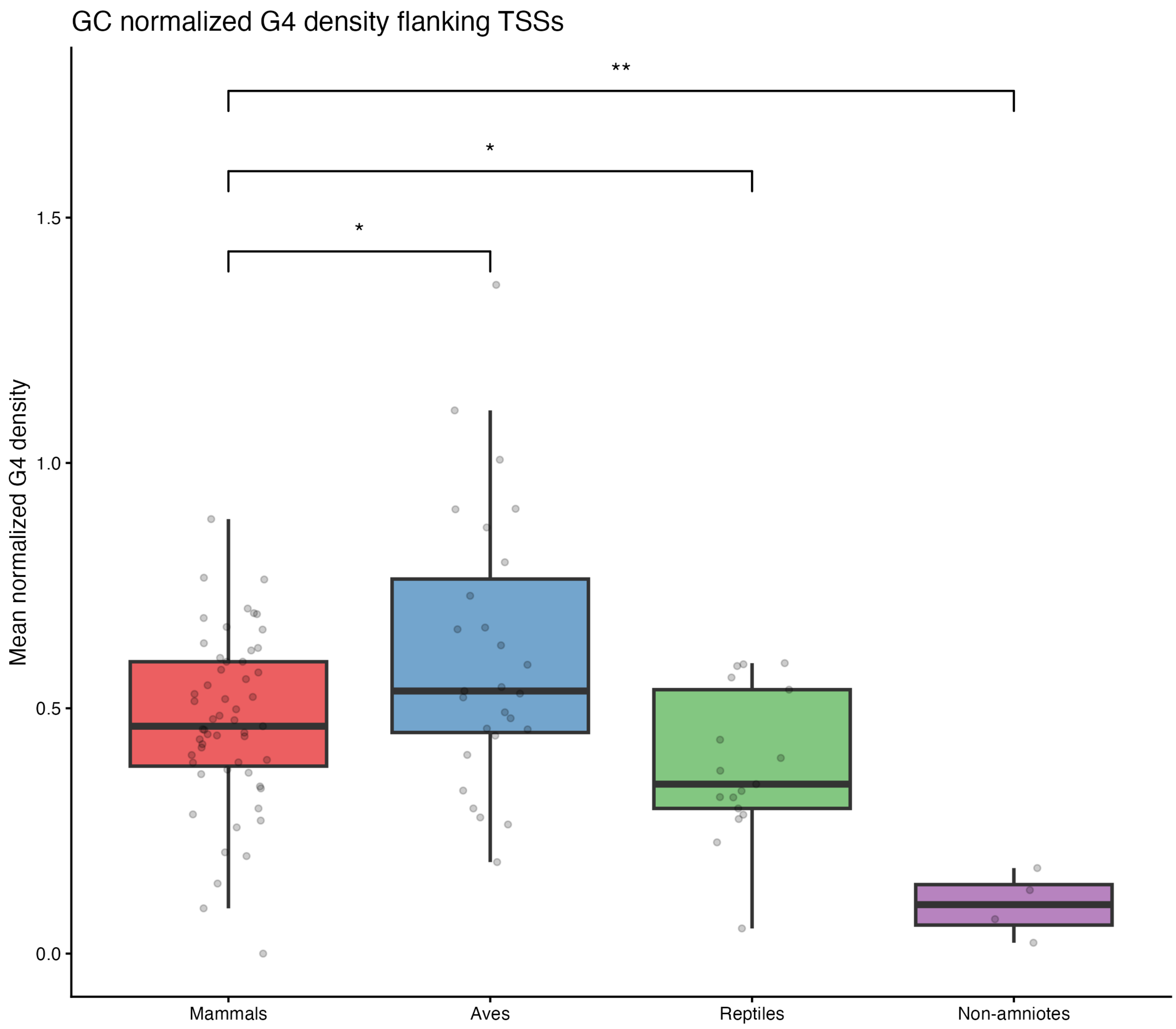

### Fig. S3

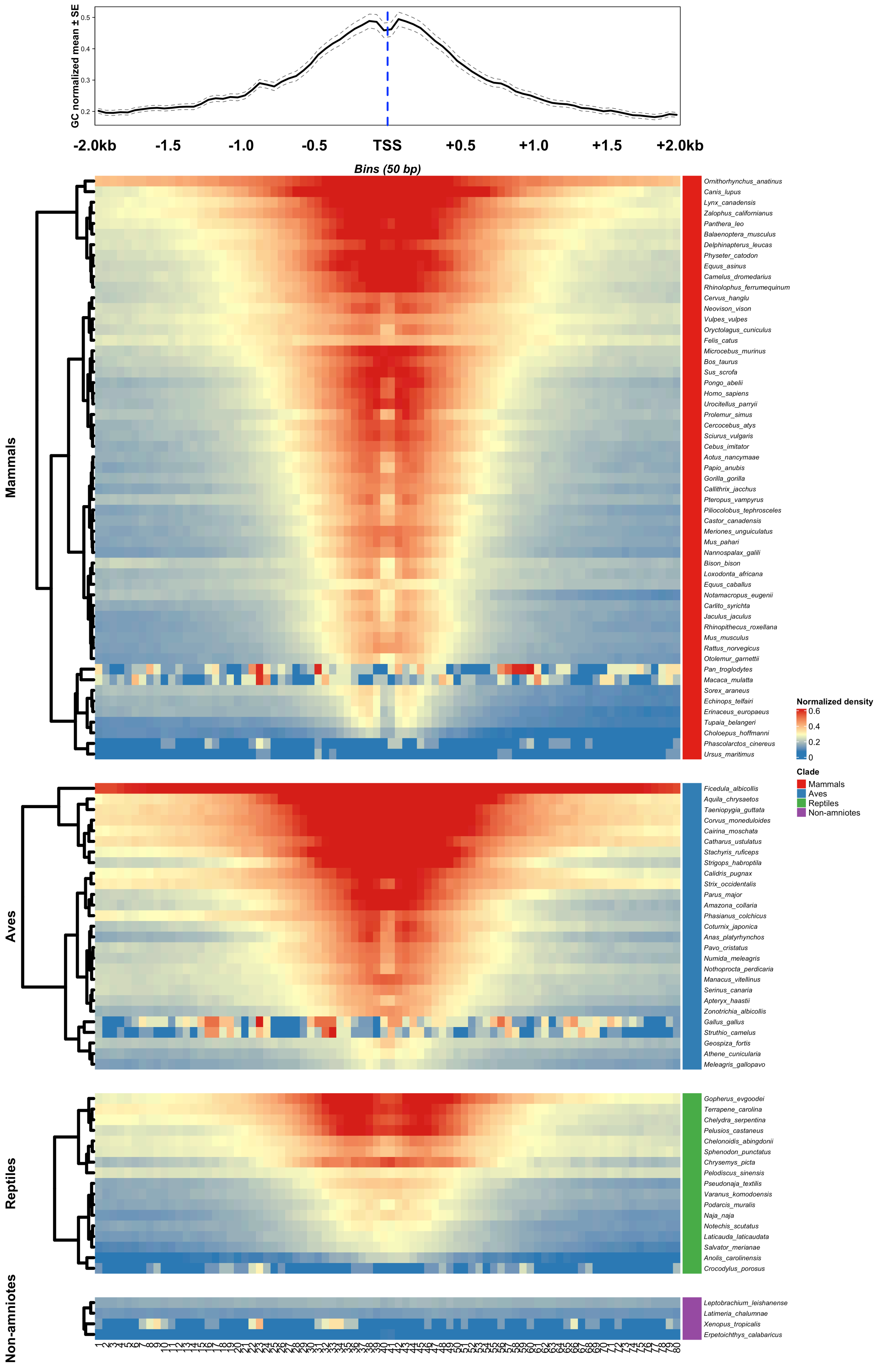

### Fig. S4

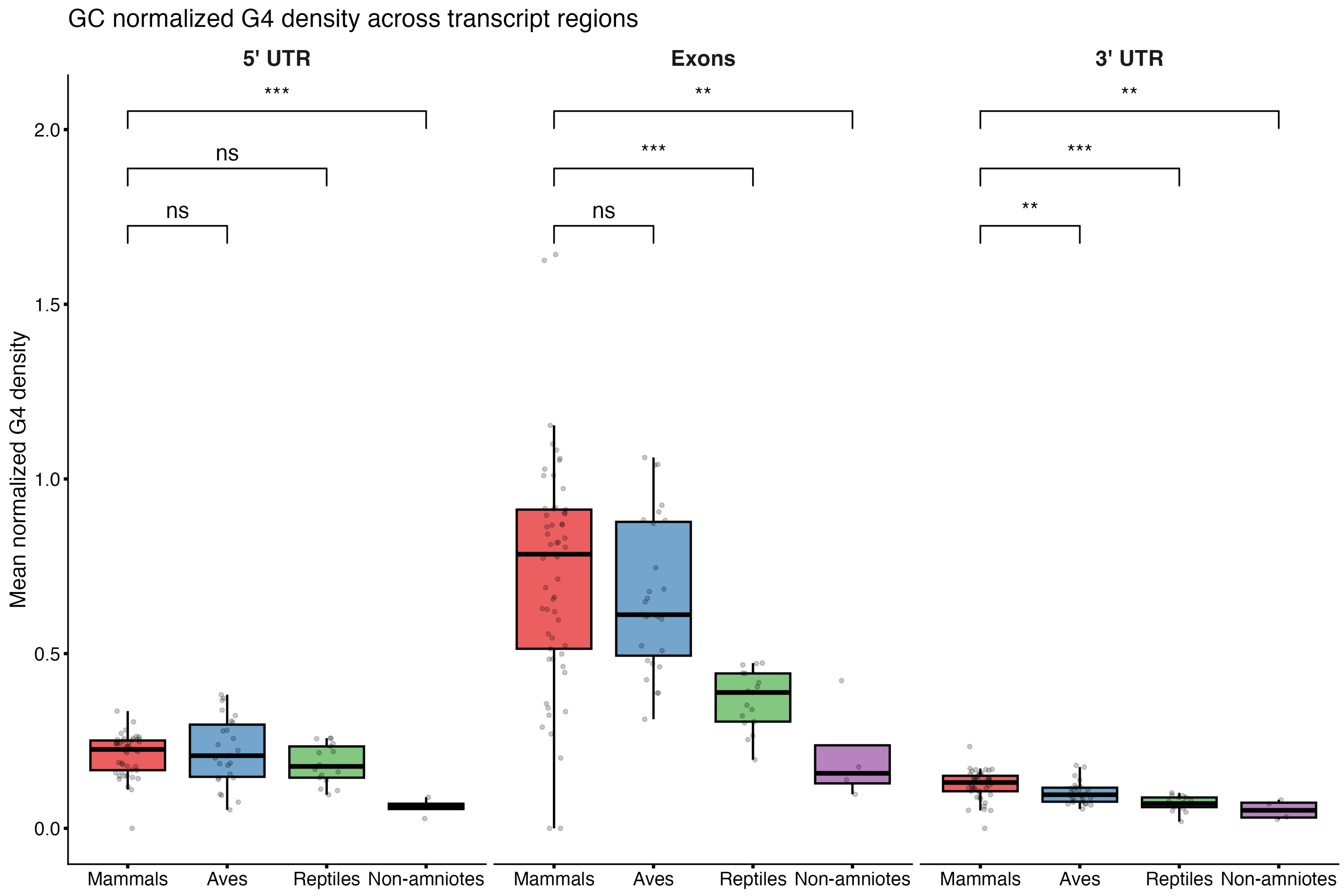

### Fig. S5

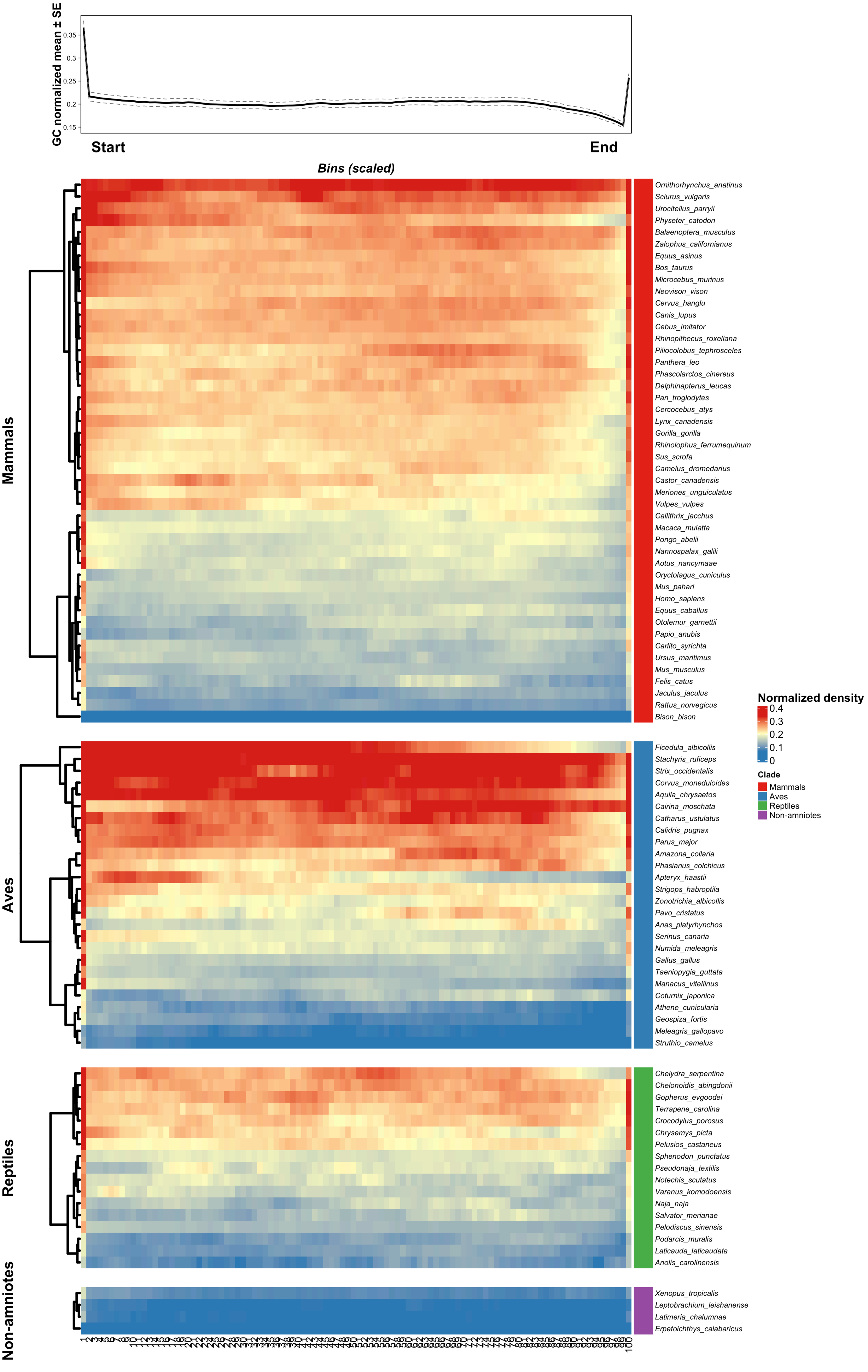

### Fig. S6

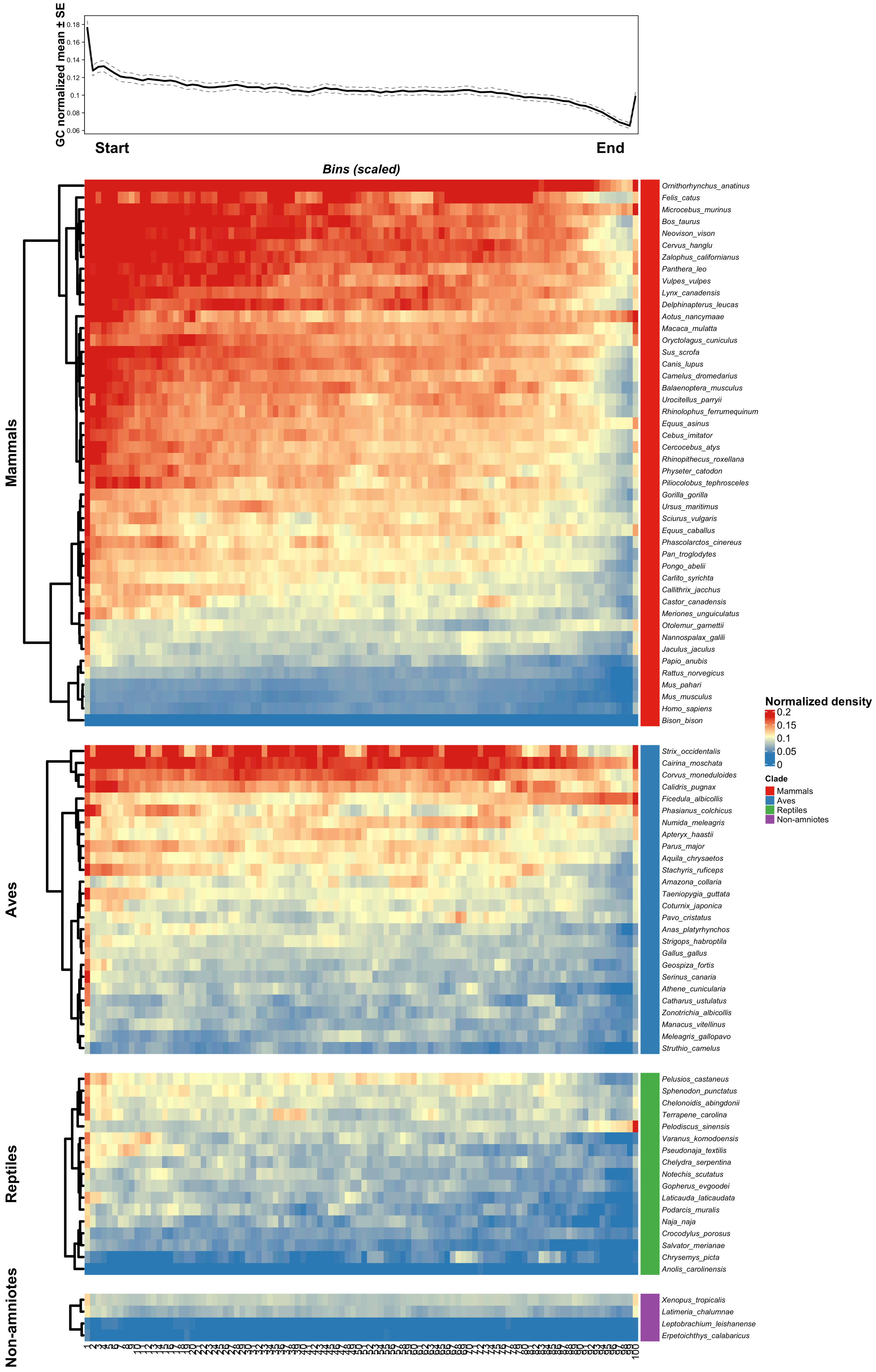

### Fig. S7

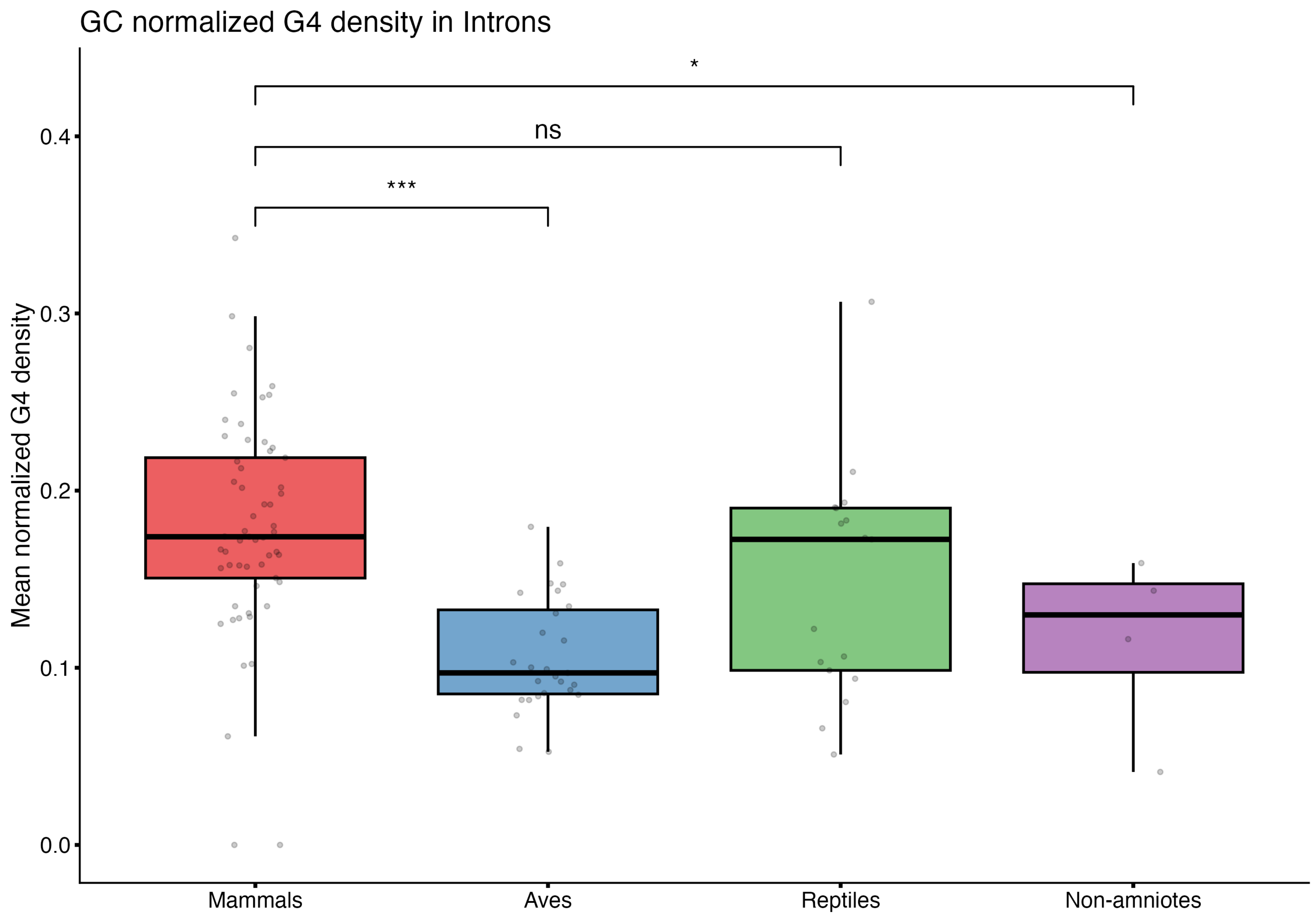

### Fig. S8

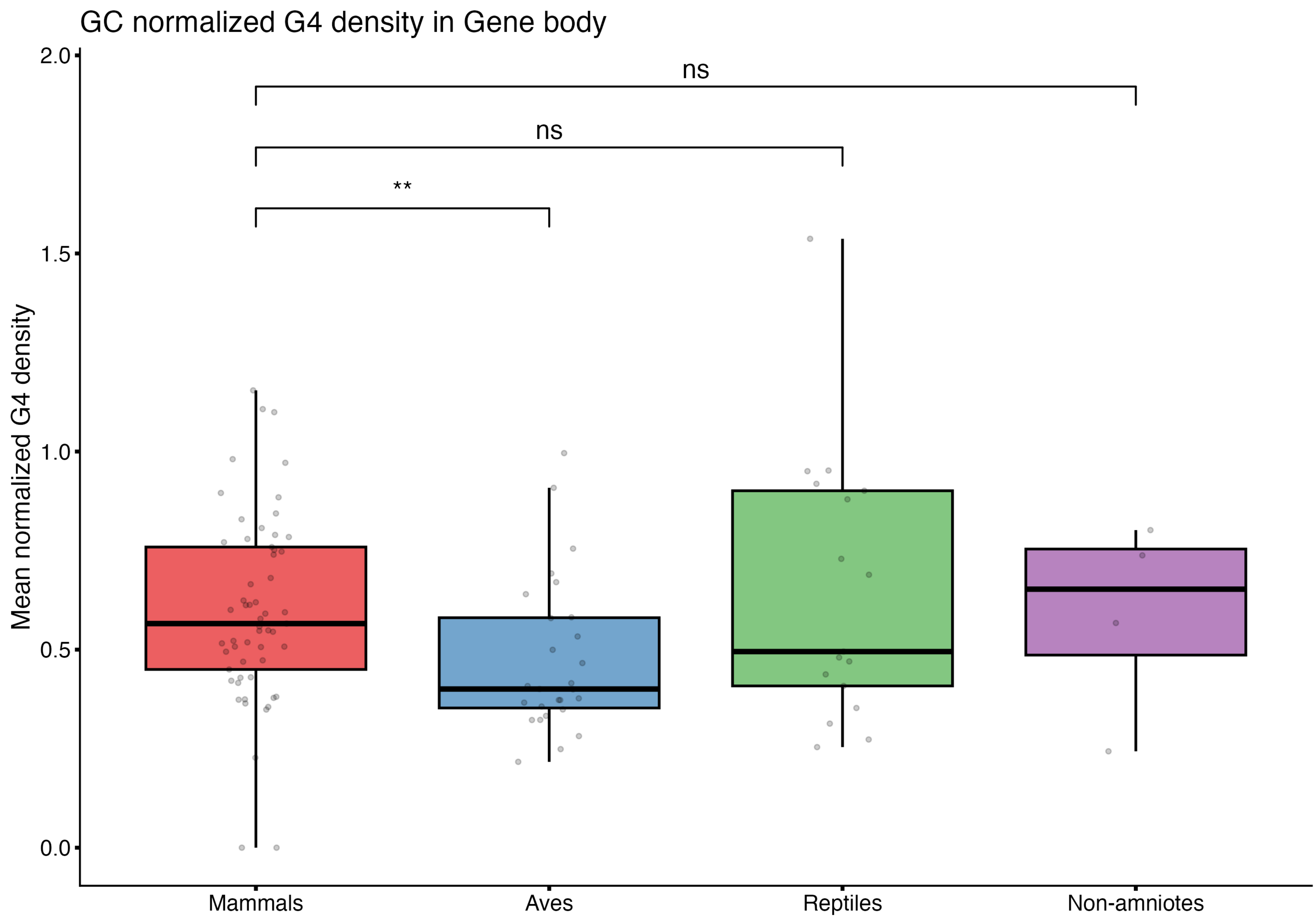

### Fig. S9

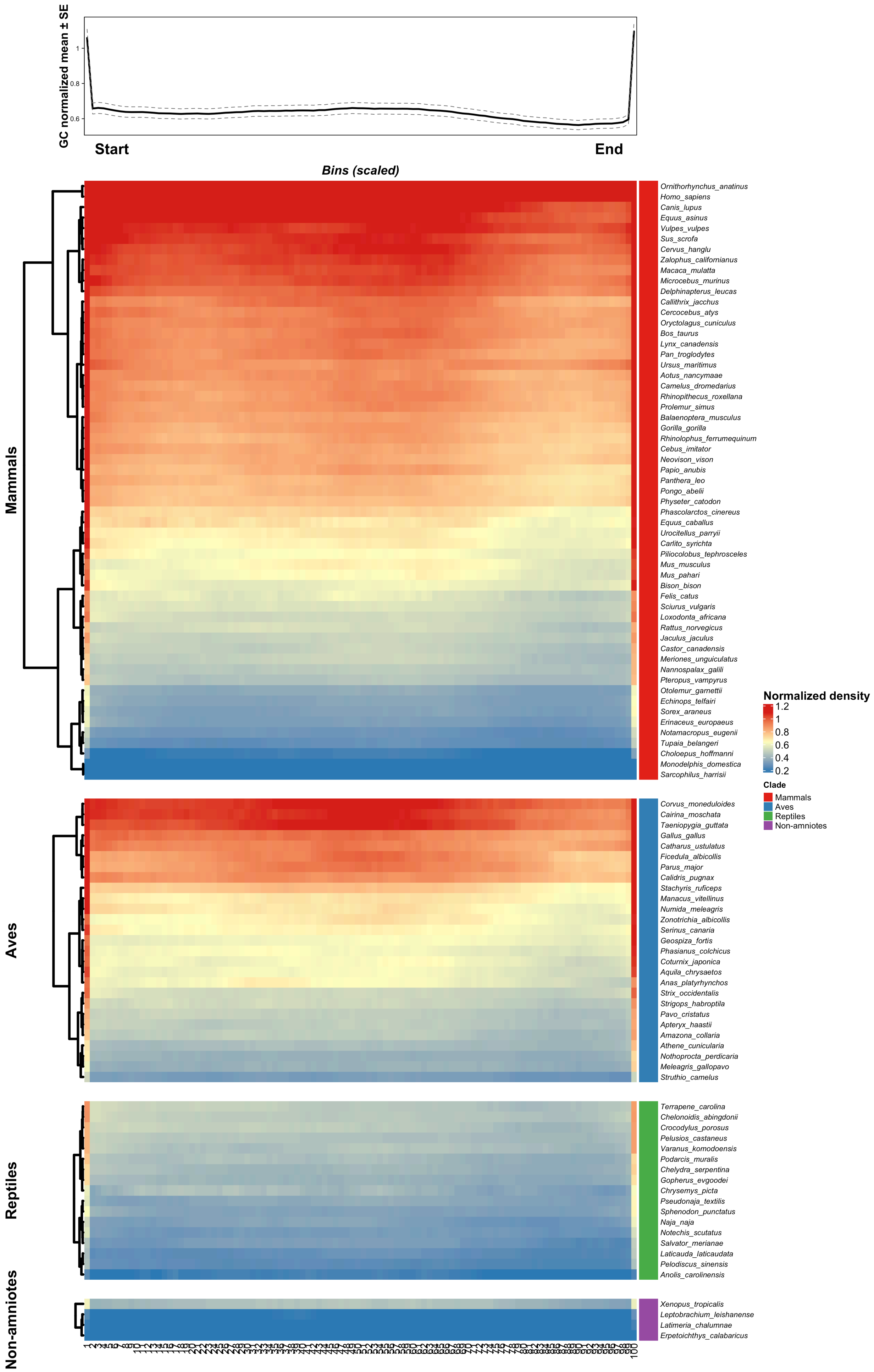

### Fig. S10

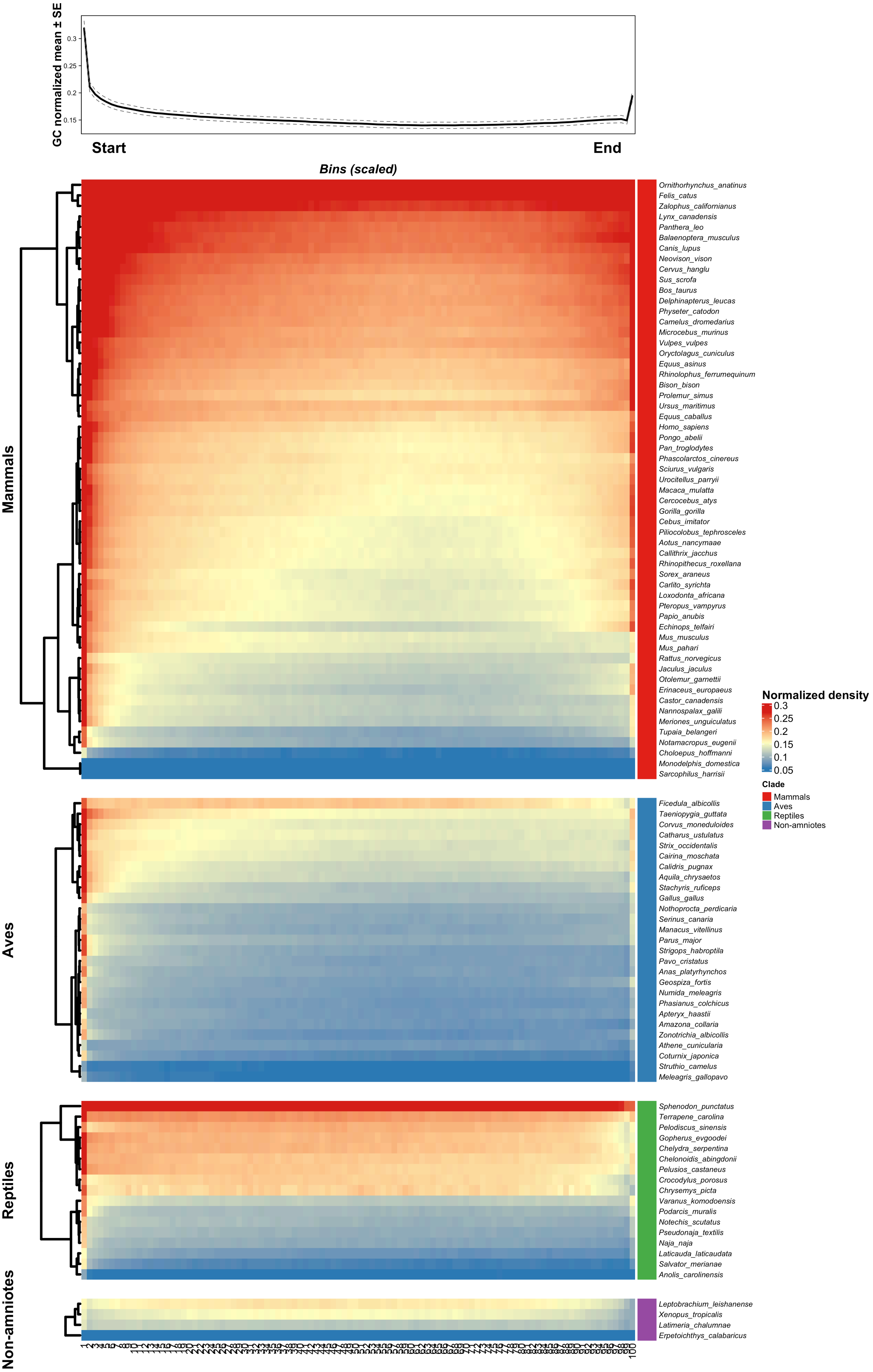

### Fig. S11

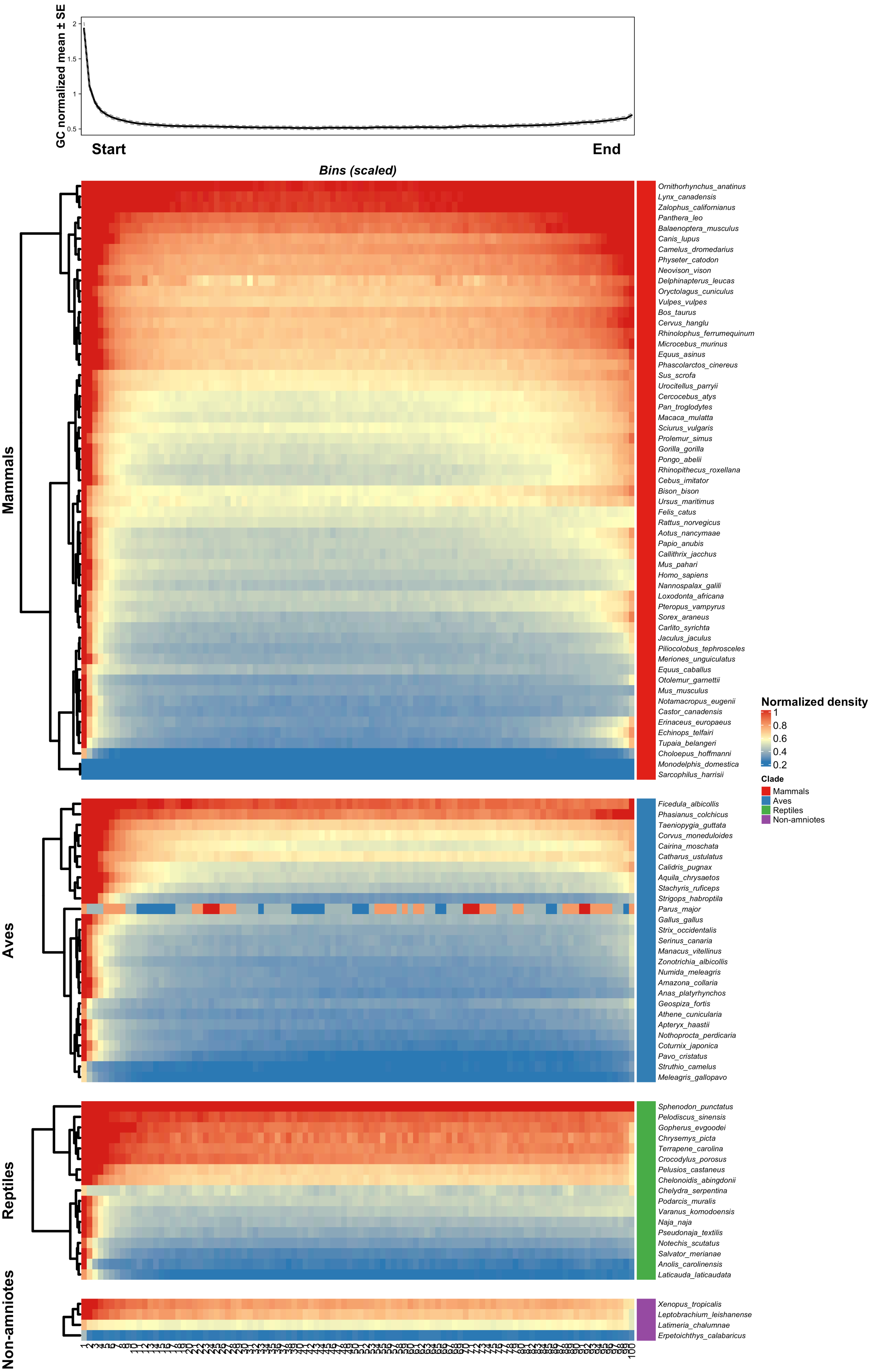

### Fig. S12

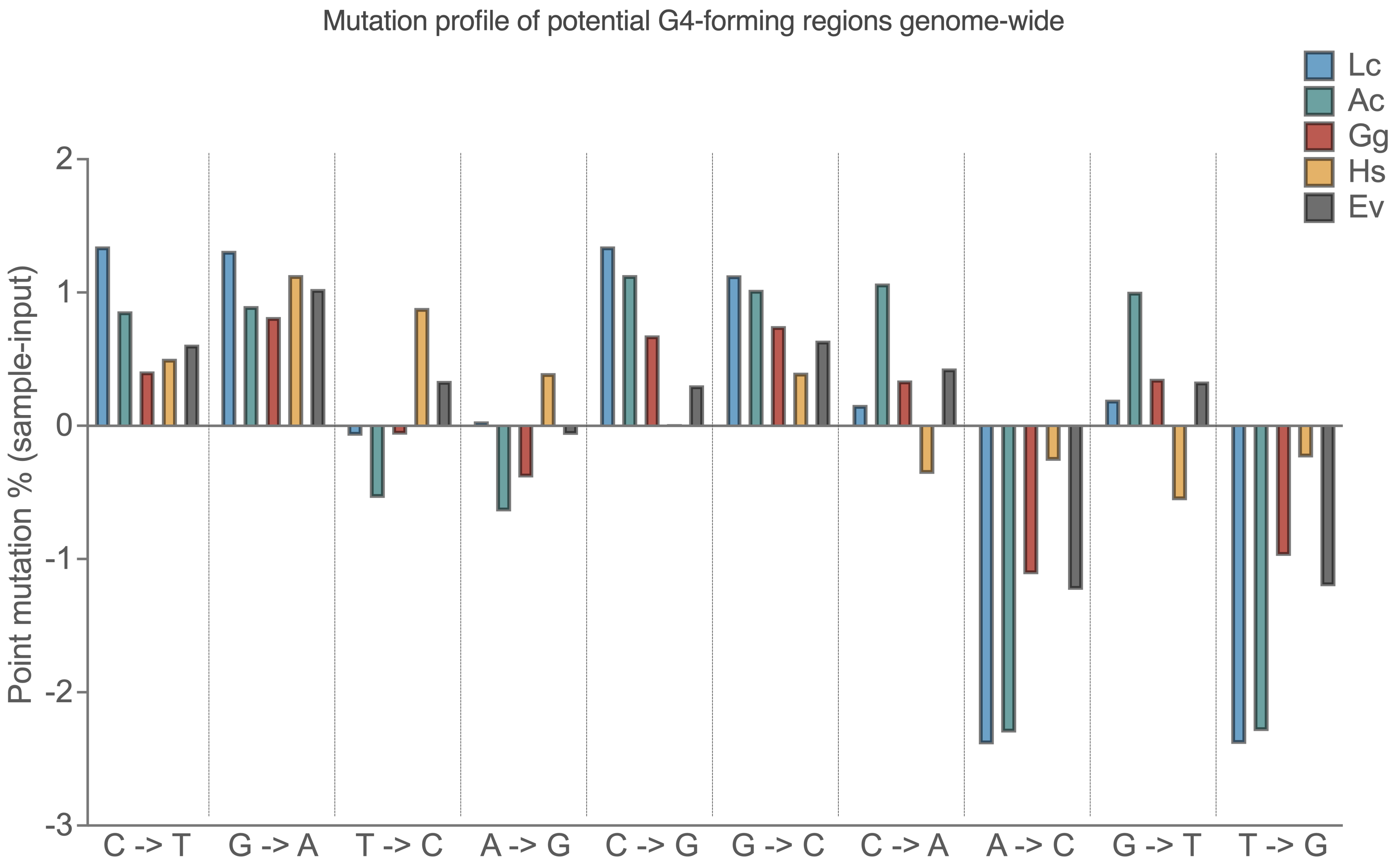

### Fig. S13

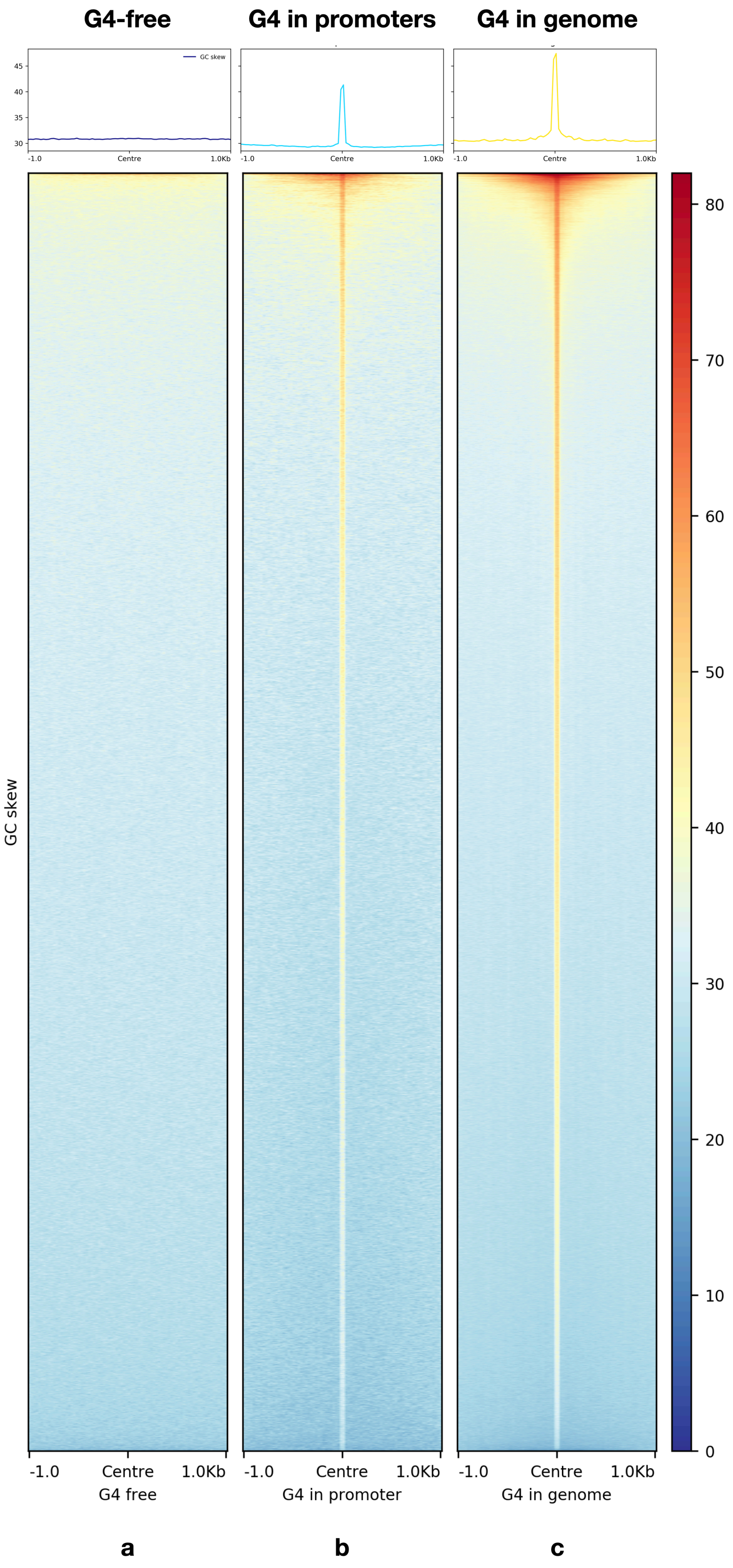

### Fig. S14

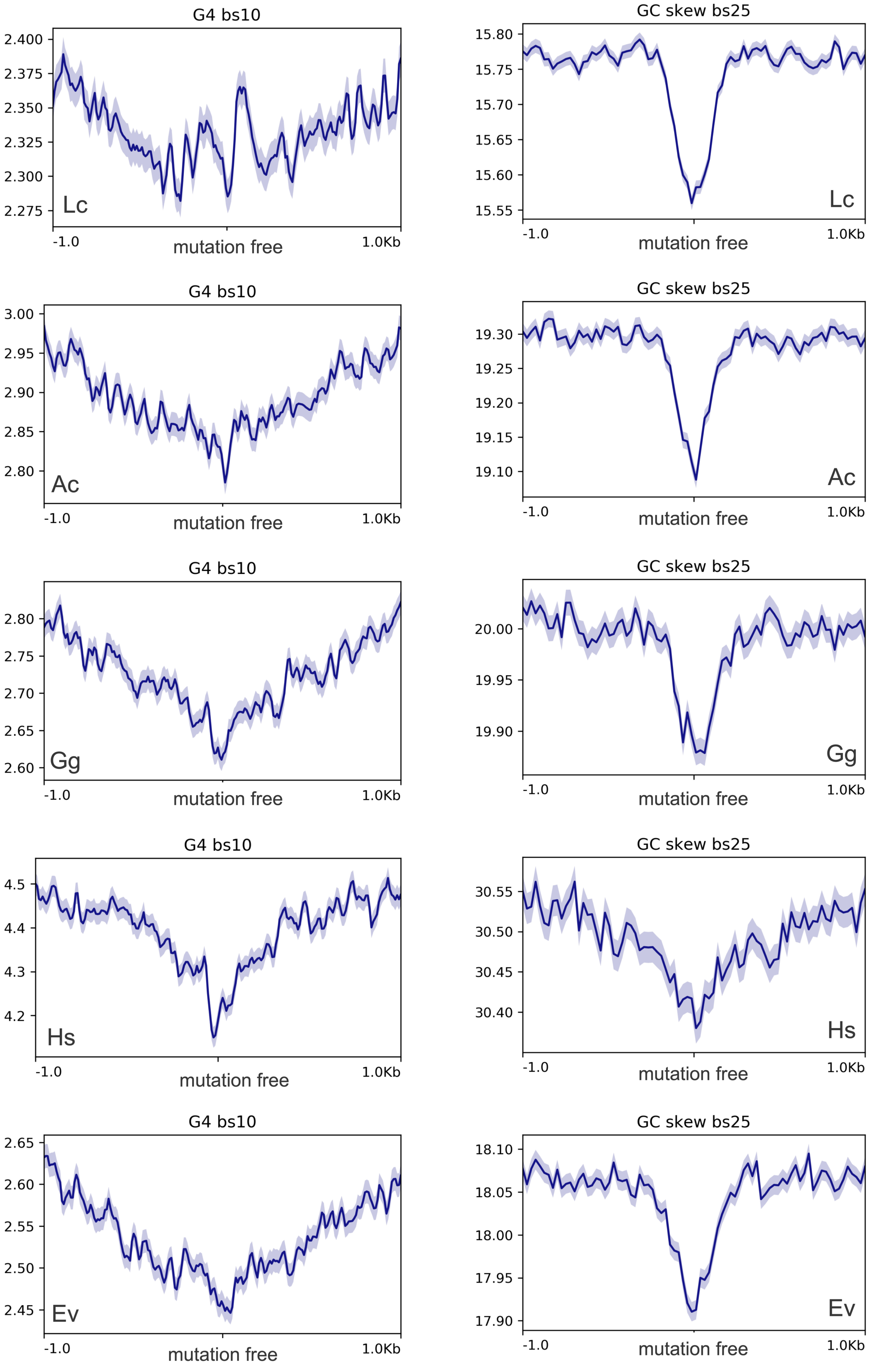

### Fig. S15

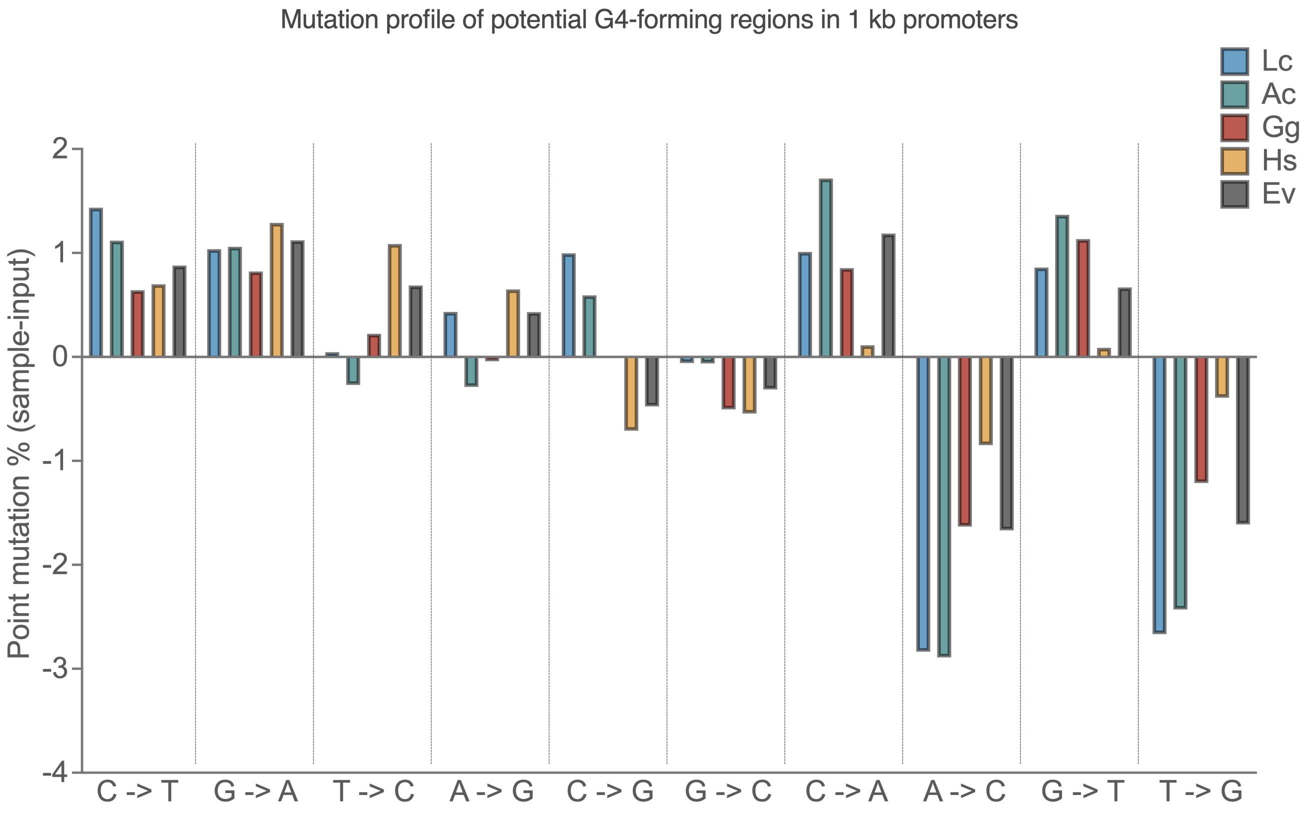

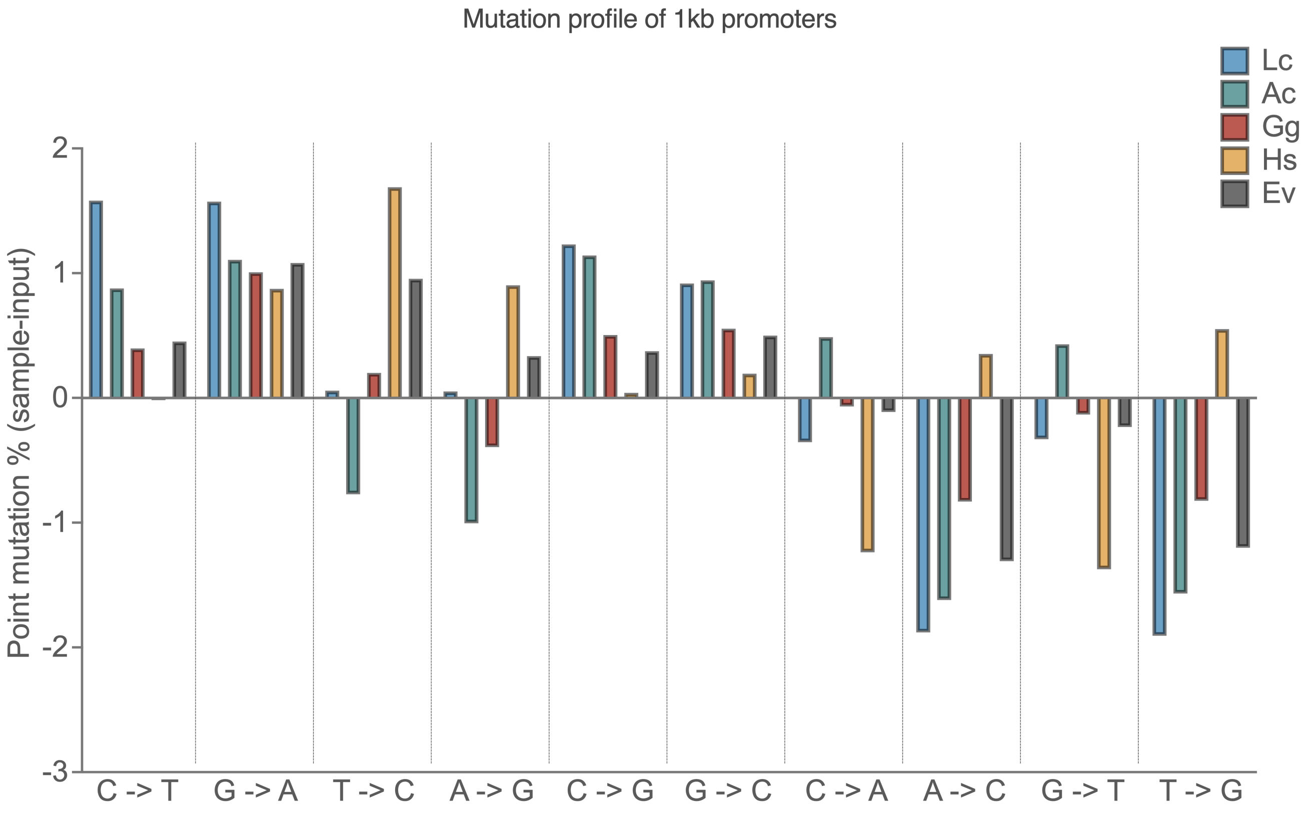

### Fig. S16

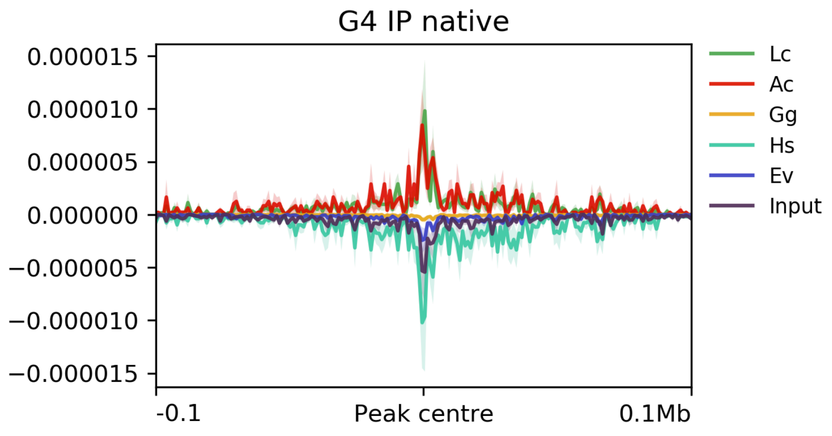

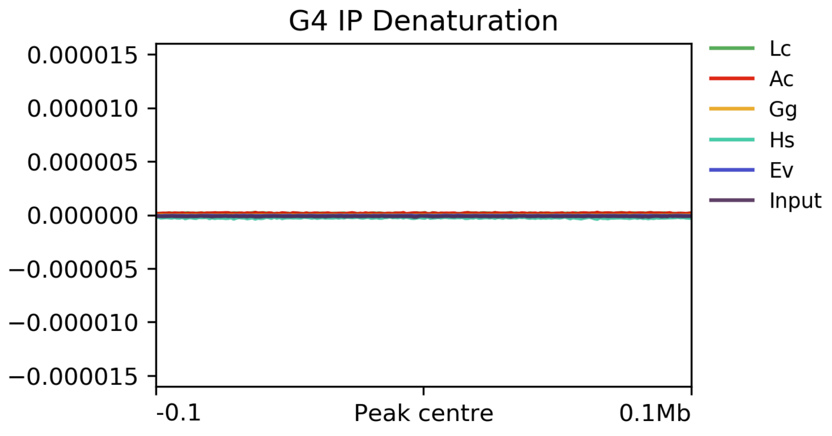

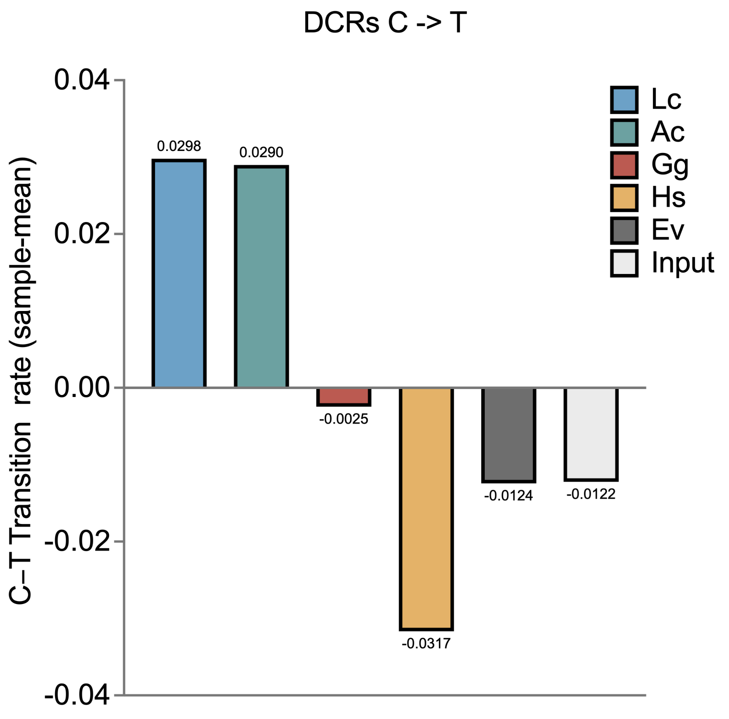

### Fig. S17

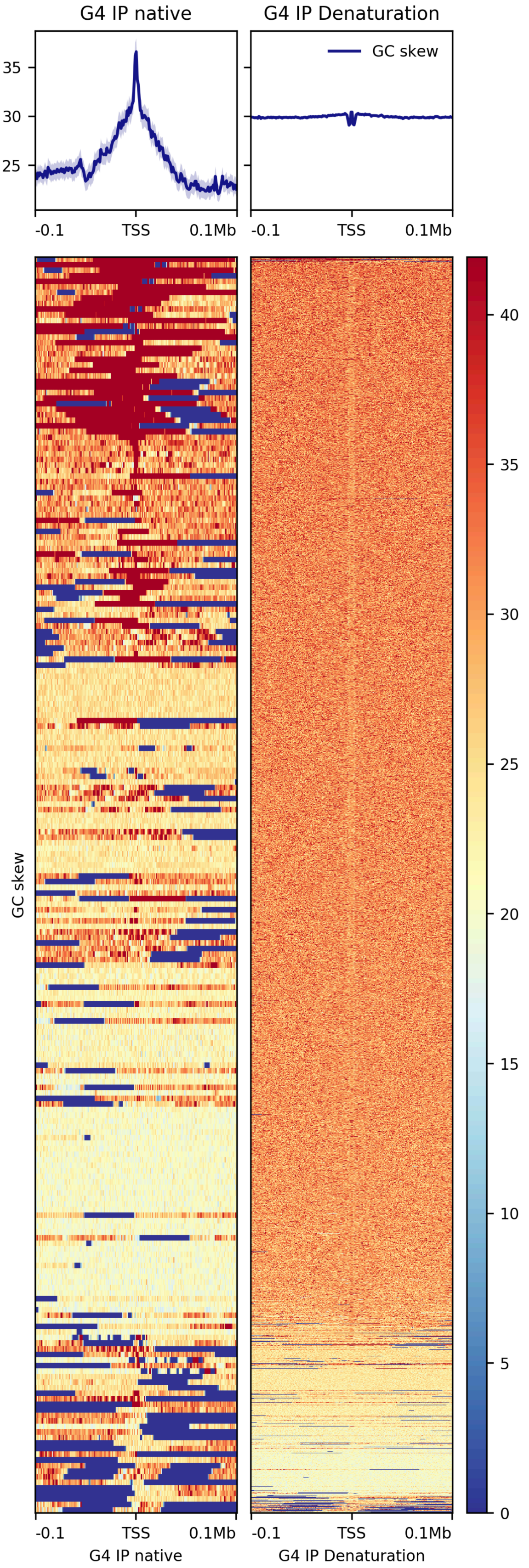

### Fig. S18

meC-to-T transition in GC-rich TFBSs

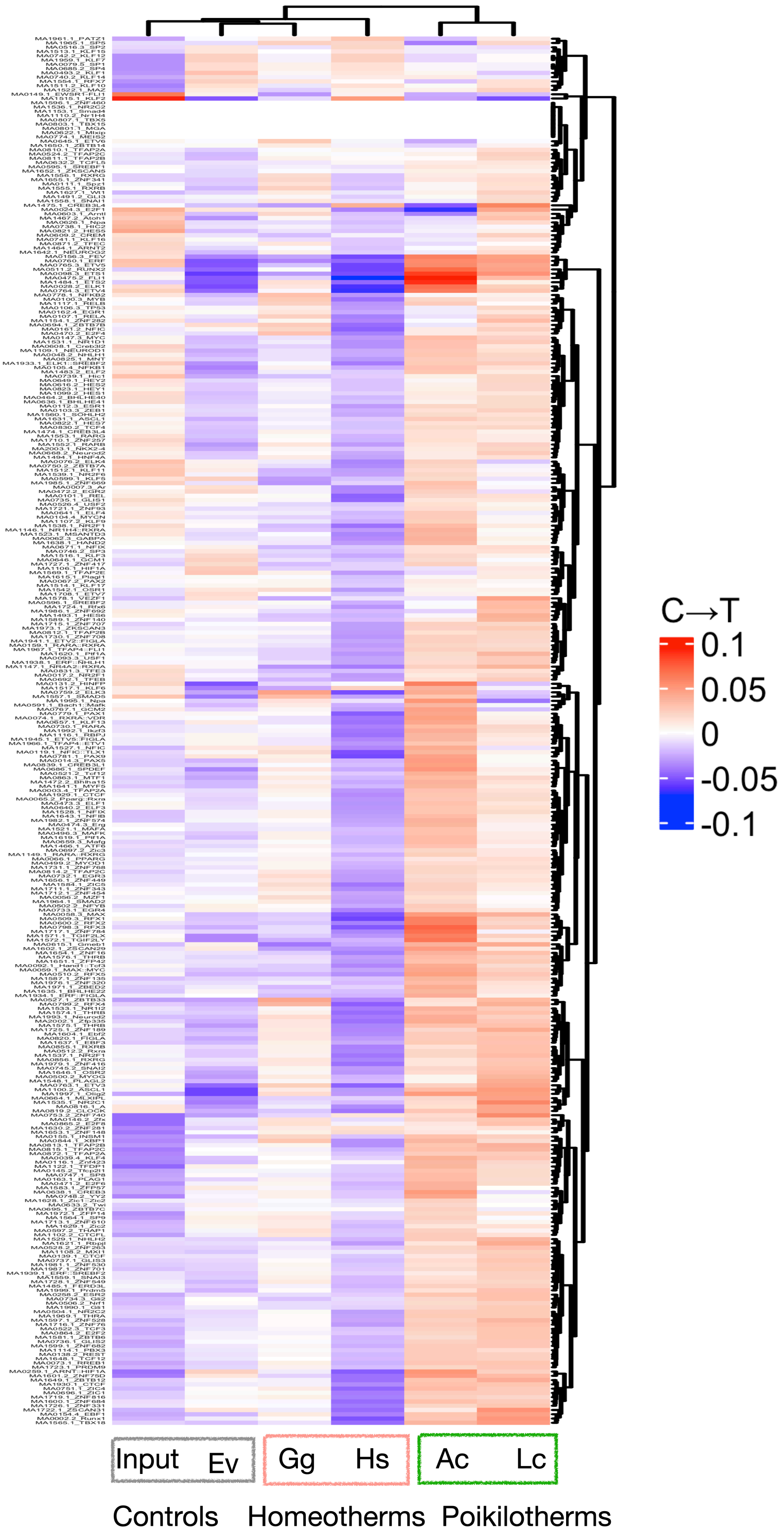

### Fig. S19

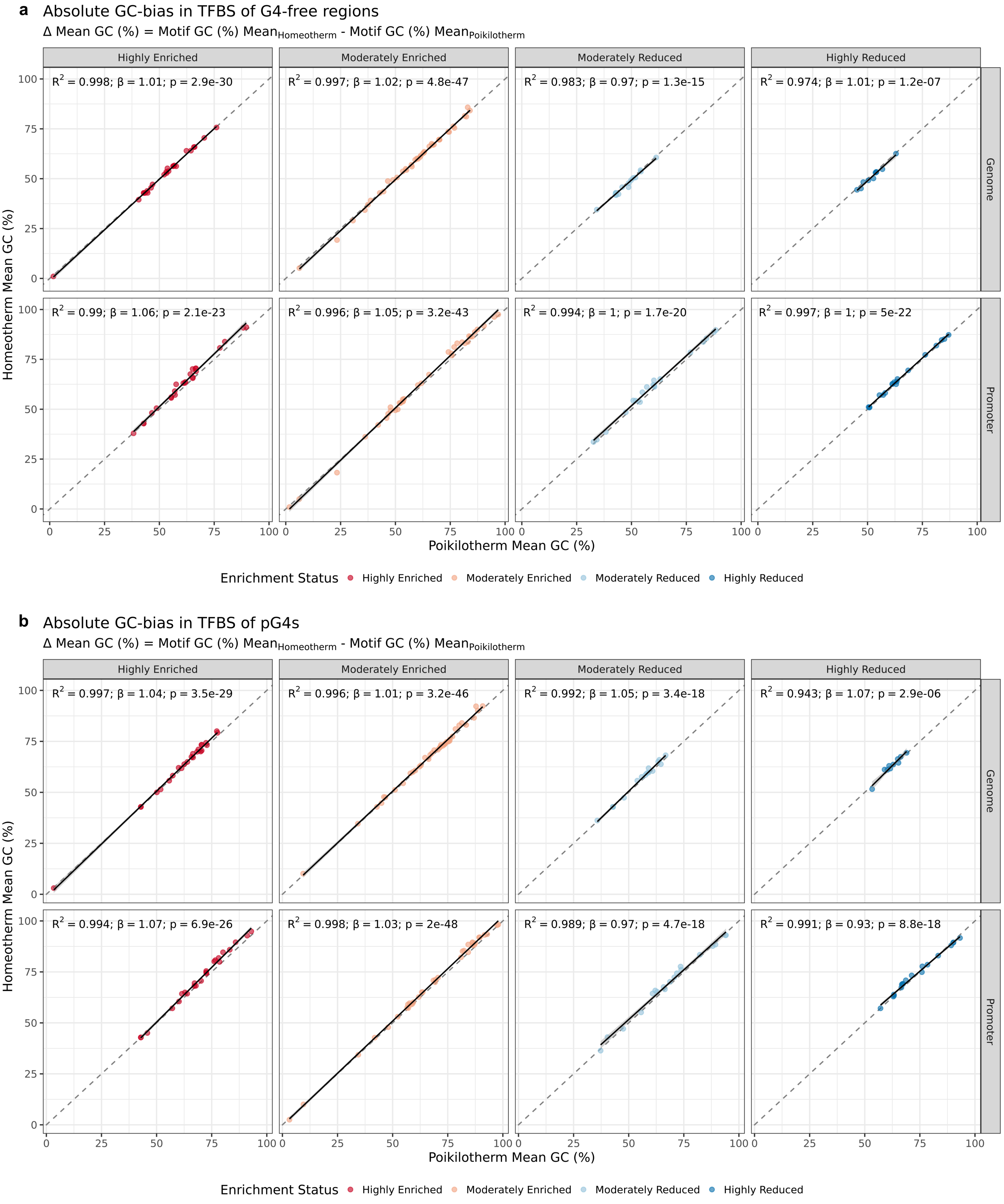
