## Supplementary material for "CGGBP1-regulated heterogeneous C–T transition rates relate with G-quadruplex potential of terrestrial vertebrate genomes": Fig. S20

### Genome : Highly Enriched in G4s

*MA0024.3 - E2F1*

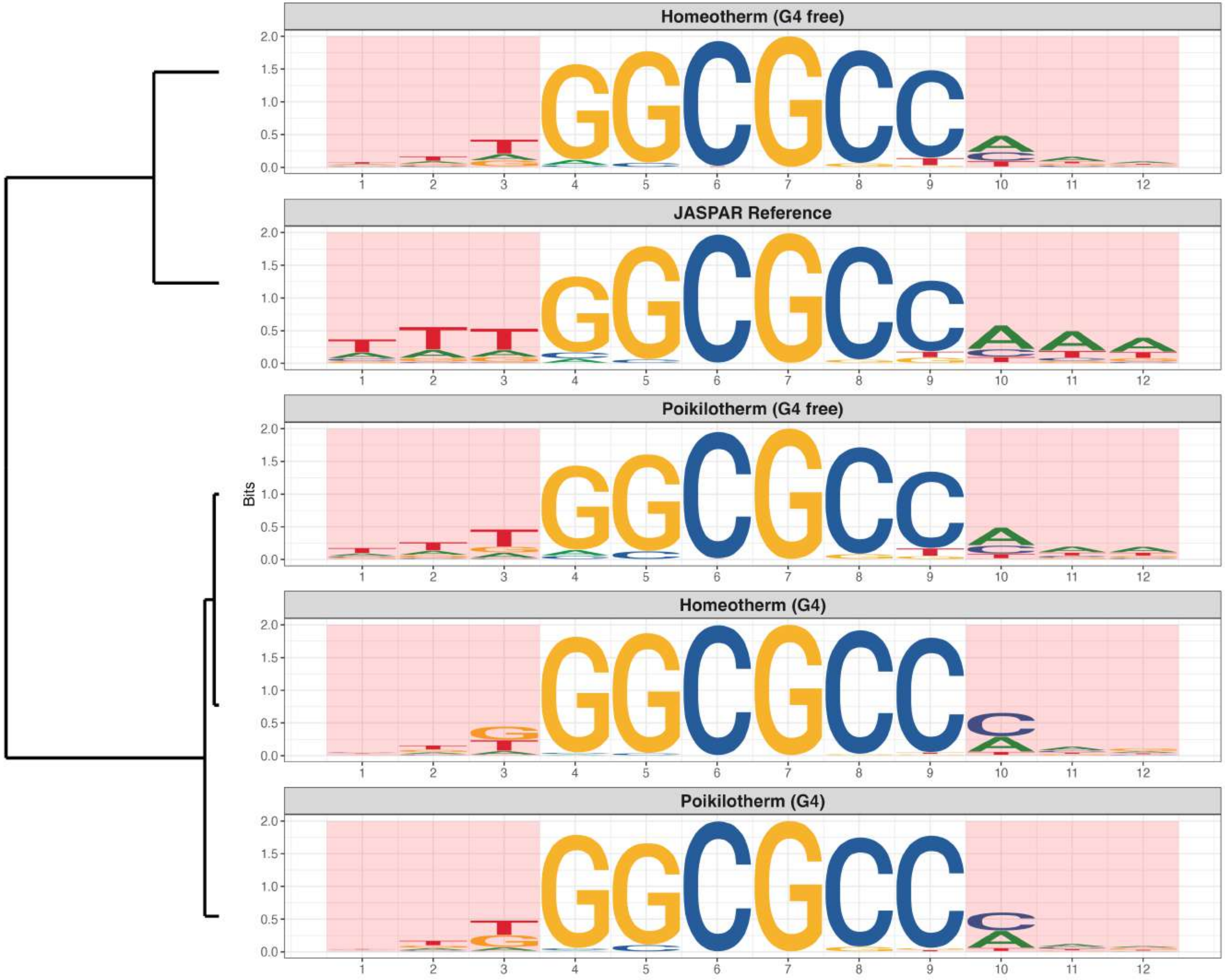

### Genome : Highly Enriched in G4s

*MA0067.3 - PAX2*

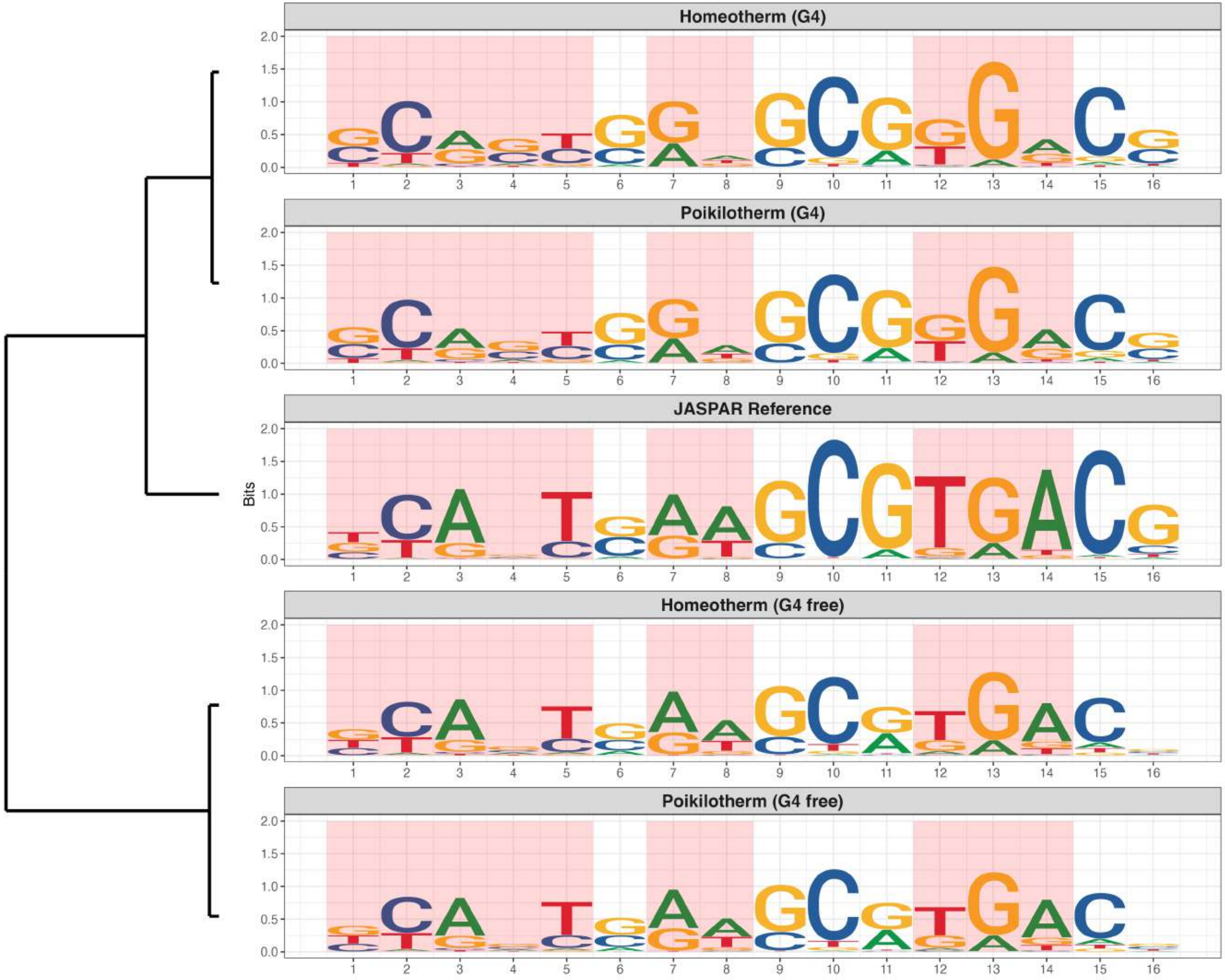

### Genome : Highly Enriched in G4s

*MA0076.3 - ELK4*

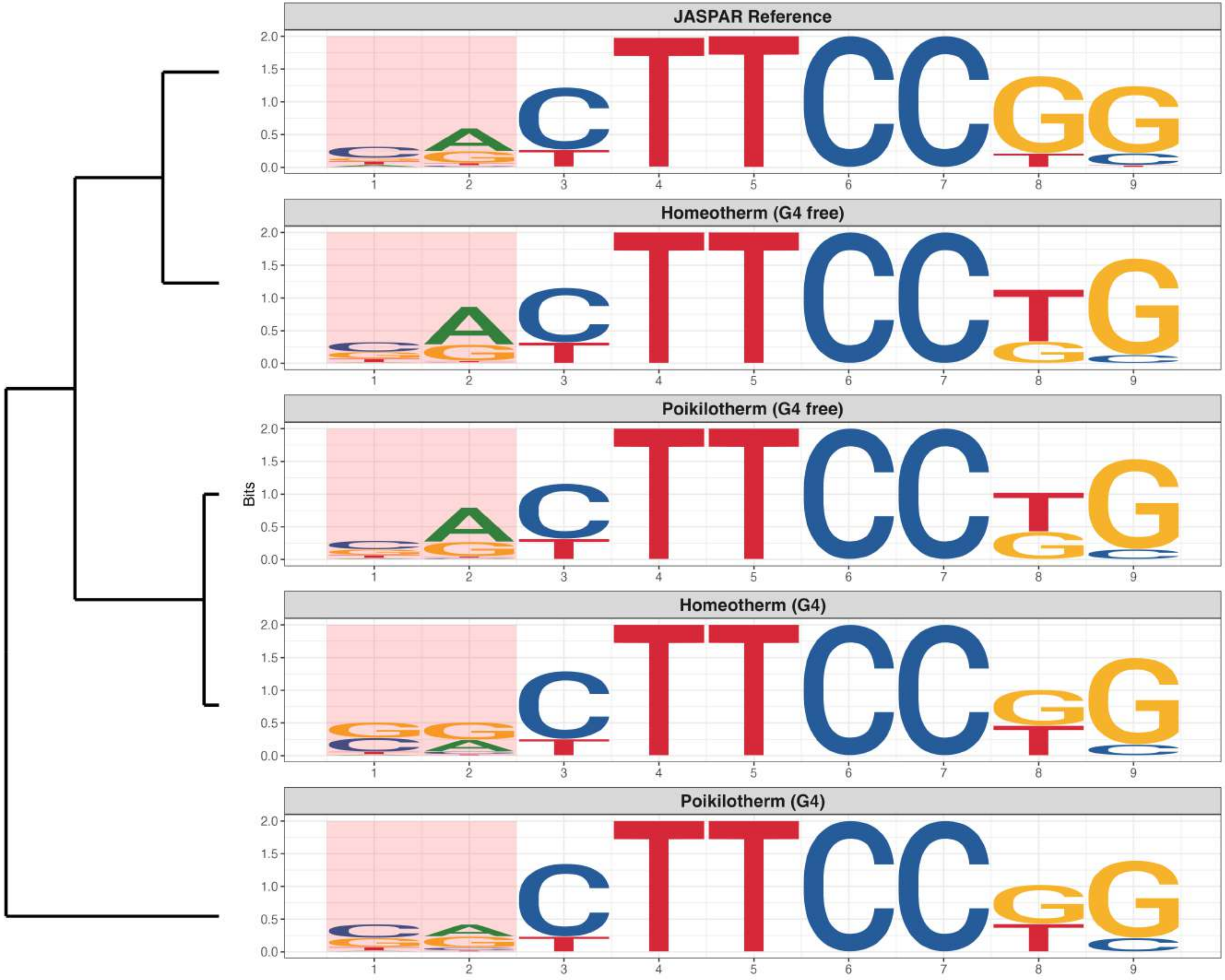

### Genome : Highly Enriched in G4s

*MA0124.3 - Nkx3-1*

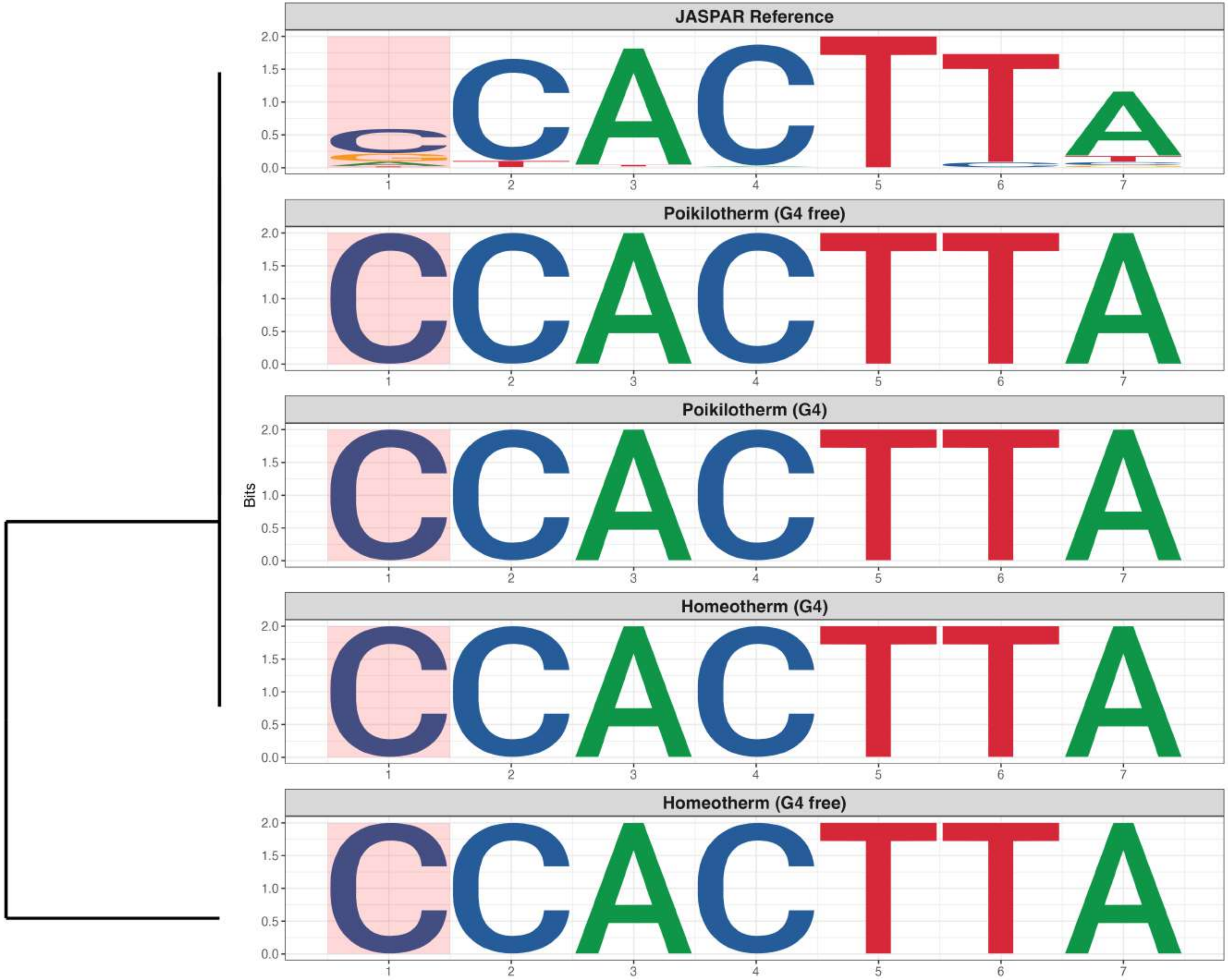

### Genome : Highly Enriched in G4s

*MA0500.3 - MYOG*

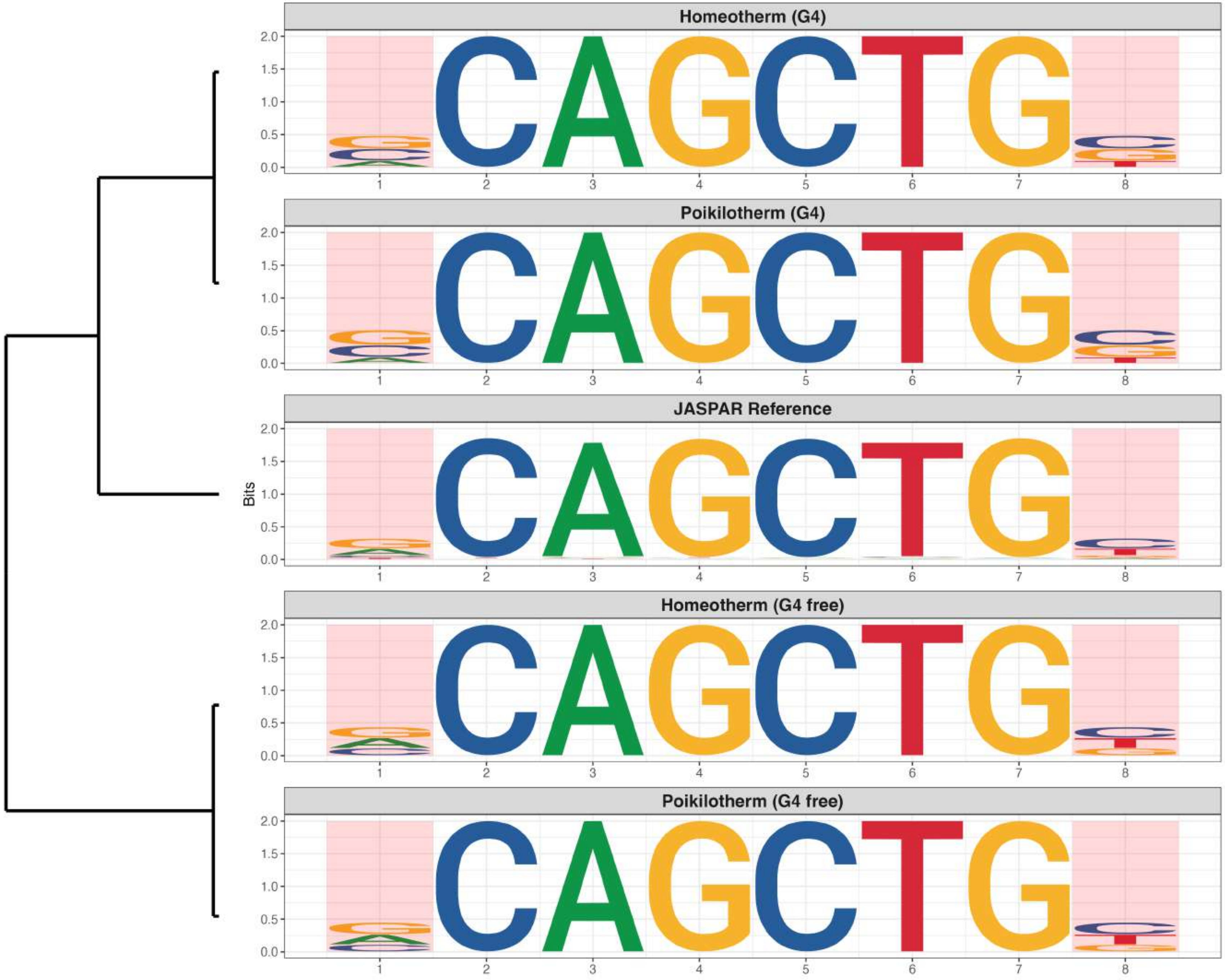

### Genome : Highly Enriched in G4s

*MA0508.4 - PRDM1*

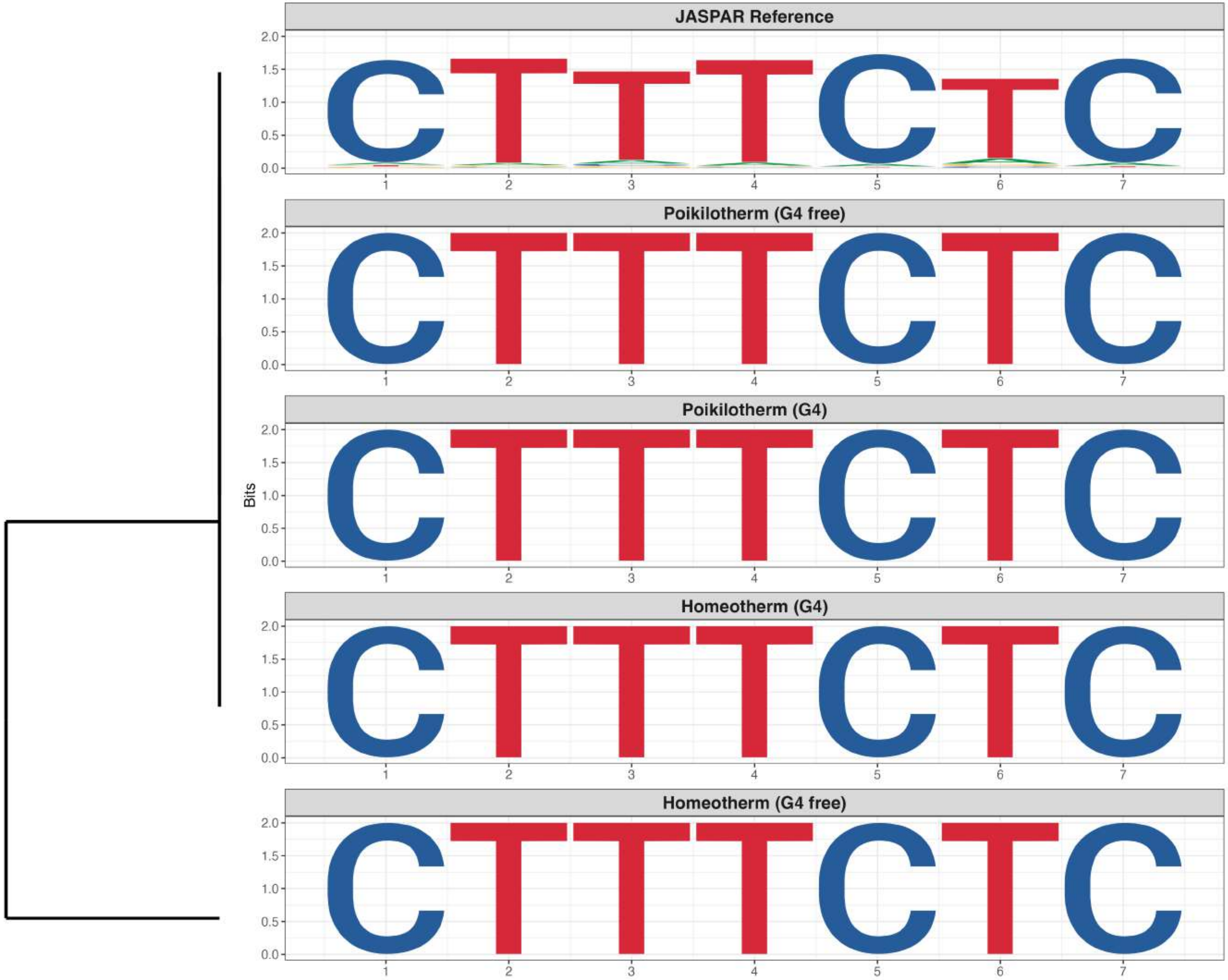

### Genome : Highly Enriched in G4s

*MA0519.2 - Stat5a::Stat5b*

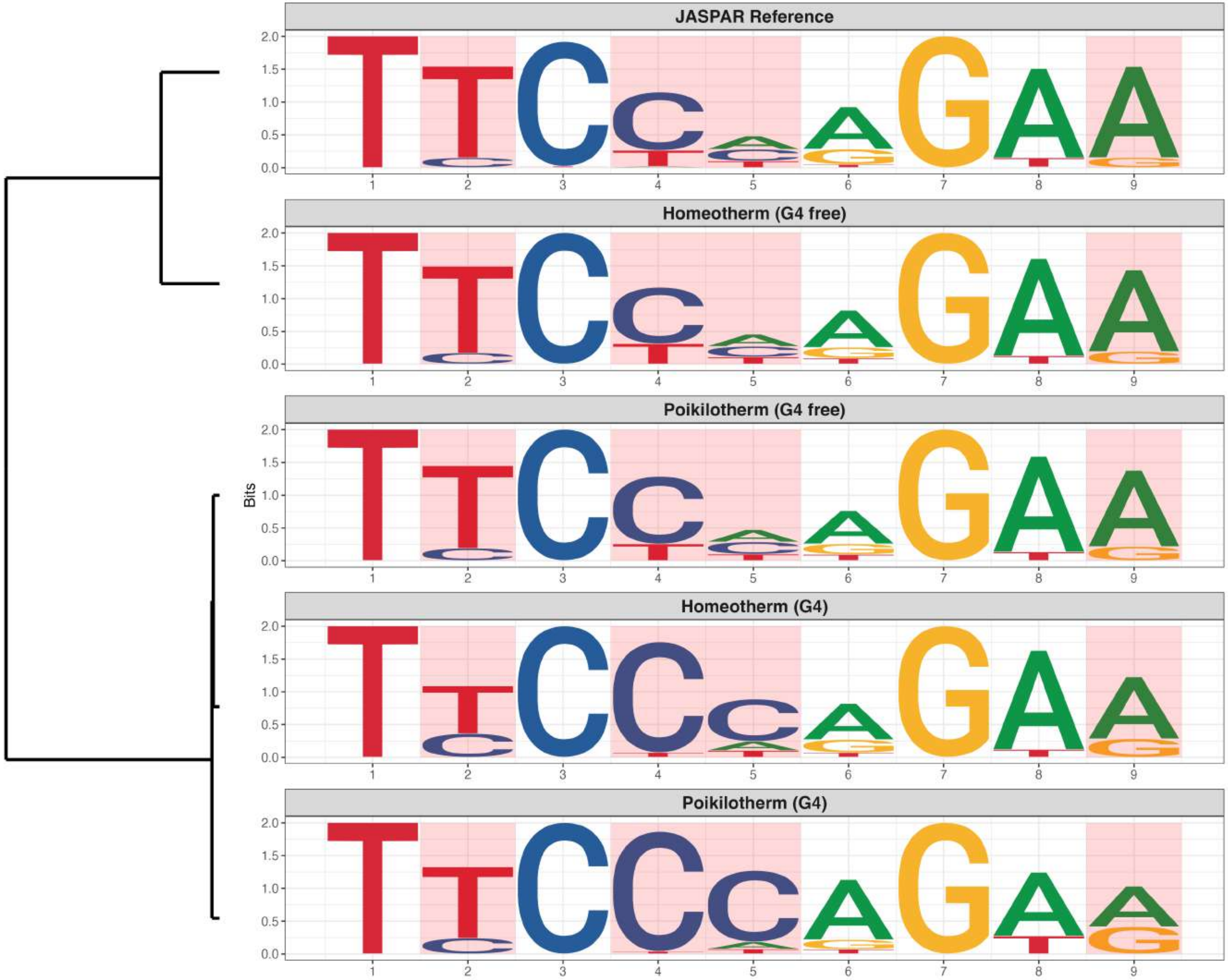

### Genome : Highly Enriched in G4s

*MA0596.1 - SREBF2*

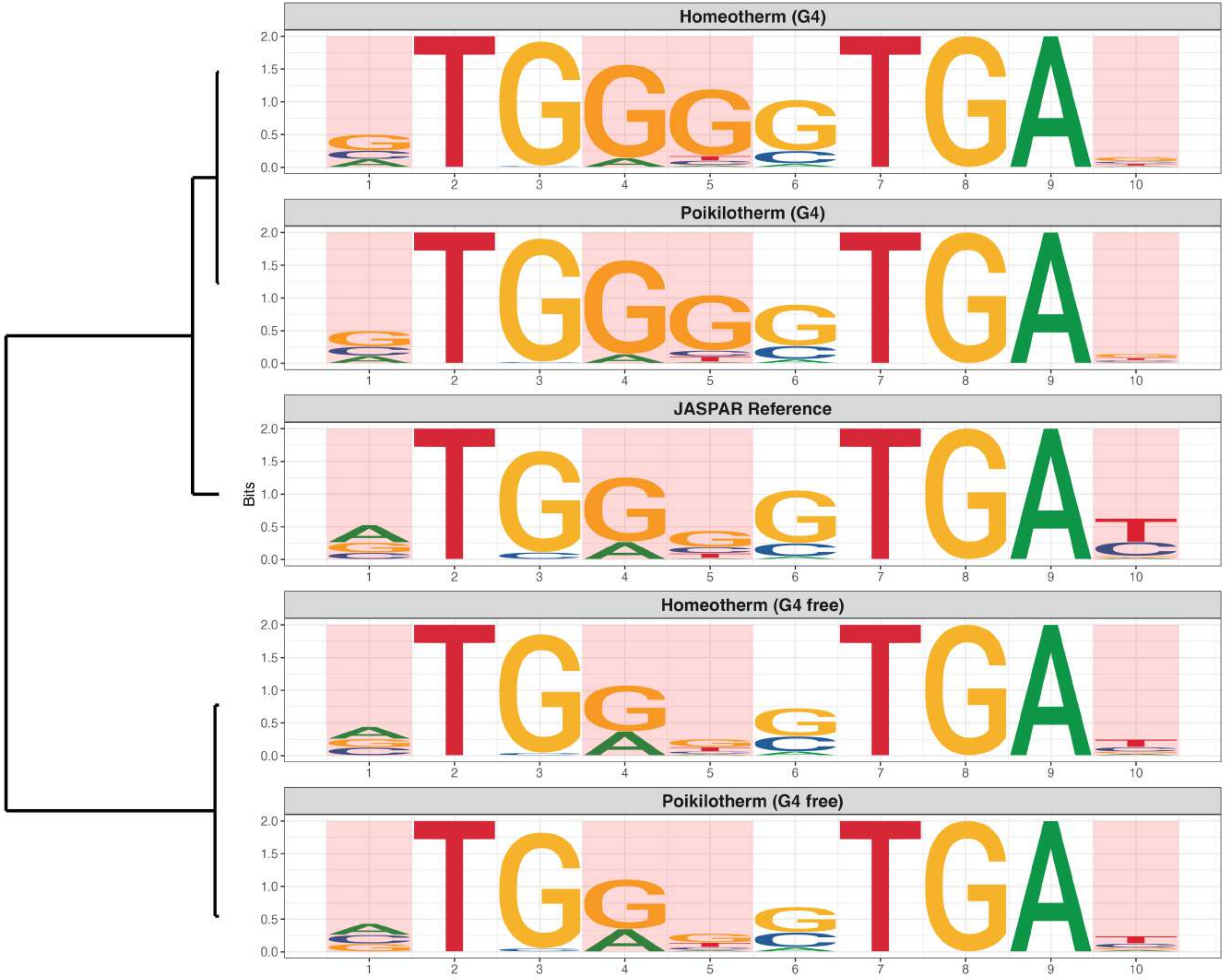

### Genome : Highly Enriched in G4s

*MA0779.2 - PAX1*

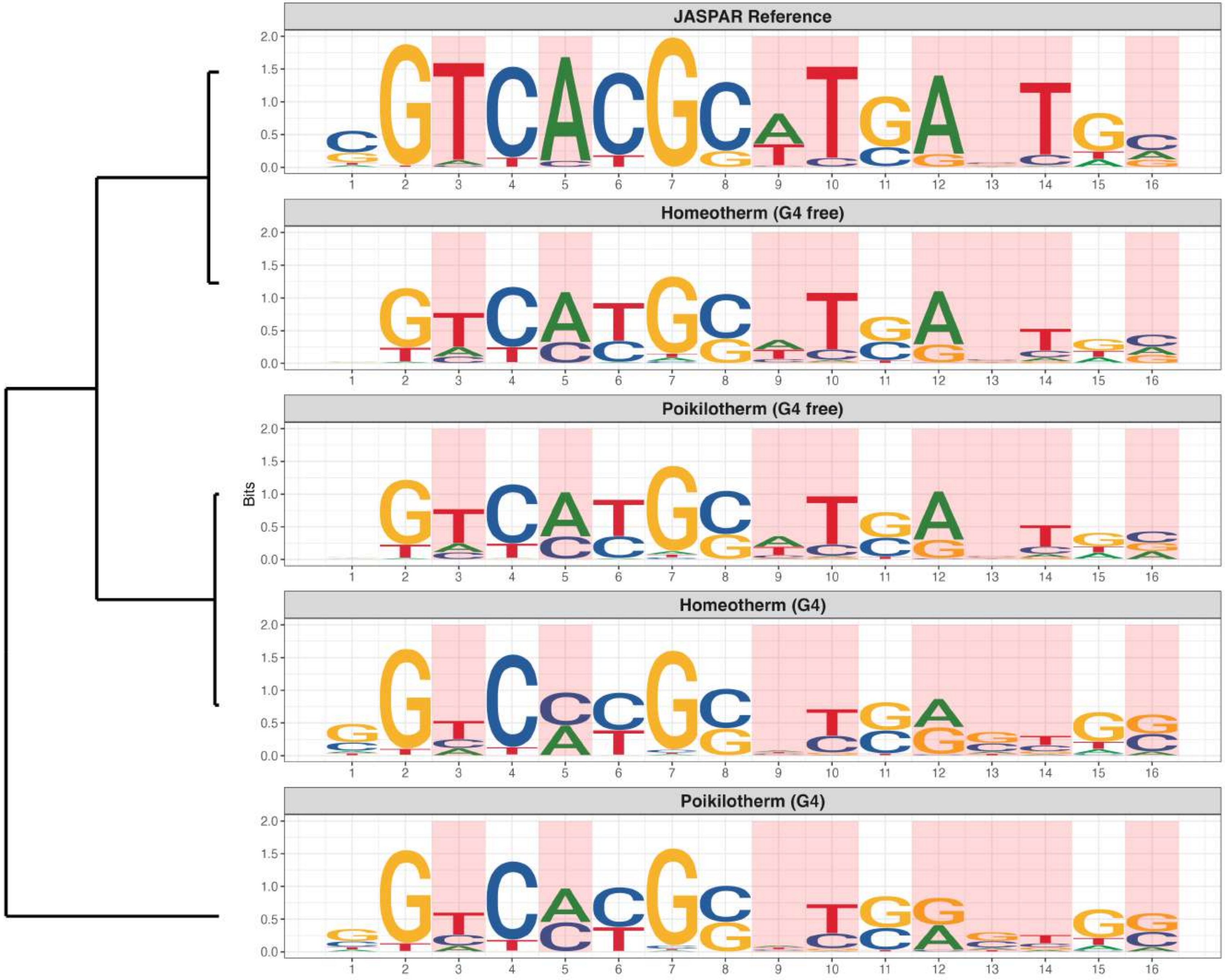

### Genome : Highly Enriched in G4s

*MA0781.2 - PAX9*

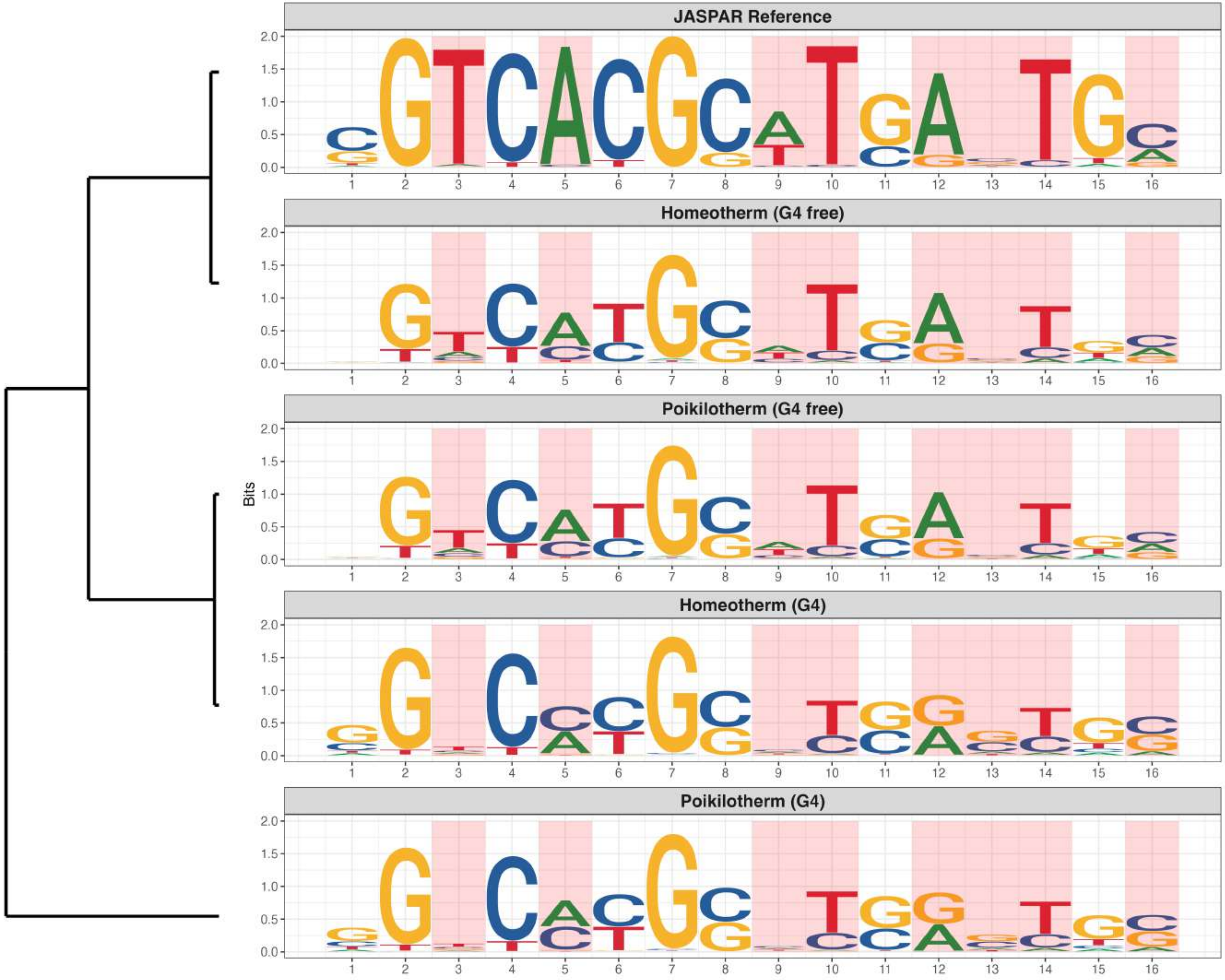

### Genome : Highly Enriched in G4s

*MA0858.1 - Rarb*

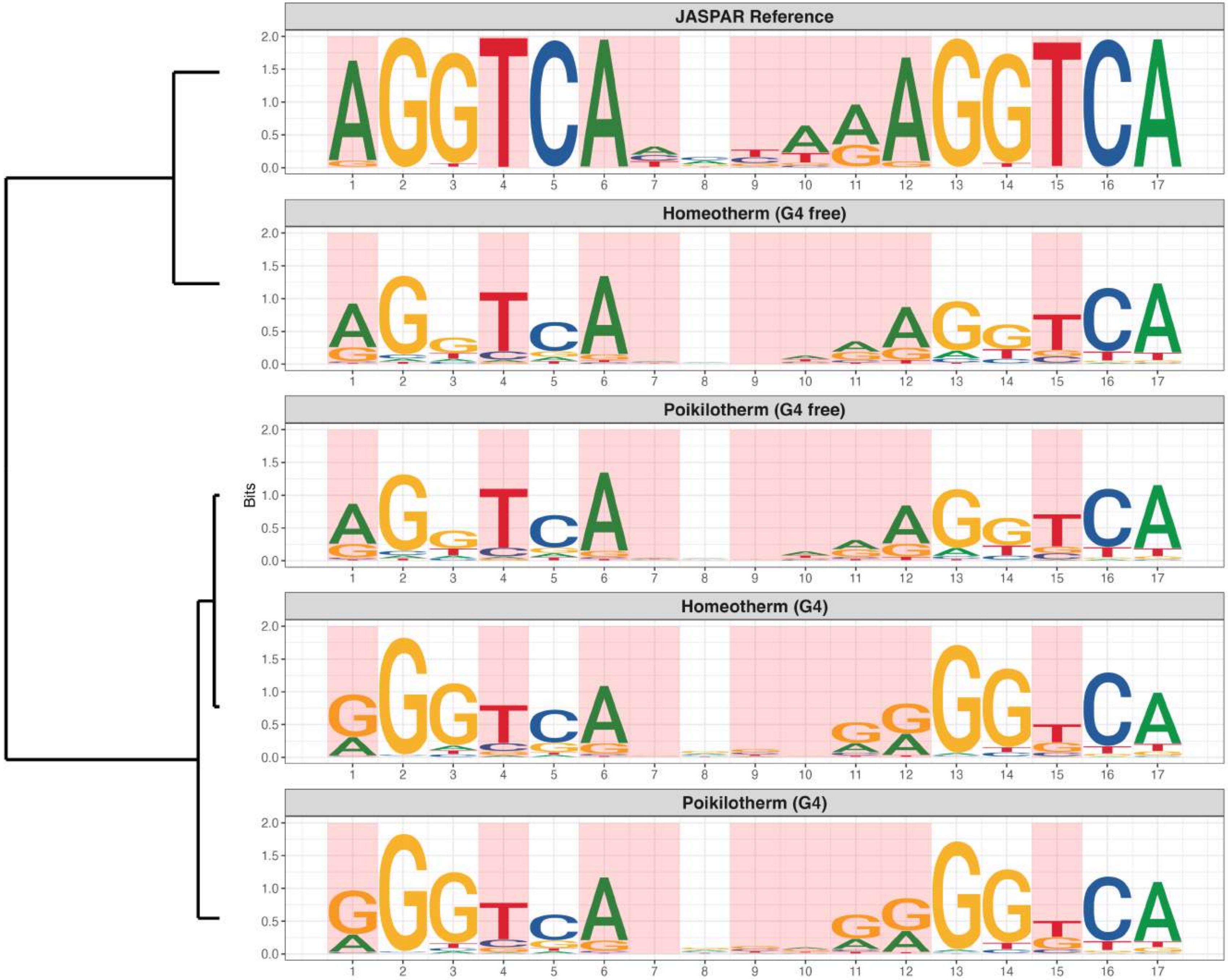

**Genome : Highly Enriched in G4s**

*MA0863.1 - MTF1*

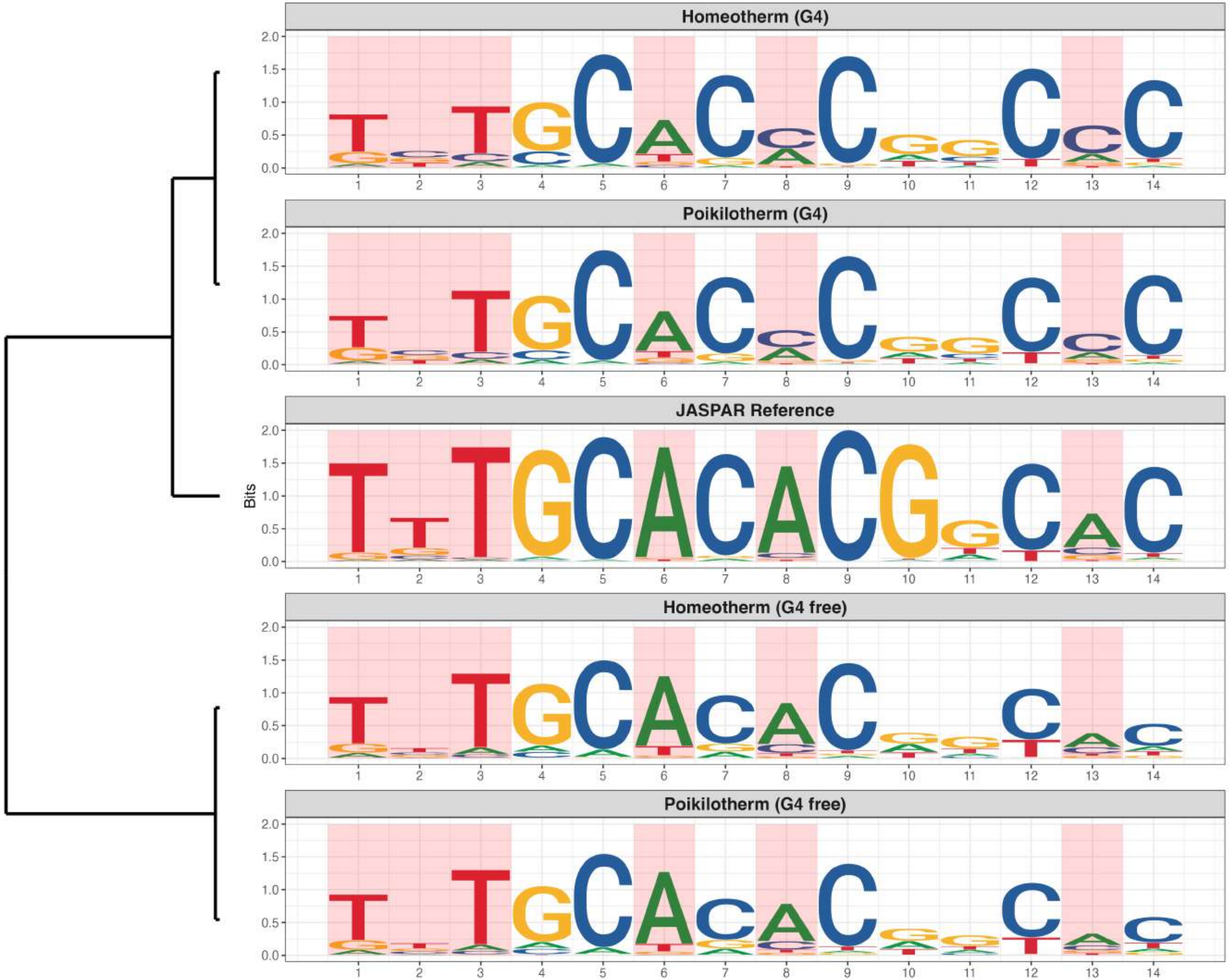

**Genome : Highly Enriched in G4s**

*MA1100.3 - ASCL1*

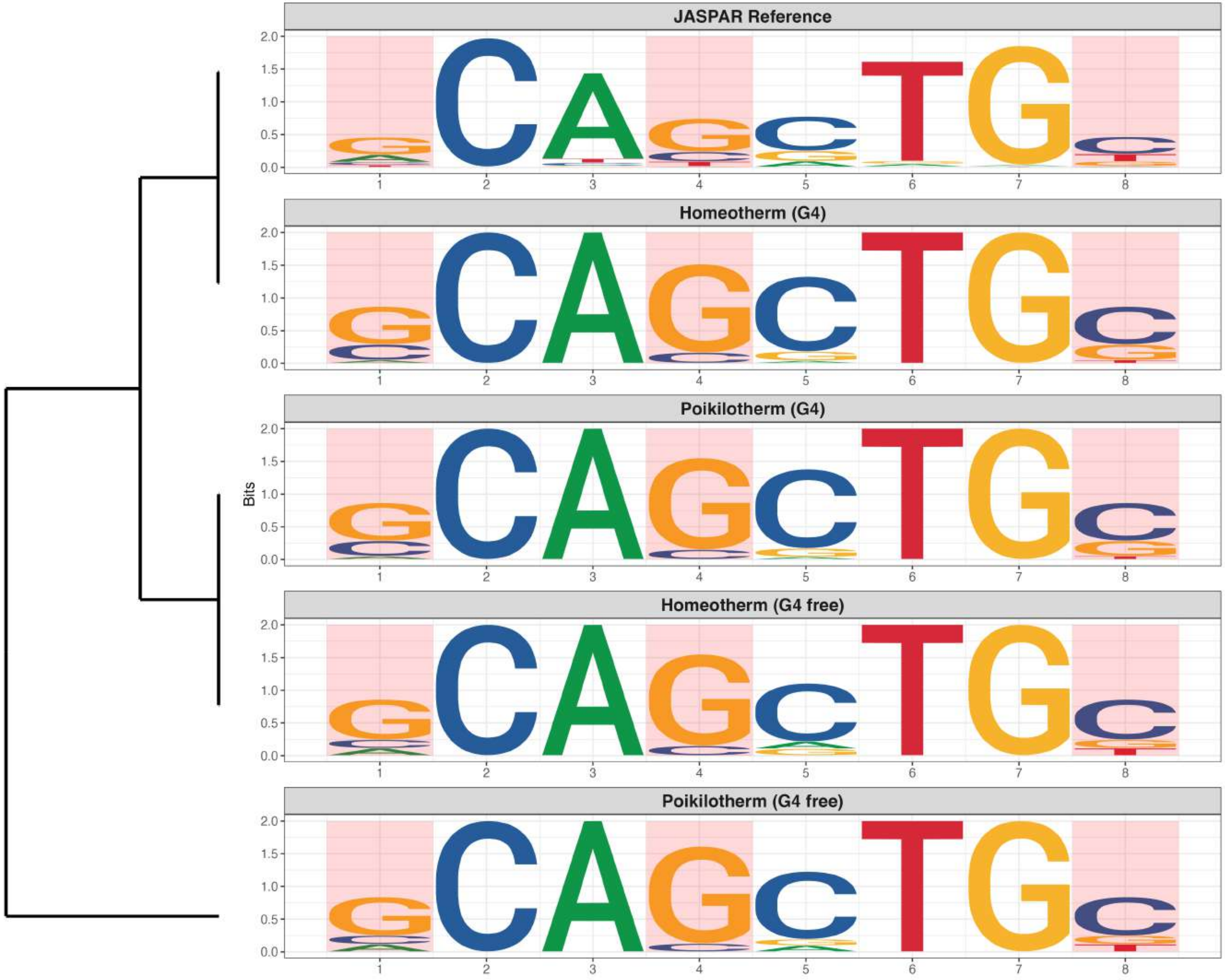

### Genome : Highly Enriched in G4s

*MA1125.2 - ZNF384*

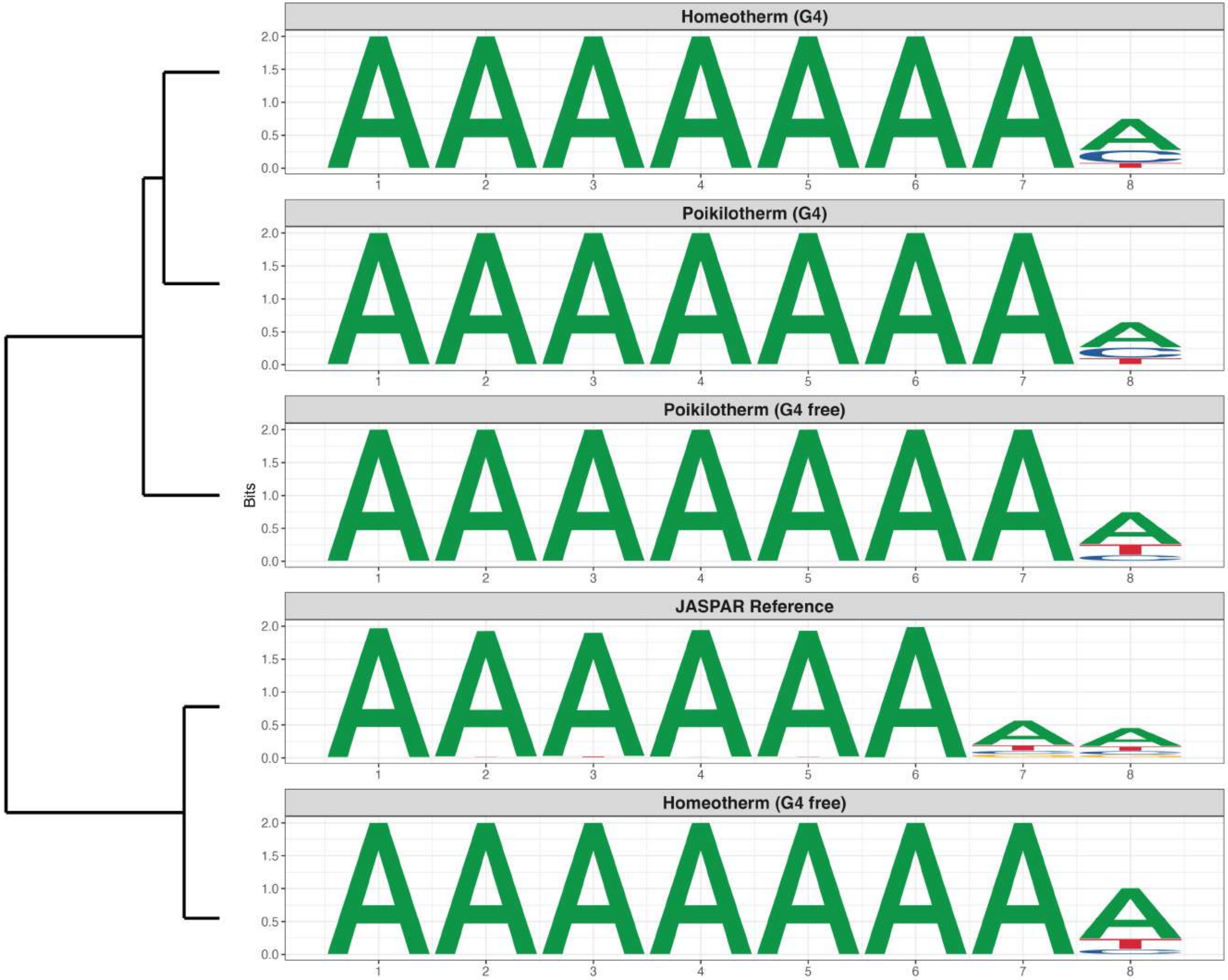

**Genome : Highly Enriched in G4s**

*MA1472.3 - Bhlha15*

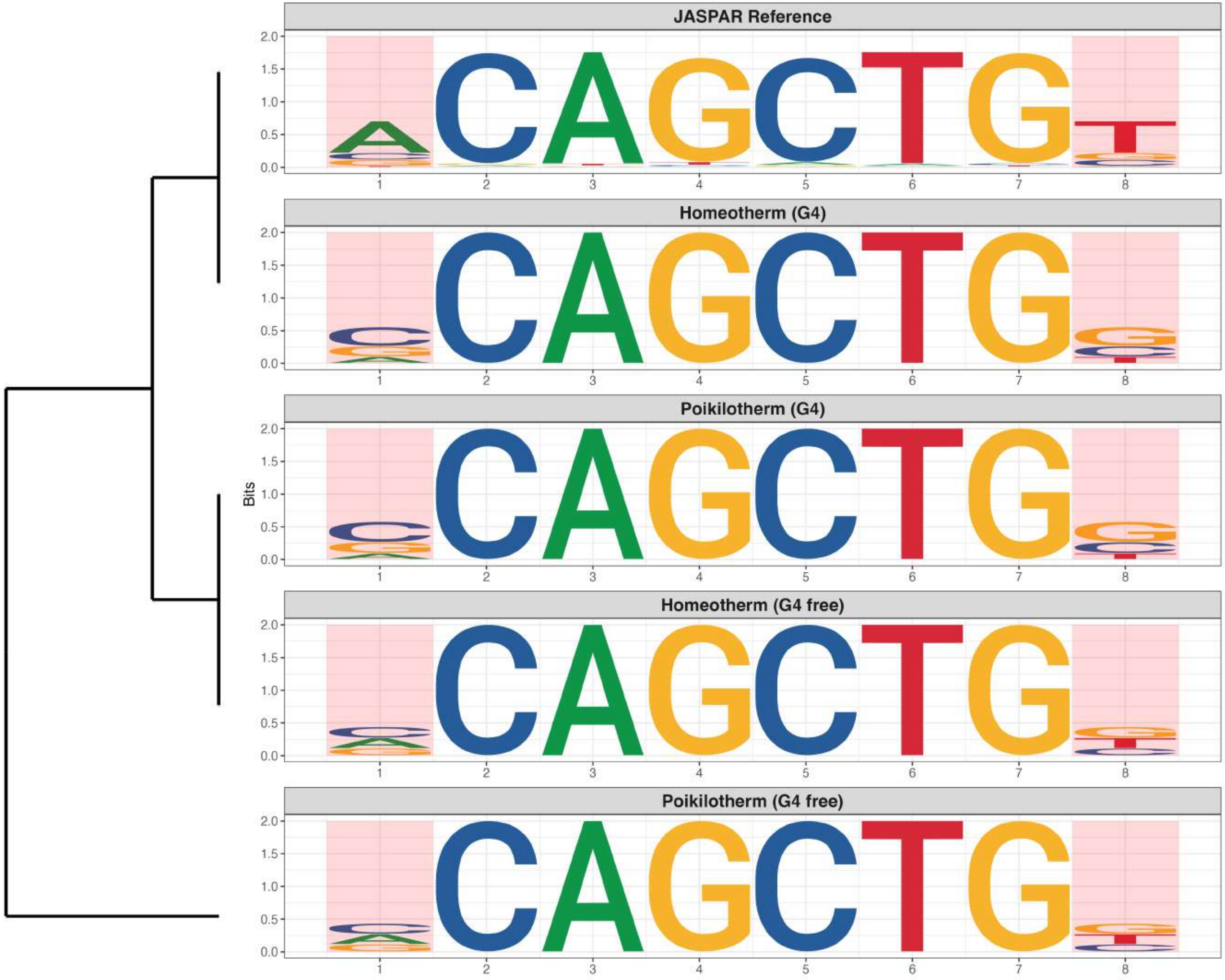

**Genome : Highly Enriched in G4s**

*MA1583.2 - ZFP57*

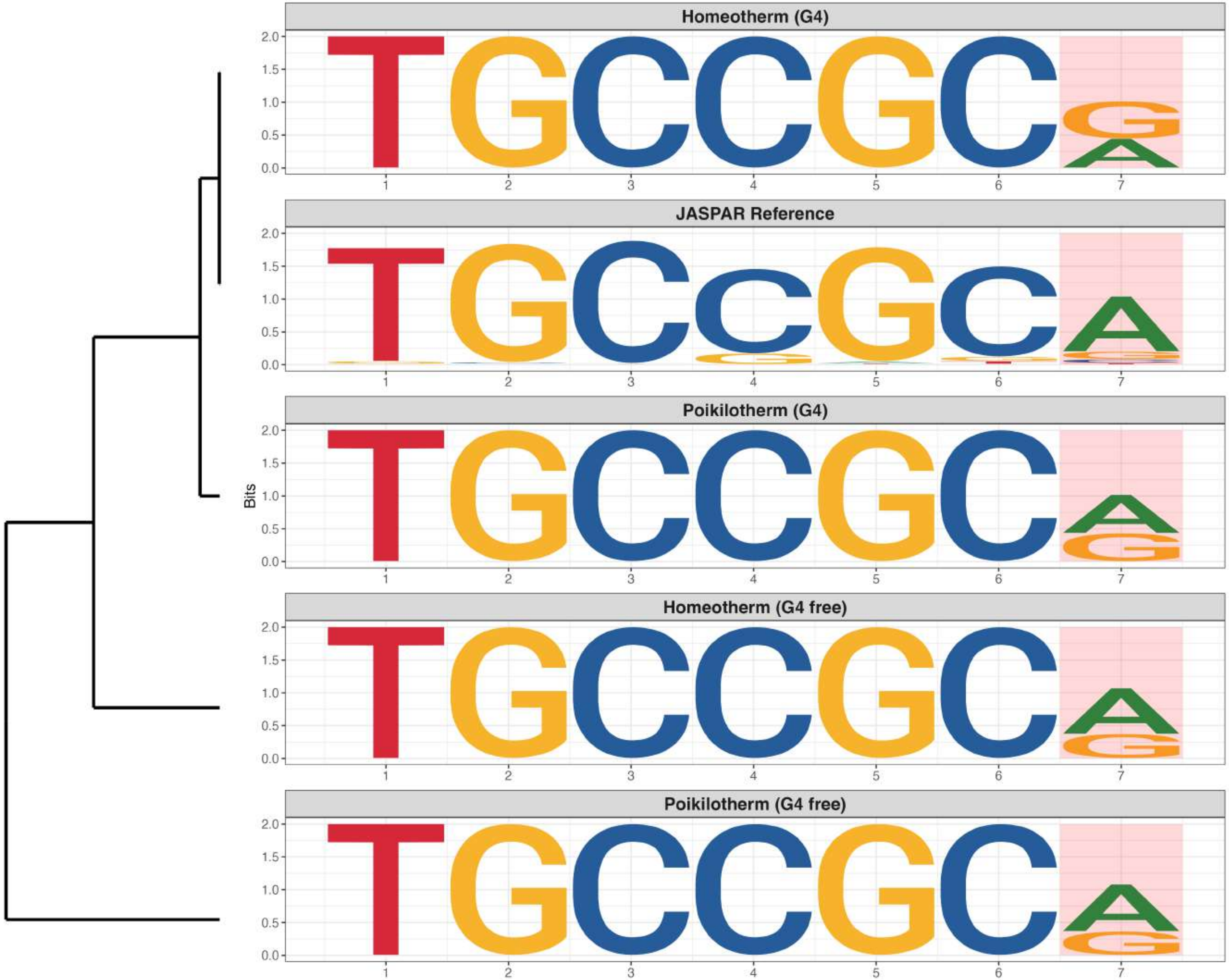

**Genome : Highly Enriched in G4s**

*MA1625.2 - Stat5b*

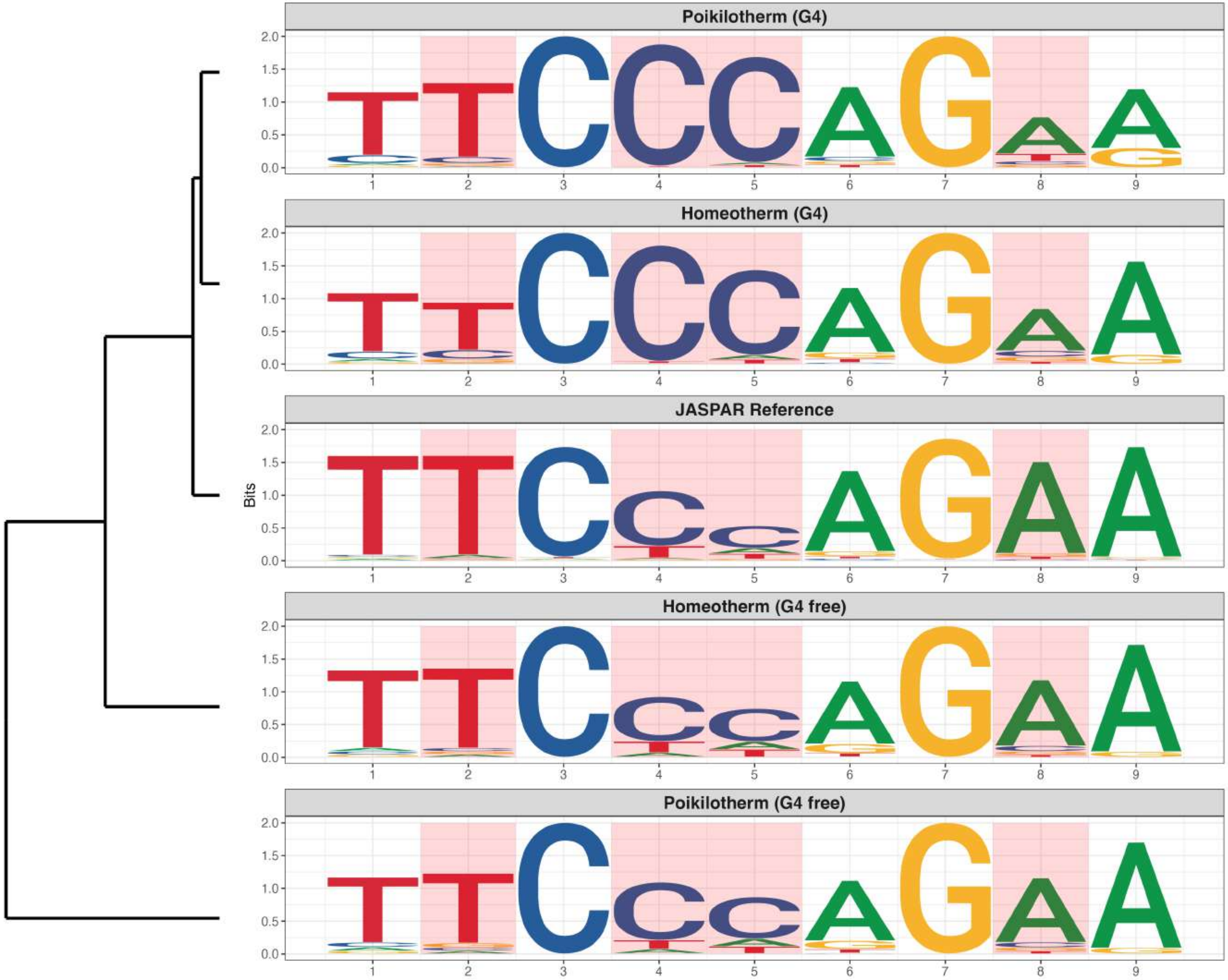

### Genome : Highly Enriched in G4s

*MA1946.2 - ETV5::FOXI1*

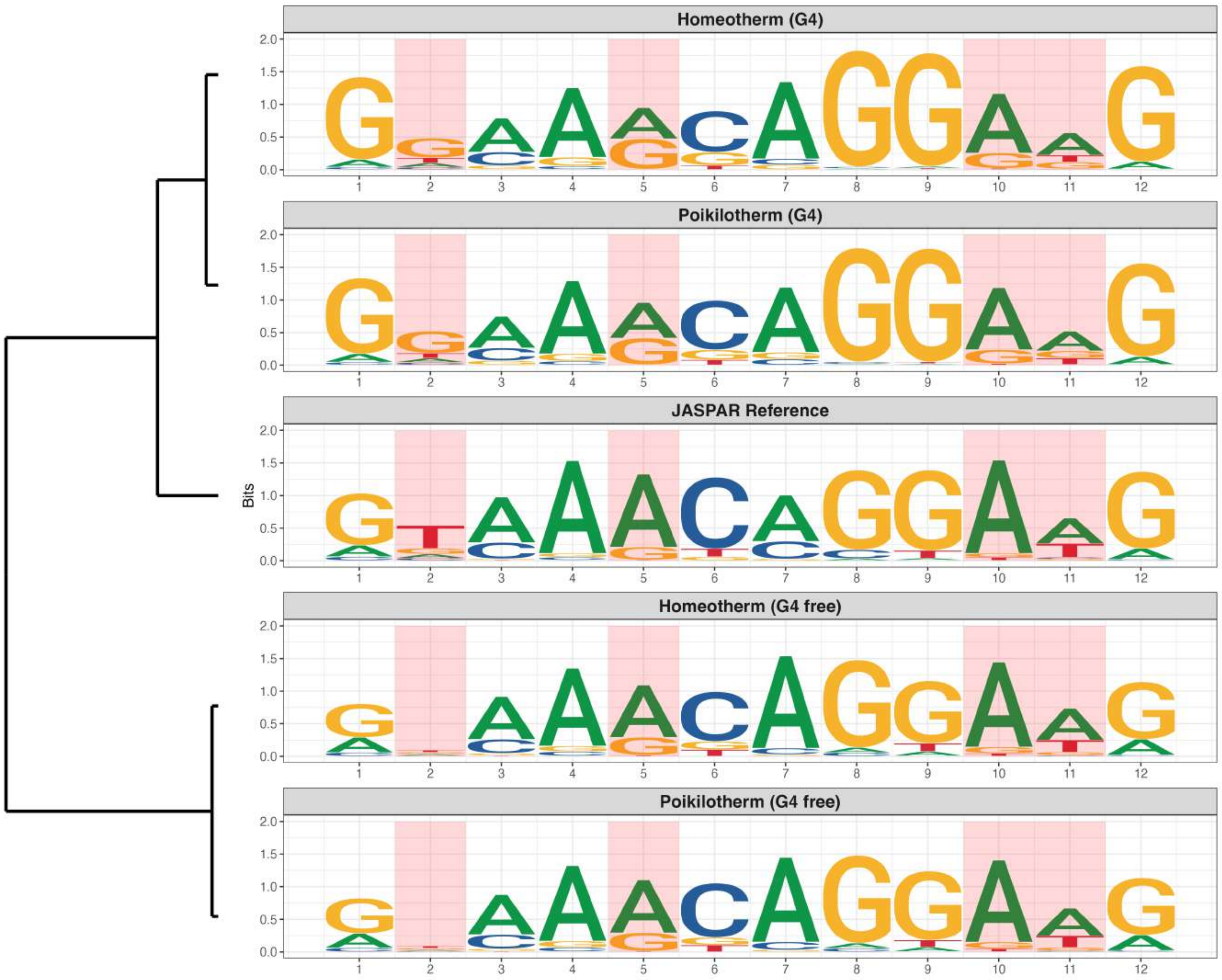

### Genome : Highly Enriched in G4s

*MA1967.2 - TFAP4::FLI1*

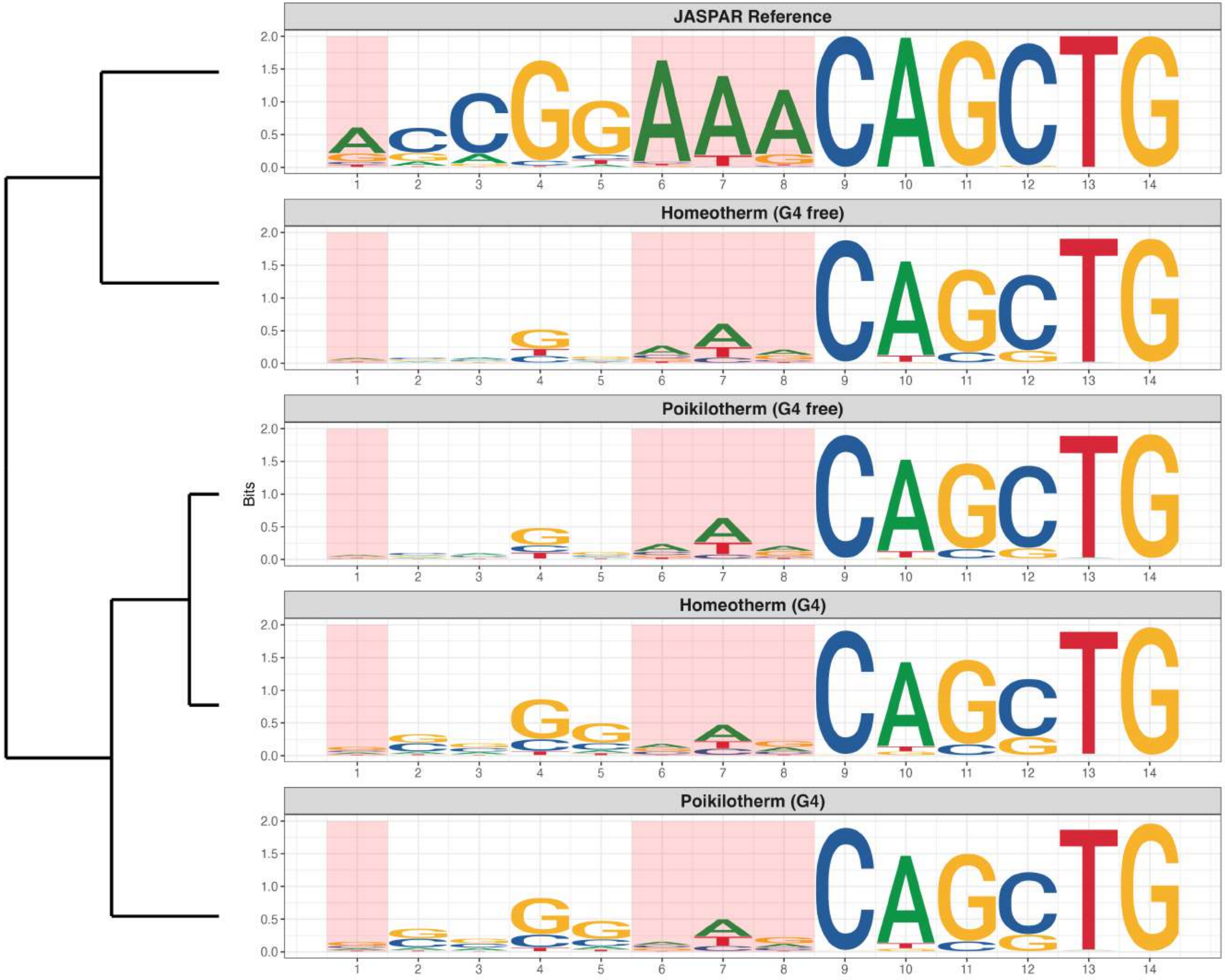

**Genome : Highly Enriched in G4s**

*MA1983.2 - ZNF582*

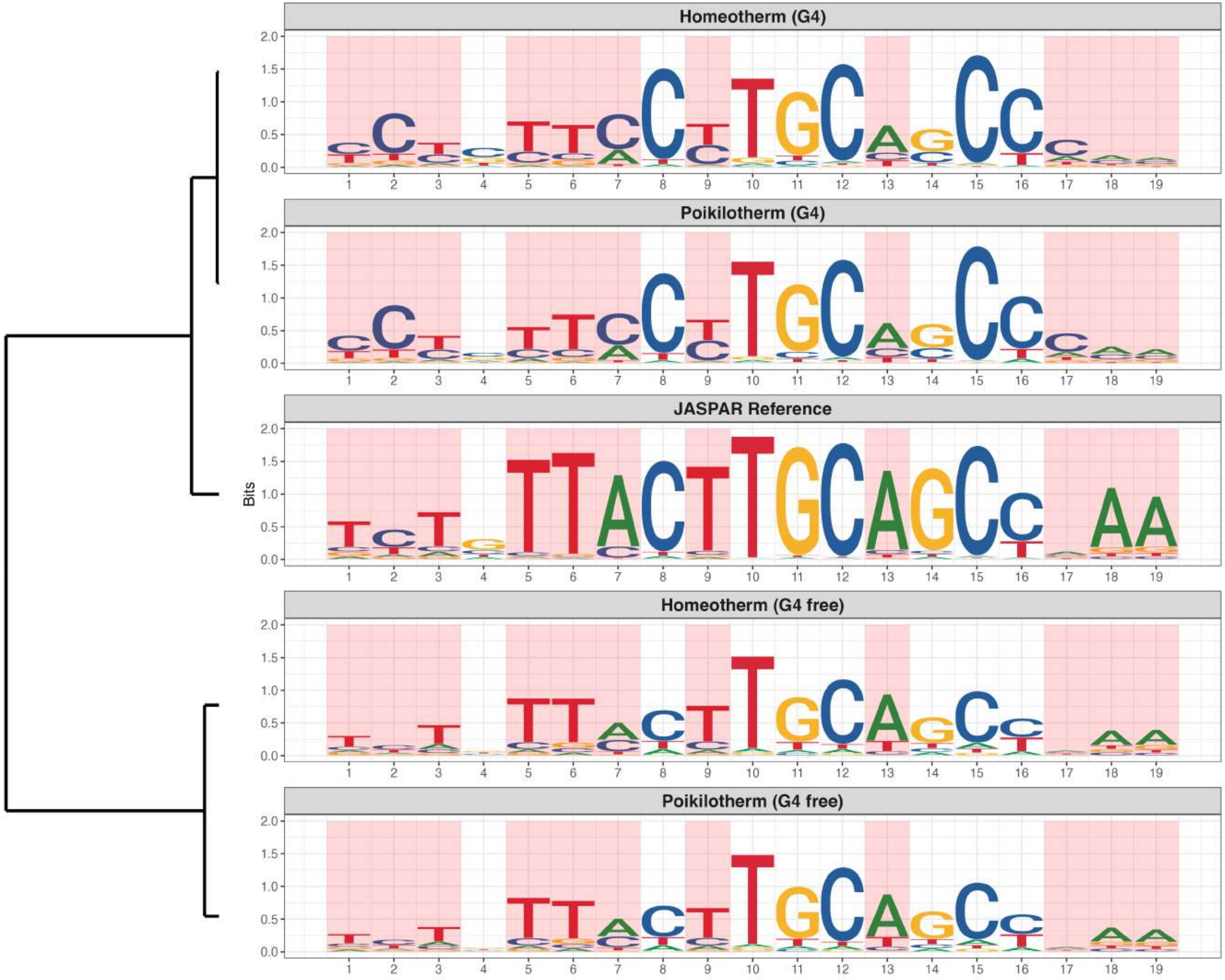

### Genome : Highly Enriched in G4s

*MA2094.1 - PAX8*

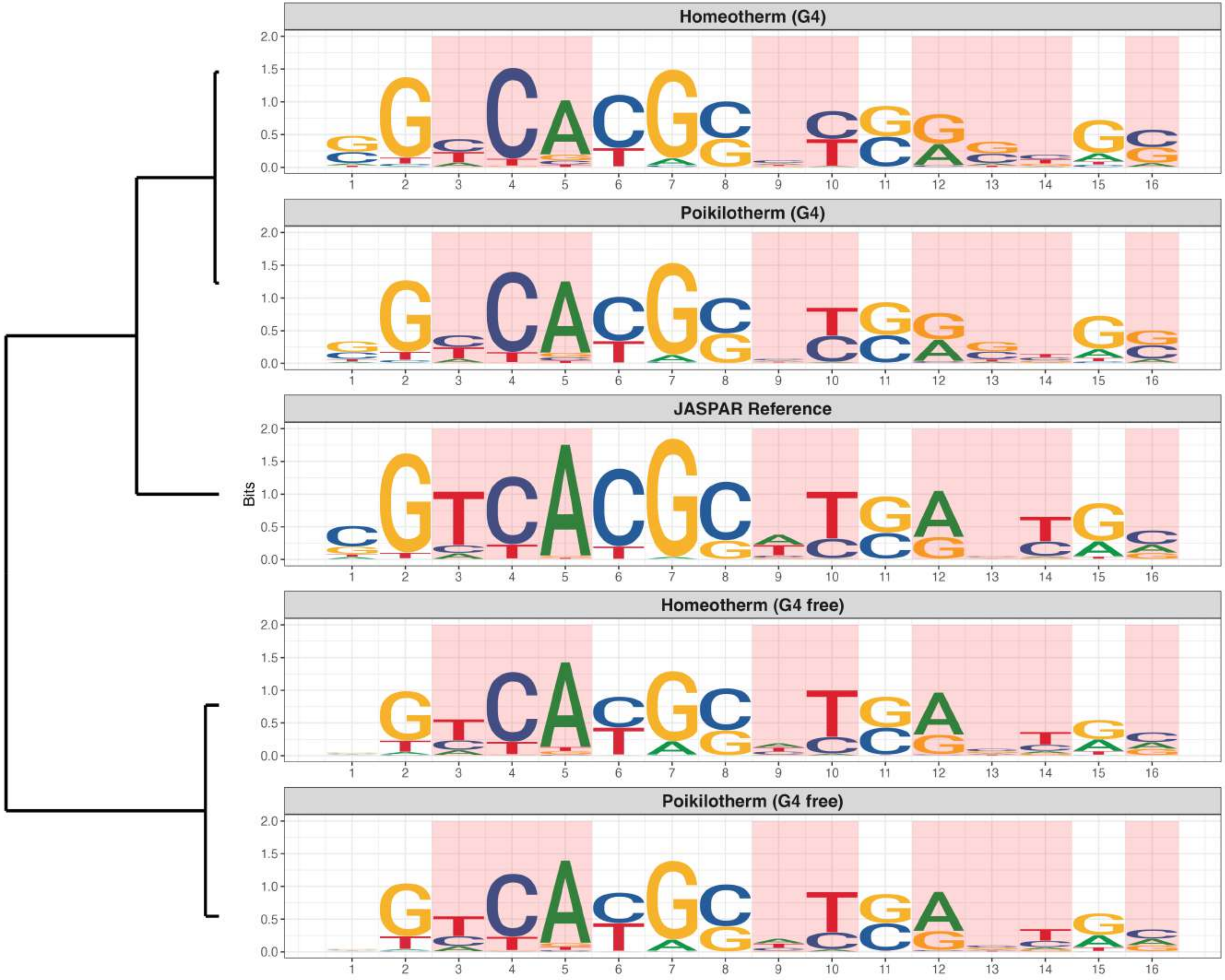

### Genome : Highly Enriched in G4s

*MA2490.1 - ZFP3*

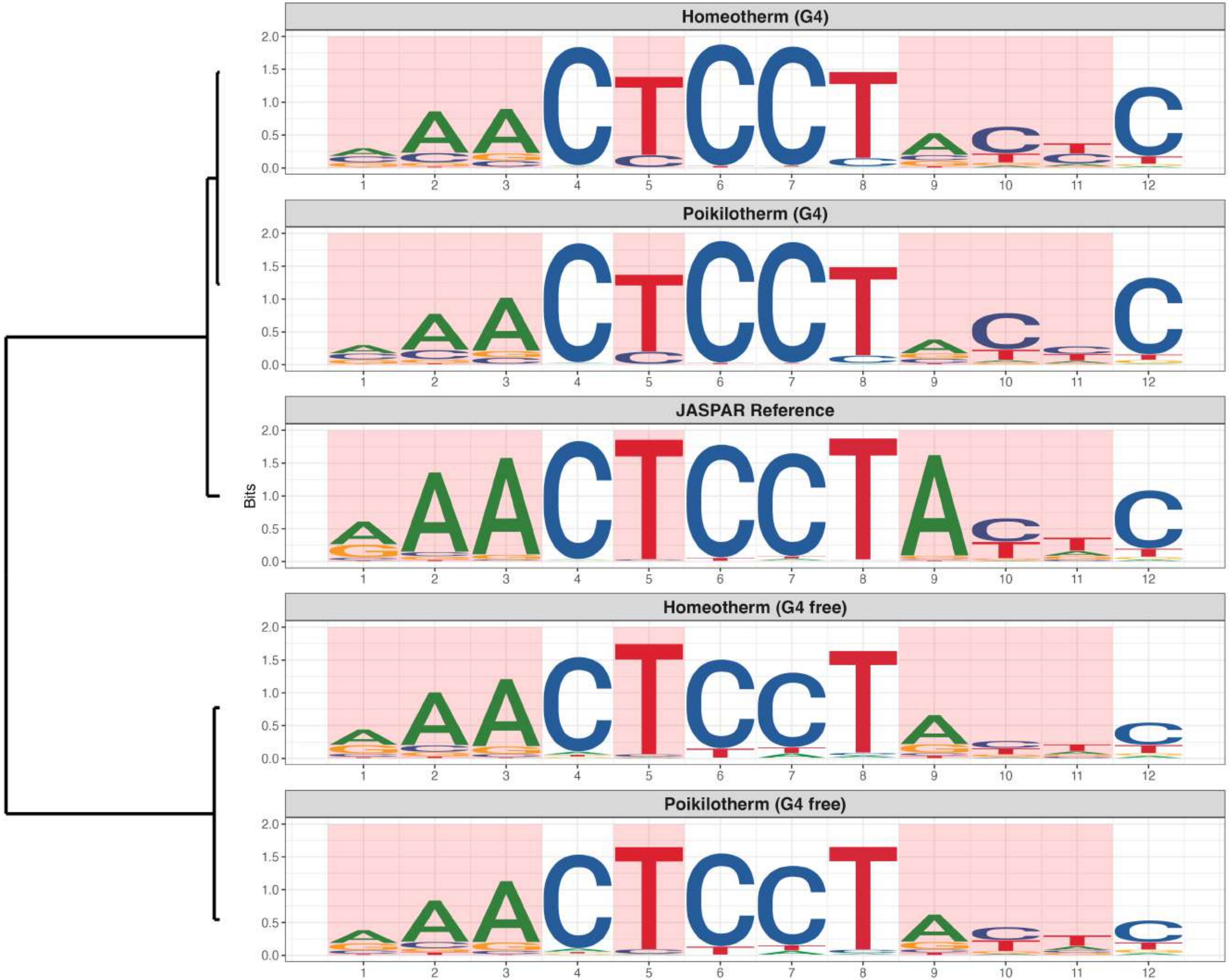

**Genome : Highly Enriched in G4s**

*MA2503.1 - Banp*

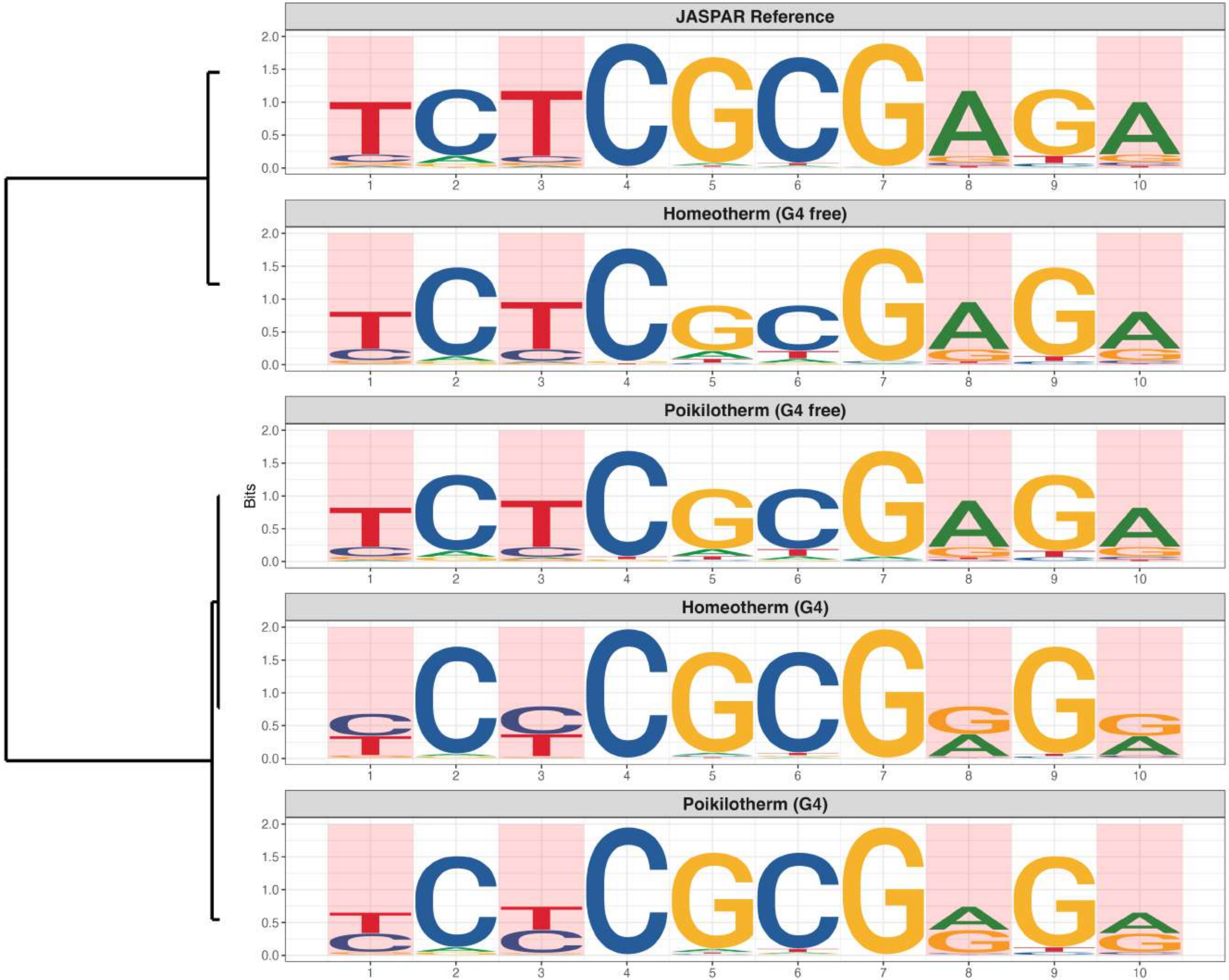

### Genome : Highly Enriched in G4s

*MA2585.1 - ZNF500*

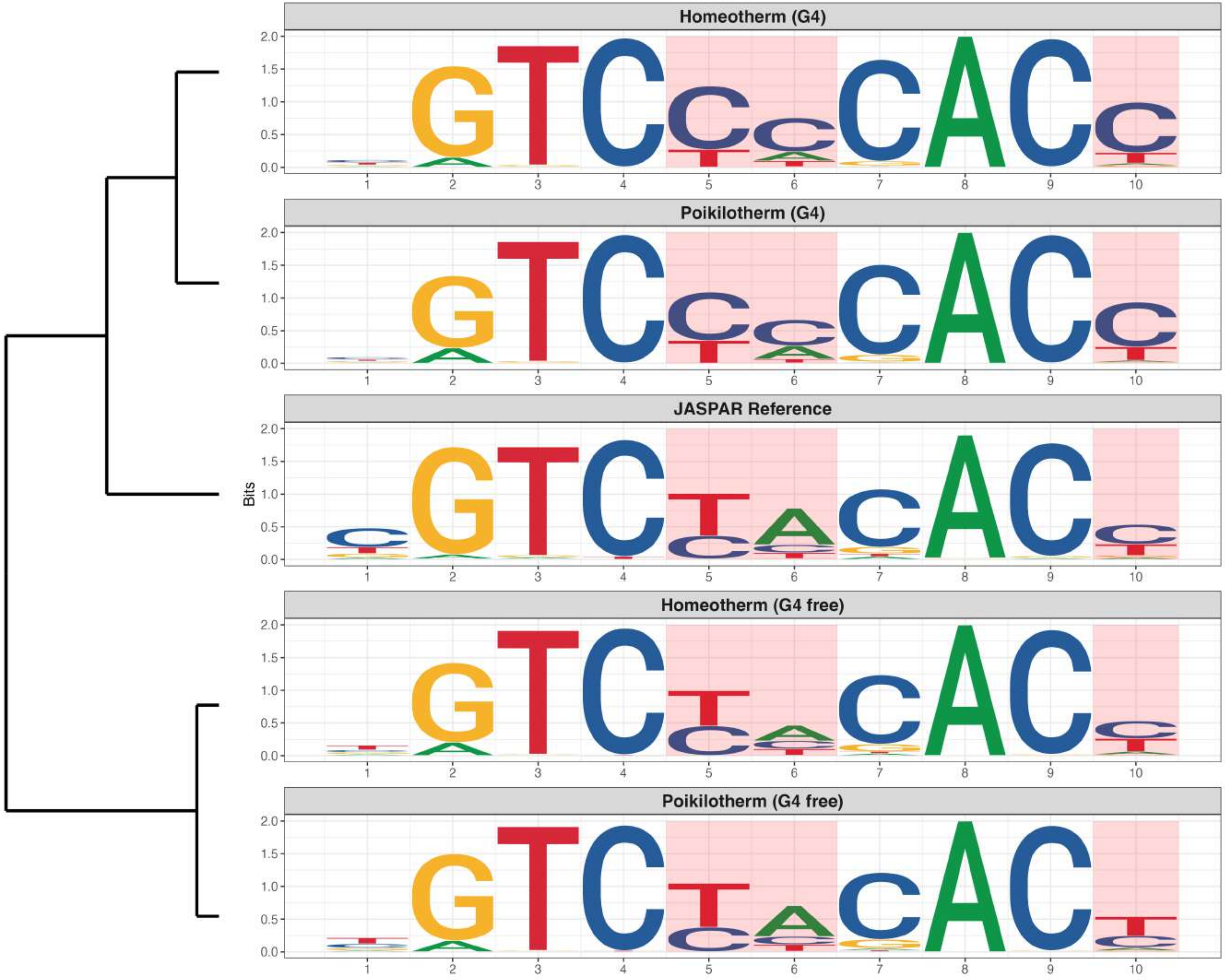

**Genome : Moderately Enriched in G4s**

*MA0028.3 - ELK1*

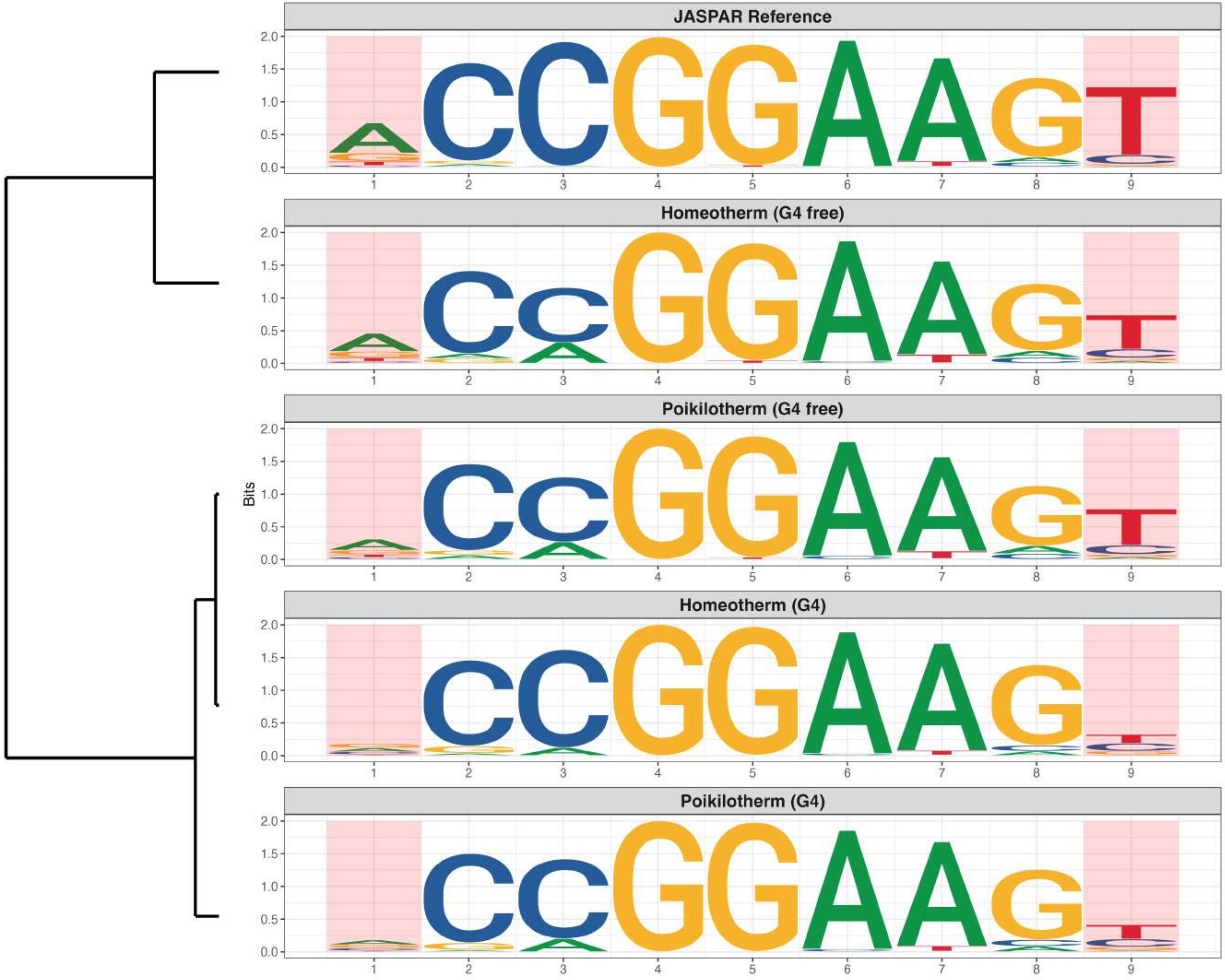

**Genome : Moderately Enriched in G4s**

*MA0104.5 - MYCN*

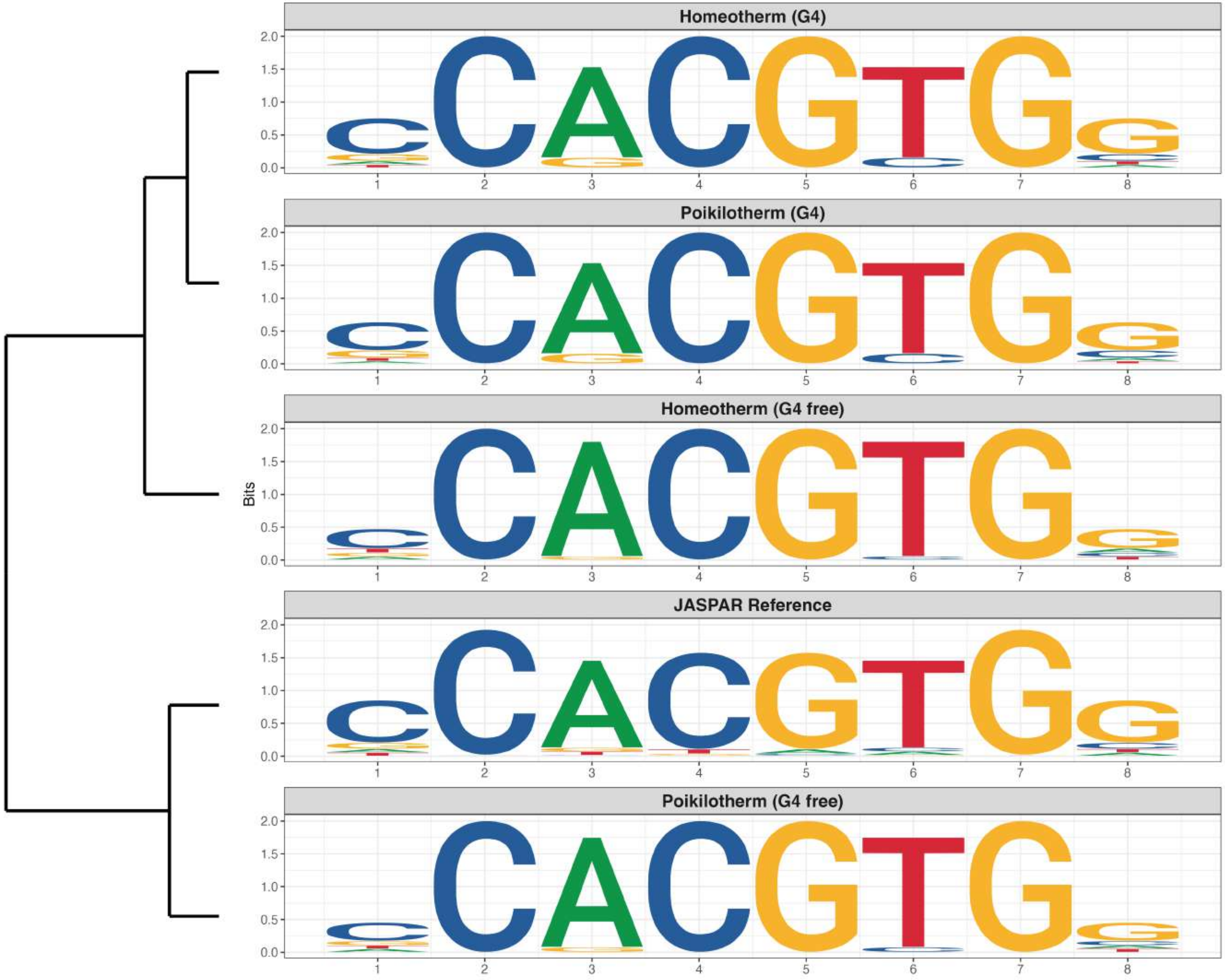

**Genome : Moderately Enriched in G4s**

*MA0131.3 - HINFP*

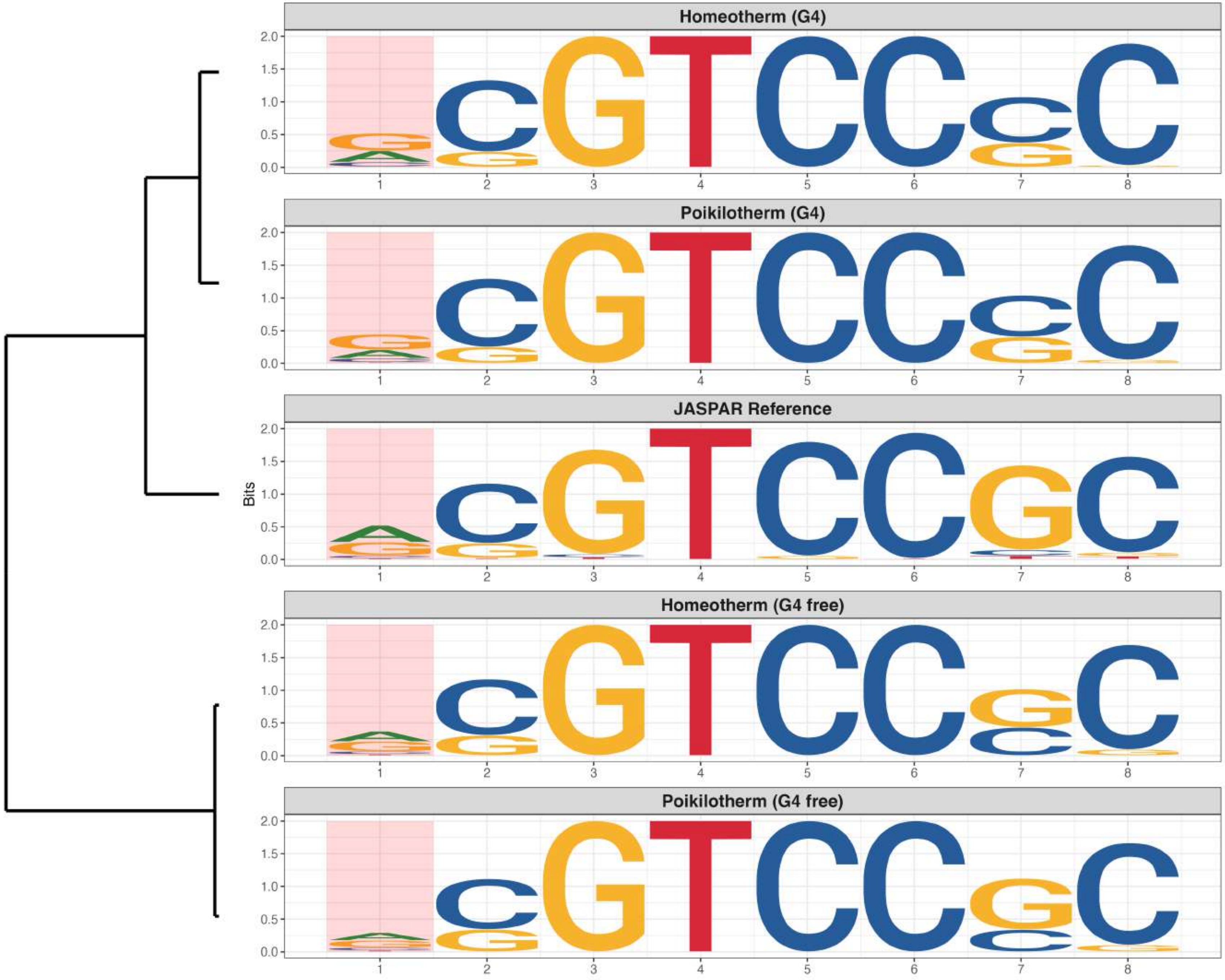

**Genome : Moderately Enriched in G4s**

*MA0478.2 - FOSL2*

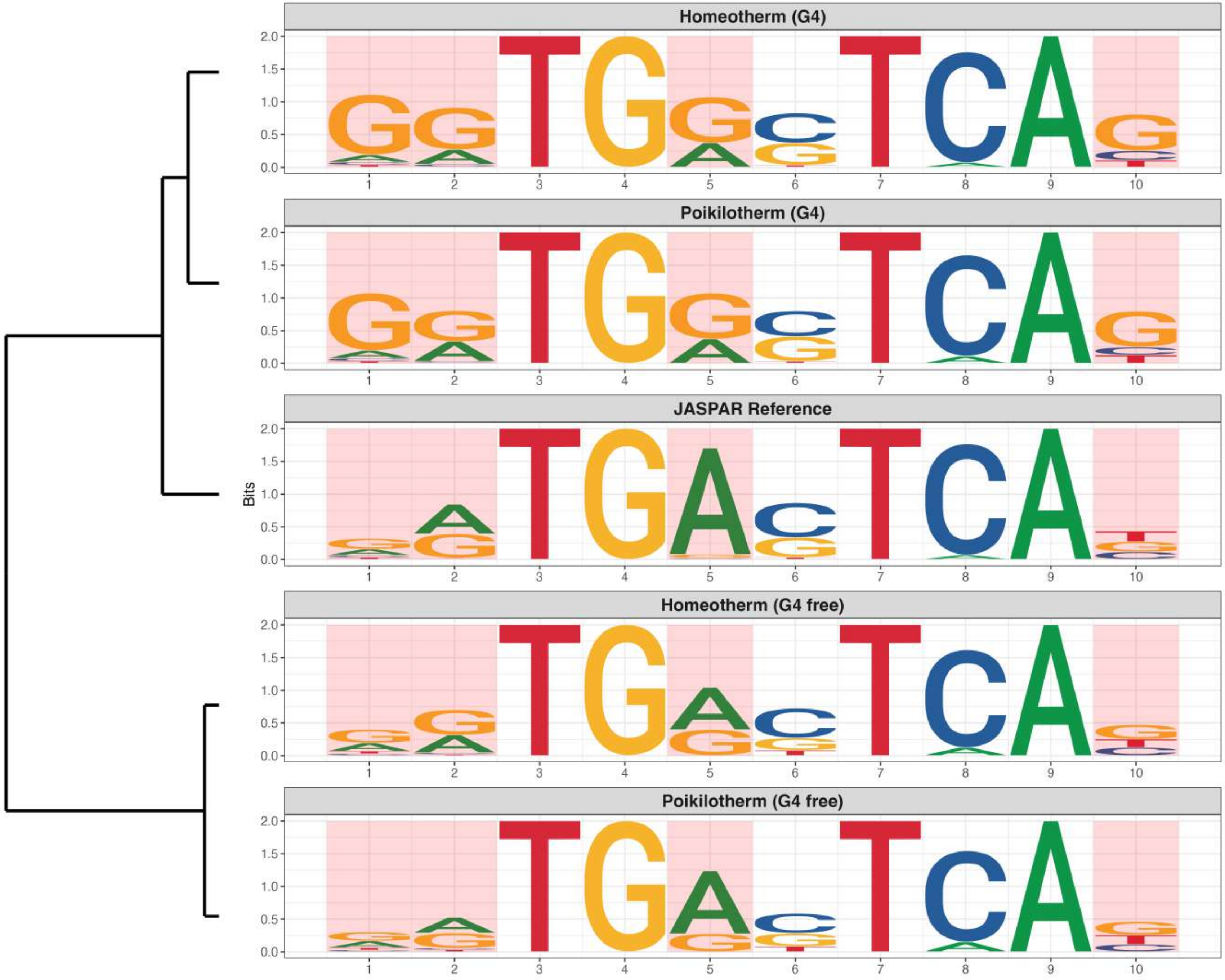

**Genome : Moderately Enriched in G4s**

*MA0499.3 - MYOD1*

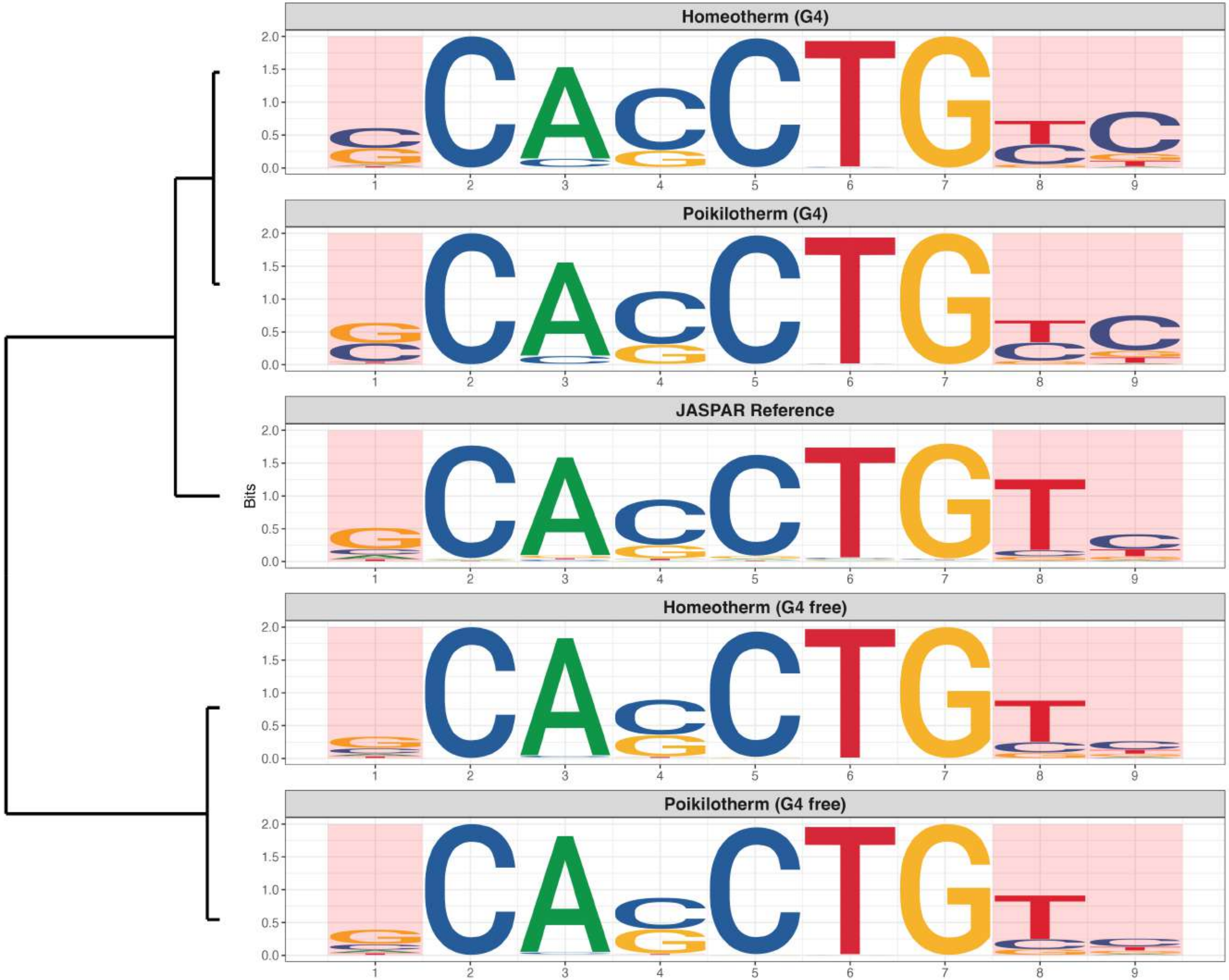

### Genome : Moderately Enriched in G4s

*MA0527.2 - ZBTB33*

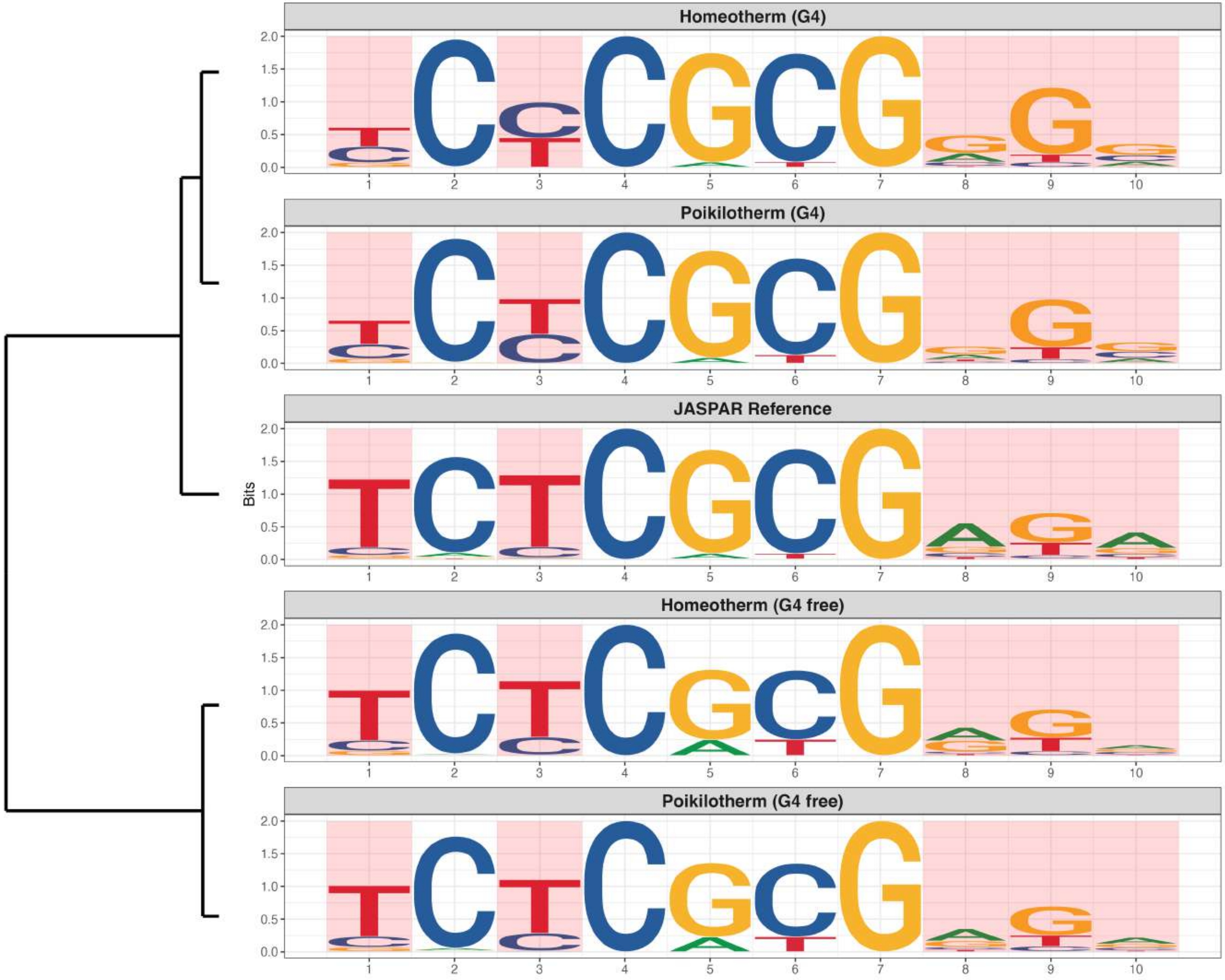

### Genome : Moderately Enriched in G4s

*MA0595.1 - SREBF1*

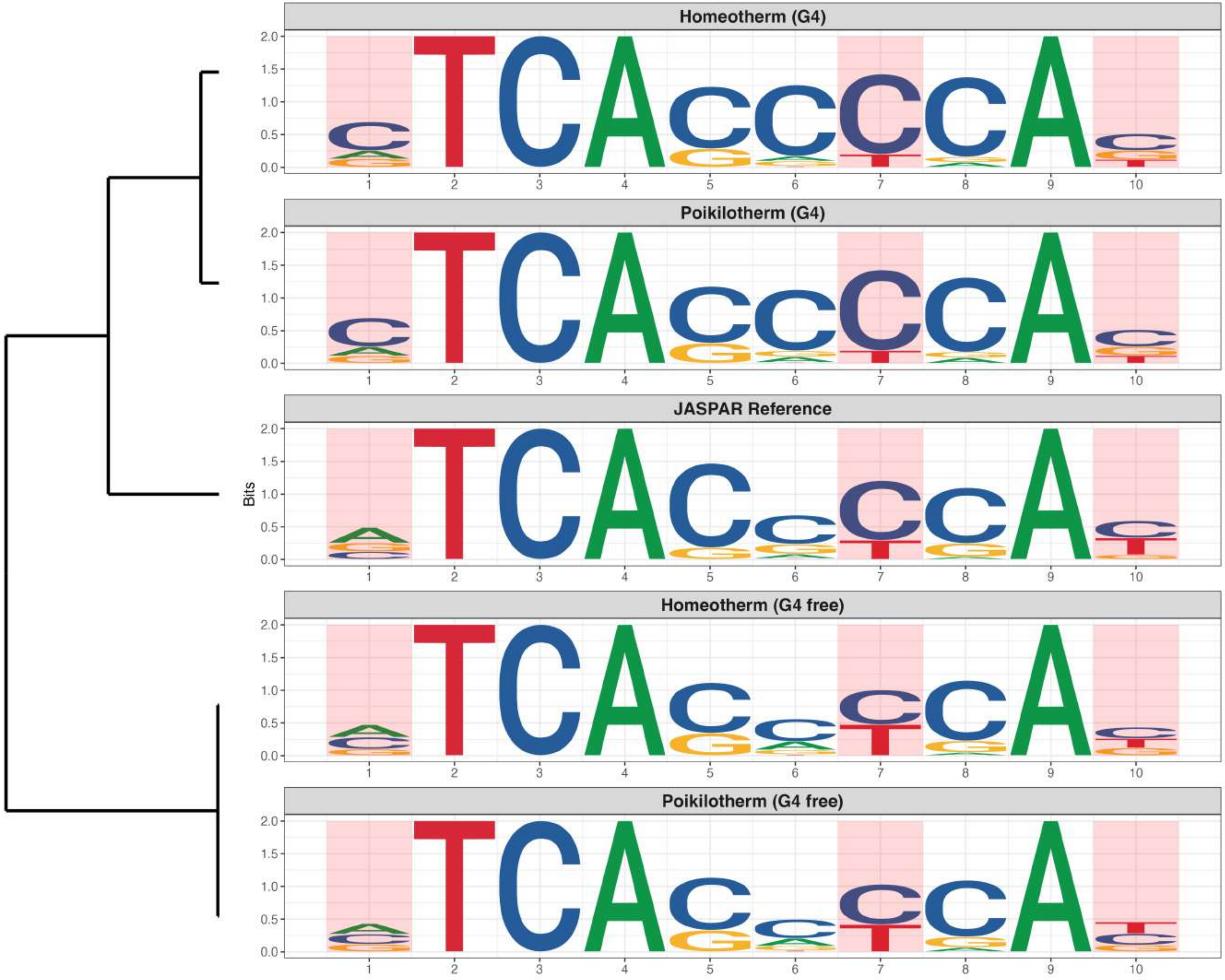

### Genome : Moderately Enriched in G4s

*MA0632.3 - TCFL5*

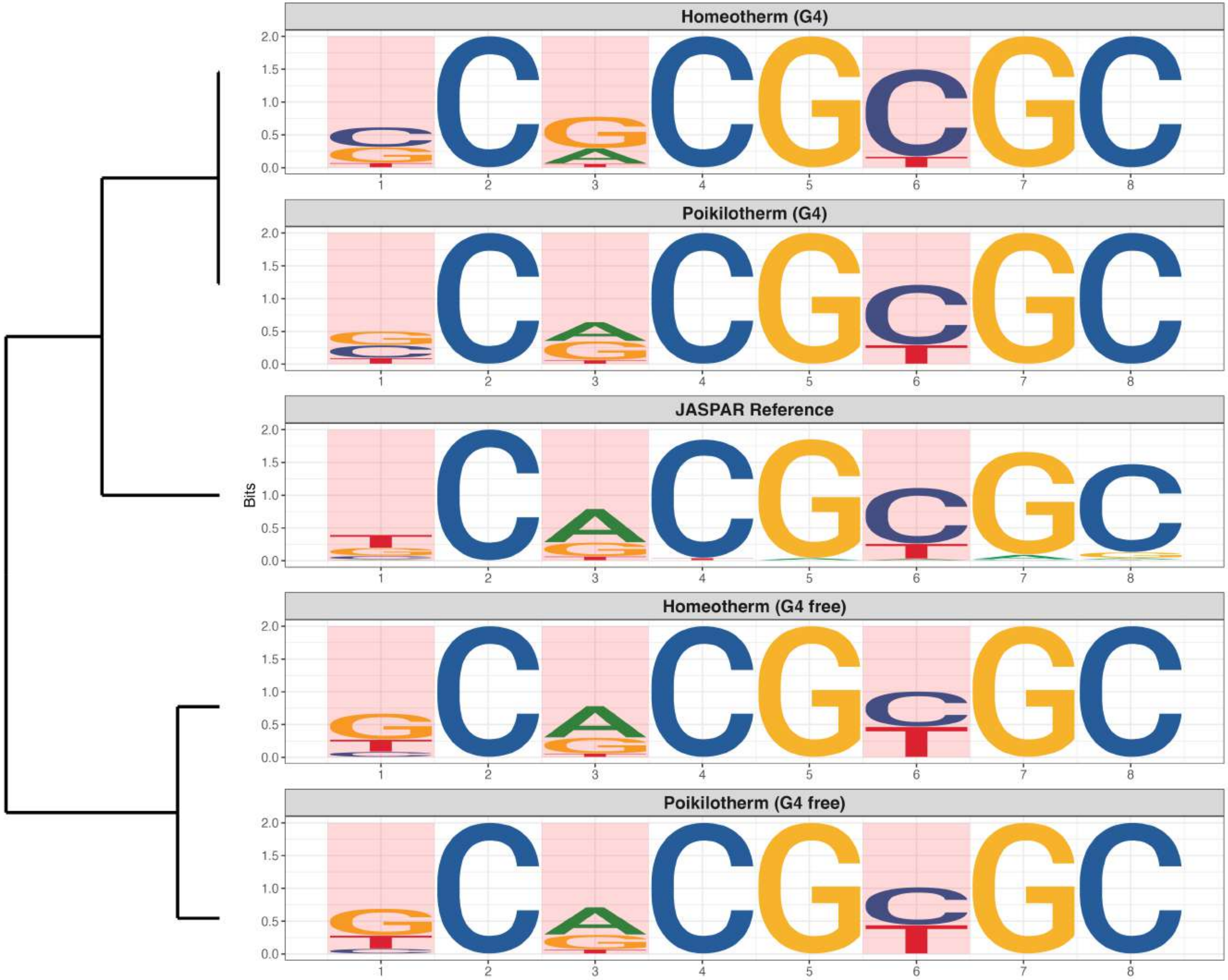

### Genome : Moderately Enriched in G4s

*MA0693.4 - Vdr*

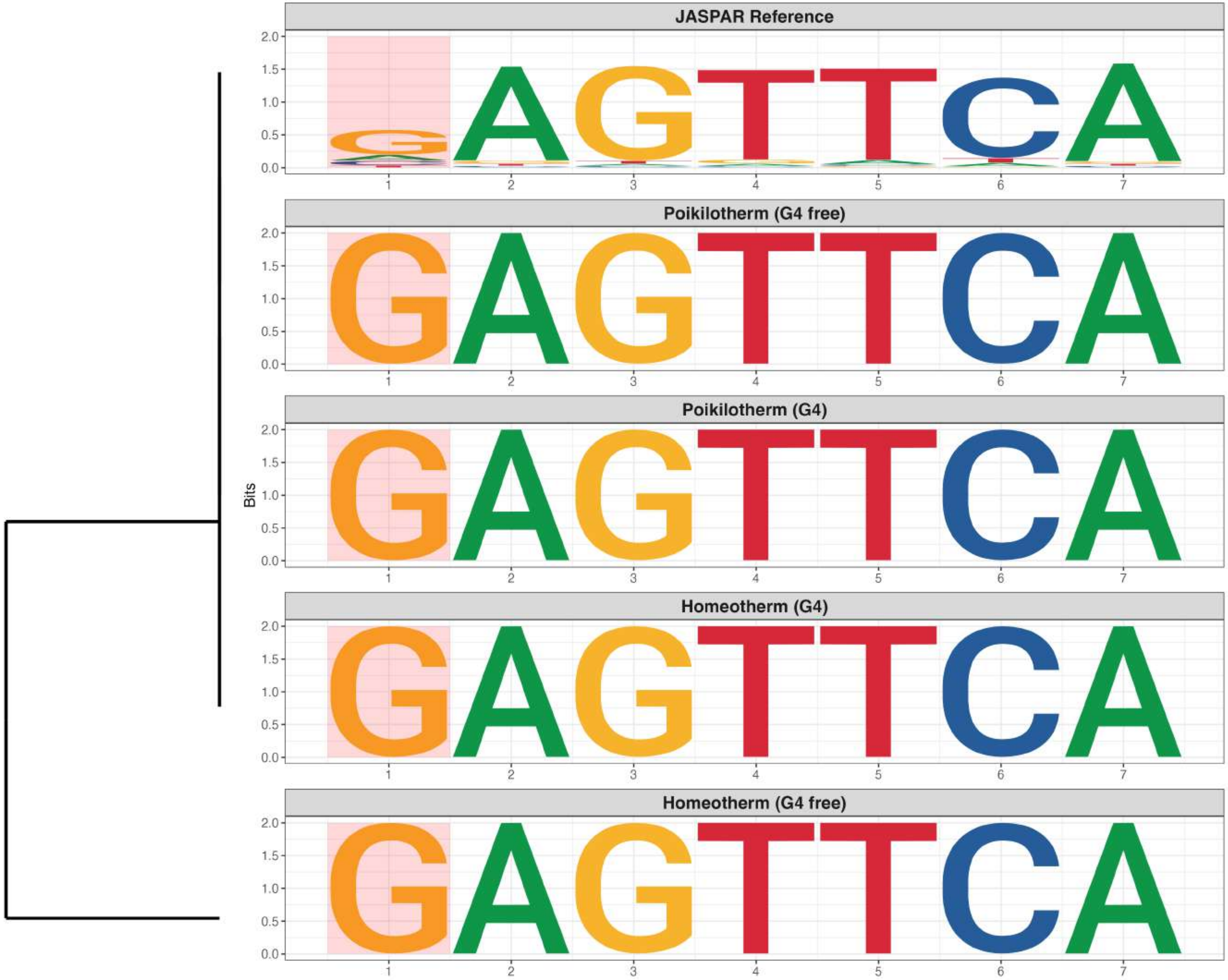

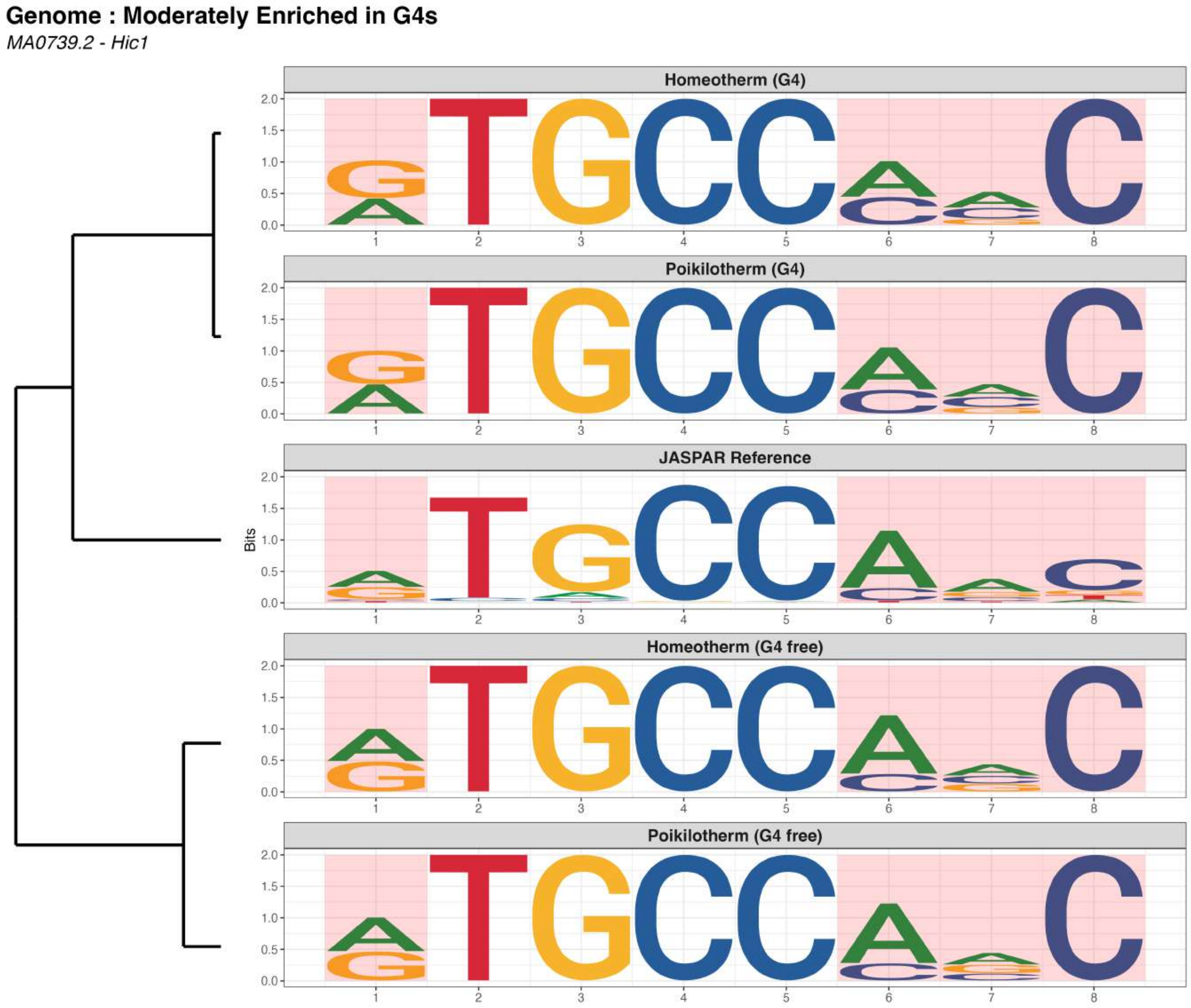
Genome : Moderately Enriched in G4s
MA0739.2 - Hic1
Homeotherm (G4)
Poikilotherm (G4)
JASPAR Reference
Homeotherm (G4 free)
Poikilotherm (G4 free)
Bits
2.0
1.5
1.0
0.5
0.0
1
2
3
4
5
6
7
8

### Genome : Moderately Enriched in G4s

*MA0765.4 - ETV5*

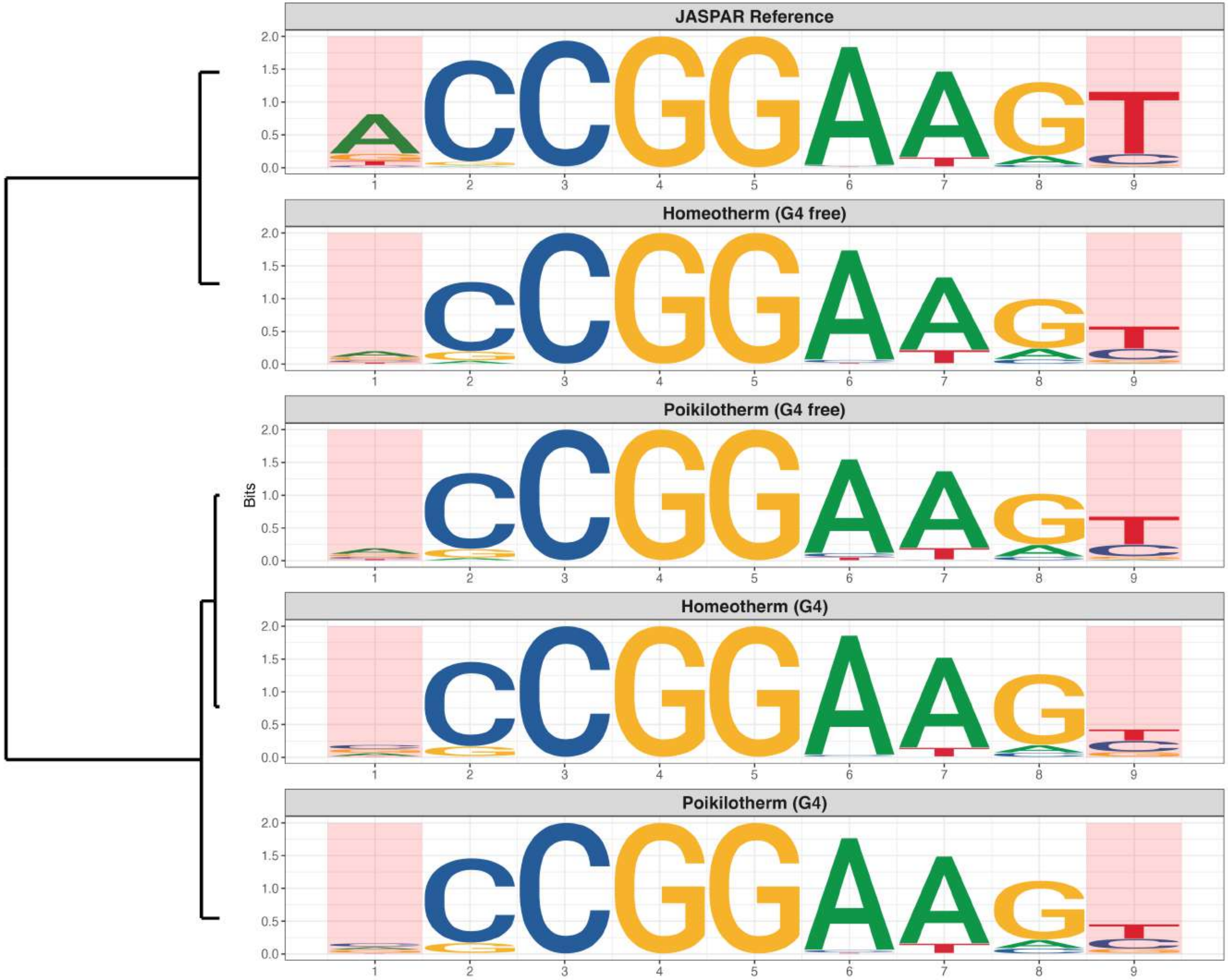

### Genome : Moderately Enriched in G4s

*MA0803.1 - TBX15*

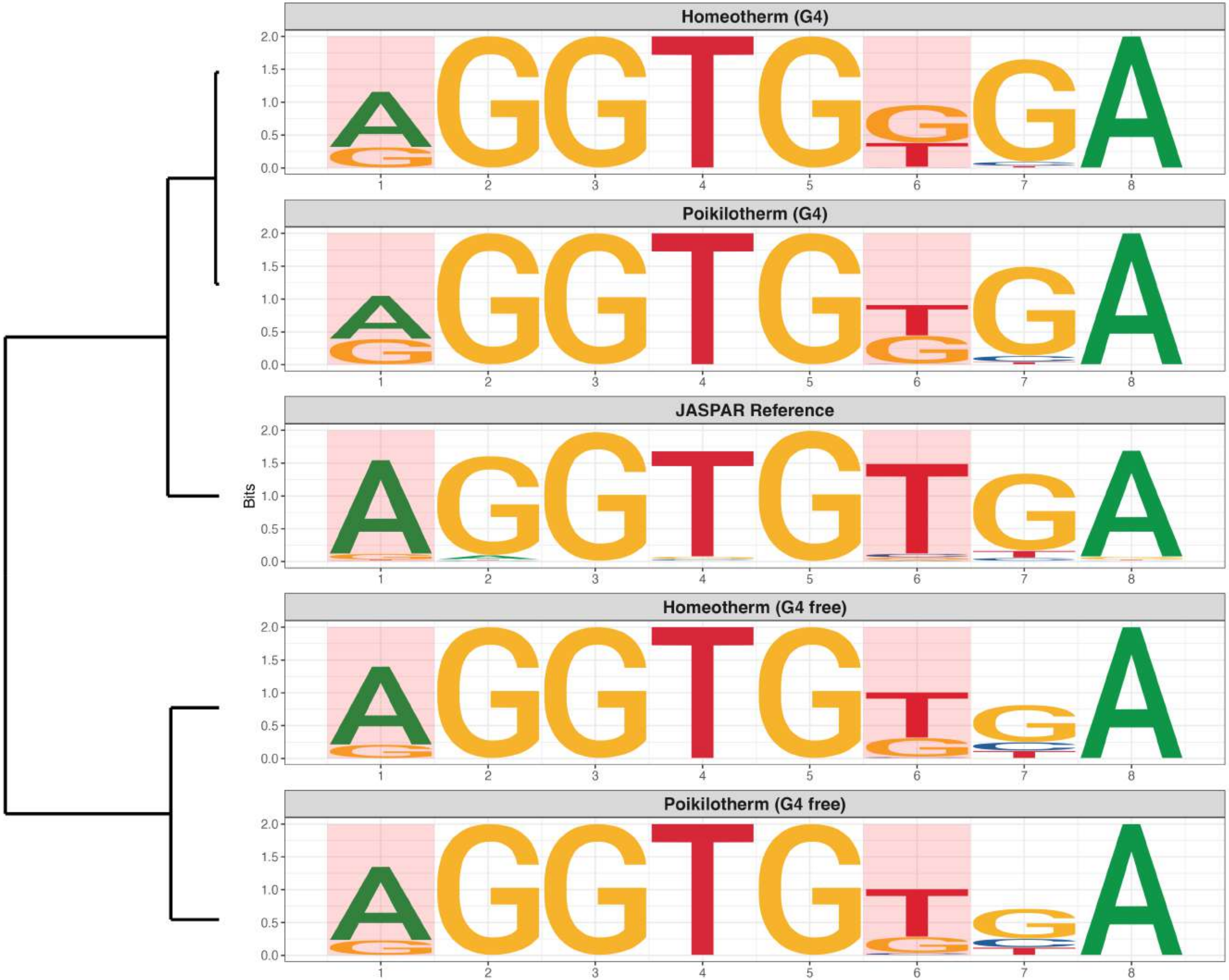

### Genome : Moderately Enriched in G4s

*MA0809.3 - TEAD4*

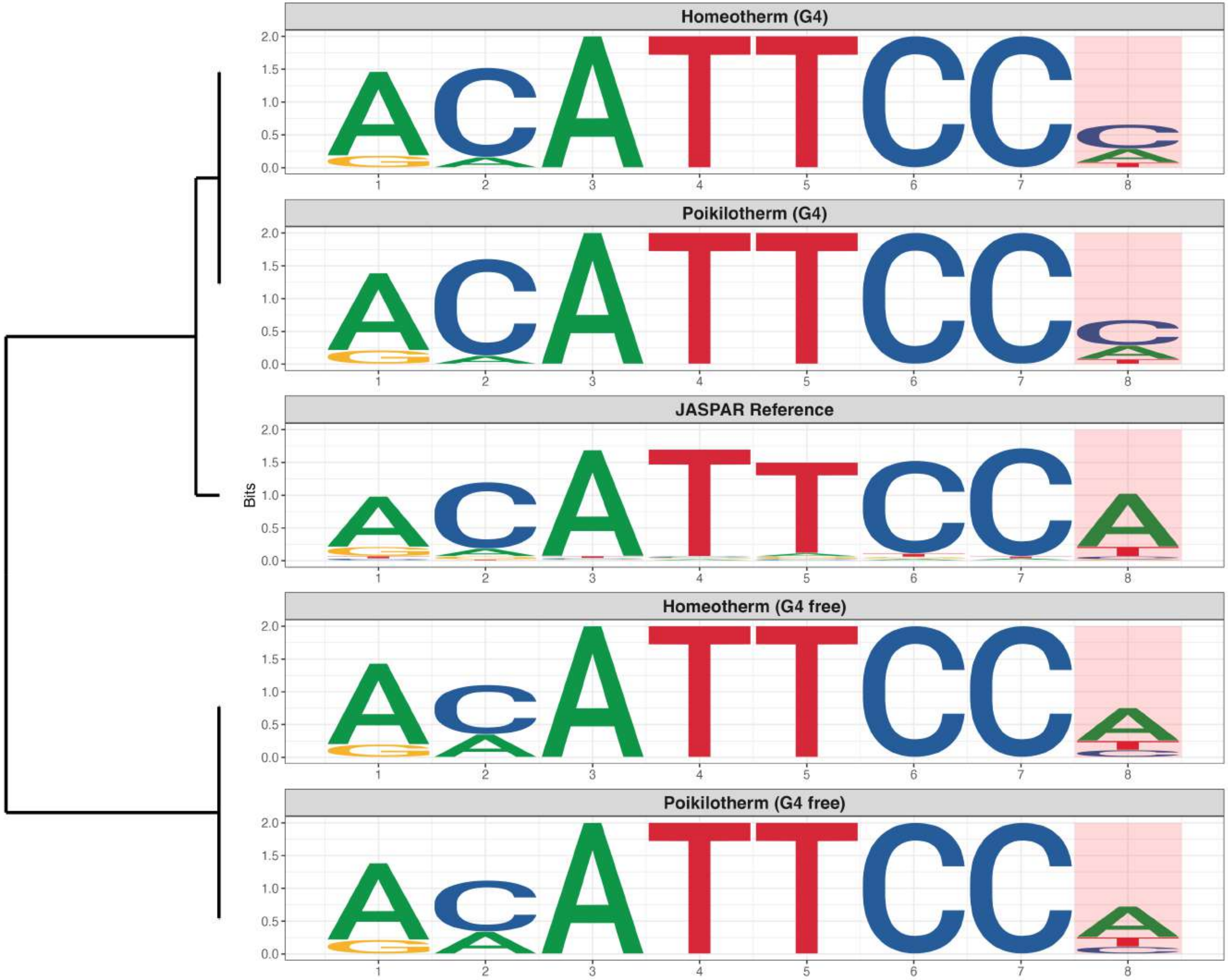

### Genome : Moderately Enriched in G4s

*MA0823.1 - HEY1*

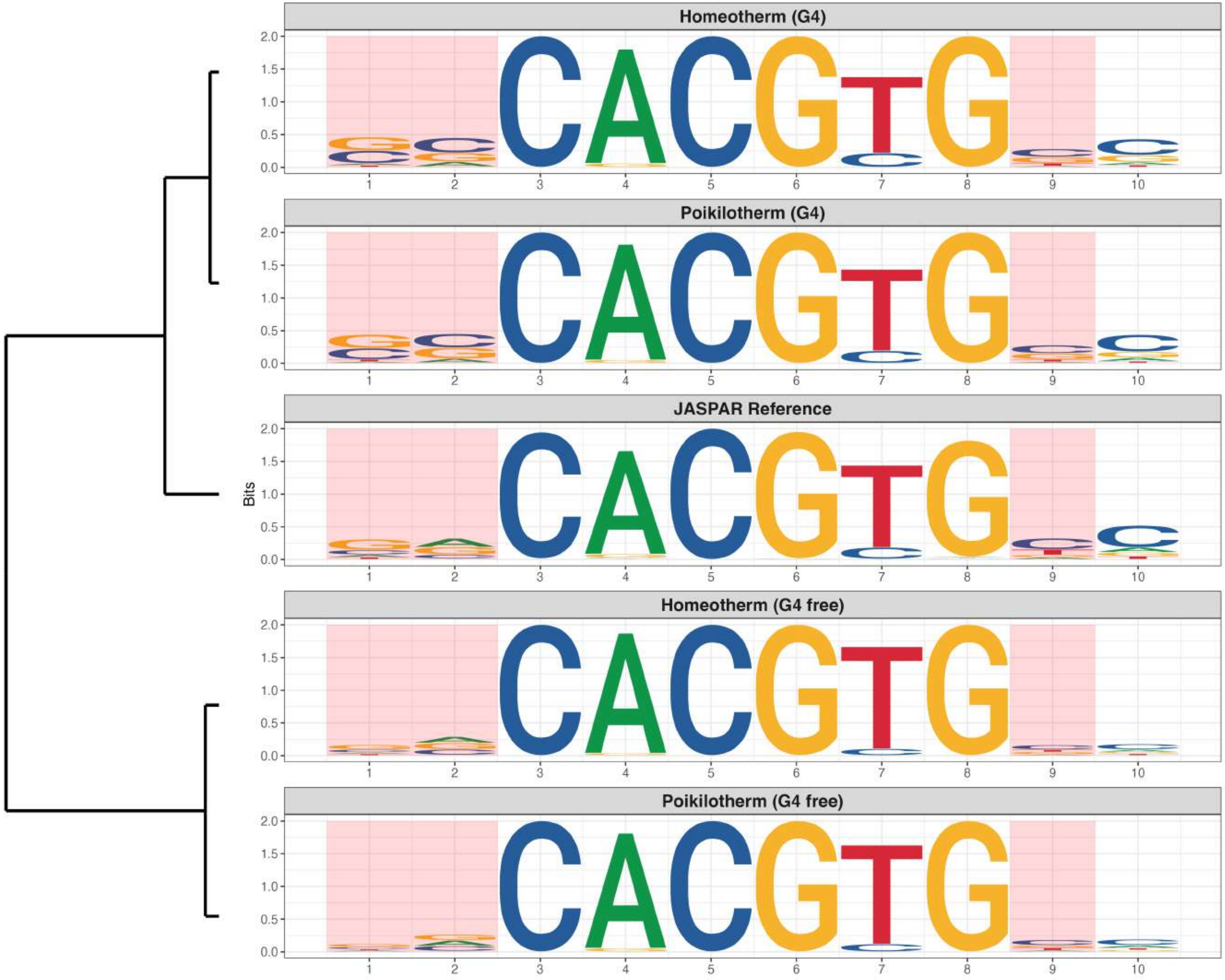

**Genome : Moderately Enriched in G4s**

*MA1099.3 - HES1*

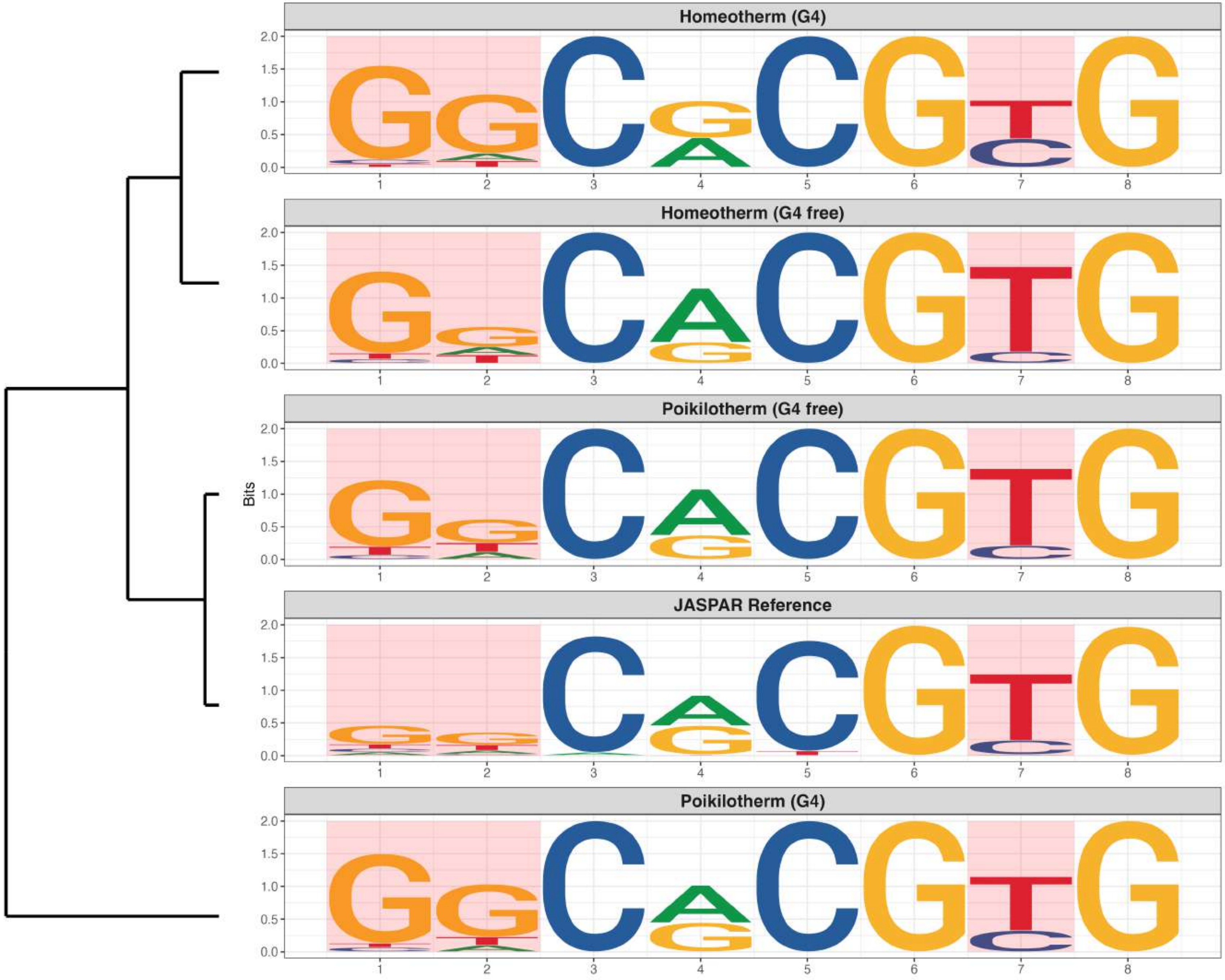

### Genome : Moderately Enriched in G4s

*MA1474.2 - CREB3L4*

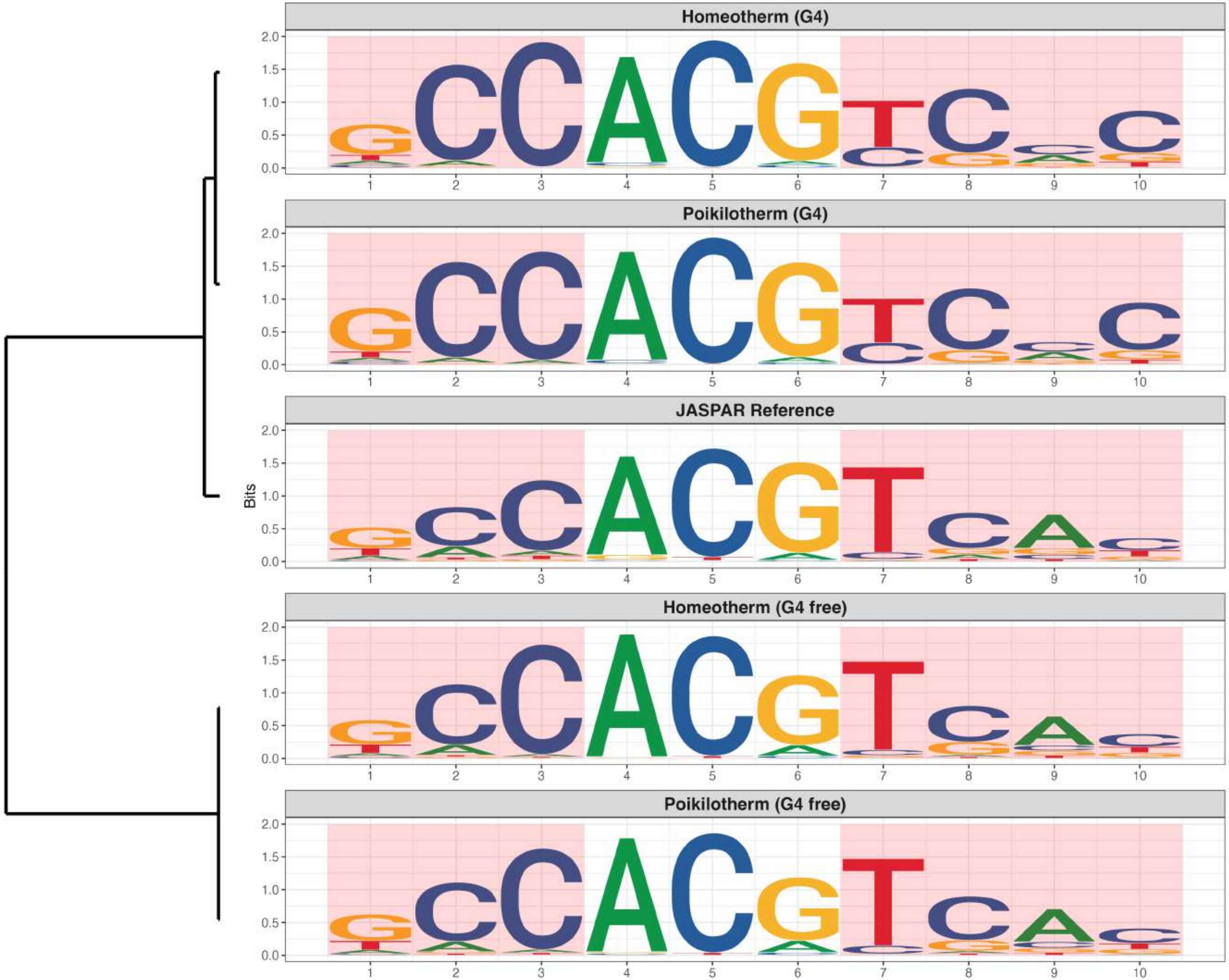

### Genome : Moderately Enriched in G4s

*MA1485.1 - FERD3L*

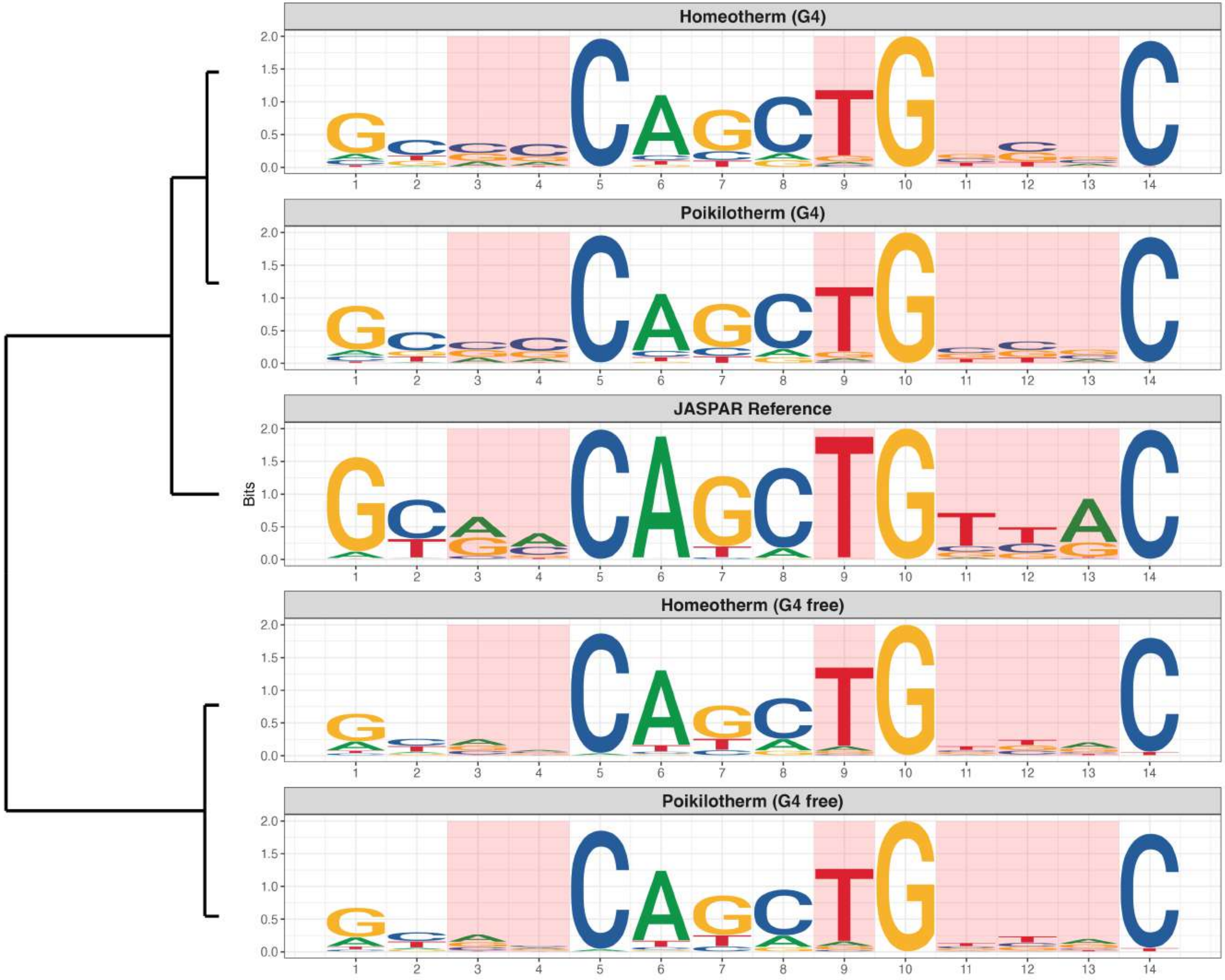

### Genome : Moderately Enriched in G4s

*MA1524.3 - Msgn1*

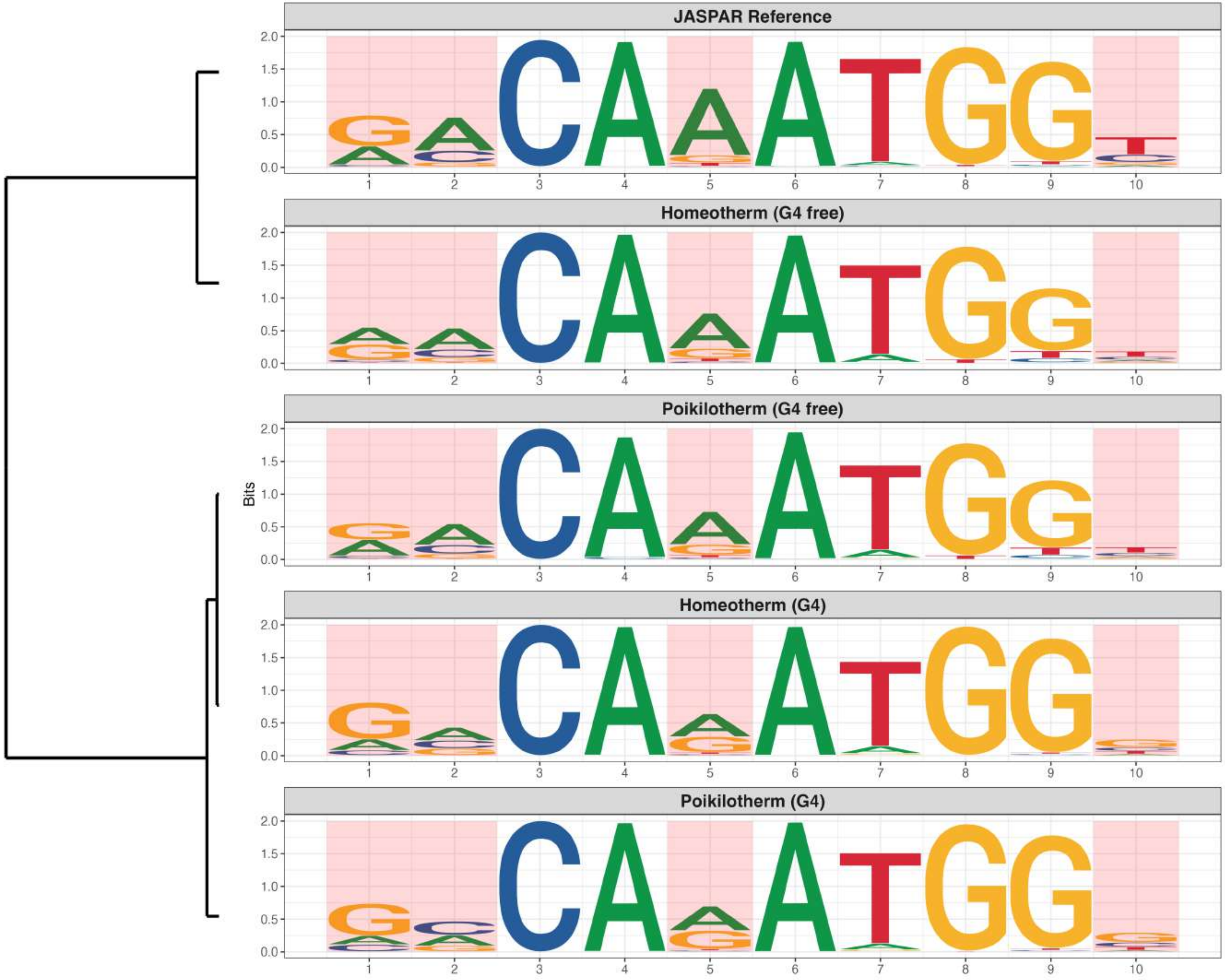

### Genome : Moderately Enriched in G4s

*MA1529.2 - NHLH2*

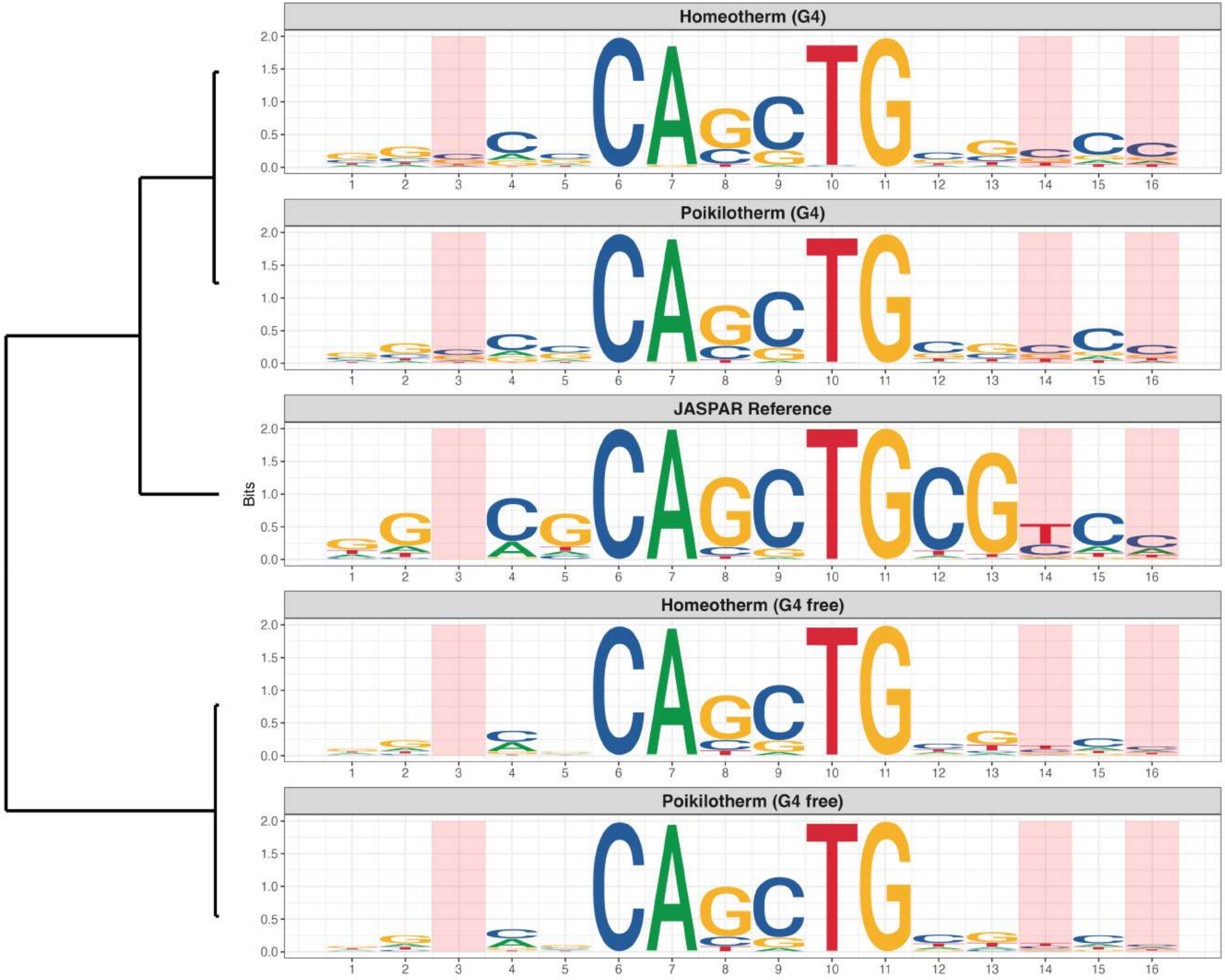

### Genome : Moderately Enriched in G4s

*MA1576.2 - THRB*

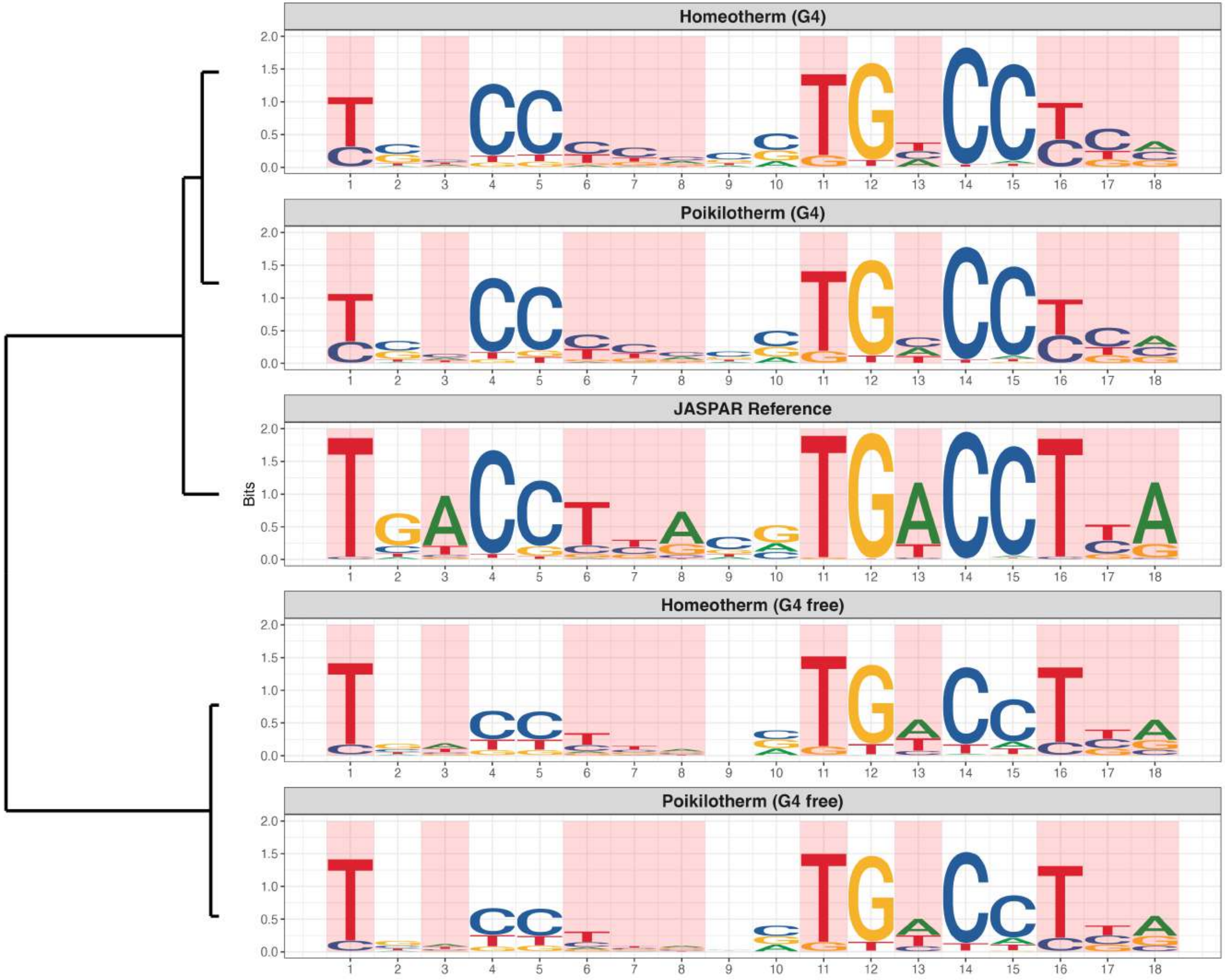

**Genome : Moderately Enriched in G4s**

*MA1630.3 - ZNF281*

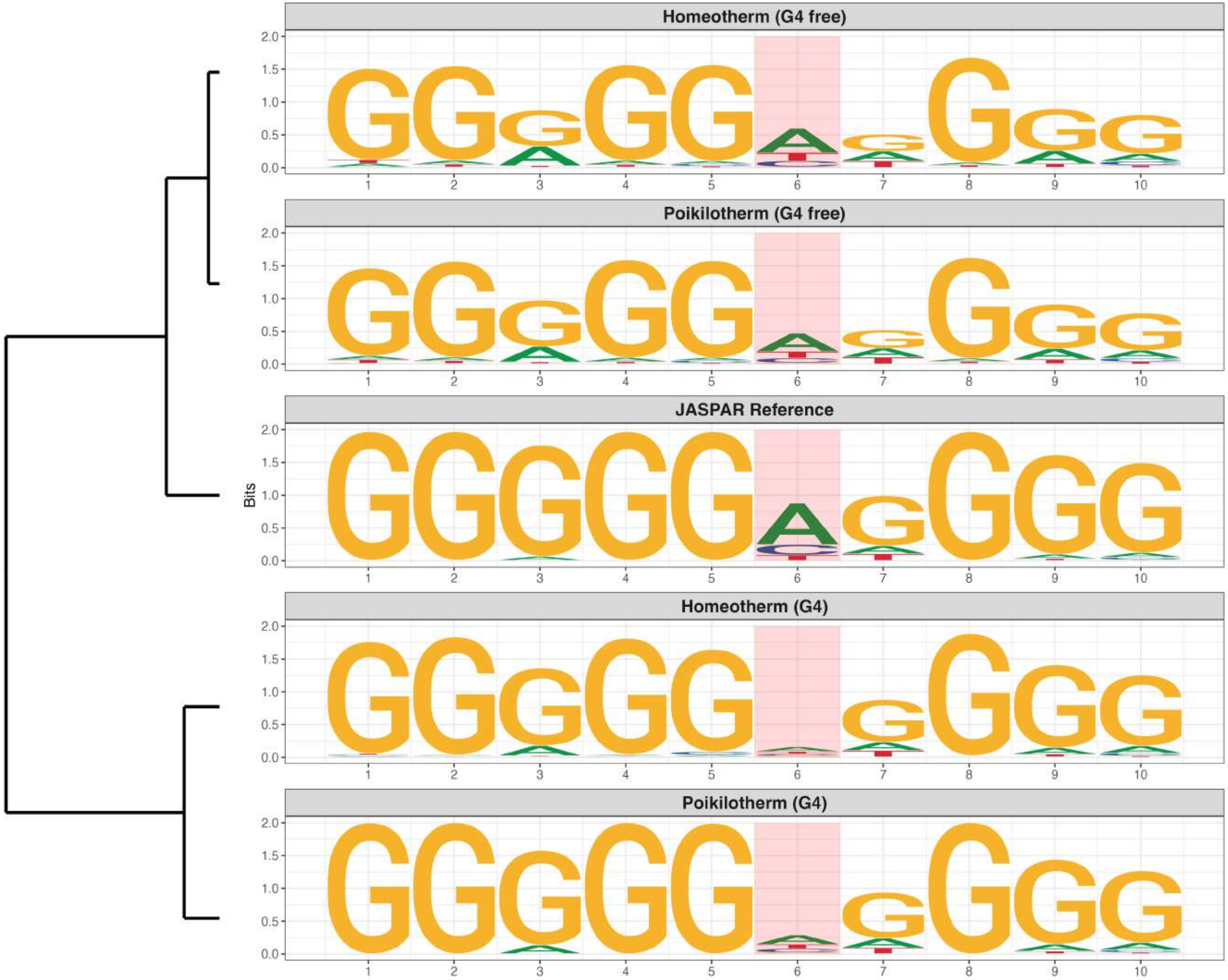

### Genome : Moderately Enriched in G4s

*MA1632.2 - ATF2*

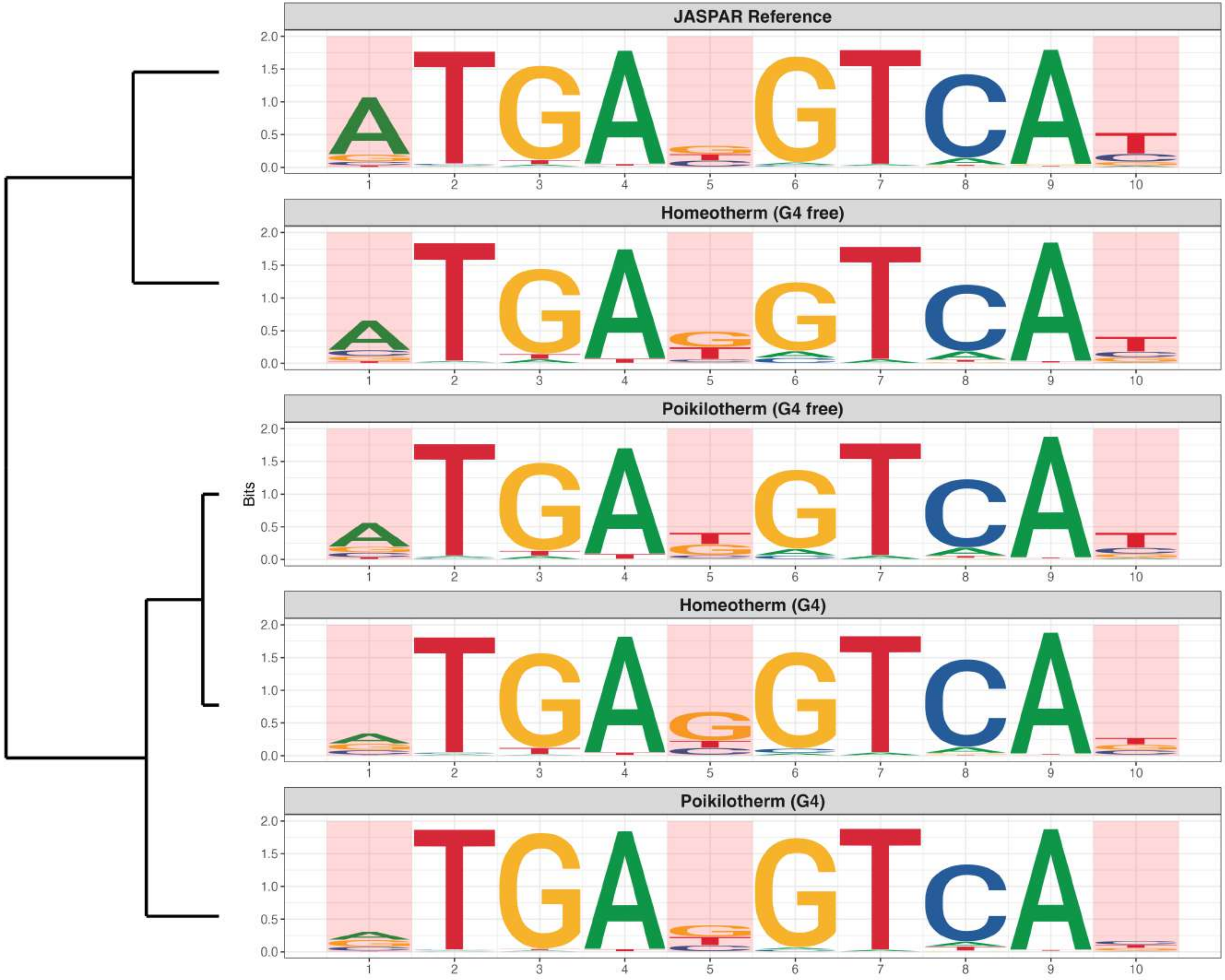

**Genome : Moderately Enriched in G4s**

*MA1725.2 - ZNF189*

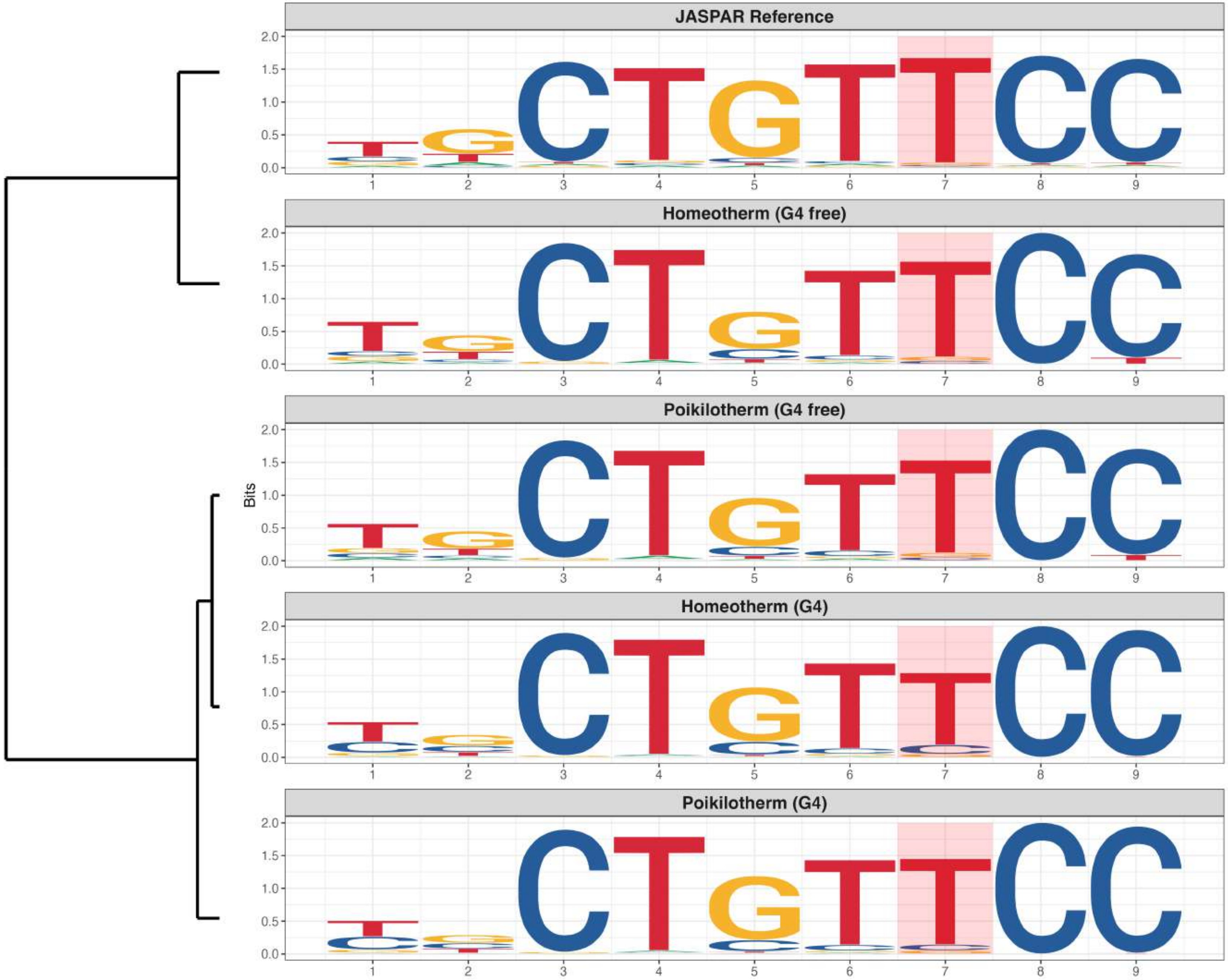

**Genome : Moderately Enriched in G4s**

*MA1727.2 - ZNF417*

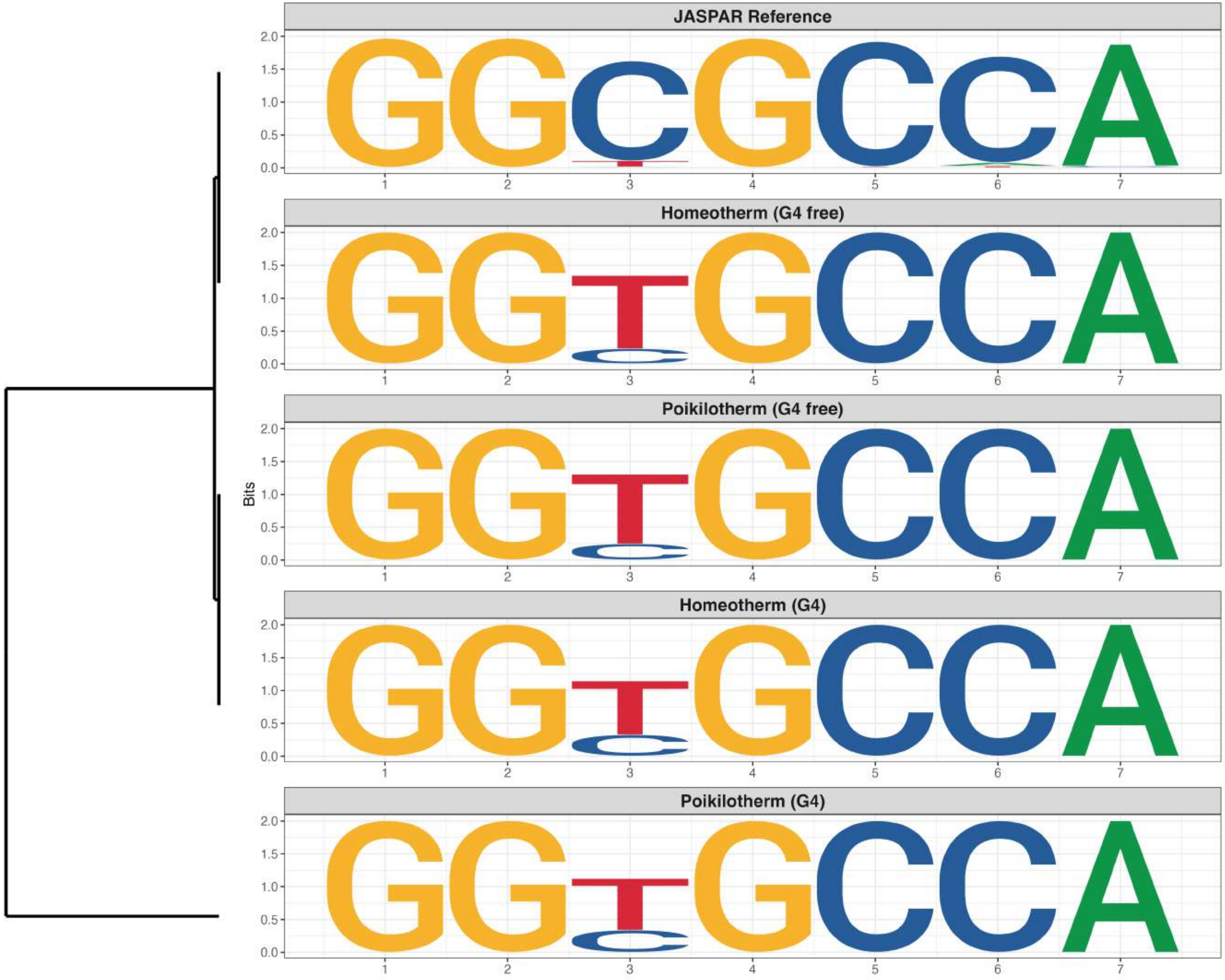

### Genome : Moderately Enriched in G4s

*MA1938.2 - ERF::NHLH1*

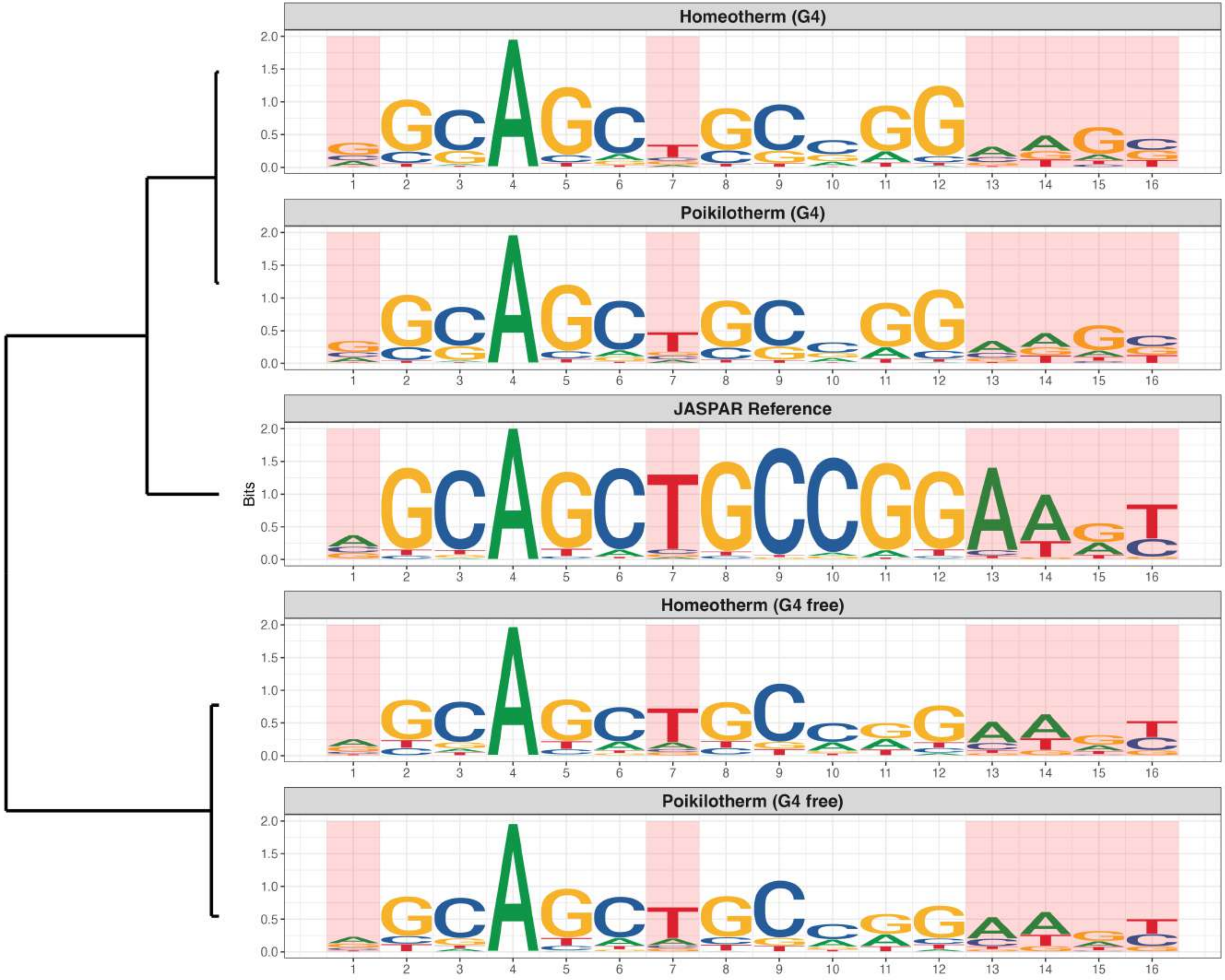

### Genome : Moderately Enriched in G4s

*MA1977.2 - ZNF324*

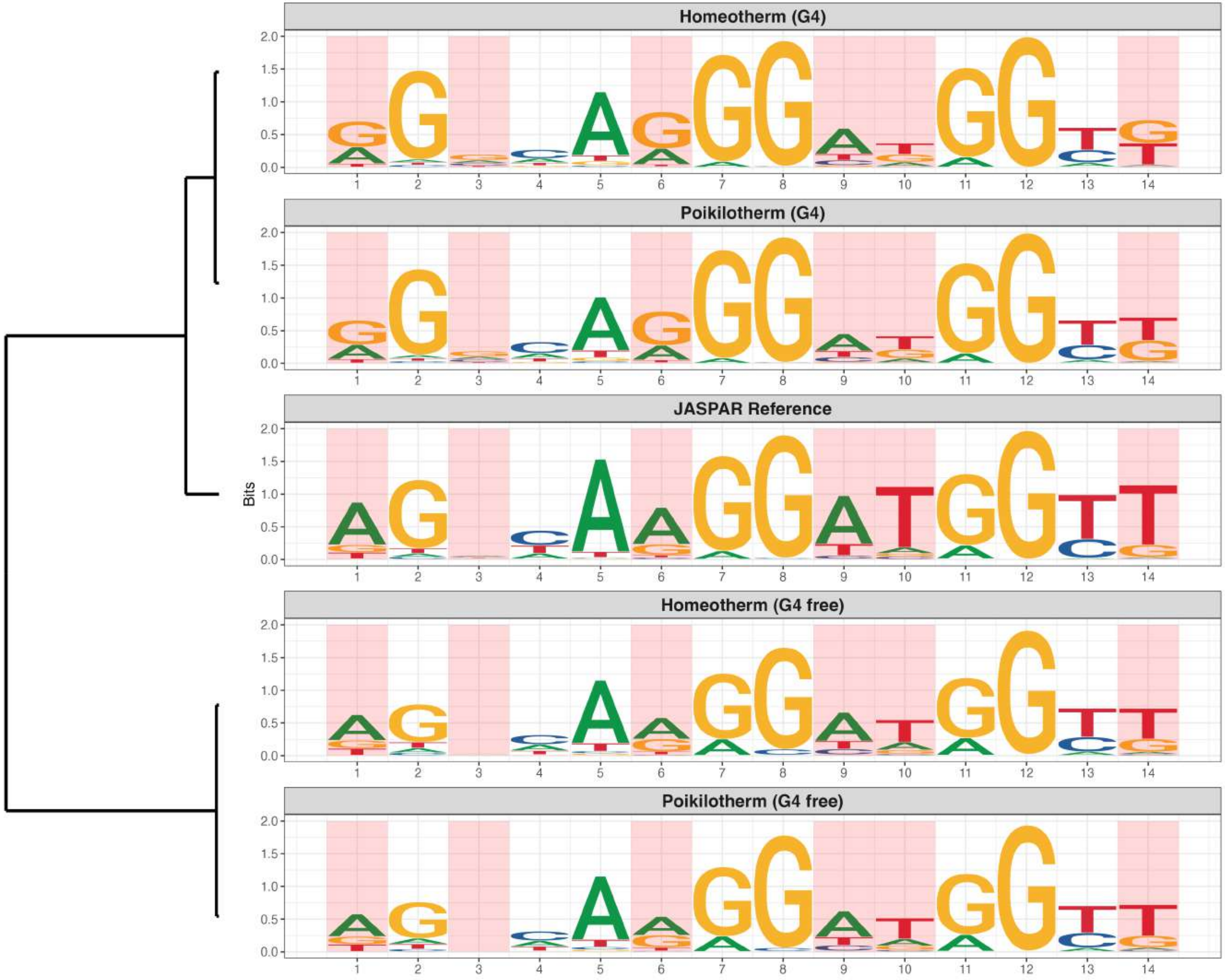

### Genome : Moderately Enriched in G4s

*MA2489.1 - ZFP28*

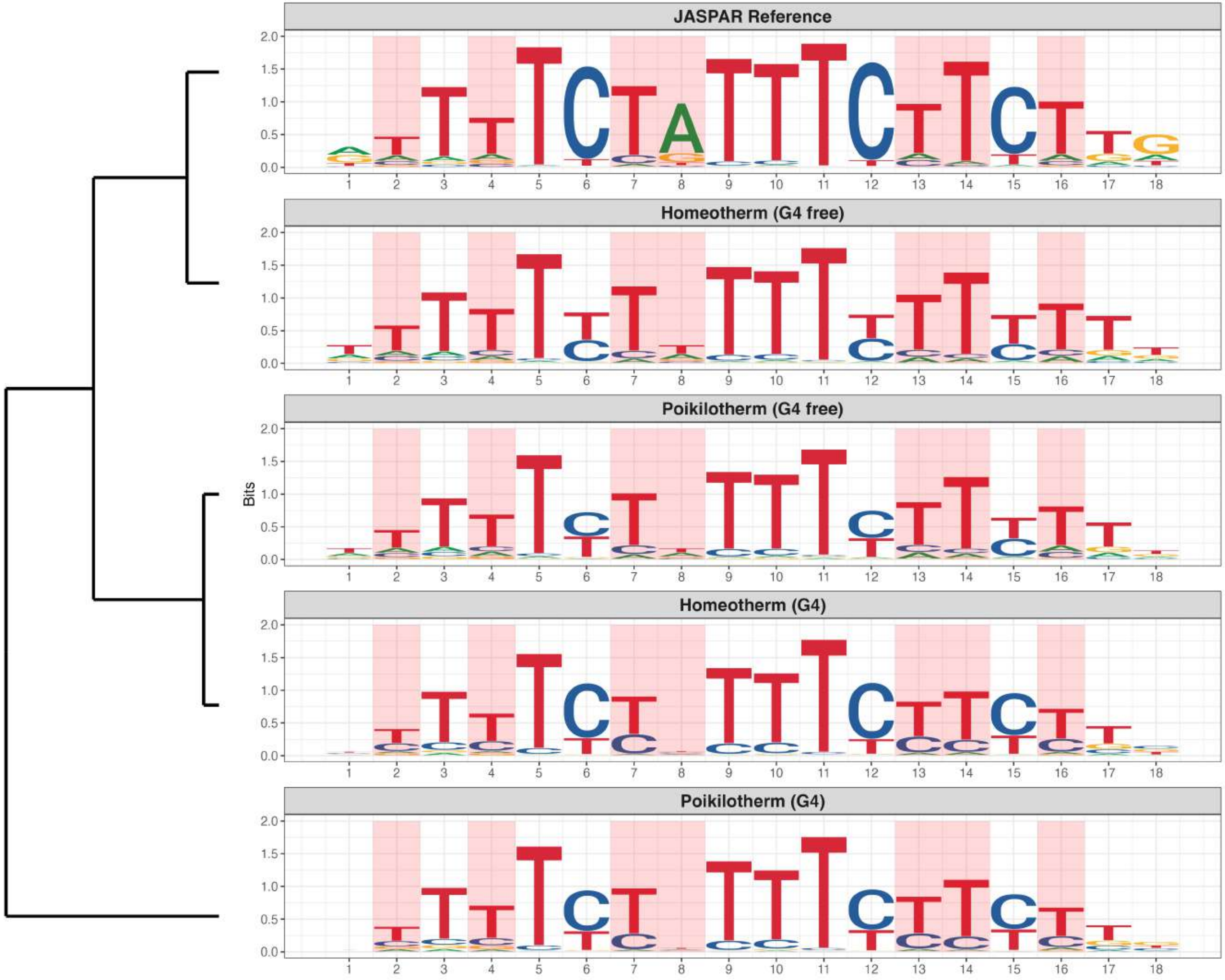

**Genome : Moderately Enriched in G4s**

*MA2538.1 - CGGBP1*

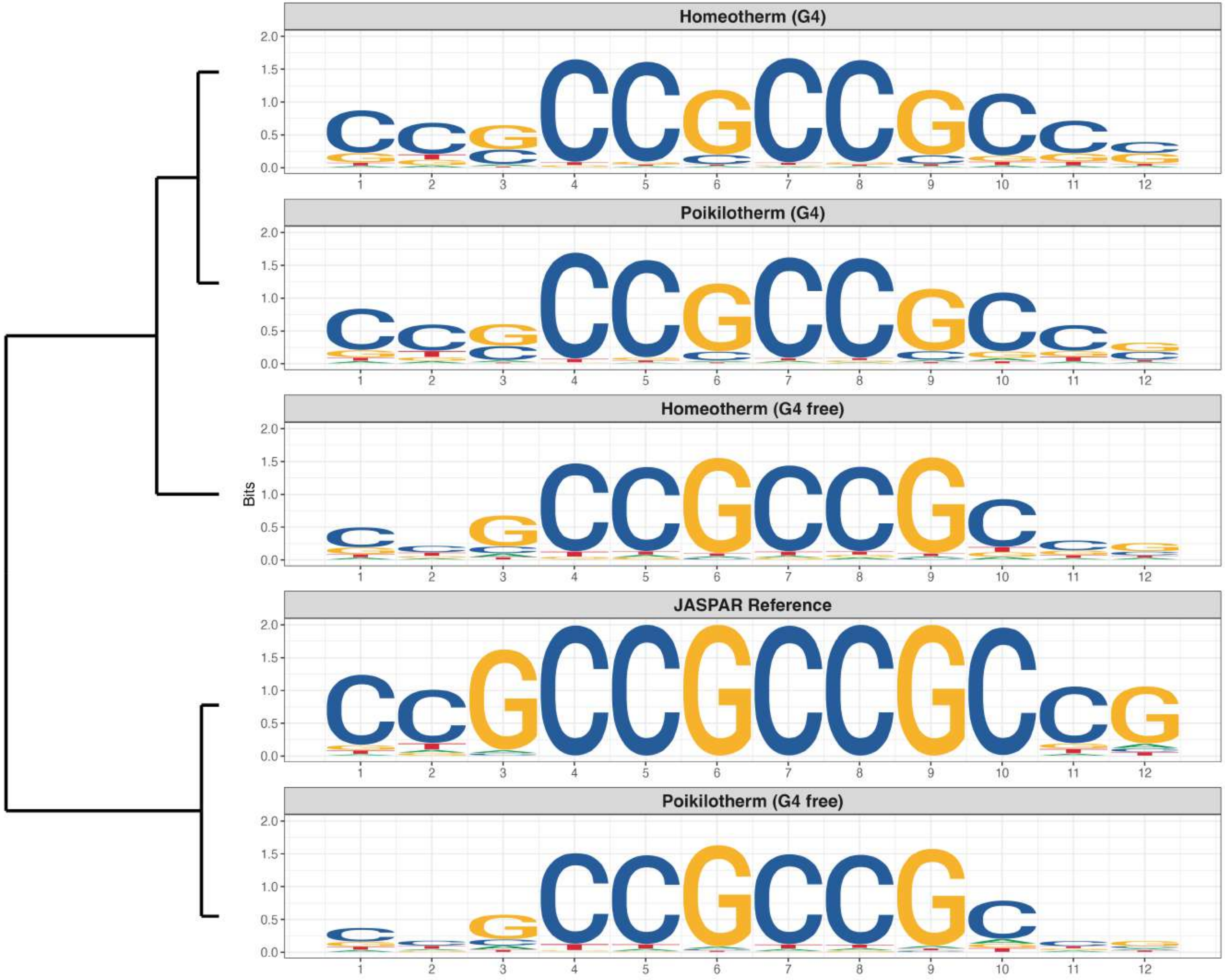

### Genome : Moderately Enriched in G4s

*MA2541.1 - CREB3L3*

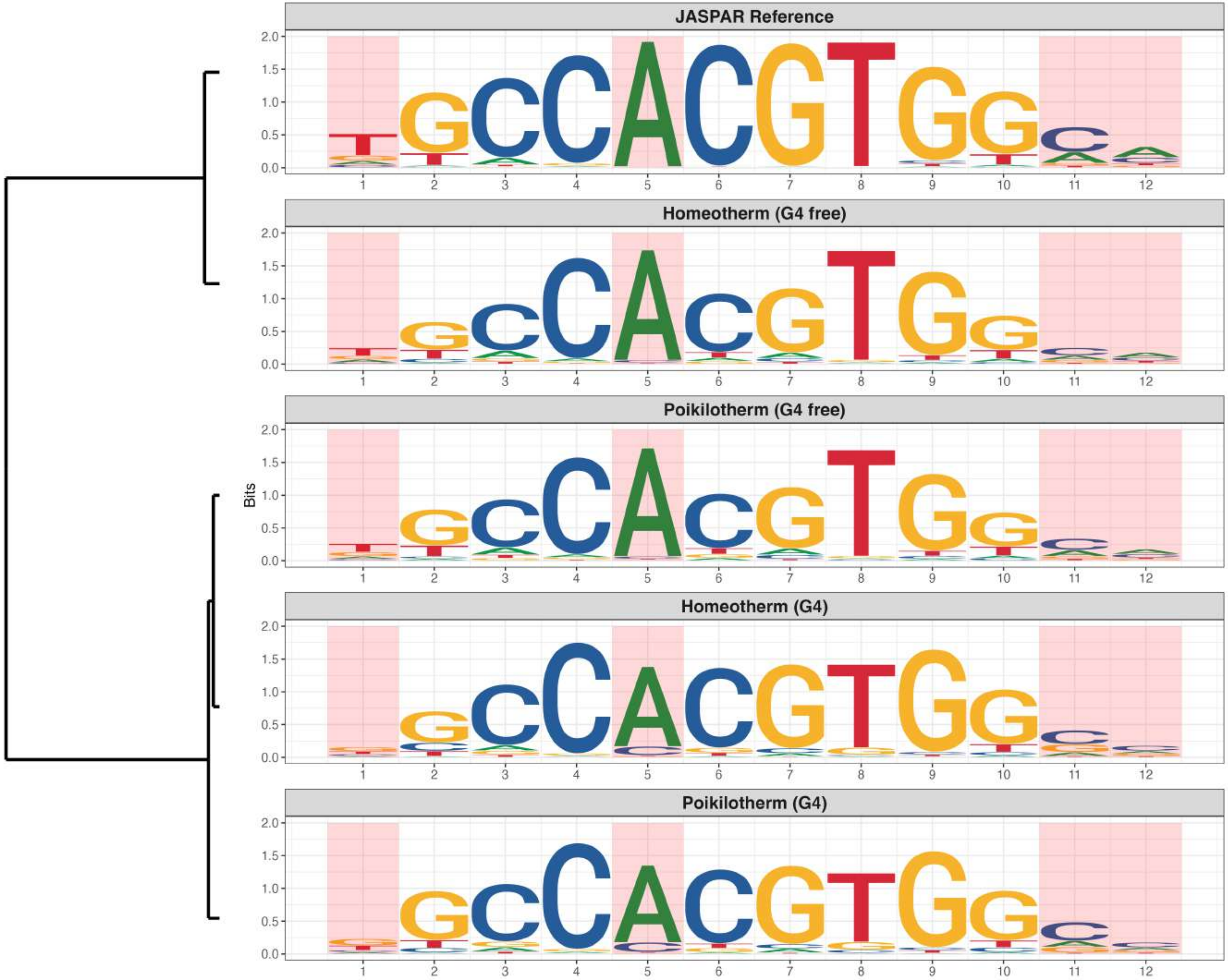

### Genome : Moderately Enriched in G4s

*MA2553.1 - ZNF407*

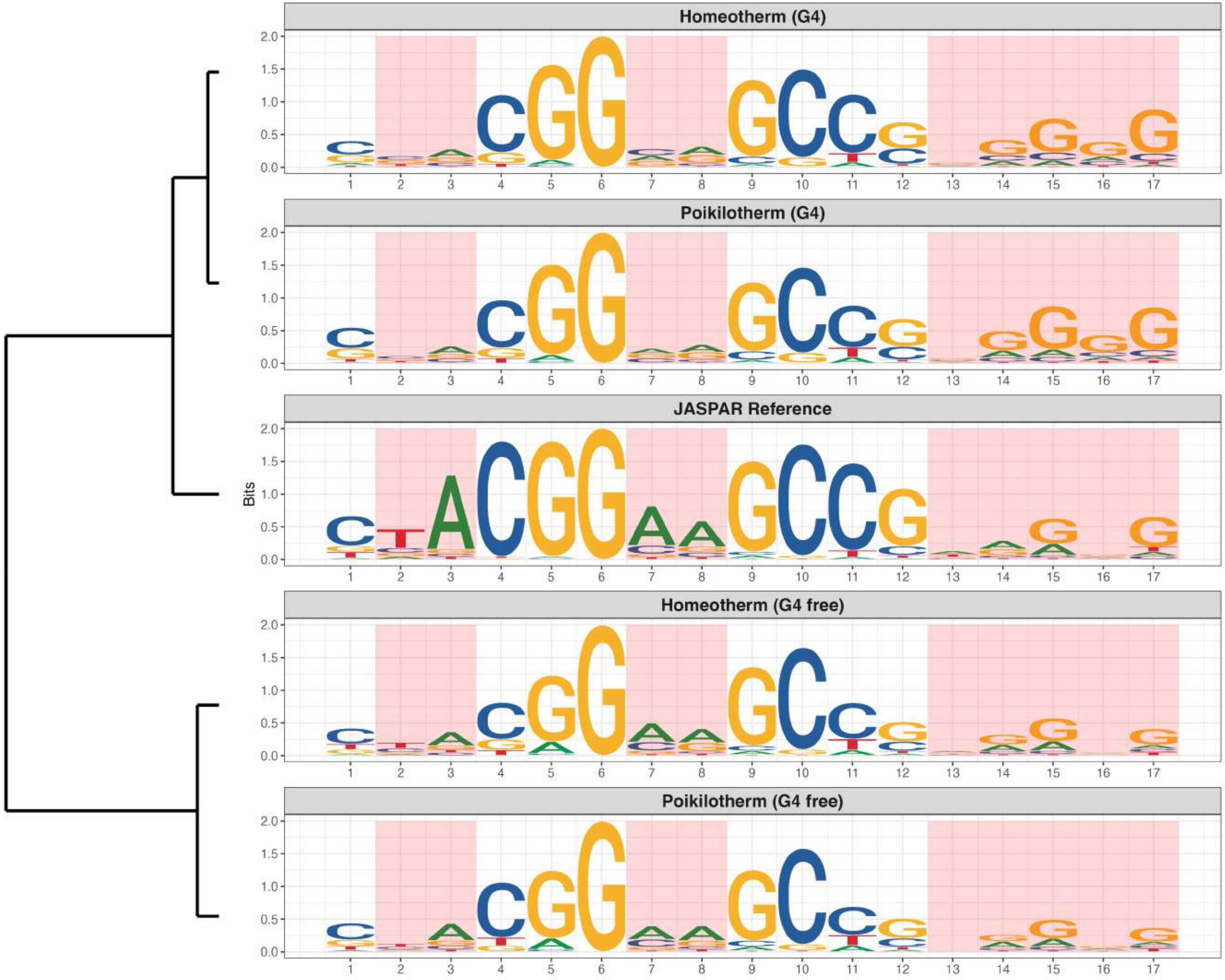

**Genome : Moderately Enriched in G4s**

*MA2554.1 - ZNF471*

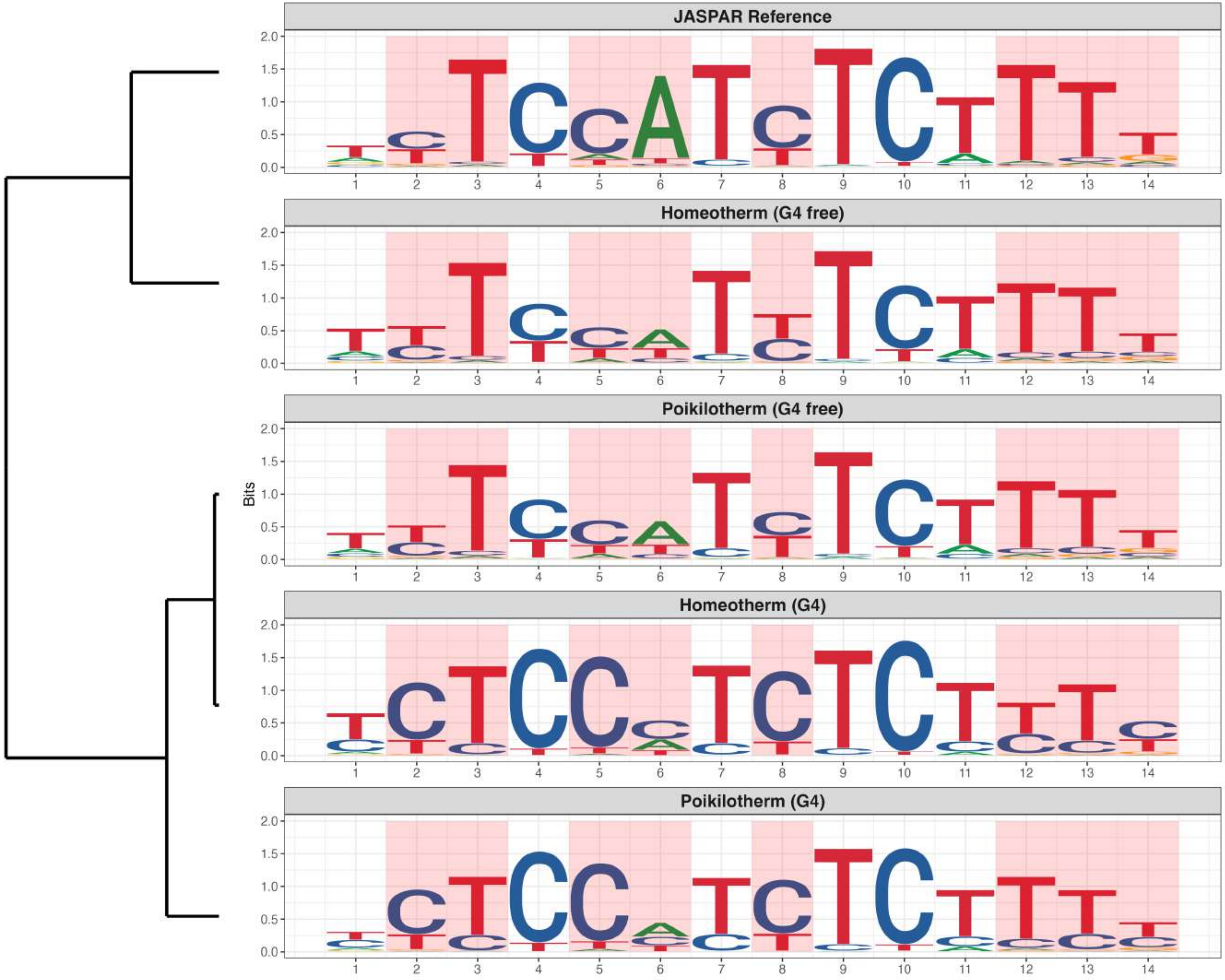

### Genome : Moderately Enriched in G4s

*MA2569.1 - ZNF676*

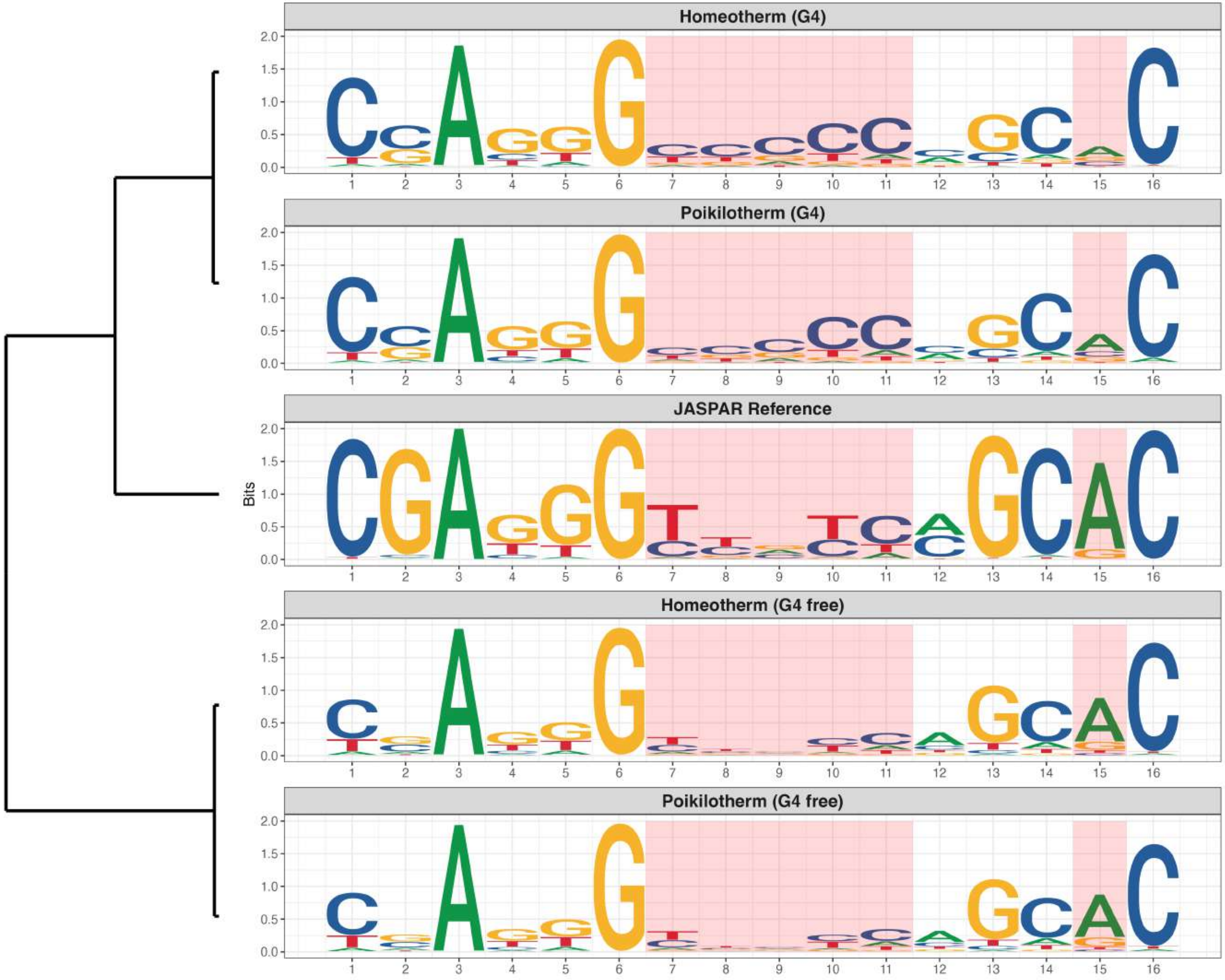

### Genome : Moderately Enriched in G4s

*MA2581.1 - ZNF66*

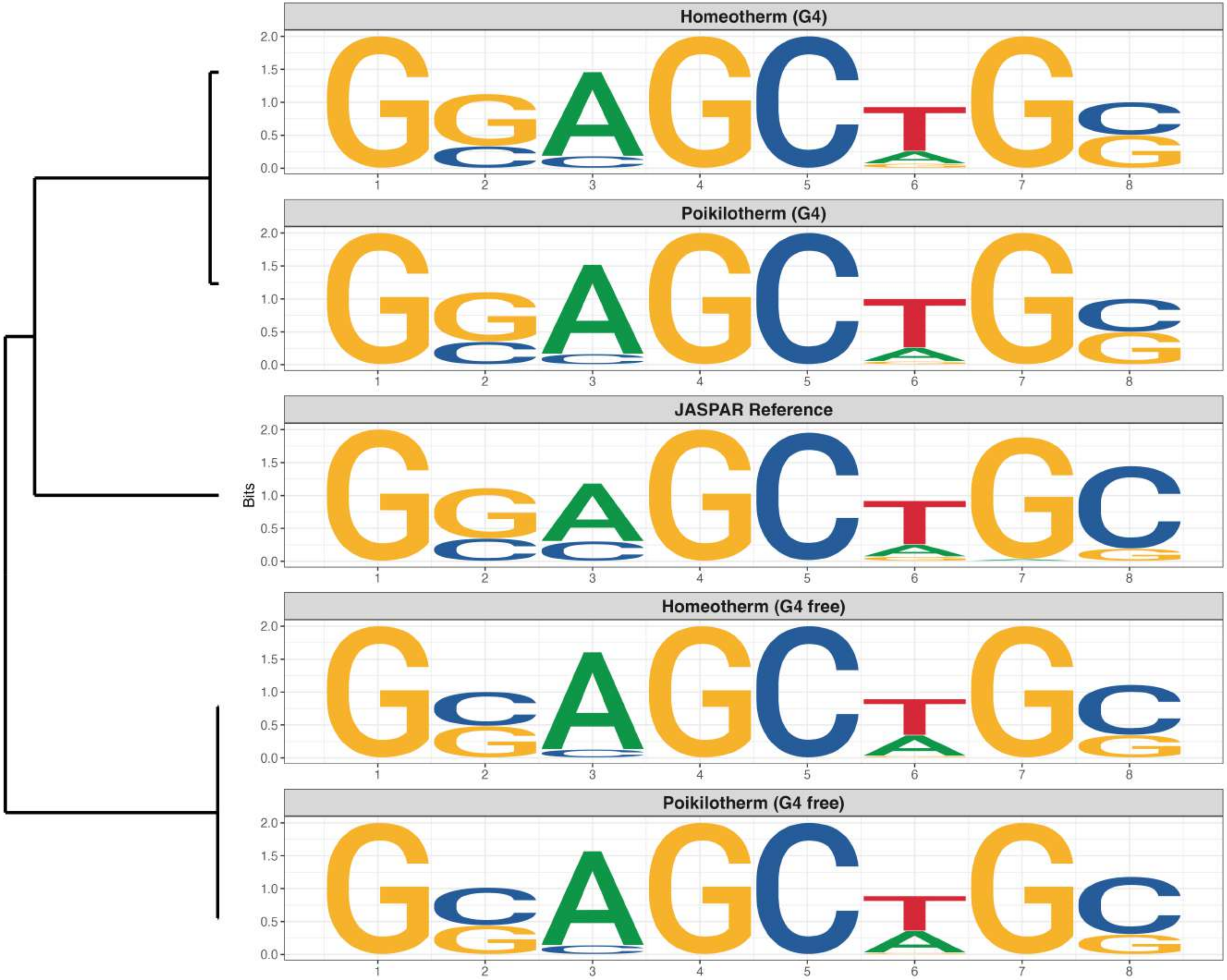

### Genome : Moderately Enriched in G4s

*MA2588.1 - FAM200B*

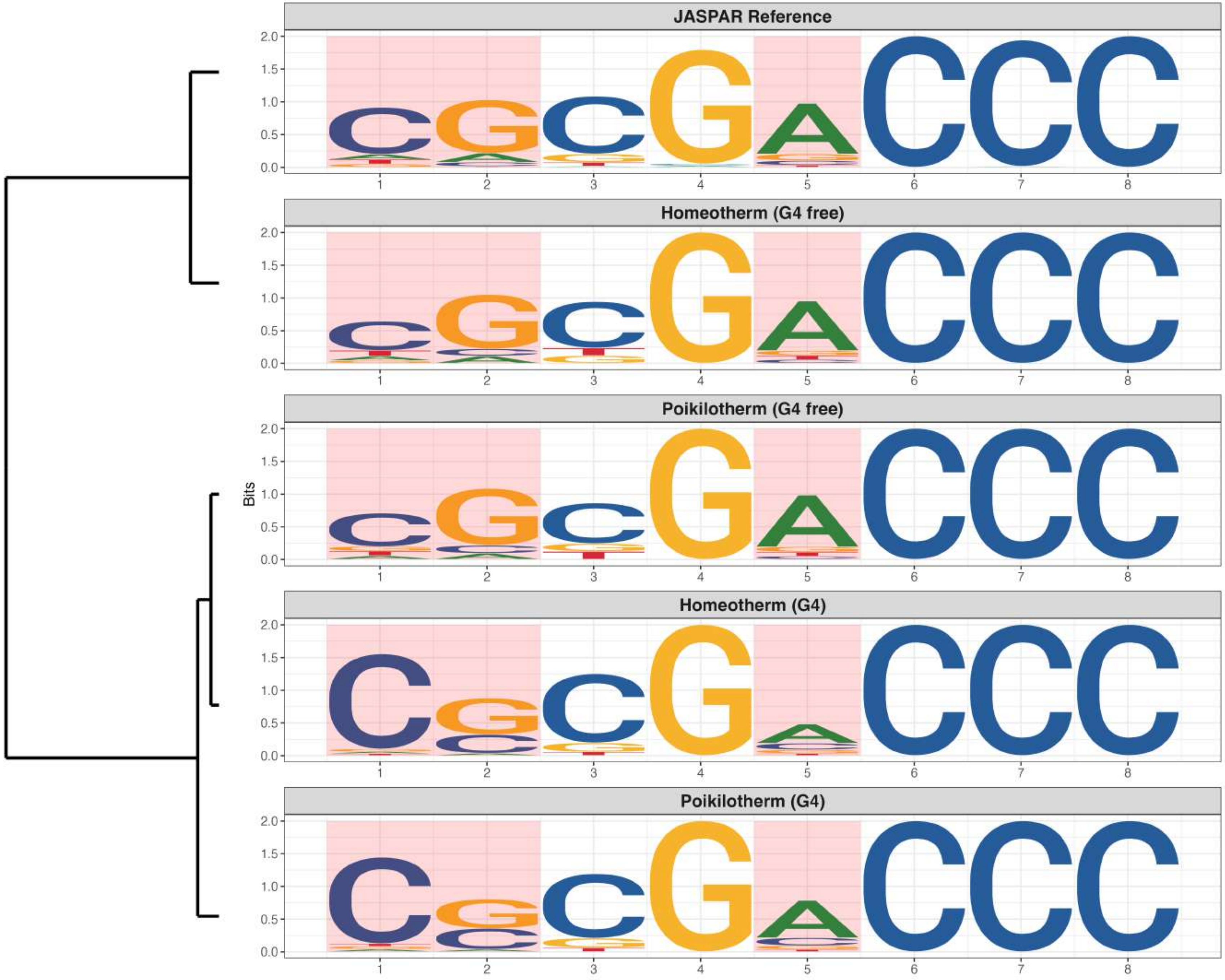

### Genome : Moderately Enriched in G4s

*MA2593.1 - ZNF362*

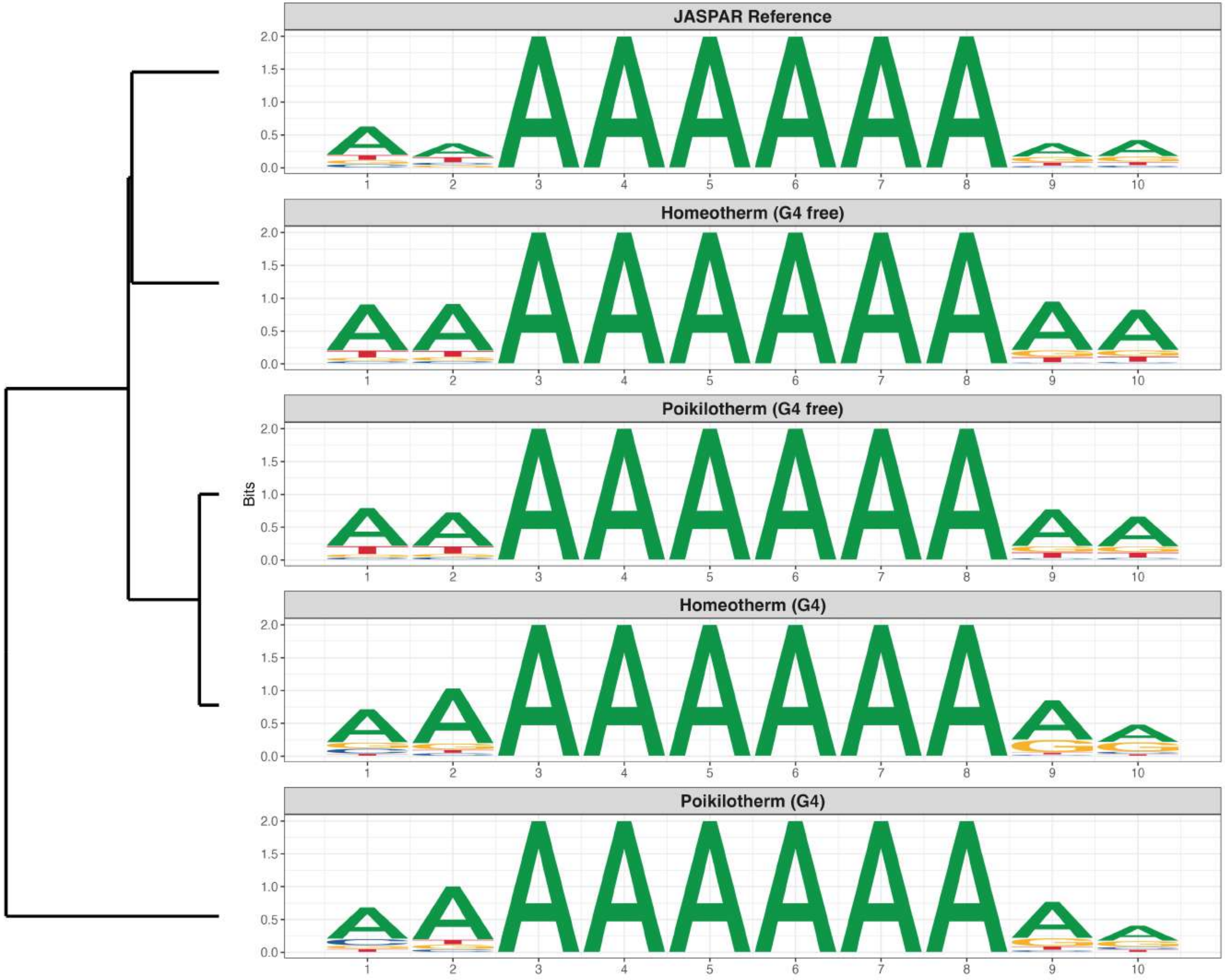

**Genome : Moderately Enriched in G4s**

*MA2594.1 - DMTF1*

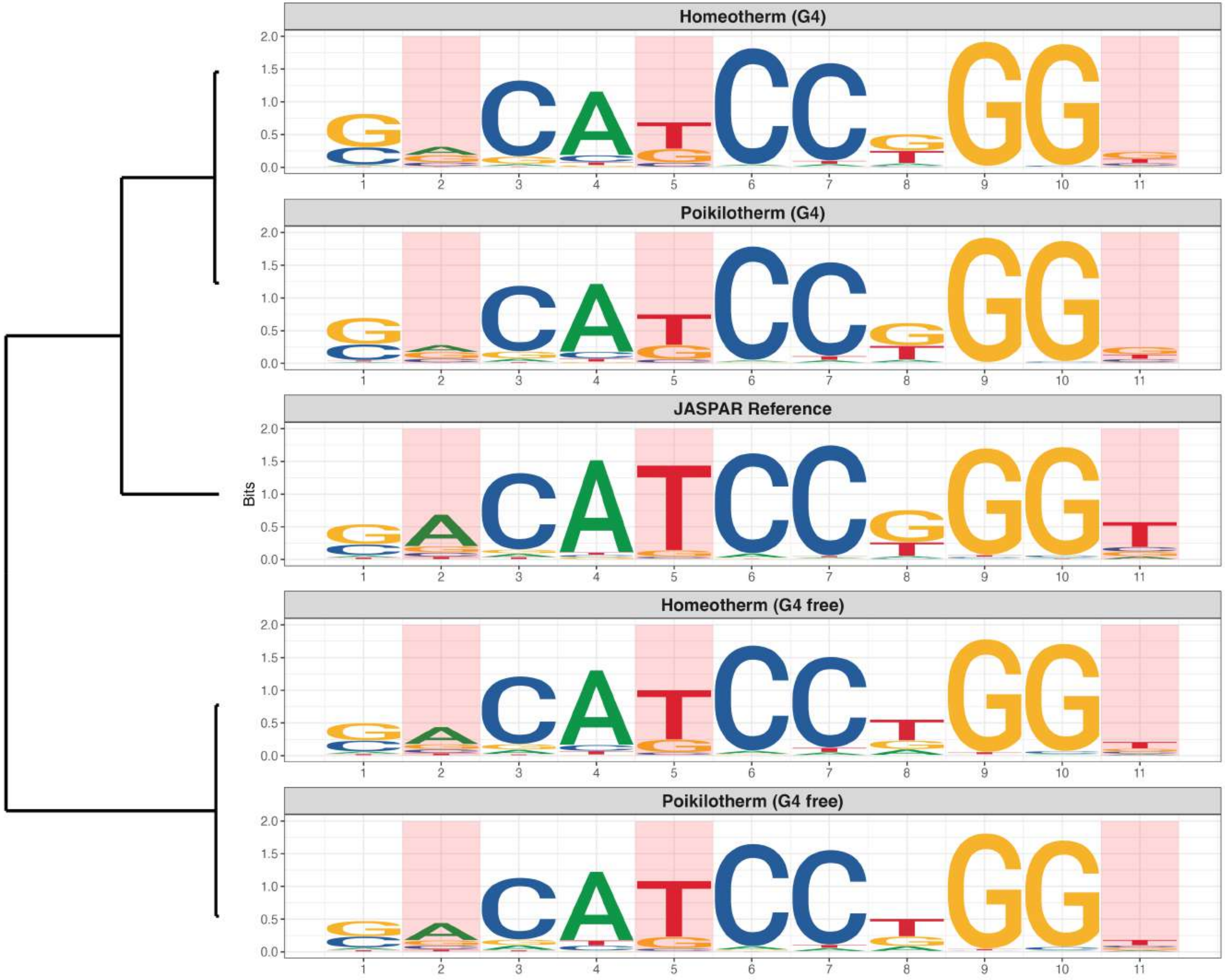

### Genome : Moderately Enriched in G4s

*MA2601.1 - USF3*

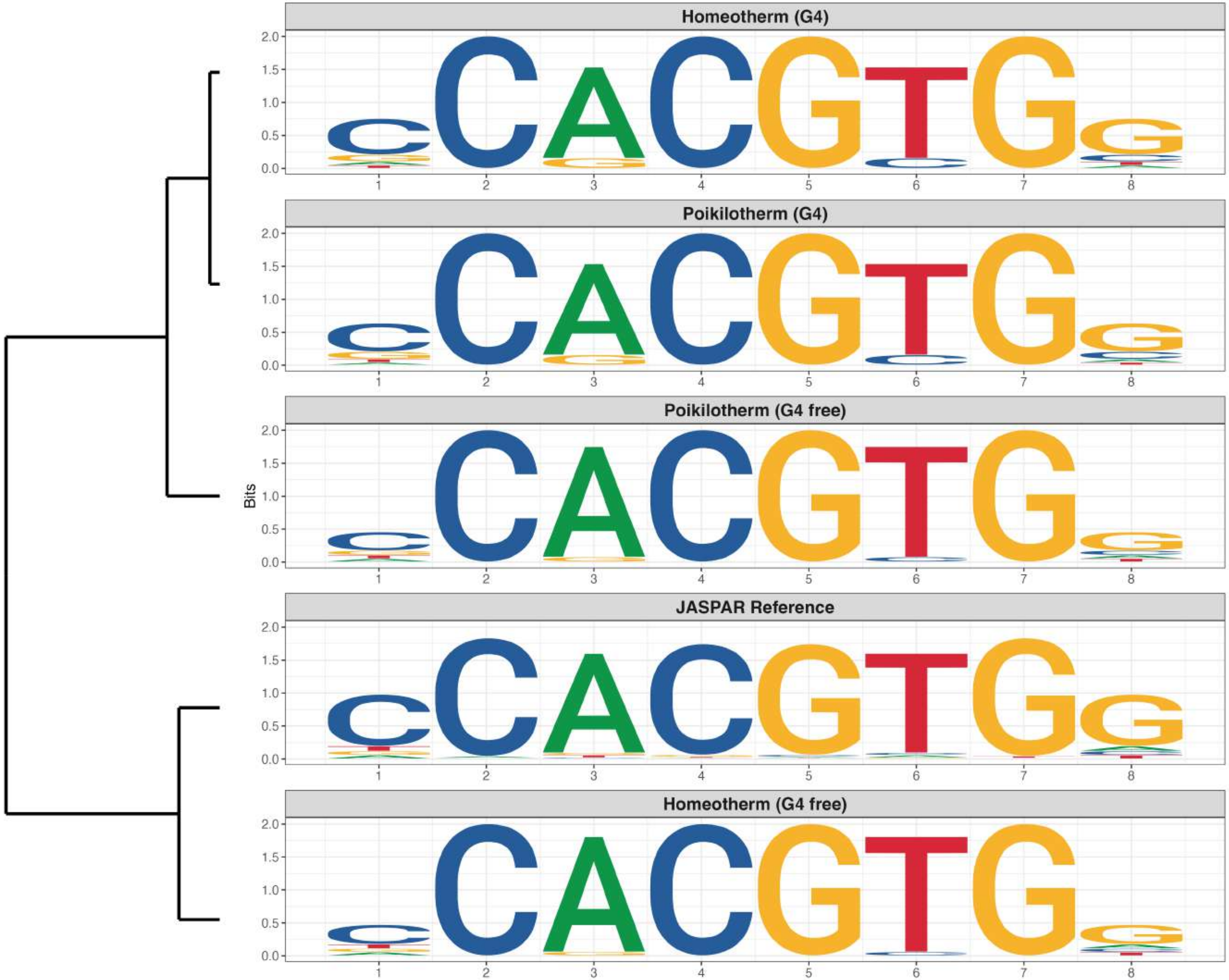

**Genome : Moderately Enriched in G4s**

*MA2628.1 - HELT*

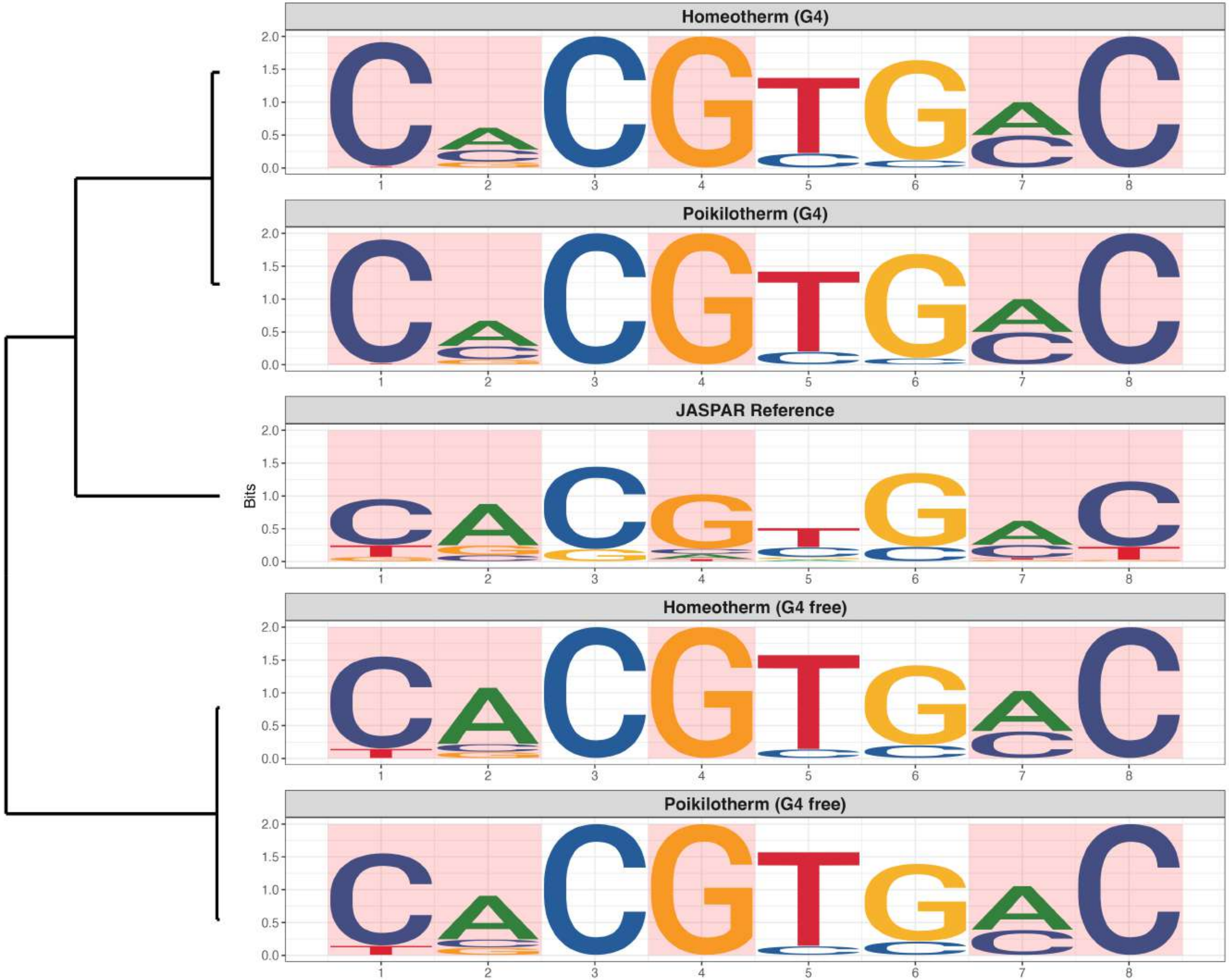

### Genome : Moderately Enriched in G4s

*MA2689.1 - ZNF470*

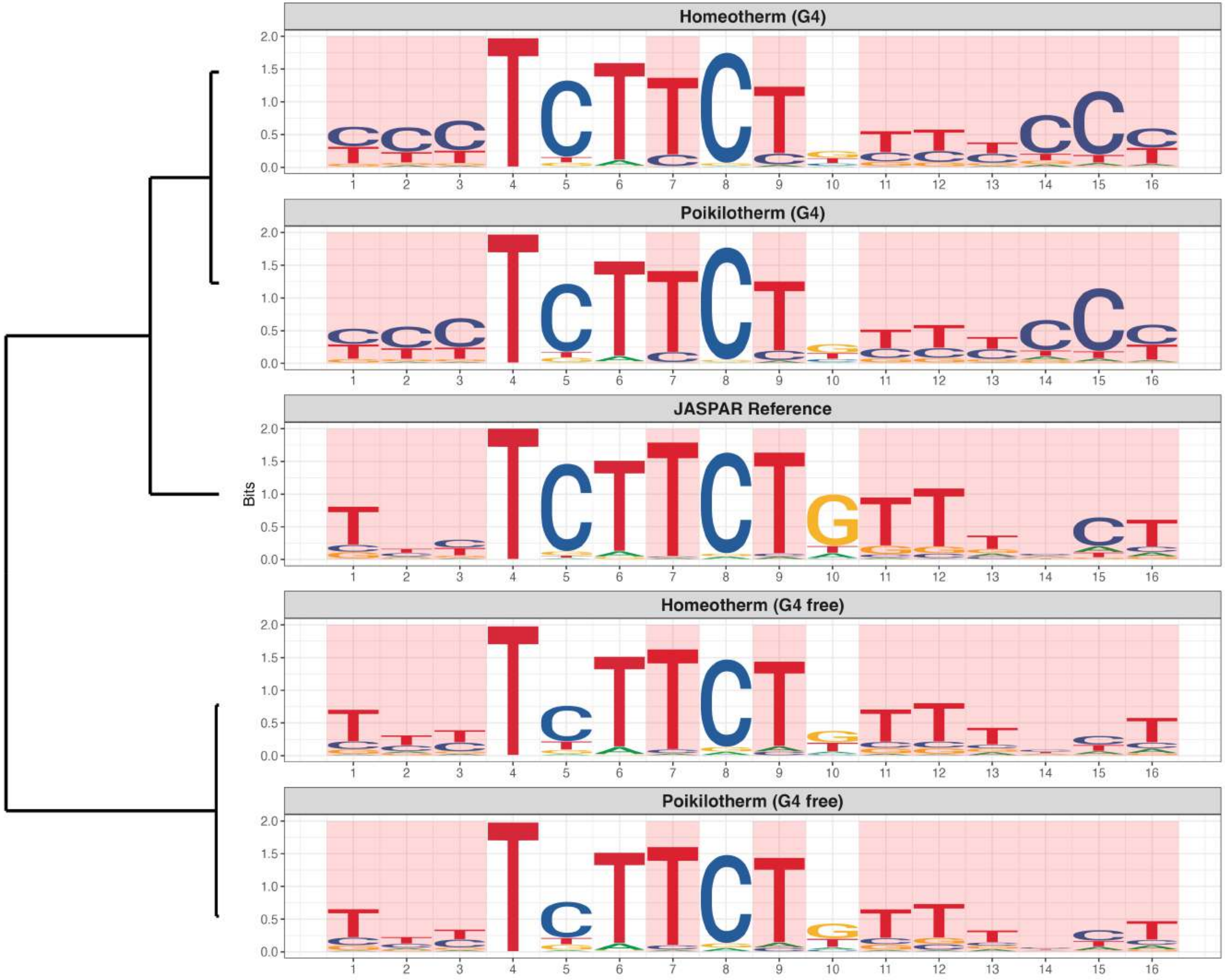

### Genome : Highly Reduced in G4s

*MA0469.4 - E2F3*

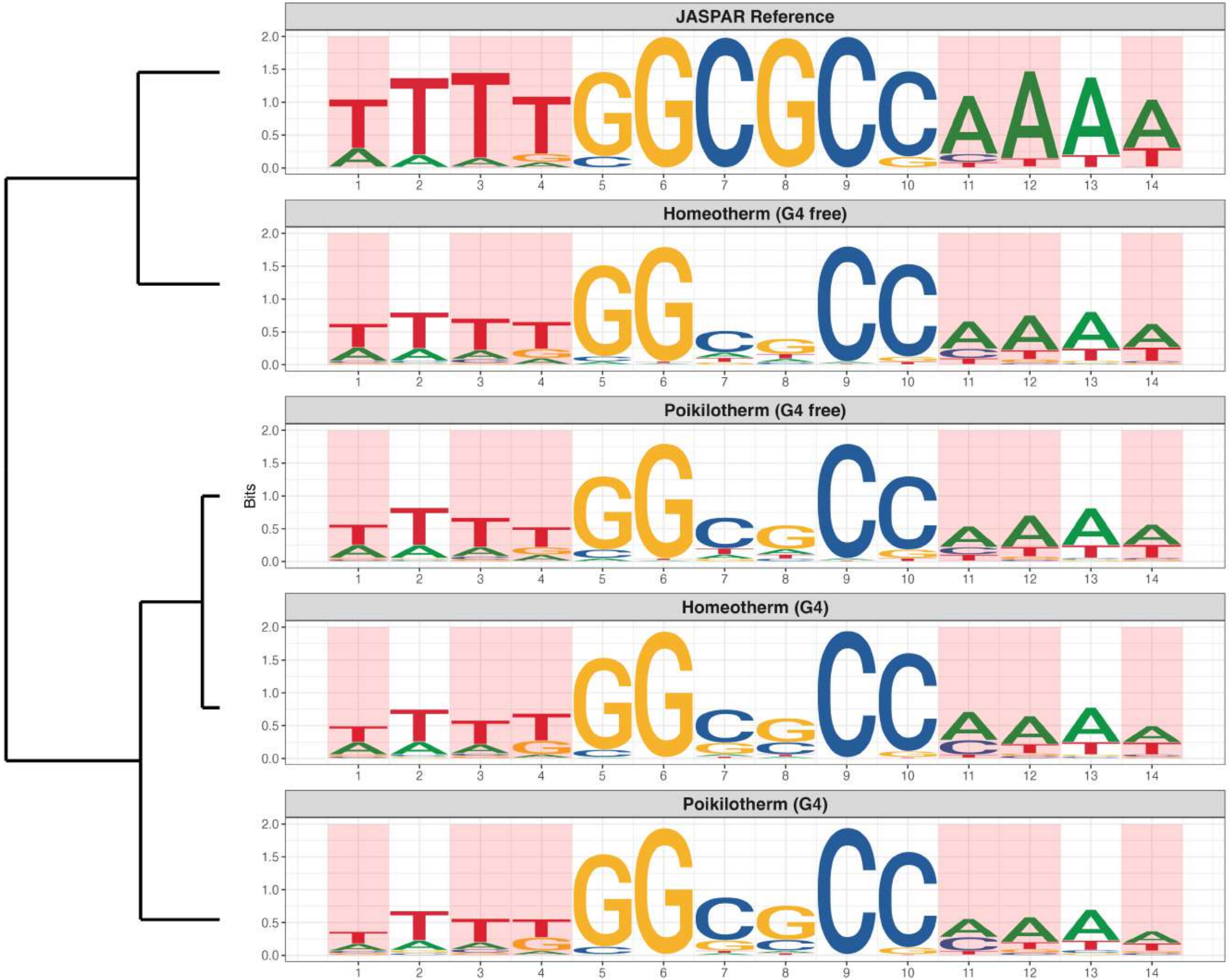

### Genome : Highly Reduced in G4s

*MA0638.2 - CREB3*

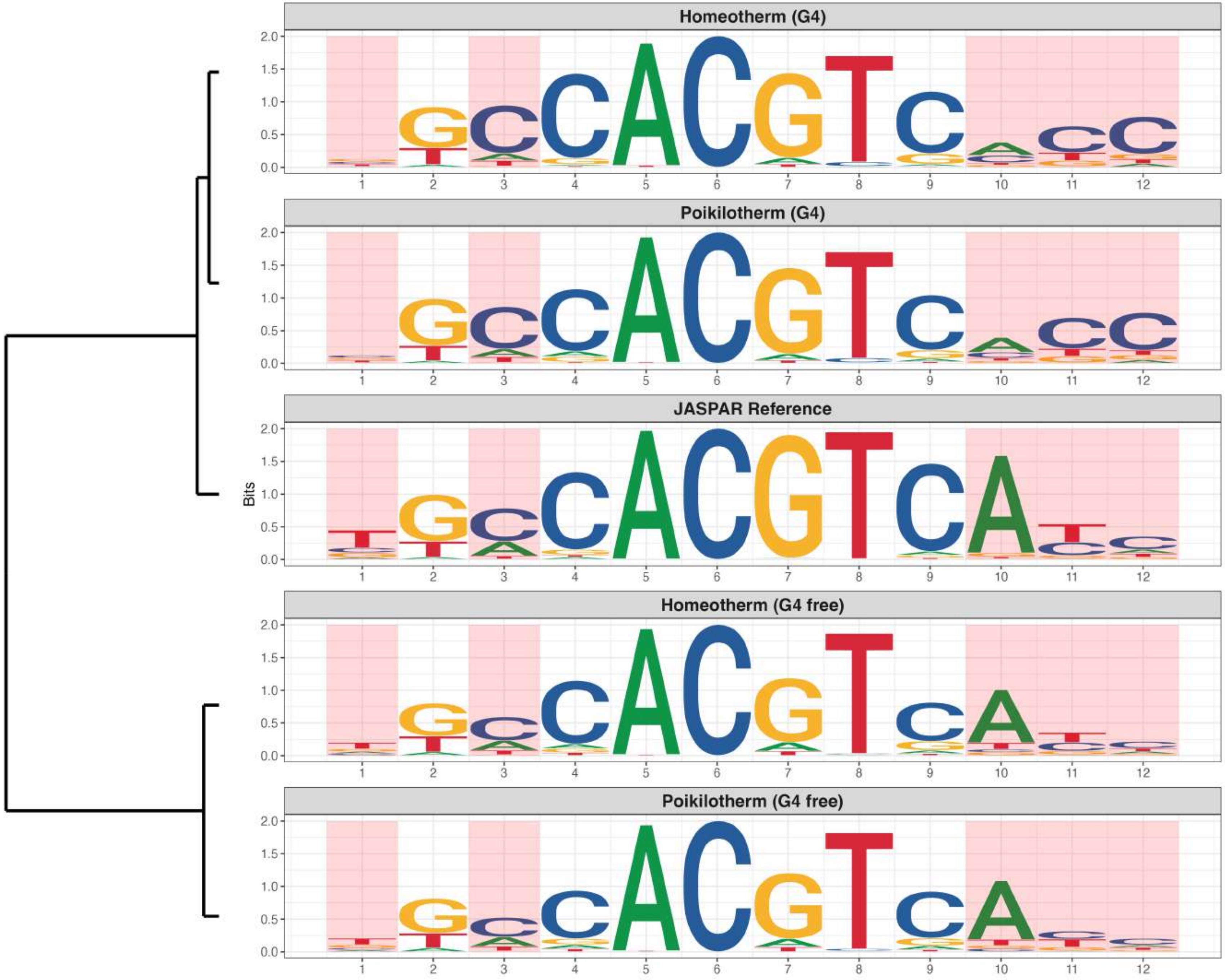

**Genome : Highly Reduced in G4s**

*MA0763.2 - ETV3*

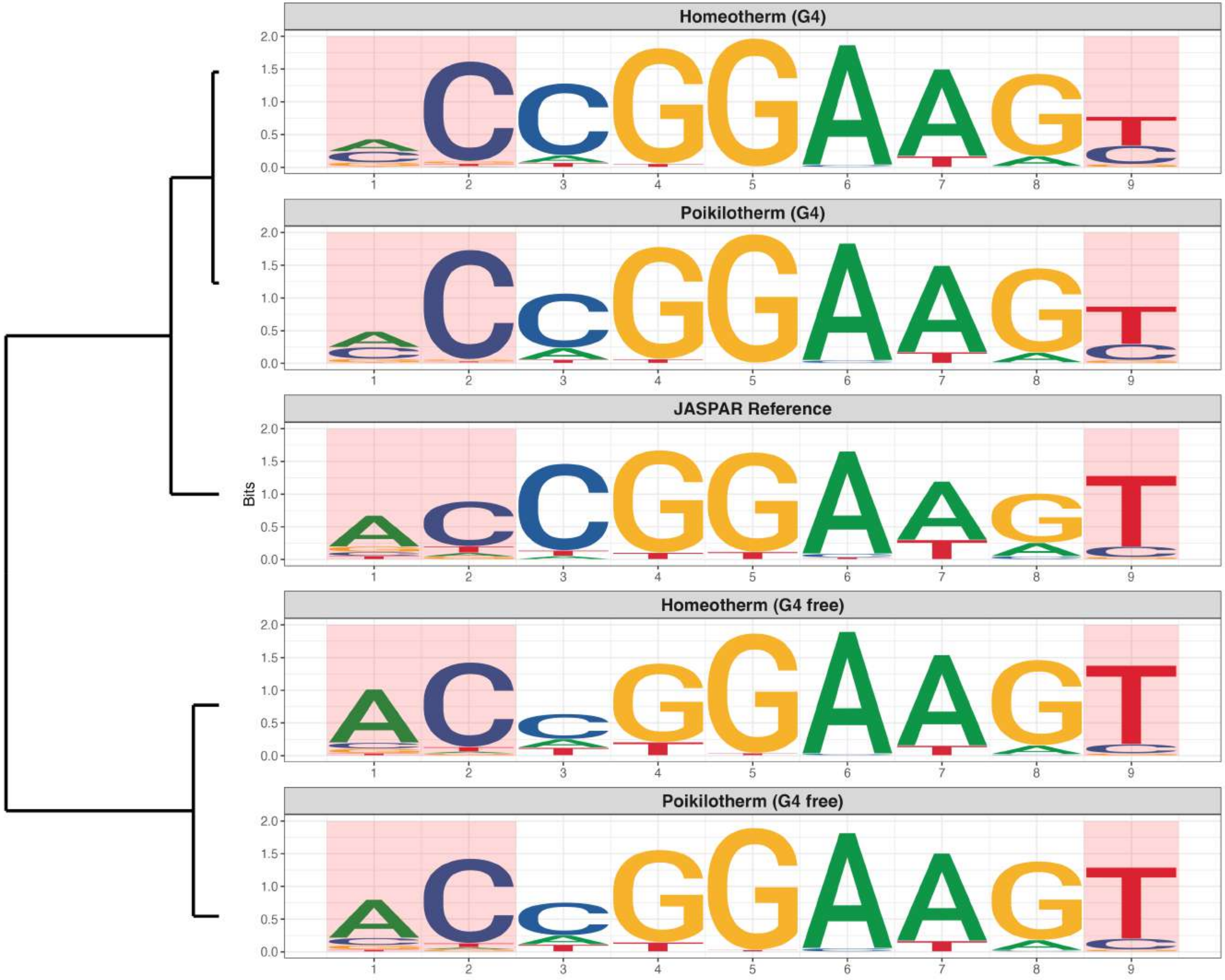

**Genome : Highly Reduced in G4s**

*MA0798.3 - RFX3*

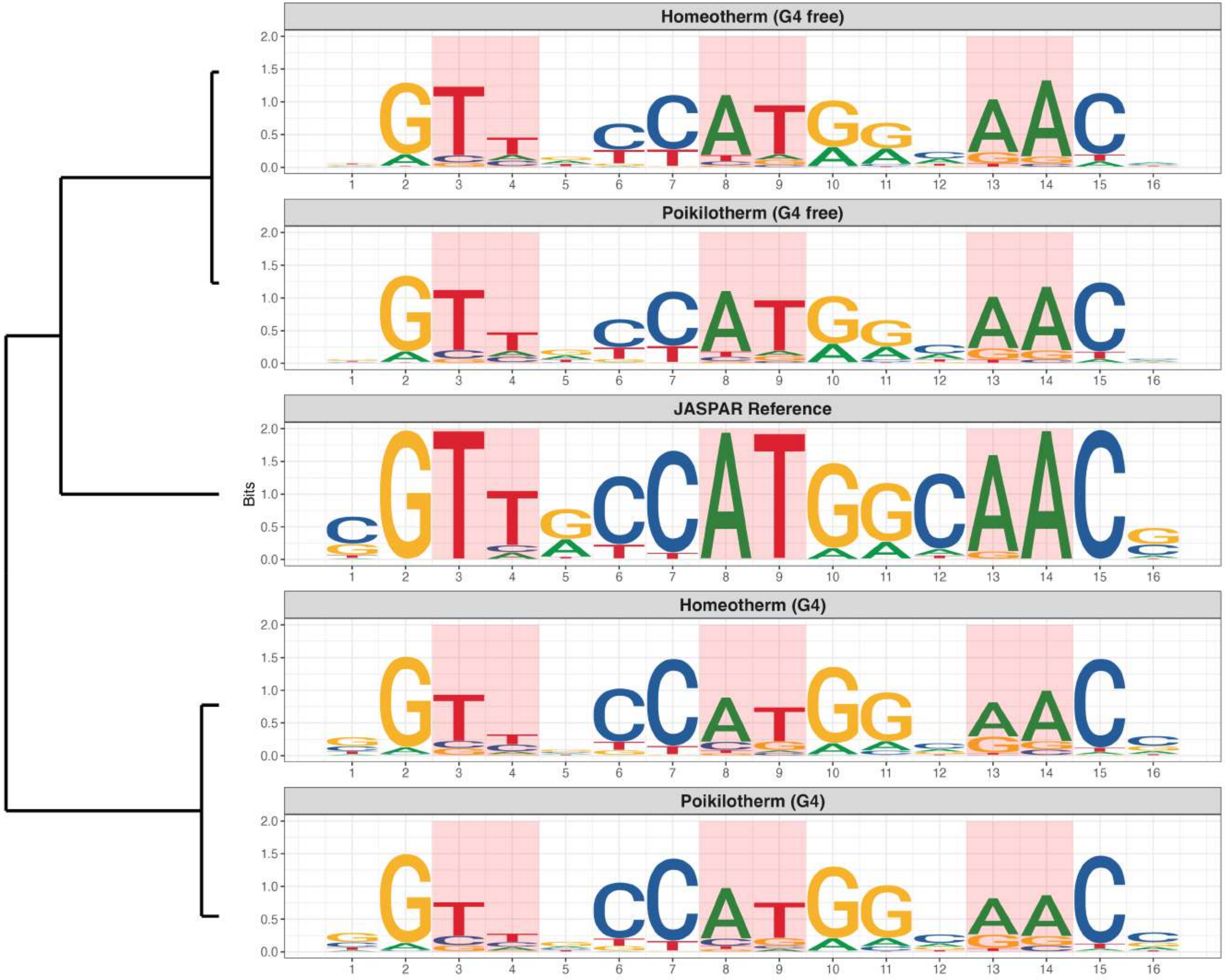

### Genome : Highly Reduced in G4s

*MA1466.2 - ATF6*

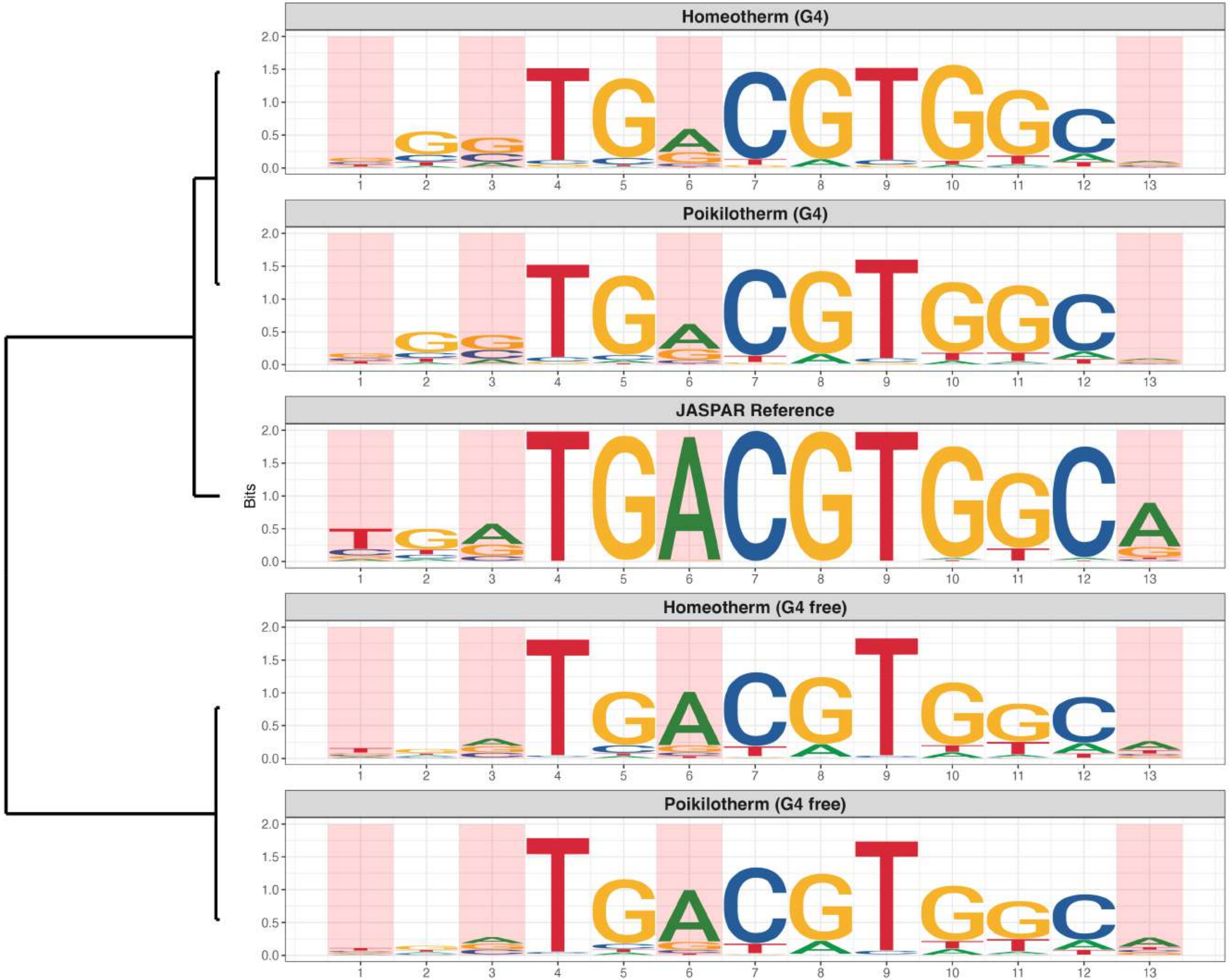

### Genome : Highly Reduced in G4s

*MA1483.3 - ELF2*

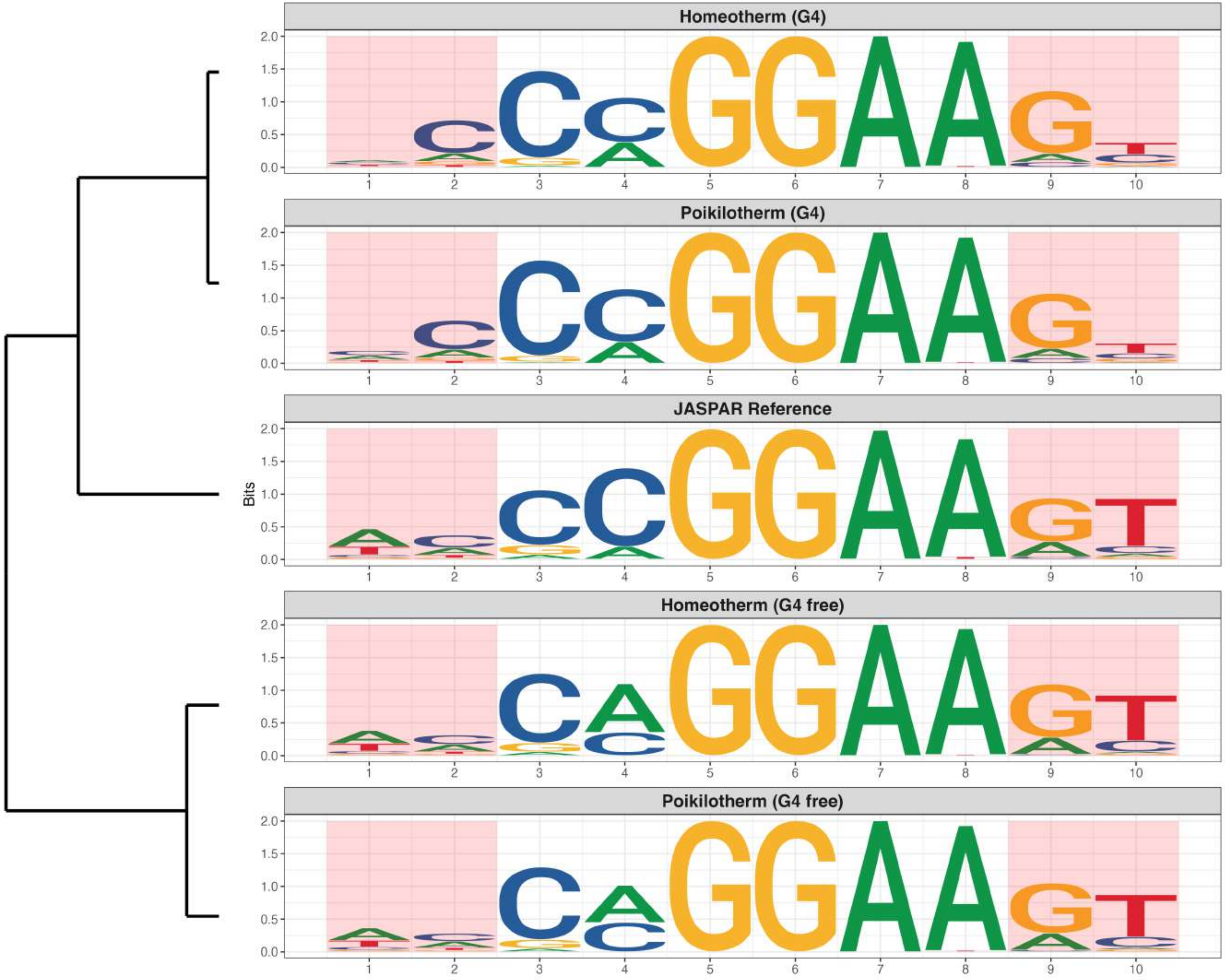

**Genome : Highly Reduced in G4s**

*MA1484.2 - ETS2*

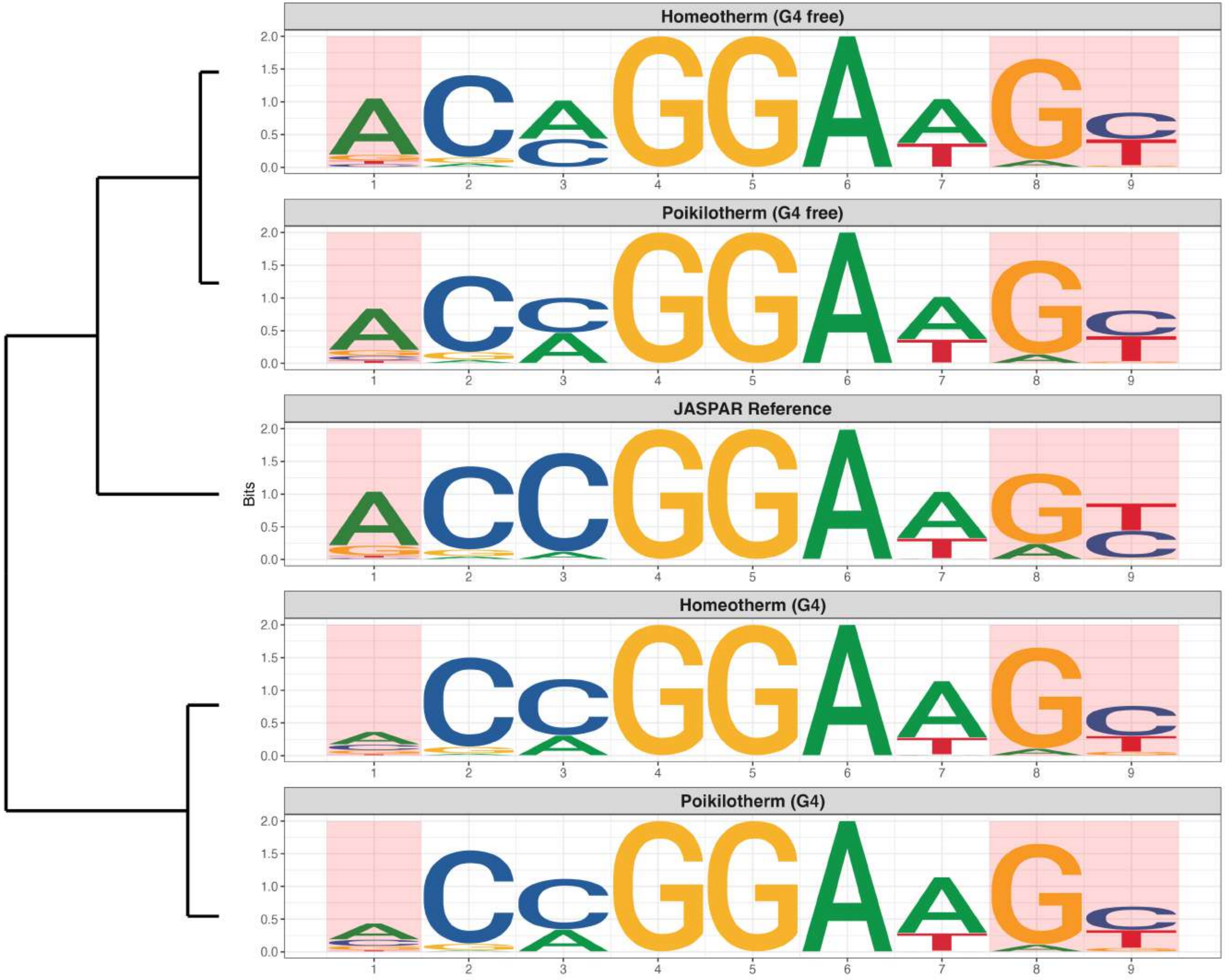

**Genome : Highly Reduced in G4s**

*MA1555.1 - RXRB*

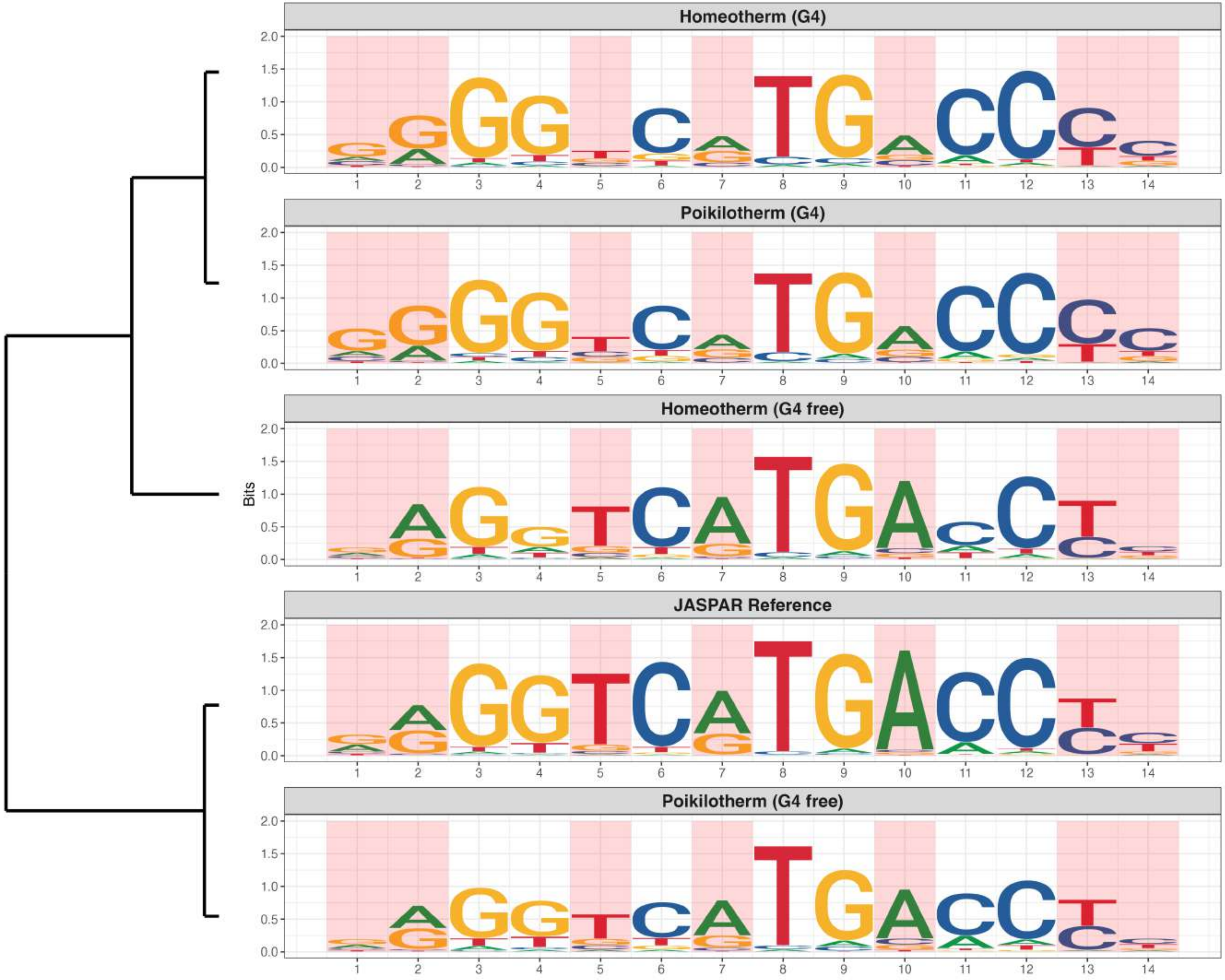

### Genome : Highly Reduced in G4s

*MA1602.2 - ZSCAN29*

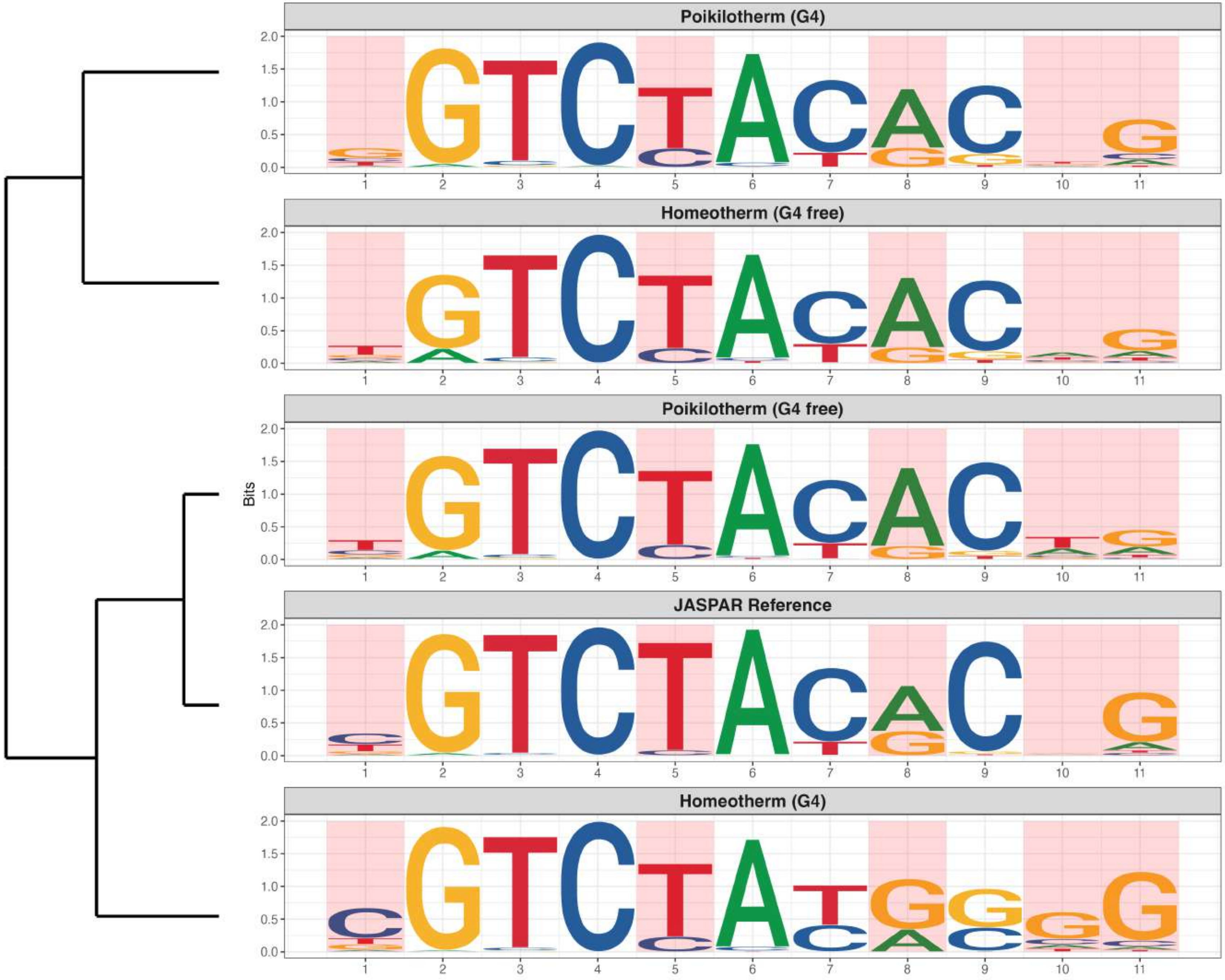

### Genome : Highly Reduced in G4s

*MA1641.2 - MYF5*

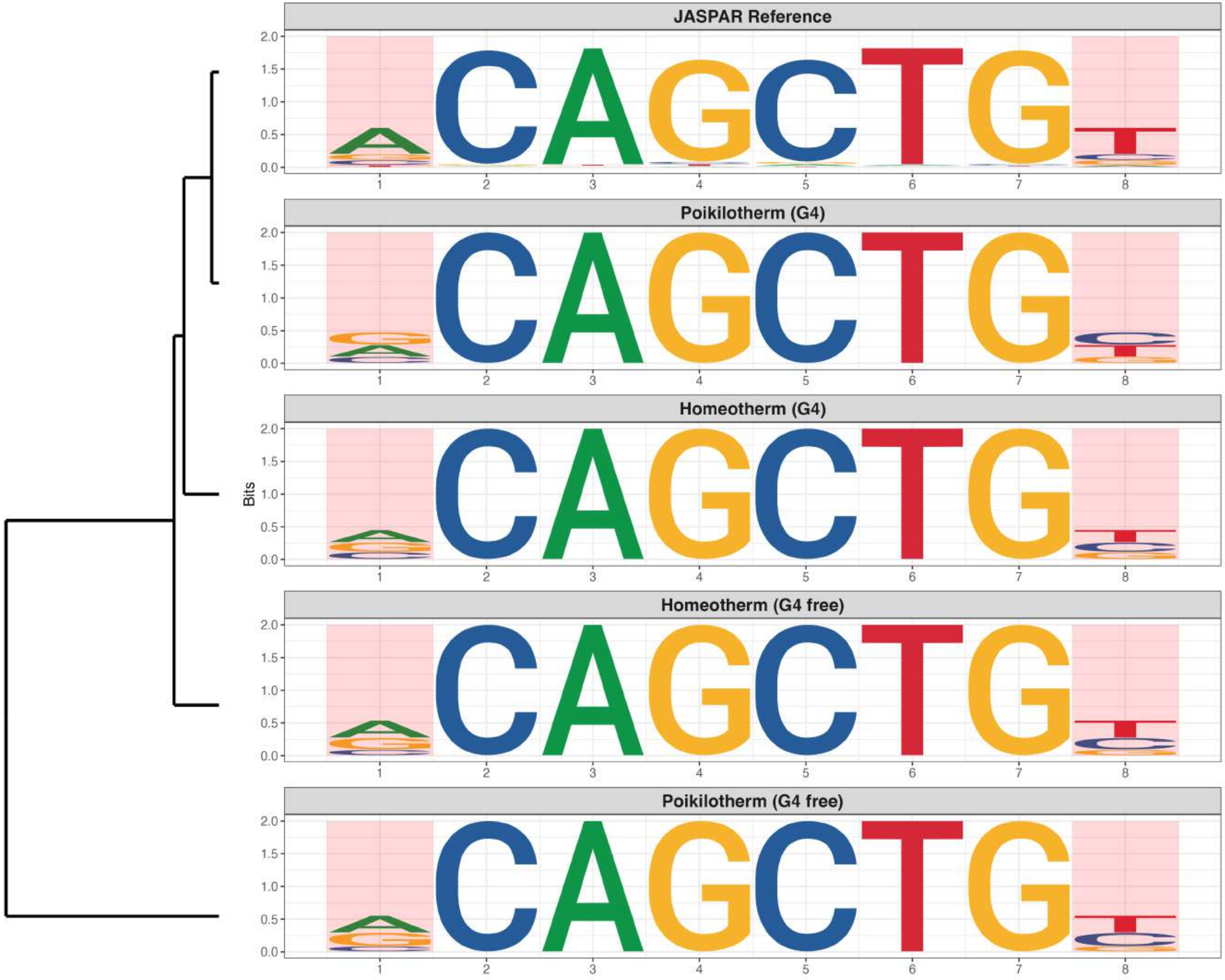

### Genome : Moderately Reduced in G4s

*MA0113.4 - NR3C1*

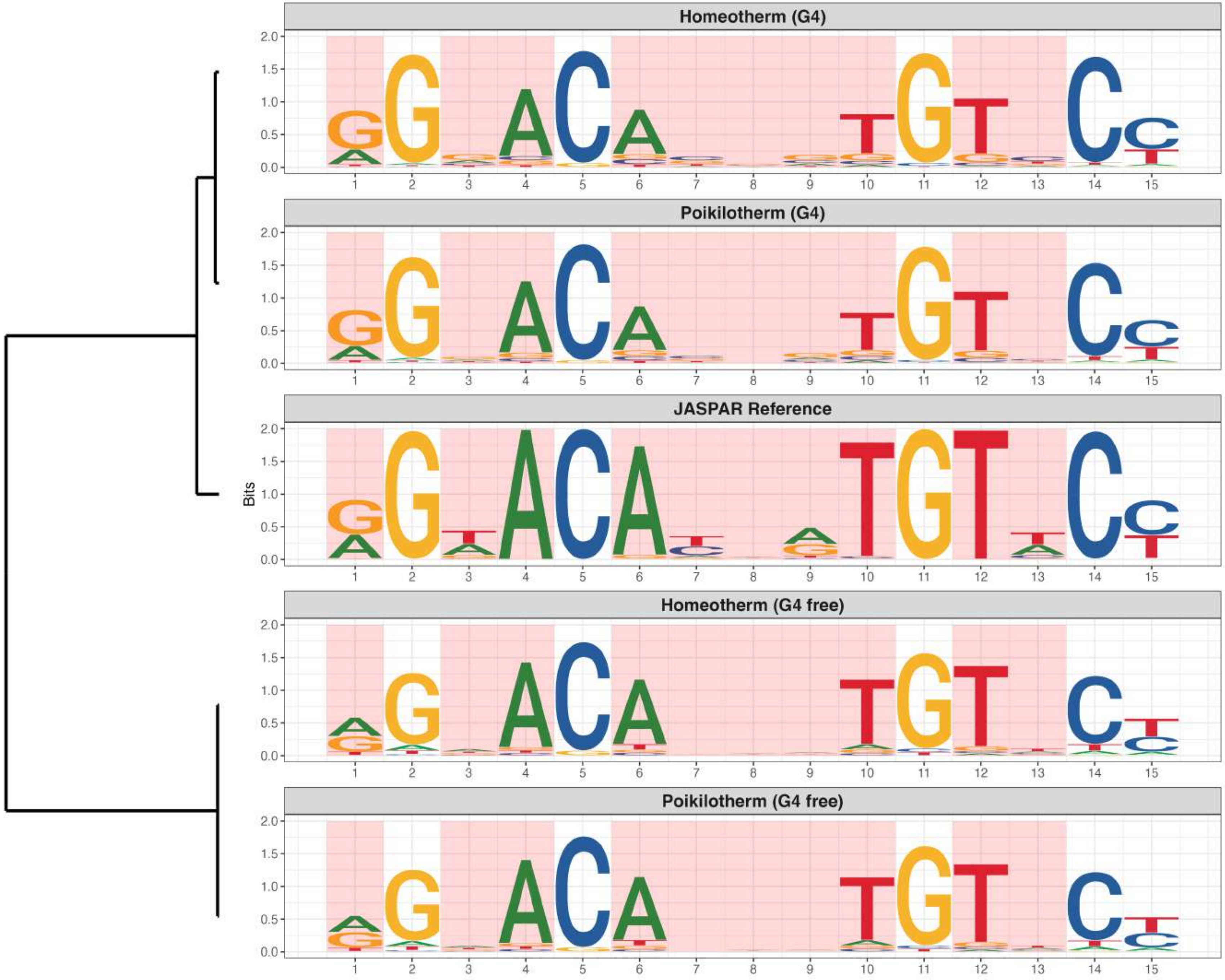

**Genome : Moderately Reduced in G4s**

*MA0143.5 - SOX2*

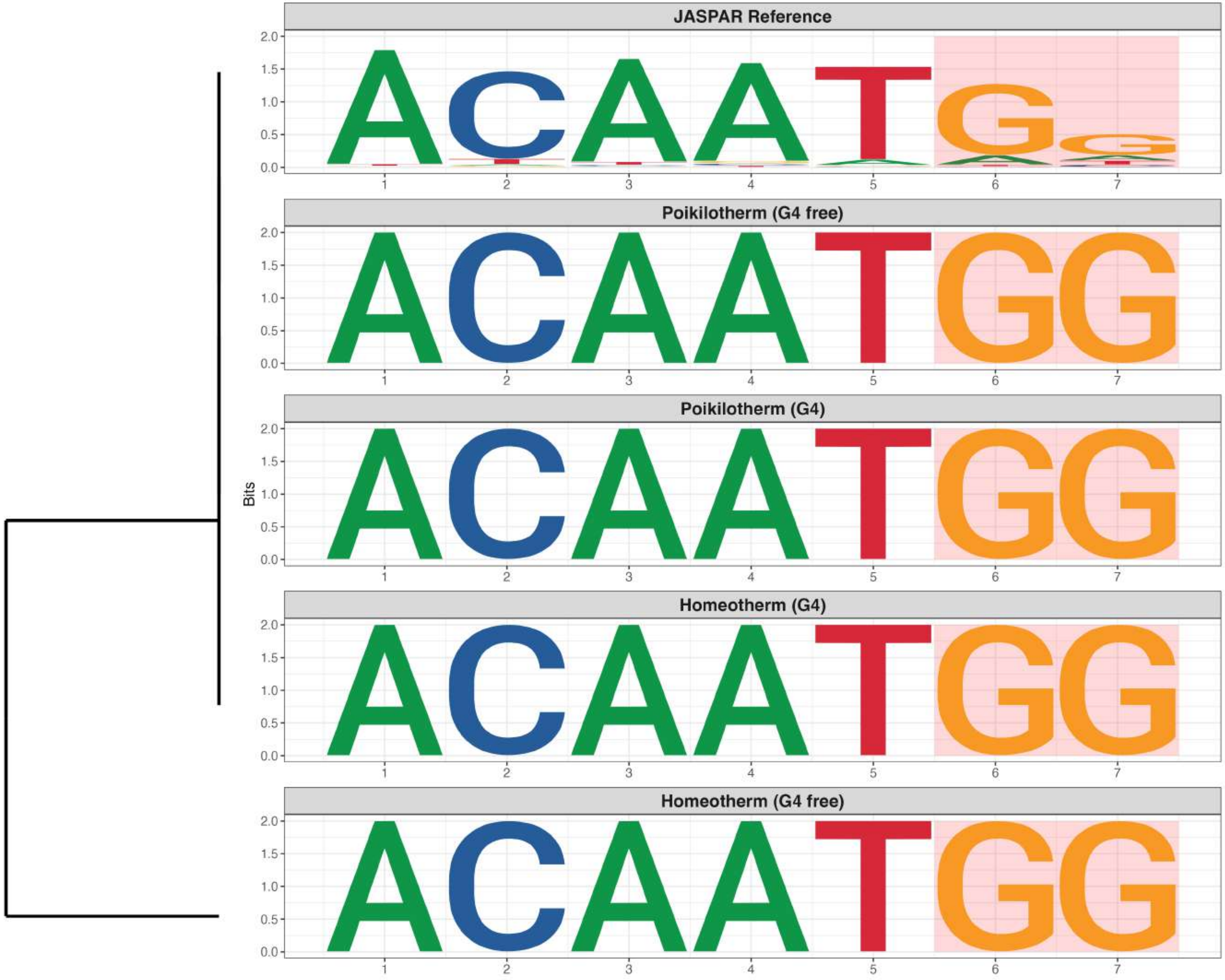

### Genome : Moderately Reduced in G4s

*MA0475.3 - FLI1*

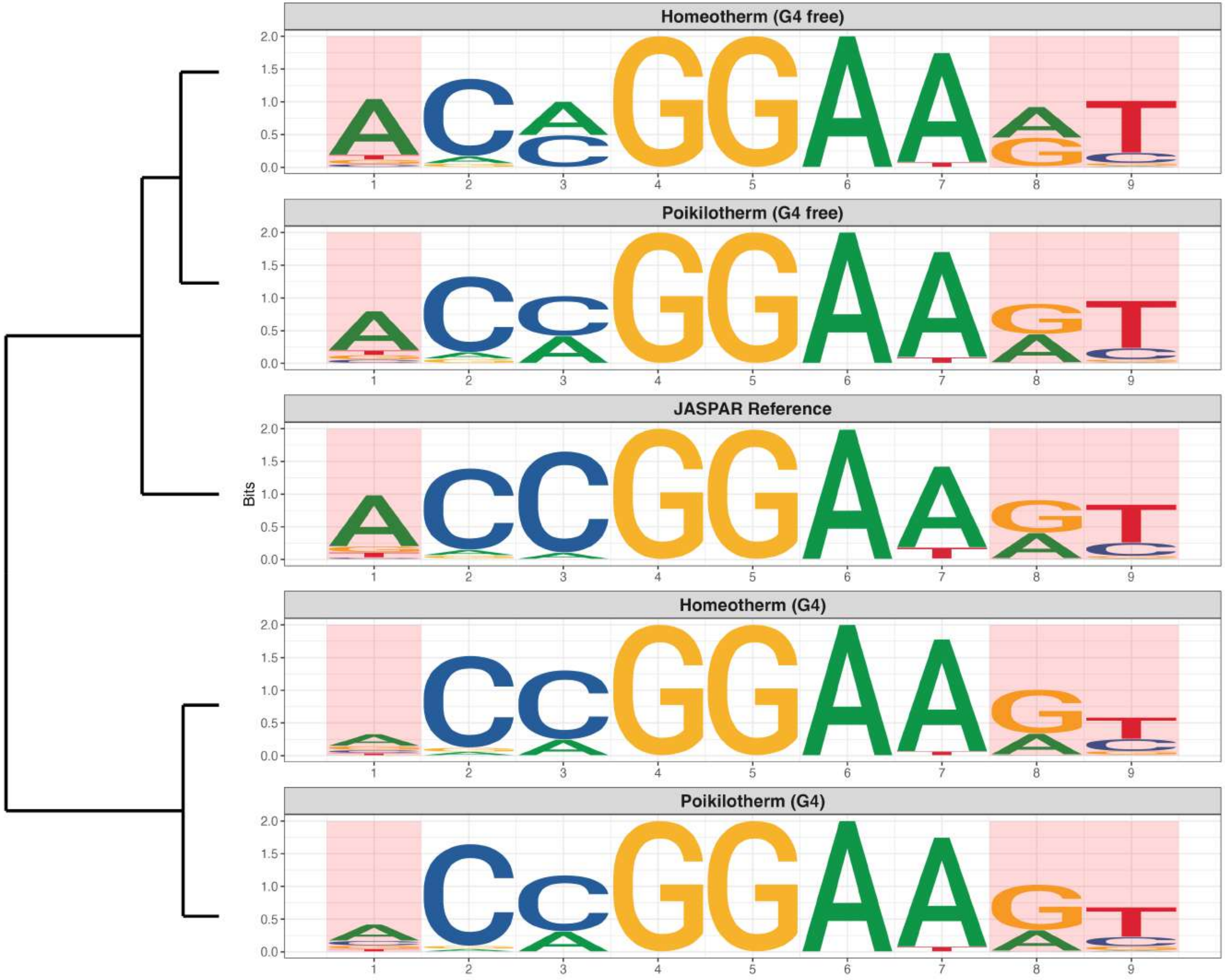

### Genome : Moderately Reduced in G4s

*MA0509.3 - RFX1*

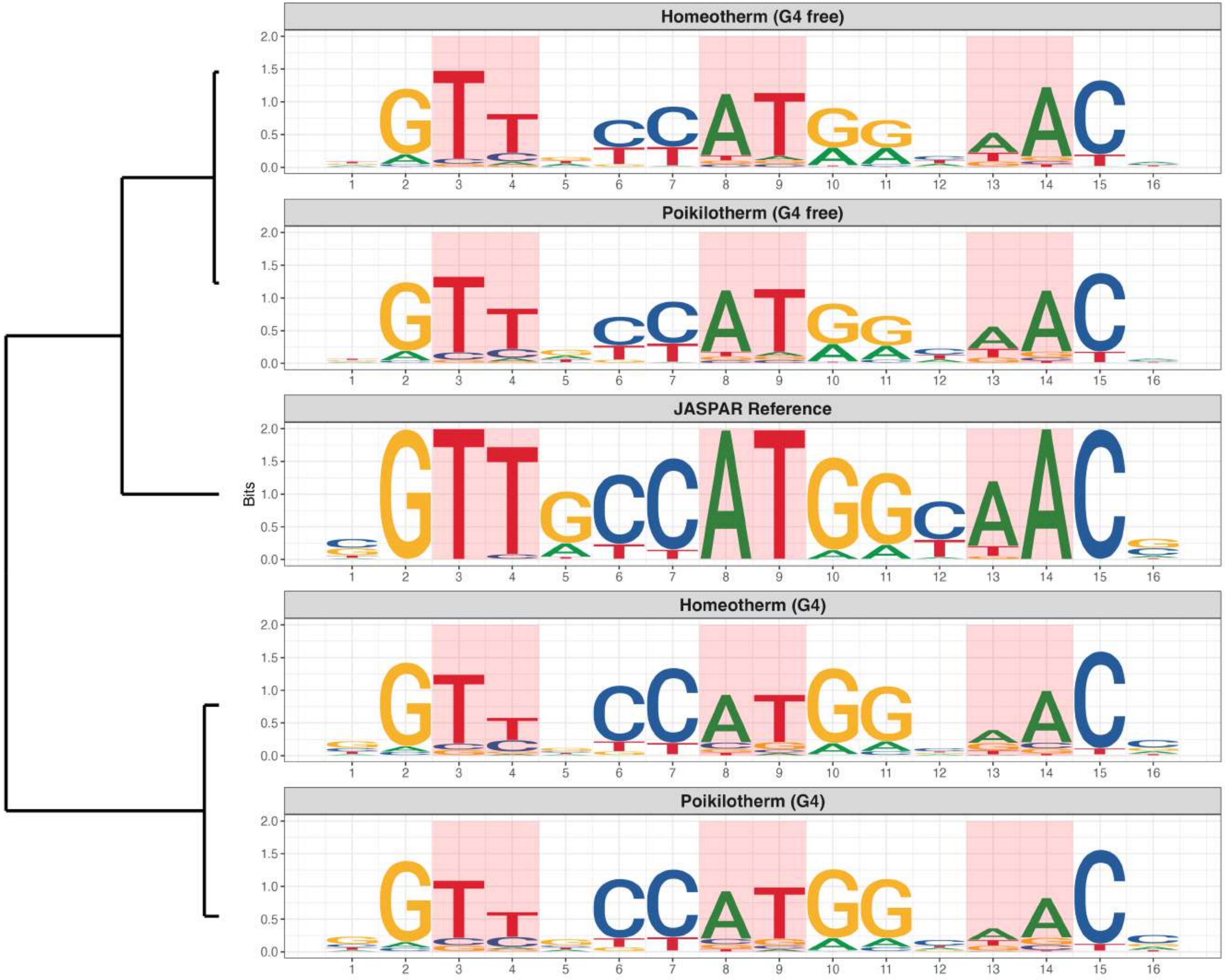

**Genome : Moderately Reduced in G4s**

*MA0514.3 - Sox3*

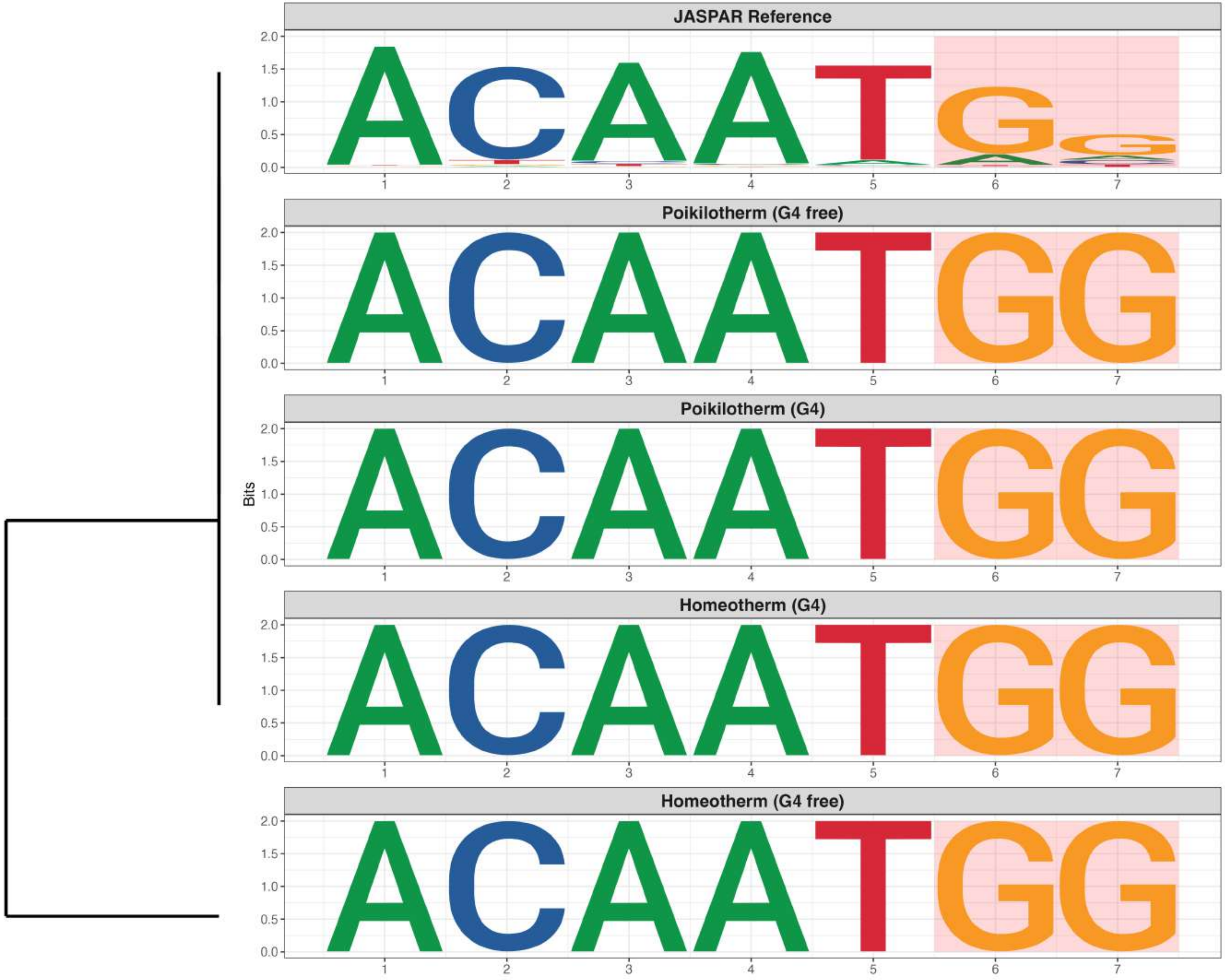

**Genome : Moderately Reduced in G4s**

*MA0594.3 - HOXA9*

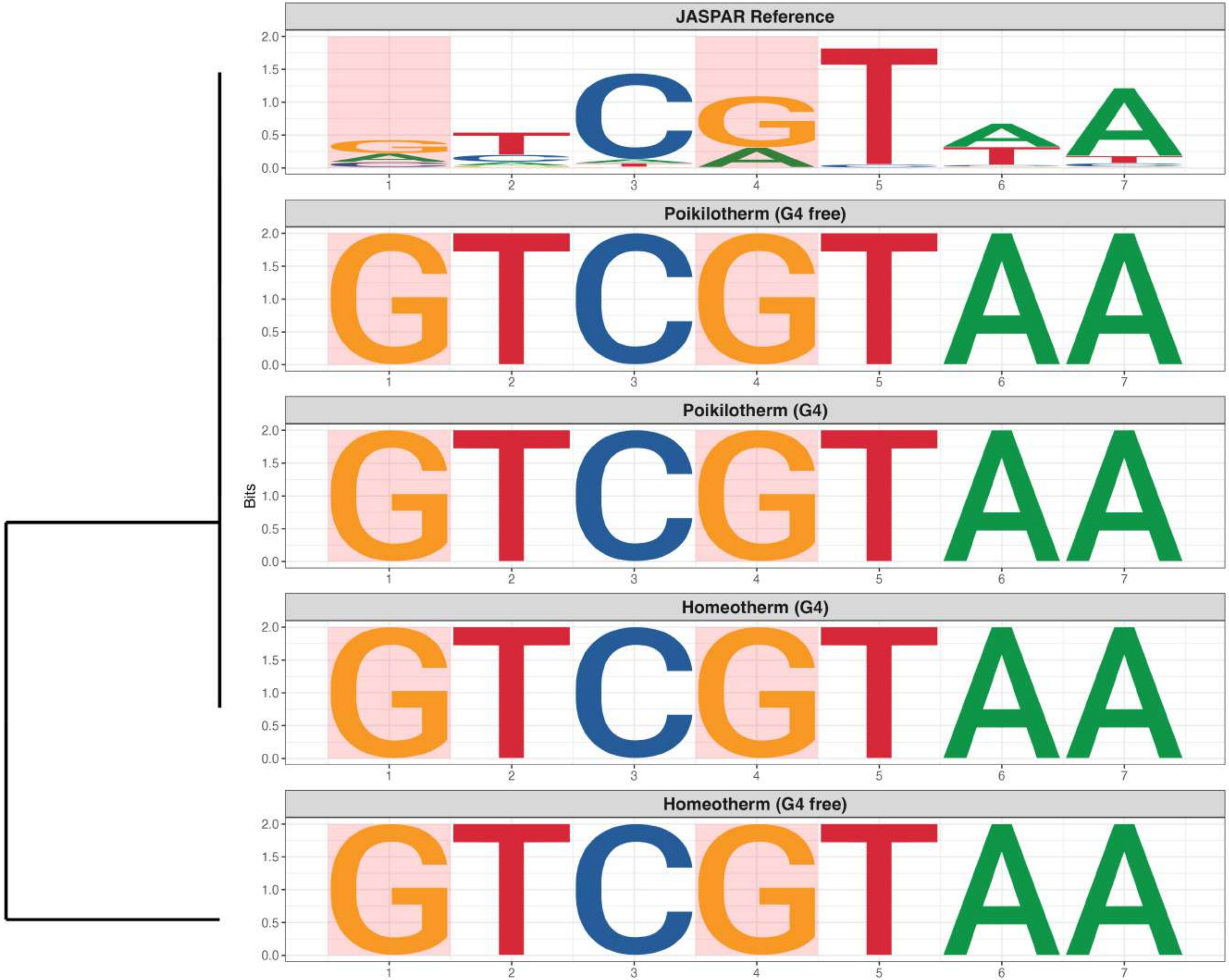

### Genome : Moderately Reduced in G4s

*MA0604.1 - Atf1*

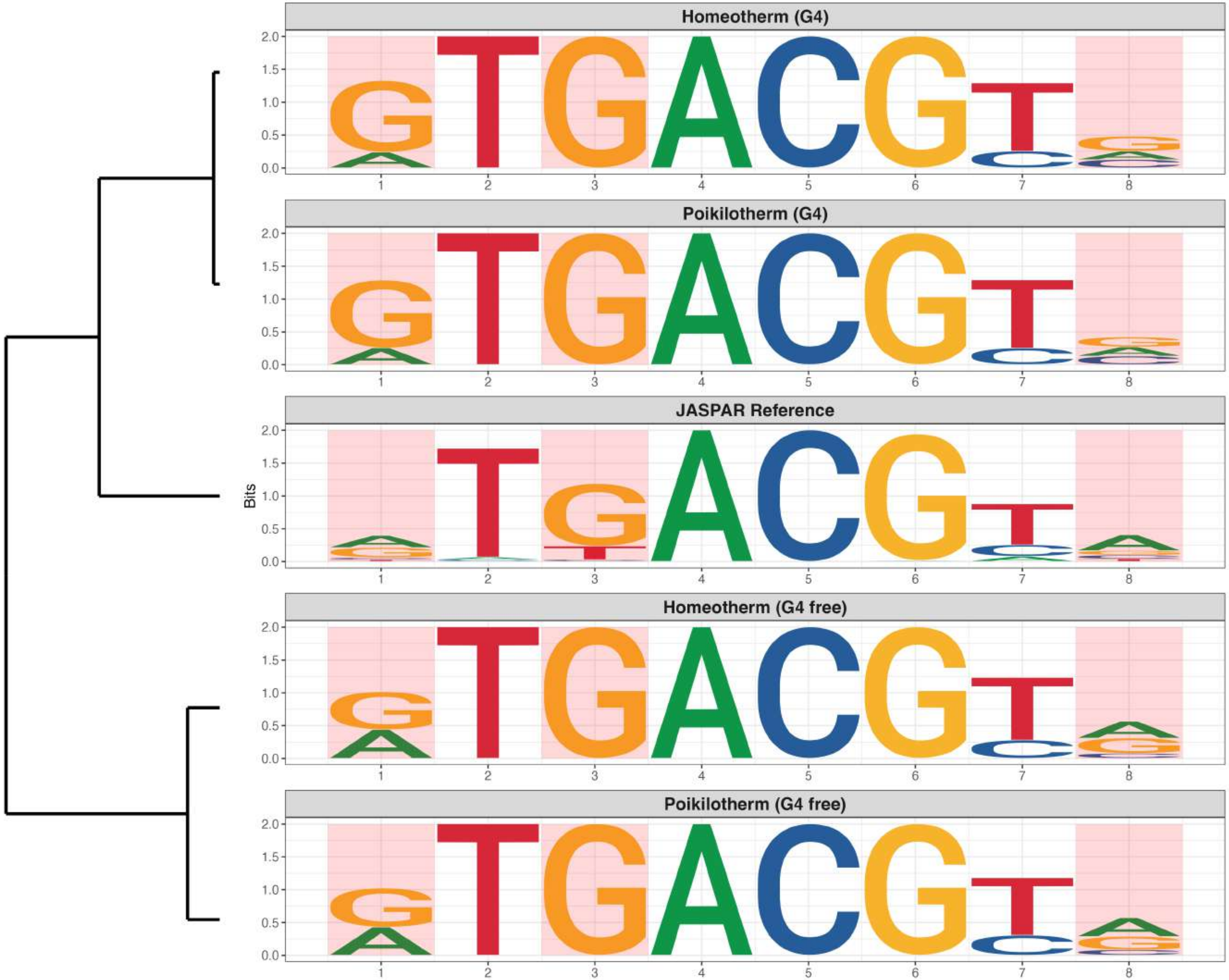

### Genome : Moderately Reduced in G4s

*MA0749.2 - ZBED1*

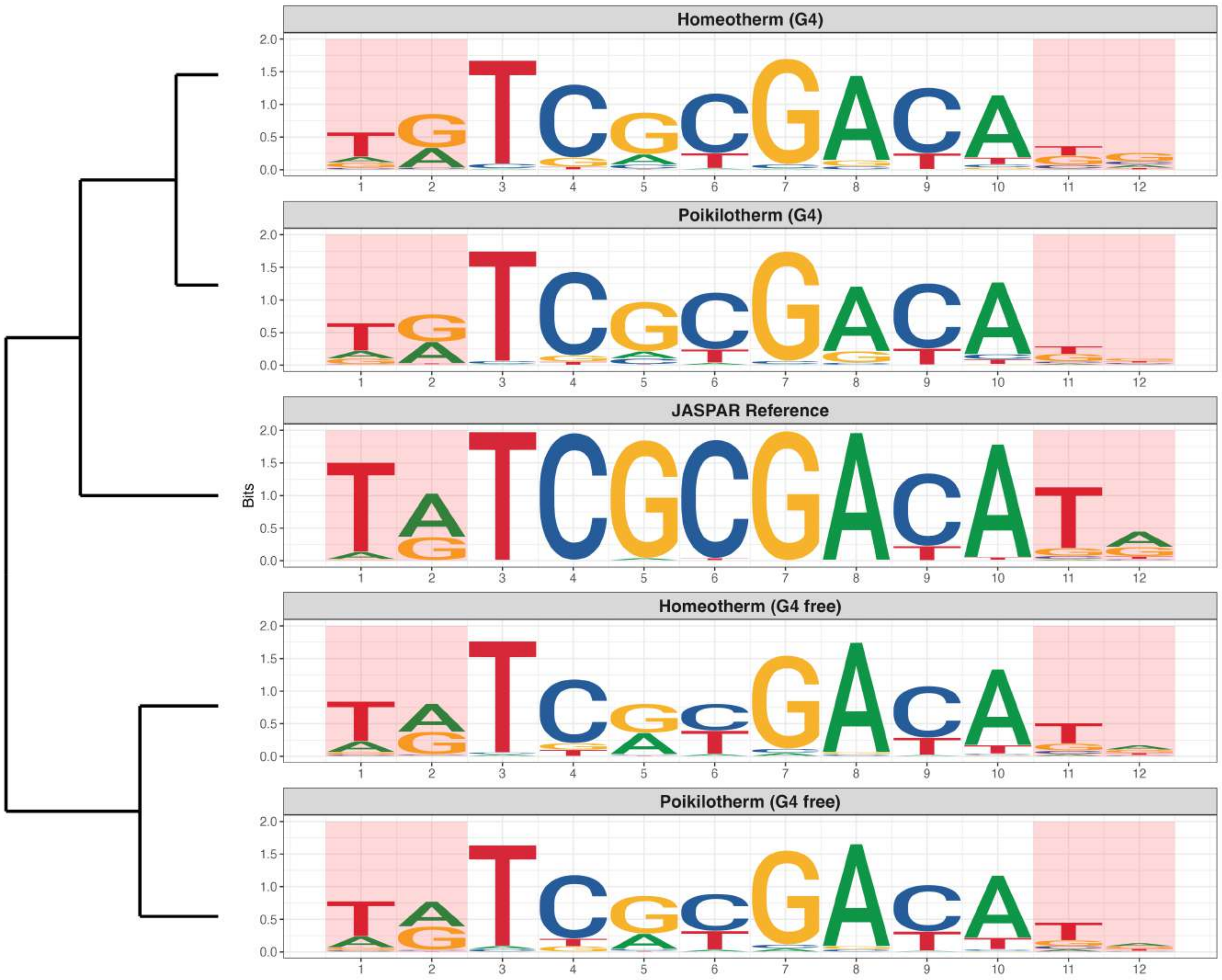

### Genome : Moderately Reduced in G4s

*MA0799.3 - RFX4*

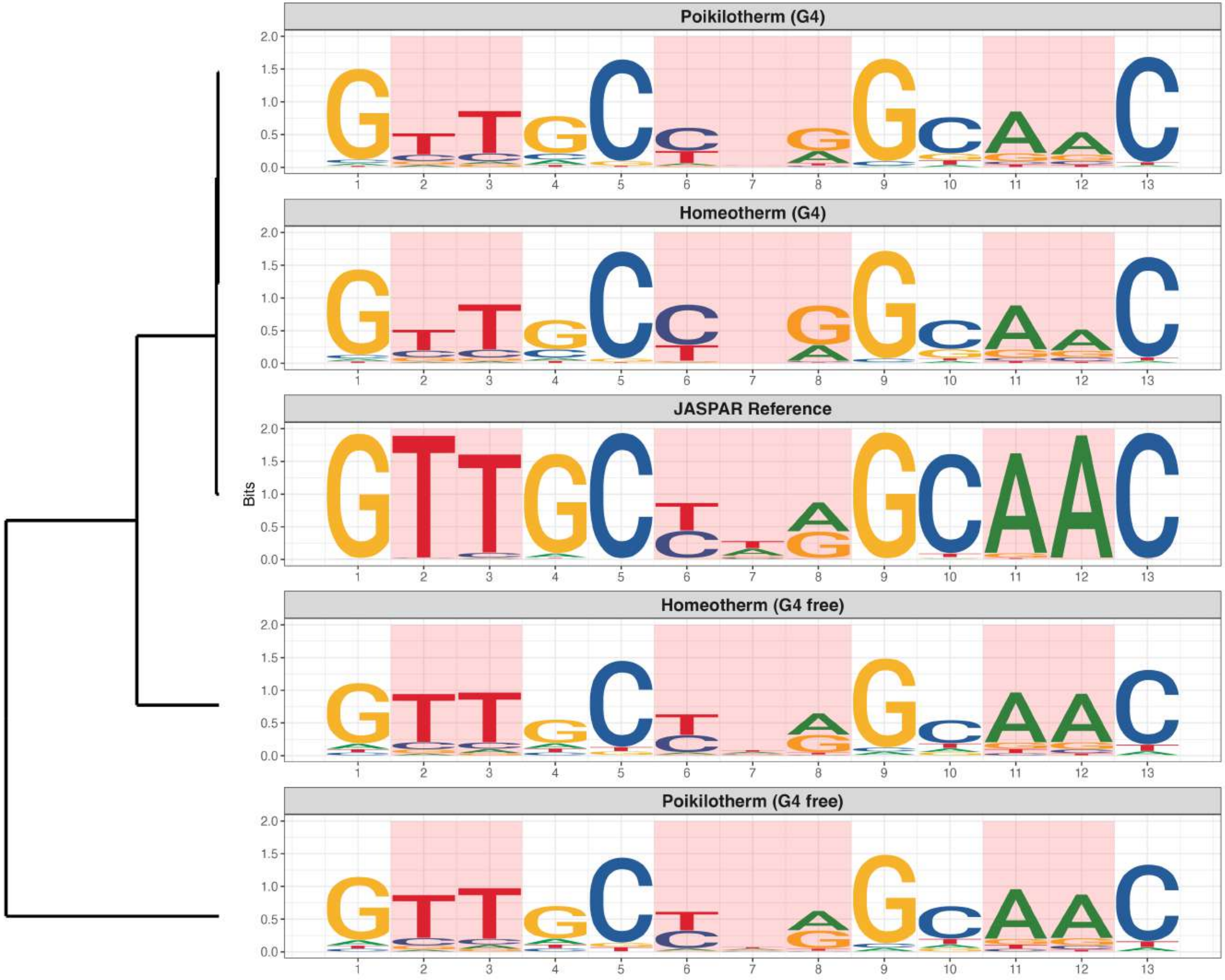

**Genome : Moderately Reduced in G4s**

*MA1120.2 - SOX13*

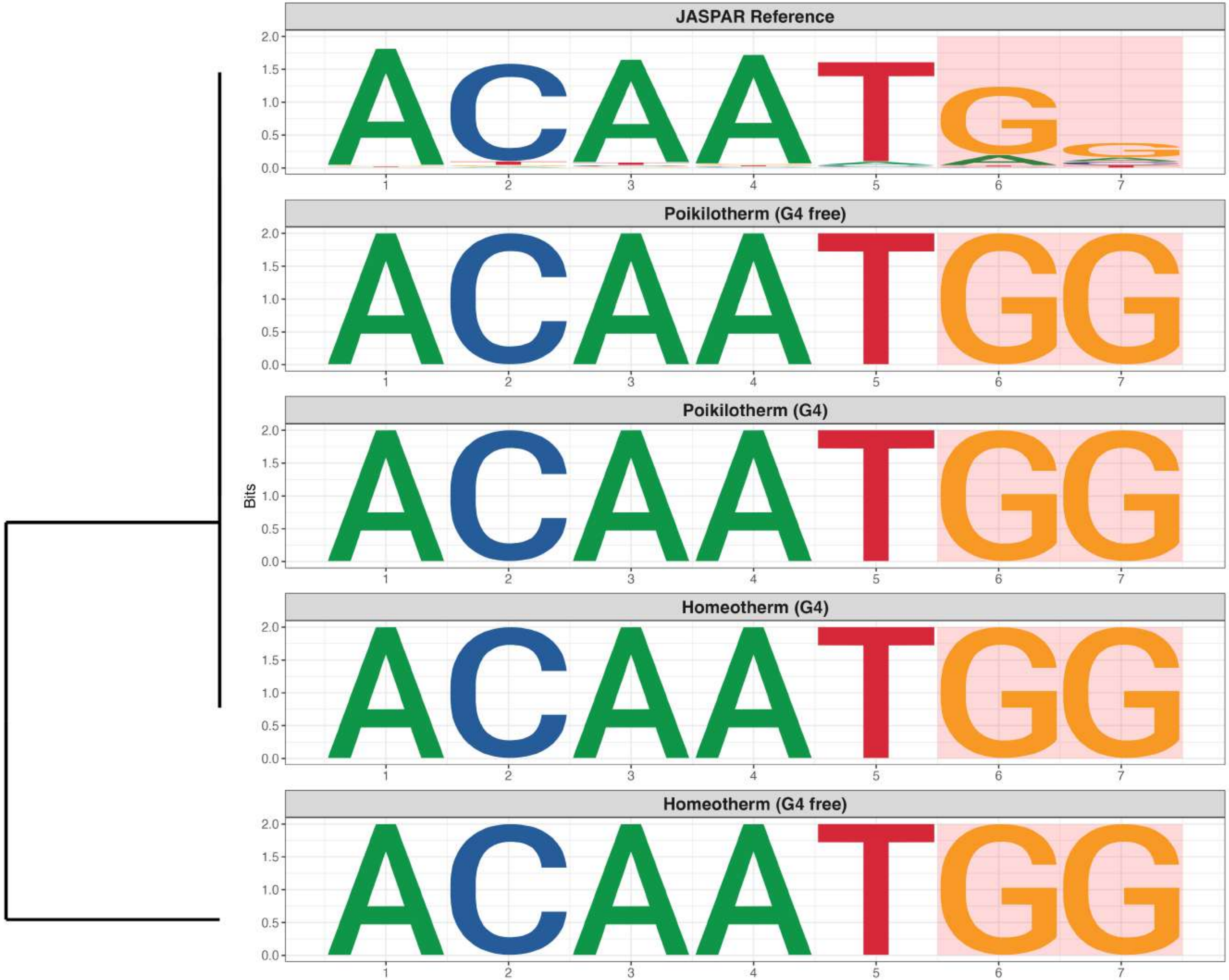

**Genome : Moderately Reduced in G4s**

*MA1532.2 - NR1D2*

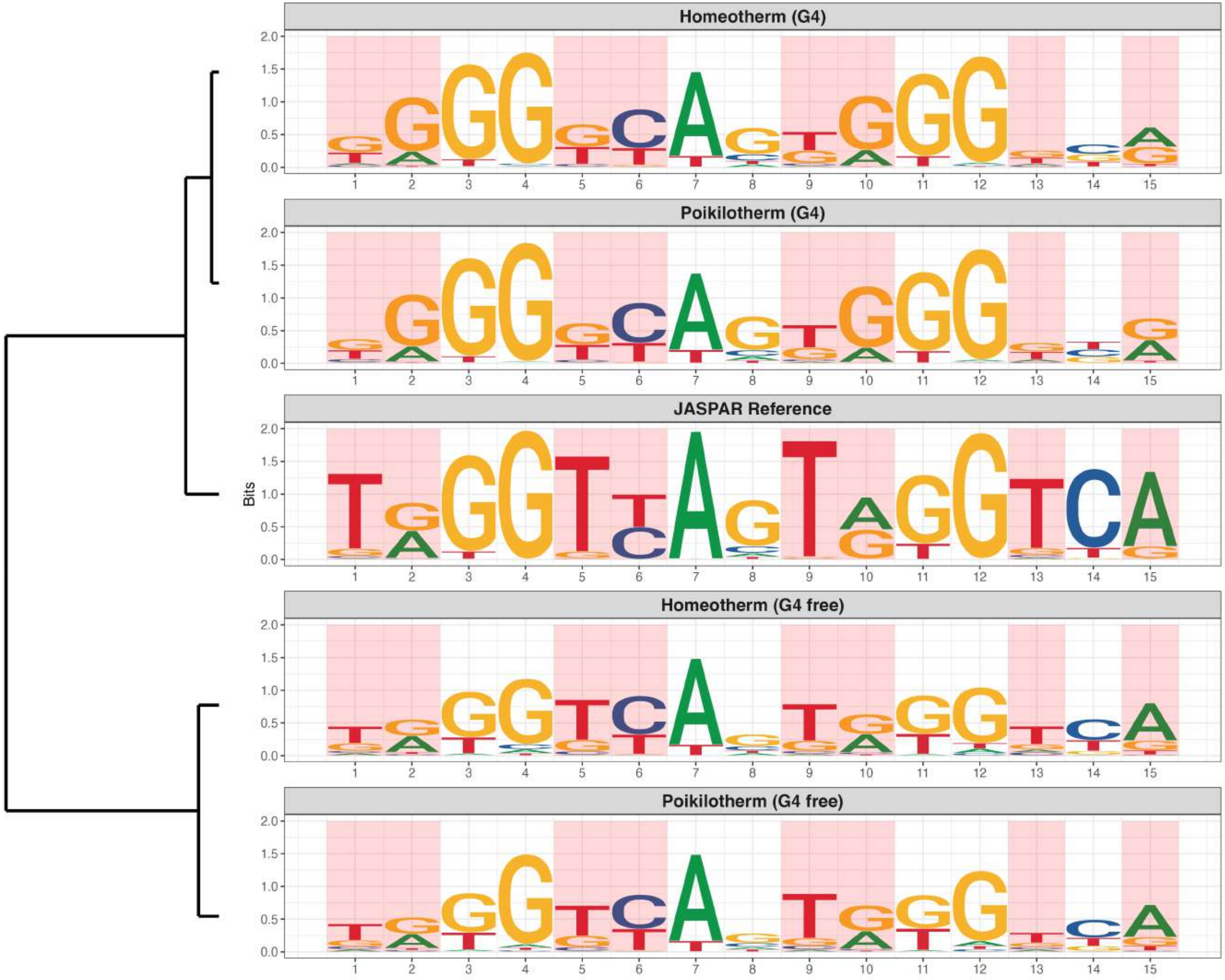

**Genome : Moderately Reduced in G4s**

*MA1544.2 - OVOL1*

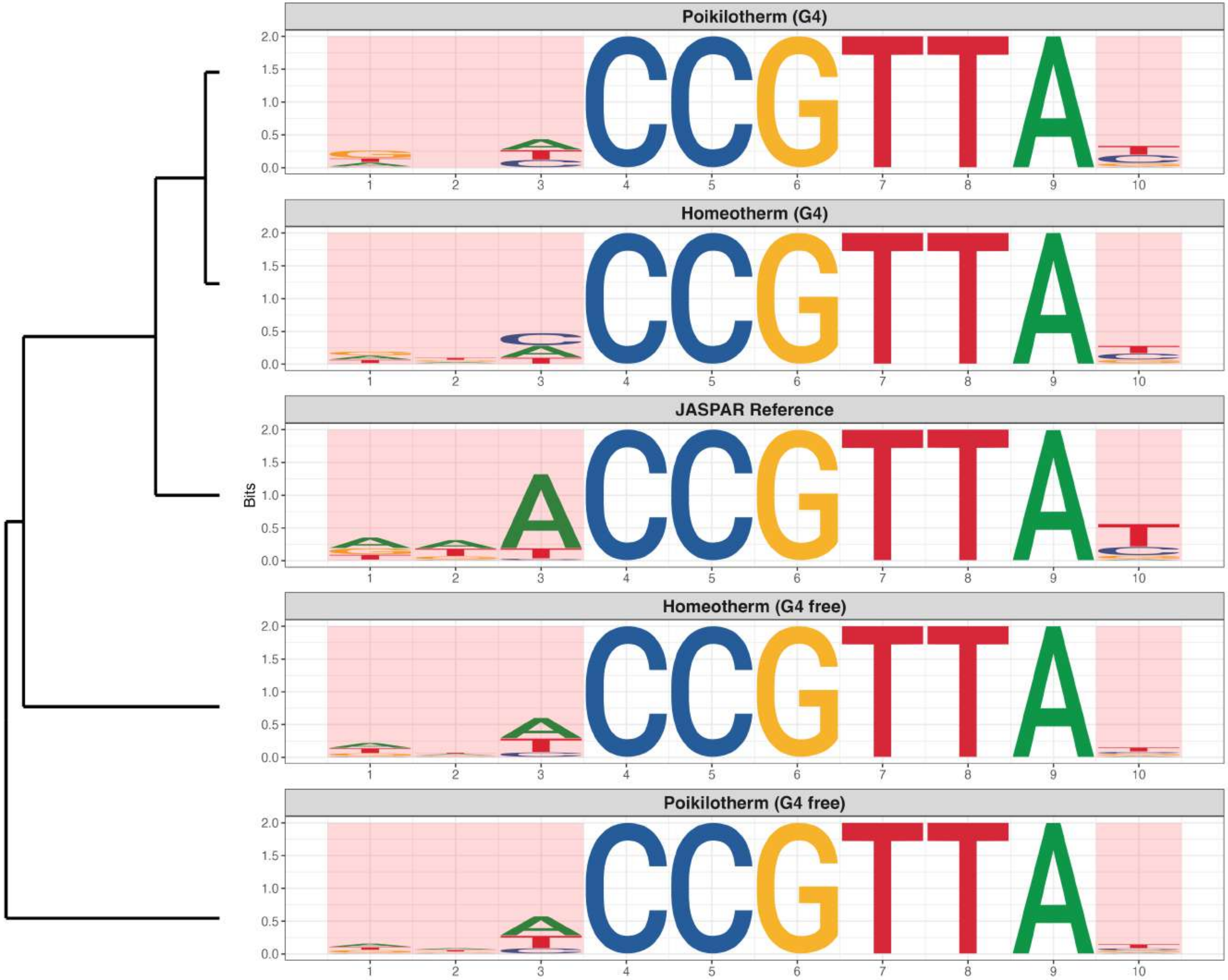

**Genome : Moderately Reduced in G4s**

*MA1545.2 - OVOL2*

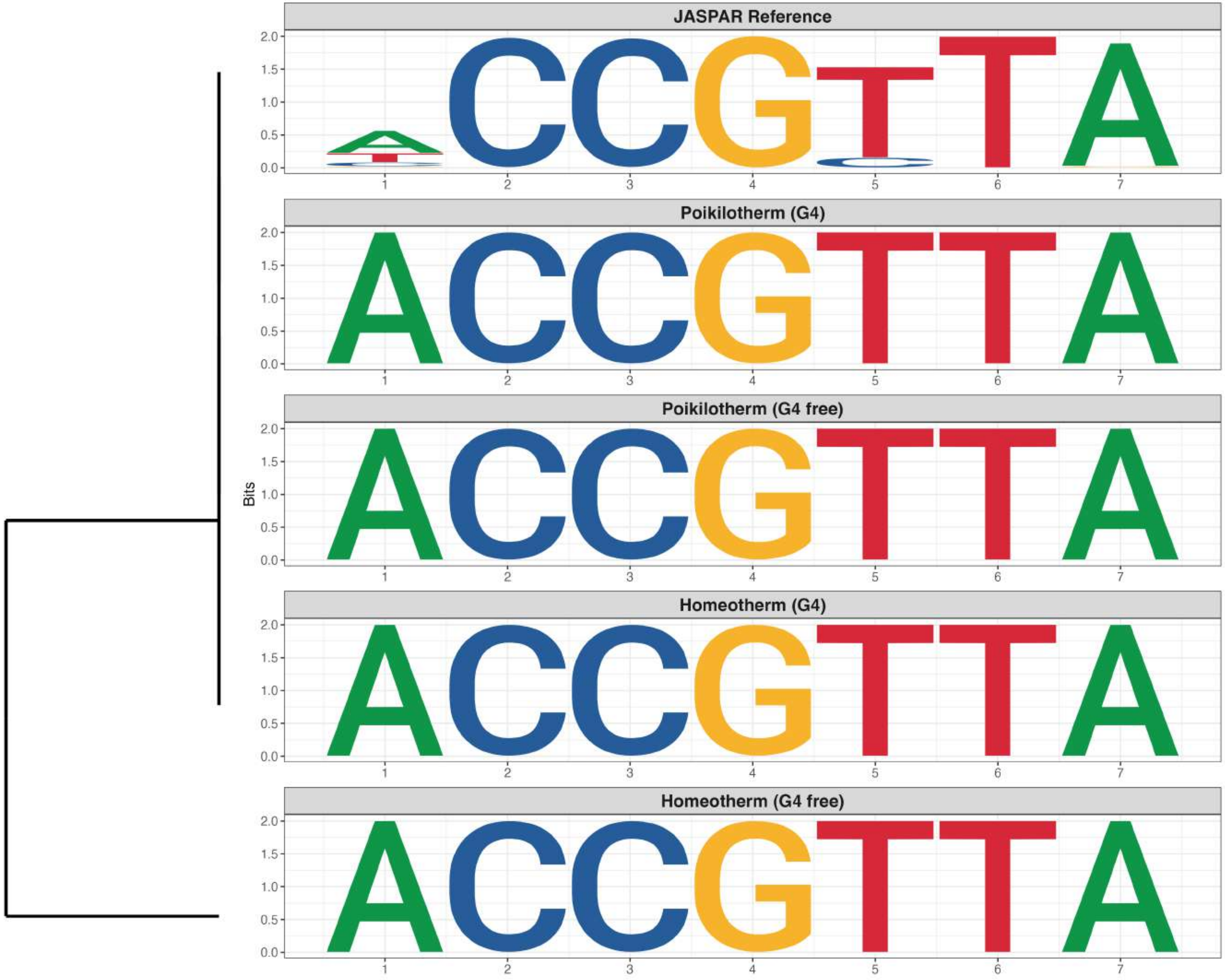

### Genome : Moderately Reduced in G4s

*MA1554.2 - RFX7*

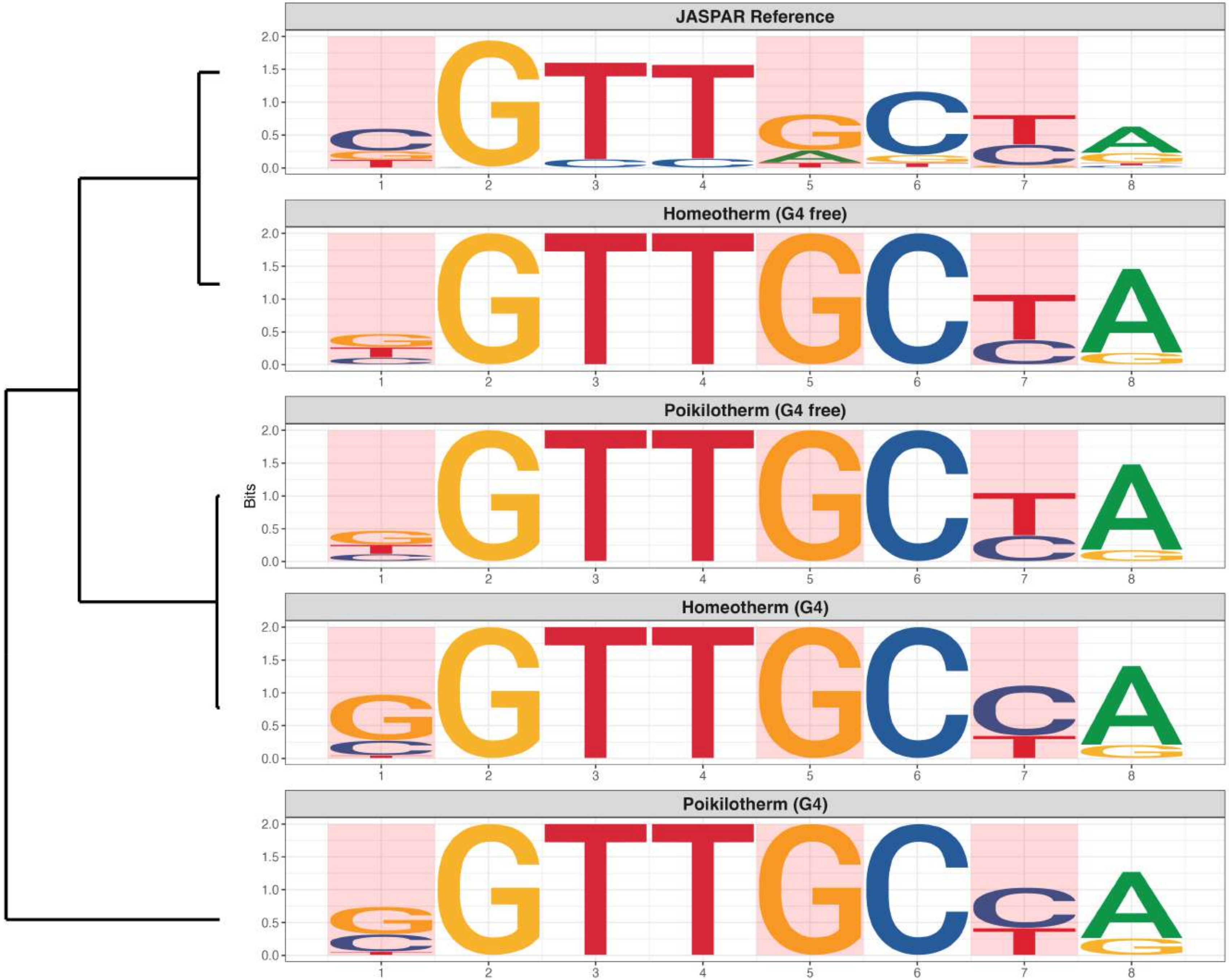

**Genome : Moderately Reduced in G4s**

*MA1589.3 - ZNF140*

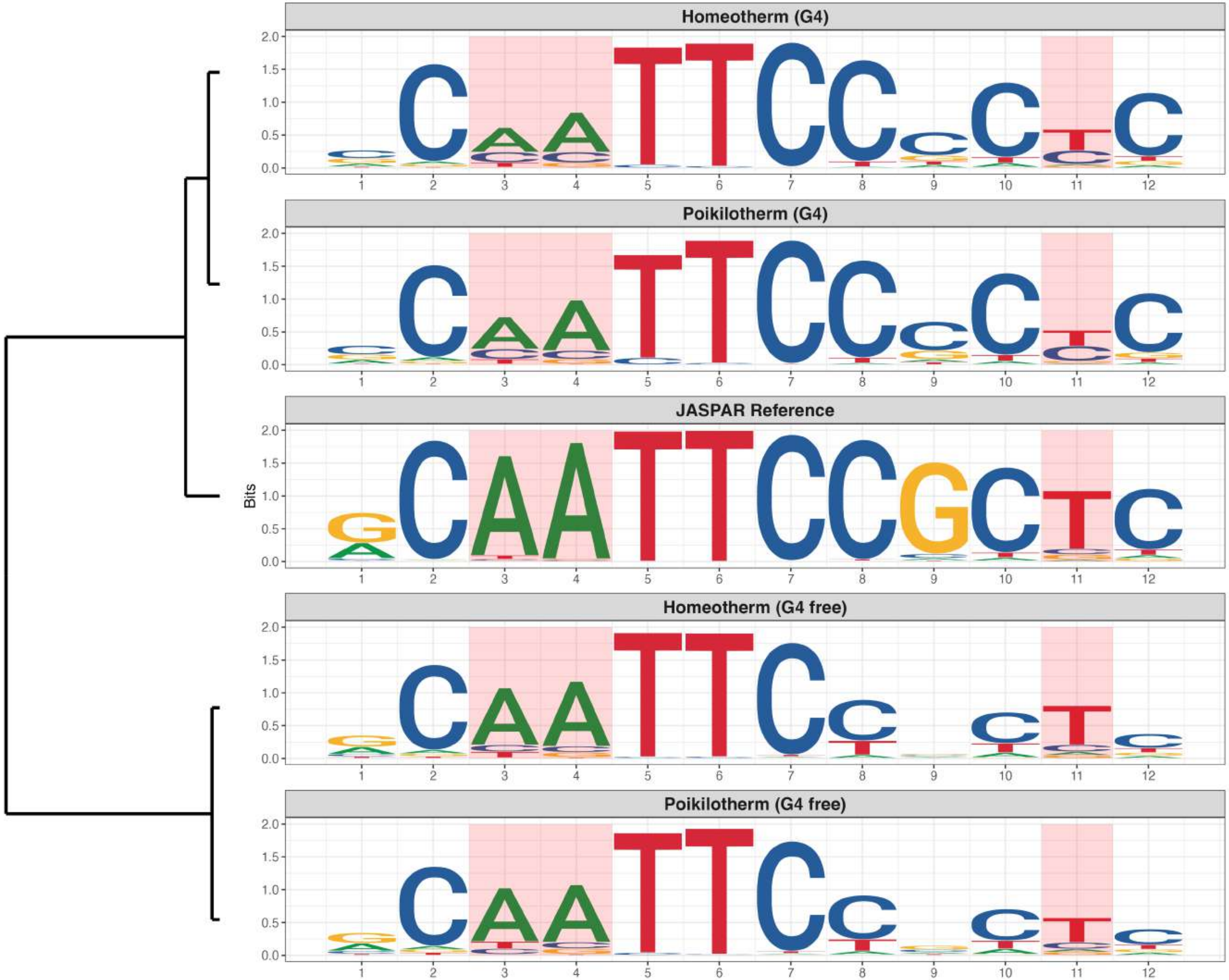

**Genome : Moderately Reduced in G4s**

*MA1619.2 - Ptf1A*

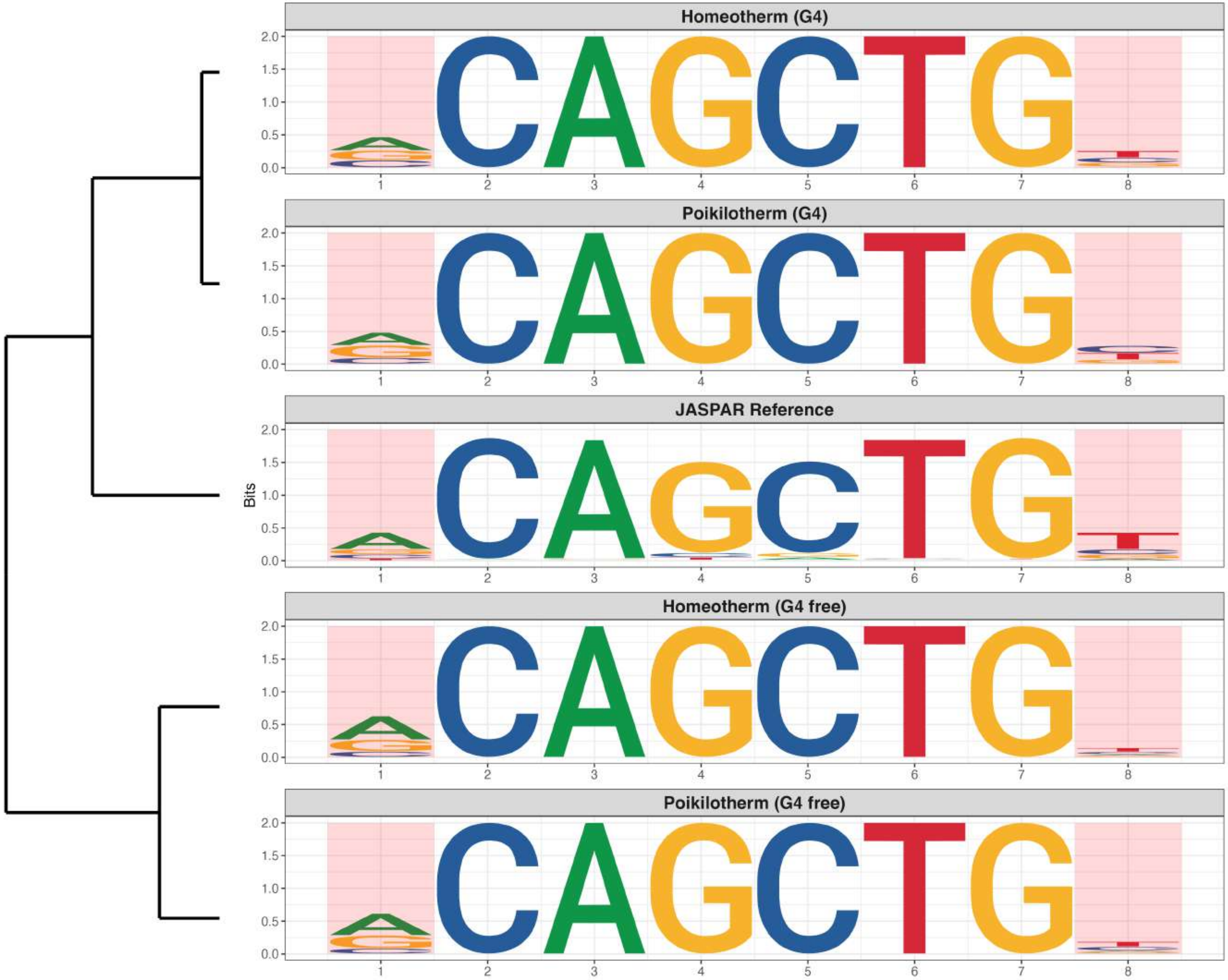

**Genome : Moderately Reduced in G4s**

*MA2102.1 - ZBTB17*

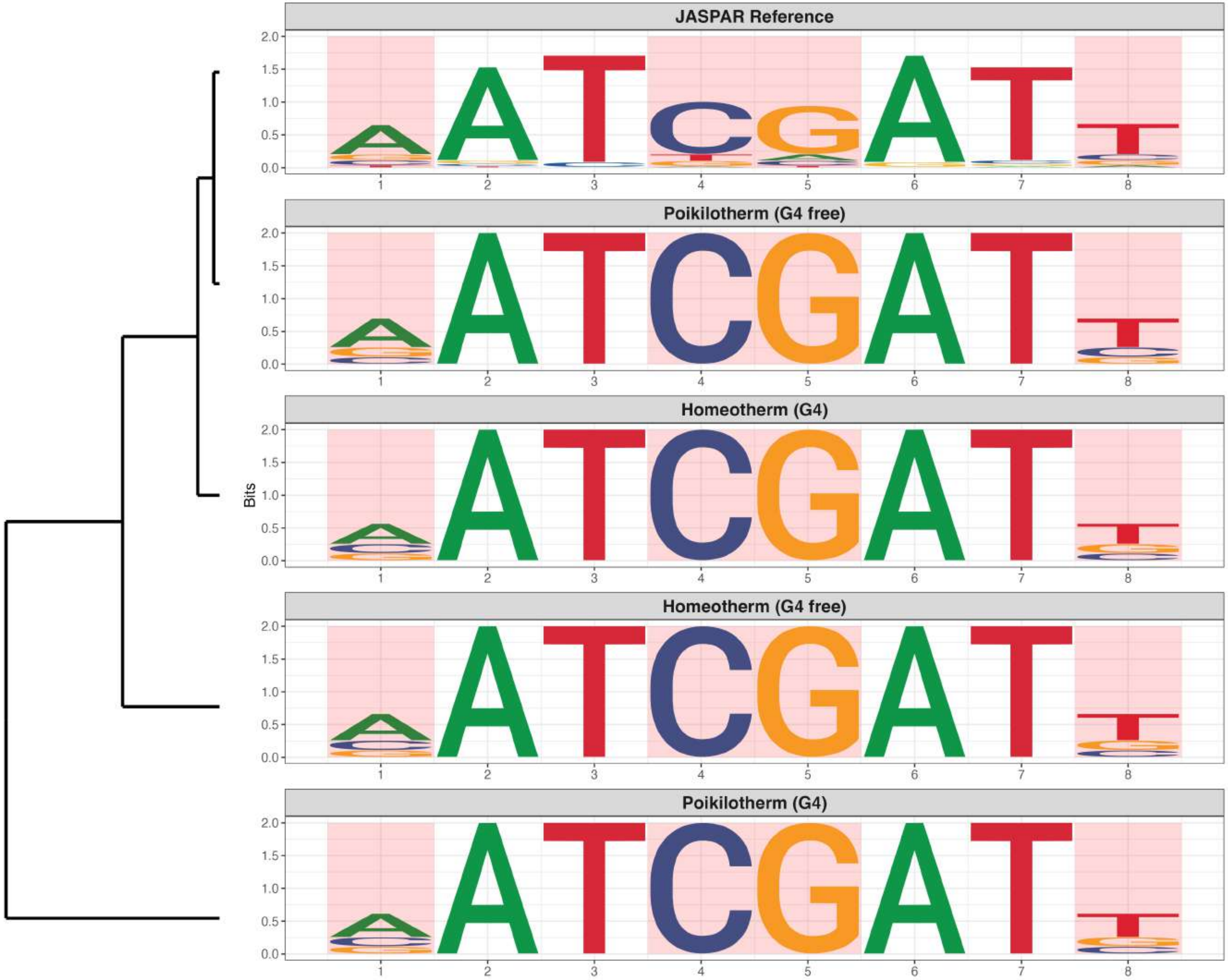

**Genome : Moderately Reduced in G4s**

*MA2590.1 - ZNF841*

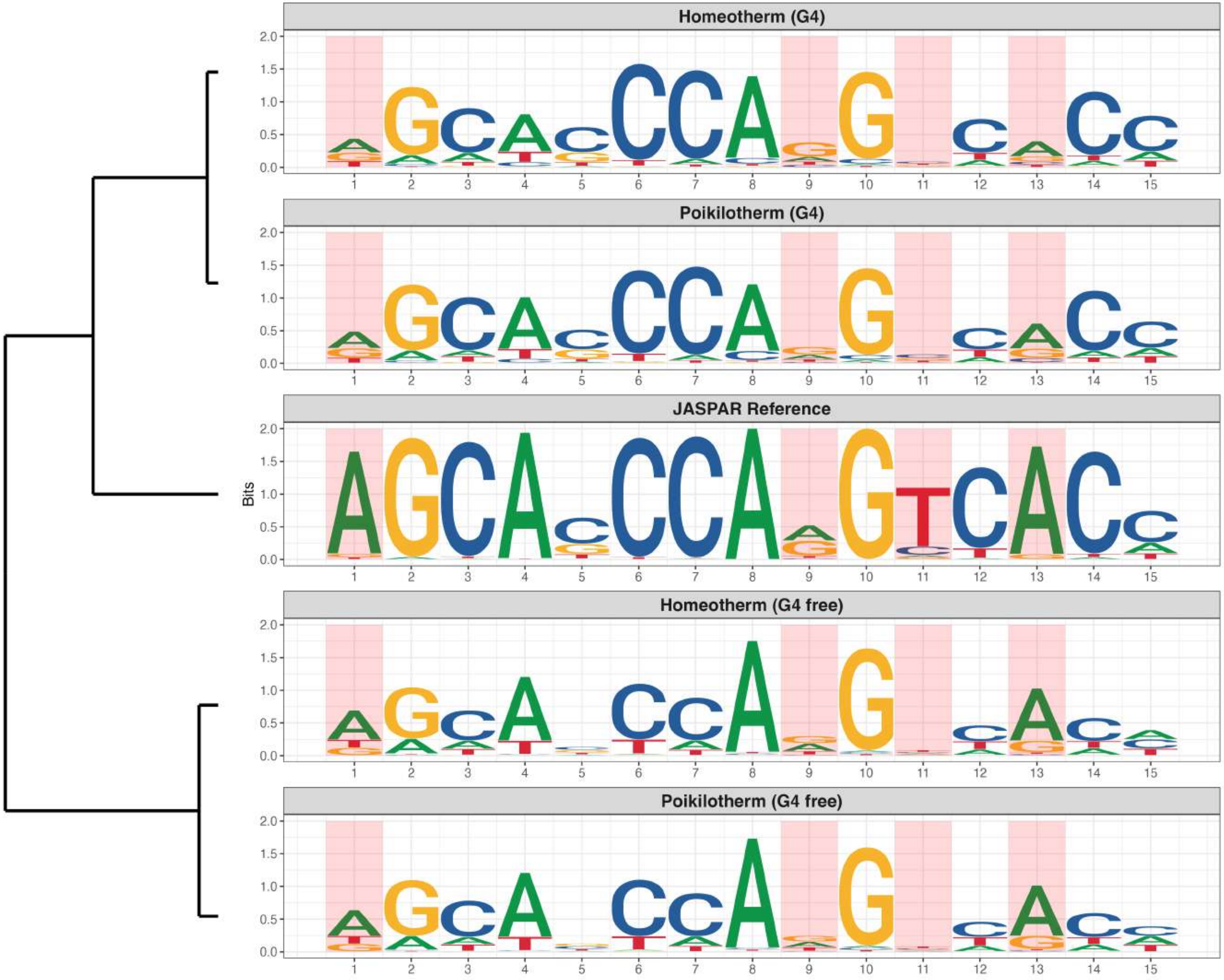
