## Supplementary material for "CGGBP1-regulated heterogeneous C–T transition rates relate with G-quadruplex potential of terrestrial vertebrate genomes": Fig. S21

### Promoter : Highly Enriched in G4s

*MA0145.2 - Tfcp2l1*

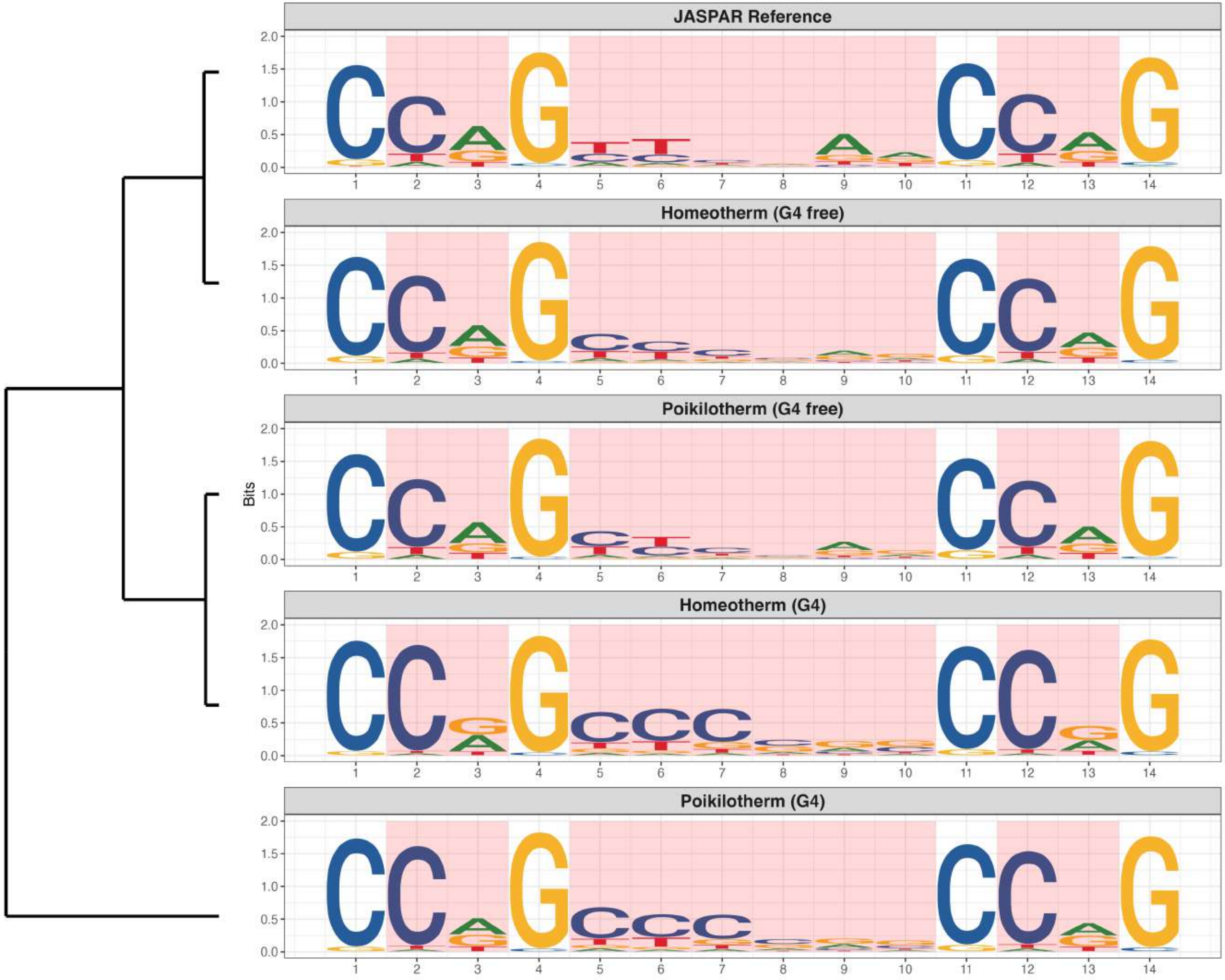

**Promoter : Highly Enriched in G4s**

*MA0258.2 - ESR2*

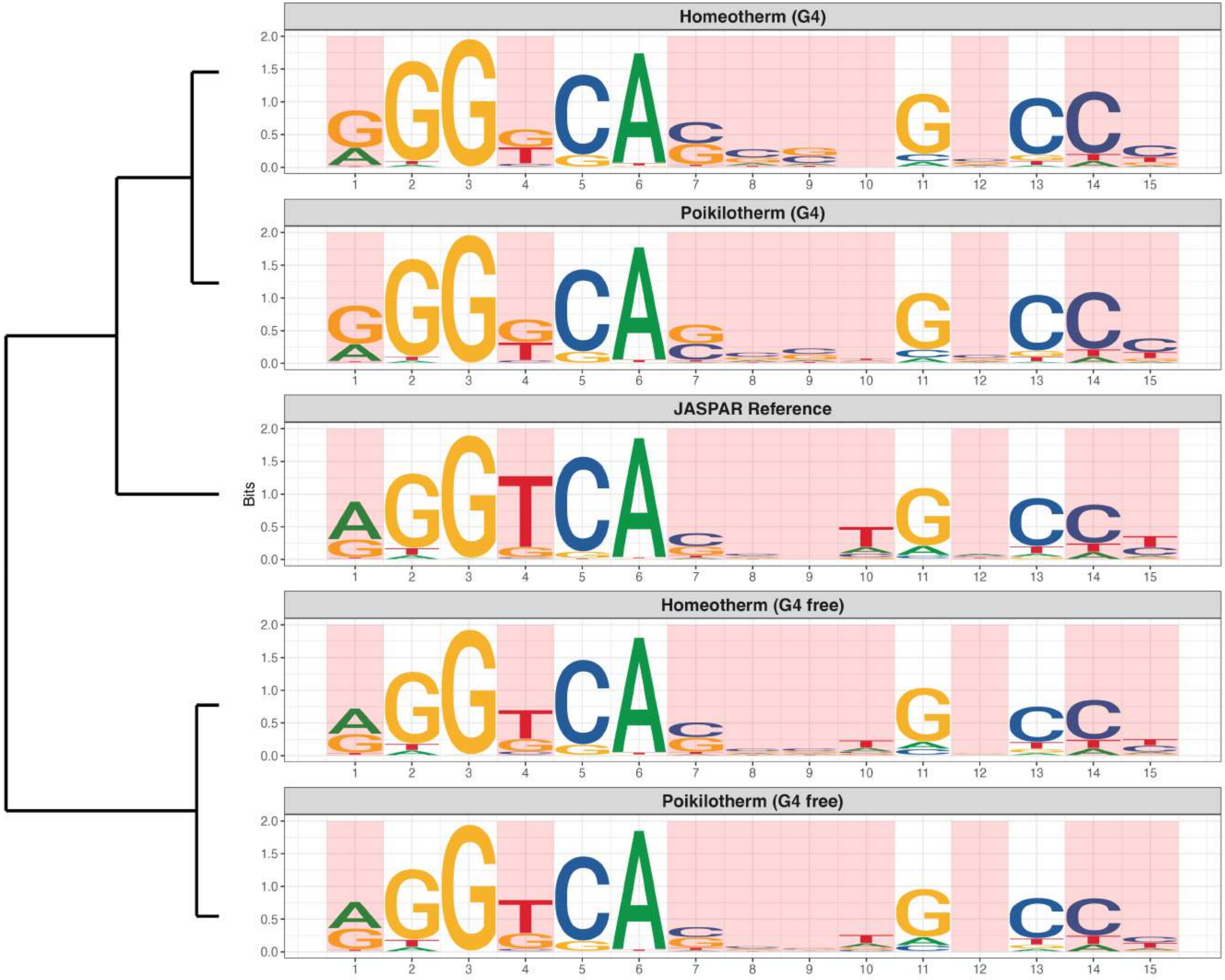

### Promoter : Highly Enriched in G4s

*MA0506.3 - Nrf1*

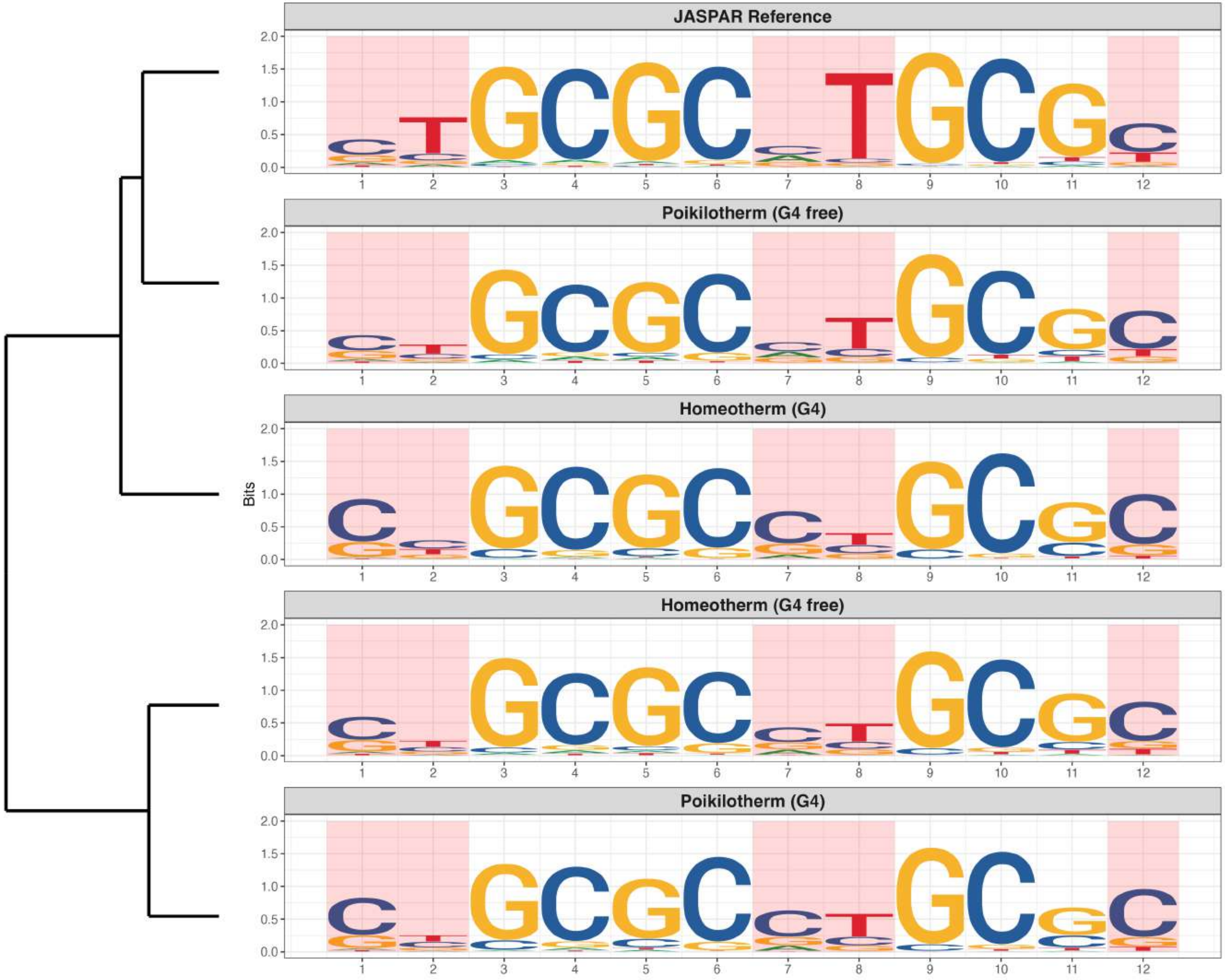

### Promoter : Highly Enriched in G4s

*MA0508.4 - PRDM1*

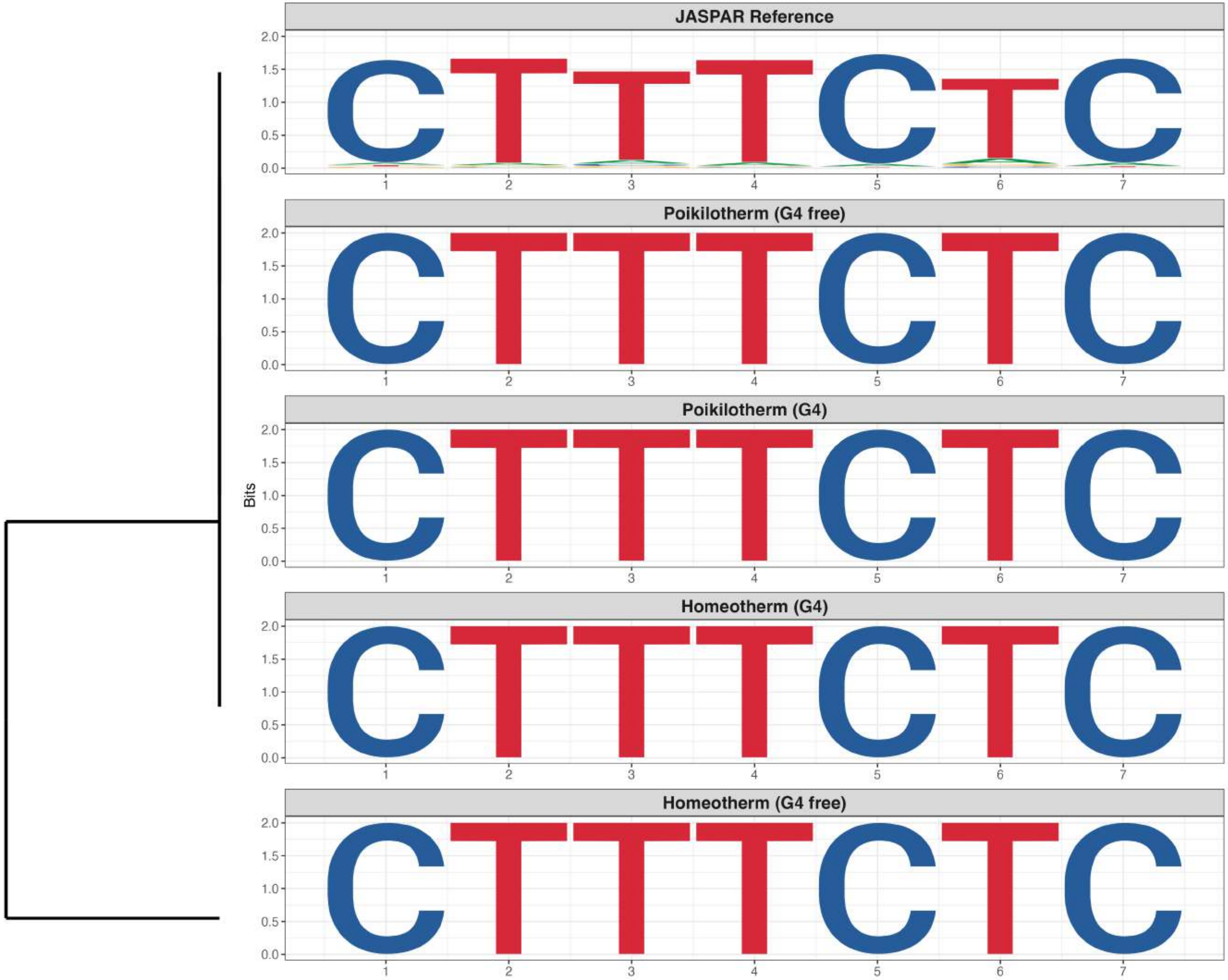

### Promoter : Highly Enriched in G4s

*MA0632.3 - TCFL5*

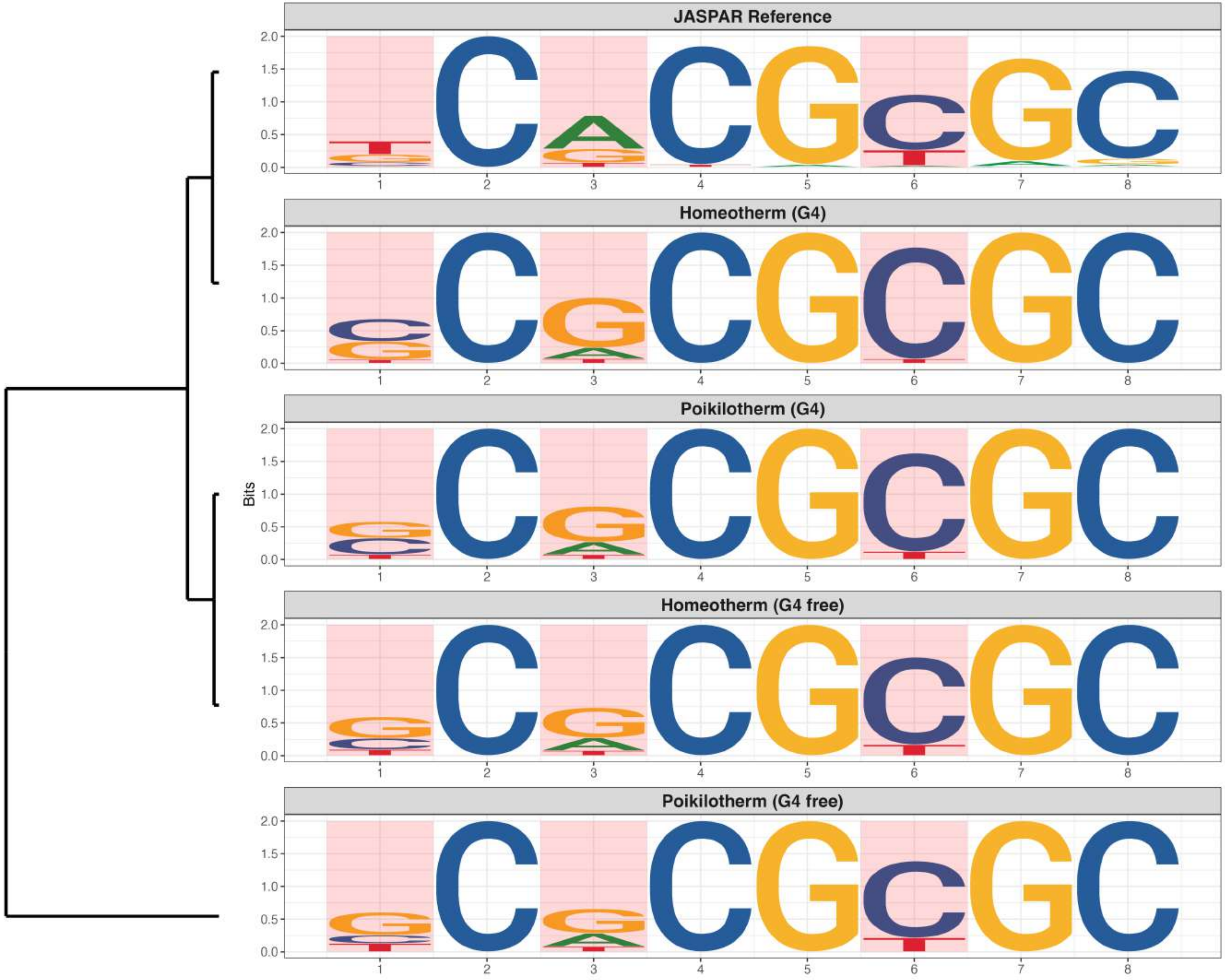

**Promoter : Highly Enriched in G4s**

*MA0646.2 - GCM1*

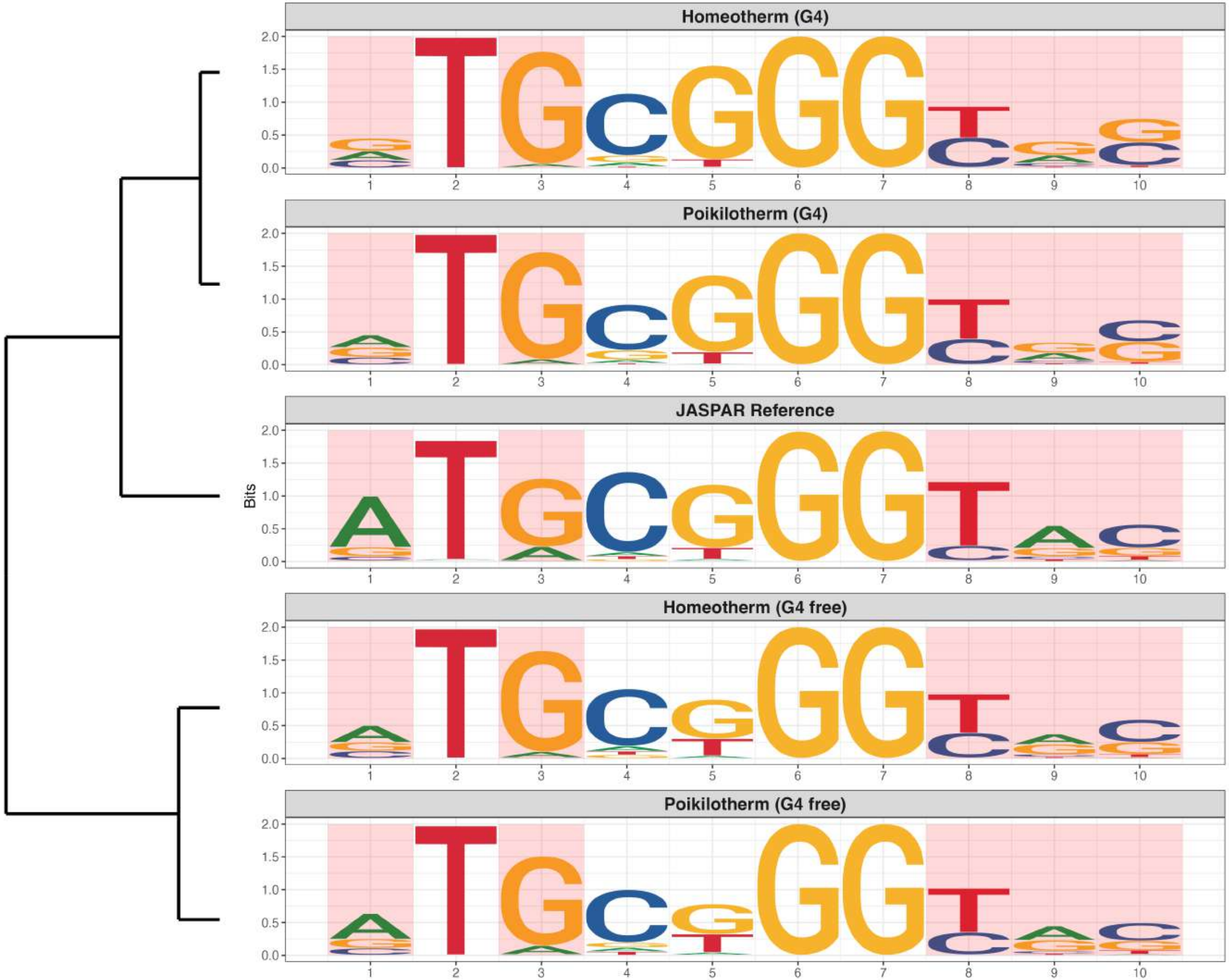

**Promoter : Highly Enriched in G4s**

*MA0712.3 - OTX2*

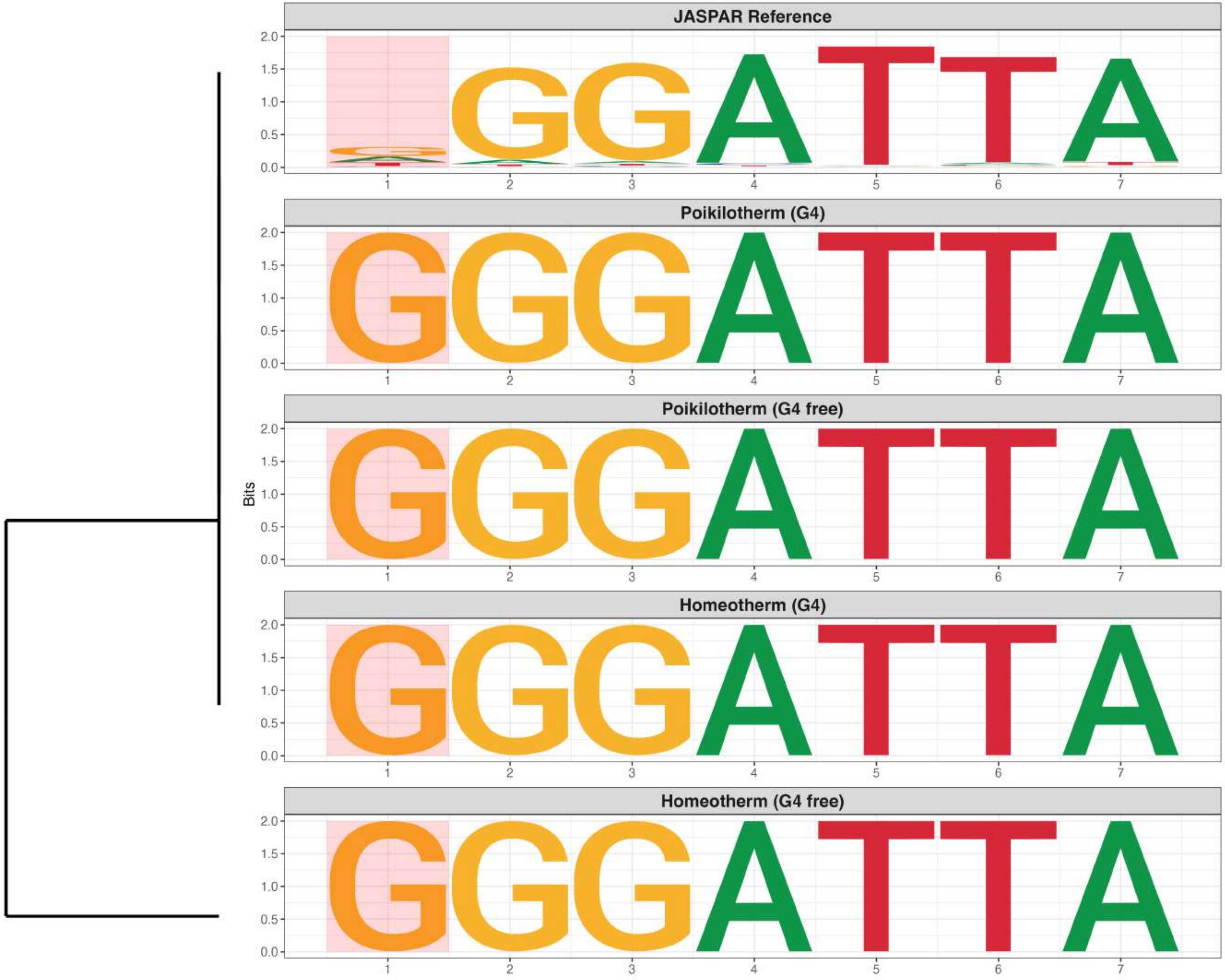

**Promoter : Highly Enriched in G4s**

*MA0728.1 - Nr2F6*

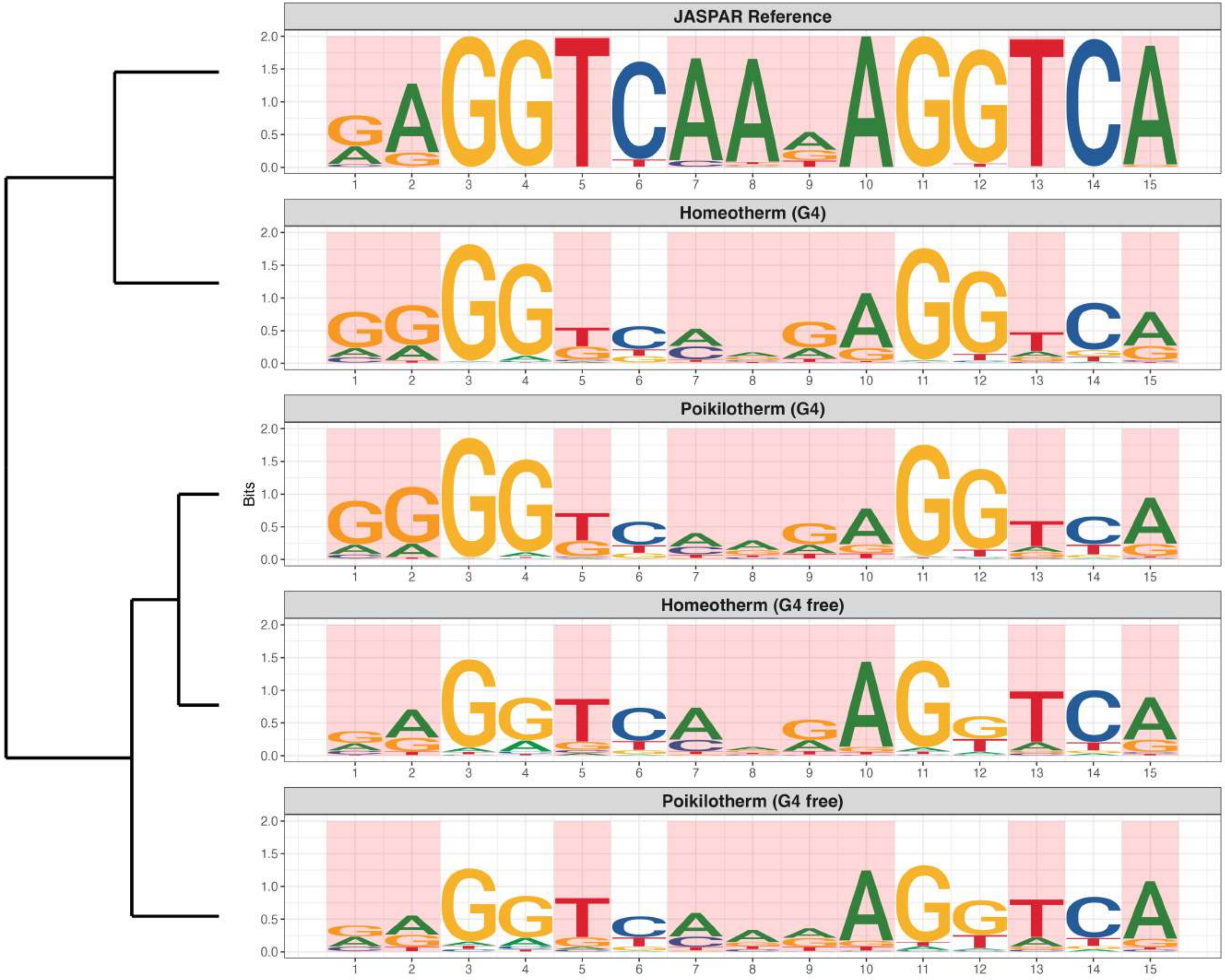

**Promoter : Highly Enriched in G4s**

*MA0729.1 - RARA*

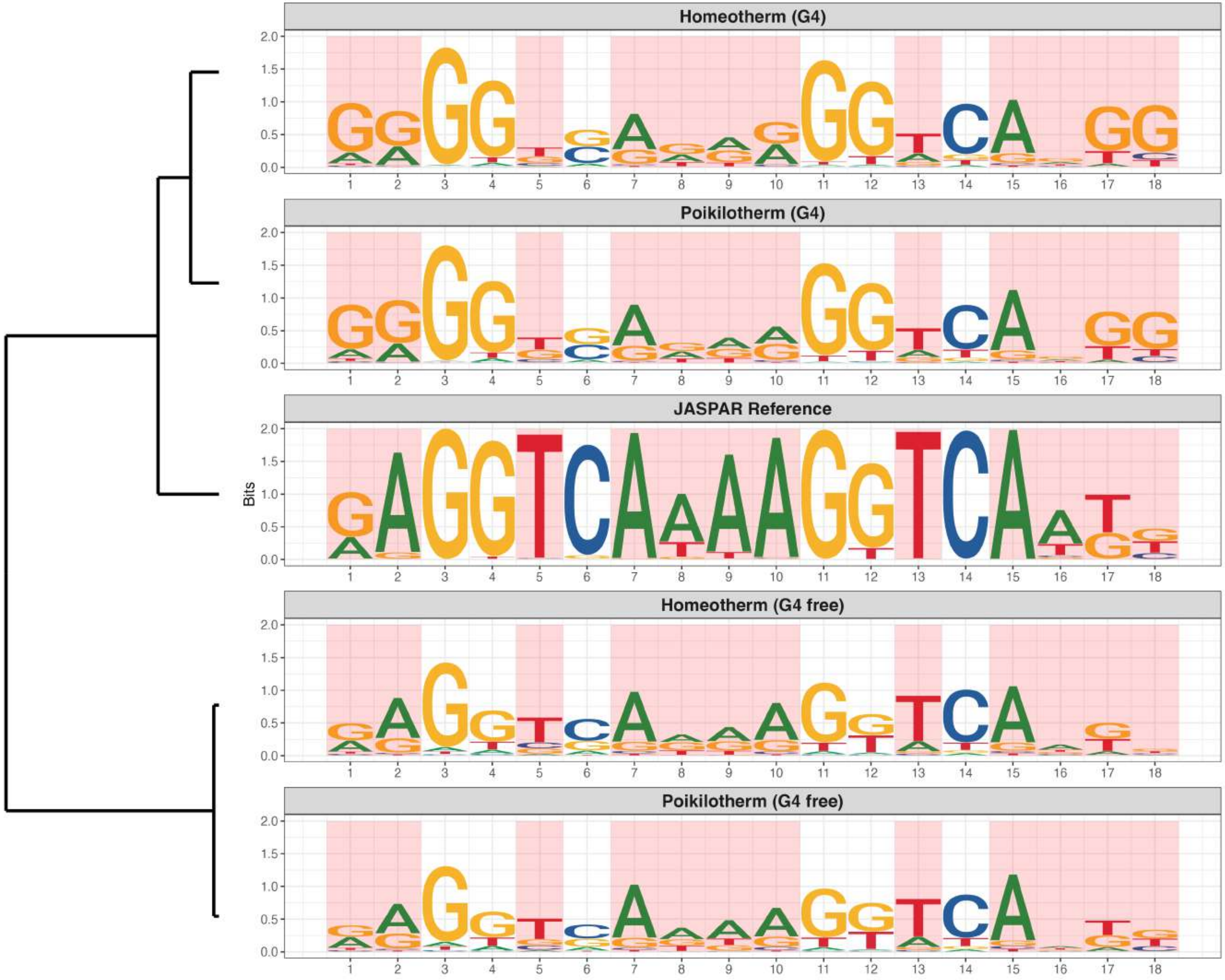

**Promoter : Highly Enriched in G4s**

*MA0745.3 - SNAI2*

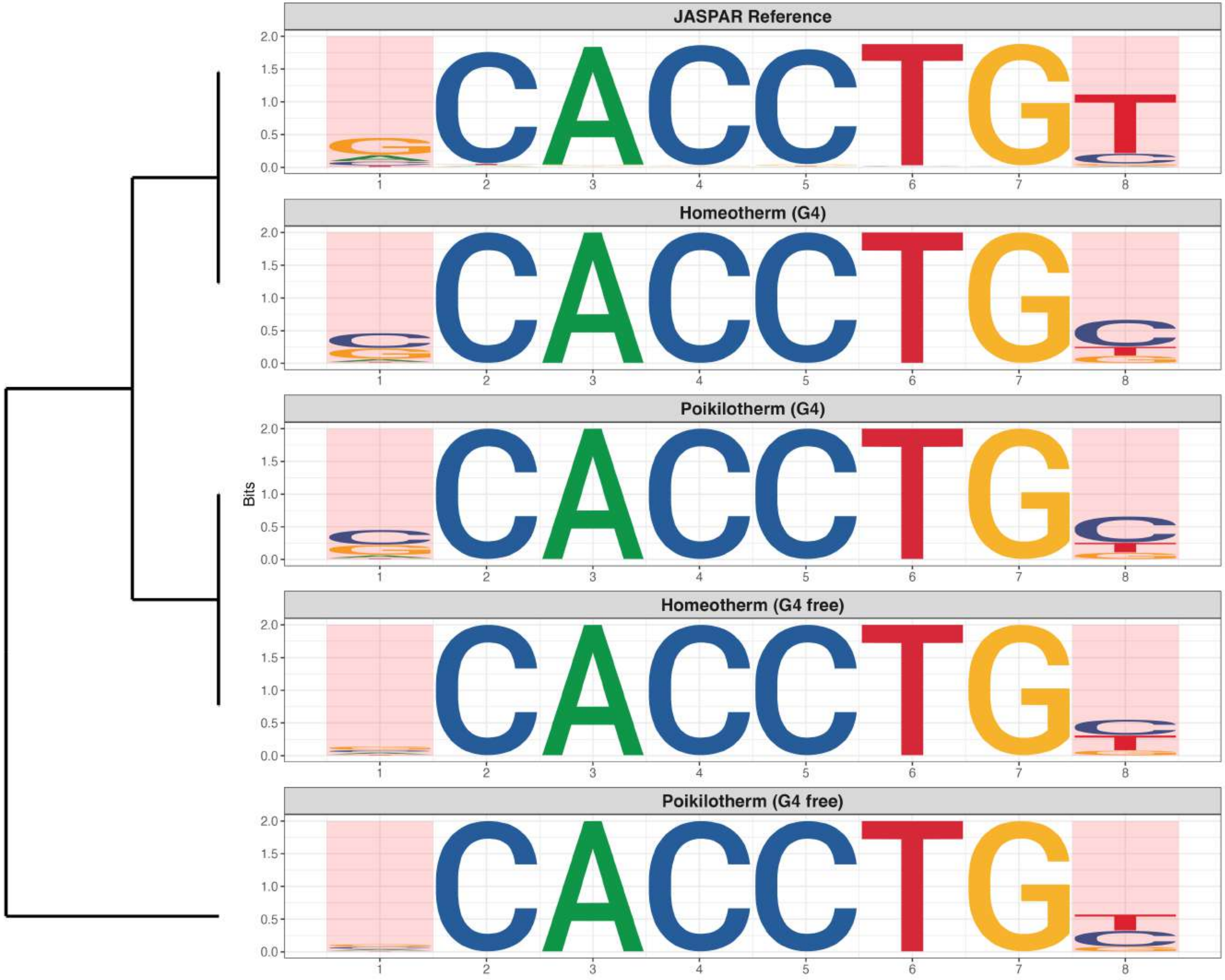

**Promoter : Highly Enriched in G4s**

*MA1474.2 - CREB3L4*

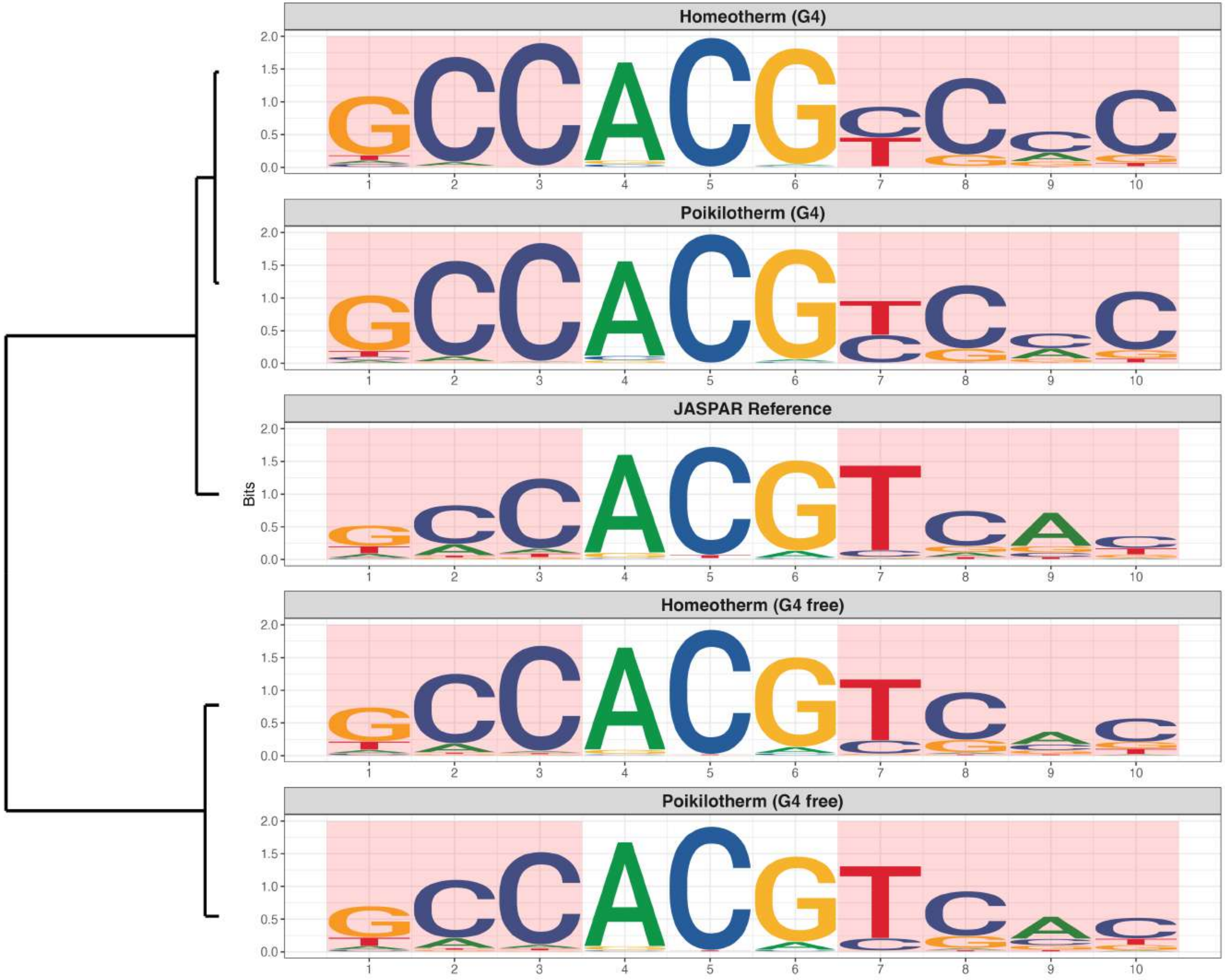

**Promoter : Highly Enriched in G4s**

*MA1523.2 - MSANTD3*

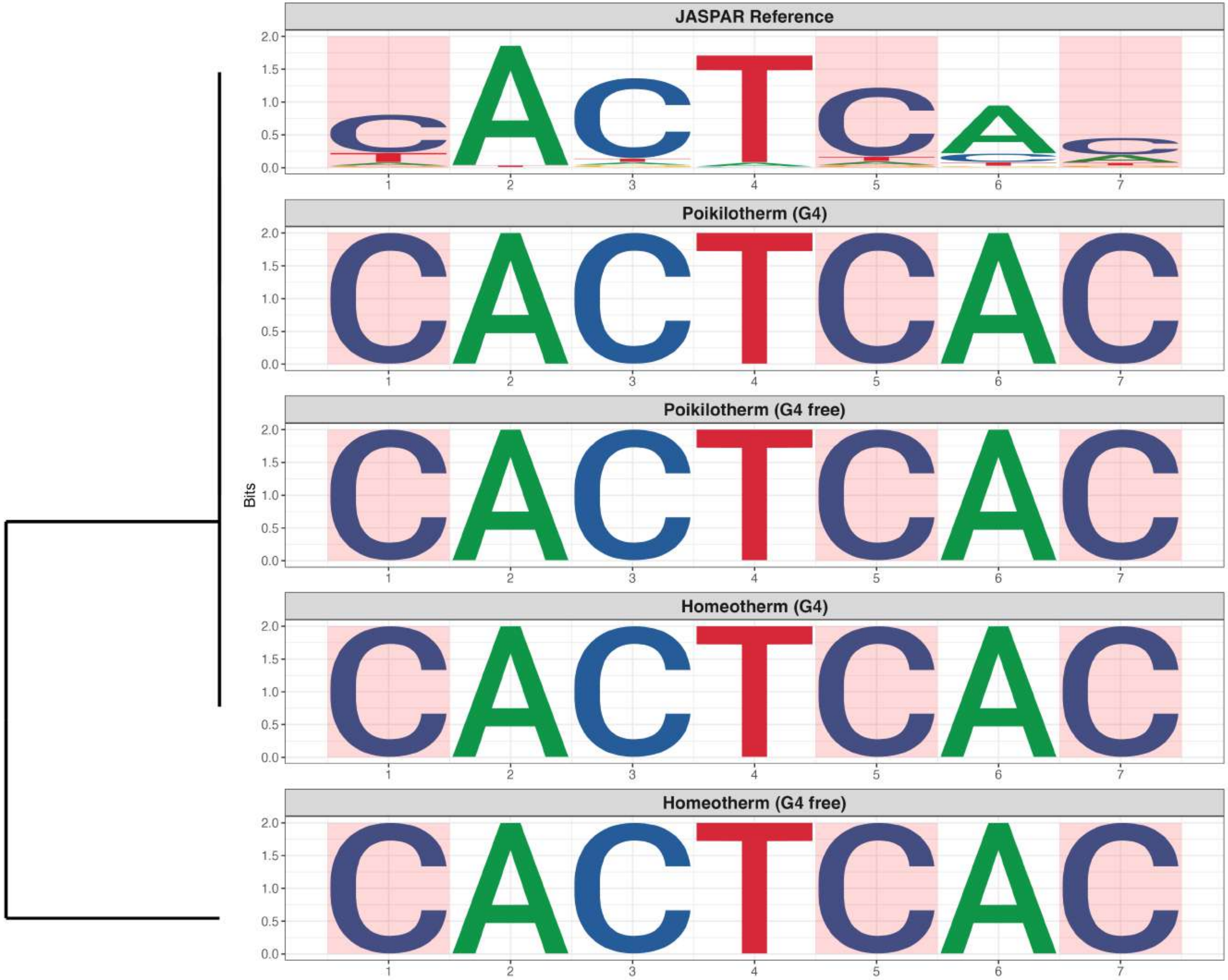

**Promoter : Highly Enriched in G4s**

*MA1528.2 - NFIX*

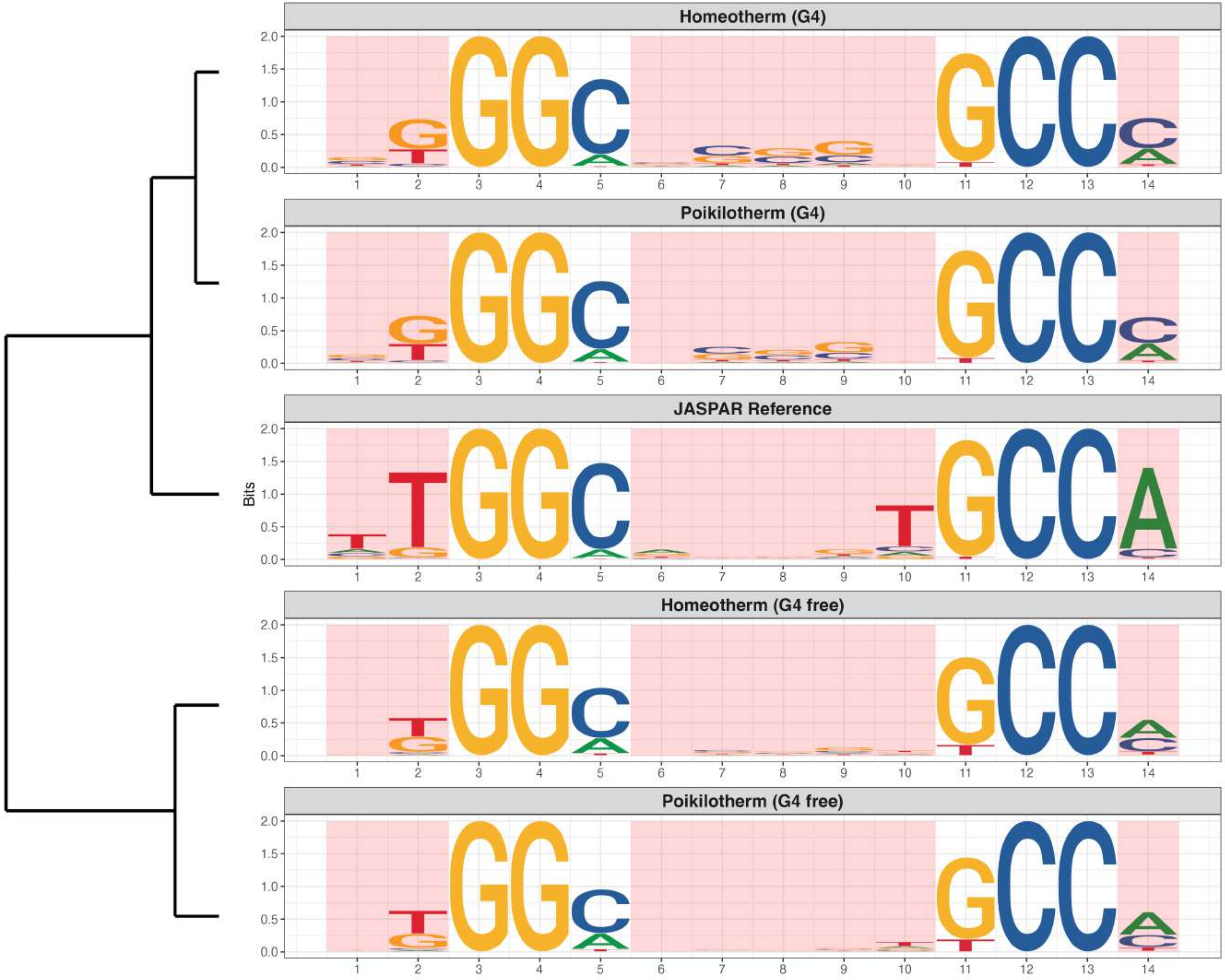

### Promoter : Highly Enriched in G4s

*MA1533.2 - NR1I2*

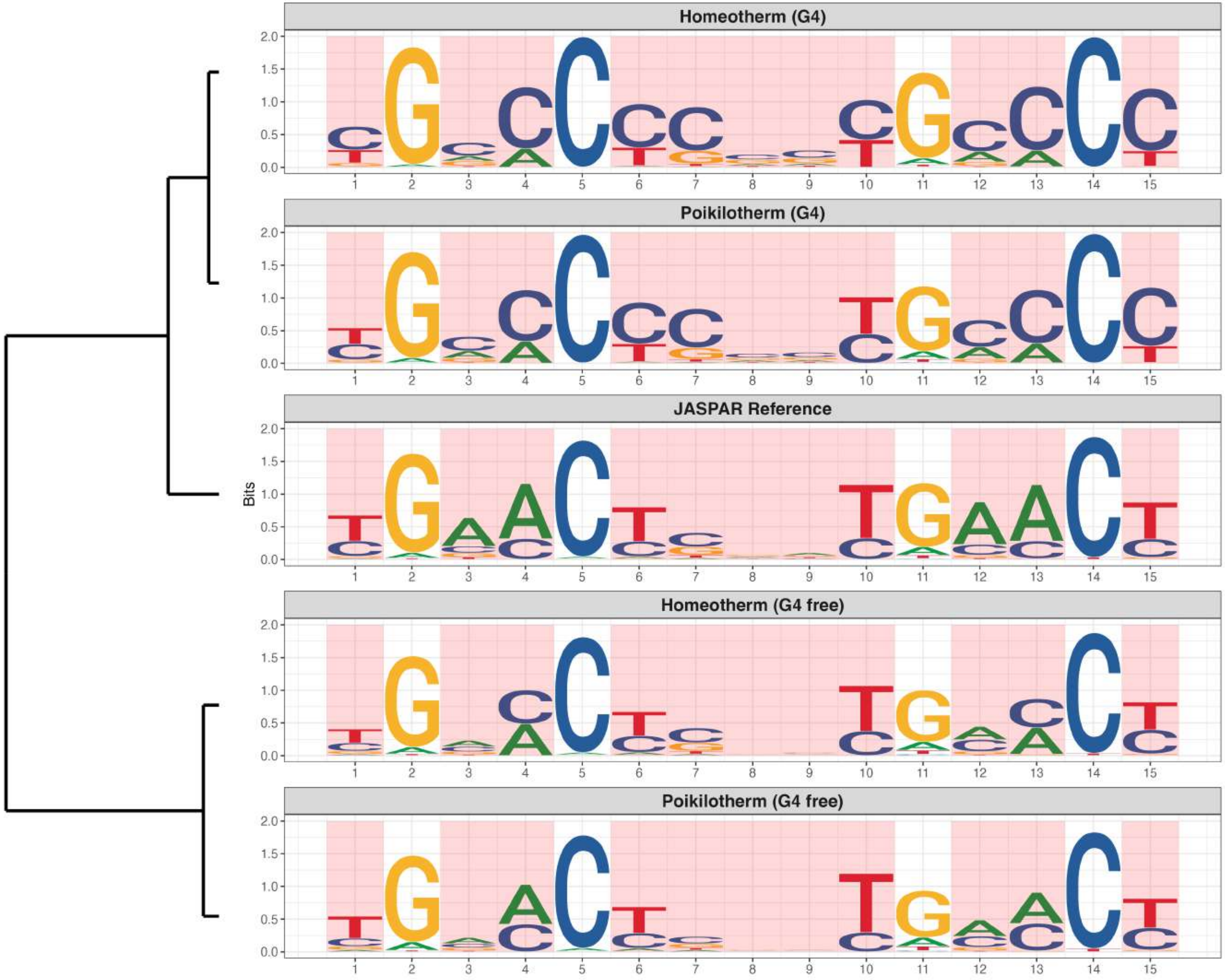

**Promoter : Highly Enriched in G4s**

*MA1558.2 - SNAI1*

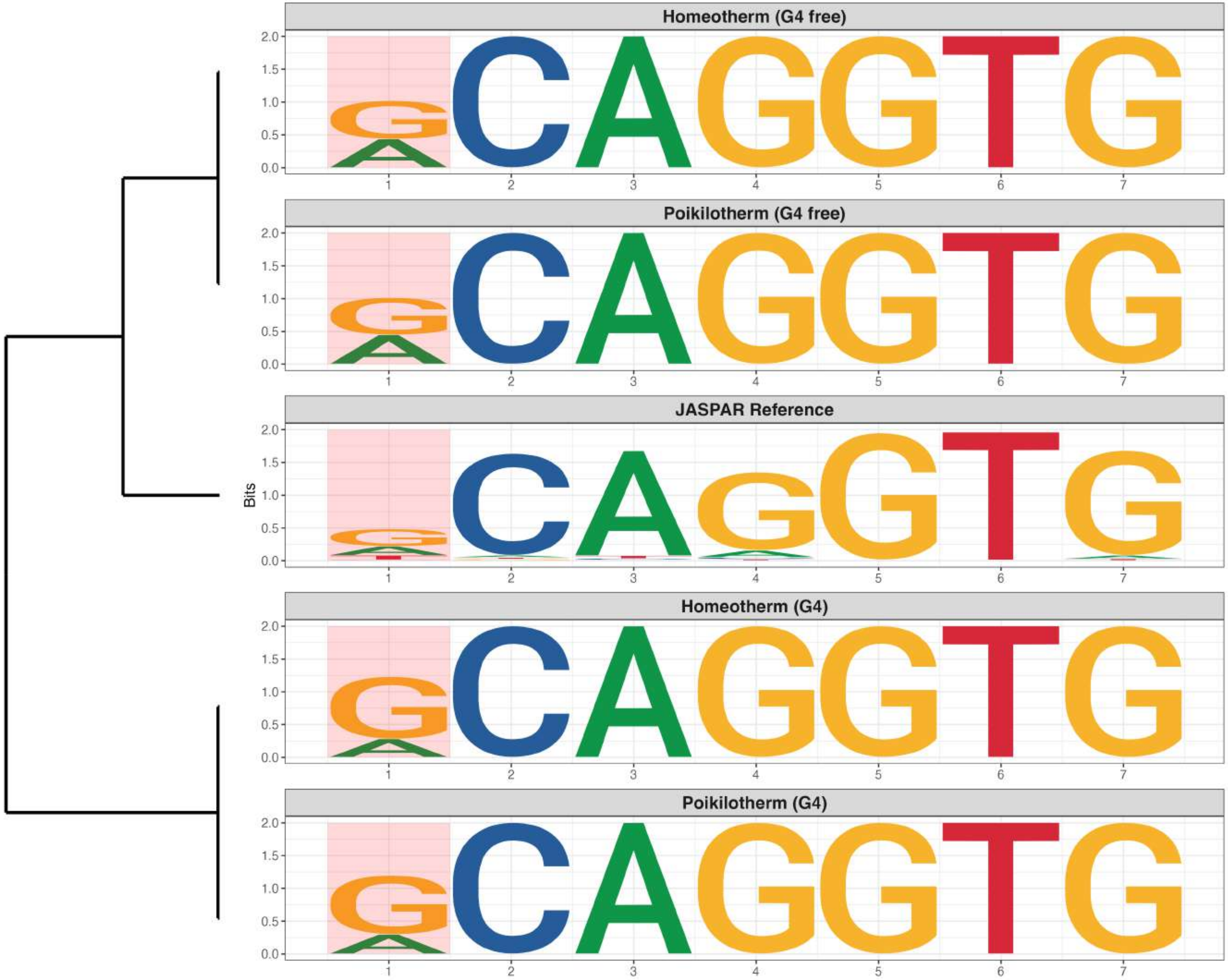

**Promoter : Highly Enriched in G4s**

*MA1645.2 - NKX2-2*

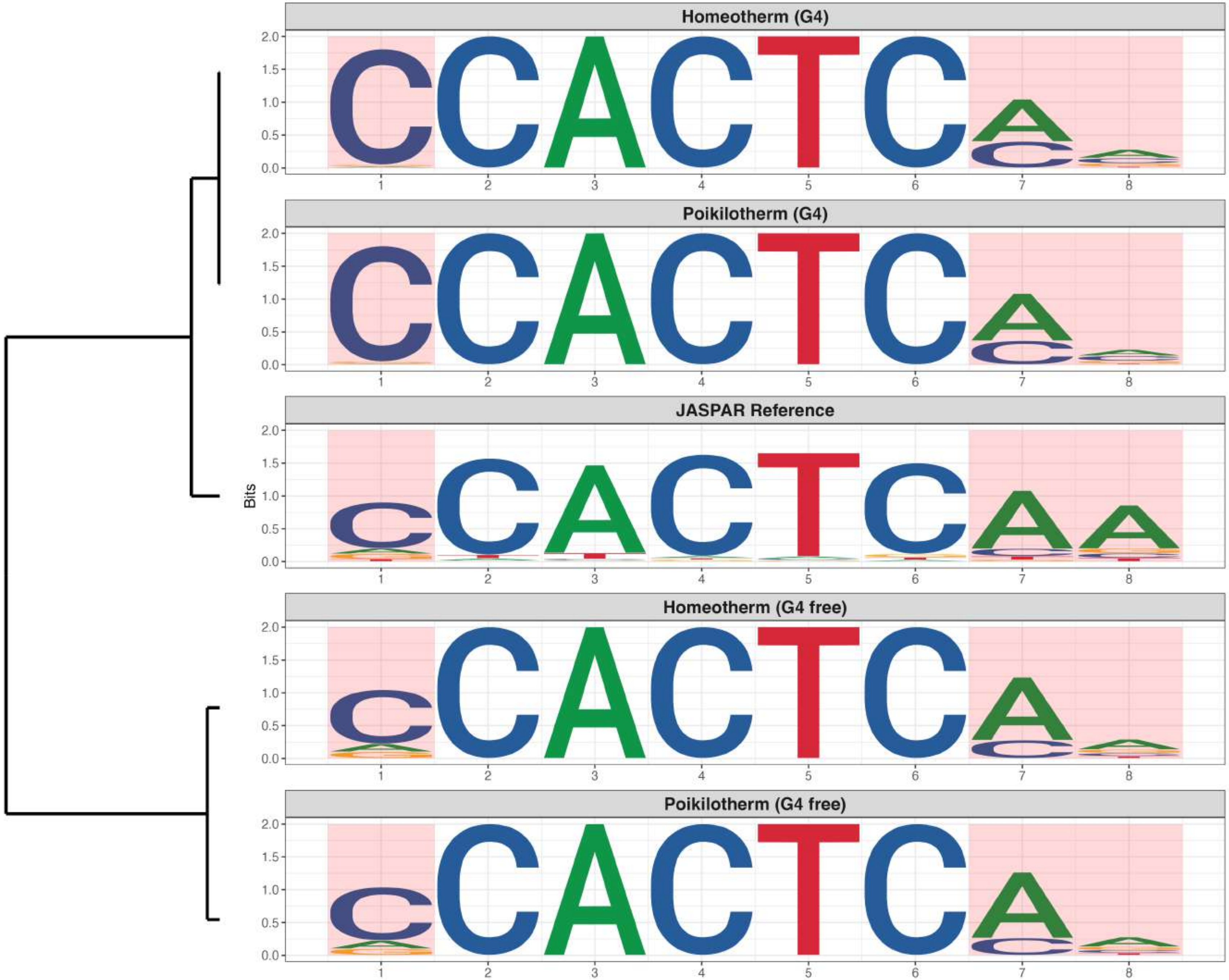

**Promoter : Highly Enriched in G4s**

*MA1712.2 - ZNF454*

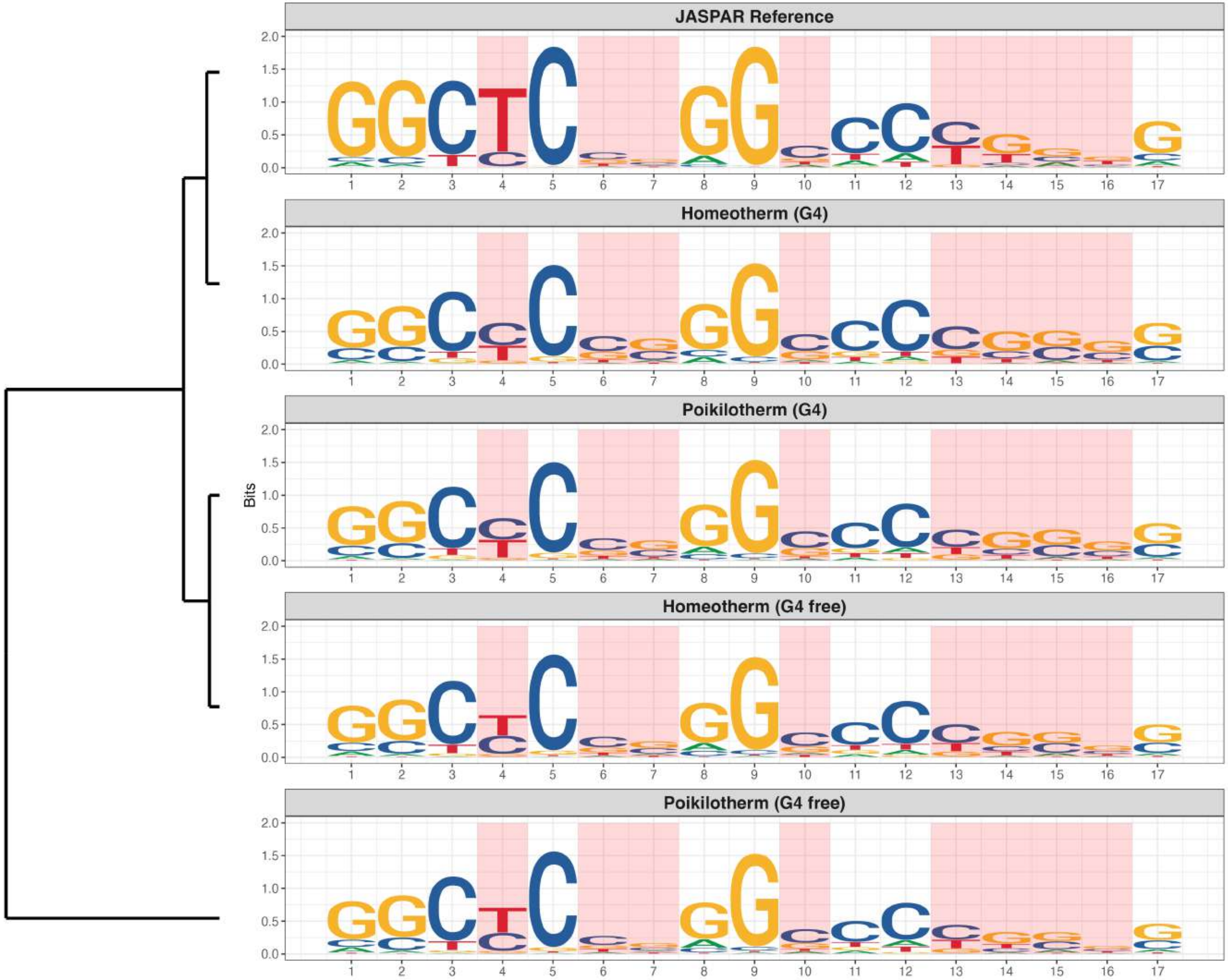

### Promoter : Highly Enriched in G4s

*MA1979.2 - ZNF416*

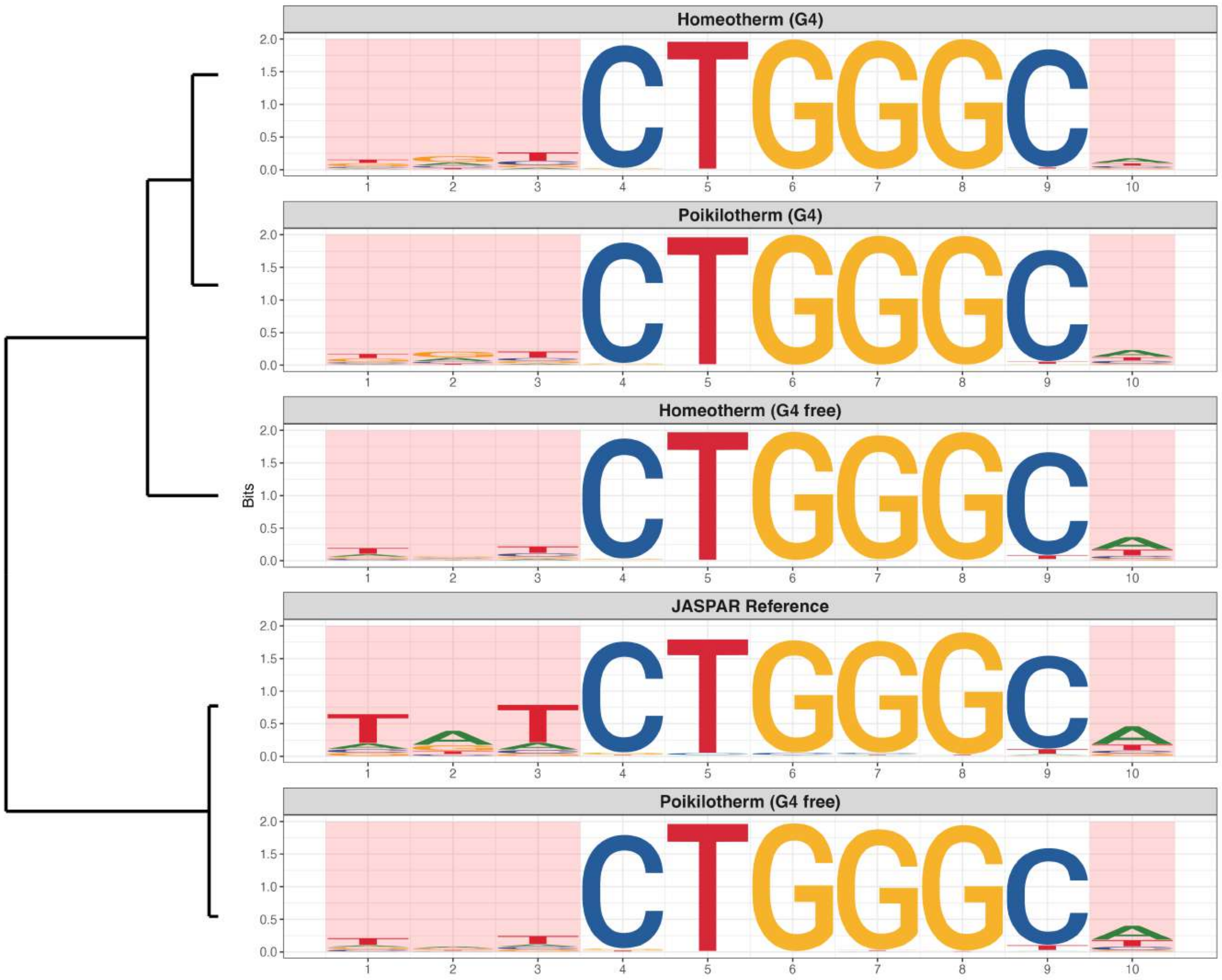

**Promoter : Highly Enriched in G4s**

*MA2528.1 - ZNF732*

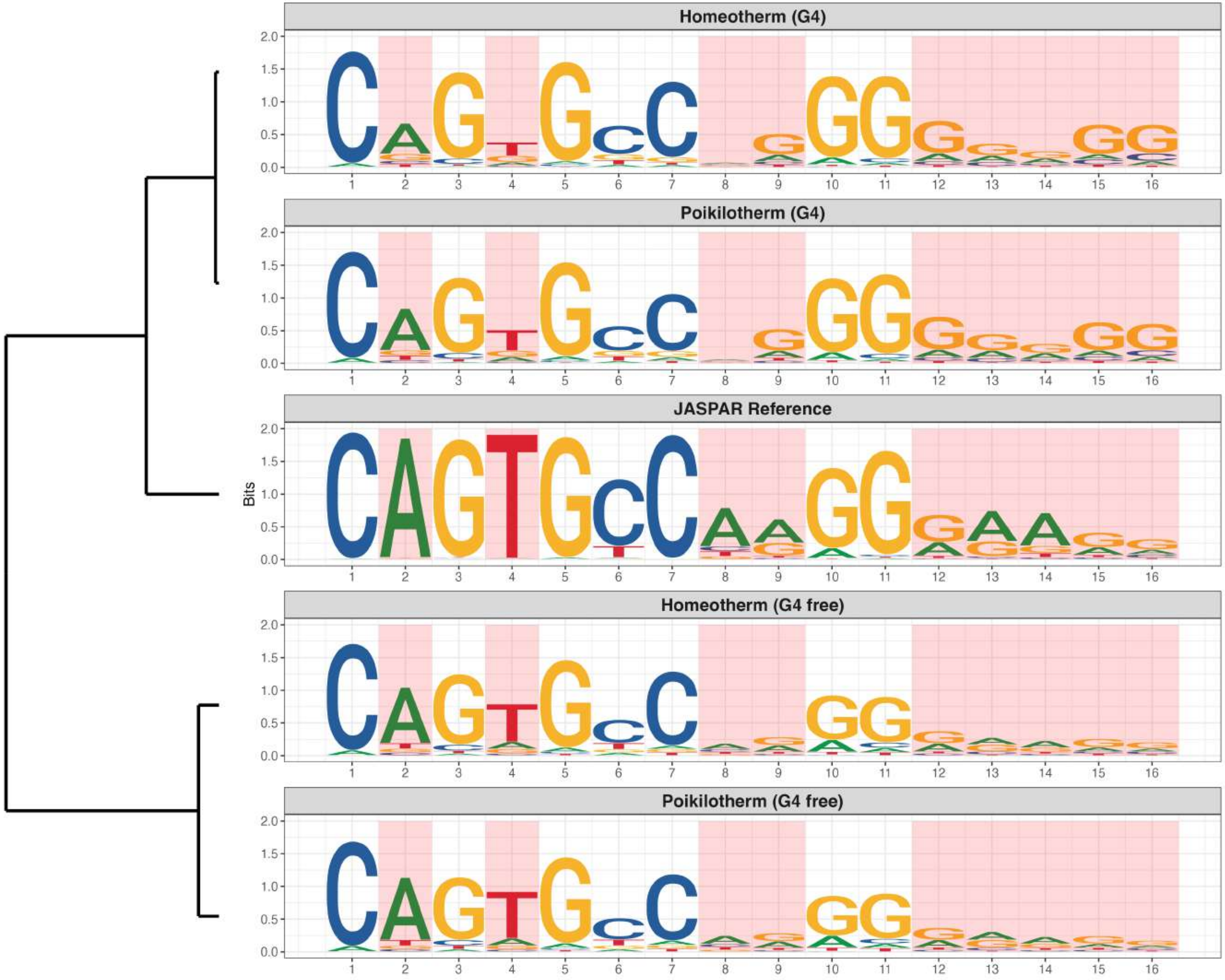

**Promoter : Highly Enriched in G4s**

*MA2545.1 - ZNF57*

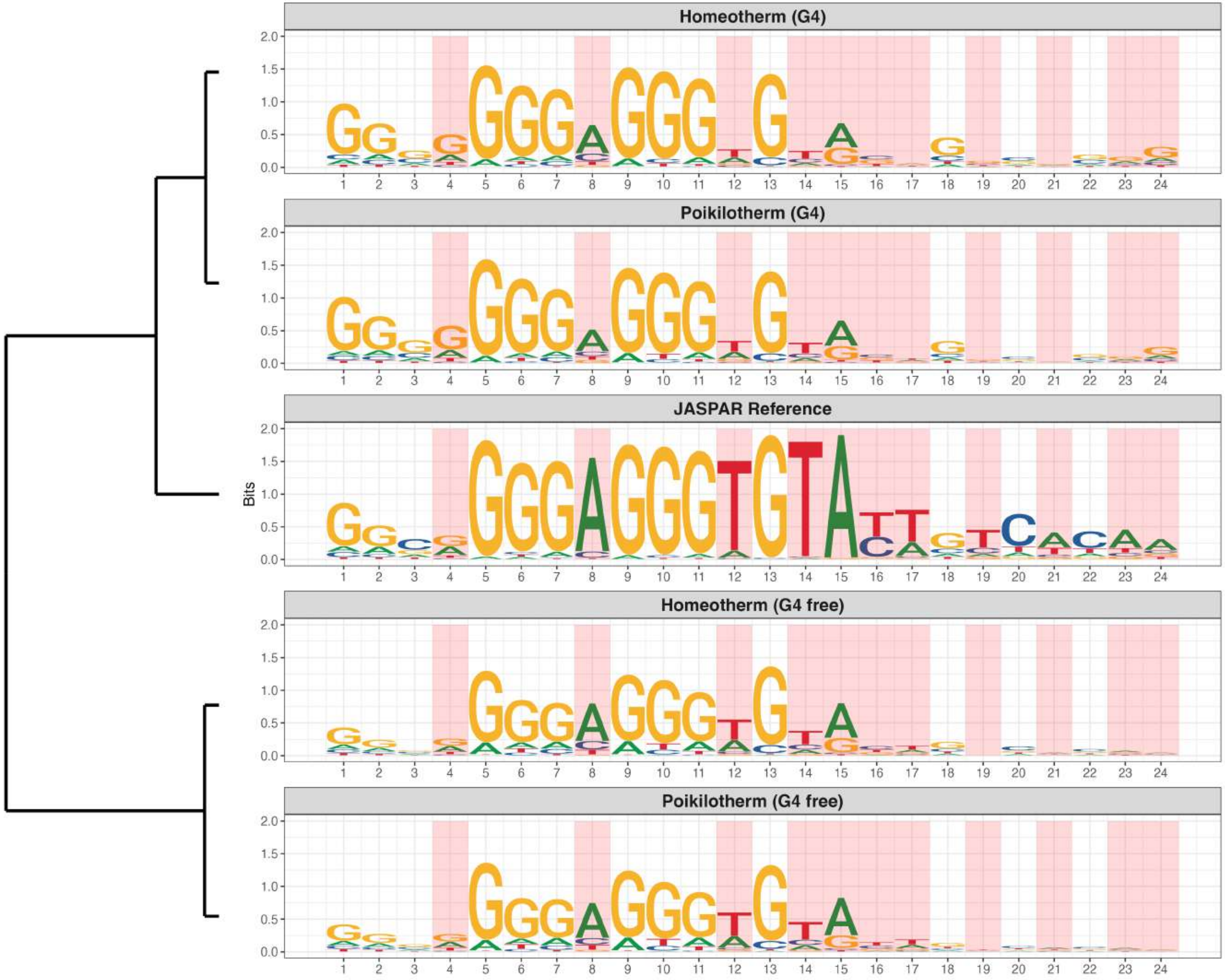

**Promoter : Highly Enriched in G4s**

*MA2573.1 - ZNF836*

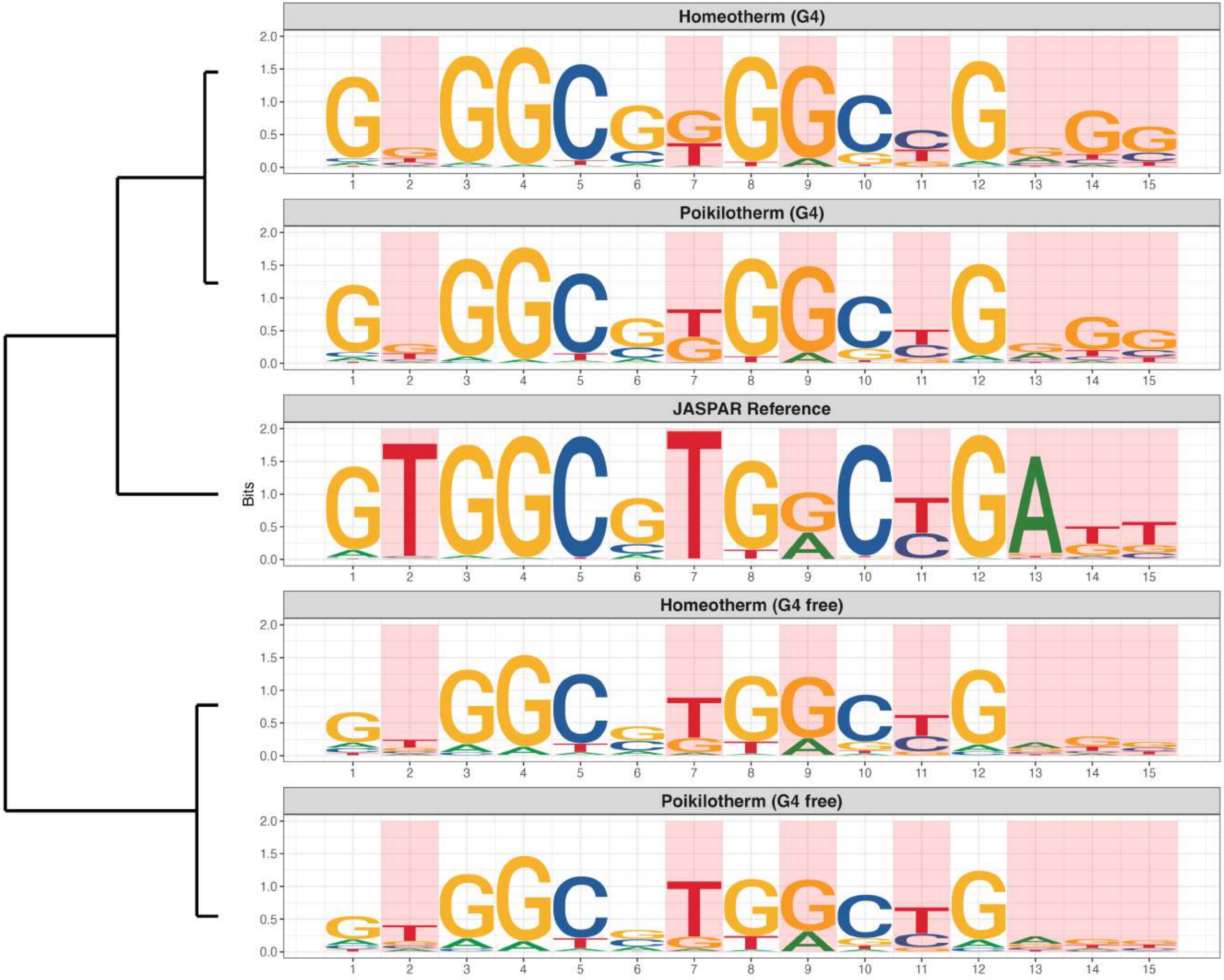

**Promoter : Highly Enriched in G4s**

*MA2582.1 - ZNF121*

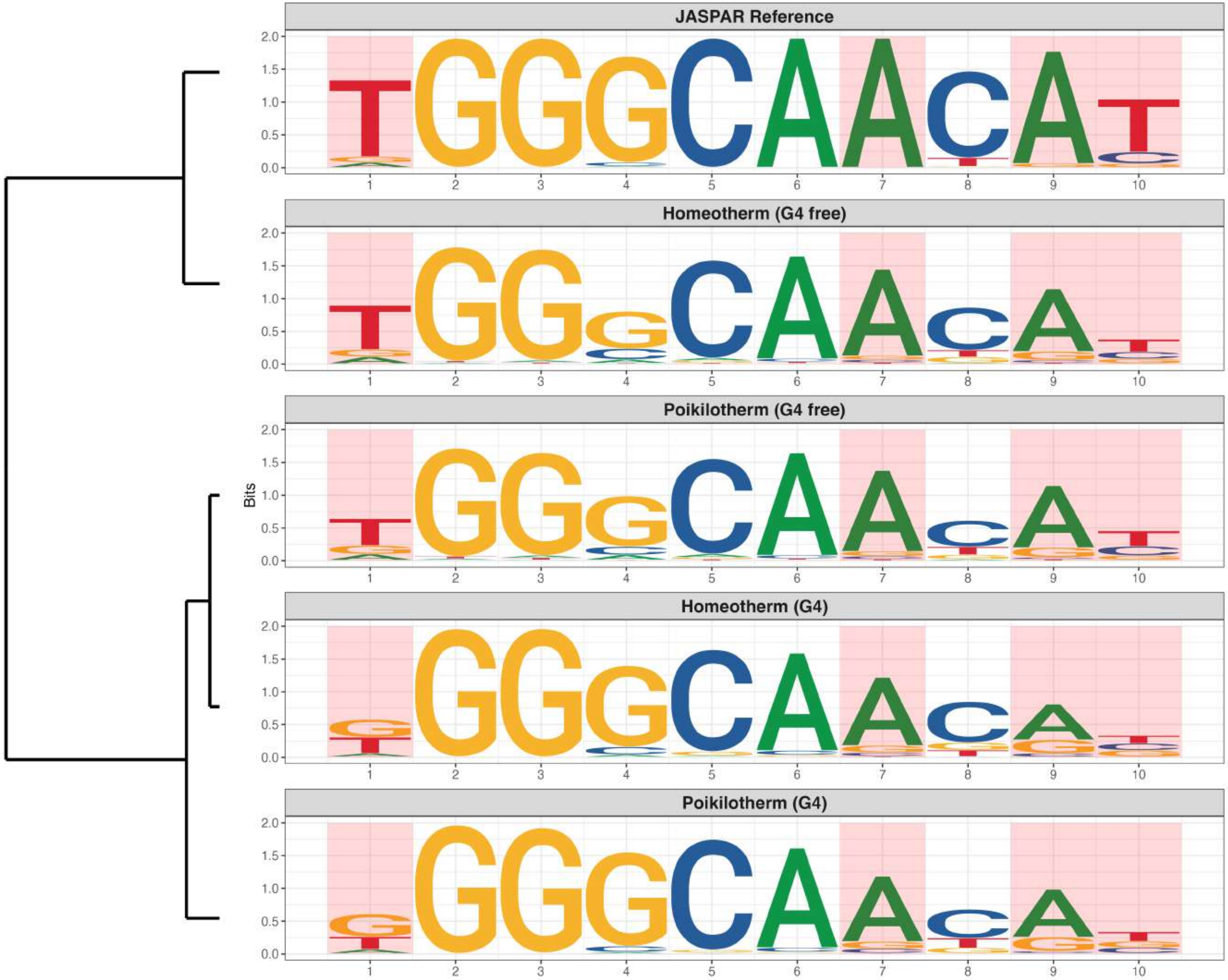

### Promoter : Highly Enriched in G4s

*MA2588.1 - FAM200B*

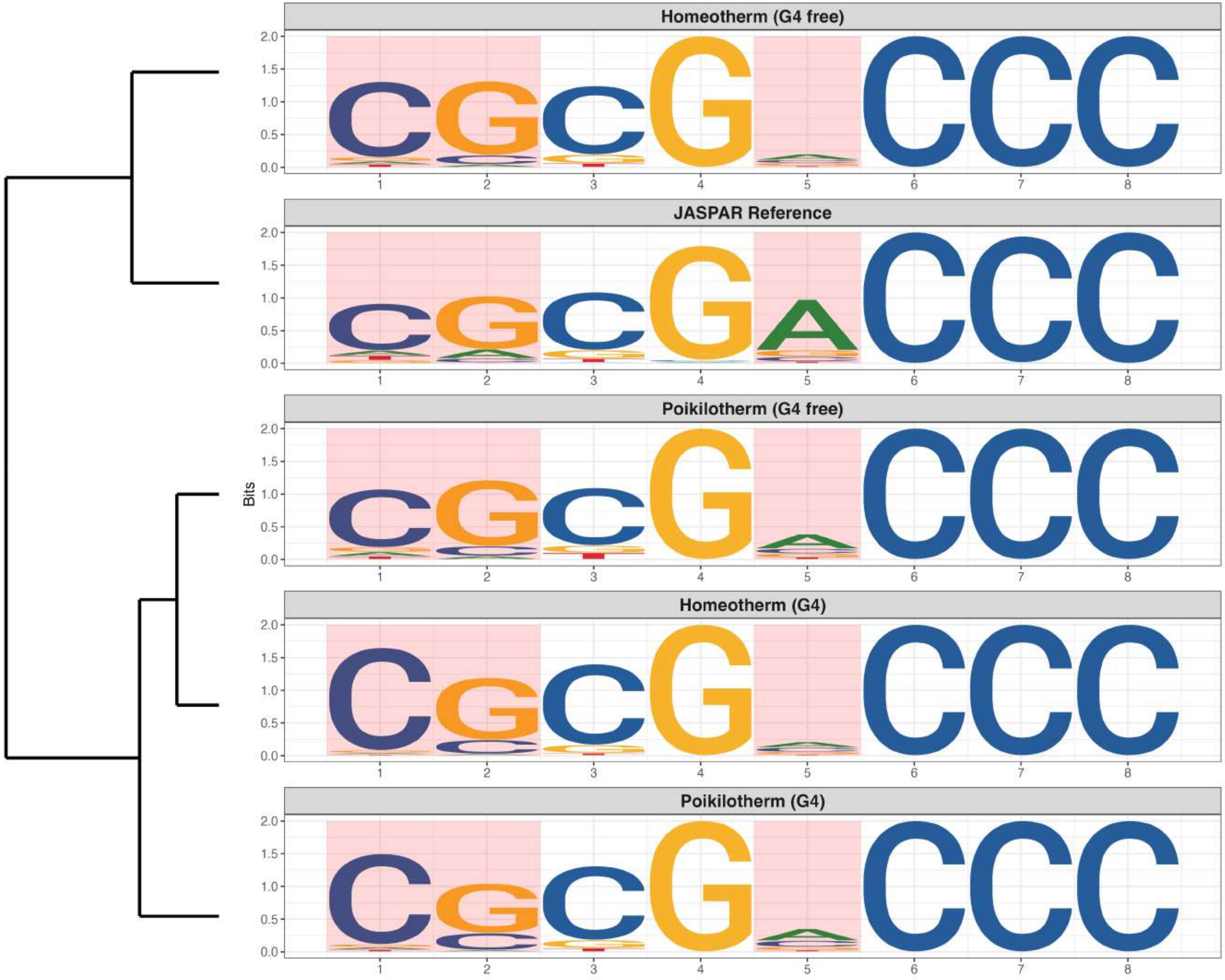

**Promoter : Highly Enriched in G4s**

*MA2597.1 - LEUTX*

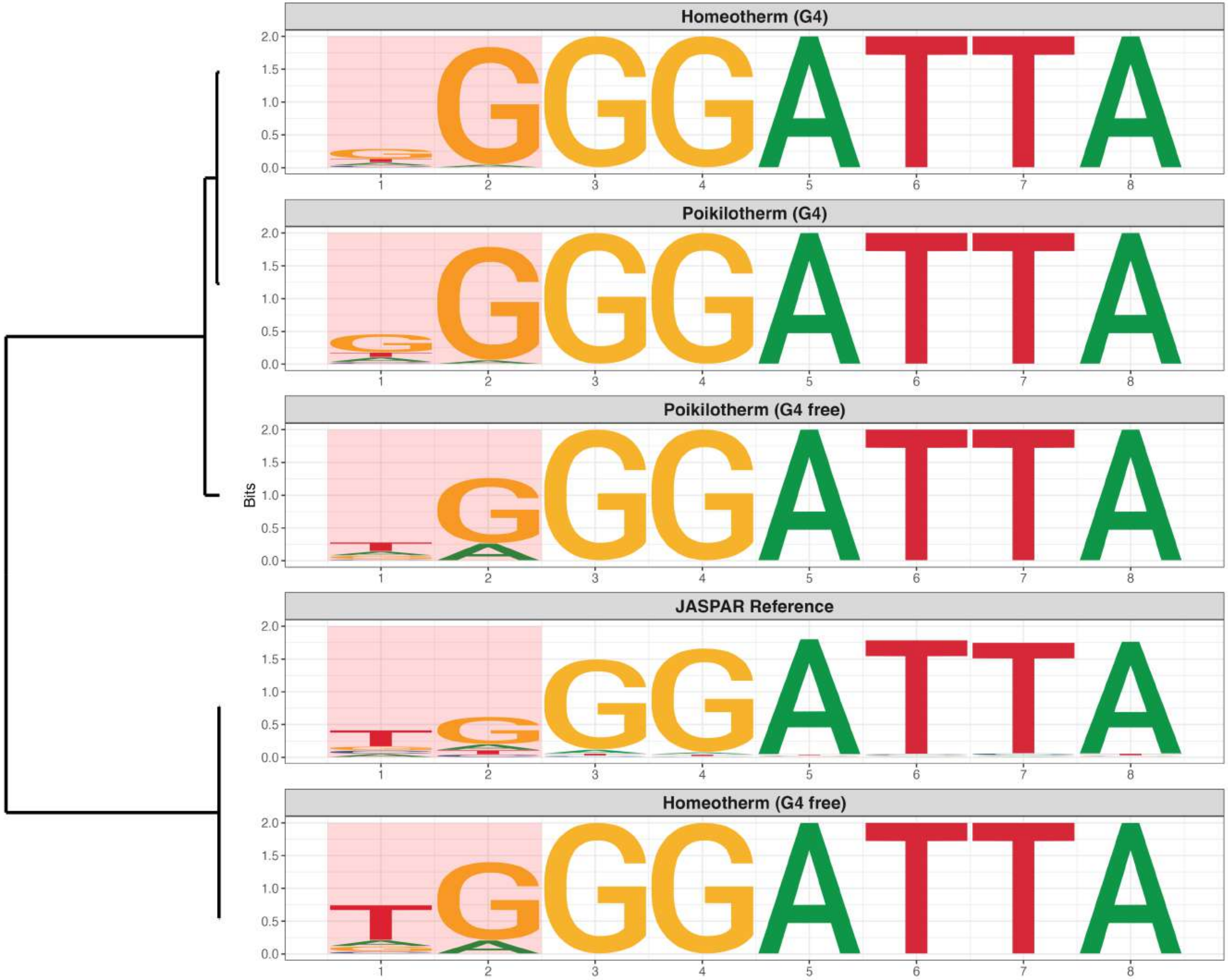

### Promoter : Moderately Enriched in G4s

*MA0131.3 - HINFP*

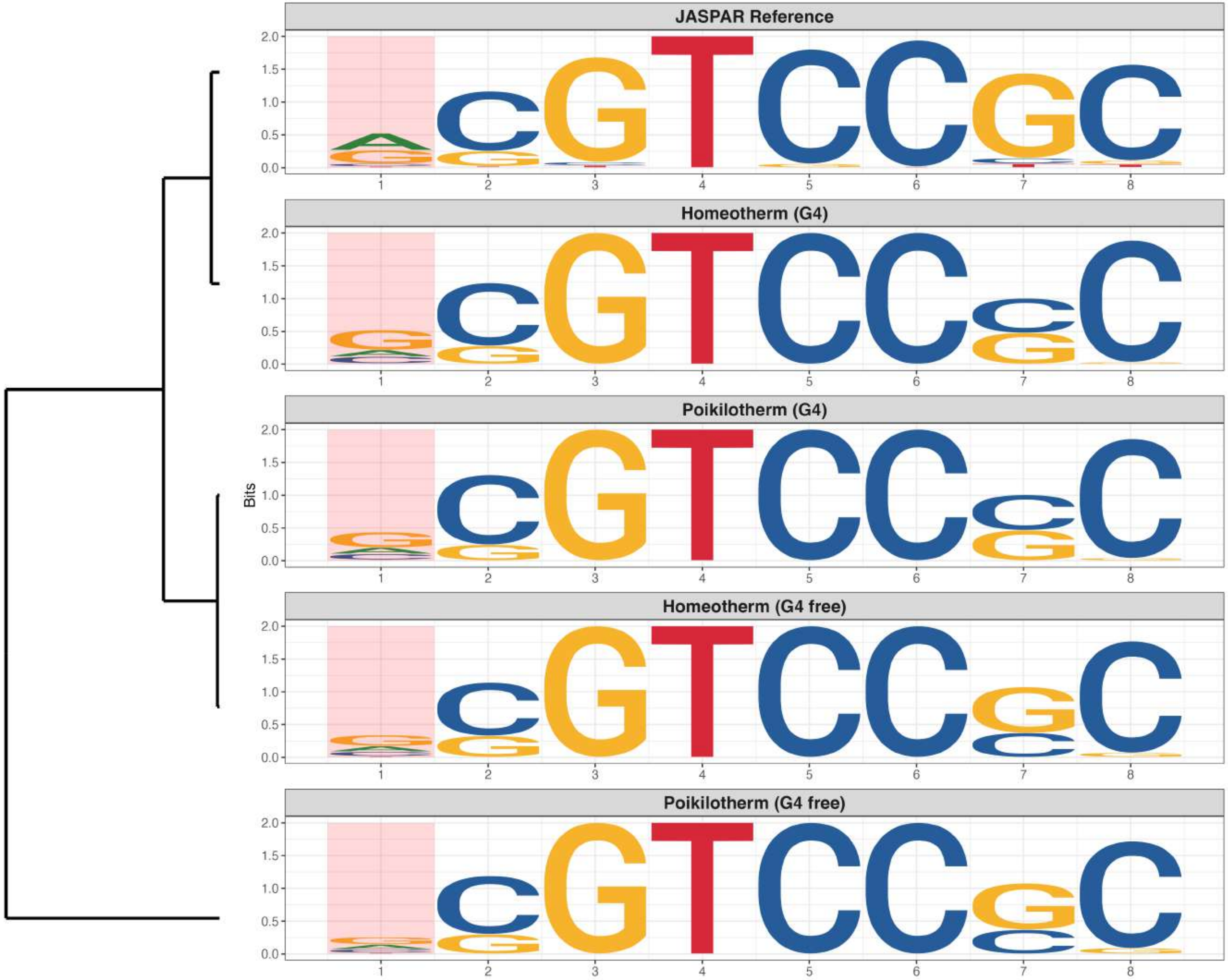

### Promoter : Moderately Enriched in G4s

*MA0136.4 - Elf5*

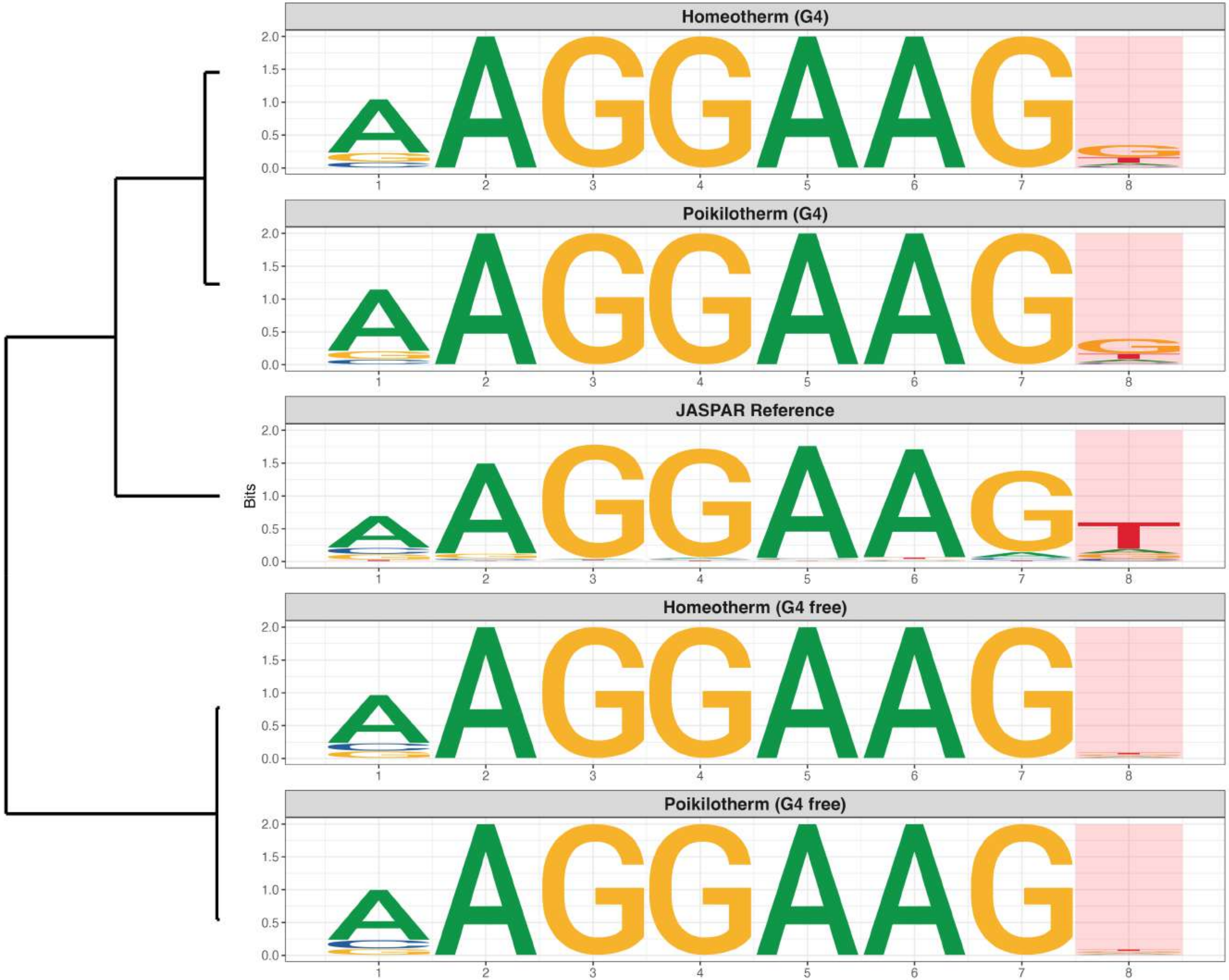

**Promoter : Moderately Enriched in G4s**

*MA0146.3 - Zfx*

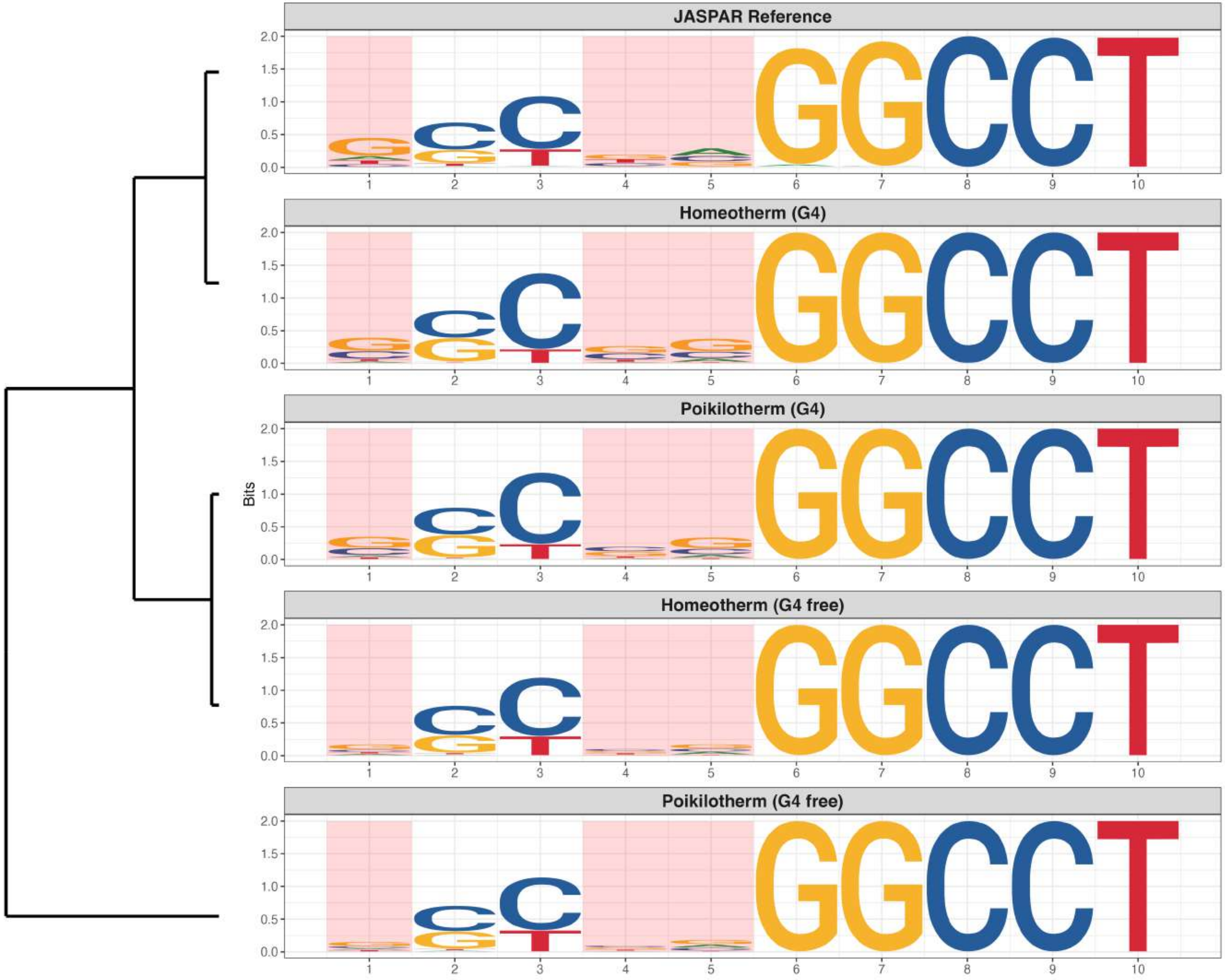

**Promoter : Moderately Enriched in G4s**

*MA0472.2 - EGR2*

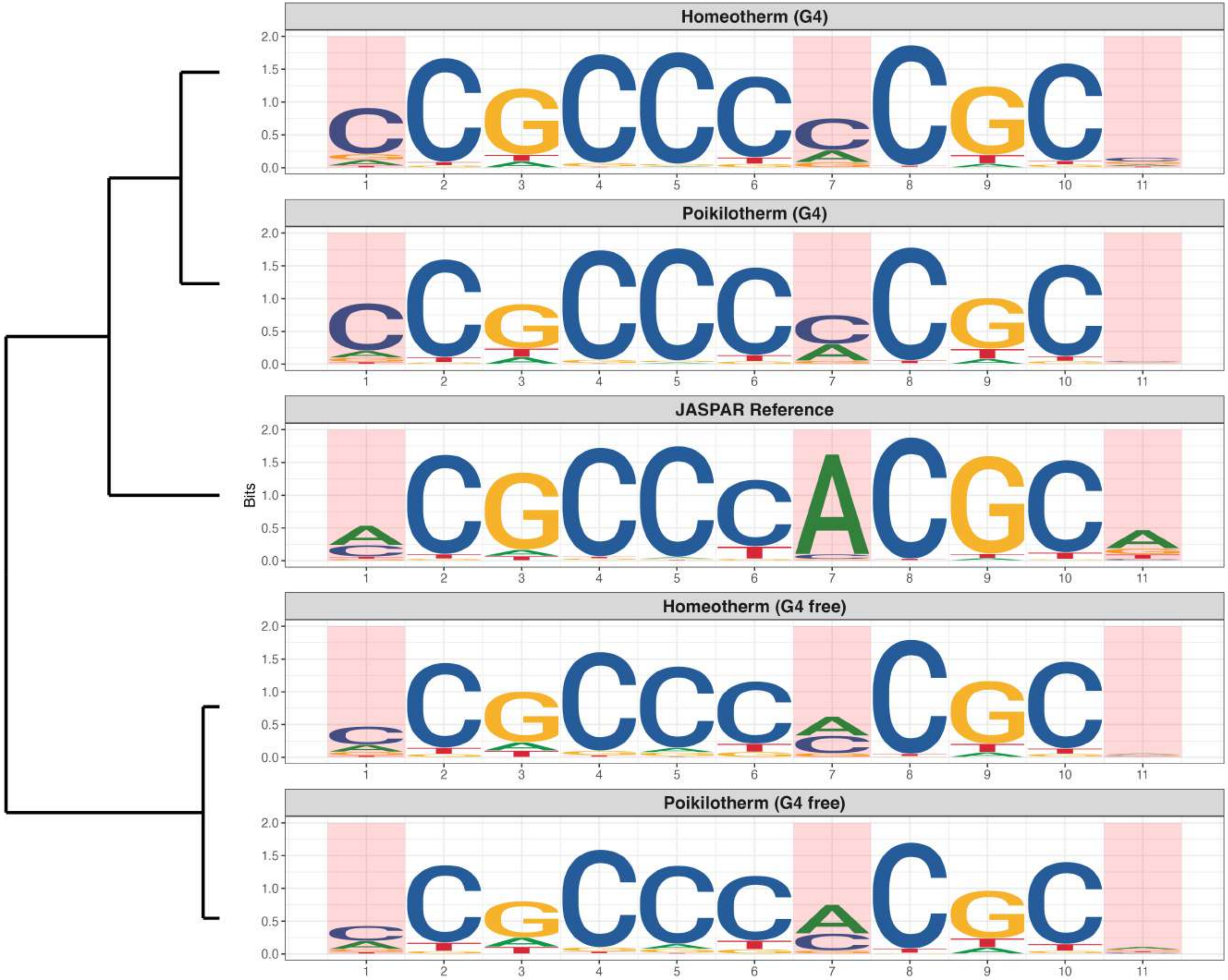

**Promoter : Moderately Enriched in G4s**

*MA0478.2 - FOSL2*

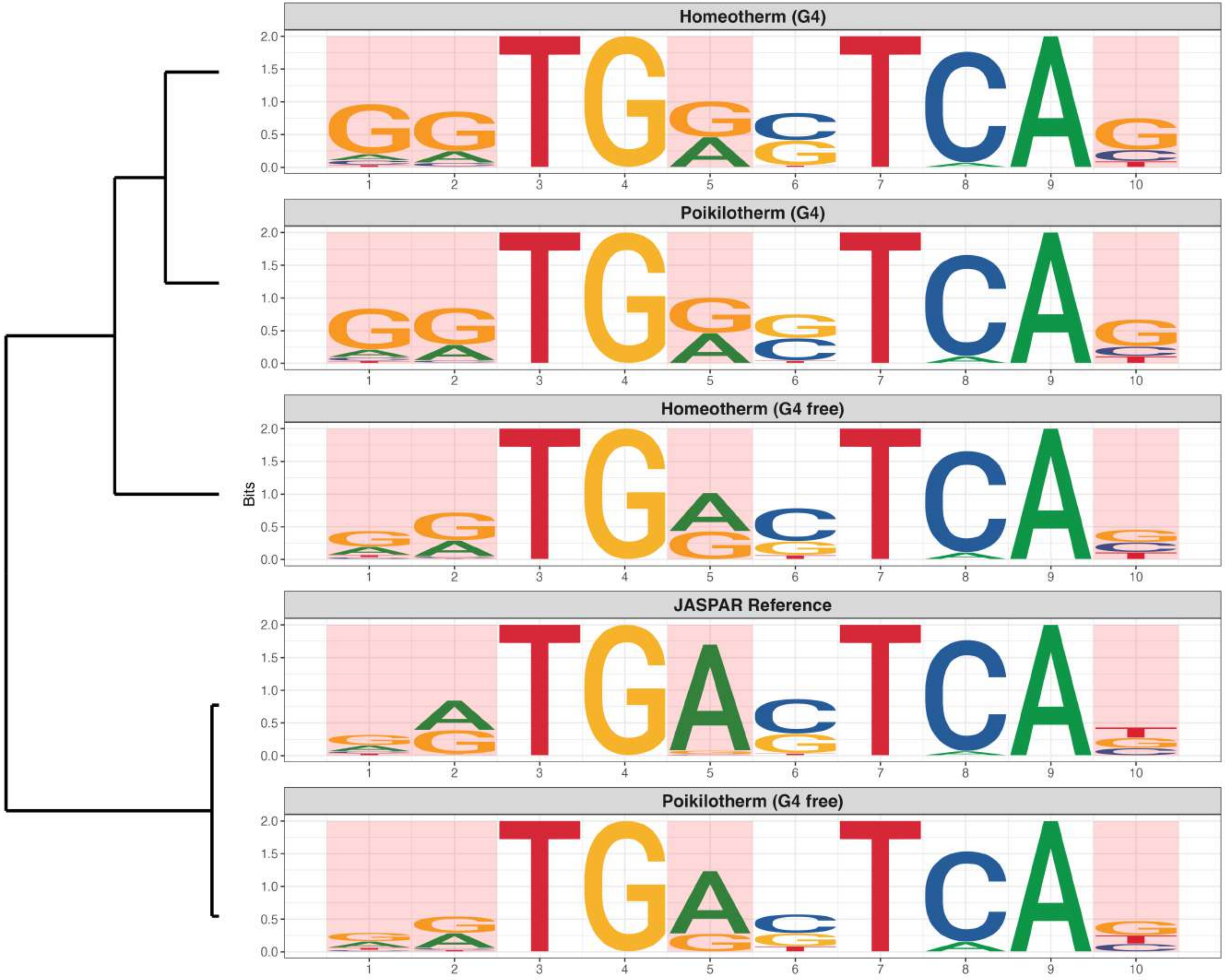

**Promoter : Moderately Enriched in G4s**

*MA0519.2 - Stat5a::Stat5b*

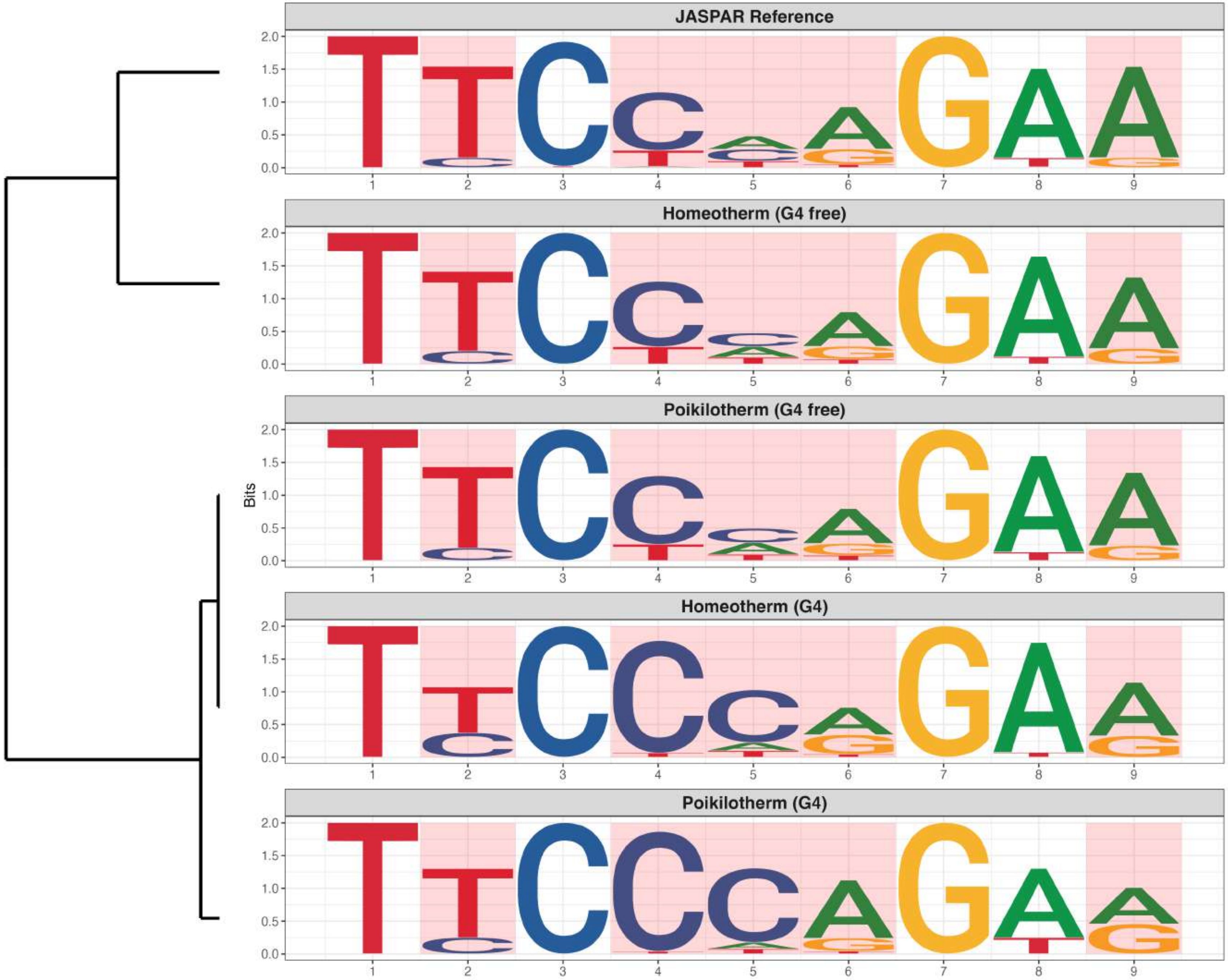

**Promoter : Moderately Enriched in G4s**

*MA0640.3 - ELF3*

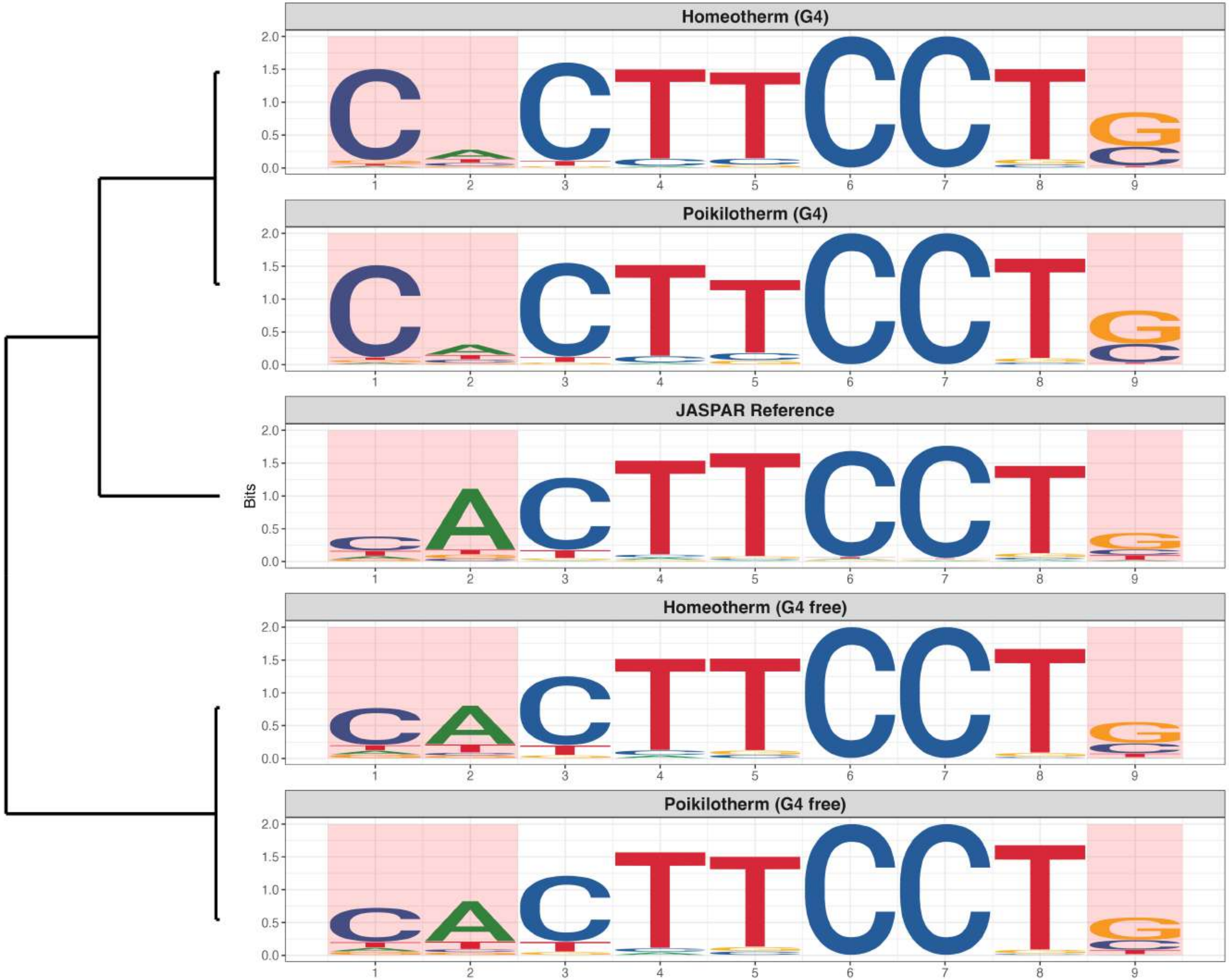

**Promoter : Moderately Enriched in G4s**

*MA0801.1 - MGA*

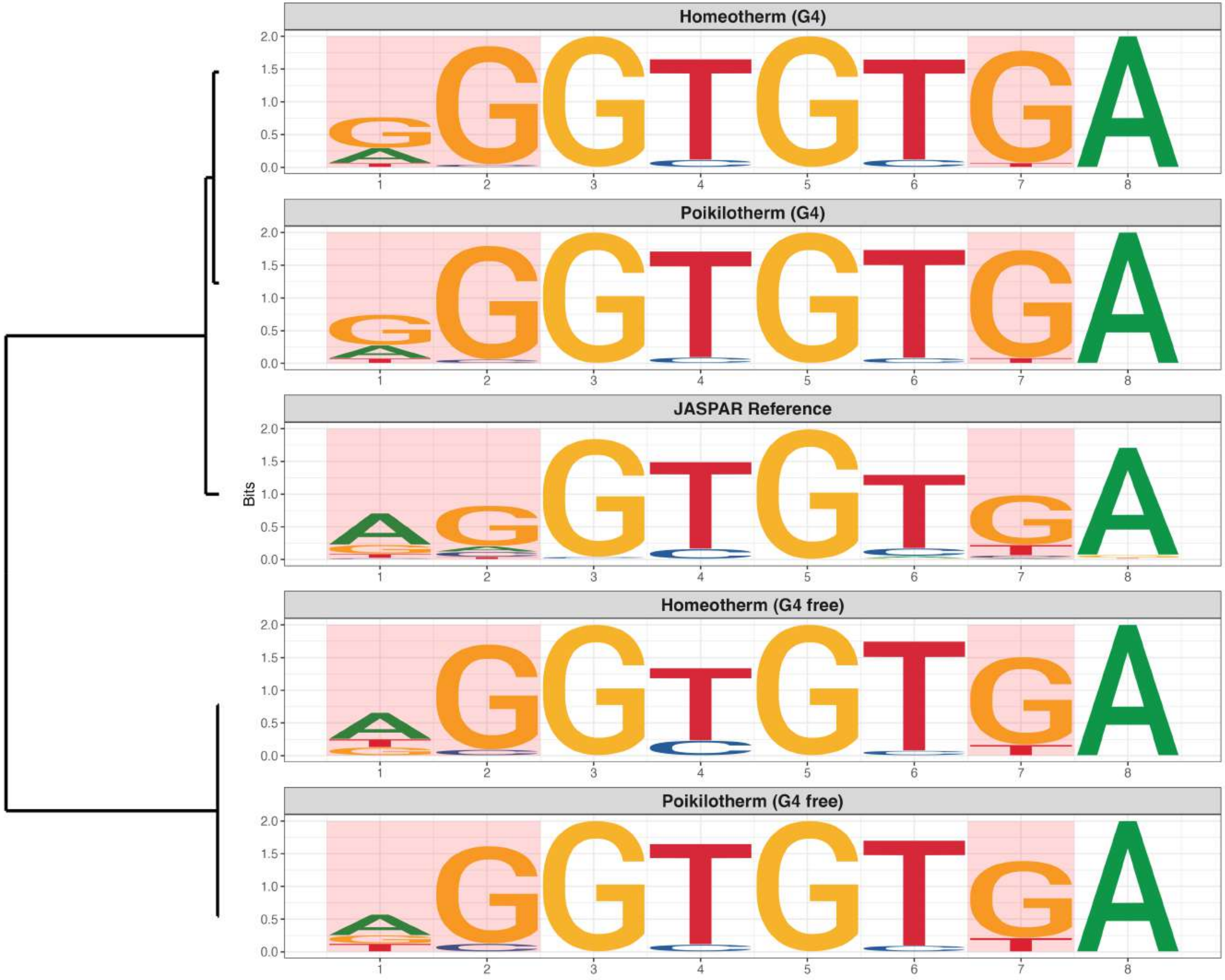

**Promoter : Moderately Enriched in G4s**

*MA0807.1 - TBX5*

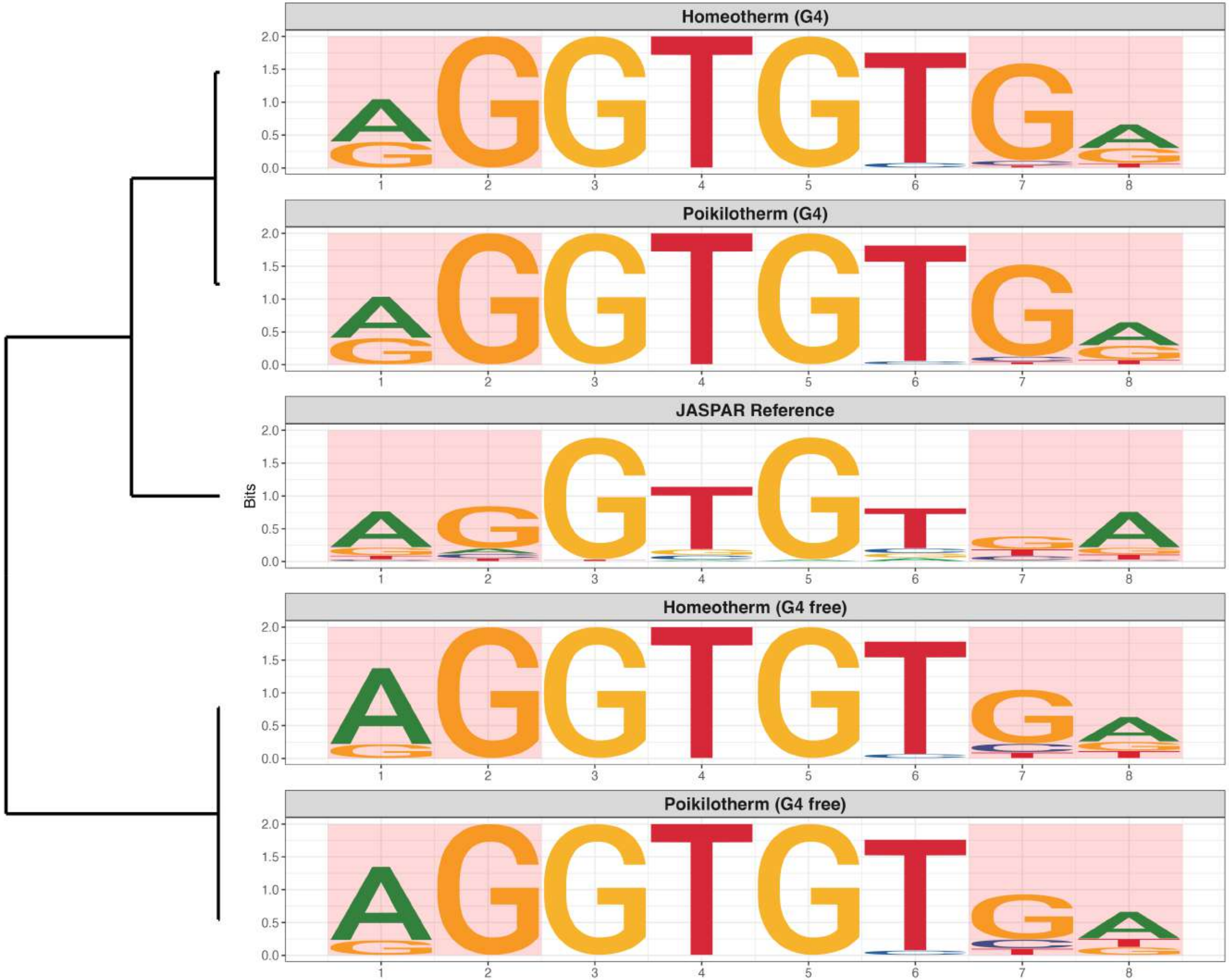

**Promoter : Moderately Enriched in G4s**

*MA0812.2 - TFAP2B*

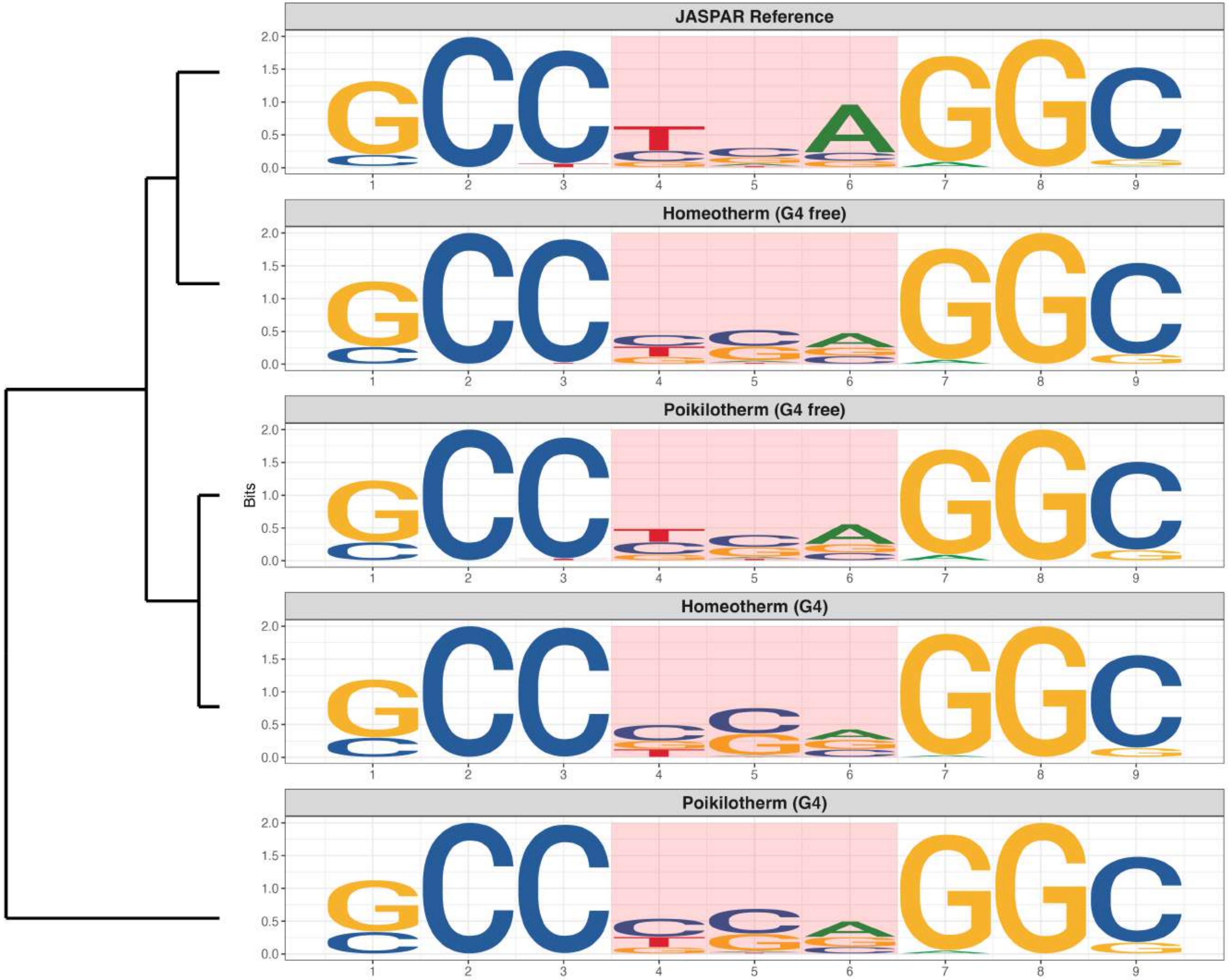

### Promoter : Moderately Enriched in G4s

*MA0814.3 - TFAP2C*

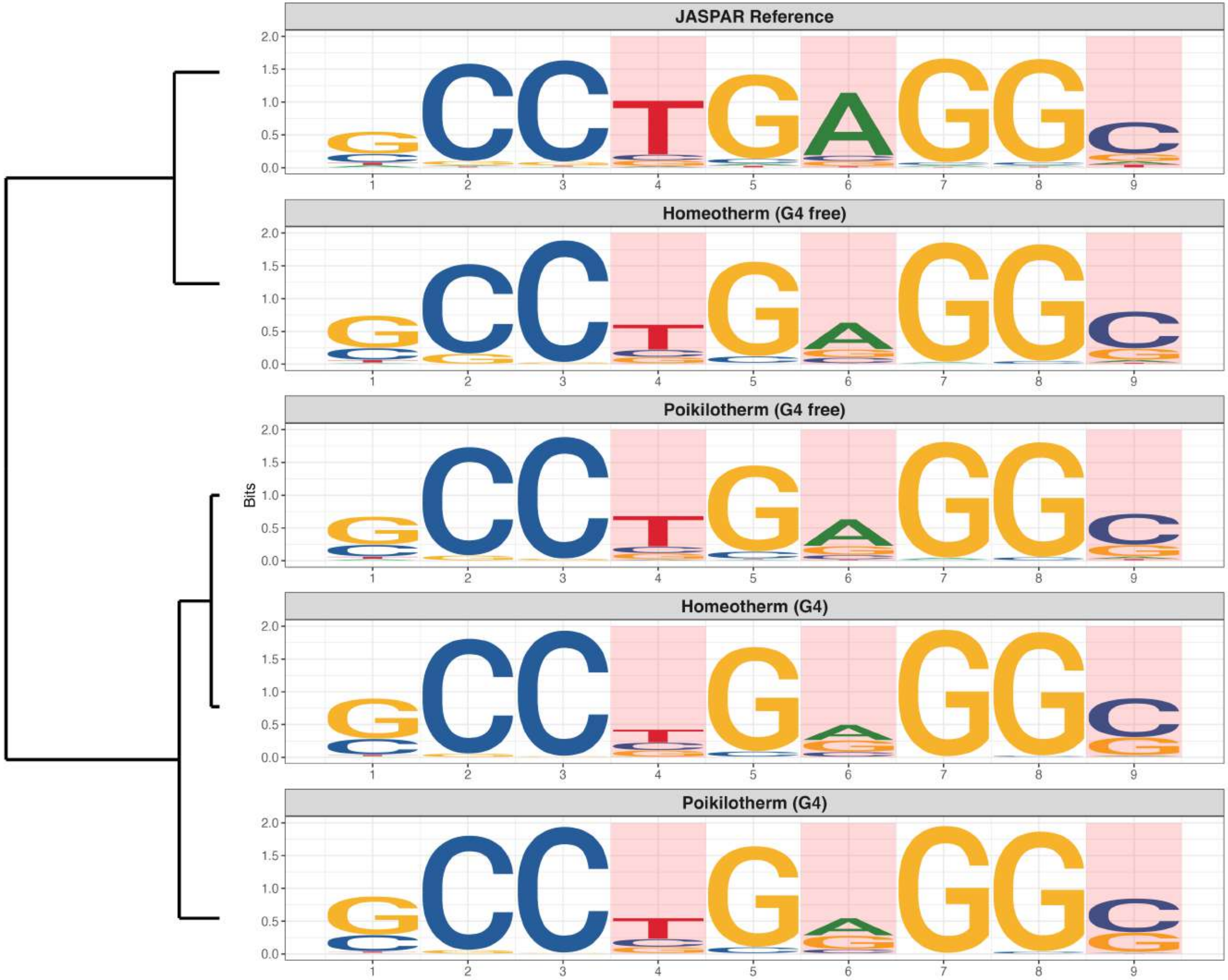

**Promoter : Moderately Enriched in G4s**

*MA0821.2 - HES5*

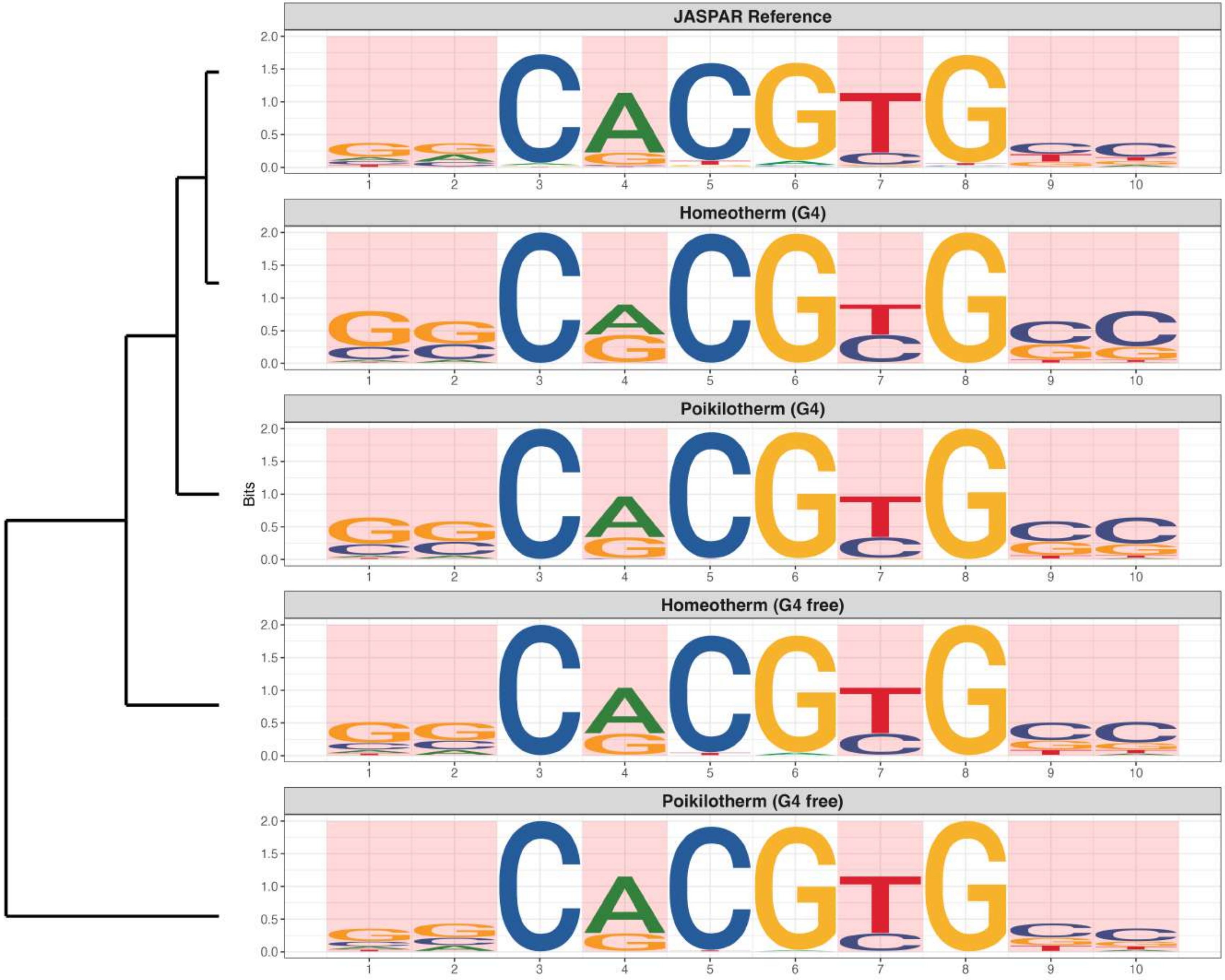

### Promoter : Moderately Enriched in G4s

*MA0857.1 - Rarb*

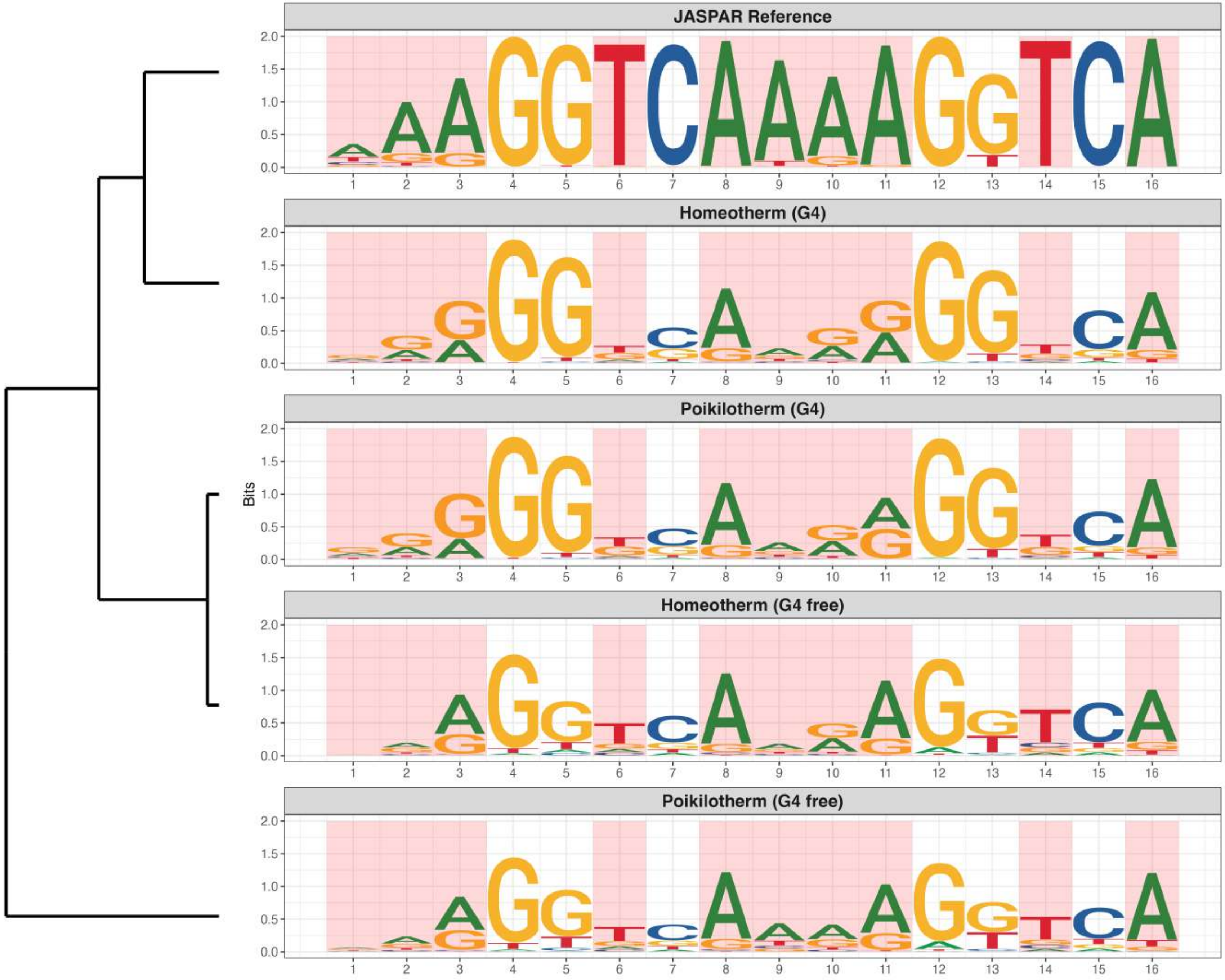

**Promoter : Moderately Enriched in G4s**

*MA1099.3 - HES1*

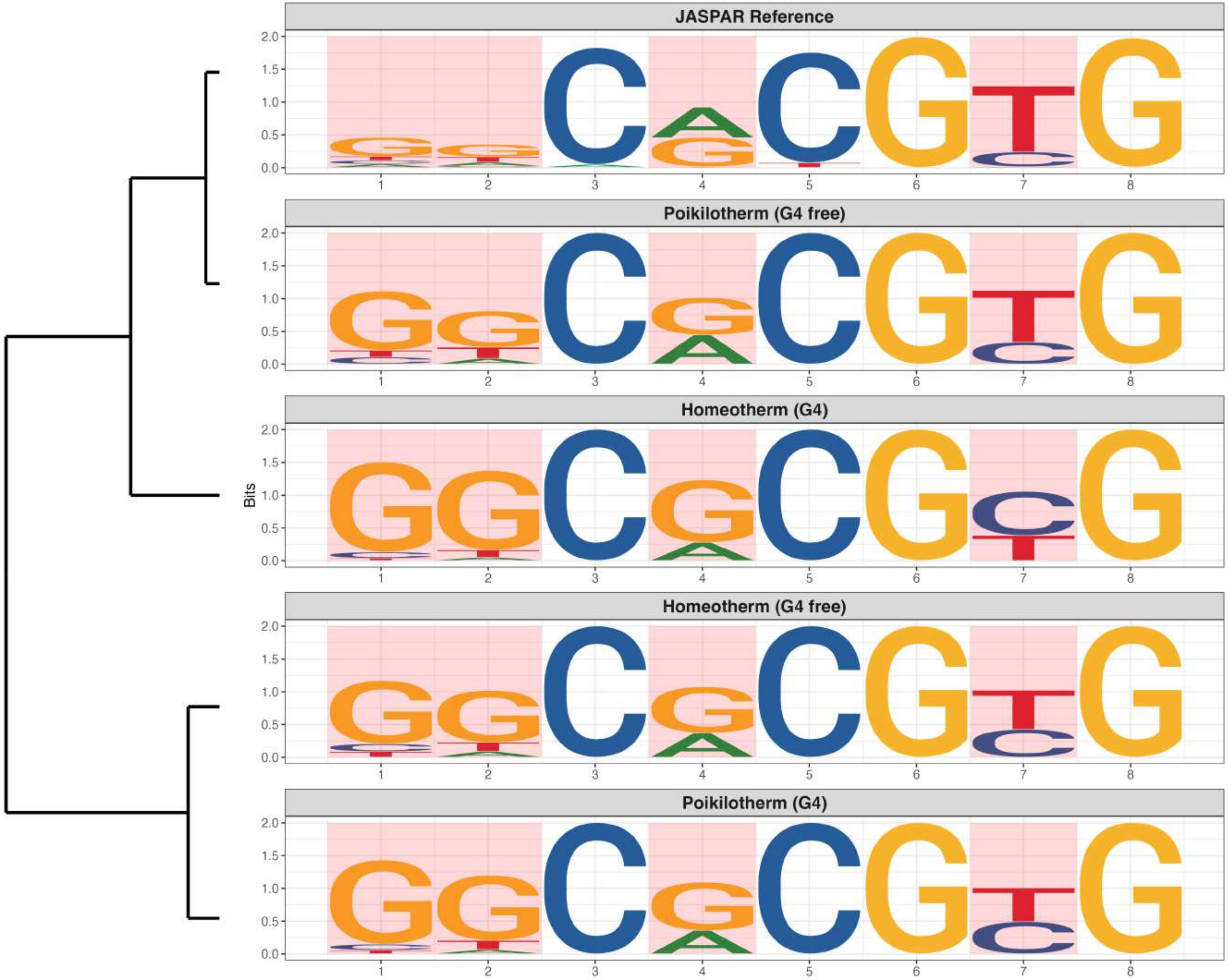

**Promoter : Moderately Enriched in G4s**

*MA1114.2 - PBX3*

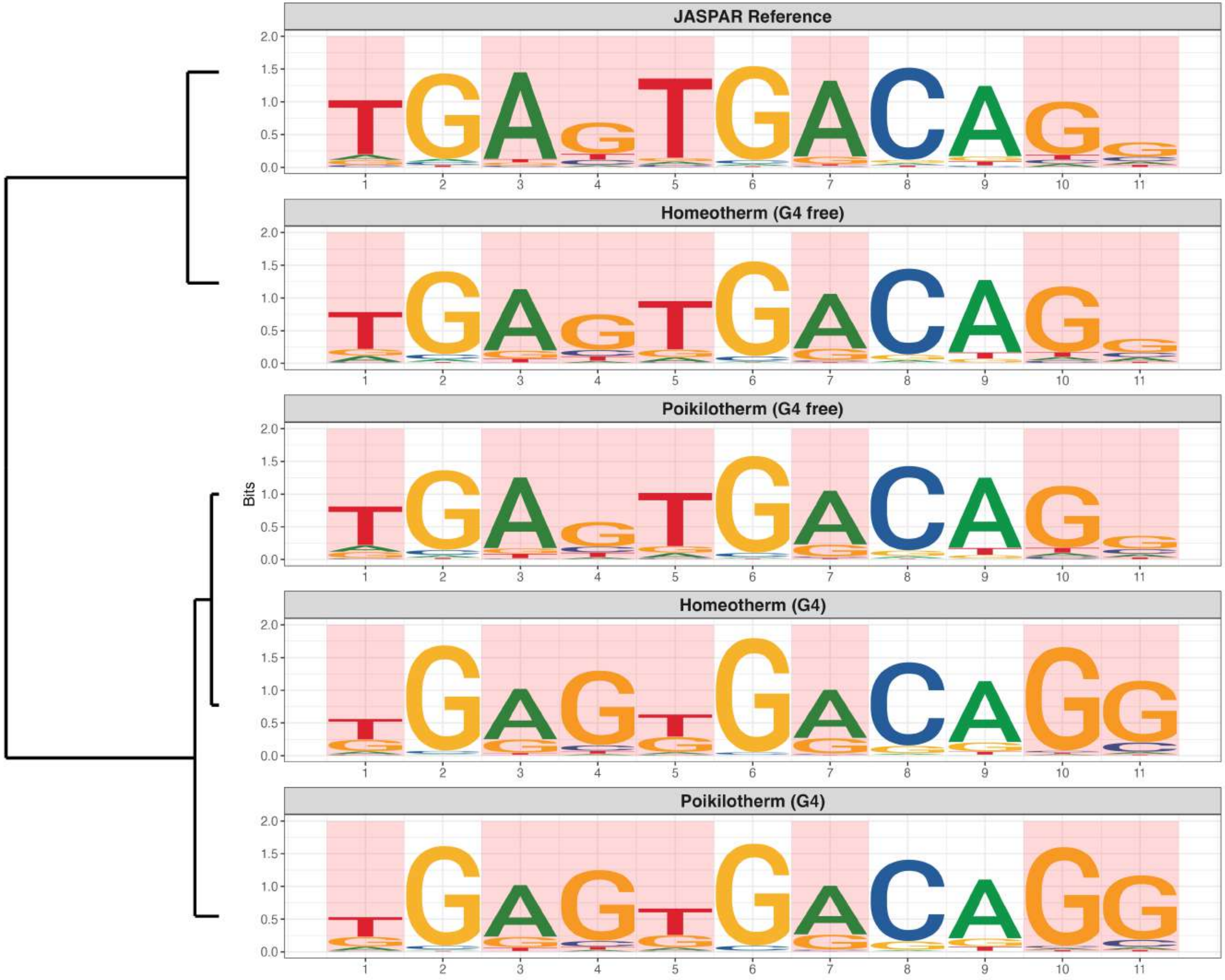

**Promoter : Moderately Enriched in G4s**

*MA1125.2 - ZNF384*

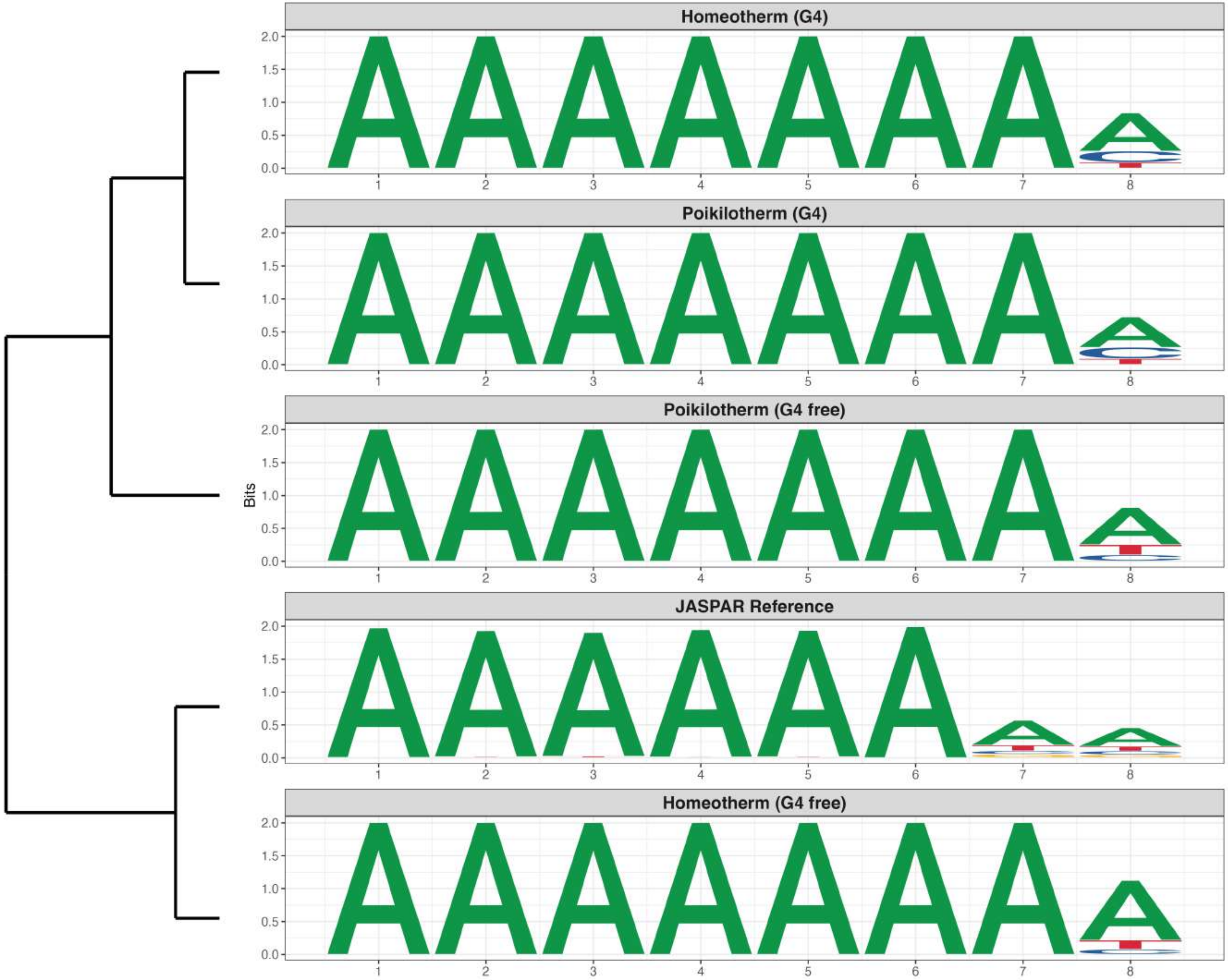

**Promoter : Moderately Enriched in G4s**

*MA1513.2 - KLF15*

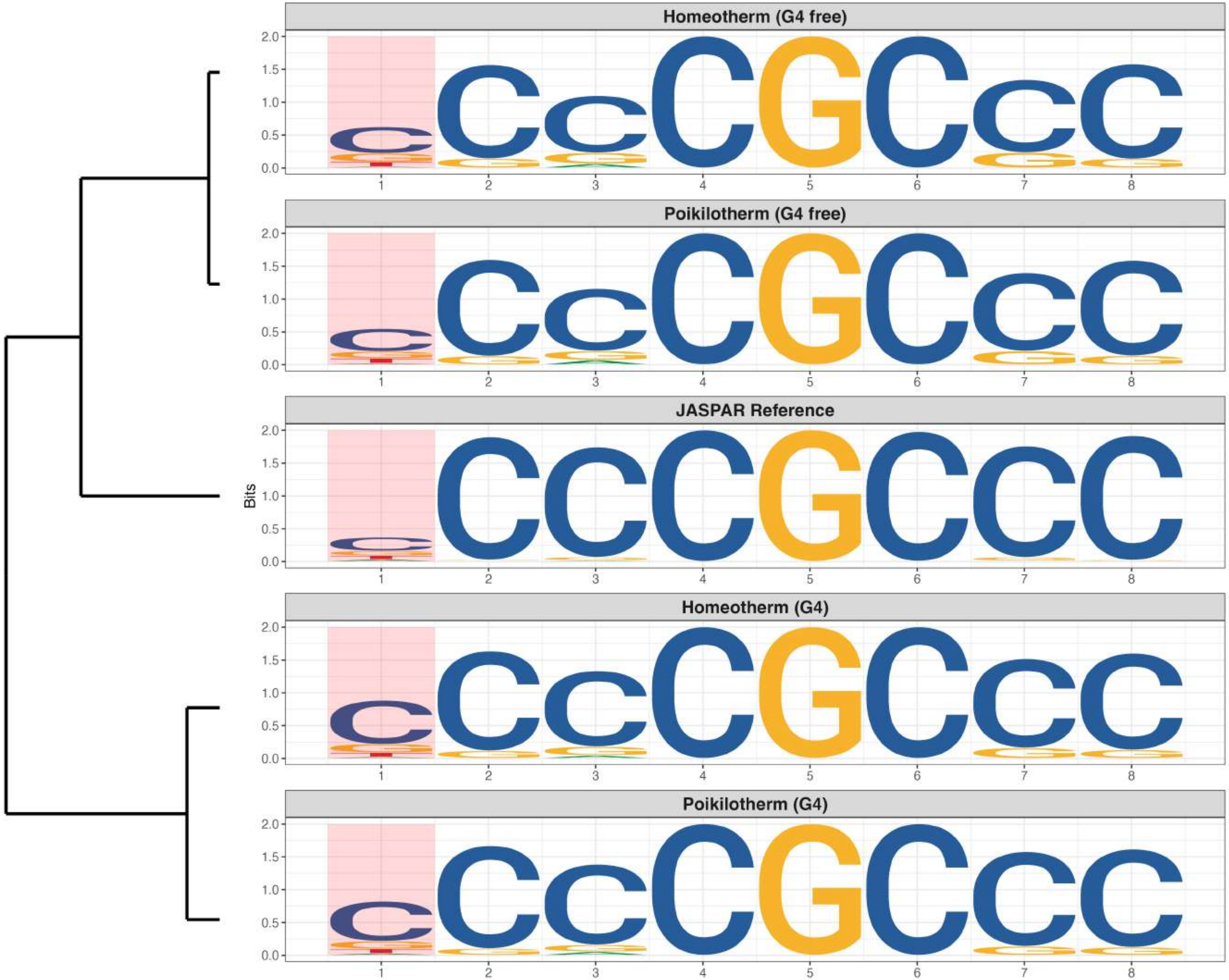

### Promoter : Moderately Enriched in G4s

*MA1516.2 - KLF3*

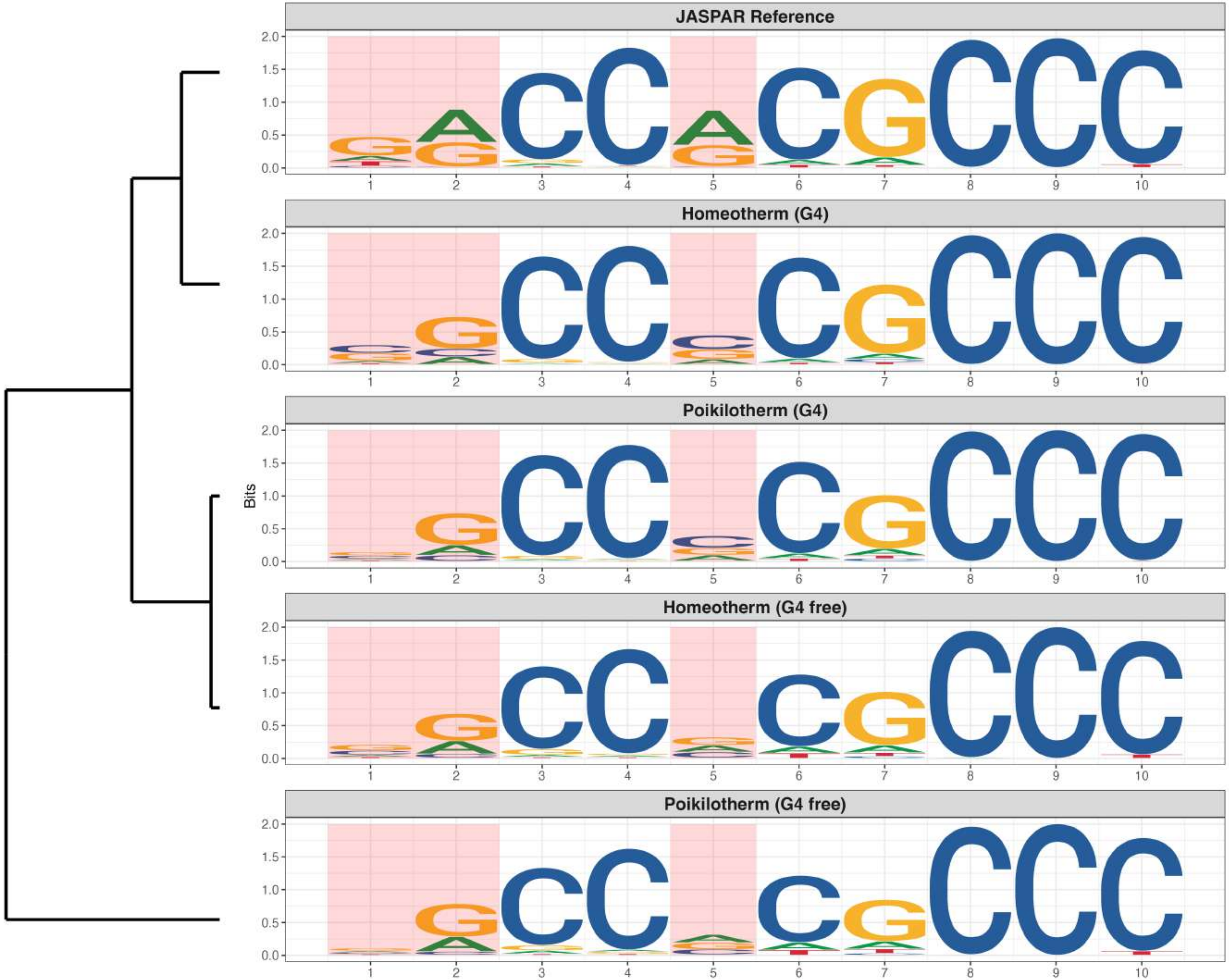

**Promoter : Moderately Enriched in G4s**

*MA1547.2 - PITX2*

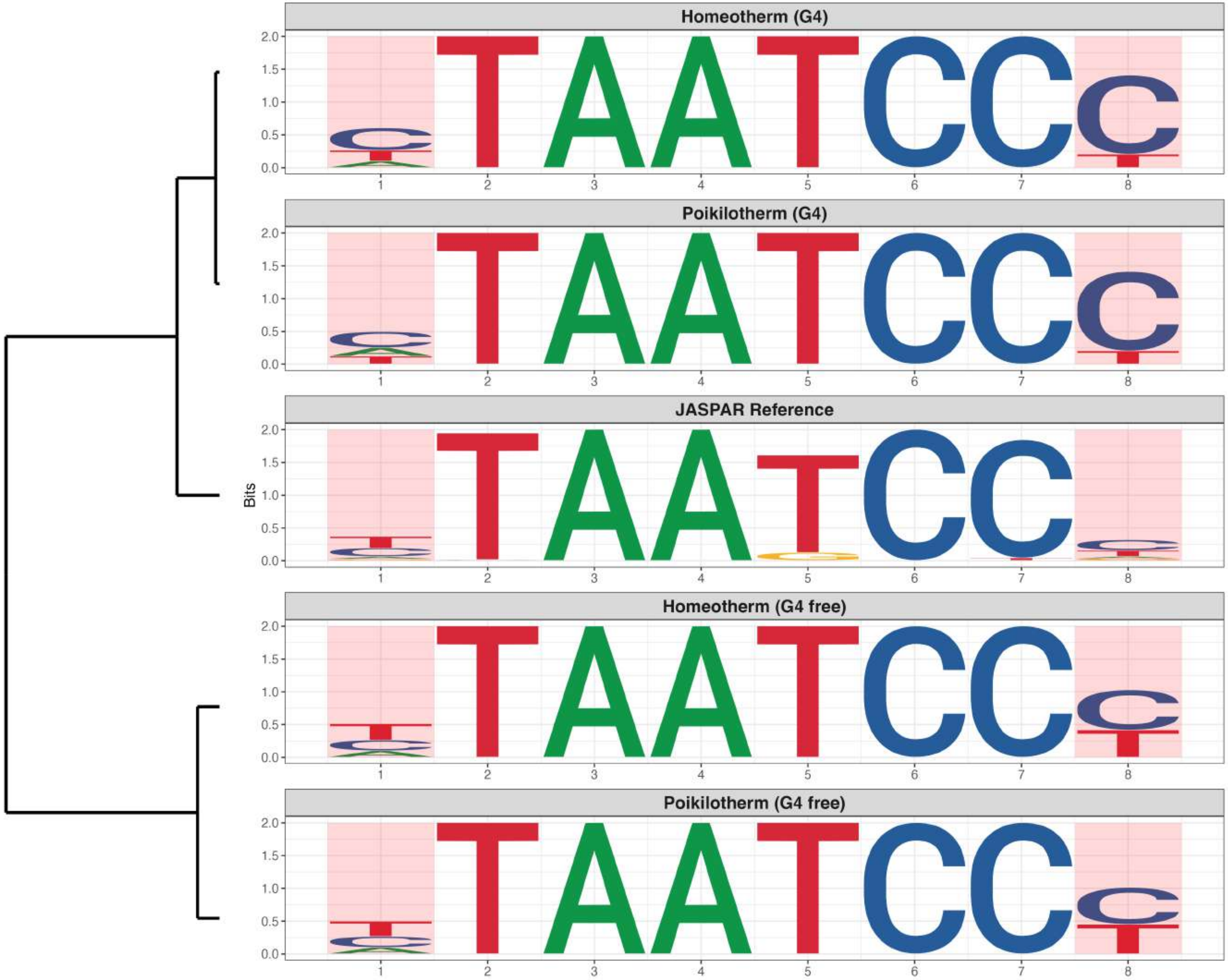

### Promoter : Moderately Enriched in G4s

*MA1553.2 - RARG*

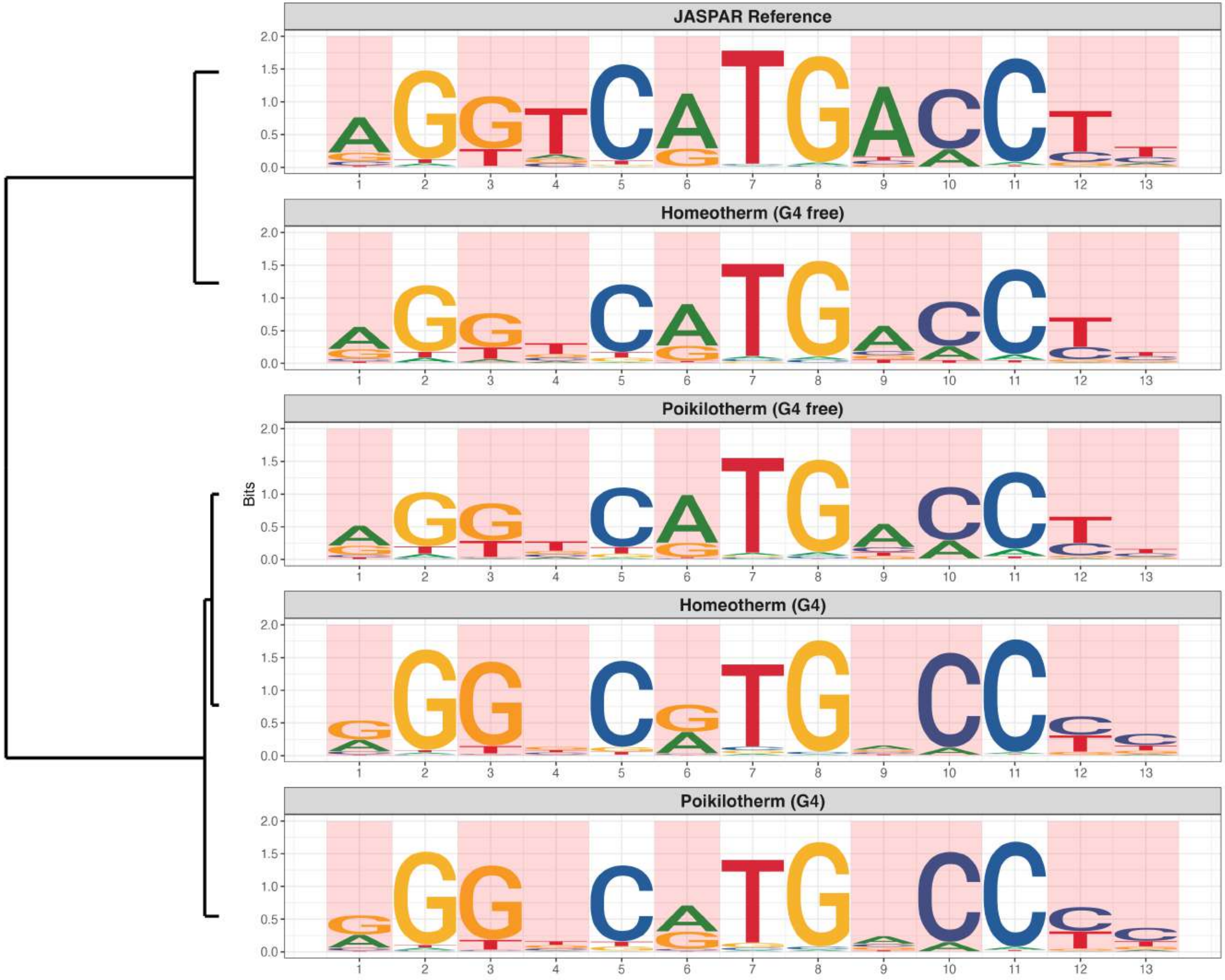

**Promoter : Moderately Enriched in G4s**

*MA1559.2 - SNAI3*

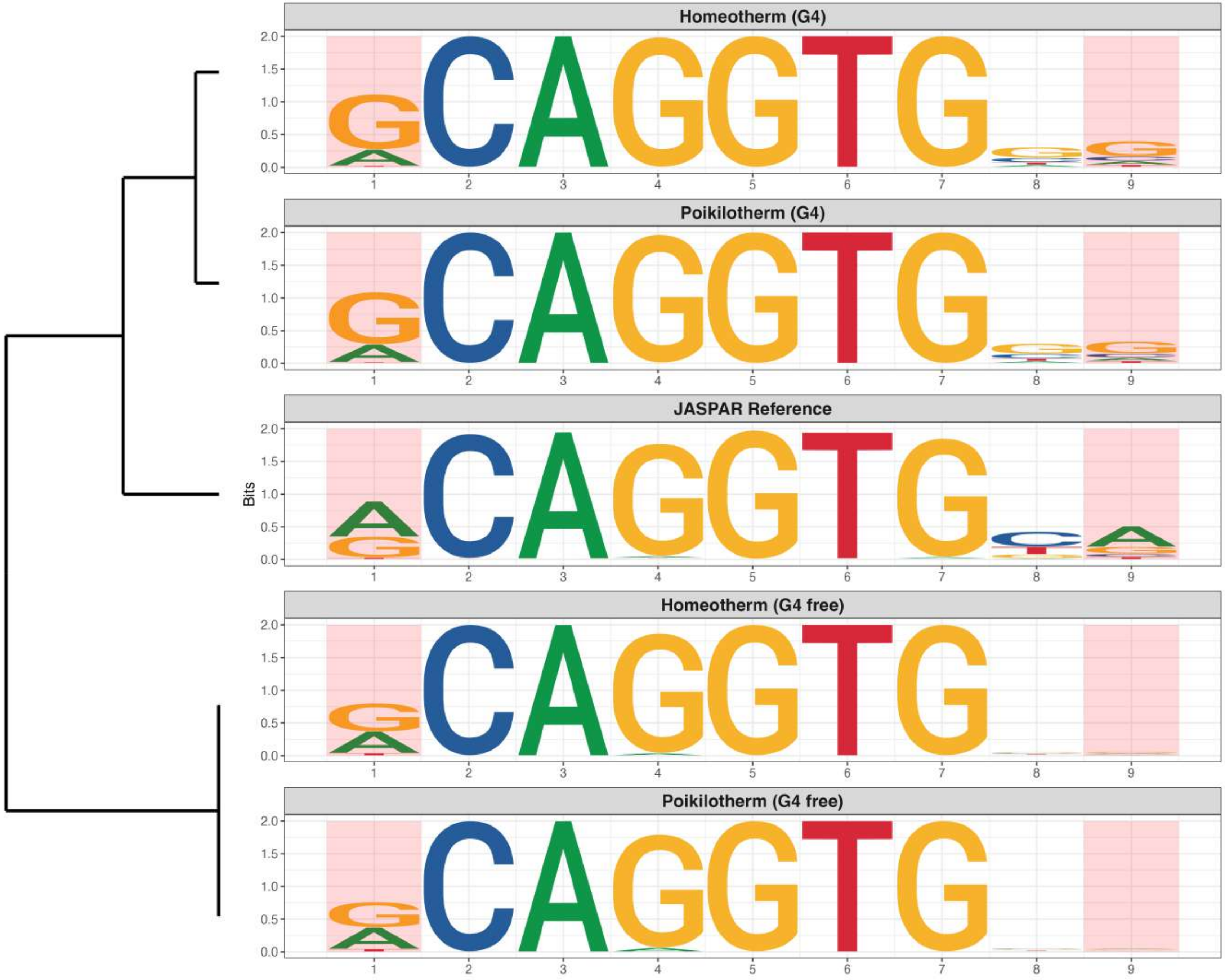

**Promoter : Moderately Enriched in G4s**

*MA1585.2 - ZKSCAN1*

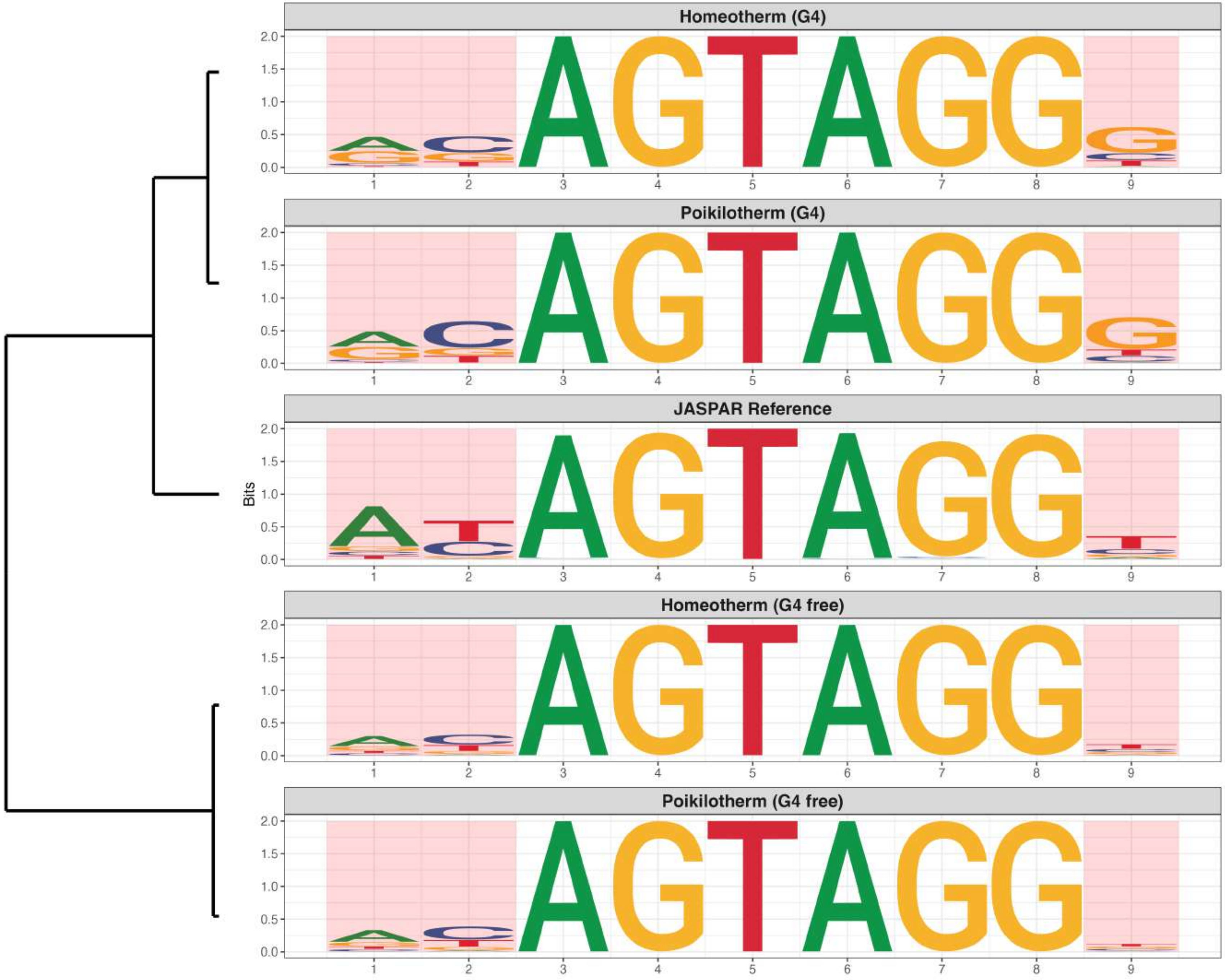

**Promoter : Moderately Enriched in G4s**

*MA1713.2 - ZNF610*

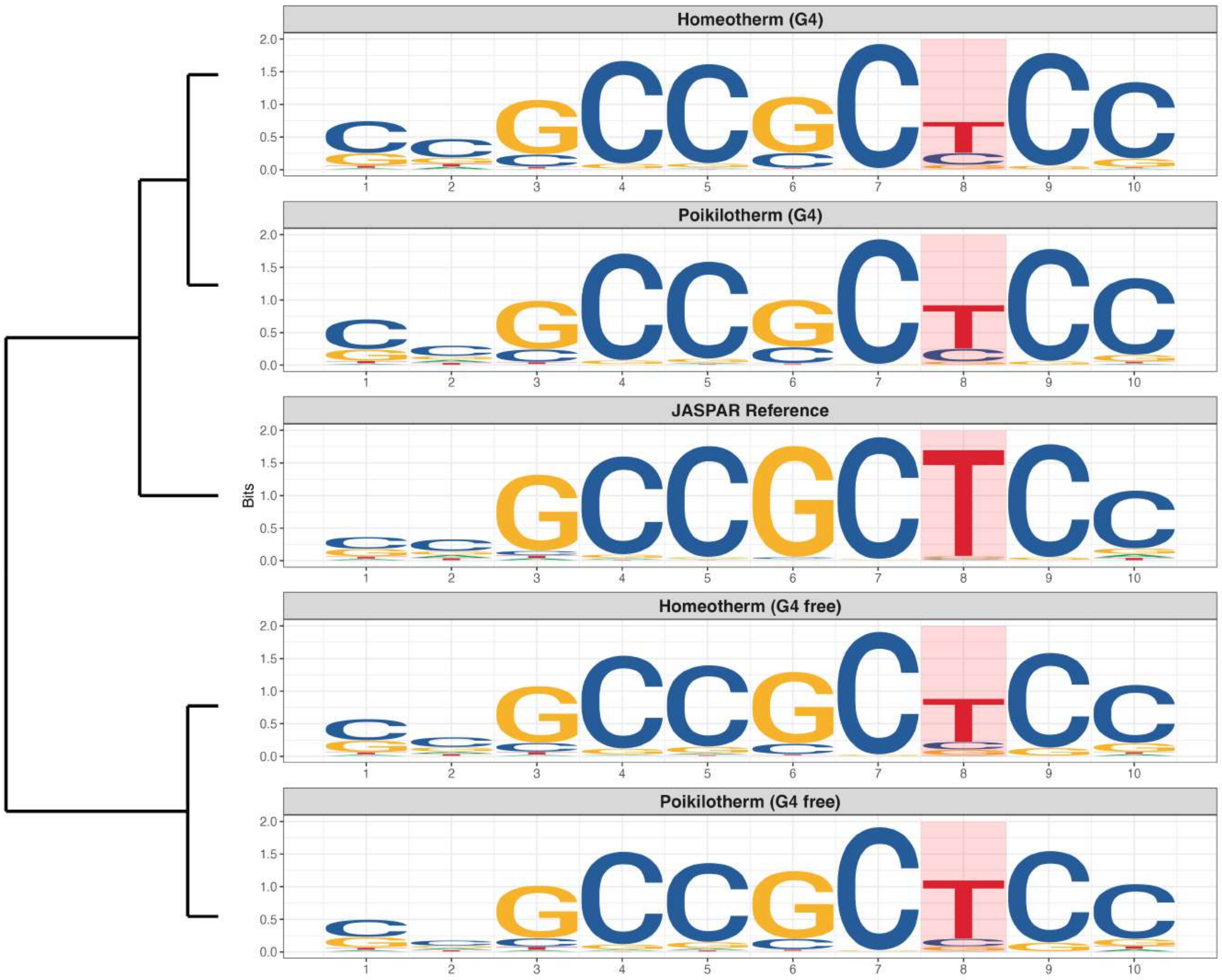

### Promoter : Moderately Enriched in G4s

*MA1721.2 - ZNF93*

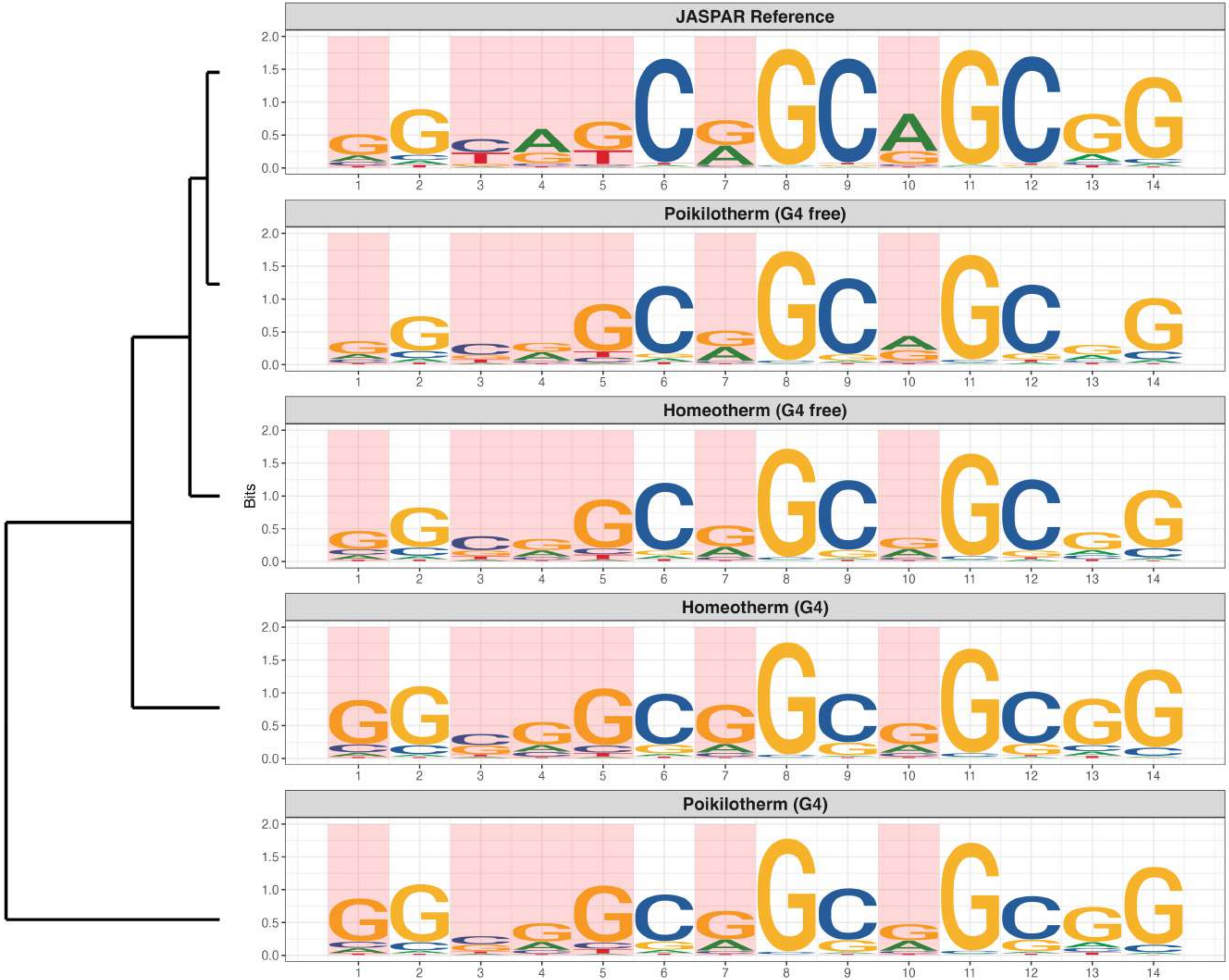

**Promoter : Moderately Enriched in G4s**

*MA1998.2 - Prdm14*

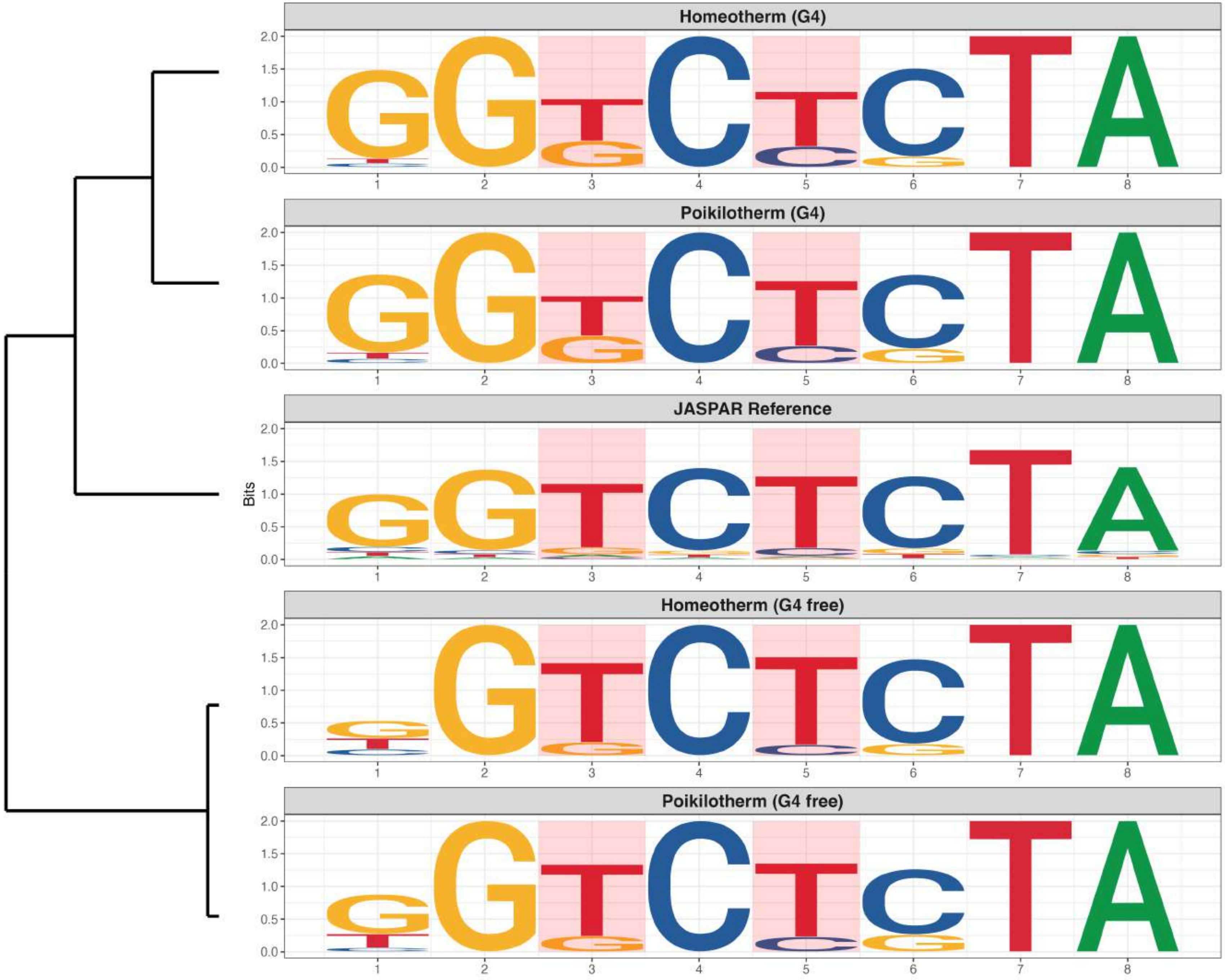

**Promoter : Moderately Enriched in G4s**

*MA2328.1 - ZBED4*

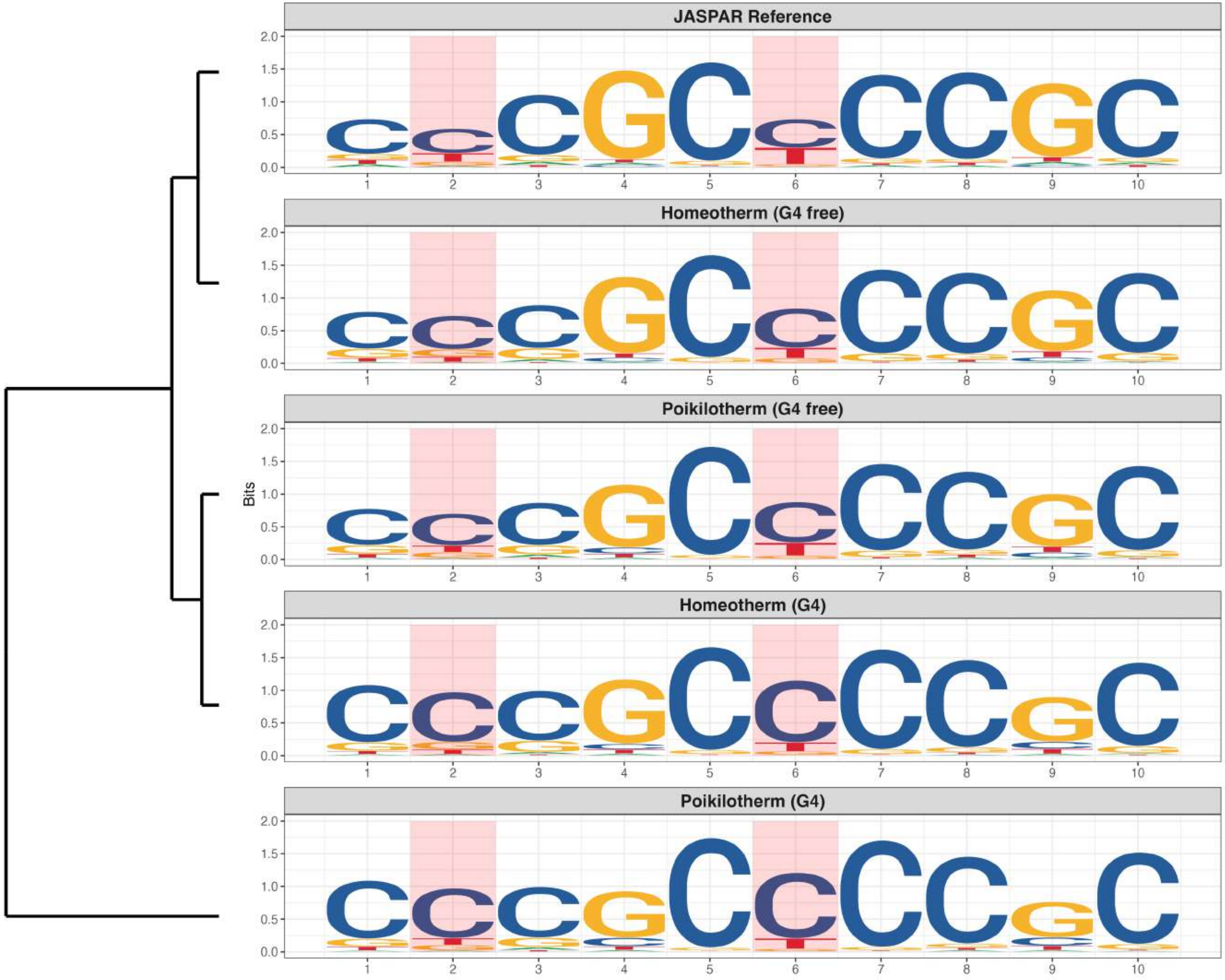

**Promoter : Moderately Enriched in G4s**

*MA2340.1 - Zbtb2*

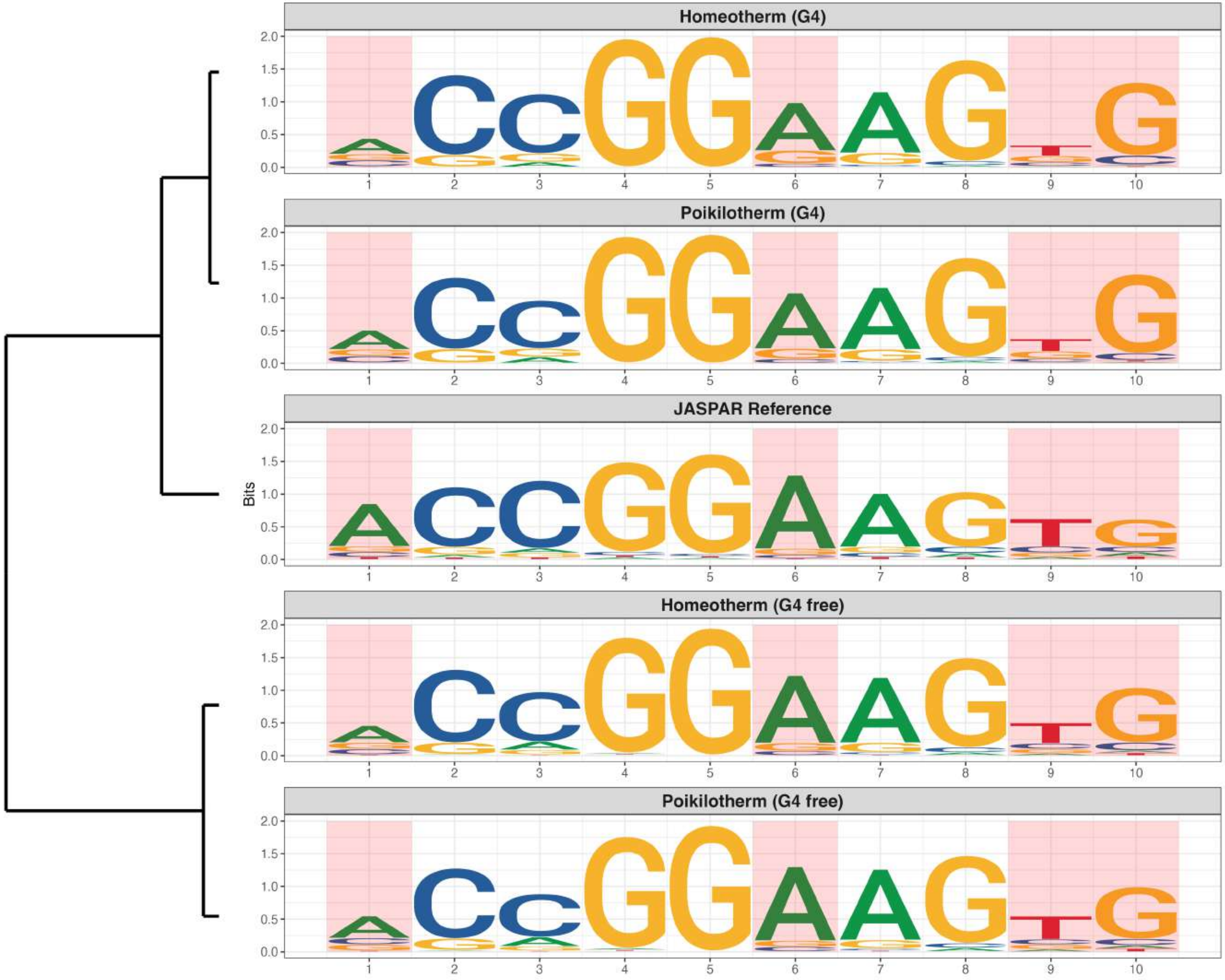

**Promoter : Moderately Enriched in G4s**

*MA2489.1 - ZFP28*

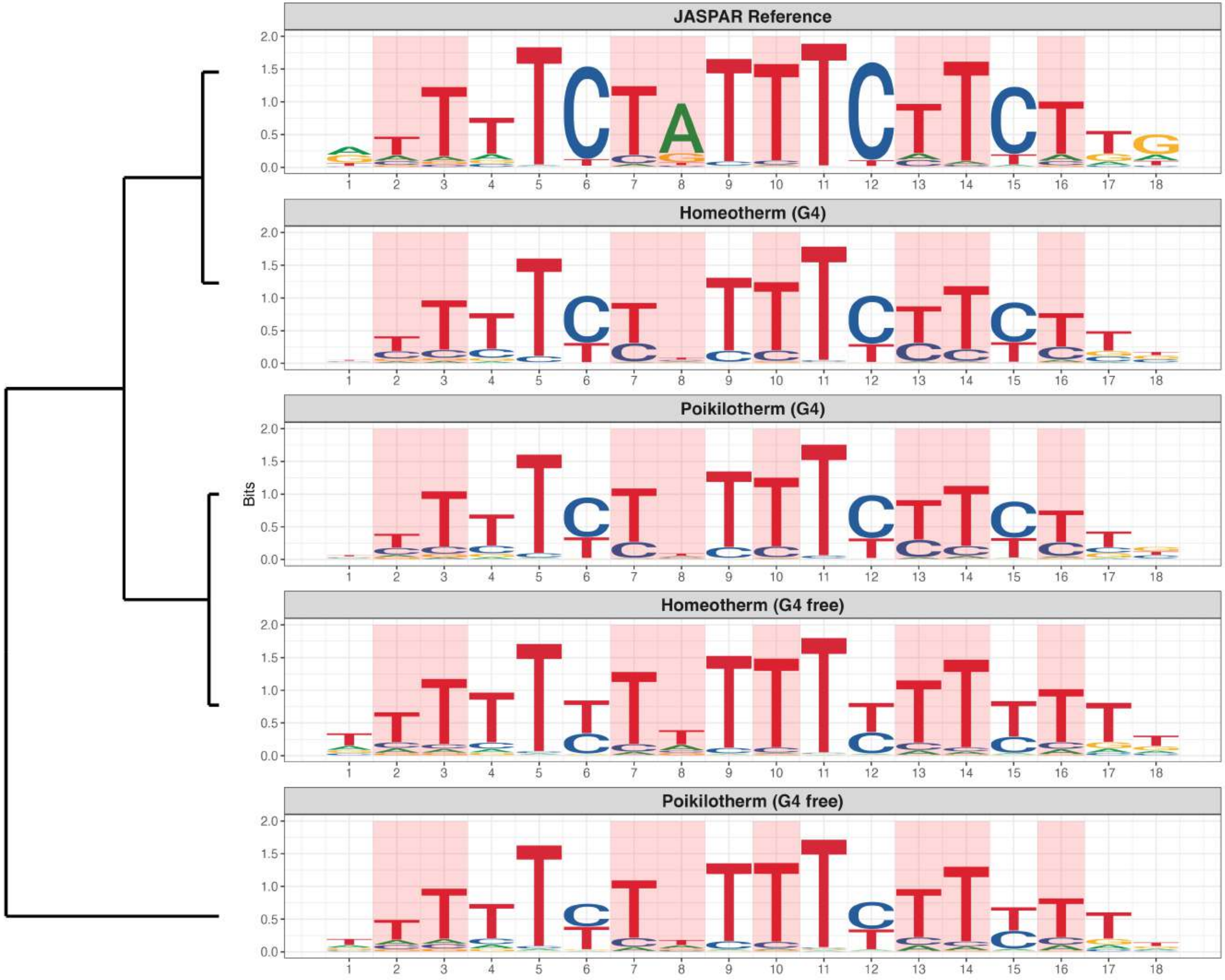

**Promoter : Moderately Enriched in G4s**

*MA2493.1 - ZNF322*

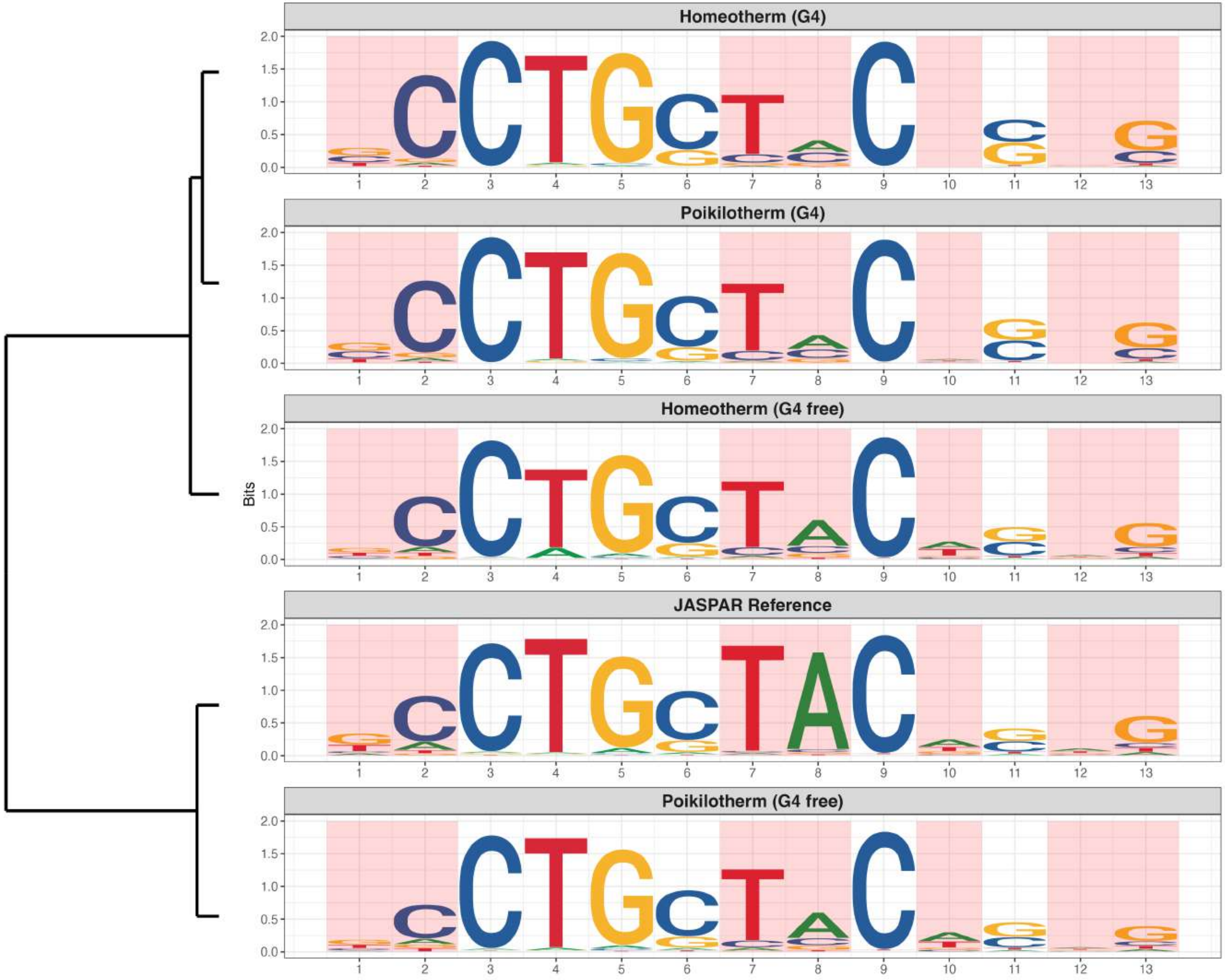

**Promoter : Moderately Enriched in G4s**

*MA2524.1 - ZNF696*

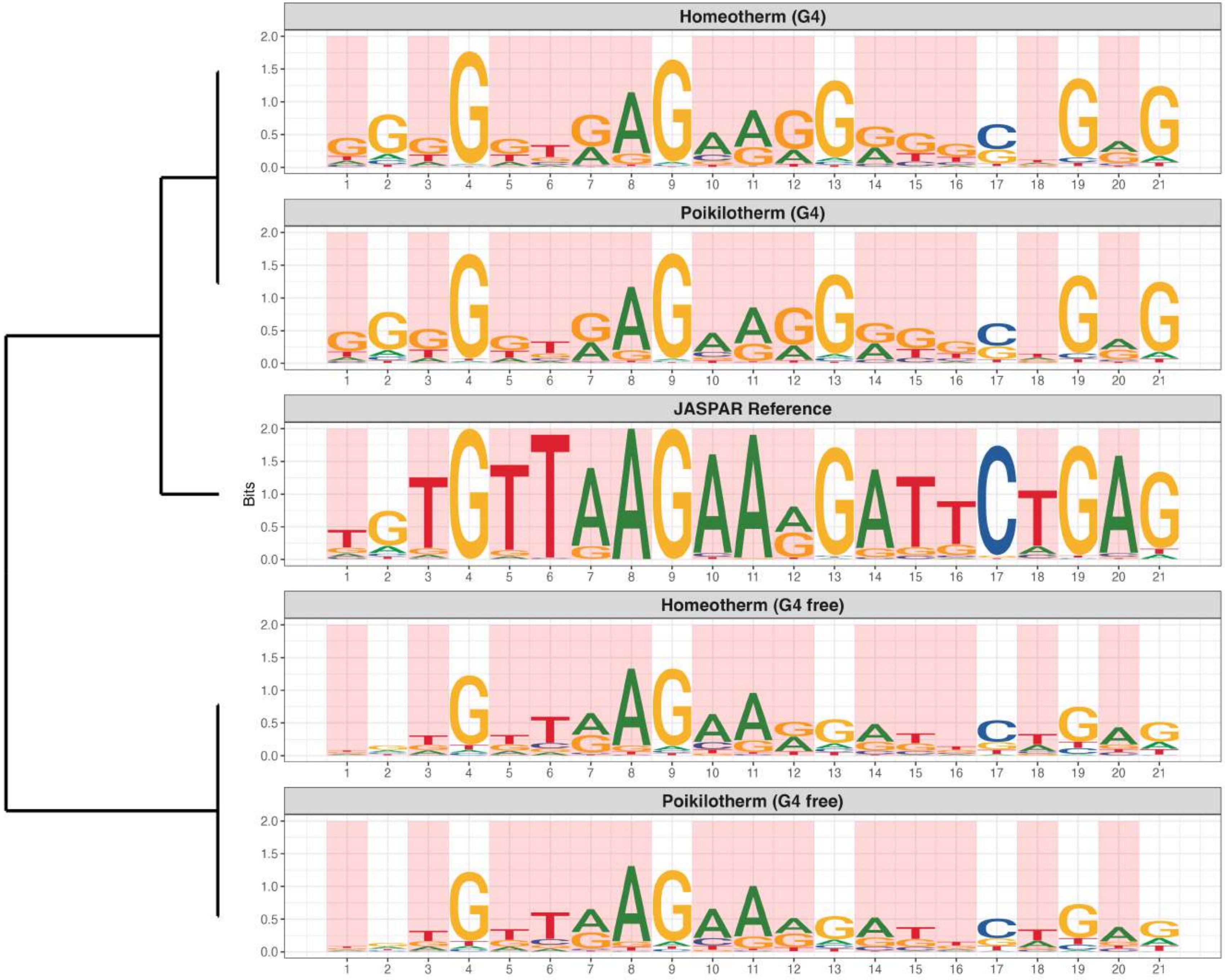

**Promoter : Moderately Enriched in G4s**

*MA2538.1 - CGGBP1*

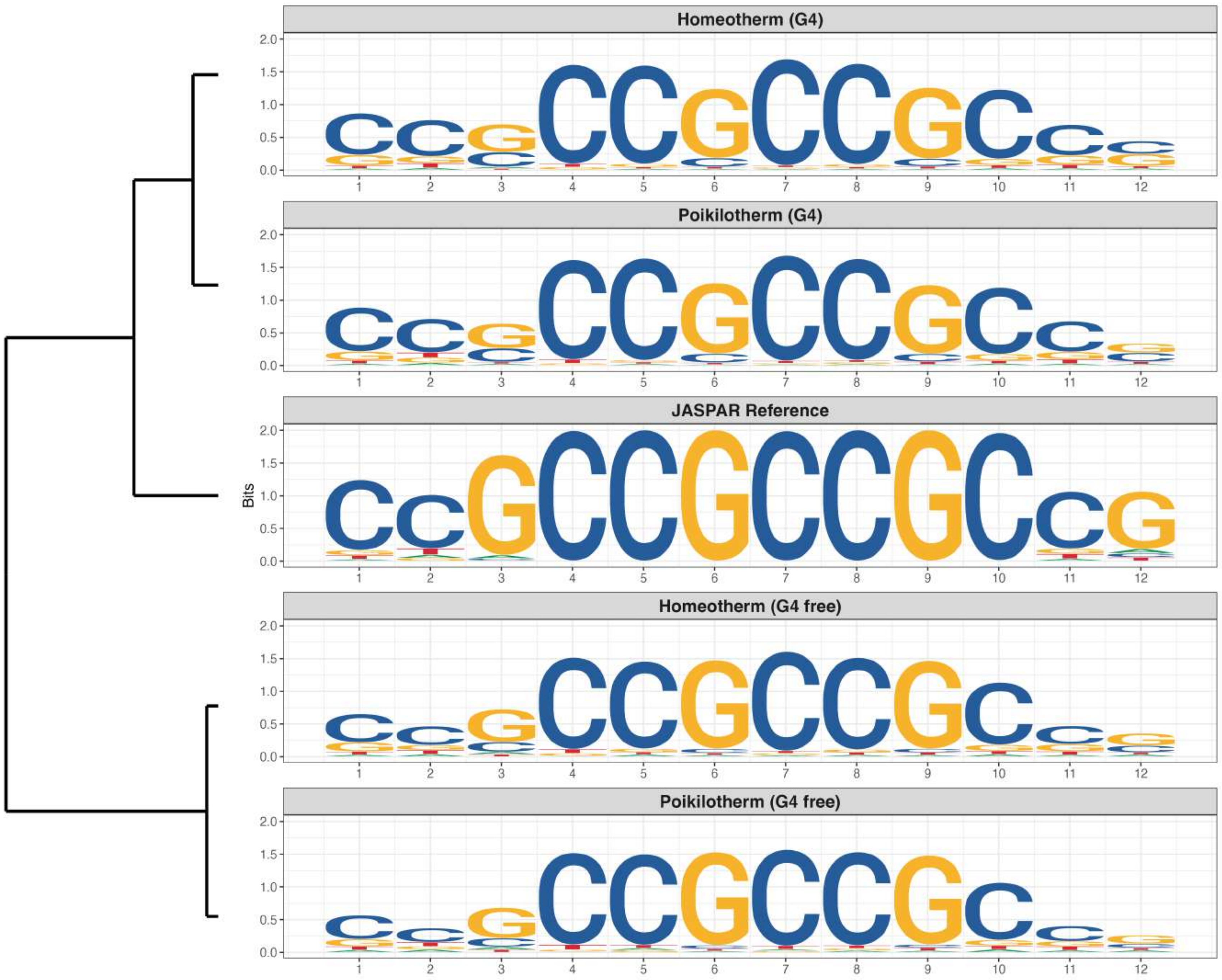

### Promoter : Moderately Enriched in G4s

*MA2539.1 - SLC2A4RG*

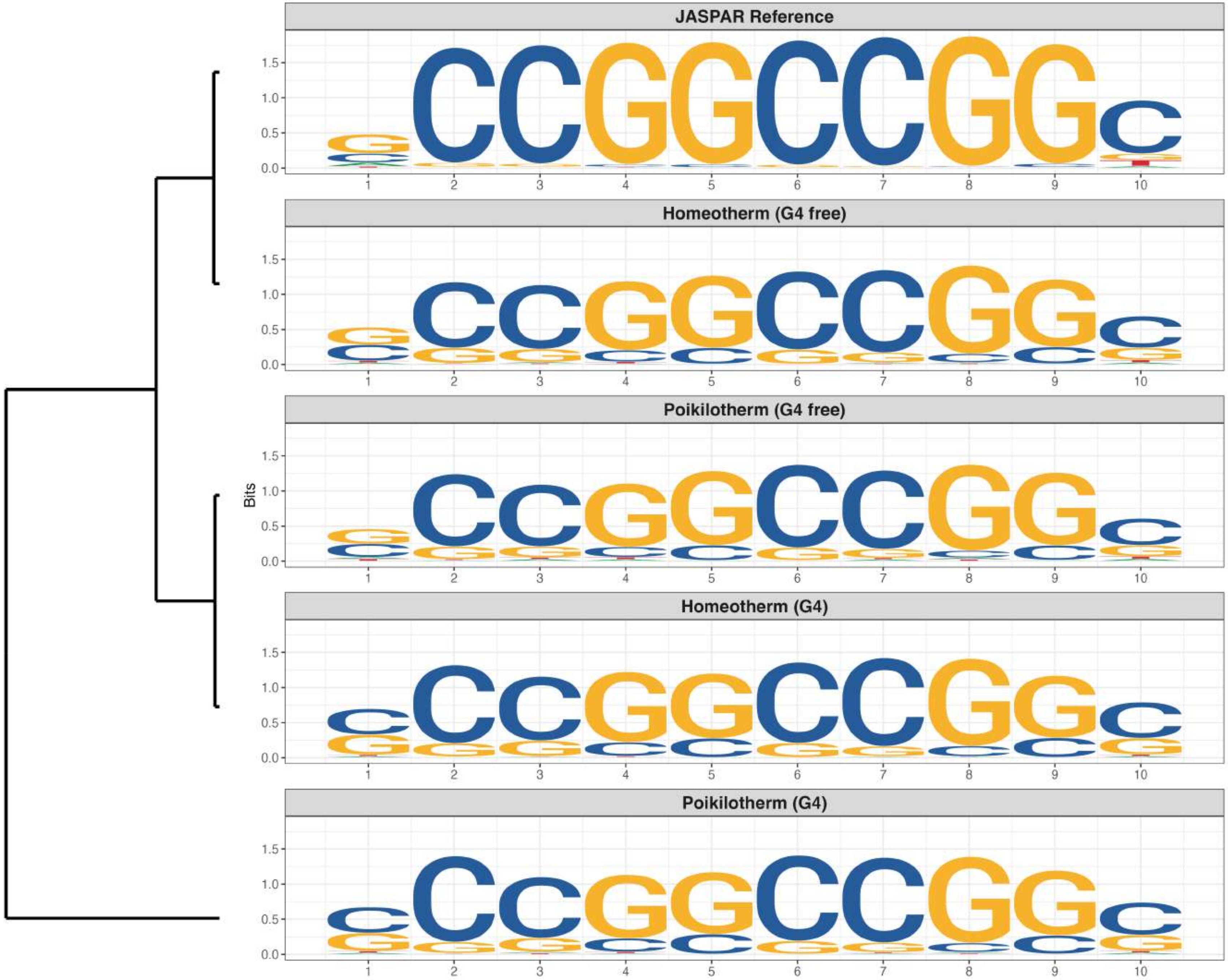

### Promoter : Moderately Enriched in G4s

*MA2553.1 - ZNF407*

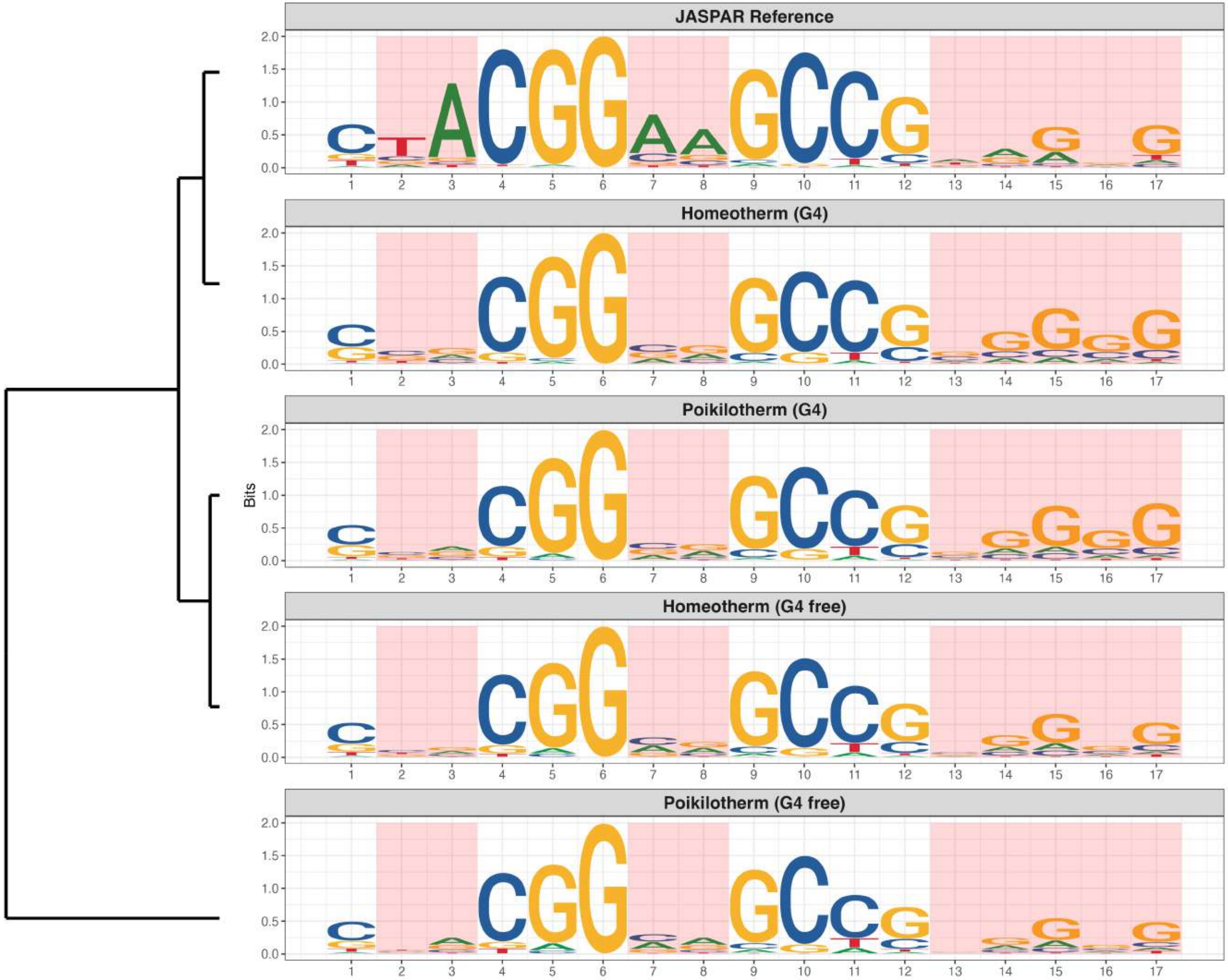

**Promoter : Moderately Enriched in G4s**

*MA2580.1 - ZNF20*

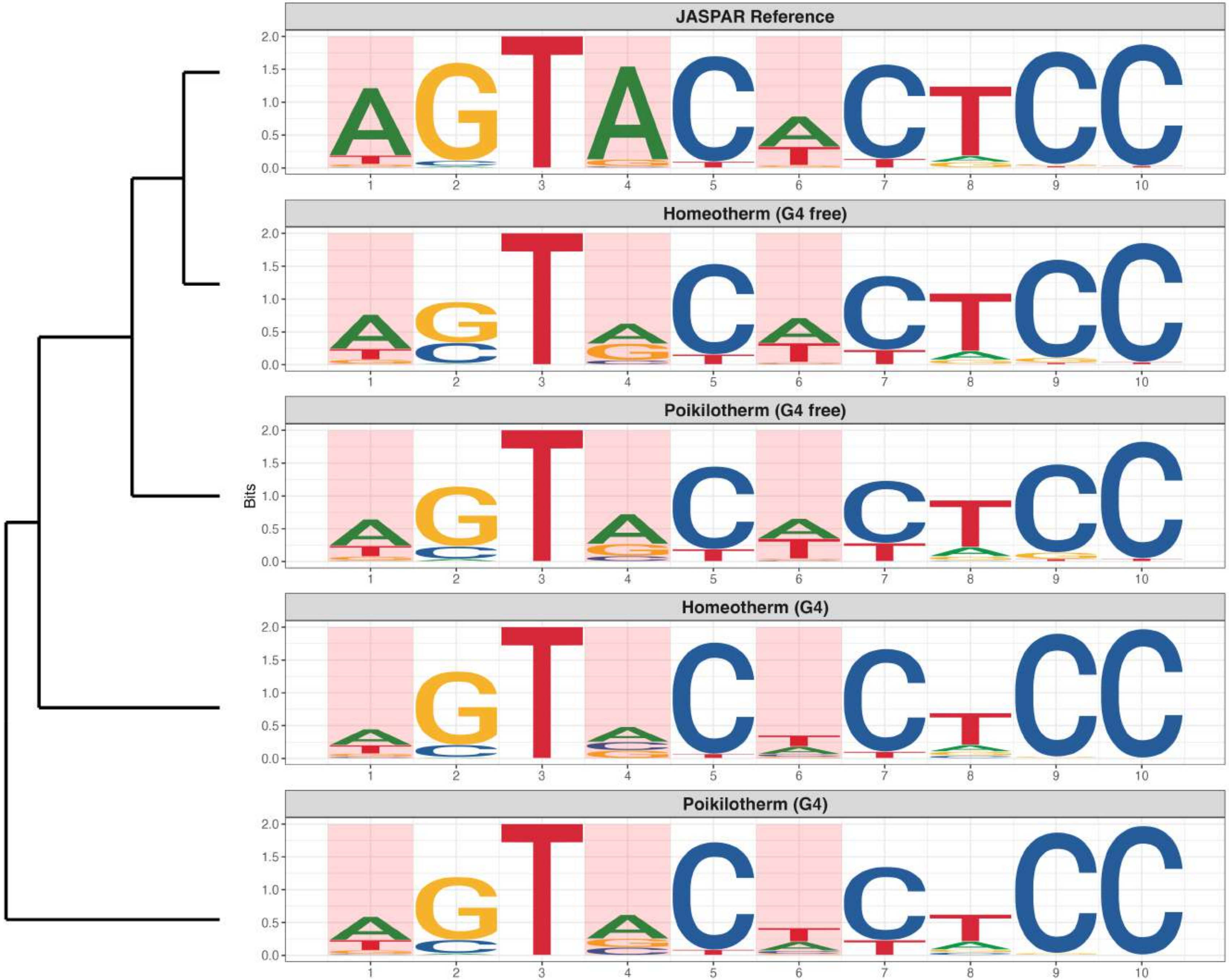

**Promoter : Moderately Enriched in G4s**

*MA2593.1 - ZNF362*

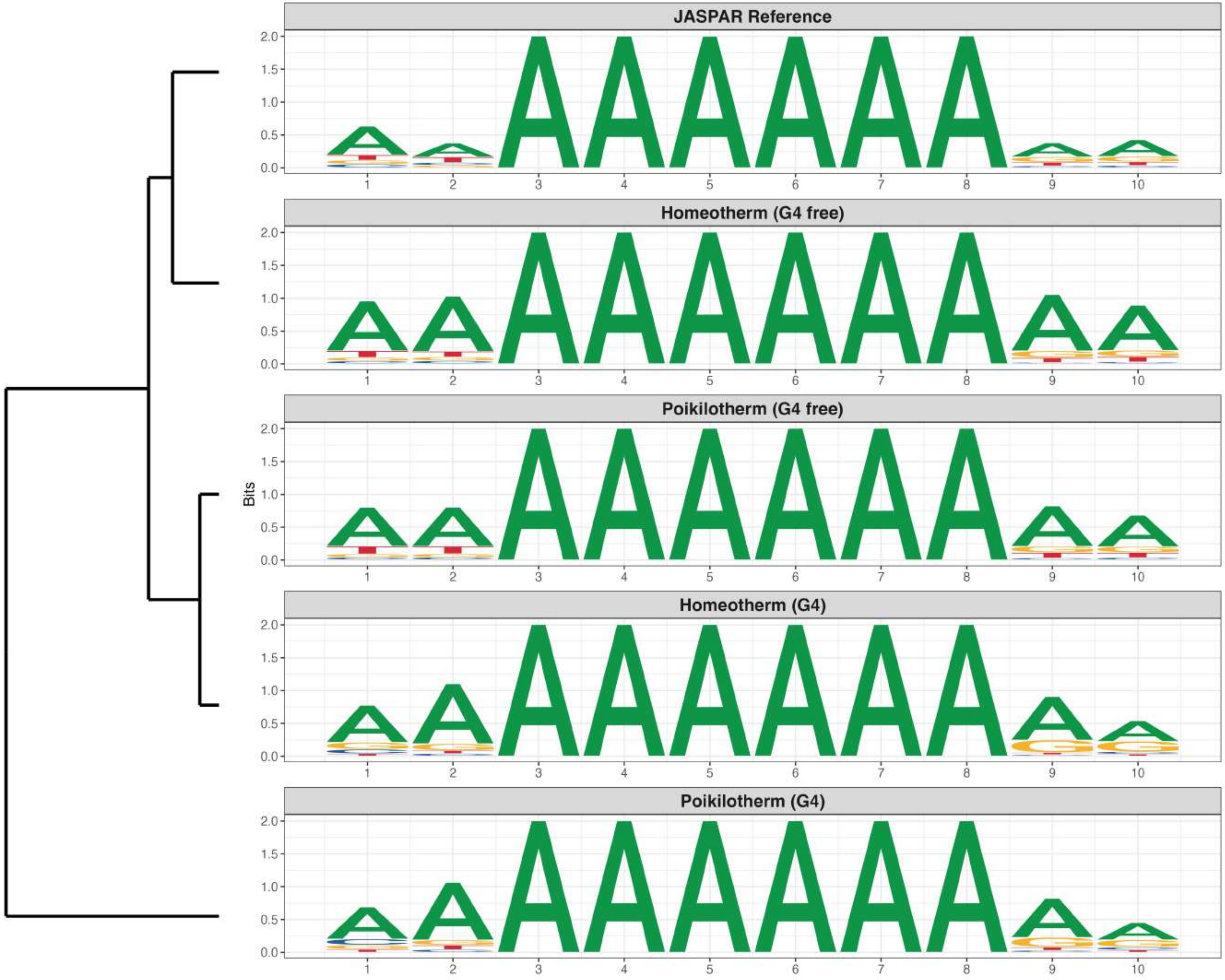

**Promoter : Moderately Enriched in G4s**

*MA2602.1 - ZBED5*

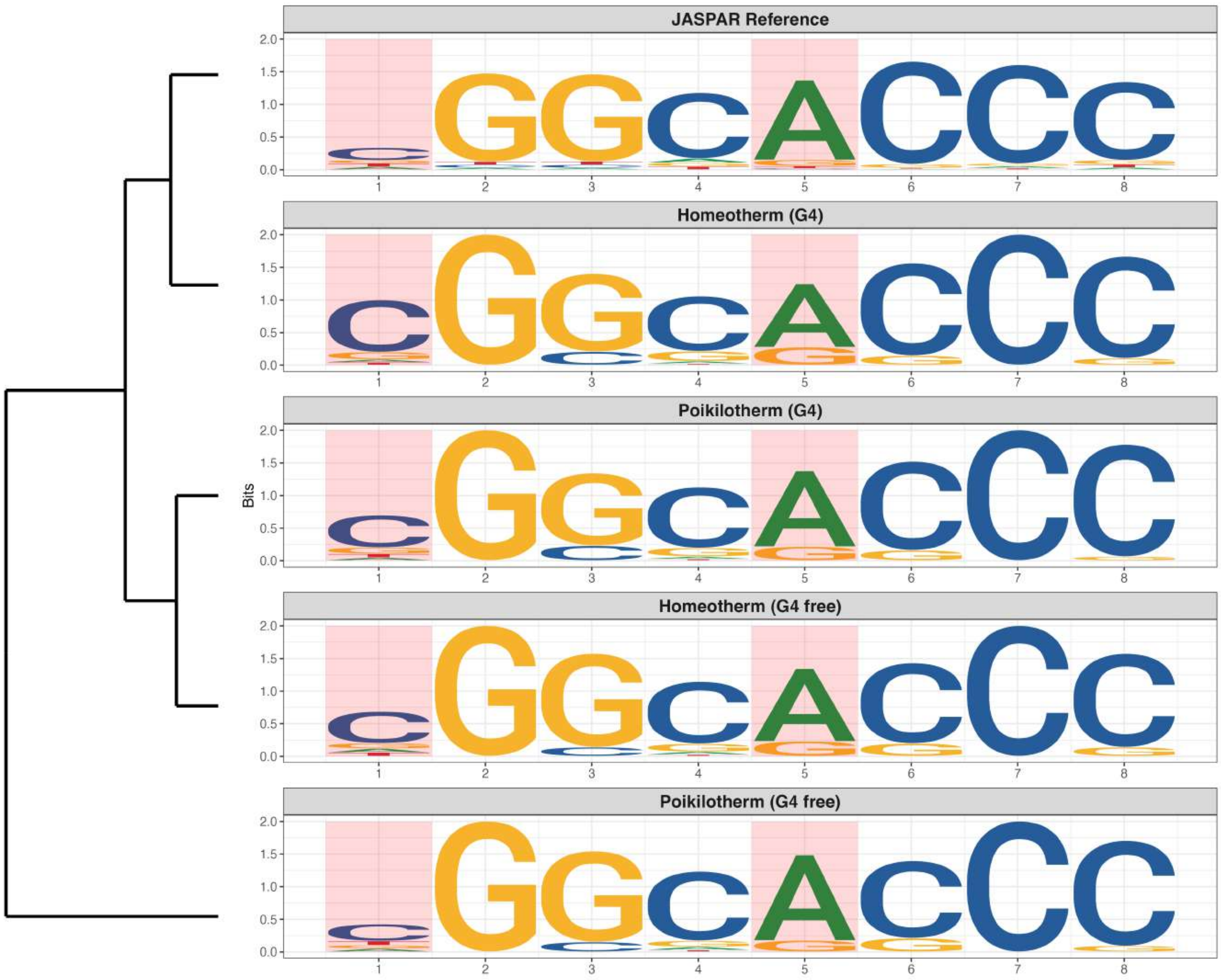

**Promoter : Highly Reduced in G4s**

*MA0028.3 - ELK1*

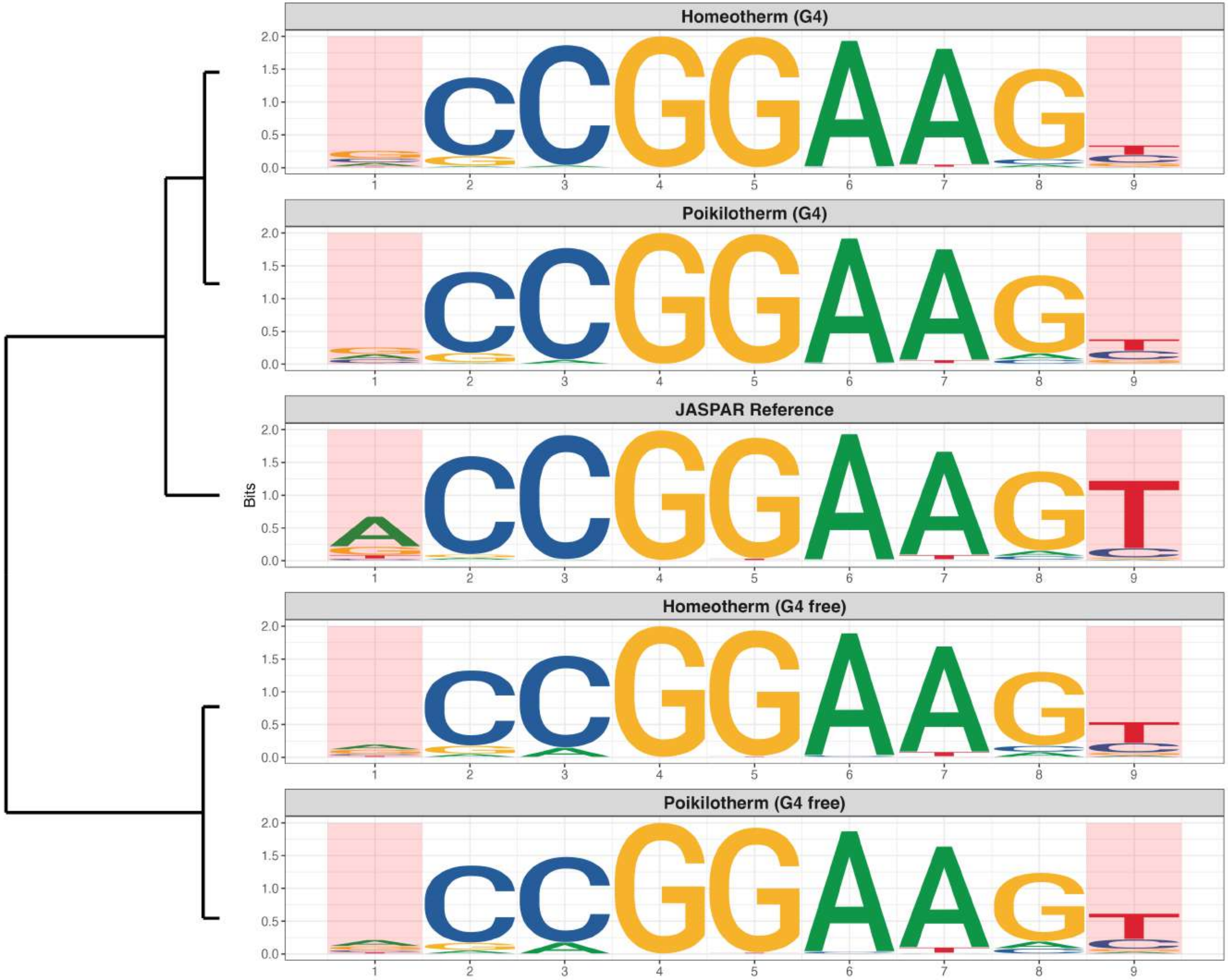

**Promoter : Highly Reduced in G4s**

*MA0073.2 - RREB1*

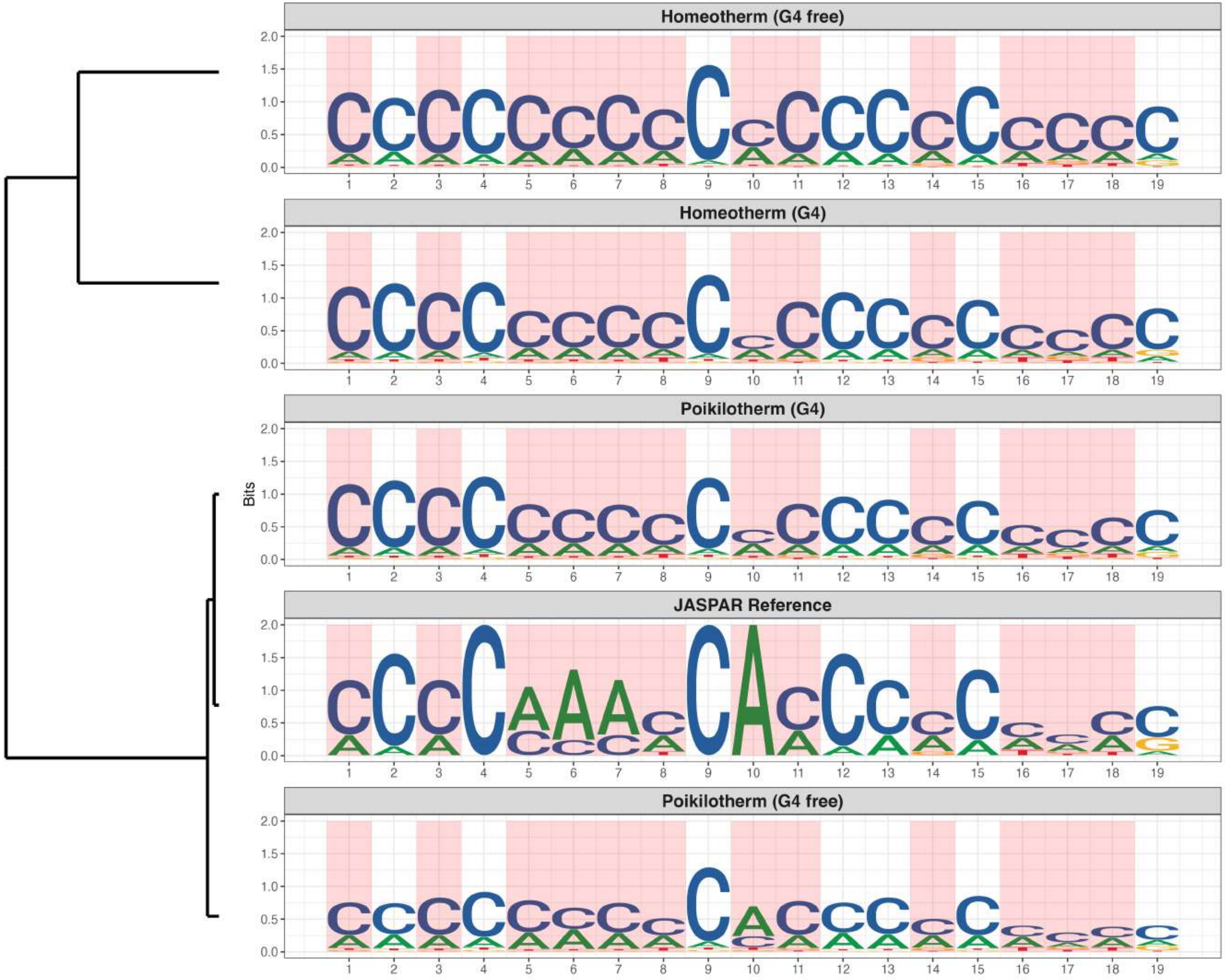

**Promoter : Highly Reduced in G4s**

*MA0080.7 - Spi1*

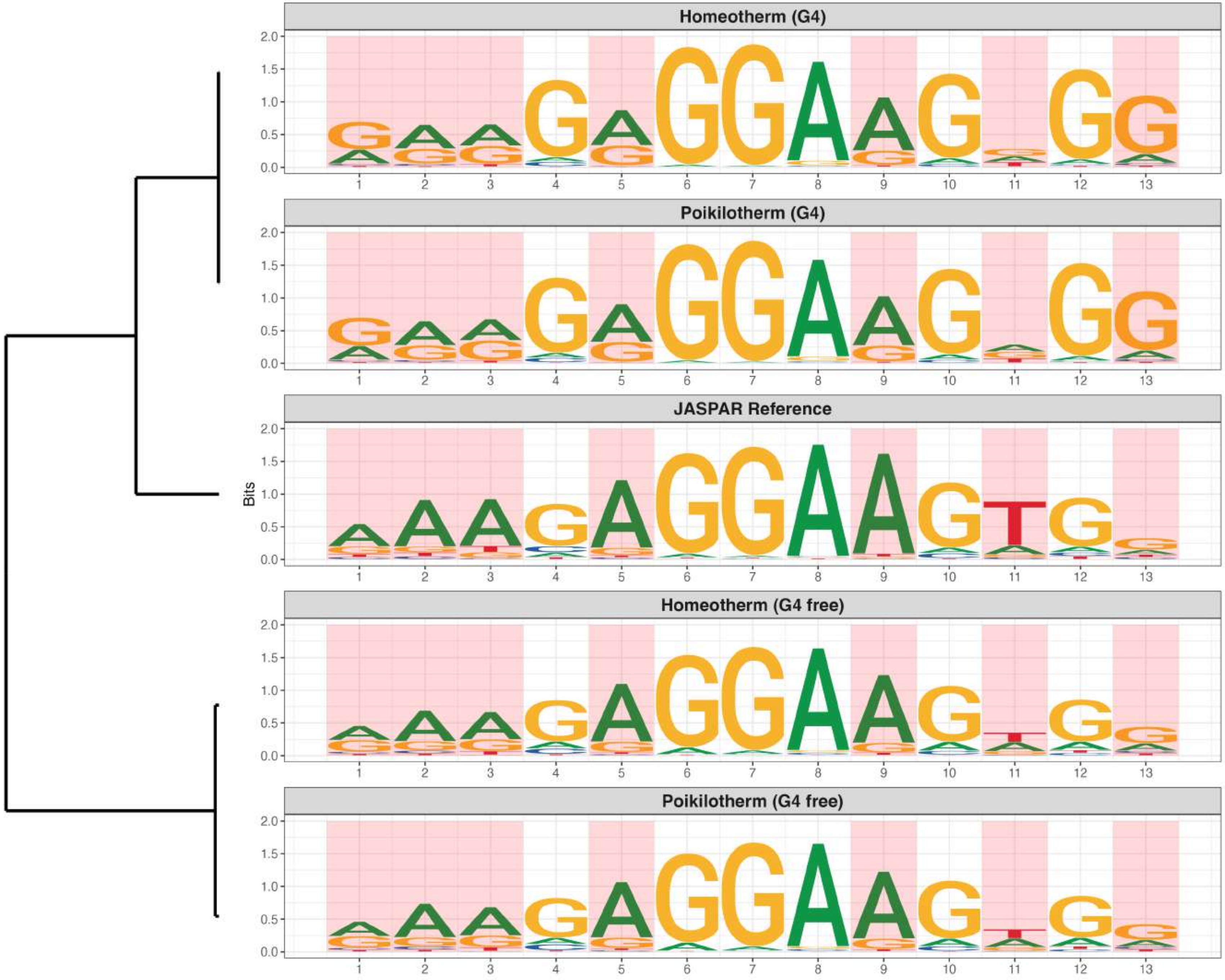

**Promoter : Highly Reduced in G4s**

*MA0081.3 - SPIB*

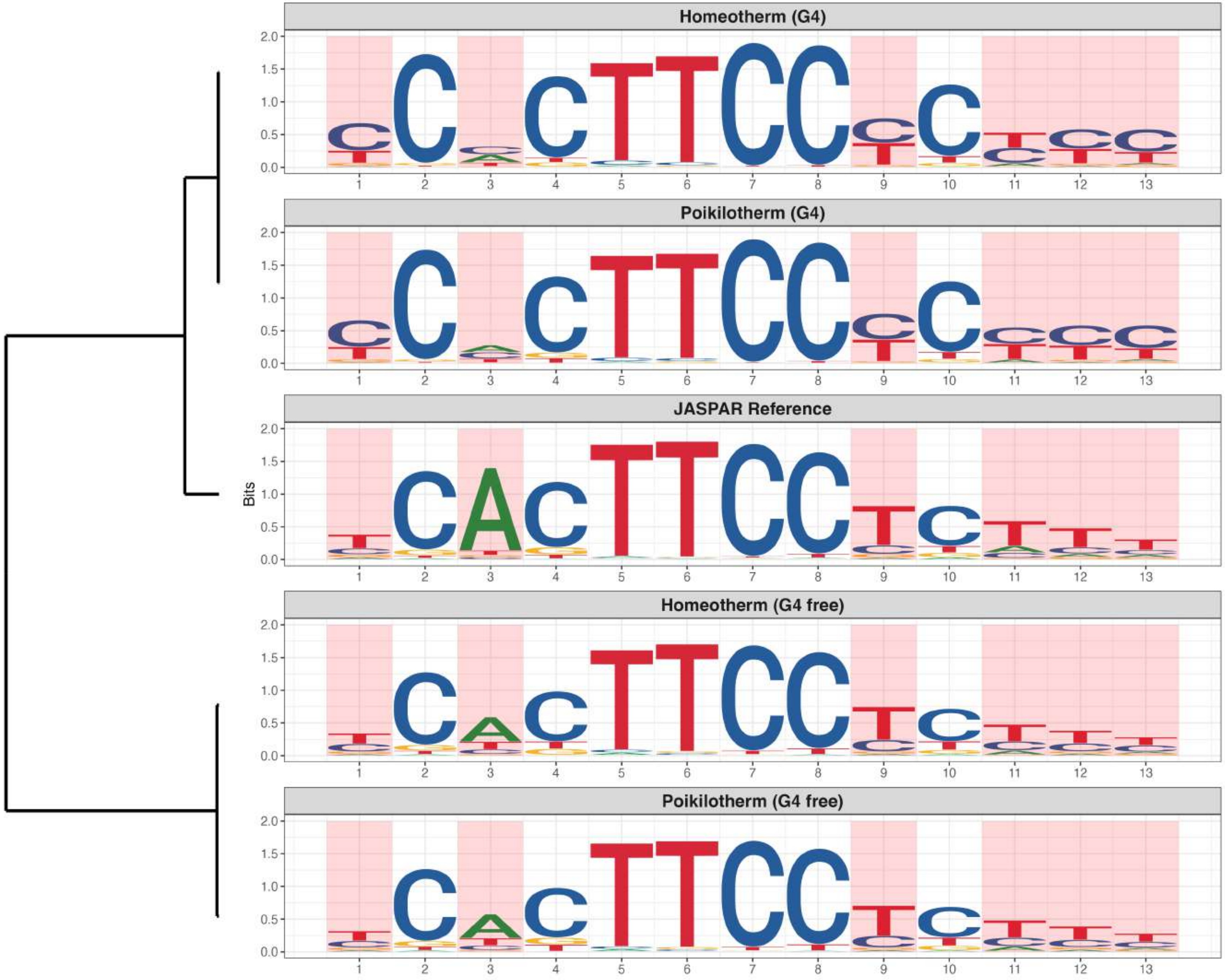

**Promoter : Highly Reduced in G4s**

*MA0111.1 - Spz1*

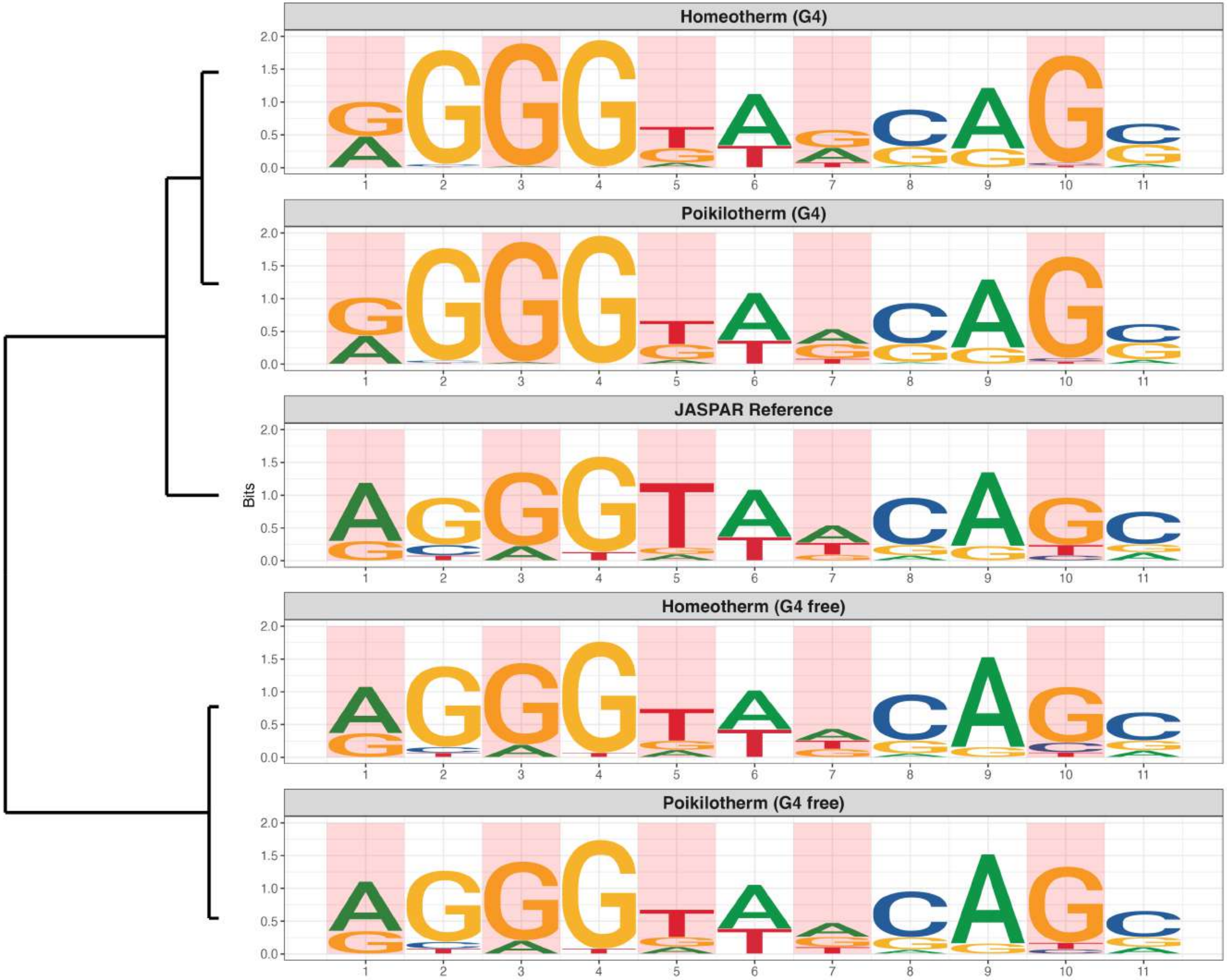

**Promoter : Highly Reduced in G4s**

*MA0163.1 - PLAG1*

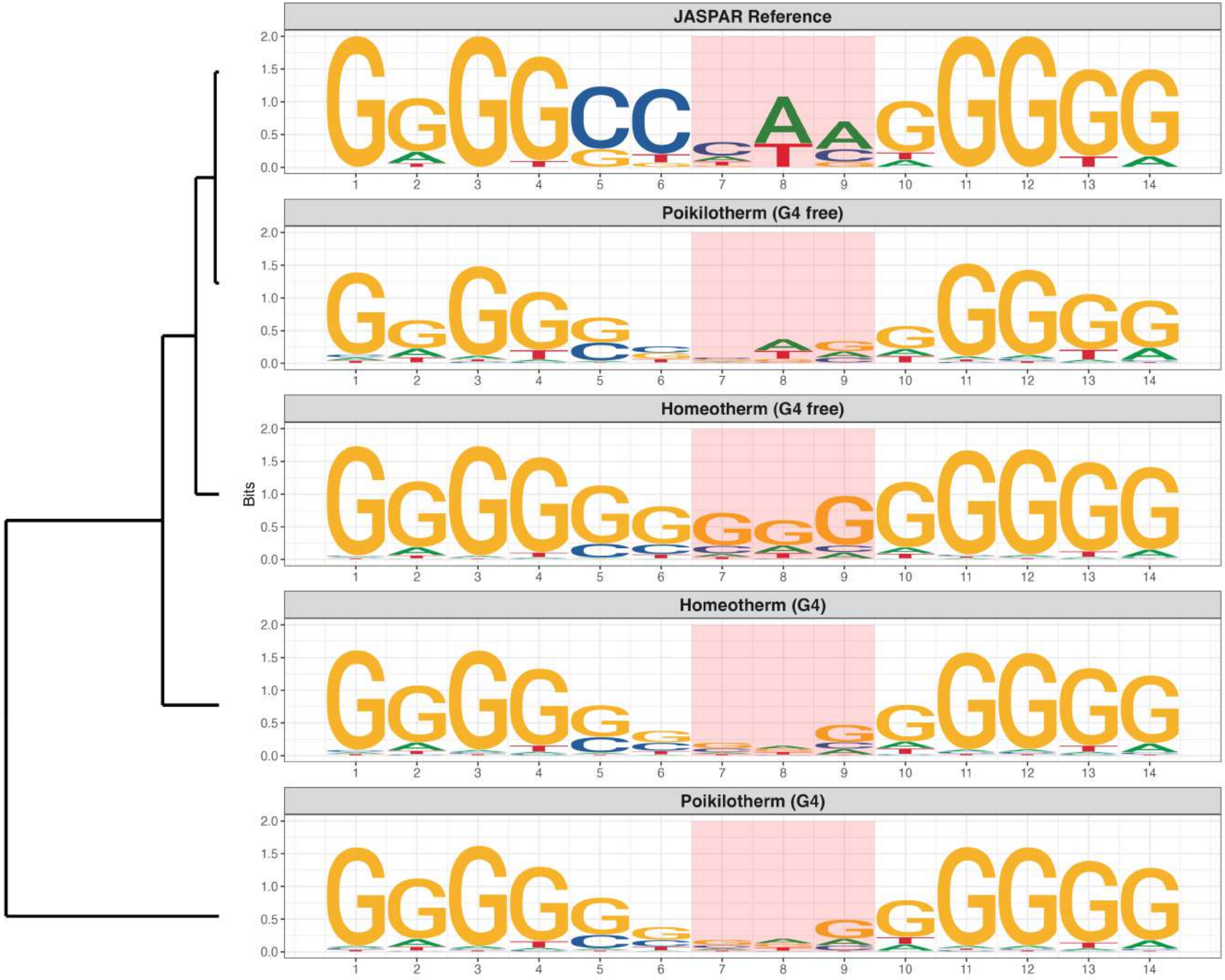

### Promoter : Highly Reduced in G4s

*MA0741.1 - KLF16*

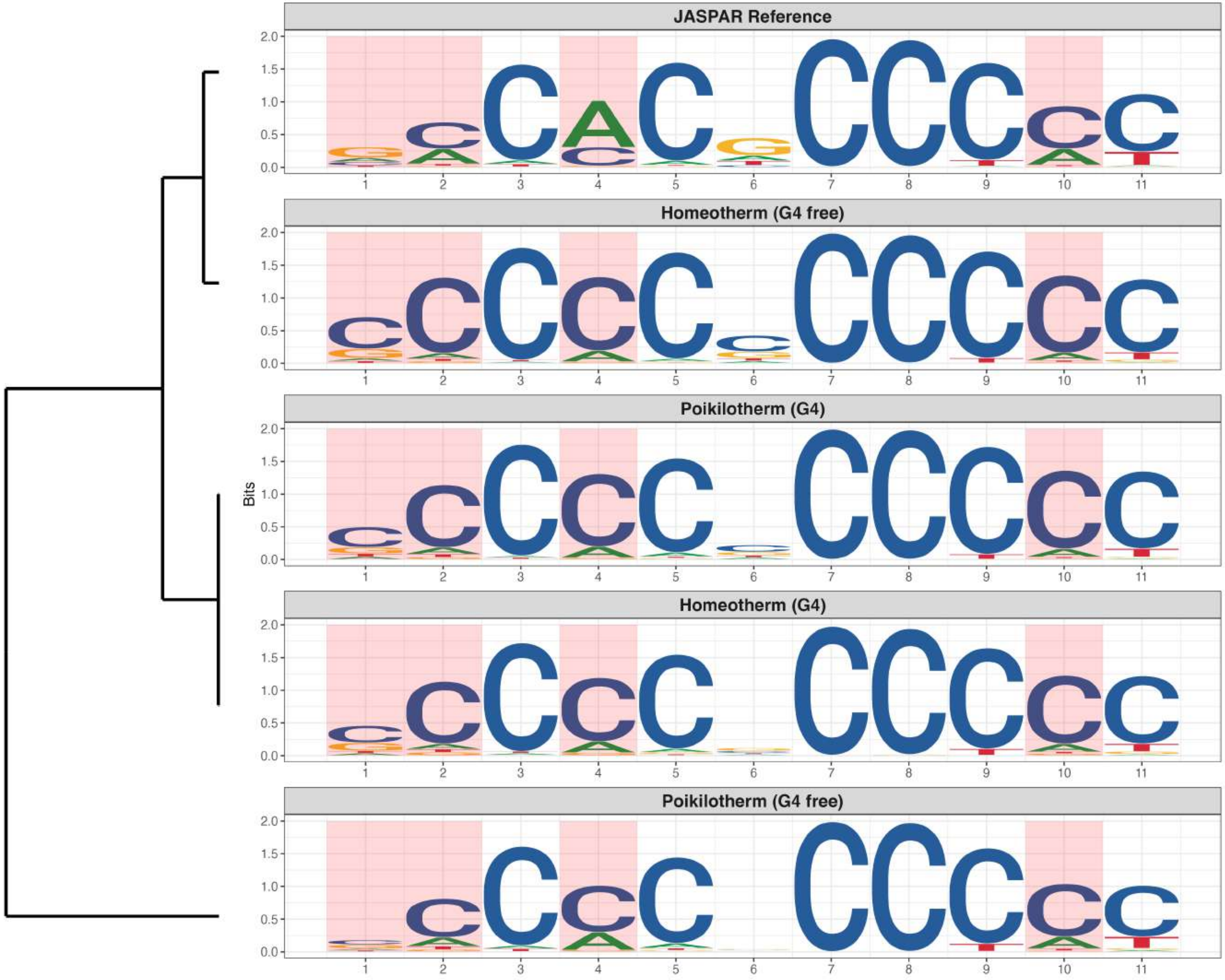

**Promoter : Highly Reduced in G4s**

*MA0747.2 - SP8*

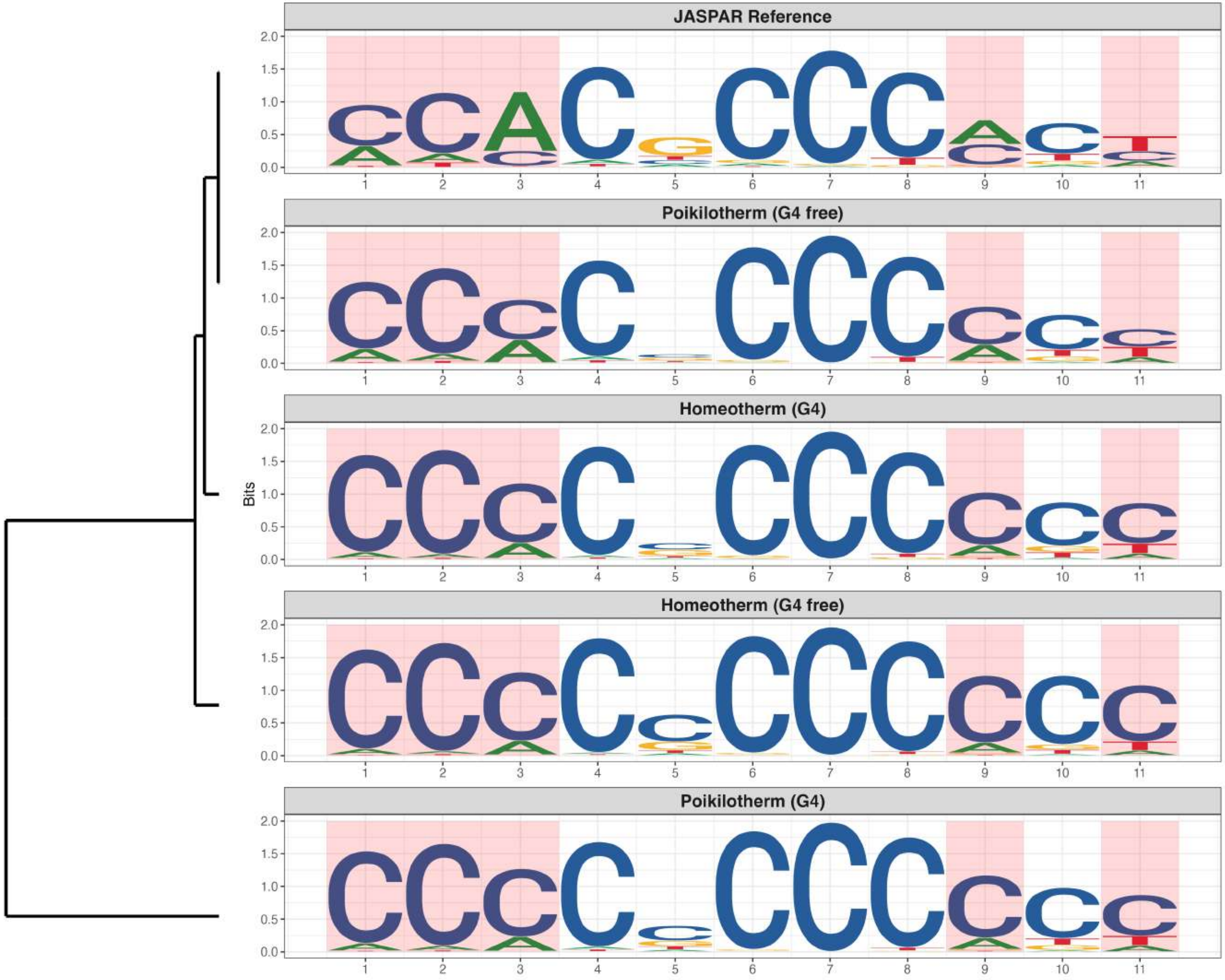

### Promoter : Highly Reduced in G4s

*MA0753.3 - ZNF740*

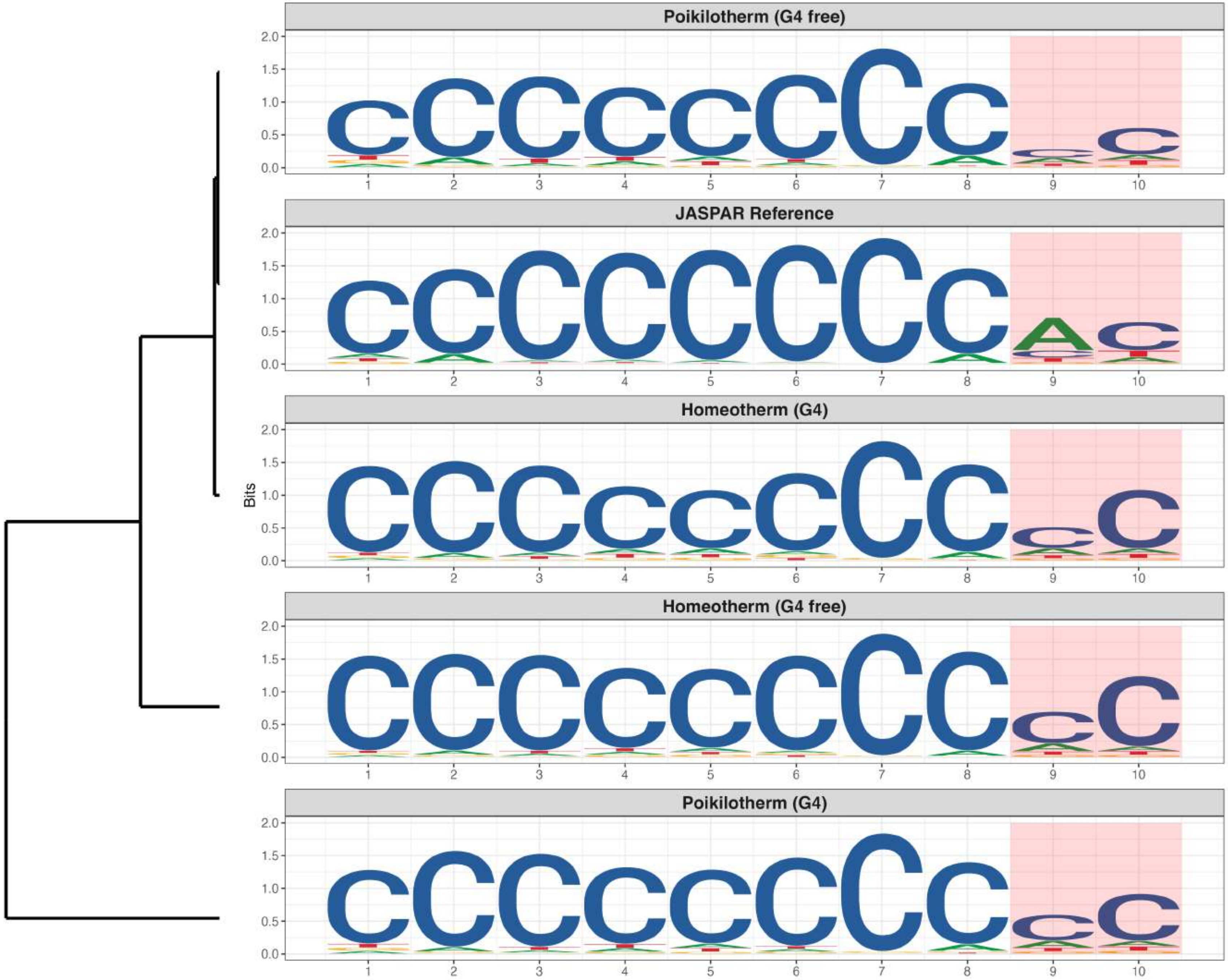

**Promoter : Highly Reduced in G4s**

*MA0764.4 - ETV4*

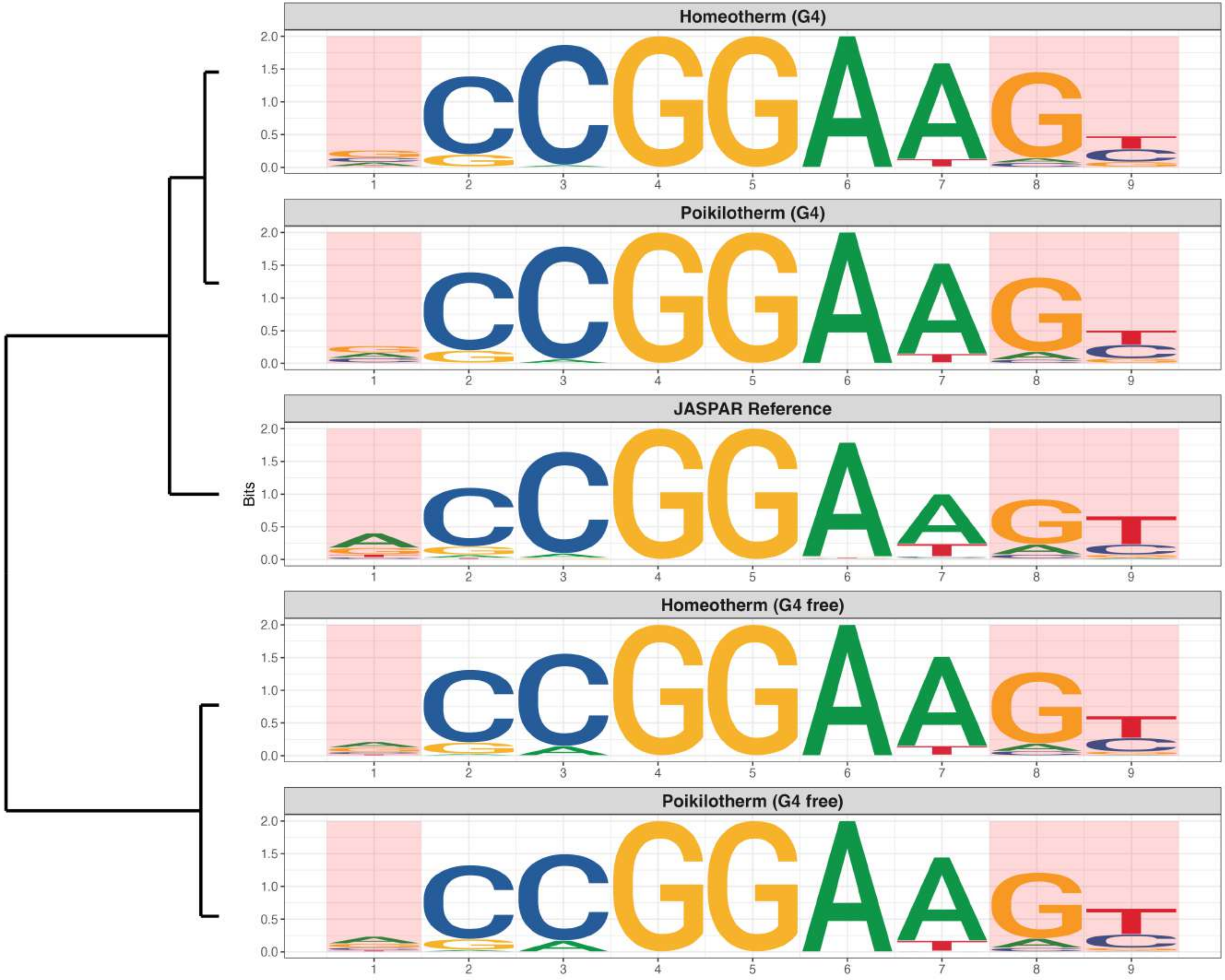

**Promoter : Highly Reduced in G4s**

*MA0765.4 - ETV5*

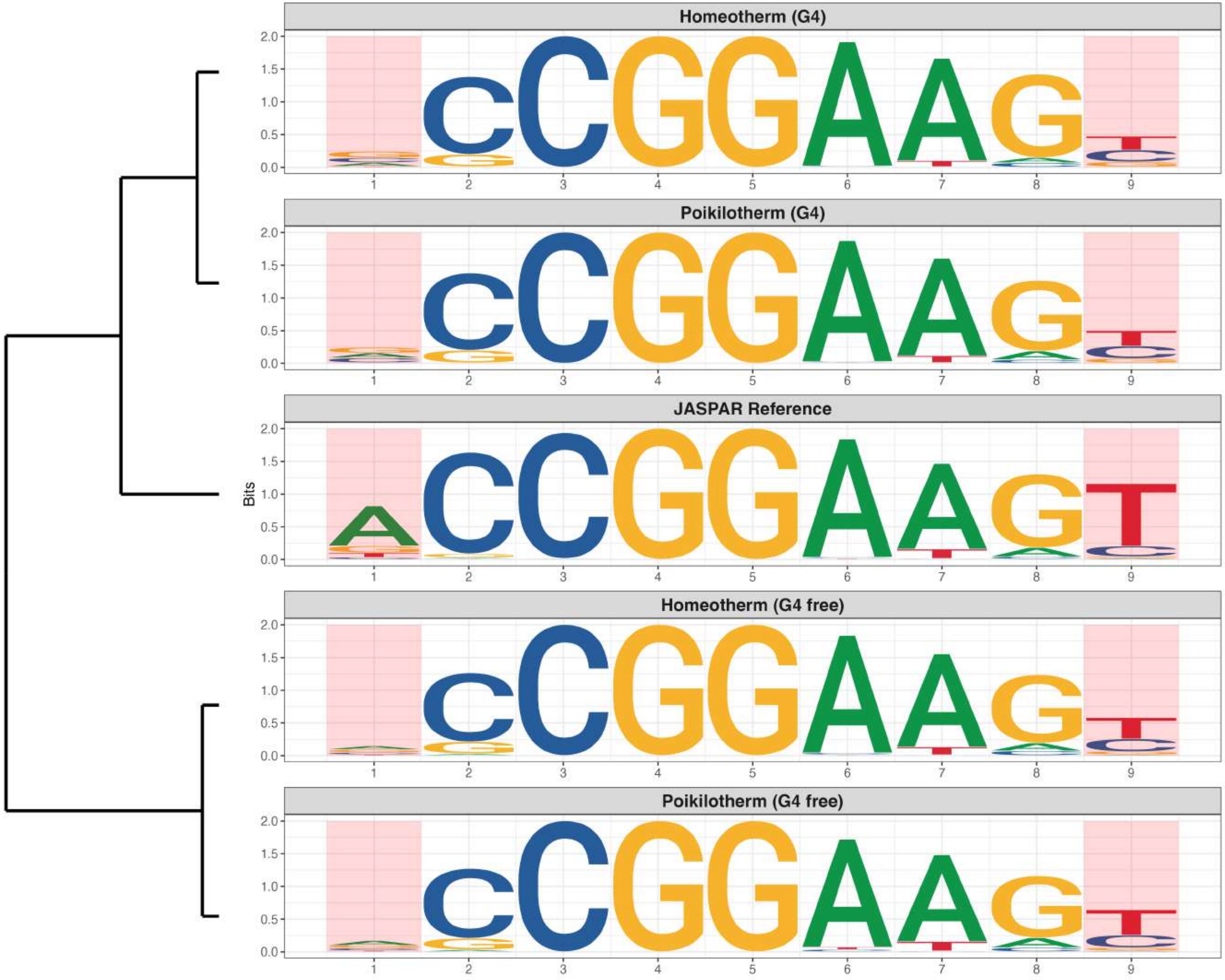

**Promoter : Highly Reduced in G4s**

*MA1117.2 - RELB*

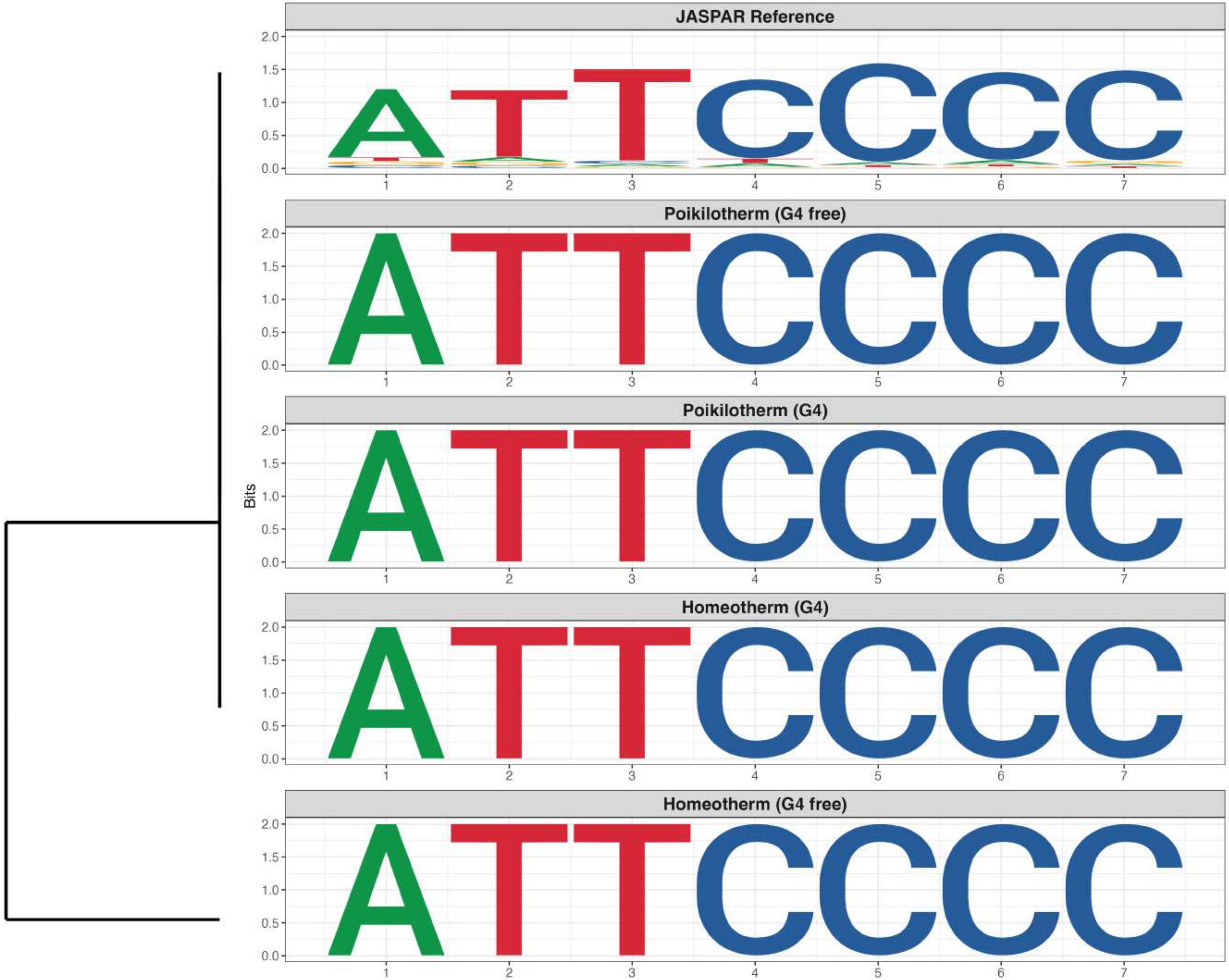

**Promoter : Highly Reduced in G4s**

*MA1532.2 - NR1D2*

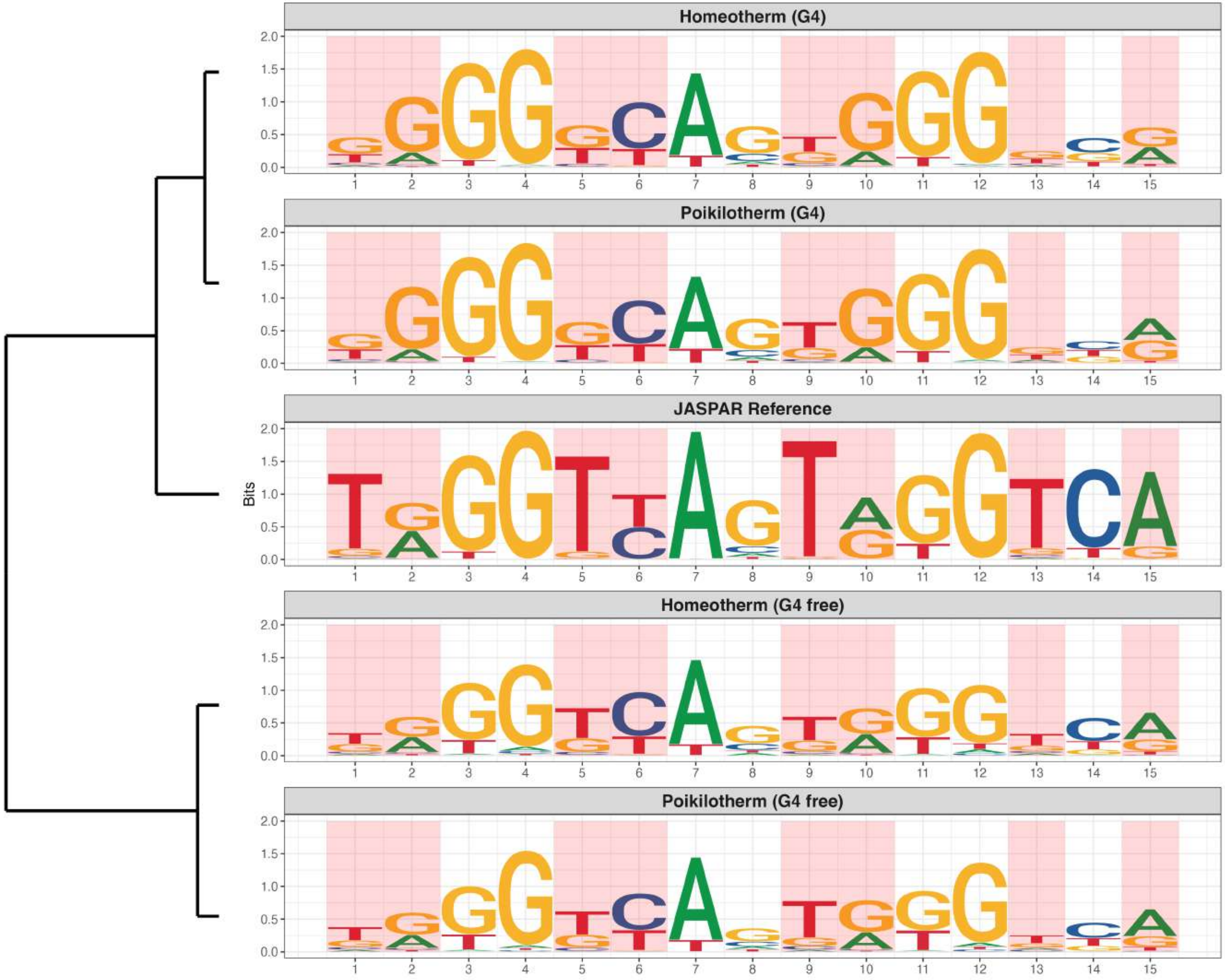

### Promoter : Highly Reduced in G4s

*MA1564.2 - SP9*

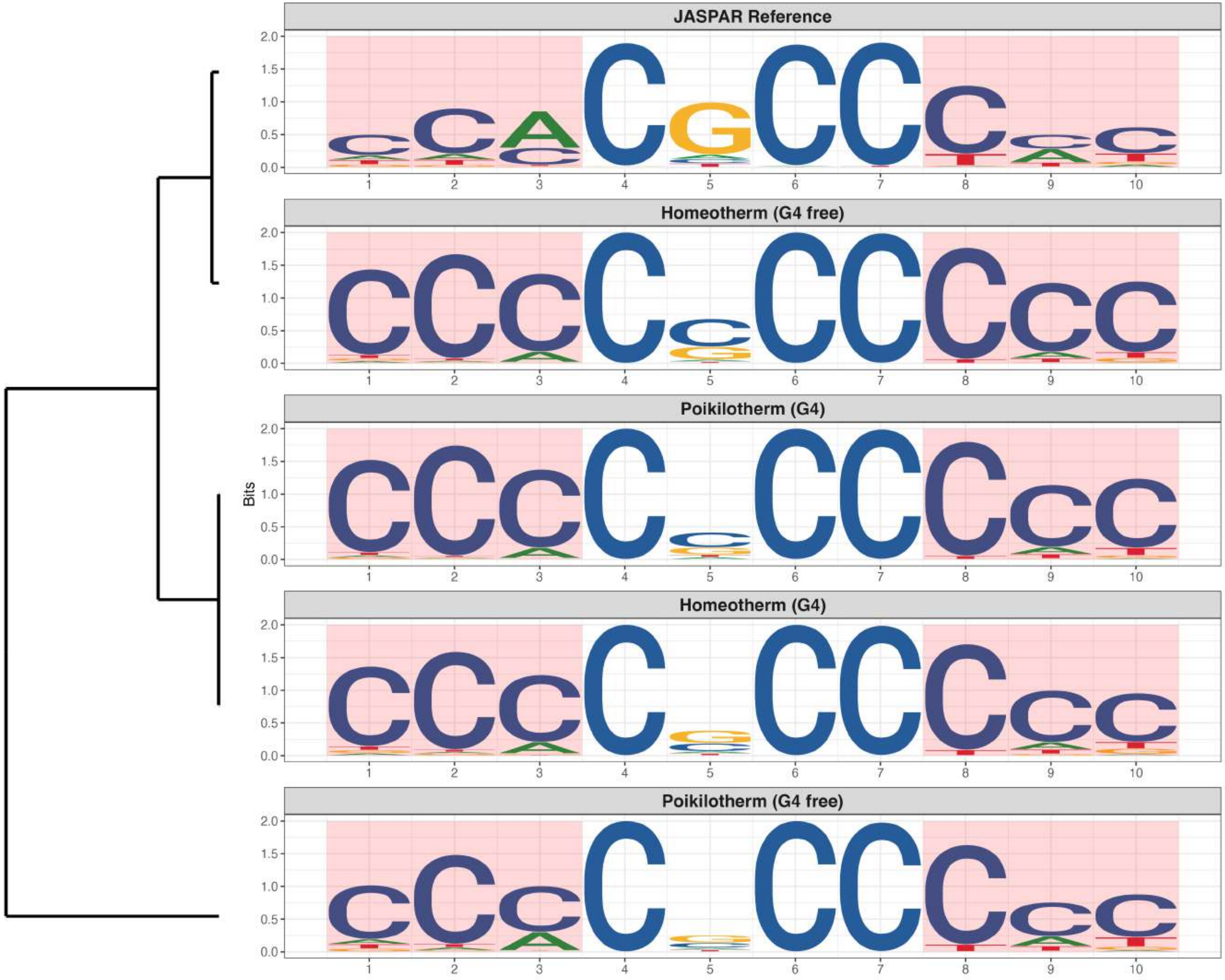

### Promoter : Highly Reduced in G4s

*MA1714.2 - ZNF675*

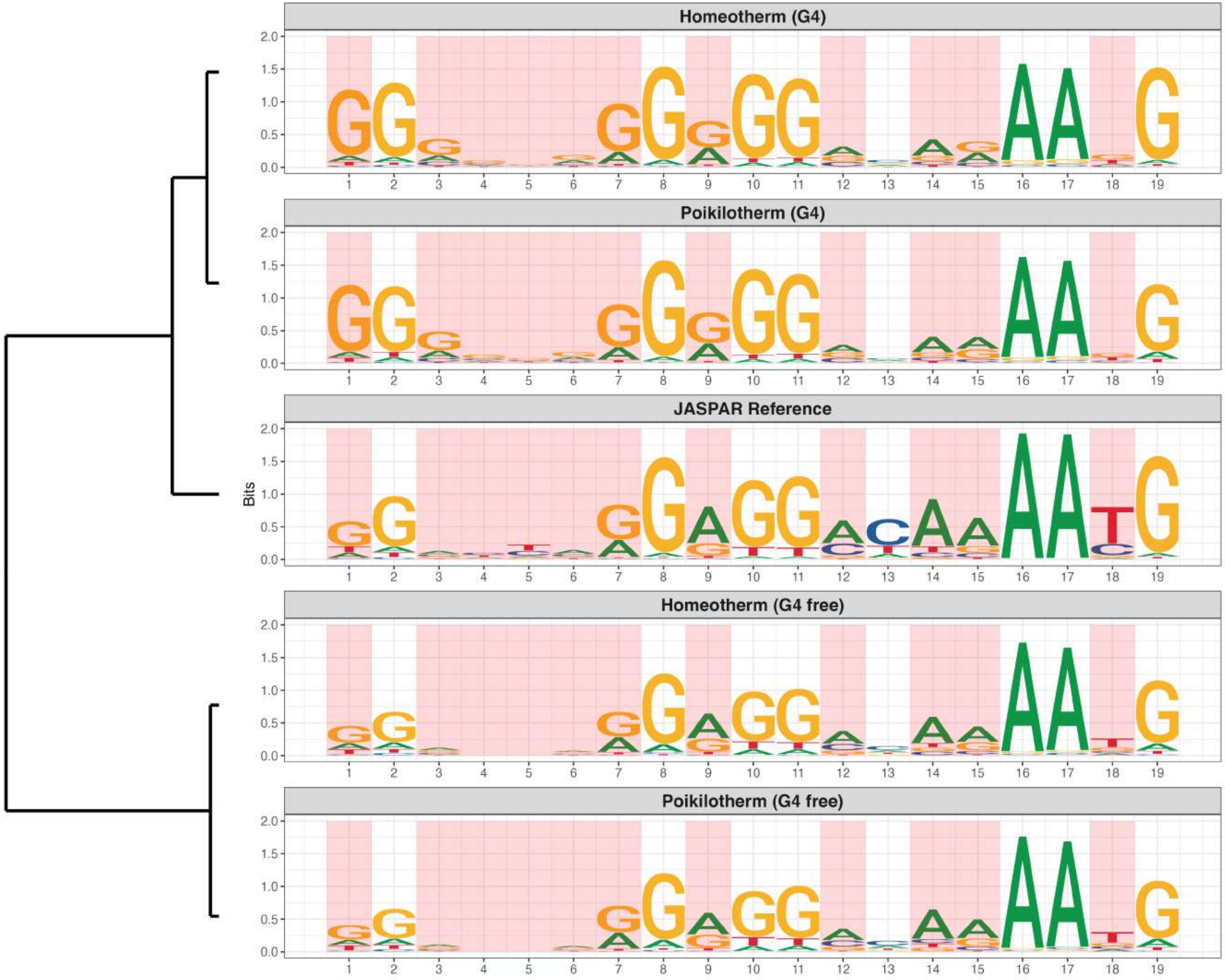

**Promoter : Highly Reduced in G4s**

*MA1987.2 - ZNF701*

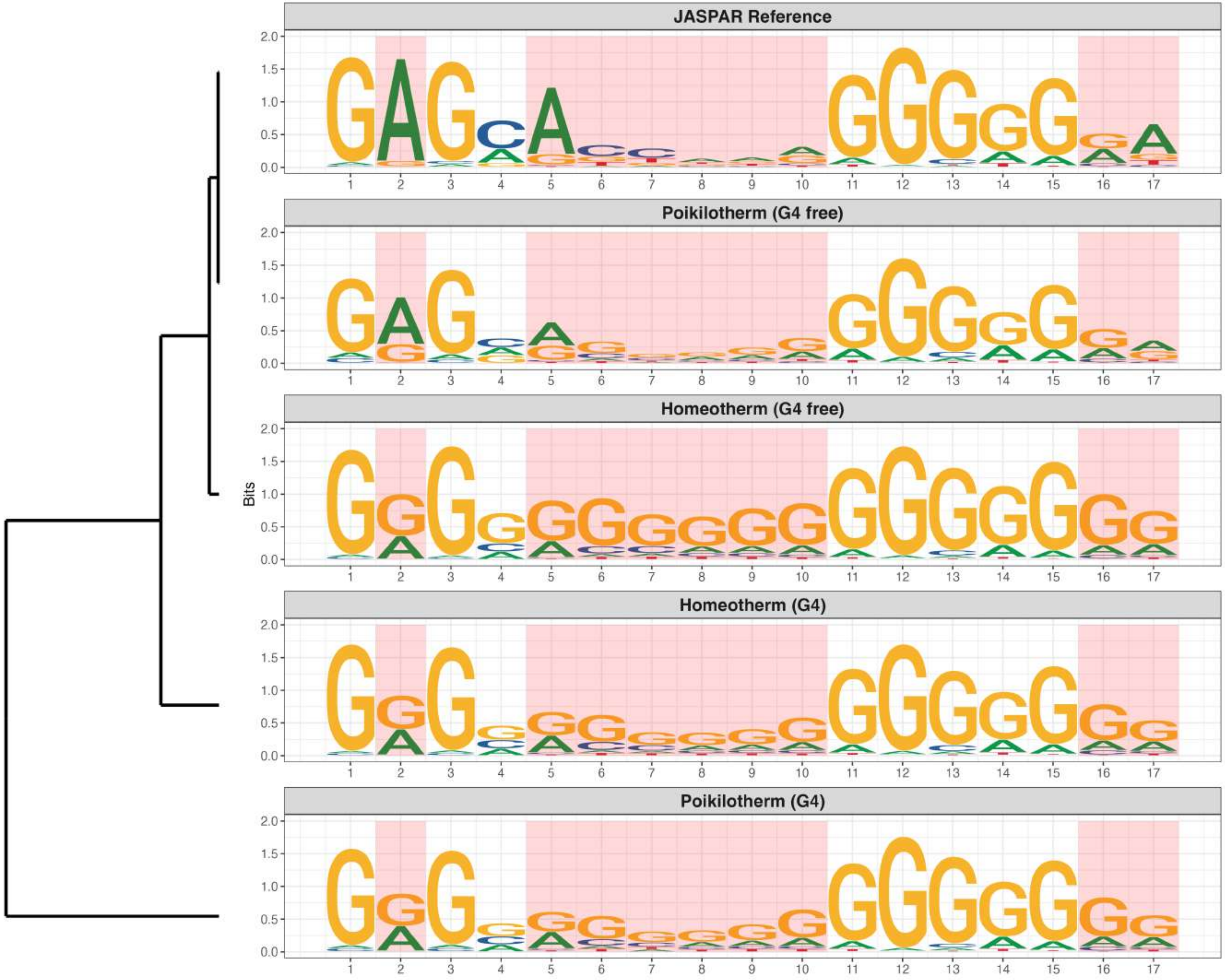

### Promoter : Highly Reduced in G4s

*MA2519.1 - ZNF606*

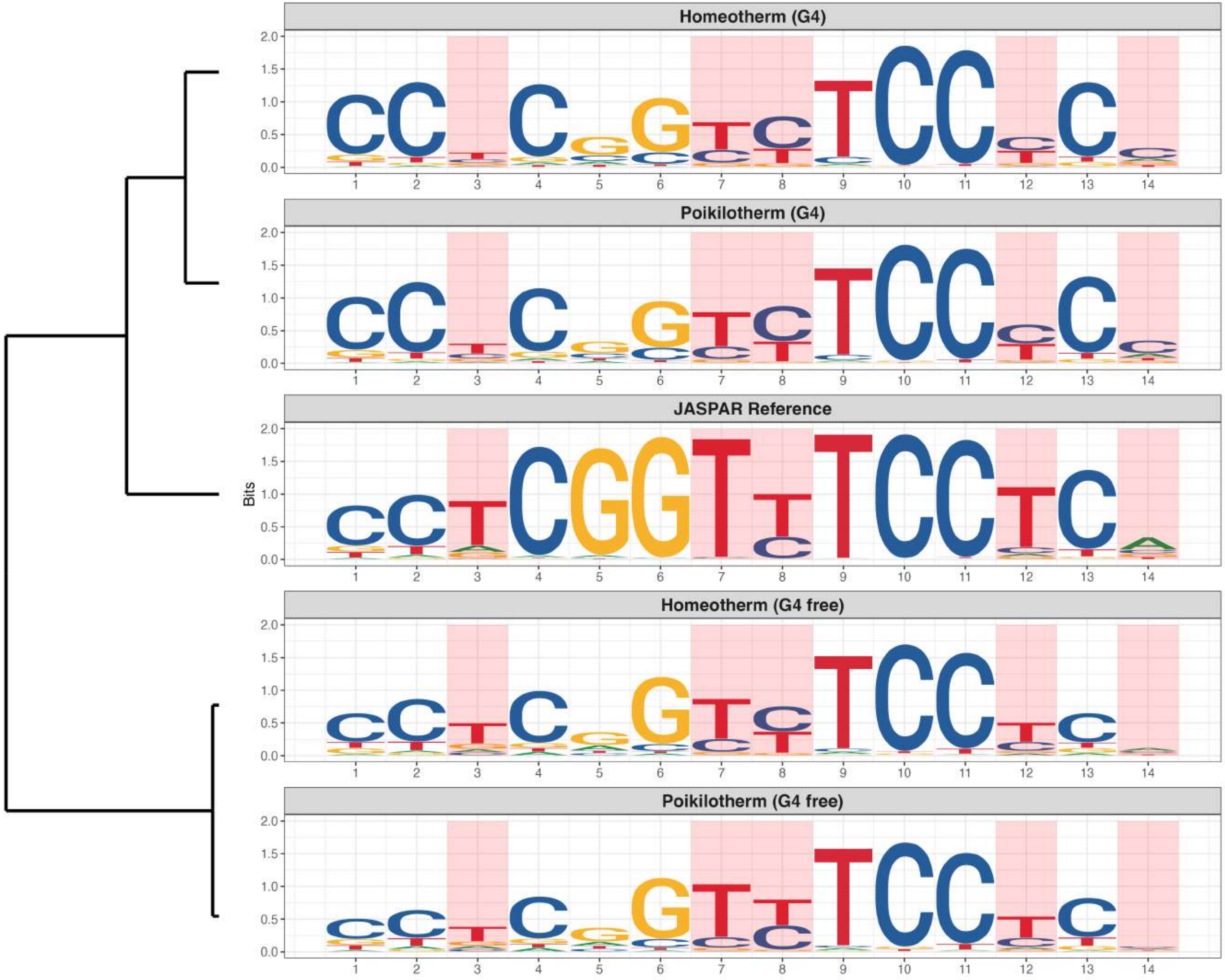

### Promoter : Highly Reduced in G4s

*MA2688.1 - ZNF347*

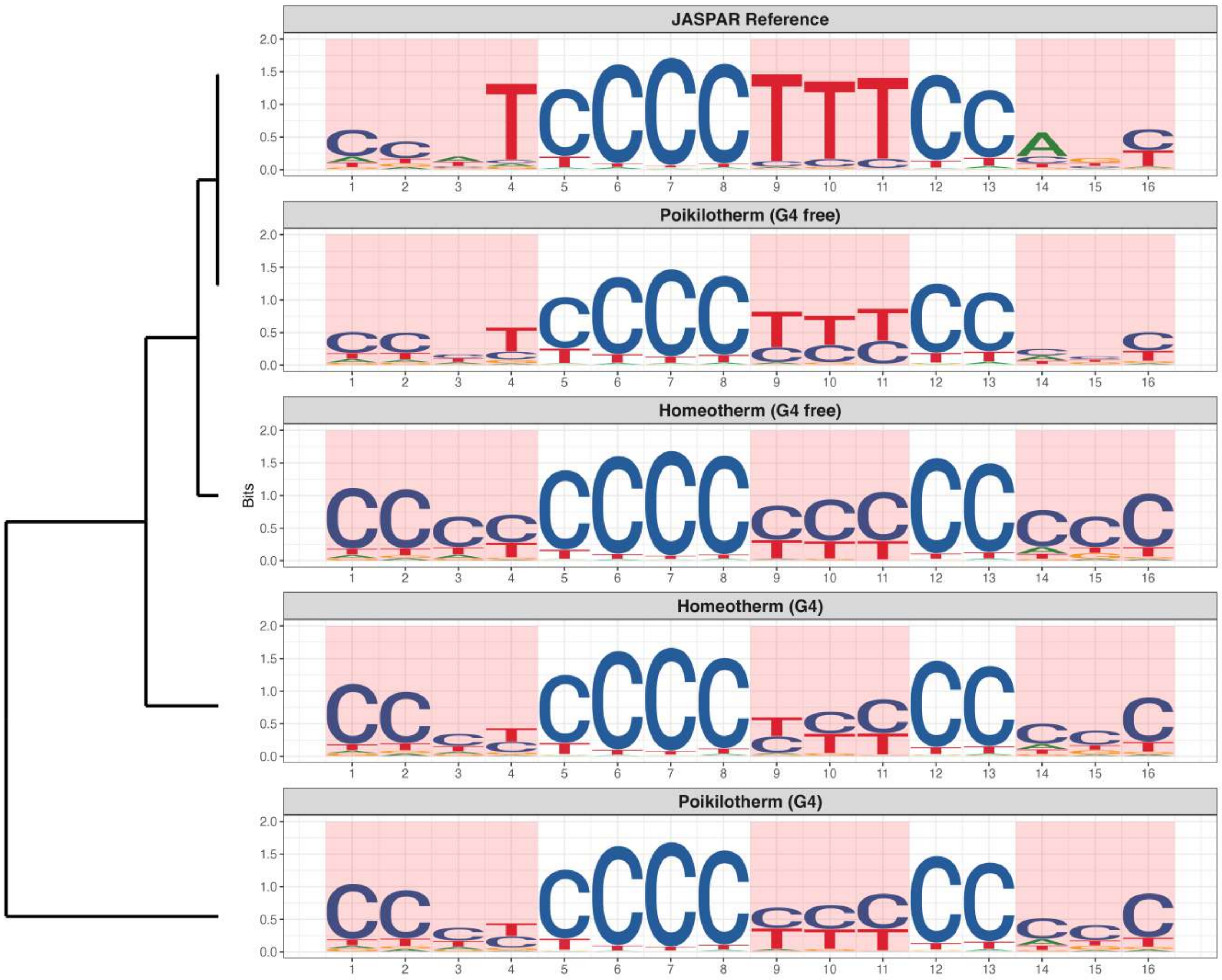

**Promoter : Moderately Reduced in G4s**

*MA0079.5 - SP1*

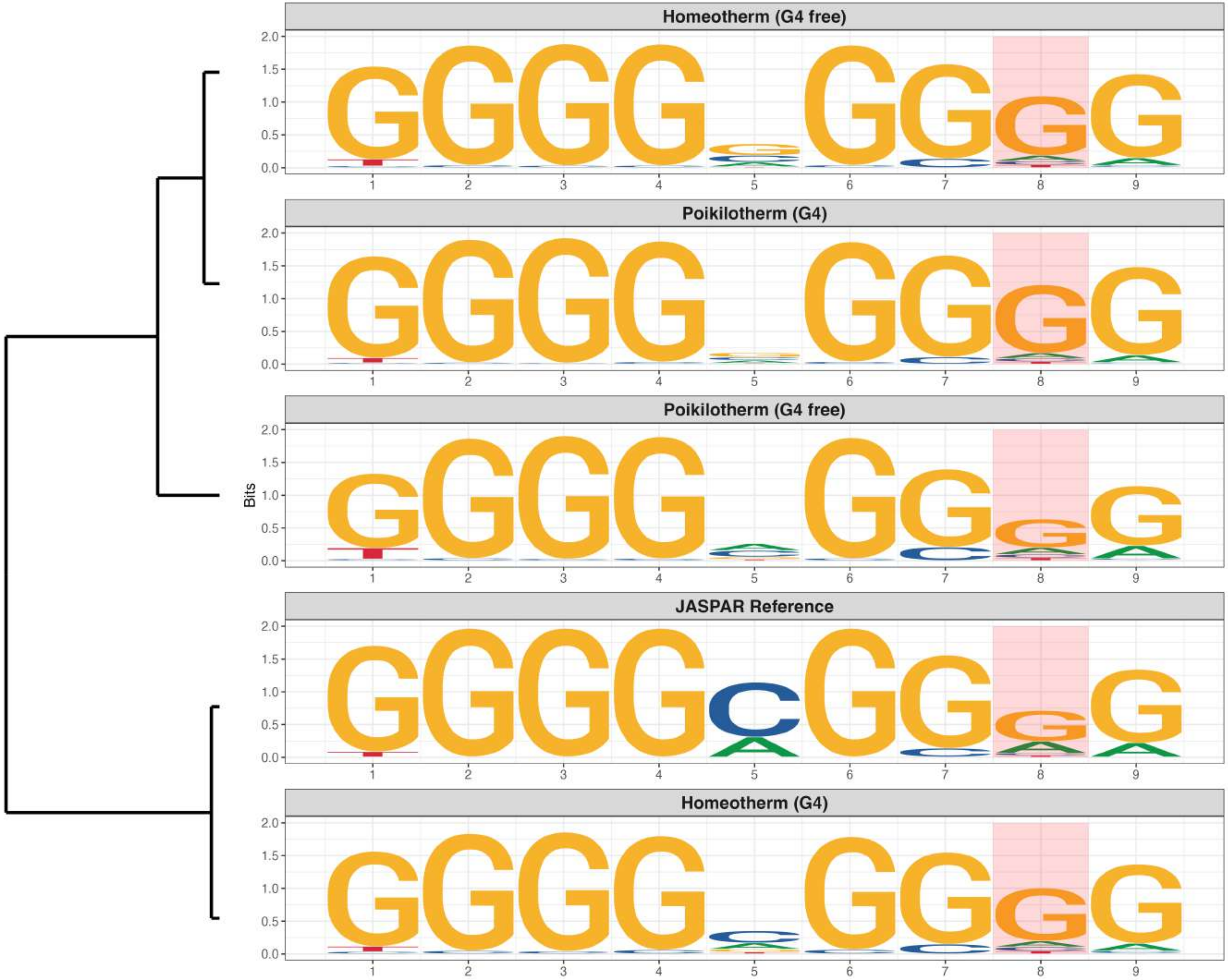

**Promoter : Moderately Reduced in G4s**

*MA0469.4 - E2F3*

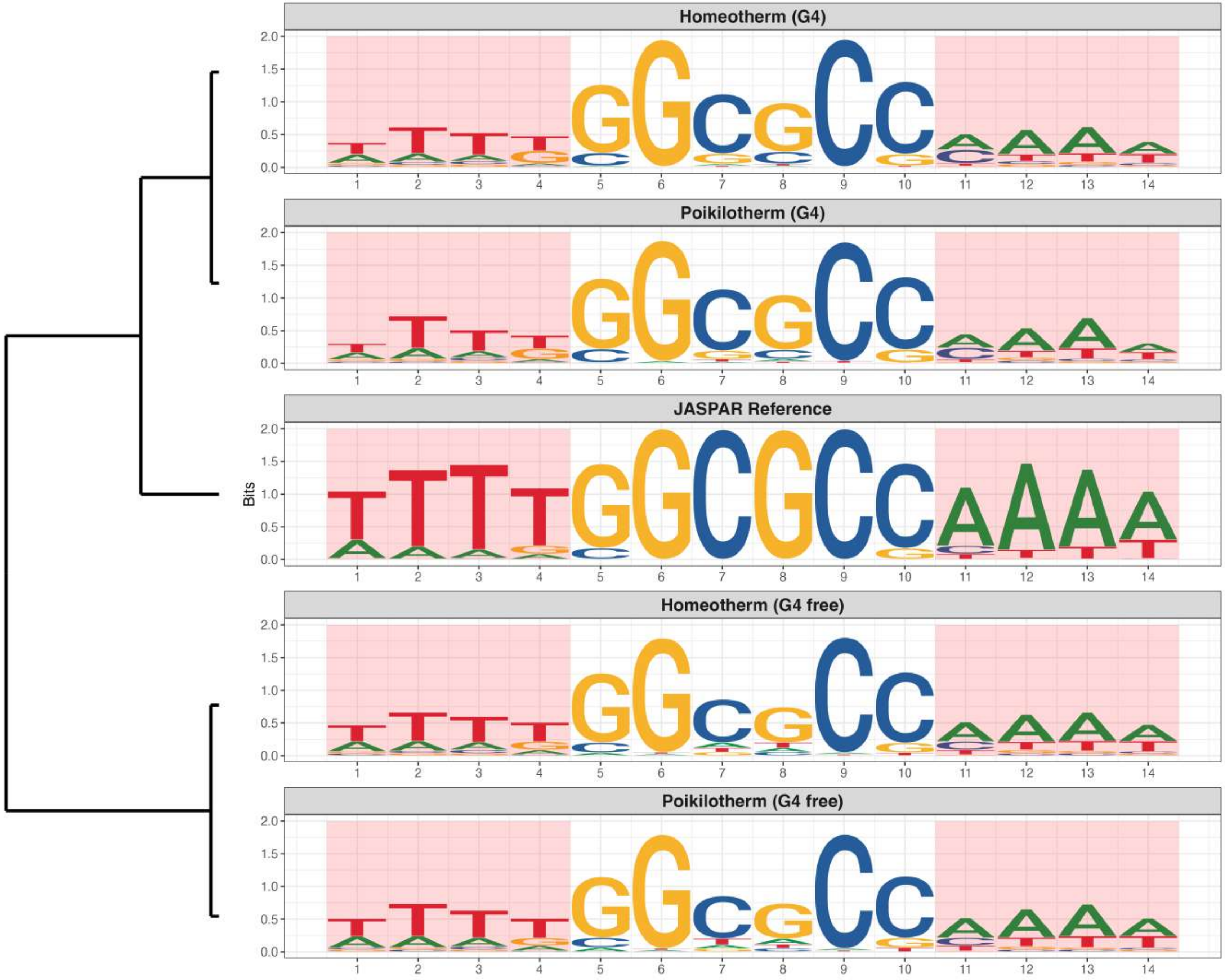

**Promoter : Moderately Reduced in G4s**

*MA0526.5 - USF2*

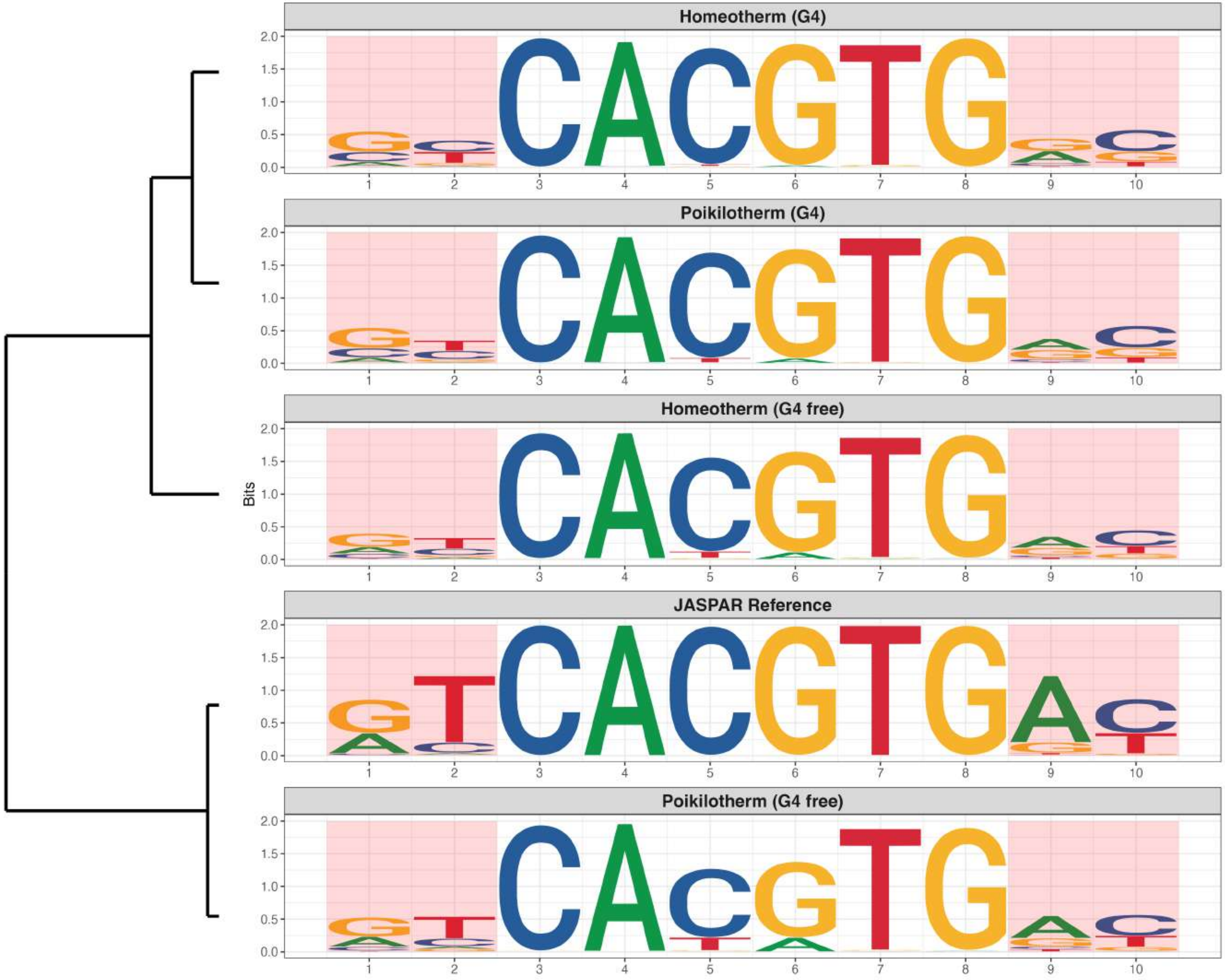

**Promoter : Moderately Reduced in G4s**

*MA0620.4 - MITF*

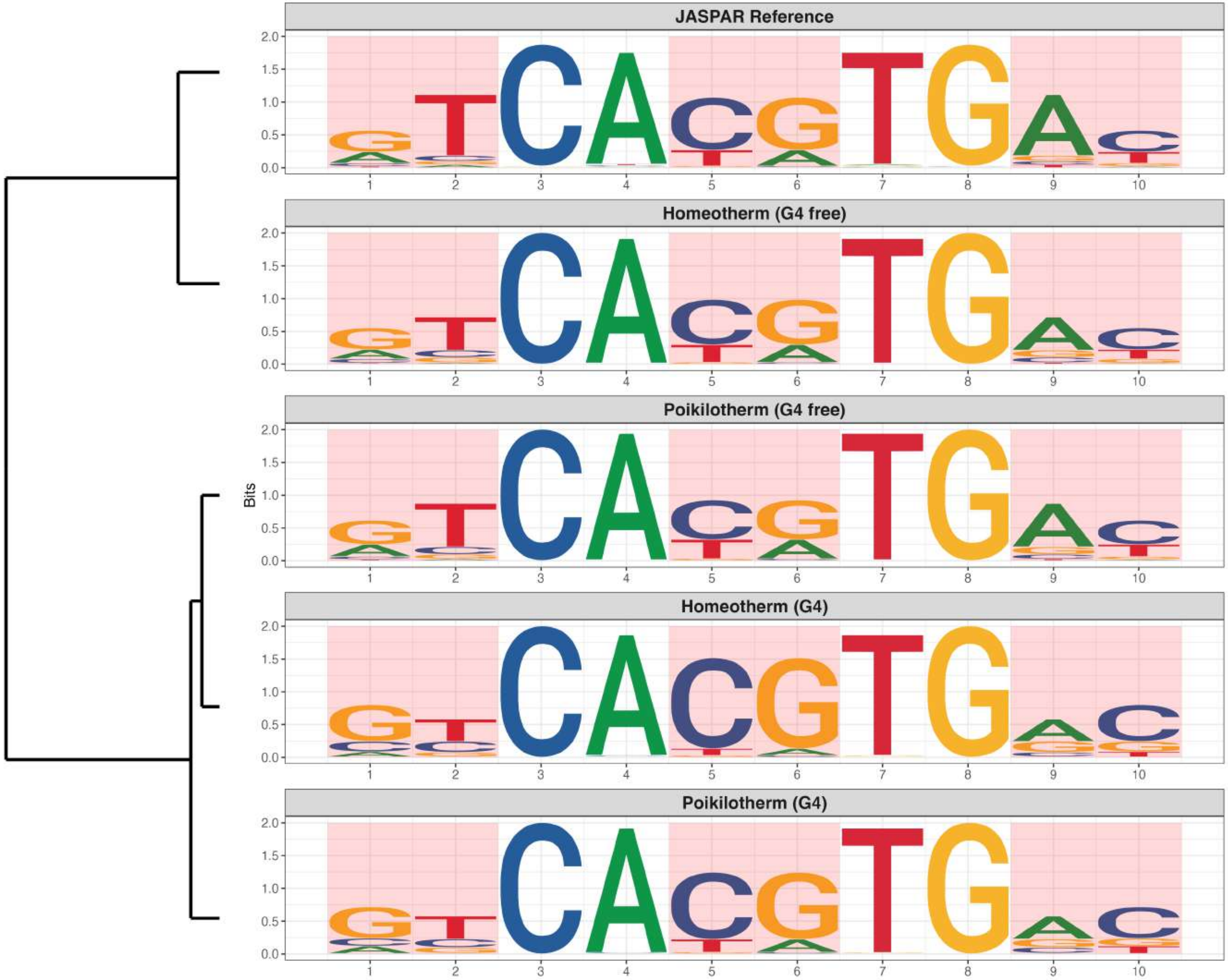

### Promoter : Moderately Reduced in G4s

*MA0685.2 - SP4*

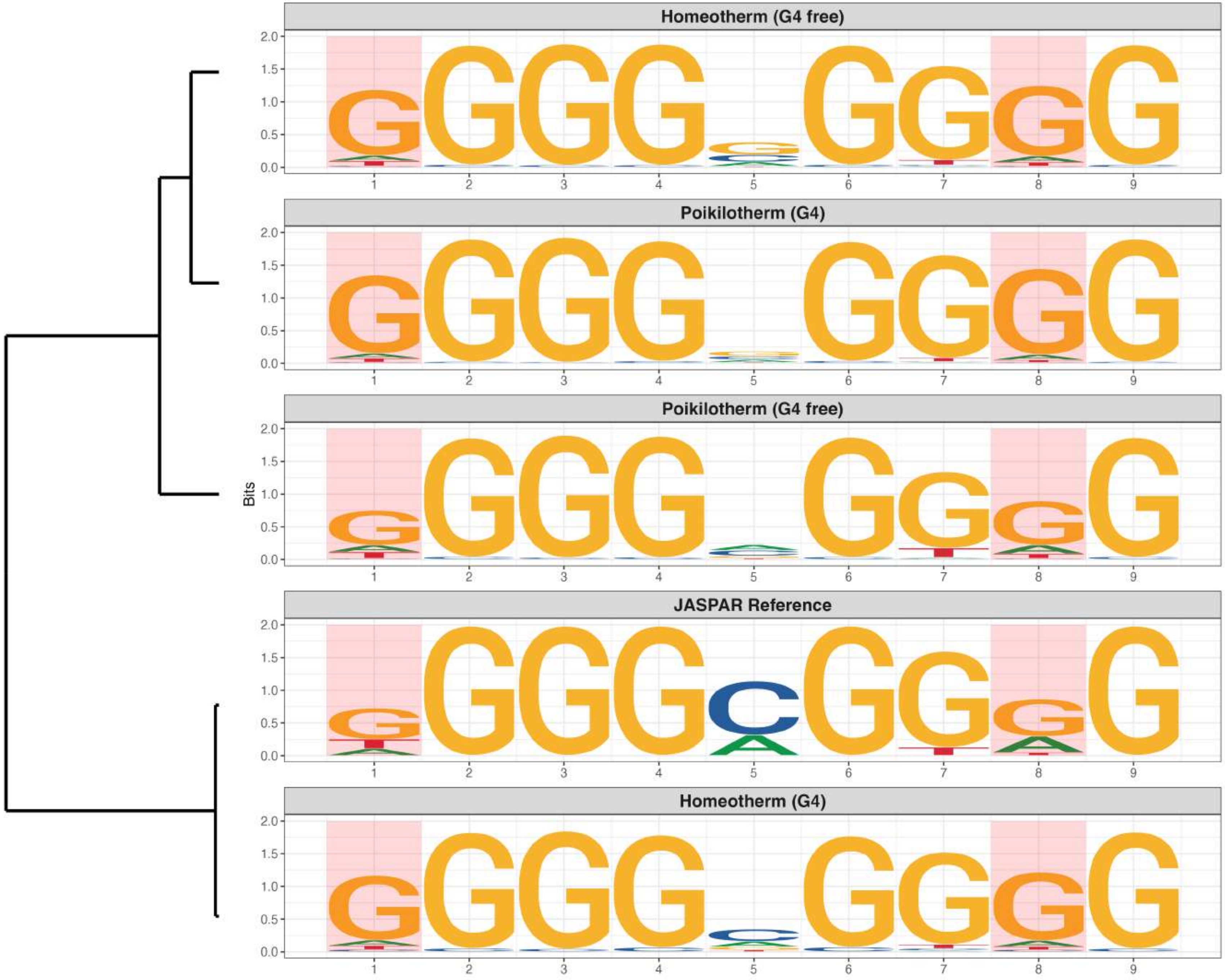

### Promoter : Moderately Reduced in G4s

*MA0736.1 - GLIS2*

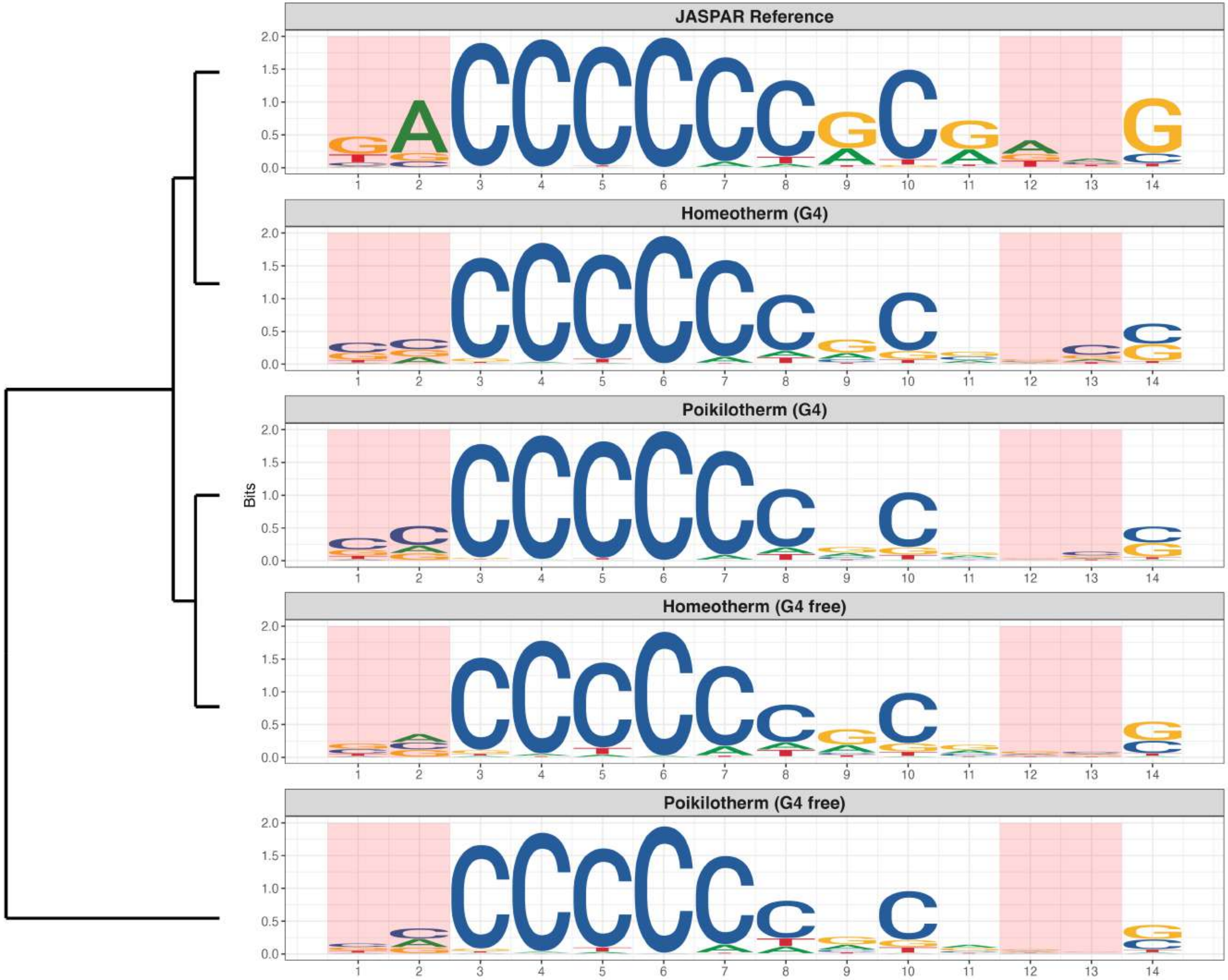

### Promoter : Moderately Reduced in G4s

*MA0781.2 - PAX9*

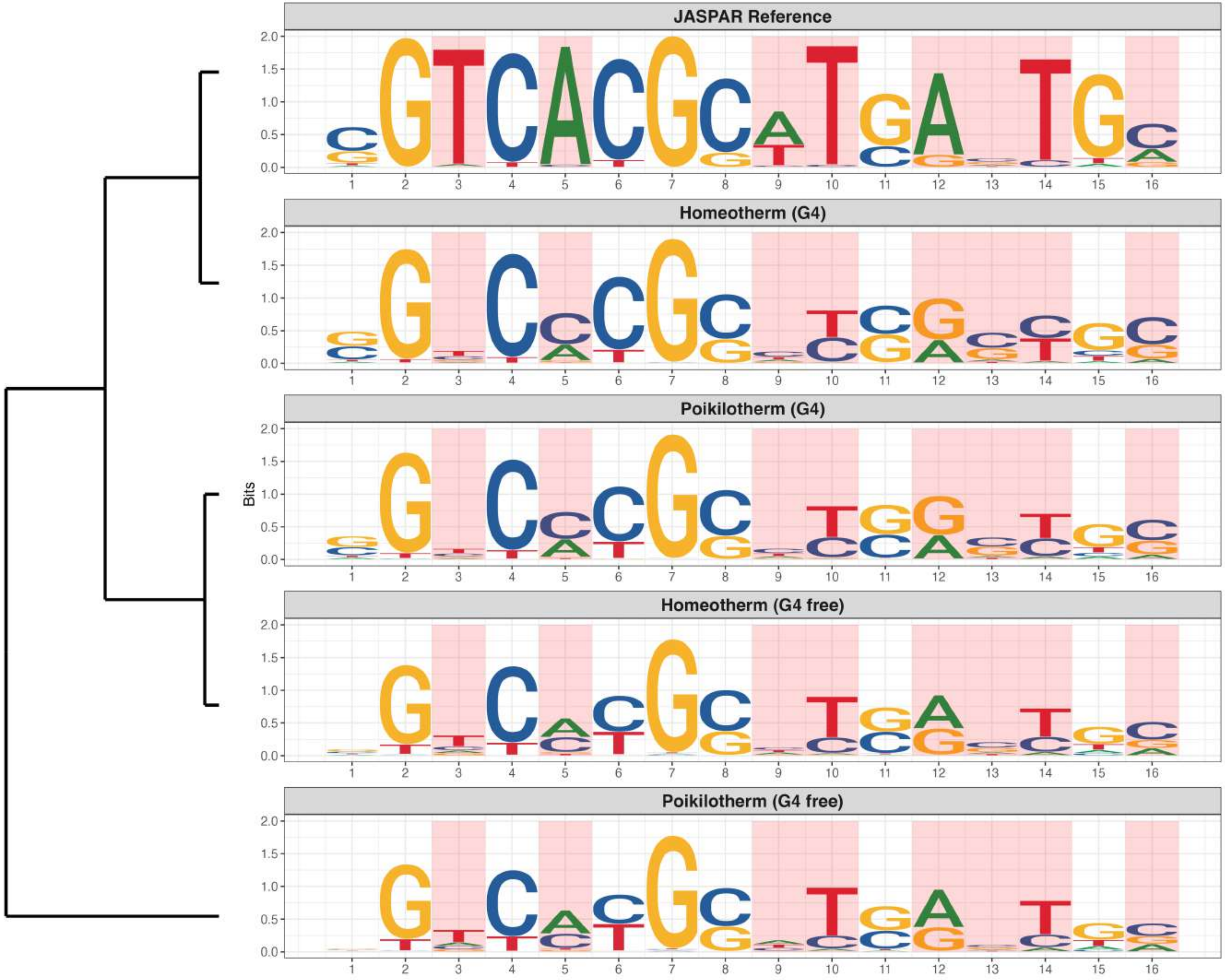

**Promoter : Moderately Reduced in G4s**

*MA0831.3 - TFE3*

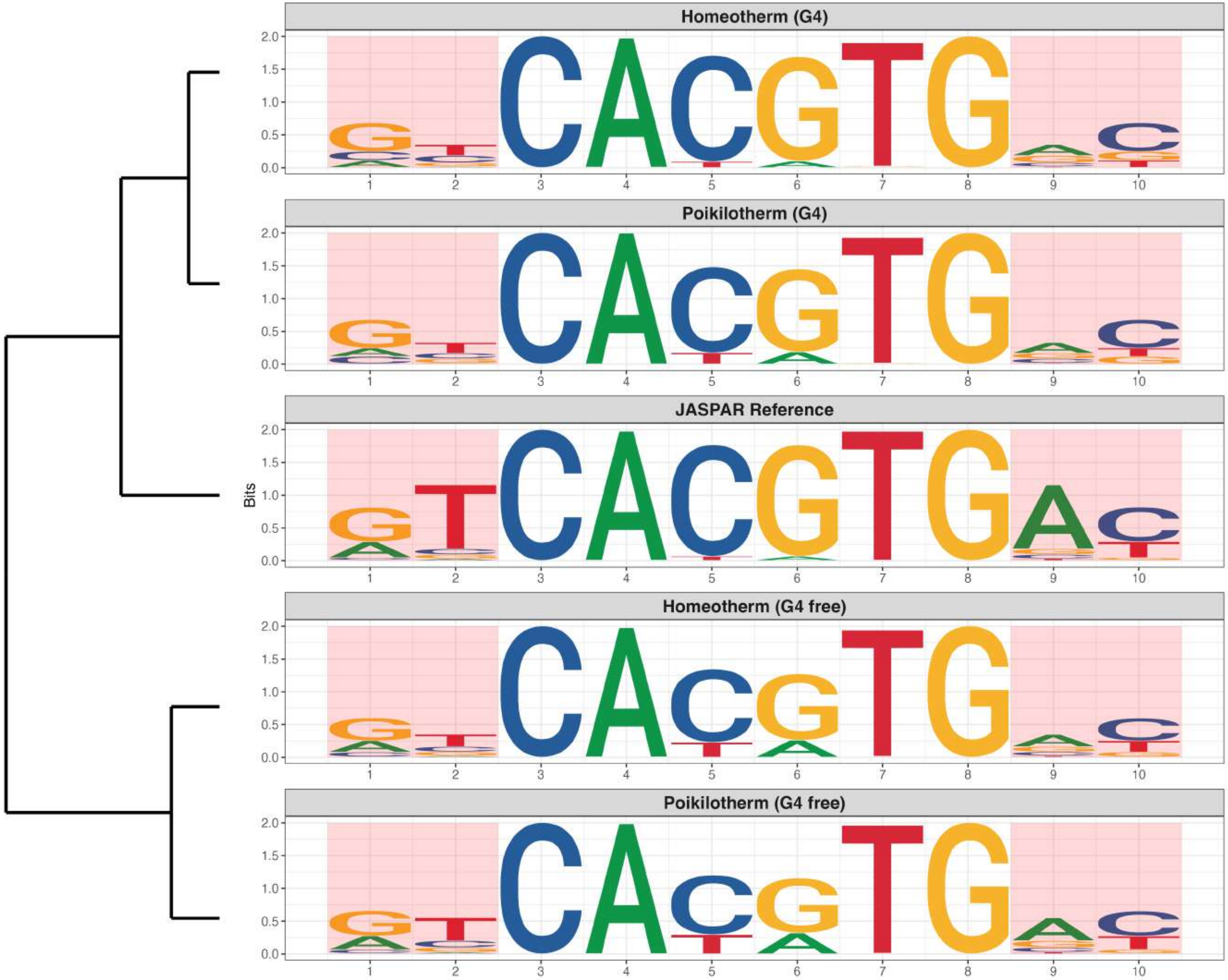

**Promoter : Moderately Reduced in G4s**

*MA0837.3 - CEBPE*

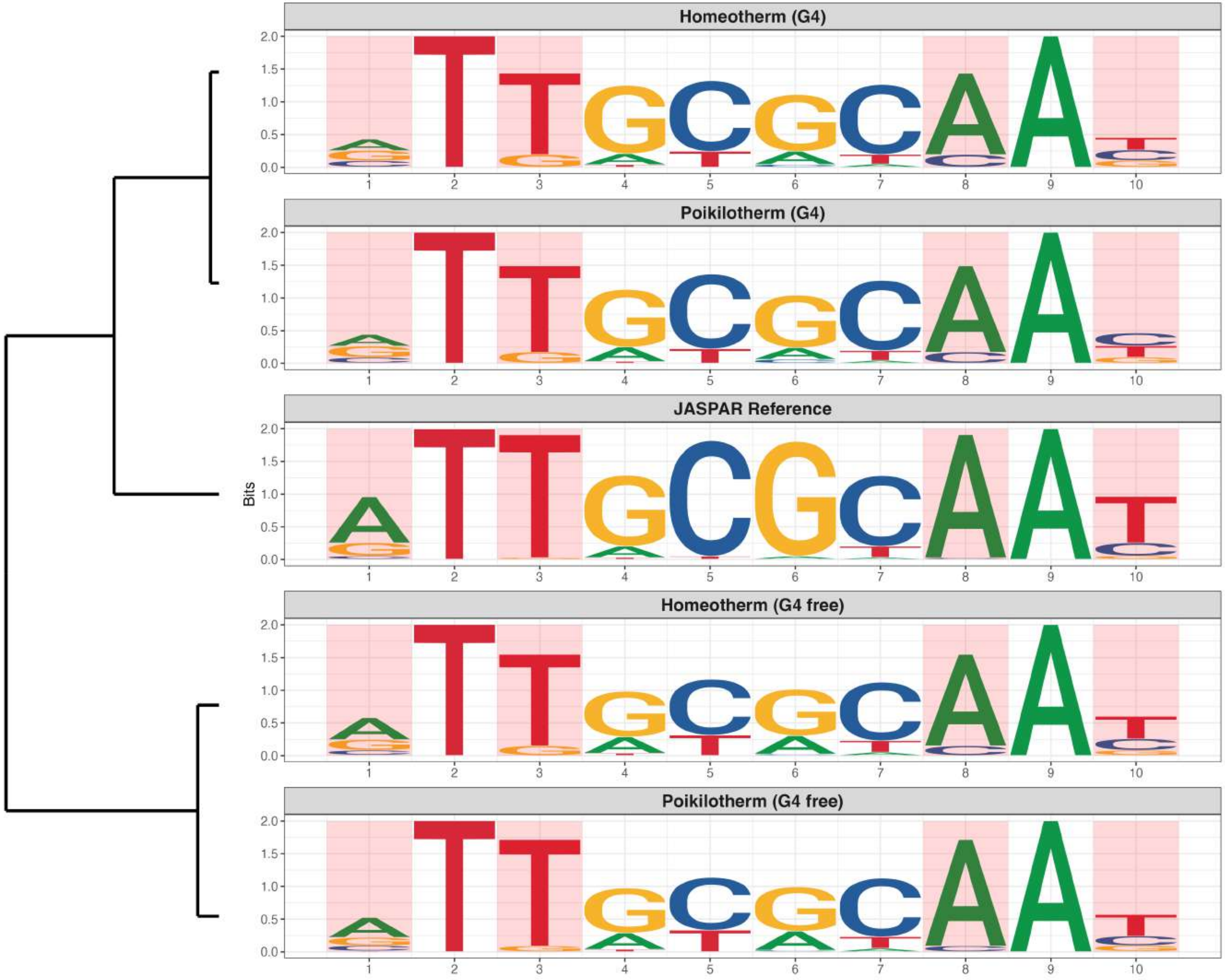

**Promoter : Moderately Reduced in G4s**

*MA1531.2 - NR1D1*

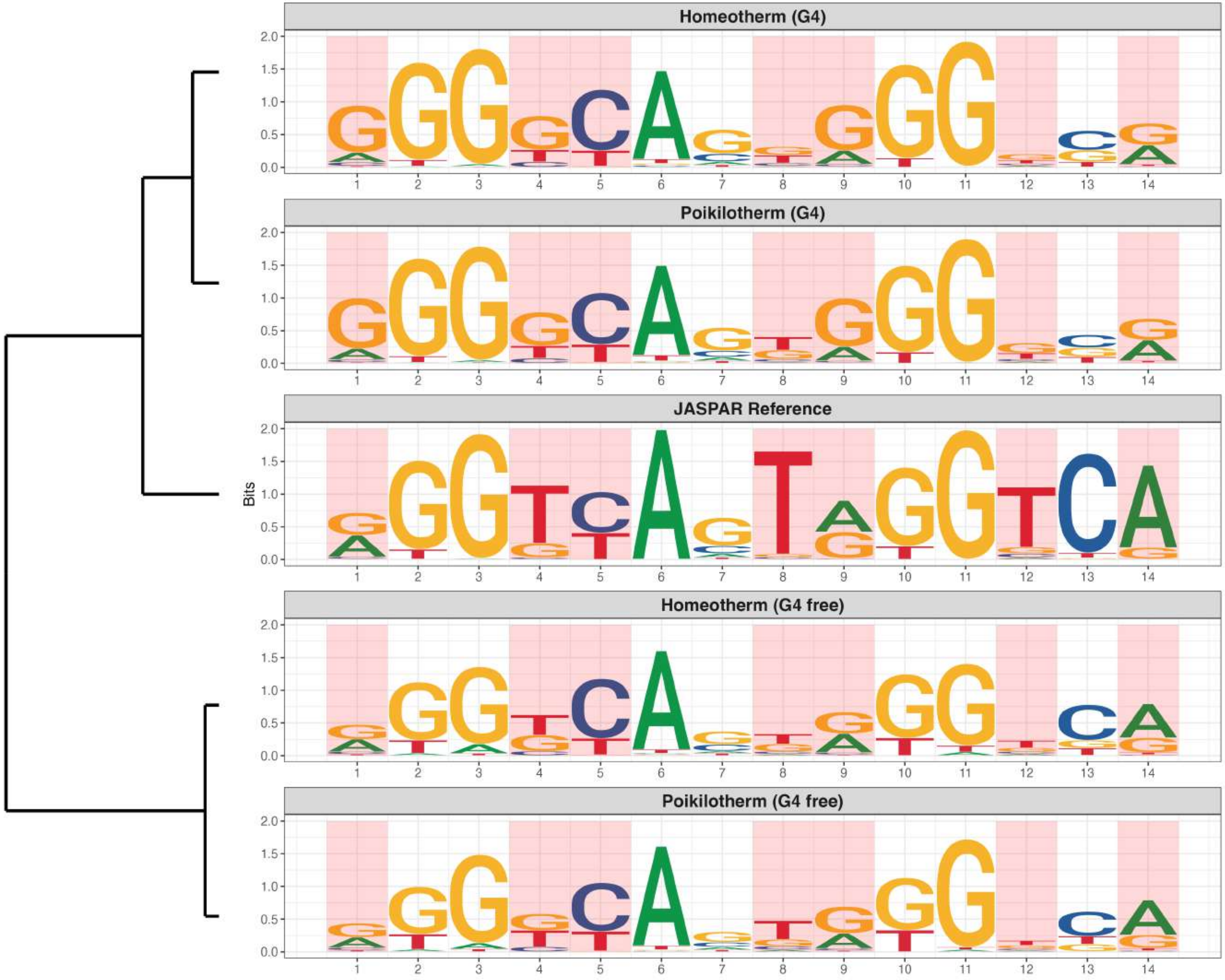

**Promoter : Moderately Reduced in G4s**

*MA1580.1 - ZBTB32*

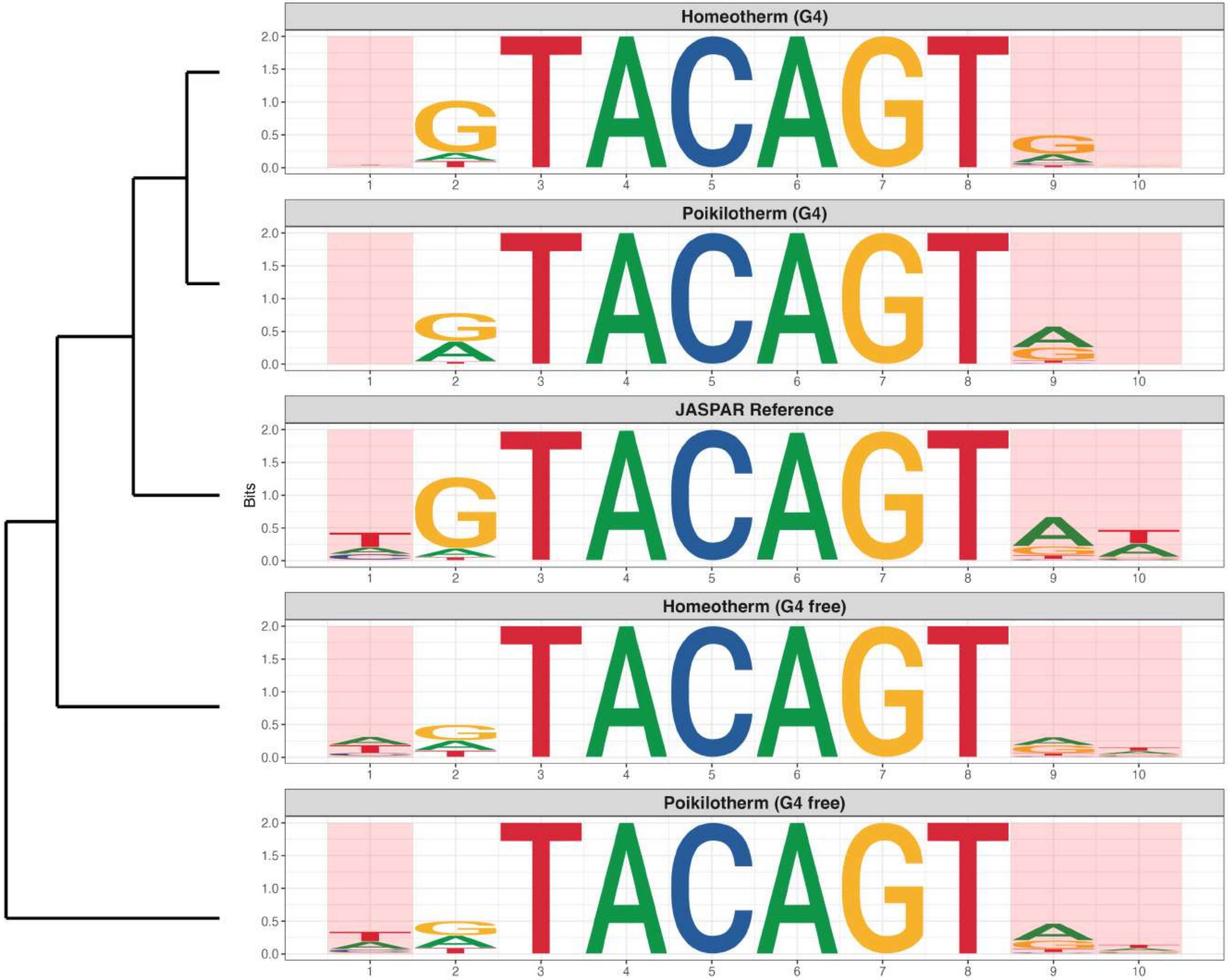

**Promoter : Moderately Reduced in G4s**

*MA1602.2 - ZSCAN29*

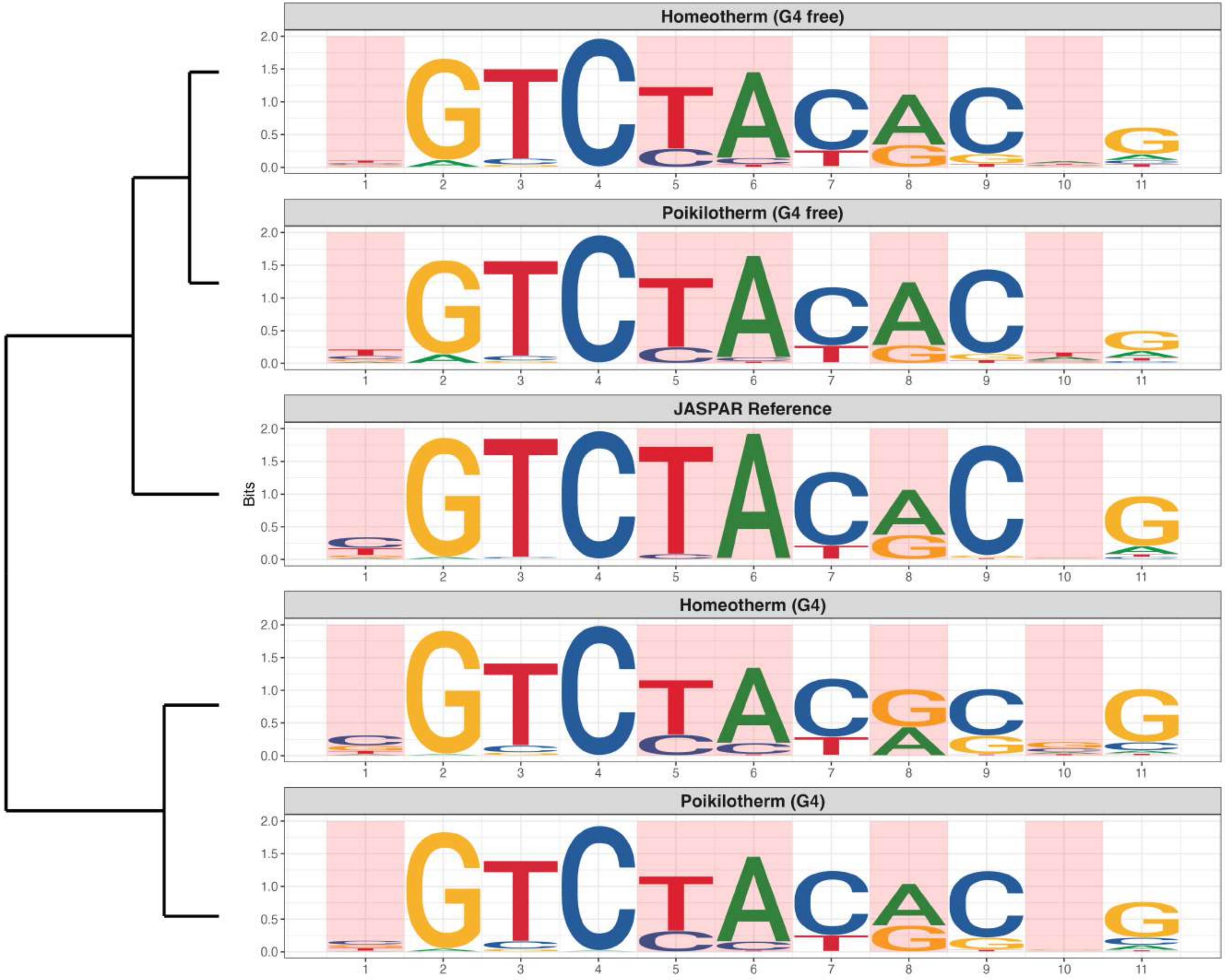

**Promoter : Moderately Reduced in G4s**

*MA1630.3 - ZNF281*

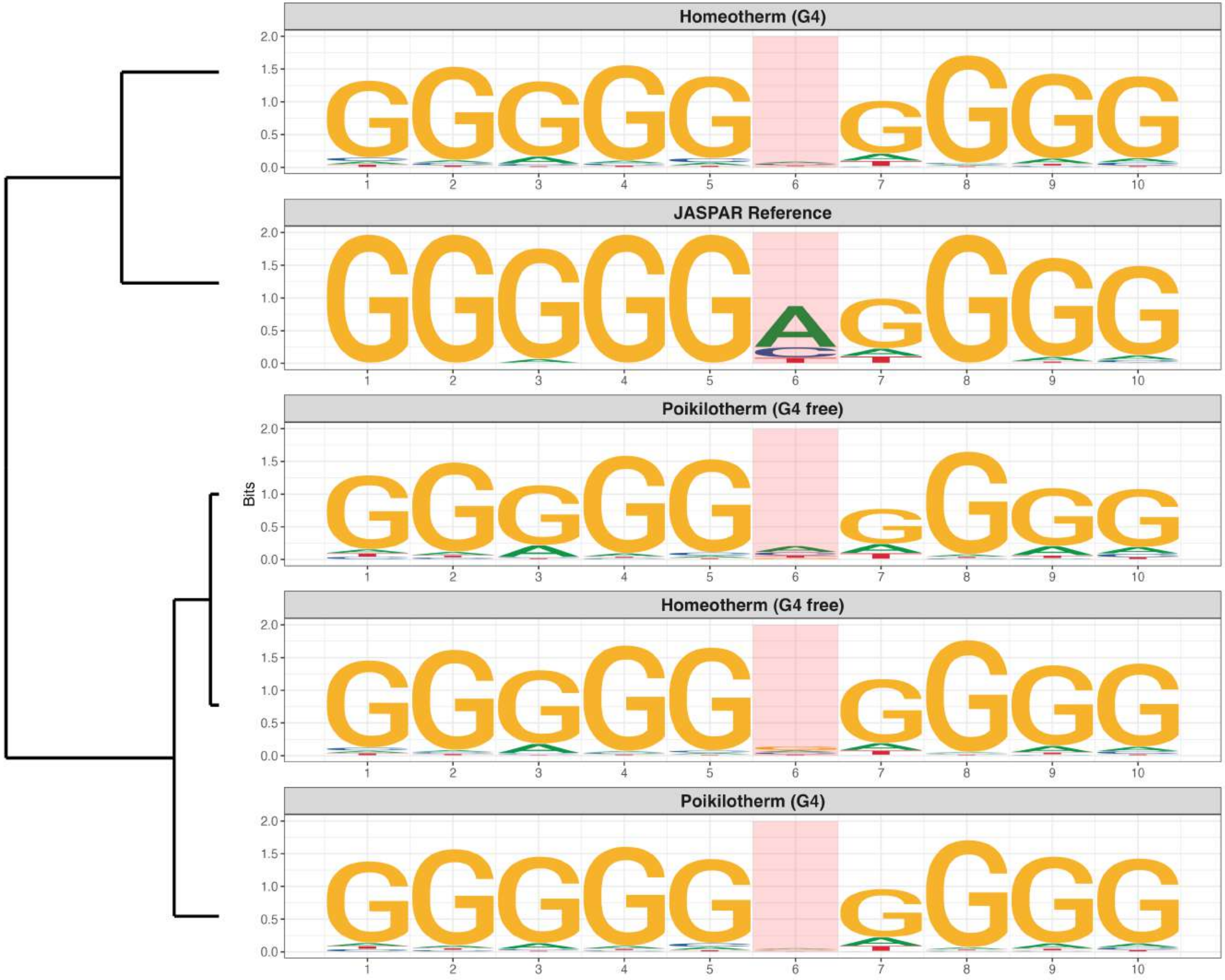

**Promoter : Moderately Reduced in G4s**

*MA1653.2 - ZNF148*

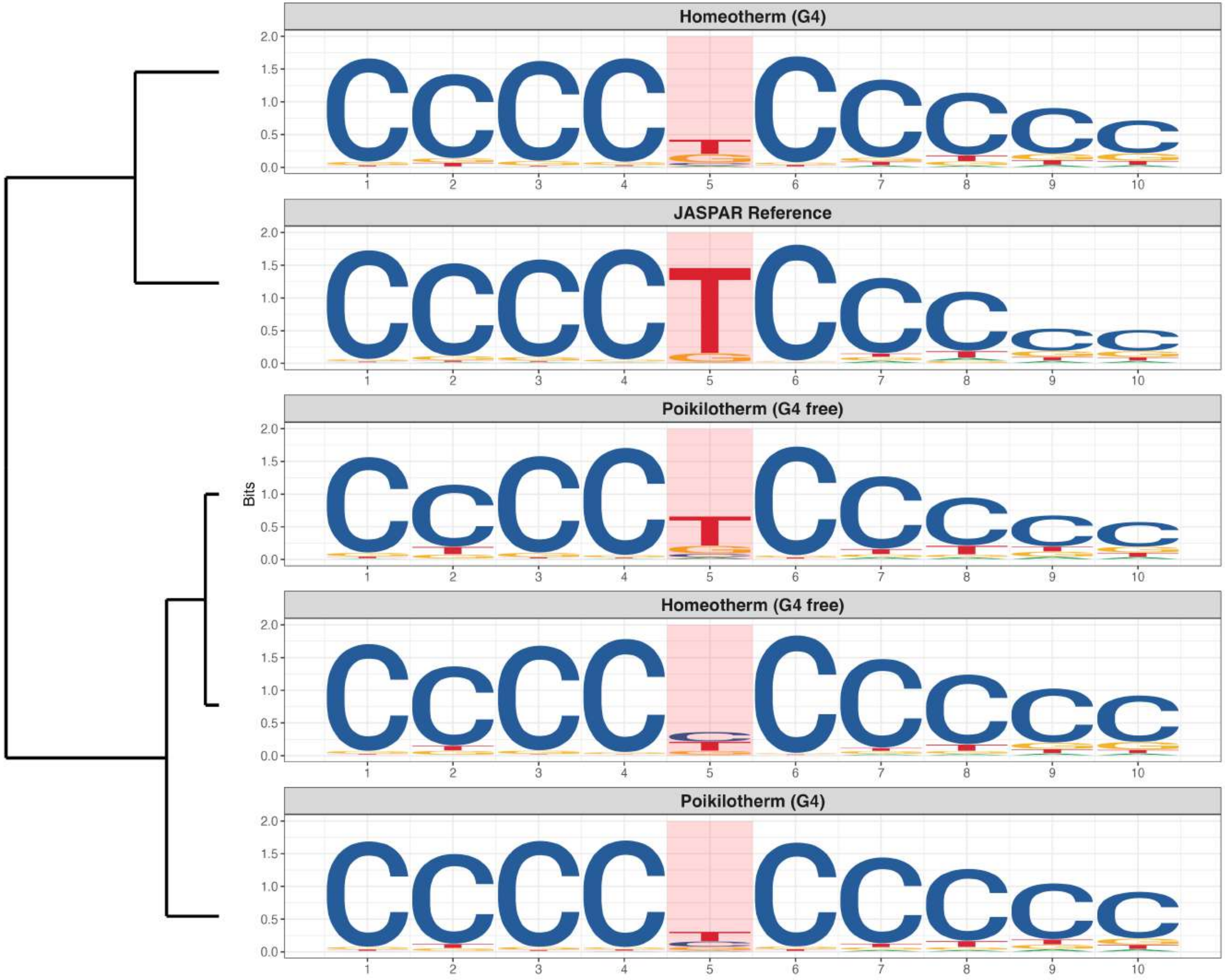

**Promoter : Moderately Reduced in G4s**

*MA1654.2 - ZNF16*

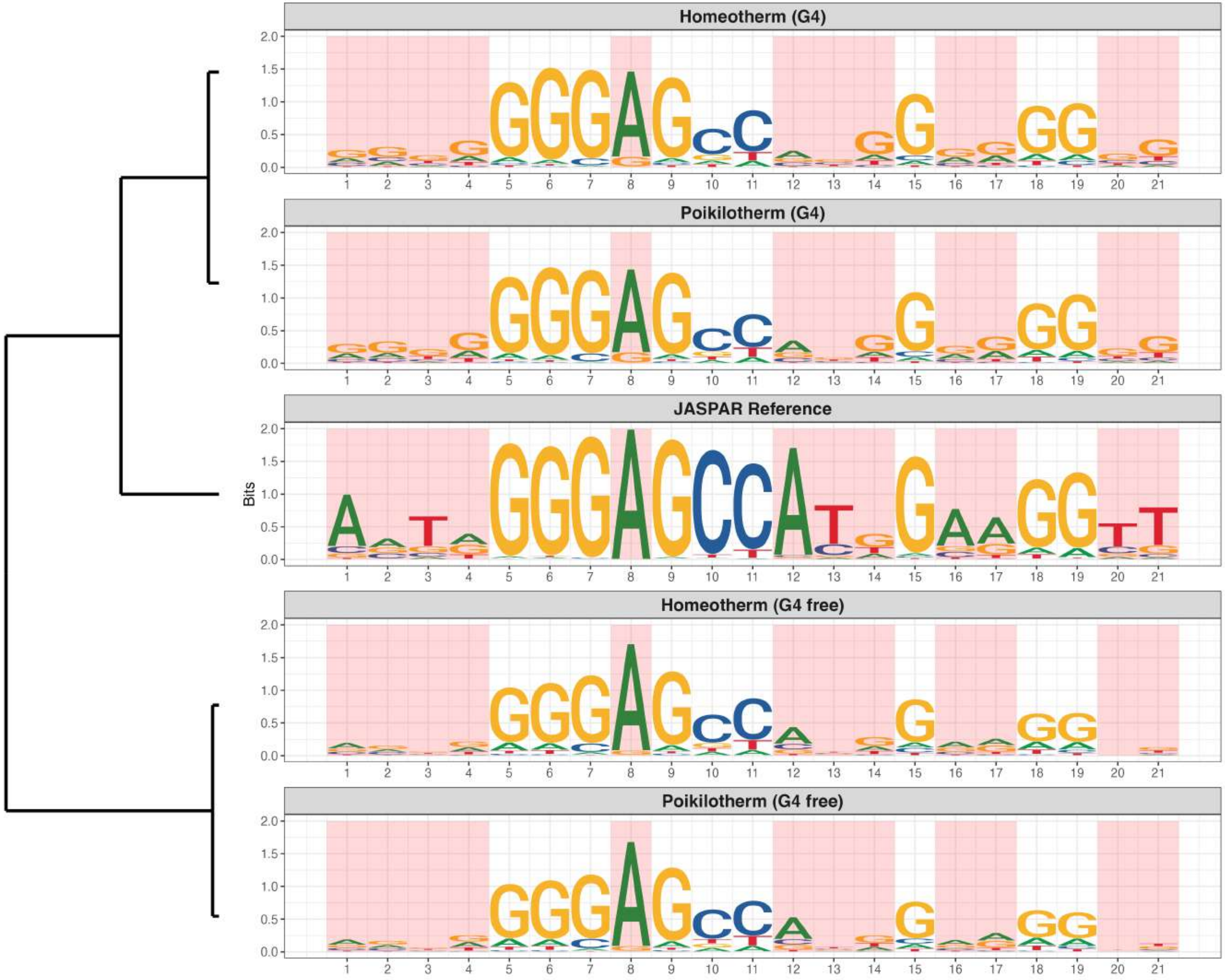

**Promoter : Moderately Reduced in G4s**

*MA2102.1 - ZBTB17*

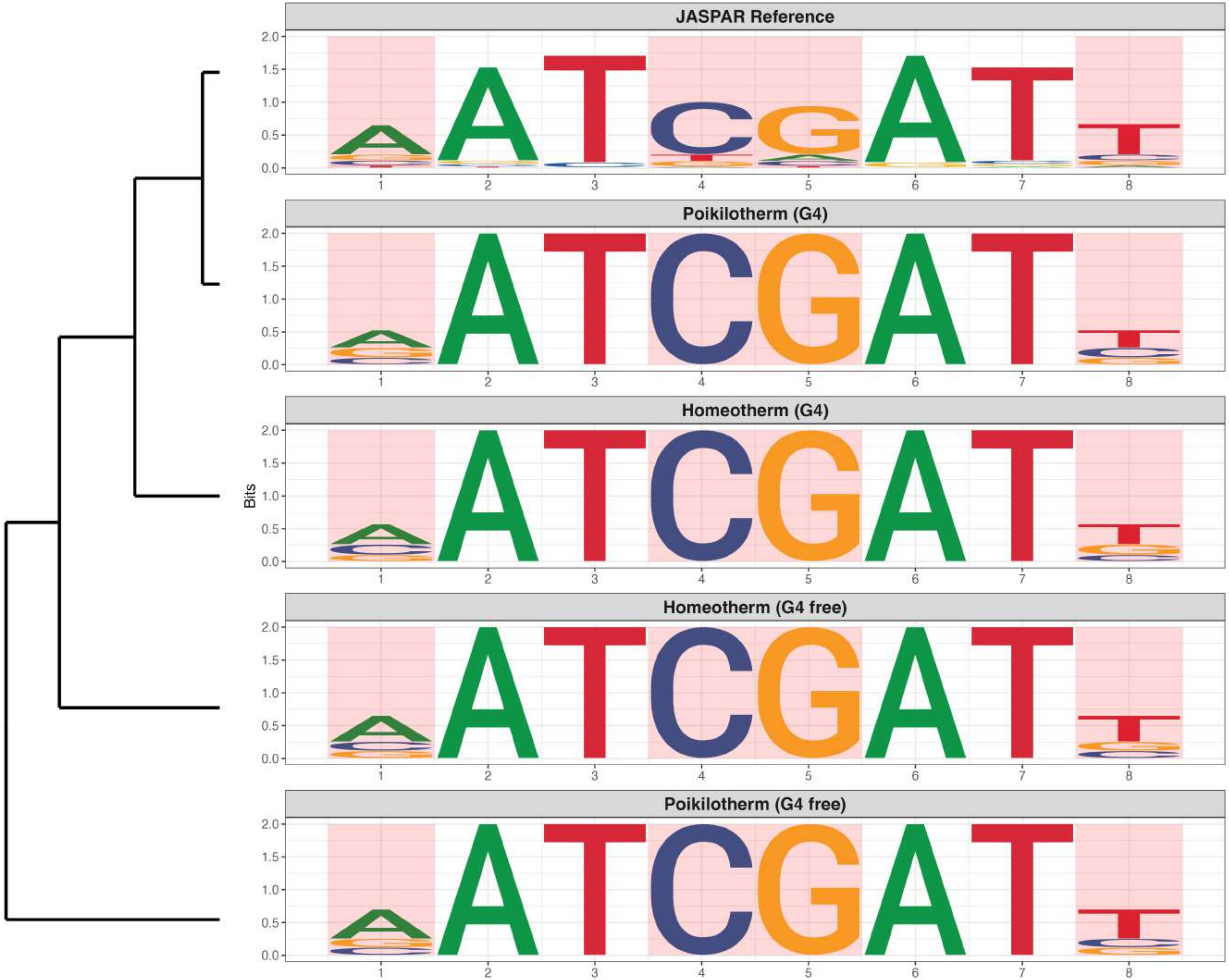

### Promoter : Moderately Reduced in G4s

*MA2494.1 - ZNF436*

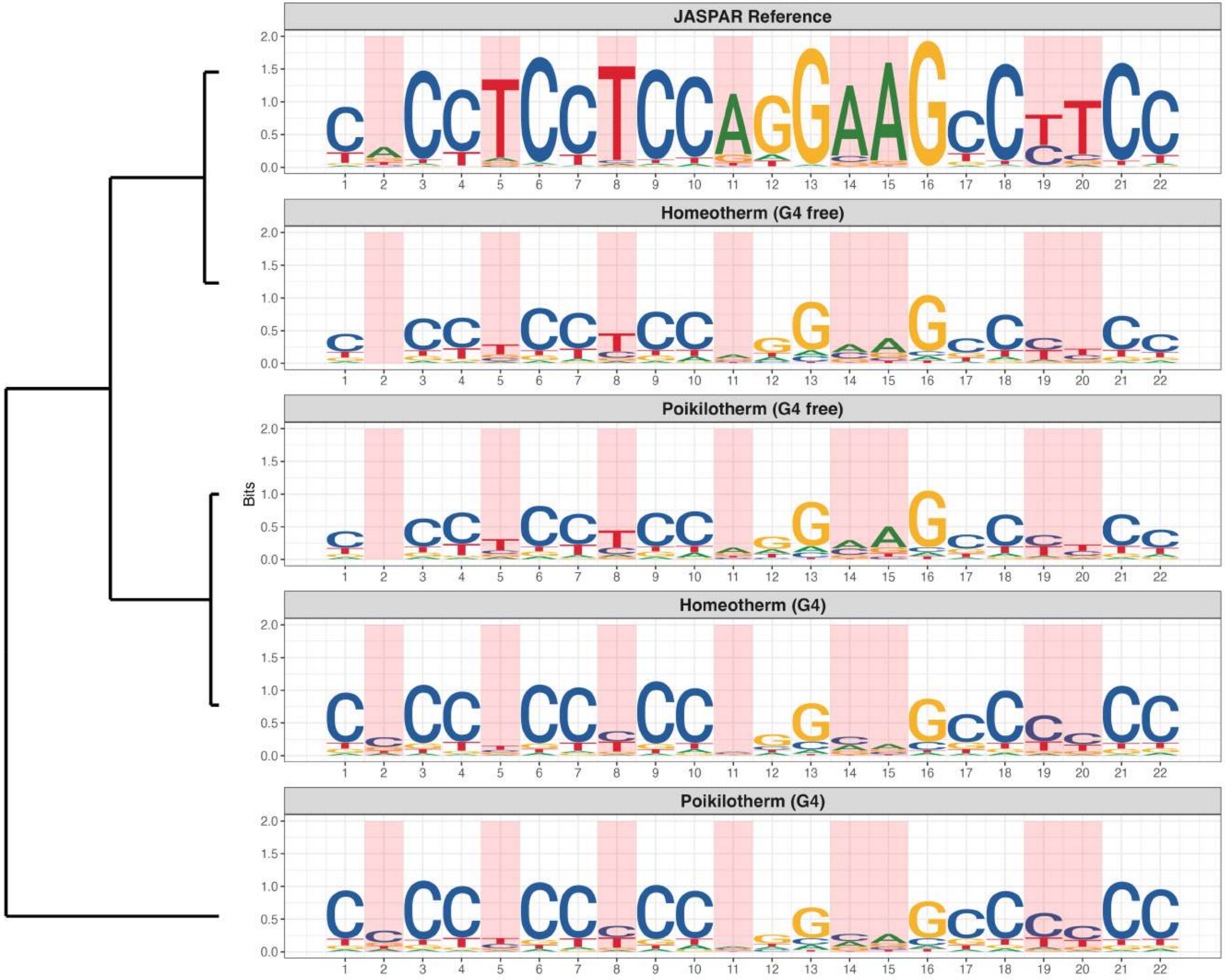

**Promoter : Moderately Reduced in G4s**

*MA2531.1 - ZNF775*

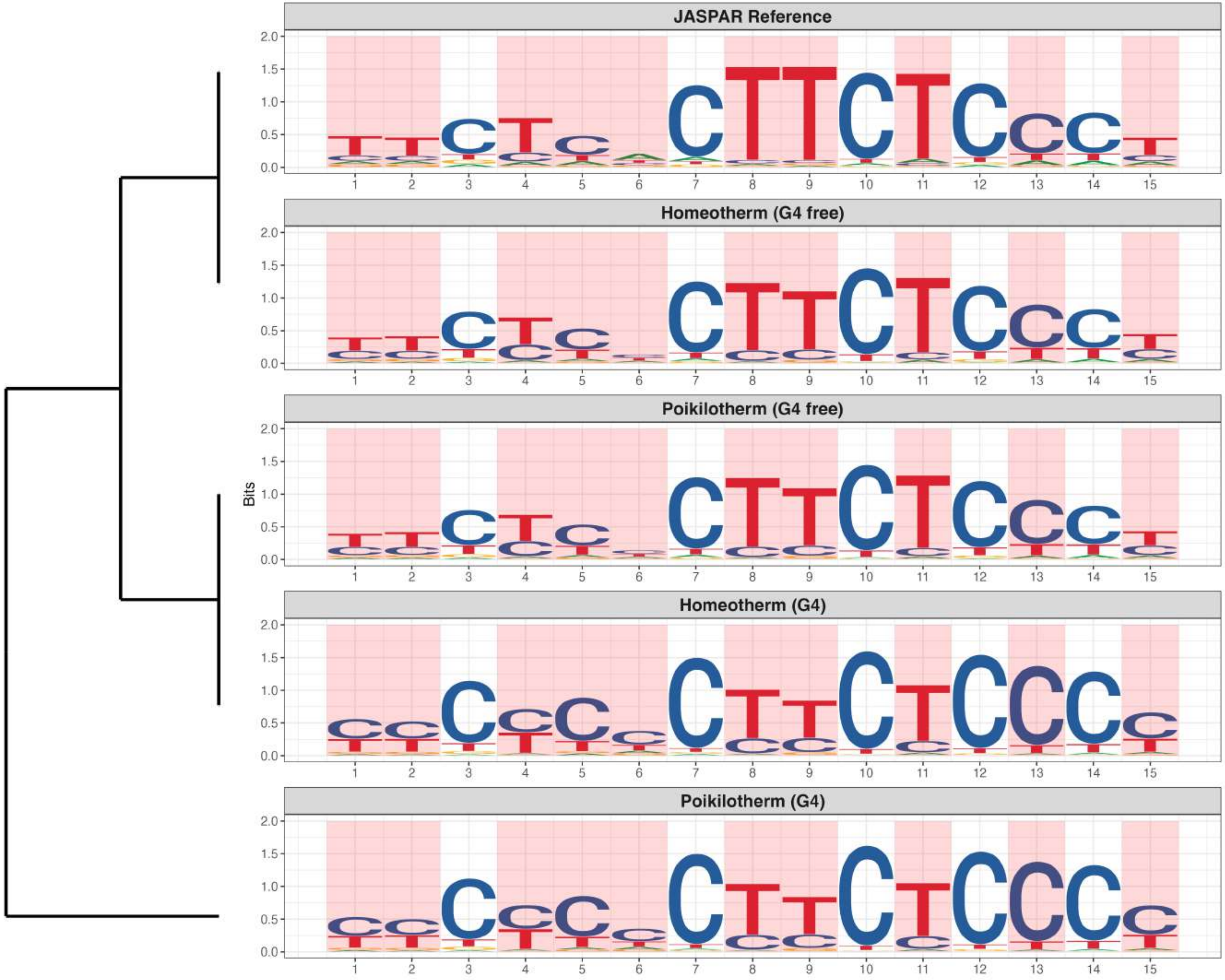

### Promoter : Moderately Reduced in G4s

*MA2534.1 - ZNF831*

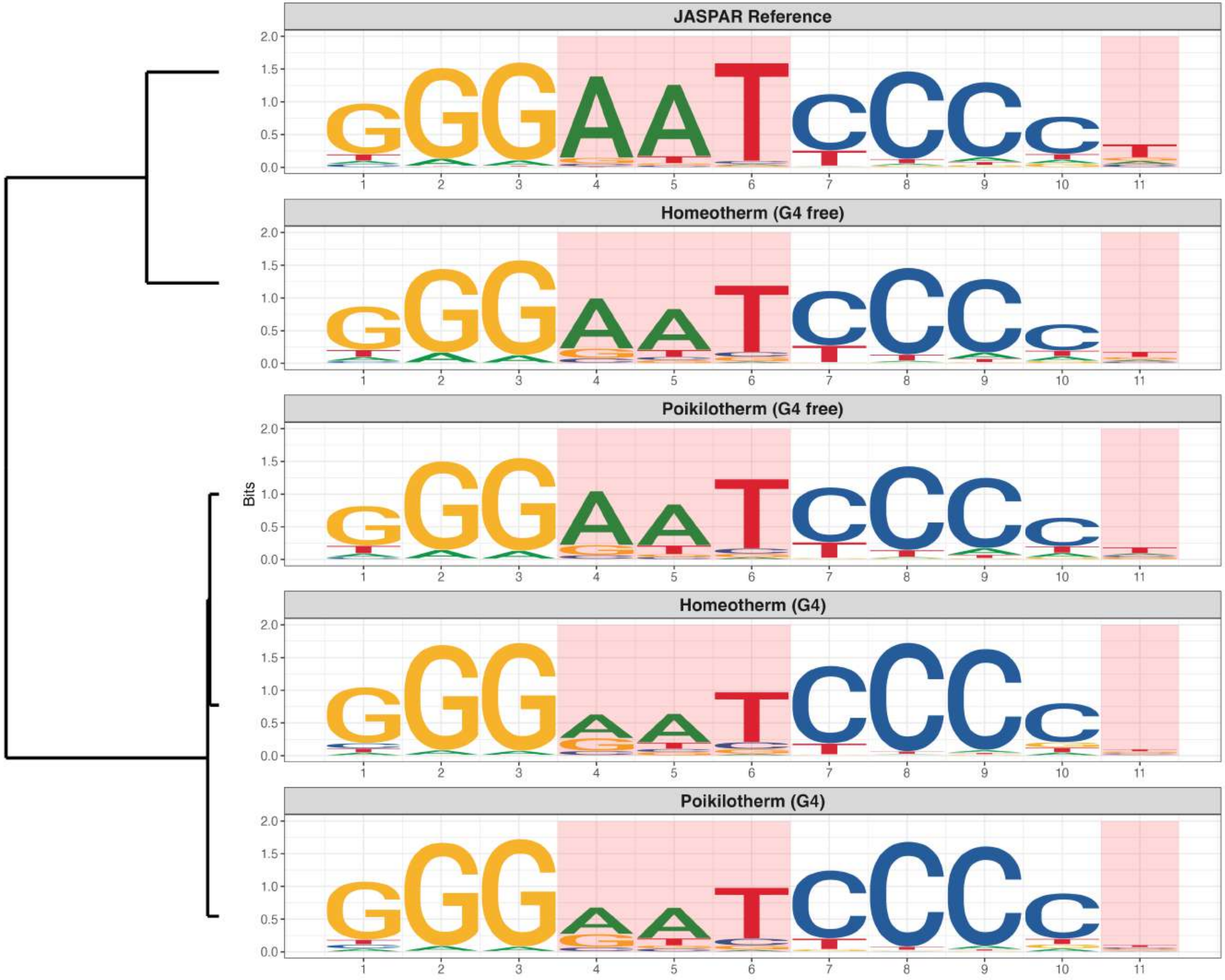
